# Natural killer cell dysfunction drives keloid pathogenesis

**DOI:** 10.1101/2024.11.20.624588

**Authors:** Ying Zhao, Qin Wei, Rui Zeng, Yan Wang, Yong Yang, Yetao Wang

## Abstract

Keloids, characterized by excessive scar tissue resulting from abnormal wound healing, are primarily driven by hyperproliferative fibroblasts and overproduction of extracellular matrix. Although human natural killer (NK) cells are known for their role in inhibiting uncontrolled cell growth through cytokines like IFN-γ and TNF-α, they also produce amphiregulin (AREG), which paradoxically promotes cell proliferation and survival. The involvement of NK cells in keloid pathogenesis, however, remains largely unexplored. This study uncovers remarkable functional changes in NK cells within both lesional skin and the blood of keloid patients. In the skin, NK cell-produced IFN-γ plays a pivotal role in limiting keloid progression by inducing fibroblast apoptosis and curbing excessive extracellular matrix production, while NK cell-derived AREG actively opposes these protective effects. Notably, TGF-β-driven fibroblasts in keloid lesions further dampen NK cell IFN-γ production, revealing a complex and dynamic cellular interplay. In the bloodstream of keloid patients, a distinct NK cell subset emerges, marked by elevated interferon-stimulated genes (ISGs) and diminished IFN-γ production, which correlates with increased plasma IFN-β levels. This elevated IFN-β serves as a key initiating factor, driving NK cell exhaustion through impaired mitochondrial function and metabolic disruption. These findings highlight a critical mechanism underlying the functional abnormalities of keloid-associated NK cells and emphasize the influence of both local and systemic factors in shaping NK cell responses in keloid pathogenesis.

**Graphical abstract:** 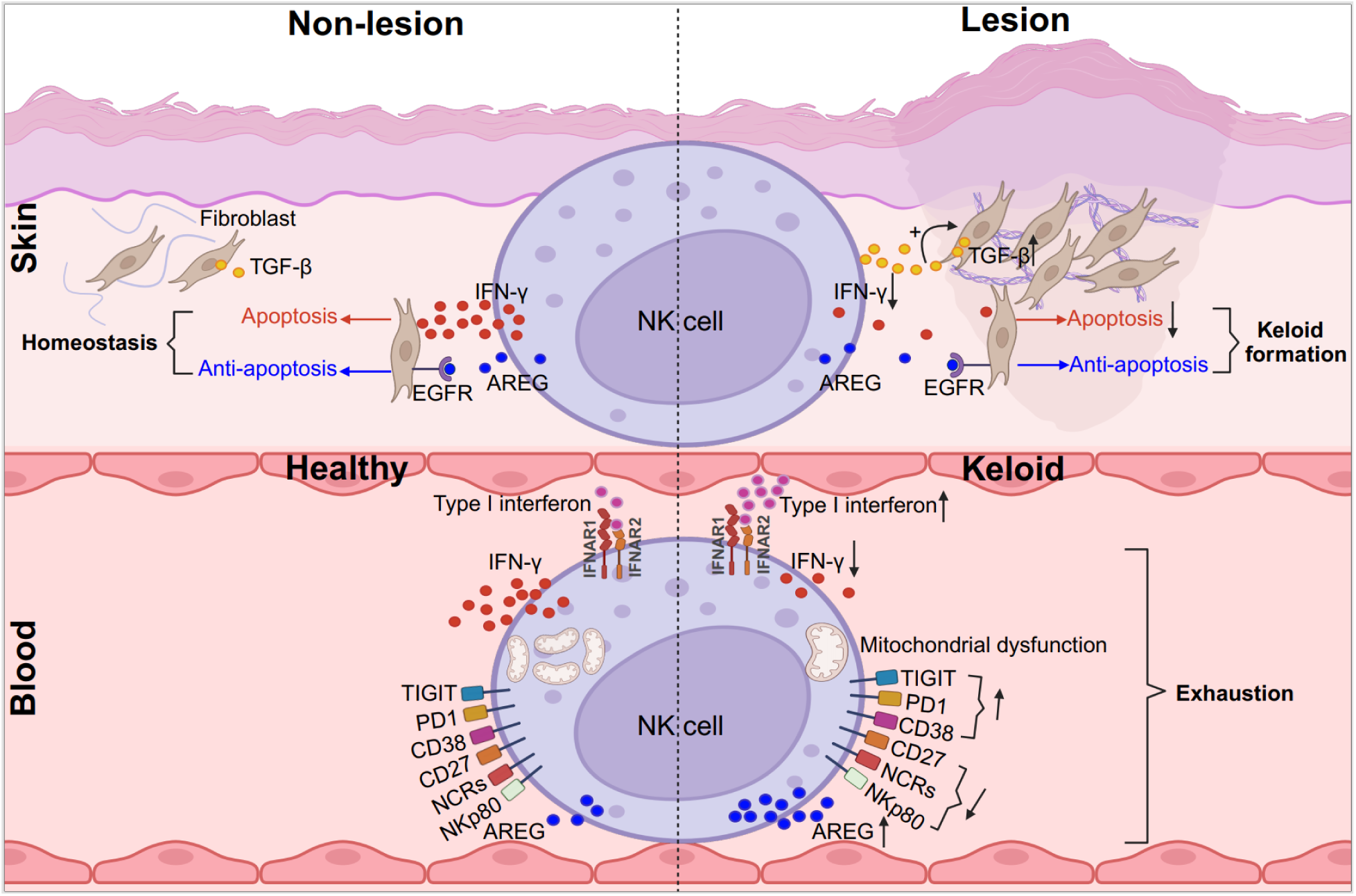

**Highlights:**

- In the skin, NK cell-derived IFN-γ is crucial for suppressing keloid fibroblast proliferation and extracellular matrix production.
- AREG from NK cells counteracts the inhibitory effects of IFN-γ on keloid progression.
- In the blood, elevated IFN-β in keloid patients drives the emergence of ISG^+^NK cells with reduced IFN-γ production.
- IFN-β induces mitochondrial dysfunction and functional exhaustion in NK cells.

## Introduction

Keloid is a debilitating fibroproliferative skin disorder arising from abnormal wound healing, characterized by elevated, thickened scars that extend beyond the original injury^1^. Documented as early as 1700 BC^2^, this ancient disease continues to affect a significant portion of the population (0.09% to 16%)^3^, with prevalence shaped by factors such as genetic predisposition, age, race, and ethnicity^2,3^. The associated pain, itching, restricted movement, cosmetic concerns, and emotional distress profoundly impact physical health and psychological well-being^4^.

Keloid formation is primarily driven by the hyperproliferation of fibroblasts and their excessive synthesis of extracellular matrix^5,6^. The crosstalk between fibroblasts and immune cells is pivotal in the development and persistence of keloids. Cytokines such as IL-1β, IL-6, IL-33, and TNF-α, produced by fibroblasts, are instrumental in shaping the local immune environment and contributing to chronic inflammation^7,8^. Conversely, growth factors and cytokines like PGDF, VEGF, IGF, TGF-β, IL-4, and IL-13, derived from M2 macrophages, Tregs, and Th2 cells, are essential for sustaining fibroblast proliferation and extracellular matrix deposition^9–11^. keloids, therefore, serve as a valuable model for investigating the interplay between fibroblasts and immune cells.

Blood acts as a primary reservoir for immune cells that are recruited to inflamed skin, accumulating evidence linking abnormalities in these blood-derived immune cells to the pathogenesis of various skin diseases. For instance, Tregs from patients with severe psoriasis exhibit an increased propensity to differentiate into pathogenic IL-17^+^RORγt^+^ cells, a phenomenon not observed in healthy individuals^12^. Additionally, the excessive upregulation of CD71 on blood T cells disrupts mTORC1 signaling, leading to enhanced differentiation of Th17 cells while suppressing Treg expansion, a mechanism implicated in the pathogenesis of systemic lupus erythematosus (SLE)^13^. In a steady state, NK cells are typically rare in healthy skin, whereas, during bacterial or viral infections, circulating natural killer (NK) cells can be rapidly recruited to the skin, where they exert effector functions and differentiate into tissue-resident NK cells in a TCF7-dependent manner^14^. Therefore, blood-derived immune cells are critically involved in the pathology of skin diseases. However, the connection between their dysfunction and keloid pathogenesis remains unclear. Gaining a deeper understanding of associated mechanisms could pave the way for new therapeutic targets.

NK cells are well-known for inhibiting hyperproliferative cell growth by secreting cytokines like IFN-γ and TNF-α, and releasing cytotoxic molecules such as perforin and granzymes, which are essential in limiting disease progression associated with uncontrolled cell growth, such as tumors^15^. The activity of NK cells is closely linked to the pathogenesis of both melanoma and nonmelanoma skin cancers^16,17^. Interestingly, unlike mouse NK cells, recent findings demonstrate that human NK cells can produce amphiregulin (AREG)^18^, a ligand of the epidermal growth factor receptor (EGFR), which promotes cell proliferation and mediates immune tolerance^19^. How these contrasting roles of NK cells influence the progression of keloids, and whether keloid-associated pathology affects the normal physiology of NK cells, remains unknown.

In this study, we uncovered the functional crosstalk between fibroblasts and NK cells in keloid pathogenesis, which likely contributes to the uncontrolled proliferation of fibroblasts observed in keloid lesions. Notably, elevated type 1 interferon (IFN) levels in the plasma of keloid patients lead to the exhaustion of NK cells, characterized by reduced metabolic activity and impaired IFN-γ production, resulting in a unique ISG^+^NK cell subset in the blood. These findings suggest that targeting NK cells may represent a promising therapeutic strategy for keloid treatment.

## Results

### Alterations in NK cell percentage and function in keloid lesional skin

To investigate the involvement of NK cells in keloid pathogenesis, NK cells from donor matched lesional (L) and adjacent non-lesional (NL) skin of keloid patients were analyzed by flow cytometry (Table S1). A 14-marker antibody panel, including CD3, CD4, TCRαβ, TCRγδ, CD19, CD20, CD22, CD34, FcεRIα, CD11c, CD303, CD123, CD1a, and CD14, was employed to exclude T cells, B cells, monocytes, and other lineage-positive cells. NK cells were identified as Lin^-^TBX21^+^ within the CD45 positive population (Figure 1A), in line with previous studies^20–24^, and the absence of CD127 excluded the involvement of other ILCs in NK cell gating (Figure S1A).

**Figure 1.**
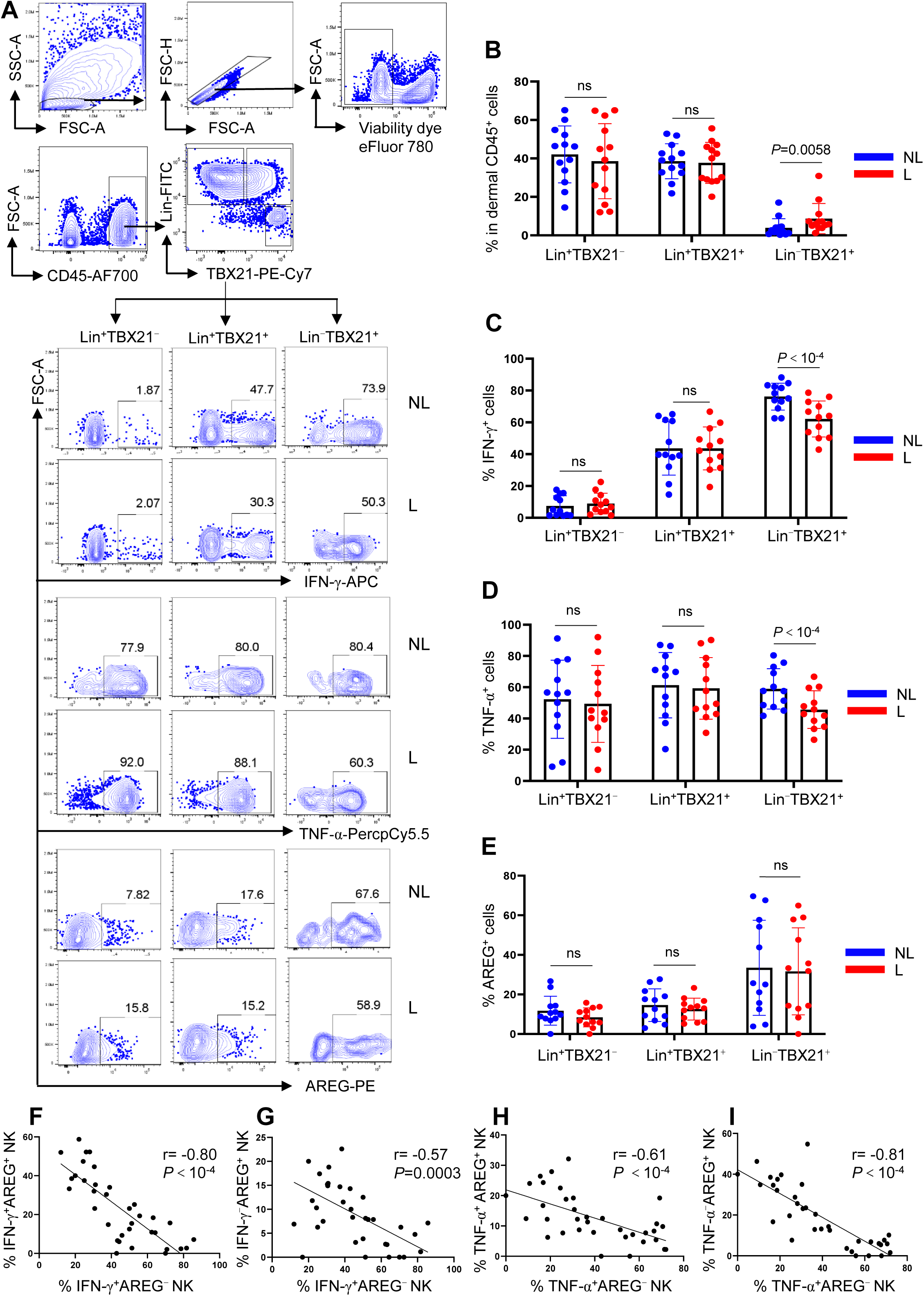
Changes in NK cell proportion and function in keloid lesional skin. (A) Flow cytometry analysis of CD45^+^, NK, IFN-γ^+^, TNF-α^+^, and AREG^+^ cell populations in the non-lesional (NL) and lesional (L) keloid dermis. Dermal cells were stimulated with PMA and ionomycin for 2 hours prior to cytokine detection. (B) Percentage of Lin^+^TBX21^-^, Lin^+^TBX21^+^ and Lin^-^TBX21^+^ populations within CD45^+^ cells in (A) (n=13). (C-E) Percentage of the IFN-γ^+^ (C), TNF-α^+^ (D) and AREG^+^ (E) cells in Lin^+^TBX21^-^, Lin^+^TBX21^+^ and Lin^-^TBX21^+^ populations (n=12). (F-I) Correlation of IFN-γ^+^AREG^-^NK cells with IFN-γ^+^AREG^+^ (F) and with IFN-γ^-^AREG^+^ (G) NK cells; and correlation of TNF-α^+^AREG^-^NK cells with TNF-α^+^AREG^+^ (H) and with TNF-α^-^AREG^+^ (I) NK cells (n=35). For (B-E), each dot represents a unique keloid donor, two-tailed paired t-test, data are mean with s.e.m., for (F-I), Pearson correlation, ns, not significant. Cell culture and stimulation conditions for each panel were detailed in Methods.

The percentage and number of CD45^+^ cells in the dermis were comparable between keloid lesional and non-lesional skin (Figures S1B and S1C). However, the proportion of NK cells within the CD45^+^ population was elevated in the lesional skin of keloid patients, this increase appeared specific to NK cells, as the proportions of ILCs (Lin^-^TBX21^-^CD127^+^), Lin^+^TBX21^+^ and Lin^+^TBX21^-^ populations remained unchanged (Figures S1A, S1D and 1B). Further analysis revealed that NK cells had the highest IFN-γ production potential compared to Lin^+^TBX21^+^ and Lin^+^TBX21^-^ populations. Nonetheless, IFN-γ and TNF-α production by NK cells were impaired in lesional skin relative to non-lesional skin (Figures 1A, 1C and 1D). These findings suggest that skin NK cells specifically respond to keloid pathology with altered composition and biological function.

In addition to anti-proliferative cytokines, human NK cells also produce AREG^22^, which promotes cell proliferation and inhibits apoptosis through EGFR signaling^19^. Among the cell populations analyzed, NK cells exhibited the highest AREG production (Figures 1A and 1E). The proportion of AREG^+^NK cells was similar between lesional and non-lesional skin in keloid patients (Figures 1A and 1E). However, AREG^+^NK cells showed an inverse correlation with AREG^-^NK cells, independent of IFN-γ and TNF-α production (Figures 1F-1I), suggesting functional differences between these subsets that may differentially influence keloid pathology.

### TGF-β-driven fibroblast inhibition of NK cell IFN-γ production in keloid lesions

To explore whether the functional differences in NK cells between keloid lesional and non-lesional skin were influenced by interactions with fibroblasts, we conducted transwell assays to evaluate the impact of fibroblast derived factors on NK cell function (Figure 2A). Keloid lesional fibroblasts, compared to non-lesional fibroblasts, selectively downregulated IFN-γ but not AREG production in NK cells from peripheral blood mononuclear cells (PBMCs) (Figures 2B and 2C). Similar inhibition was observed in magnetic beads enriched NK cells (Figure 2D, NK cell purity >95%), confirming that lesional fibroblasts suppress NK cell function independently of other cell types (Figures 2D and 2E).

**Figure 2.**
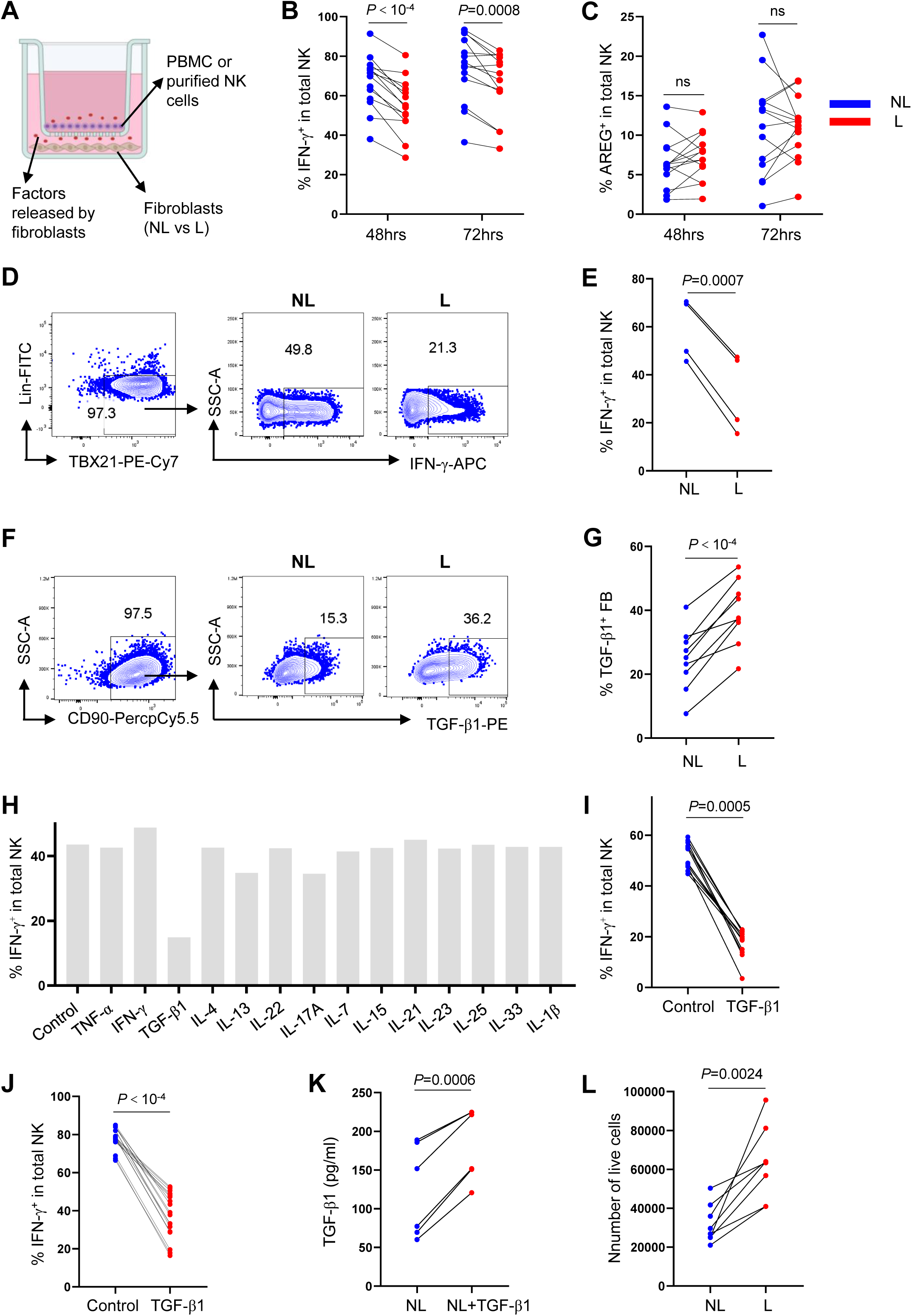
TGF-β-induced fibroblast suppression of NK cell IFN-γ production in keloid lesions. (A) PBMCs or magnetic beads enriched NK cells were co-cultured with keloid NL and L fibroblasts in transwell. (B, C) Percentage of IFN-γ^+^ (B, n=14) and AREG^+^ (C, n=14) cells among total NK cells after PBMCs were cultured with NL or L fibroblasts for 48 or 72 hrs, as in (A). (D) Flow cytometry of IFN-γ^+^ cells in enriched NK cells cultured with NL or L fibroblasts for 48 hrs, as in (A). (E) Percentage of IFN-γ^+^ cells in (D) (n=4). (F) Flow cytometry of TGF-β1^+^ cells in keloid NL and L fibroblasts after culturing in DMEM with golgi block for 16hrs. (G) Percentage of TGF-β1^+^ cells in (F) (n=9). (H) NL fibroblasts were stimulated with the indicated cytokines for 24 hrs. After removal of the stimulants, NL fibroblasts were cultured with PBMCs for 48 hrs in transwells, and IFN-γ^+^ cells among NK cells were analyzed. (I, J) NL fibroblasts were treated with or without TGF-β1, then cultured with PBMCs (I, n=12) or enriched NK cells (J, n=18) as in (H). IFN-γ^+^ cells among NK cells were analyzed. (K) NL fibroblasts were treated as in (I, J). After three washes, the fibroblasts were cultured in DMEM for 48 hrs, and the TGF-β1 concentration in the supernatant was measured by ELISA (n=6). (L) Enriched NK cells (n=8) were stimulated with IL-12 + IL-15 + IL-18 for 16 hrs. After removal of the stimulants, the NK cells were cultured with NL or L fibroblasts (start at 10^5^ cells) for 48 hours in transwells, and the number of live fibroblasts was analyzed. For (B, C, E, G, J-K), two-tailed paired t-test, for (I), Wilcoxon matched-pairs signed rank test, data are mean with s.e.m., ns, not significant. Cell culture and stimulation conditions for each panel were detailed in Methods.

TGF-β plays pivotal roles in promoting fibroblast proliferation, collagen deposition, and immune suppression in keloid development and progression^10,25^. To determine whether differences in fibroblast behavior were linked to TGF-β expression, we compared TGF-β1 production between lesional and non-lesional fibroblasts. Our results showed that keloid lesional fibroblasts produced higher levels of TGF-β1 than donor-matched non-lesional fibroblasts (Figures 2F and 2G), which likely contributes to the enhanced inhibitory effect of lesional fibroblasts on NK cell IFN-γ production (Figures 2B and 2E).

RNA-Seq data revealed that lesional keloid fibroblasts expressed higher TGF-β levels than normal skin fibroblasts (Figures S2A and S2B). To assess whether cytokines in keloid lesions drive normal fibroblasts toward a more immunosuppressive phenotype with increased TGF-β expression (Figures 2B-2G), non-lesional fibroblasts were treated with 14 cytokines elevated in keloid or inflammatory skin diseases (Figure 2H)^10,11,26^. After removing these stimulatory conditions, transwell assays showed that only TGF-β1 treated fibroblasts inhibited NK cell IFN-γ production (Figures 2H-2J), correlating with increased fibroblast TGF-β1 production following TGF-β1 treatment (Figures 2K, S2C, and S2D). Furthermore, keloid lesional fibroblasts were more resistant to NK cell induced growth restriction than non-lesional fibroblasts (Figures 2L and S2E). Thus, fibroblast produced TGF-β establishes a positive feedback loop that enhances its suppressive effect on NK cell IFN-γ production and contributes to keloid persistence.

### Effects of NK cell derived IFN-γ and AREG on keloid fibroblasts

Next, we examined the biological effects of different combinations of AREG and IFN-γ produced by NK cells on fibroblast viability and extracellular matrix production. IFN-γ^-^AREG^-^, IFN-γ^+^AREG^-^, IFN-γ^-^AREG^+^ and IFN-γ^+^AREG^+^ NK cells were generated (Figures 3A and 3B), and their conditioned medium was tested on keloid lesional fibroblasts.

**Figure 3.**
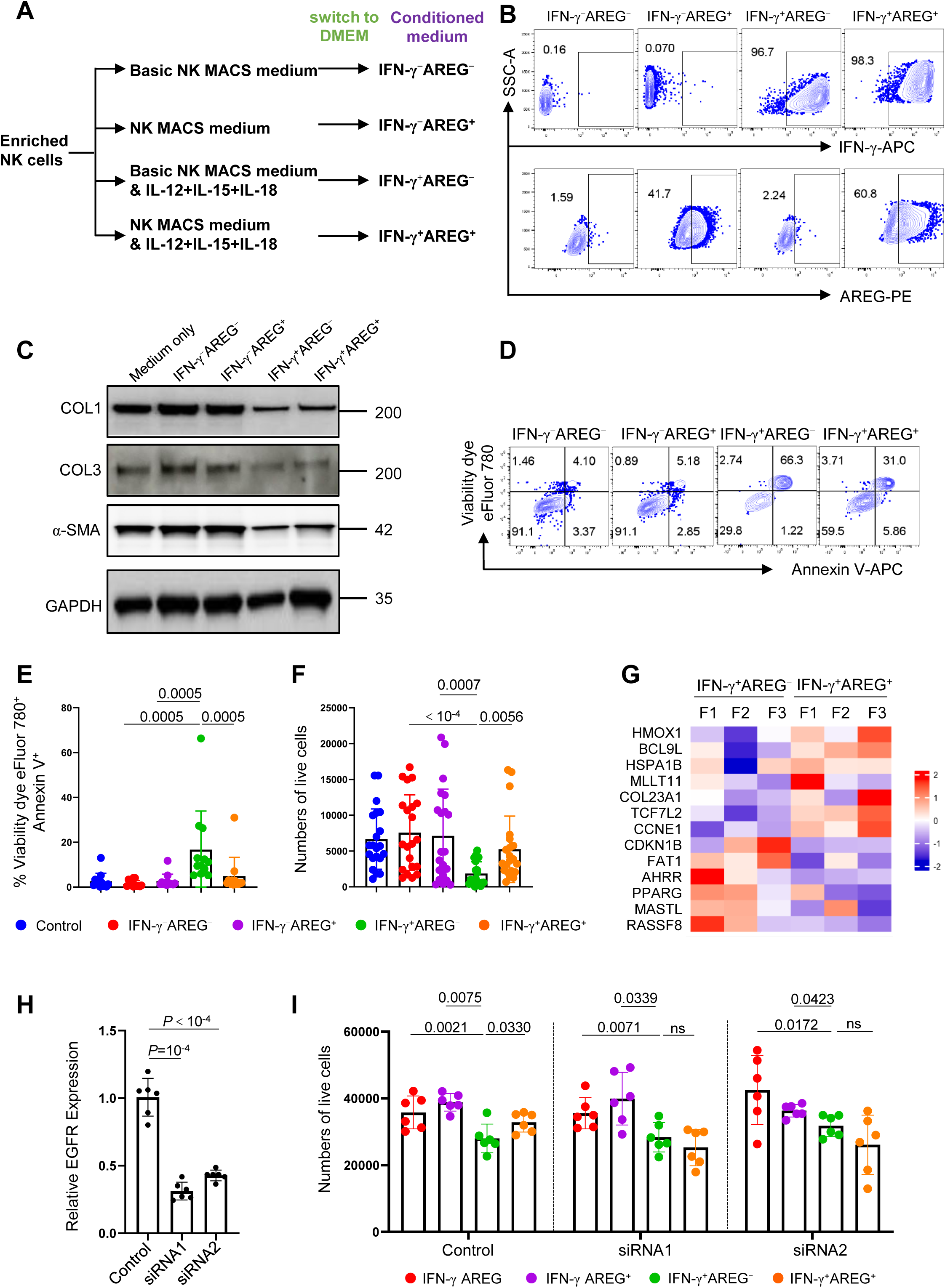
Impact of NK cell derived IFN-γ and AREG on keloid fibroblasts. (A) IFN-γ^-^AREG^-^, IFN-γ^+^AREG^-^, IFN-γ^-^AREG^+^ and IFN-γ^+^AREG^+^ NK cells were generated under indicated culture conditions. The culture medium was then replaced with DMEM and maintained for 48 hours. The conditioned DMEM was used for fibroblast culture. (B) Flow cytometry detection of IFN-γ and AREG in NK cells from the indicated culture conditions in (A). (C) Keloid fibroblasts were cultured in conditioned mediums in (A) for 24 hrs, collagen 1 (COL1), collagen 3 (COL3), α-SMA and GAPDH were detected by western blot. (D) Fibroblasts were cultured in conditioned mediums in (A) for 48 hours, stained with a viability dye and anti-Annexin V antibody, and analyzed by flow cytometry. (E, F) Percentage of the viability dye^+^Annexin V^+^ fibroblasts (E, n=12) and number of live fibroblasts (F, n=21) cultured in conditioned medium as in (D). (G) Heatmap of DEgenes in keloid fibroblasts cultured in IFN-γ^+^AREG^-^ vs IFN-γ^+^AREG^+^ conditioned medium (*P*<0.05, |log_2_FC|>0.5, n=3). (H) EGFR expression relative to β-actin in keloid fibroblasts transfected with control (non-targeting siRNA) or EGFR-targeting siRNA, measured by quantitative reverse transcription PCR (n=6). (I) Number of live cells in control and EGFR knockdown fibroblasts cultured in NK cell-conditioned medium as in (D) (n=6). All fibroblasts used in the experiments were derived from keloid lesional skin. For (E), Wilcoxon matched-pairs signed rank test, for (F, H, I), two-tailed paired t-test, data are mean with s.e.m., ns, not significant. Cell culture and stimulation conditions for each panel were detailed in Methods.

Conditioned medium containing IFN-γ (IFN-γ^+^AREG^-^ and IFN-γ^+^AREG^+^) significantly reduced fibroblast production of key factors involved in keloid fibrosis, including collagen 1 (COL1), collagen 3 (COL3), and α-SMA (Figures 3C and S3A-S3C)^27^. IFN-γ^+^AREG^-^ medium led to increased apoptosis and reduced cell viability compared to controls (medium only and IFN-γ^-^AREG^-^ medium) (Figures 3D-3F and S3D). Notably, IFN-γ^+^AREG^+^ medium resulted in less apoptosis and greater survival than IFN-γ^+^AREG^-^ medium, despite comparable NK cell IFN-γ production between the two conditions (Figures 3D-3F, S3D, and S3E). The protective effect of AREG was linked to increased expression of anti-apoptotic genes (HMOX1, HSPA1B, BCL9L, MLLT11, COL23A1, TCF7L2, CCNE1) (Figure 3G and Table S2)^28–34^, and disruption of AREG-EGFR signaling by EGFR knockdown impaired AREG’s protection against IFN-γ-induced apoptosis (Figures 3H, 3I and S3F). Thus, NK cell-derived IFN-γ limits extracellular matrix production and induces apoptosis, while AREG counteracts IFN-γ-induced apoptosis in keloid lesional fibroblasts.

### NK cell-derived IFN-γ and AREG differentially affect keloid progression in mouse xenografts

Unlike human NK cells, our previous study demonstrated that the chromatin of AREG region in mouse NK cells is not accessible, thus no transcripts of AREG in mouse NK cells, which indicated a major functional difference between human and mouse NK cells^18^. To further investigate the effects of human NK cell-derived IFN-γ and AREG observed in vitro (Figure 3), we engrafted NCG mice (NOD/ShiLtJGpt-*Prkdc^em26Cd52^*Il2rg*^em26Cd^*^22^*/Gpt*), which lack NK, T and B cells, with keloid tissue obtained from clinical surgery and evaluated keloid restriction following intralesional injection of wild-type, IFNG knockout, or AREG knockout human NK cells (Figures 4A, S4A and S4B).

**Figure 4.**
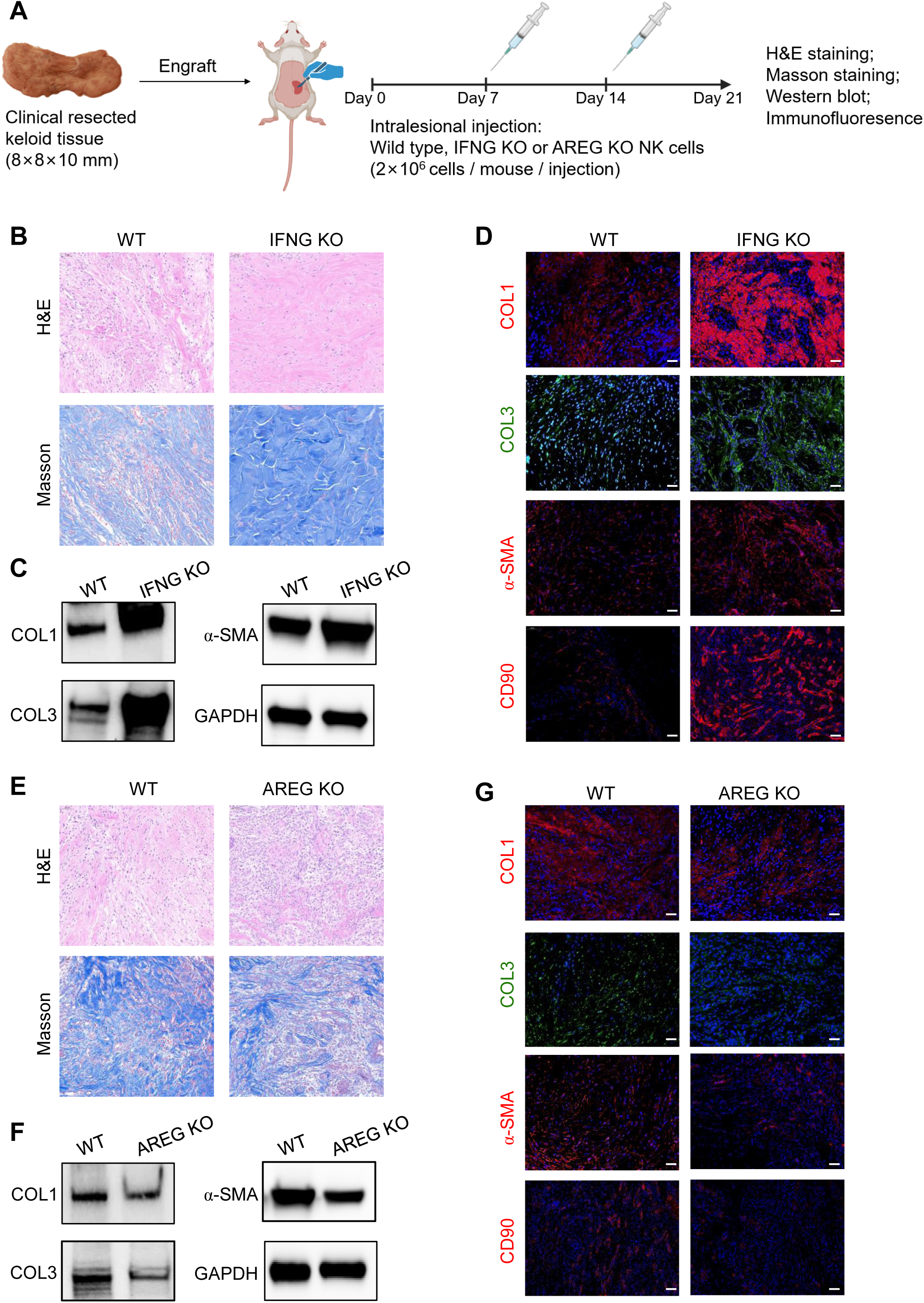
Differential effects of NK cell-derived IFN-γ and AREG on keloid progression in mouse xenografts. (A) Experimental schematic of keloid engraftment following human NK cell injection in NCG mice (n=5 for each group). (B) H&E and Masson staining of engrafted keloid tissues injected with wild-type or IFNG KO NK cells. (C) Western blot detection of COL1, COL3, α-SMA, and GAPDH in keloid tissues from (B). (D) Immunofluorescence detection of COL1, COL3, α-SMA, and CD90 in keloid tissues from (B). (E) H&E and Masson staining of engrafted keloid tissues injected with wild-type or AREG KO NK cells. (F) Western blot detection of COL1, COL3, α-SMA, and GAPDH in keloid tissues from (E). (G) Immunofluorescence detection of COL1, COL3, α-SMA, and CD90 in keloid tissues from (E).

Compared to keloid tissues injected with wild-type NK cells, those injected with IFNG knockout NK cells exhibited elongated, spindle-shaped fibroblasts, excessive collagen deposition, and a disorganized extracellular matrix, as shown by Hematoxylin and Eosin (H&E) and Masson staining (Figure 4B). Western blot and immunofluorescence analyses further revealed elevated levels of COL1, COL3, α-SMA, and CD90 in tissues injected with IFNG knockout NK cells (Figures 4C, 4D, and S4C-S4E). These findings underscore the critical role of NK cell-derived IFN-γ in limiting fibroblast pathology in keloids.

In contrast, keloid tissues injected with AREG knockout NK cells displayed reduced collagen deposition and a more organized fibroblast morphology compared to those injected with wild-type NK cells (Figure 4E). Additionally, levels of COL1, COL3, α-SMA, and CD90 were decreased (Figures 4F, 4G, and S4F-S4H), indicating that AREG knockout in NK cells enhances their ability to restrict keloid progression.

### Alterations in NK cell percentage and function in the blood of keloid patients

Blood is the major reservoir for NK cells recruited to inflamed skin^14^. To assess whether blood NK cells were associated with keloid pathology, we compared their percentage and cytokine production between healthy donors and keloid patients (Figures 5A and Table S1). The percentage and number of NK cells in the blood were similar between the two groups (Figure 5B and S5A). However, like in lesional skin, IFN-γ and TNF-α production by blood NK cells were downregulated in keloid patients compared to healthy donors (Figures 5C, 5D, S5B and S5C). While AREG^+^NK cells showed no difference between lesional and non-lesional skin (Figure 1E), they were upregulated in the blood of keloid patients compared to healthy donors (Figures 5E and S5D).

**Figure 5.**
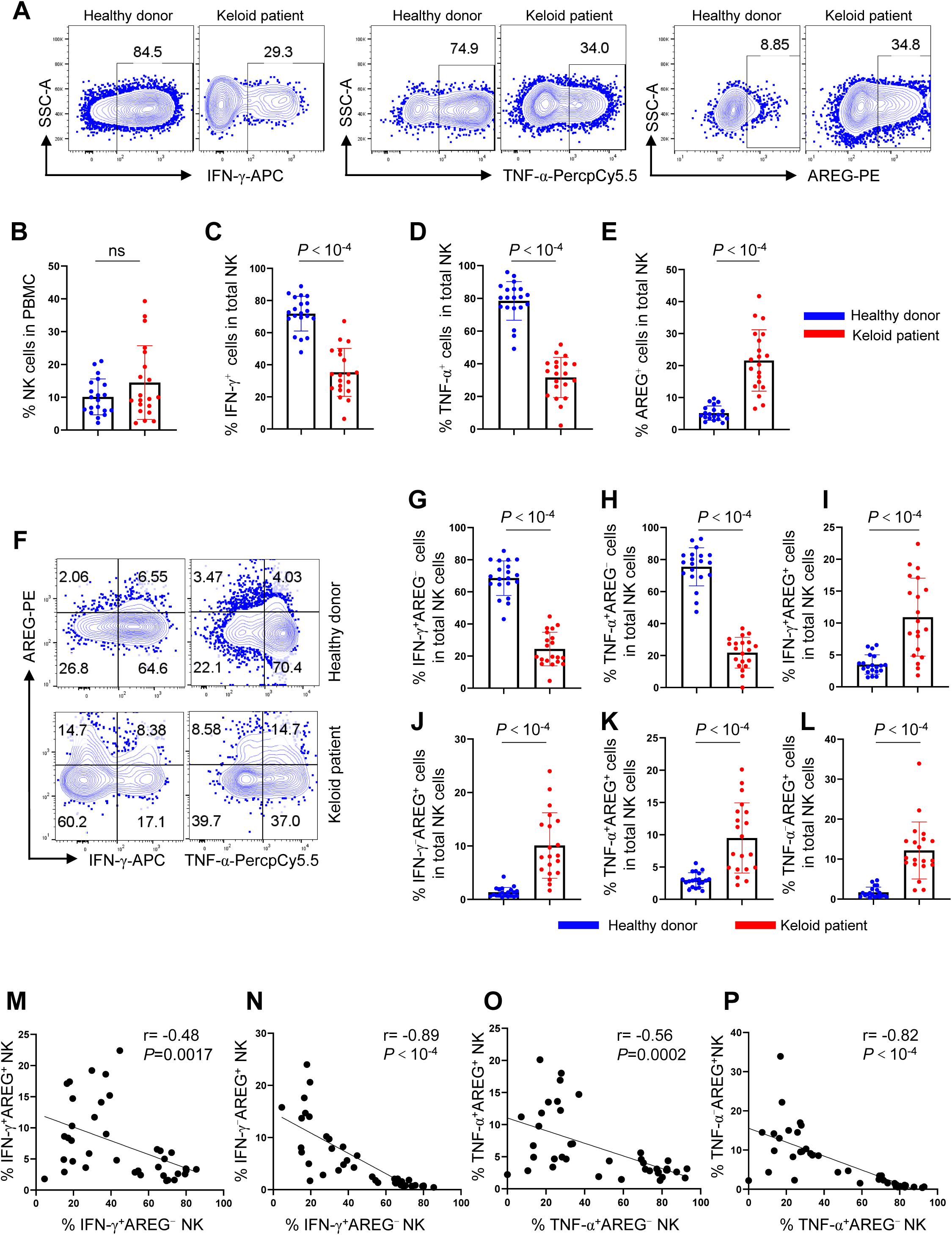
Alterations in NK cell frequency and function in keloid patient blood. (A) PBMCs from healthy and keloid individuals were stimulated with PMA and ionomycin for 2 hrs, and the production of IFN-γ, TNF-α, and AREG in NK cells was detected by flow cytometry. (B) Percentage of NK cells in PBMCs from healthy and keloid individuals (n=20). (C-E) Percentage of IFN-γ^+^ (C), TNF-α^+^ (D) and AREG^+^ (E) NK cells shown in (A) (n=20). (F) Flow cytometry detection of IFN-γ vs AREG or TNF-α vs AREG in NK cells in (A). (G-L) Percentage of IFN-γ^+^AREG^-^ (G), TNF-α^+^AREG^-^ (H), IFN-γ^+^AREG^+^ (I), IFN-γ^-^AREG^+^ (J), TNF-α^+^AREG^+^ (K), TNF-α^-^AREG^+^ (L) NK cells detected in (F) (n=20). (M-P) Correlation of IFN-γ^+^AREG^-^ NK cells with IFN-γ^+^AREG^+^ (M) (n=40) and with IFN-γ^-^AREG^+^ NK cells (N) (n=40); and correlation of TNF-α^+^AREG^-^NK cells with TNF-α^+^AREG^+^ (O) (n=40) and with TNF-α^-^AREG^+^ NK cells (P) (n=40). For (C-E, G-I, K), two-tailed unpaired t-test, for (B, J, L), Mann-Whitney test, data are mean with s.e.m., for (M-P), Spearman correlation, ns, not significant. Cell culture and stimulation conditions for each panel were detailed in Methods.

Blood NK cells with distinct cytokine production profiles were further characterized (Figure 5F). IFN-γ⁺AREG⁻NK cells, which showed the strongest inhibitory effect on fibroblast hyperproliferation and extracellular matrix production (Figures 3C-3F), were reduced in the blood of keloid patients (Figures 5G and S5E), with a similar trend observed for TNF-α⁺AREG⁻ NK cells (Figure 5H and S5F). In contrast, AREG⁺NK cells, regardless of IFN-γ or TNF-α co-expression, were increased (Figures 5I-5L and S5G-S5J). An inverse correlation between these differentially affected NK cell populations, similar to skin NK cells, was observed in the blood (Figures 5M-5P). This shift in cytokine production likely reflects the influence of keloid pathology on blood NK cells.

### ISG^+^NK cells represent a distinct NK cell subset in the blood of keloid patients

Single-cell sequencing was performed on magnetic beads enriched blood NK cells from healthy donors and keloid patients to further understand the impact of keloid pathology on blood NK cells (Methods, Figures S6A and 6A). NK cell identity was confirmed by the expression of NK cell markers and the absence of markers from other cell types (Figure S6B). A total of 16,558 high-quality NK cells were identified, which formed five distinct clusters in the uniform manifold approximation and projection (UMAP) analysis (Figures 6A).

**Figure 6.**
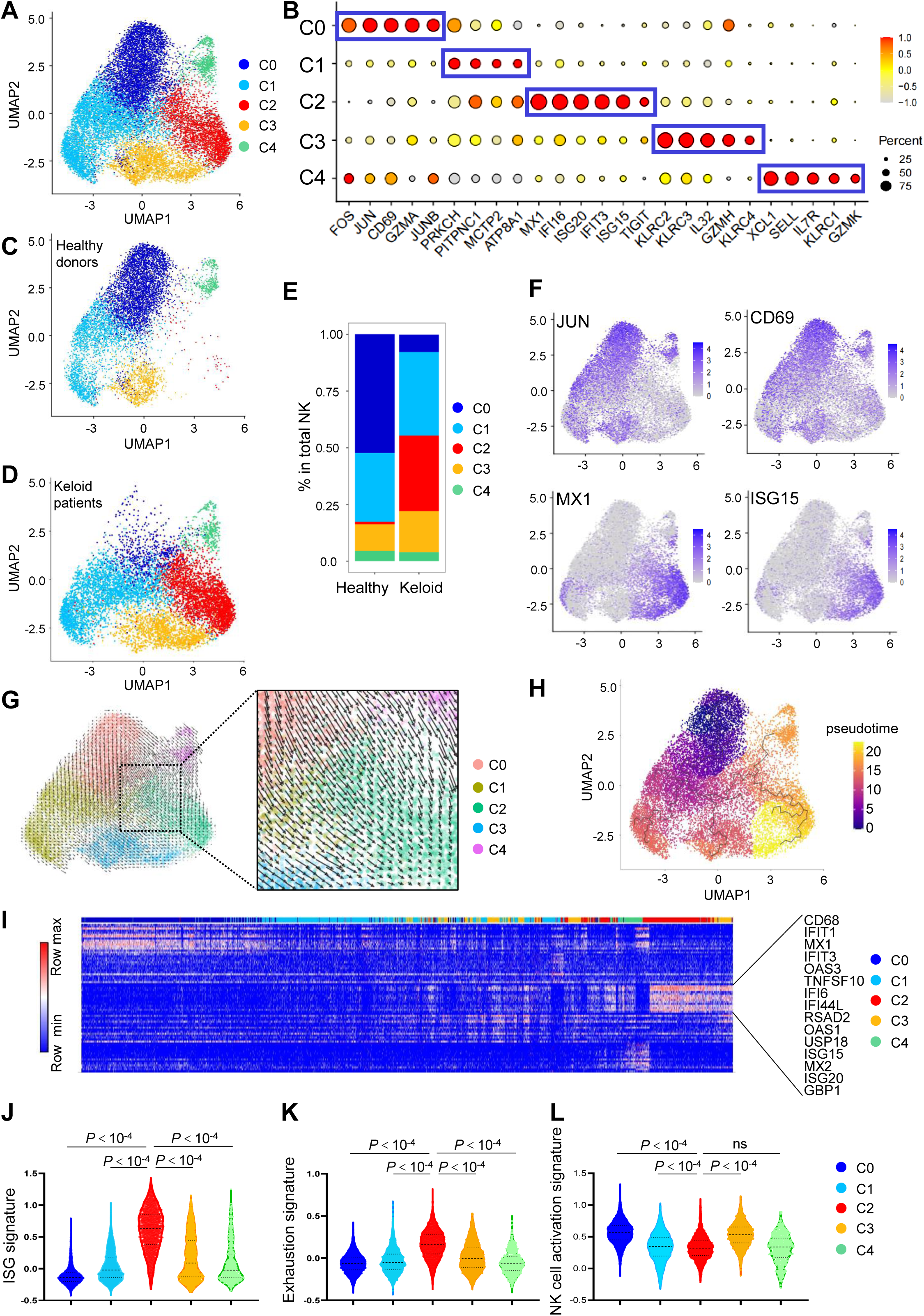
ISG^+^NK cells form a unique NK cell subset in keloid patient blood. (A) UMAP of blood NK cells from healthy donors and keloid patients. (B) Dotplot of highly expressed genes for each NK cell cluster. (C, D) UMAP of NK cells from healthy donors (C) and keloid patients (D). (E) Proportion of NK cell clusters in healthy donors and keloid patients. (F) Expression of JUN, CD69, MX1 and ISG15 in NK cell clusters. (G) RNA velocity of NK cell clusters projected on UMAP, with vectors indicating predicted gene expression trajectories (left) and a magnified view of the selected region, highlighting finer RNA velocity dynamics within clusters (right). (H) Pseudotime analysis of NK cell clusters using Monocle 3. Distinct cell clusters are color-coded by transcriptional state, with black lines representing inferred pseudotime trajectories. (I) Heatmap showing the top 15 upregulated DE genes for each NK cell cluster along the pseudotime trajectory. (J-L) Gene set score analysis of ISG (J), exhaustion (K), and NK cell activation (L) signatures across NK cell clusters. Gene set score was calculated by AddMouduleScore in Seurat, the *P* value was determined by Mann-Whitney test.

Cluster 0 (C0) NK cells exhibited high expression of the tissue resident marker CD69 and AP-1 family members (FOS, JUN, JUNB). Cluster 1 (C1) NK cells showed high expression of PRKCH and PITPNC1, which are associated with regulating NK cell and CD8^+^T cell effector functions^35,36^. Cluster 2 (C2) NK cells were distinguished by high expression of interferon-stimulated genes (ISGs), such as MX1, IFIT3, ISG15, and ISG20, along with TIGIT, which is expressed on exhausted NK and T cells^37^. Cluster 3 (C3) NK cells expressed high levels of NK cell activating receptors (KLRC2, KLRC3, KLRC4) and the pro-inflammatory cytokine IL-32. Cluster 4 (C4) NK cells, representing CD56^hi^NK cells, displayed high expression of SELL, IL7R, GZMK, XCL1, and GZMK (Figures 6B and Table S3)^21,22^. Notably, C2 NK cells, characterized by high ISG expression, were almost exclusively derived from the blood of keloid patients, whereas C0 NK cells, marked by high expression of CD69 and AP-1 family genes, were predominantly from healthy individuals (Figures 6C-6F and Table S3).

To gain insights into the cellular dynamics of NK cell clusters under steady state and how they might be affected by keloid pathology, RNA velocity analysis was performed^38^. The results revealed a clear directional flow from C0 (dominant in healthy donors), as well as C1 and C4 NK cells, toward C2 NK cells (dominant in keloid patients) (Figure 6G). Consistent with this, pseudotime analysis placed C0 NK cells at the initial stage and C2 NK cells at the terminal stage along the trajectory (Figure 6H)^39,40^. A heatmap of pseudotime-ordered single cells showed that C2 NK cells, enriched with ISGs, clustered distinctly at the terminal stage (Figure 6I). Gene set score analysis revealed that C2 NK cells exhibited the highest ISG and exhaustion signatures^41^ among all NK cell clusters, while displaying the lowest NK cell activation signatures among CD56^dim^NK cell clusters (C0-C3) (Figures 6J-6L and Table S4). Additionally, all NK cell clusters from keloid patients displayed higher ISG and exhaustion signatures, alongside lower activation signatures, compared to those from healthy donors (Figures S6C-S6E). Moreover, commonly upregulated genes in each NK cell cluster from keloid patients were enriched with type 1 IFN response pathways (Figure S6F), suggesting that the type 1 IFN-associated cytokine milieu in the blood of keloid patients may drive normal NK cells toward a specialized ISG^+^NK population with an exhausted phenotype.

### Elevated IFN-β in keloid patient blood is responsible for increased ISG^+^NK cells and downregulated IFN-γ production

Consistent with gene set score analysis (Figure 6J), C2 NK cells exhibited the highest expression of type 1 IFN receptors (IFNAR1, IFNAR2) and downstream transcription factors (IRF1, IRF7, IRF9, STAT1, STAT2), confirming their enhanced responsiveness to type 1 IFNs (Figure 7A). Conversely, C2 NK cells had the lowest IFNG expression across all NK cell clusters (Figure 7A), indicating that ISG^+^NK cells derived from the type 1 IFN-enriched environment in keloid patients exhibit impaired IFN-γ production. Indeed, IFN-β induced significant upregulation of ISG15 in NK cells, with ISG15^+^NK cells showing reduced IFN-γ production compared to ISG15^-^NK cells (Figures 7B and 7C).

**Figure 7.**
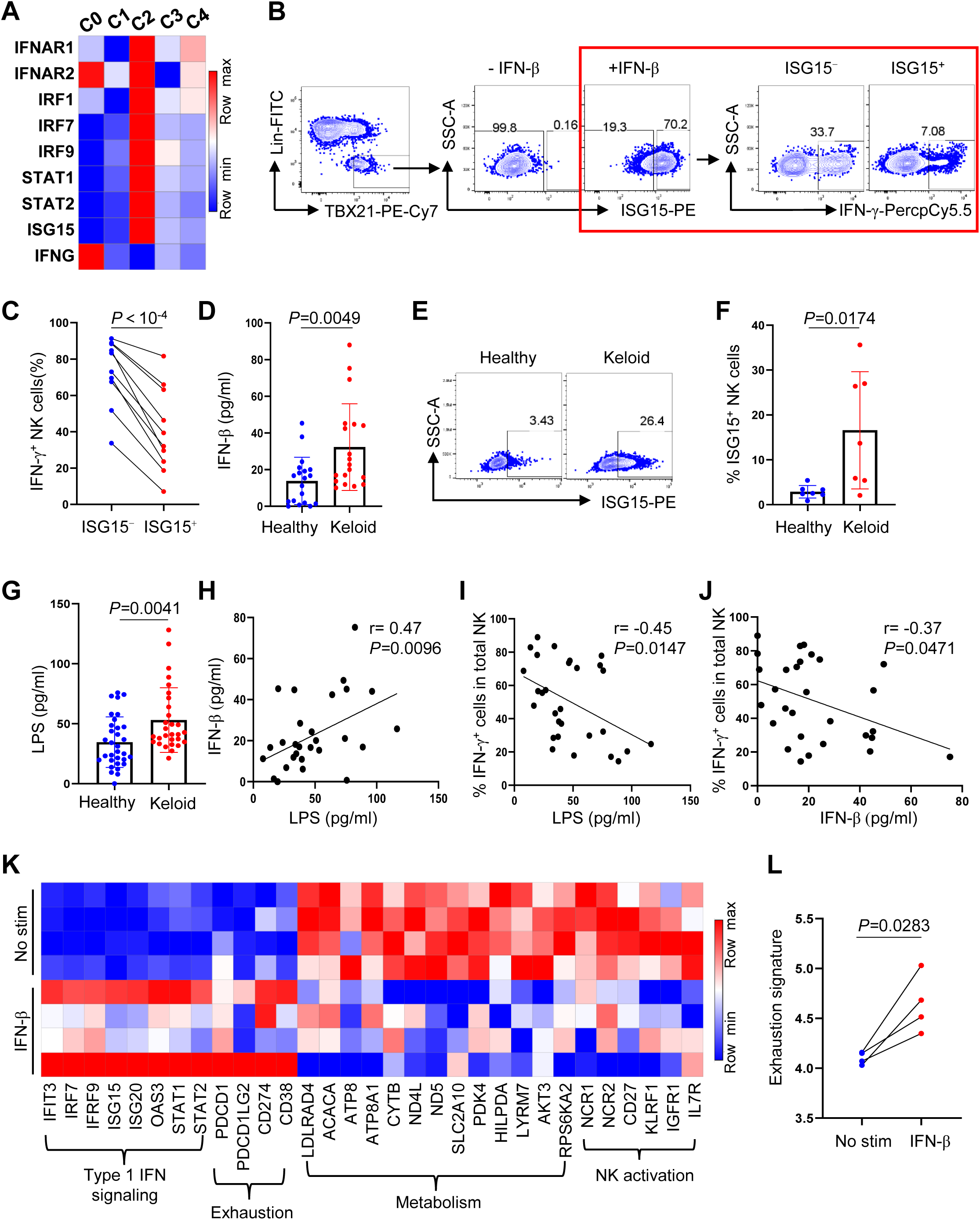
Elevated IFN-β in keloid patient blood drives increased ISG^+^NK cells and reduced IFN-γ production. (A) Heatmap of type I IFN signaling associated genes and IFNG across NK cell clusters. (B) PBMCs from healthy donors were pre-treated with IFN-β for 16 hours, followed by stimulation with IL-12, IL-15, and IL-18 for 16 hours. IFN-γ production in ISG15^-^ and ISG15^+^ NK cells was measured by flow cytometry. (C) Percentage of IFN-γ^+^ cells in ISG15^-^ and ISG15^+^ NK cells in (B) (n=10). (D) Plasma concentration of IFN-β in healthy donors and keloid patients measured by ELISA (n=19). (E) PBMCs from healthy donors and keloid patients were incubated with golgi block for 6 hrs, and ISG15 in NK cells was measured by flow cytometry. (F) Percentage of ISG^+^NK cells in (E) (n=7). (G) Plasma concentration of LPS in healthy donors (n=32) and keloid patients (n=29) measured by ELISA. (H, I) Correlation of plasma LPS concentration with IFN-β levels (H, n=29) and IFN-γ^+^NK cells (I, n=29). (J) Correlation of plasma IFN-β concentration with IFN-γ^+^NK cells (n=29). (K) Heatmap of DEgenes associated with type 1 IFN signaling, exhaustion, metabolism and NK cell activation in NK cells cultured with or without IFN-β for 7 days (n=4). (L) Gene set score analysis of exhaustion signature of NK cells cultured with or without IFN-β in (K). For (C, L), two-tailed paired t-test, for (D, F), two-tailed unpaired t-test, for (G), Mann-Whitney test, data are mean with s.e.m., for (H-J), Spearman correlation, ns, not significant. Cell culture and stimulation conditions for each panel were detailed in Methods.

Plasma levels of IFN-β were consistently elevated in keloid patients (Figure 7D), who also showed a higher percentage of ISG15^+^NK cells without type 1 IFN stimulation compared to healthy donors (Figures 7E and 7F). This increase in IFN-β is likely attributable to systemic inflammation, as keloid patients exhibited elevated plasma LPS levels (Figure 7G), which positively correlated with IFN-β levels (Figure 7H). Importantly, both plasma LPS and IFN-β levels negatively correlated with the IFN-γ productivity of NK cells (Figures 7I and 7J). Bulk RNA-Seq analysis revealed that IFN-β-treated NK cells displayed an exhausted transcriptional profile, characterized by upregulation of exhaustion genes^42,43^, and downregulation of genes associated with metabolic fitness^44–48^, NK cell activation^49–51^, and proliferation (Figures 7K, 7L and Table S5). These findings highlight the significant impact of type 1 IFN on normal NK cell physiology in the blood of keloid patients.

### Type 1 IFN induces NK cell exhaustion by disrupting mitochondrial function and metabolism

We hypothesized that elevated type 1 IFN in the blood of keloid patients induces functional exhaustion of NK cells based on the following evidence: (1) IFN-β induces ISG15 expression in NK cells, and ISG15^+^NK cells demonstrate reduced IFN-γ production compared to ISG15^-^NK cells (Figures 7B and 7C); (2) plasma IFN-β levels negatively correlate with the percentage of IFN-γ^+^ NK cells (Figure 7J); (3) ISG15^+^NK cells enriched in the blood of keloid patients, or NK cells treated with IFN-β, exhibit an exhaustion-associated transcriptional profile (Figures 6K, 7K, and 7L); and (4) NK cells from keloid patients display reduced IFN-γ production (Figure 5C).

Indeed, IFN-β treatment increased the proportion of TIGIT^+^NK cells while reducing the percentage of IFN-γ^+^NK cells (Figures 8A–8C). Consistent with these results in PBMCs, we observed elevated TIGIT expression, along with suppressed IFN-γ production and reduced proliferation, in magnetic bead-enriched NK cells exposed to IFN-β (Figures 8D–8I). Notably, knockout of type 1 IFN signaling components (IFNAR1, IRF1, and IRF9) in NK cells reversed these IFN-β-induced changes (Figures 8J–8O and S7A-S7C). These findings demonstrate that IFN-β drives an exhausted phenotype in NK cells.

**Figure 8.**
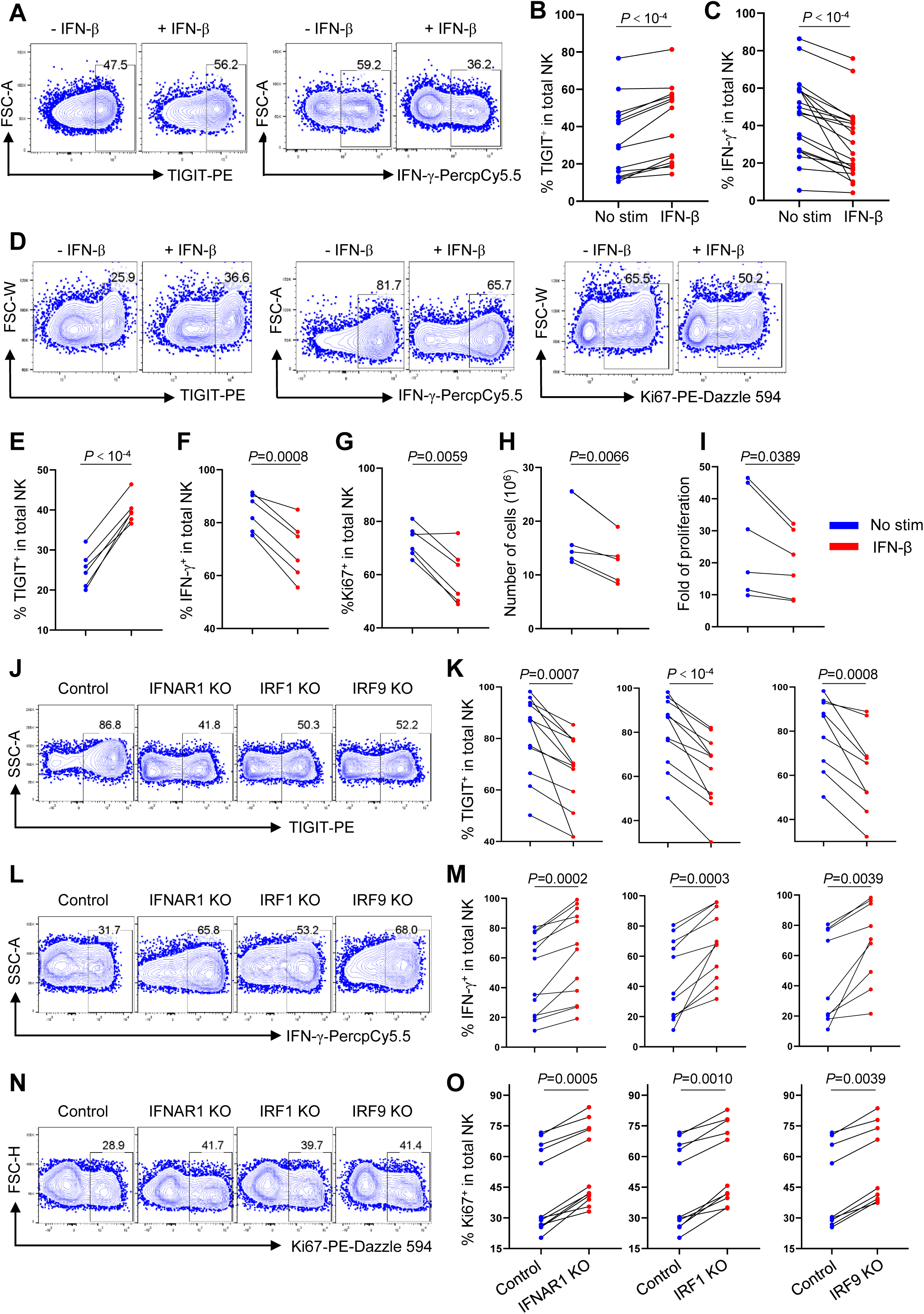
Type 1 IFN signaling induces an exhausted phenotype in NK cells. (A) PBMCs from healthy donors were cultured with or without IFN-β for 7 days, then stimulated with IL-12, IL-15, and IL-18 for 16 hrs. TIGIT and IFN-γ in NK cells were measured by flow cytometry. (B, C) Percentage of TIGIT^+^ (B, n=14) and IFN-γ^+^ (C, n=20) NK cells in (A). (D) Enriched NK cells from healthy donors were cultured with or without IFN-β for 7 days, then stimulated with IL-12, IL-15, and IL-18 for 4 hrs. TIGIT, IFN-γ and Ki67 in NK cells were detected by flow cytometry. (E-I) Percentage of TIGIT^+^ (E), IFN-γ^+^ (F), Ki67^+^ (G) NK cells, as well as NK cell counts (H) and fold of proliferation (I) after 7 days culture in (D) (n=6). (J-O) Control (targeting CD19), IFNAR1, IRF1 and IRF9 knockout NK cells were cultured in presence of IFN-β for 7 days. TIGIT (J, K), IFN-γ (L, M) and Ki67 (N, O) in NK cells were assessed by flow cytometry (IFNAR1 KO, n=12; IRF1 KO, n=11; IRF9 KO, n=9). For (B, C, E-I, K, M: IFNAR1 KO and IRF1 KO), two-tailed paired t-test, for (O, M: IRF9 KO), Wilcoxon matched-pairs signed rank test, data are mean with s.e.m.. Cell culture and stimulation conditions for each panel were detailed in Methods.

Gene set variation analysis (GSVA), in combination with differentially expressed gene analysis, revealed a downregulation of metabolic pathways and genes related to glycolysis and oxidative phosphorylation in ISG^+^ or IFN-β-treated NK cells compared to ISG^-^ or IFN-β-untreated NK cells (Figures S7D and 7K). Accordingly, Seahorse assays showed a mild but significant reduction in basal and compensatory glycolysis in NK cells treated with IFN-β under unstimulated conditions (without IL-12+IL-15+IL-18) (Figures 9A-9C). Moreover, IFN-β treatment decreased both basal and maximal oxygen consumption rates (OCR) in NK cells, with or without IL-12+IL-15+IL-18 stimulation (Figures 9D-9F). Consequently, mitochondrial ATP production and spare respiratory capacity (SRC) were impaired in IFN-β-treated NK cells (Figures 9G and 9H). Electron microscopy further revealed significant mitochondrial abnormalities, including reduced mitochondrial numbers and enlarged or elongated morphology (Figure 9I-9K). These findings suggest that IFN-β induces mitochondrial dysfunction and contributes to NK cell exhaustion by decreasing their metabolic activity.

**Figure 9.**
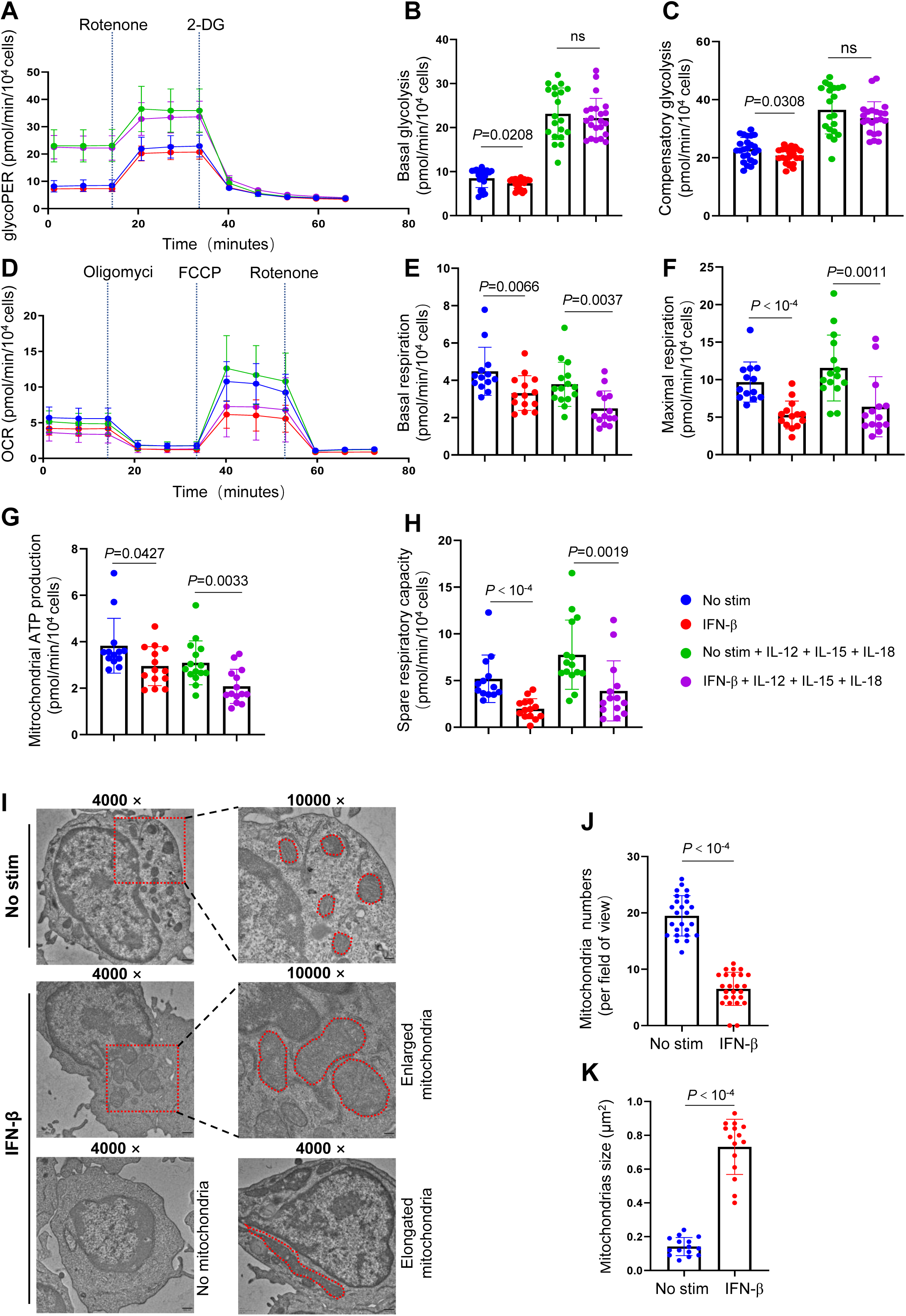
Type 1 IFN induces NK cell exhaustion by impairing mitochondrial function and metabolism (A-C) Glycolytic proton efflux rate (glycoPER) of enriched NK cells cultured in NK MACS medium alone (n=24) or supplemented with IFN-β (n=24), IL-12/15/18 (n=20), or IFN-β + IL-12/15/18 (n=22), measured by Seahorse. (B, C) Basal (B) and compensatory (C) glycolysis were calculated from (A). (D) Oxygen consumption rate (OCR) of enriched NK cells cultured in NK MACS medium alone (n=13) or supplemented with IFN-β (n=14), IL-12/15/18 (n=15), or IFN-β + IL-12/15/18 (n=14), measured by Seahorse. (E-H) Basal (E) and maximal (F) respiration, mitochondrial ATP production (G) and spare respiratory capacity (H) were calculated from (D). (I) Transmission electron microscopy of NK cells treated with or without IFN-β for 7 days. The black scale bar in the bottom right corner represents 500 nm at 4000x and 200 nm at 10000x. For (B, C, J, K), two-tailed unpaired t-test, for (E-H), Mann-Whitney test, data are mean with s.e.m., ns, not significant. Cell culture and stimulation conditions for each panel were detailed in Methods.

## Discussion

The crosstalk between immune and non-immune cells in the skin plays a critical role in the progression of cutaneous disorders^52,53^. In keloid lesions, NK cells produced IFN-γ, which inhibits fibroblast proliferation and extracellular matrix secretion was reduced (Figures 1C, 3 and 4). This reduction is likely due to increased TGF-β from lesional fibroblasts (Figures 2F-2K). In contrast, AREG levels, which counteract the effect of IFN-γ, remain unchanged (Figures 1E, 3 and 4), suggesting that the increased TGF-β in keloid skin disrupts the equilibrium between IFN-γ and AREG production by NK cells. Targeting TGF-β signaling has proven effective in fibrosis-related diseases^25^, and a combined strategy that enhances NK cell IFN-γ production while reducing AREG levels may offer additional therapeutic benefits for keloid treatment.

We observed decreased IFN-γ production by NK cells in keloid lesional skin compared to non-lesional skin, and in the blood of keloid patients versus healthy donors (Figures 1C and 5C). In contrast, AREG production by NK cells was elevated only in the blood of keloid patients (Figure 5E). Since blood is the primary source of skin NK cells^14,54^, these findings indicate that pathological changes in NK cells may be already present even in non-lesional areas of keloid skin, and that this localized immune dysregulation could contribute to keloid development.

The elevated plasma concentrations of IFN-β and LPS in keloid patients suggest systemic inflammation (Figures 7D and 7G), potentially linked to compromised gut epithelium integrity and bacterial translocation into the bloodstream^21,55^. It would be intriguing to determine whether immune cells that support gut homeostasis, such as Th17 cells and ILC3s, which secrete IL-22 to maintain gut epithelium integrity and induce antimicrobial peptides^21,56^, are downregulated in the intestines of keloid patients, and whether such a disturbance in gut homeostasis is associated with keloid pathogenesis.

The exhausted phenotype of NK cells is observed in various pathological conditions, including tumors^37,57^, metabolic diseases^58^, and viral infections^59,60^, often linked to hypoxia, inhibitory ligands, metabolic dysregulation, and mitochondrial defects^61,62^. However, the characteristics and causes of NK cell exhaustion in keloids remain unclear. The finding that elevated IFN-β in the plasma of keloid patients induces NK cell exhaustion through mitochondrial defects and metabolic dysfunction may represent a common mechanism (Figure 9). Reducing systemic inflammation or inhibiting type 1 IFN signaling could help sustain NK cell effectiveness, potentially benefiting keloid treatment and enhancing NK cell-based immunotherapies.

## Supporting information

suppemental tables

## Acknowledgments

This study was supported by National Natural Science Foundation of China (82471876), CAMS Innovation Fund for Medical Sciences (CIFMS) (2024-I2M-3-005) and Hospital for Skin Diseases, Institute of Dermatology, Chinese Academy of Medical Sciences and Peking Union Medical College grant (3301030103119) to Yetao W. National key research and development program (2022YFC2504700, 2022YFC2504701, 2022YFC2504705) to Yan W. All staff at the biobank of Institute of Dermatology, Chinese Academy of Medical Sciences, Jiangsu Biobank of Clinical Resources assisted with clinical sample collection.

## Author contributions

Yetao Wang: conceptualization; experiment design; data analysis; resources; supervision; validation; investigation; visualization; methodology; data curation; software; writing – original draft, review and editing; project administration; funding acquisition. Ying Zhao: experiment design; performing experiments; data analysis; validation; visualization; methodology; data curation; writing – methods, review and editing. Qin Wei: data analysis; methodology. Rui zeng: data analysis. Yan Wang: resources. Yong Yang: resources.

## Declaration of interests

All authors declare no competing interests.

## Supplemental figure legends

**Figure S1.**
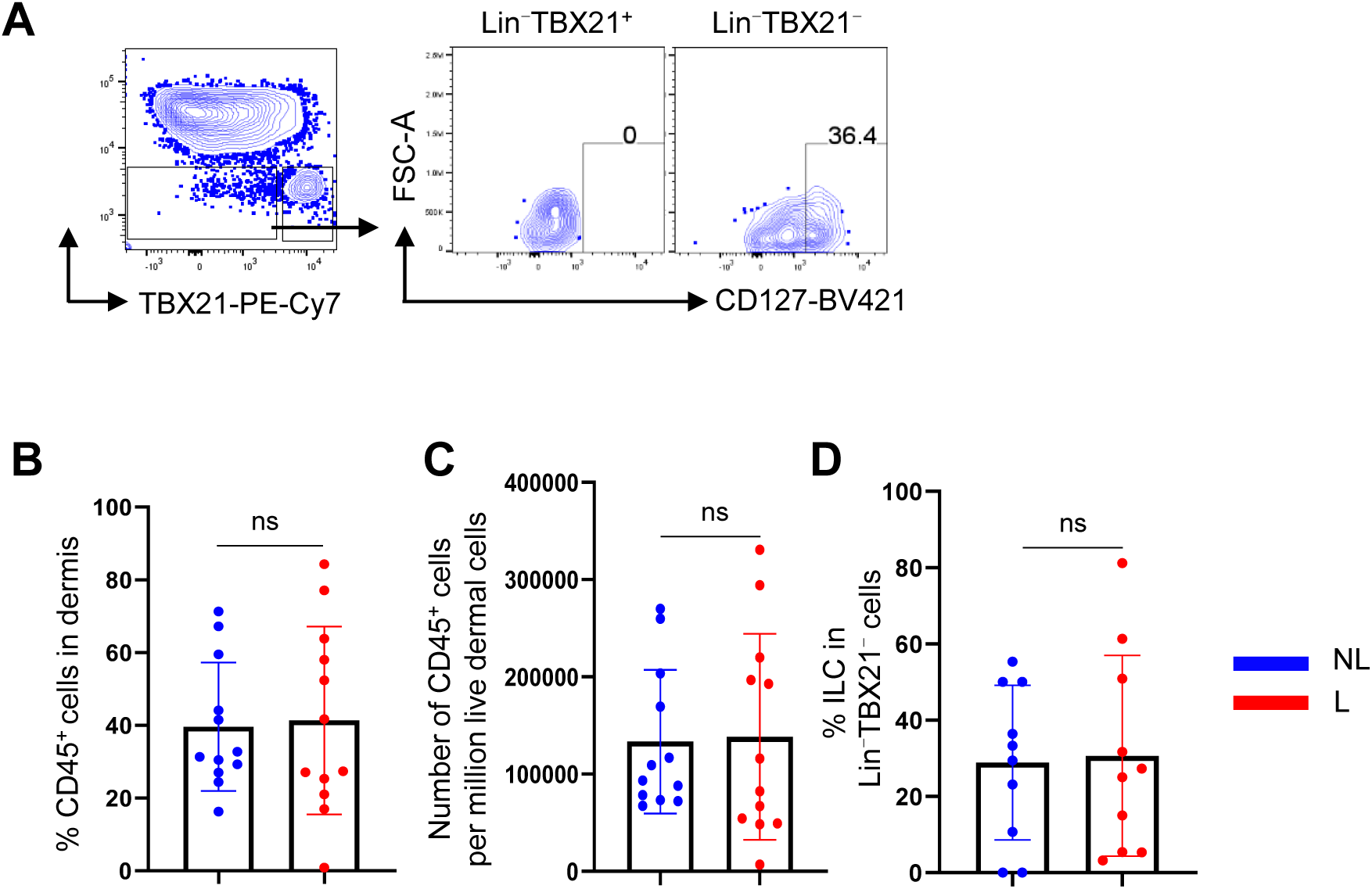
Detection of ILCs and CD45^+^ cells in keloid NL and L dermis. (A) Percentage of CD127^+^ cells in CD45^+^Lin^-^TBX21^+^ and CD45^+^Lin^-^TBX21^-^ cells in dermis detected by flow cytometry. (B, C) Percentage (B) and number (C) of CD45^+^ cells in keloid NL and L dermis (n=12). (D) Percentage of ILCs in CD45^+^Lin^-^TBX21^-^ cells in keloid NL and L dermis (n=10). For (B-D), two-tailed paired t-test, data are mean with s.e.m., ns, not significant.

**Figure S2.**
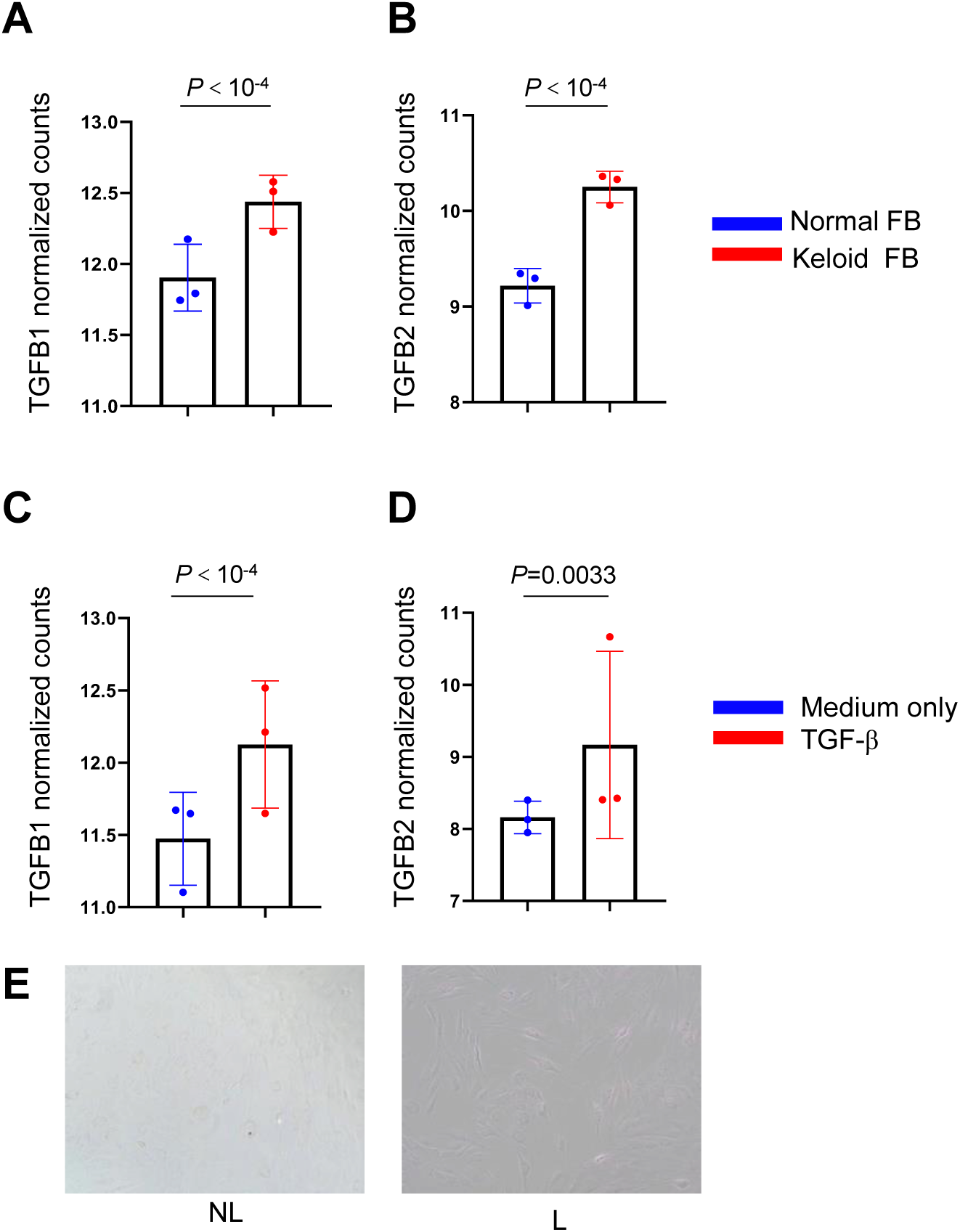
Detection of TGFB1/2 expression and microscopy of keloid NL and L fibroblasts cultured with NK cell-conditioned medium. (A, B) Normalized counts of TGFB1 (A) and TGFB2 (B) in fibroblast from healthy donors and keloid patients (n=3), based on GSE117887. (C, D) Normalized counts of TGFB1 (A) and TGFB2 (B) in fibroblast from healthy donors treated with or without TGF-β, based on GSE222916. (E) Microscopy of keloid NL and L fibroblasts co-cultured with NK cells stimulated with IL-12/15/18 for 48 hrs. Cell culture and stimulation conditions for each panel were detailed in Methods. For (A-D), *P* value was determined by DESeq2.

**Figure S3.**
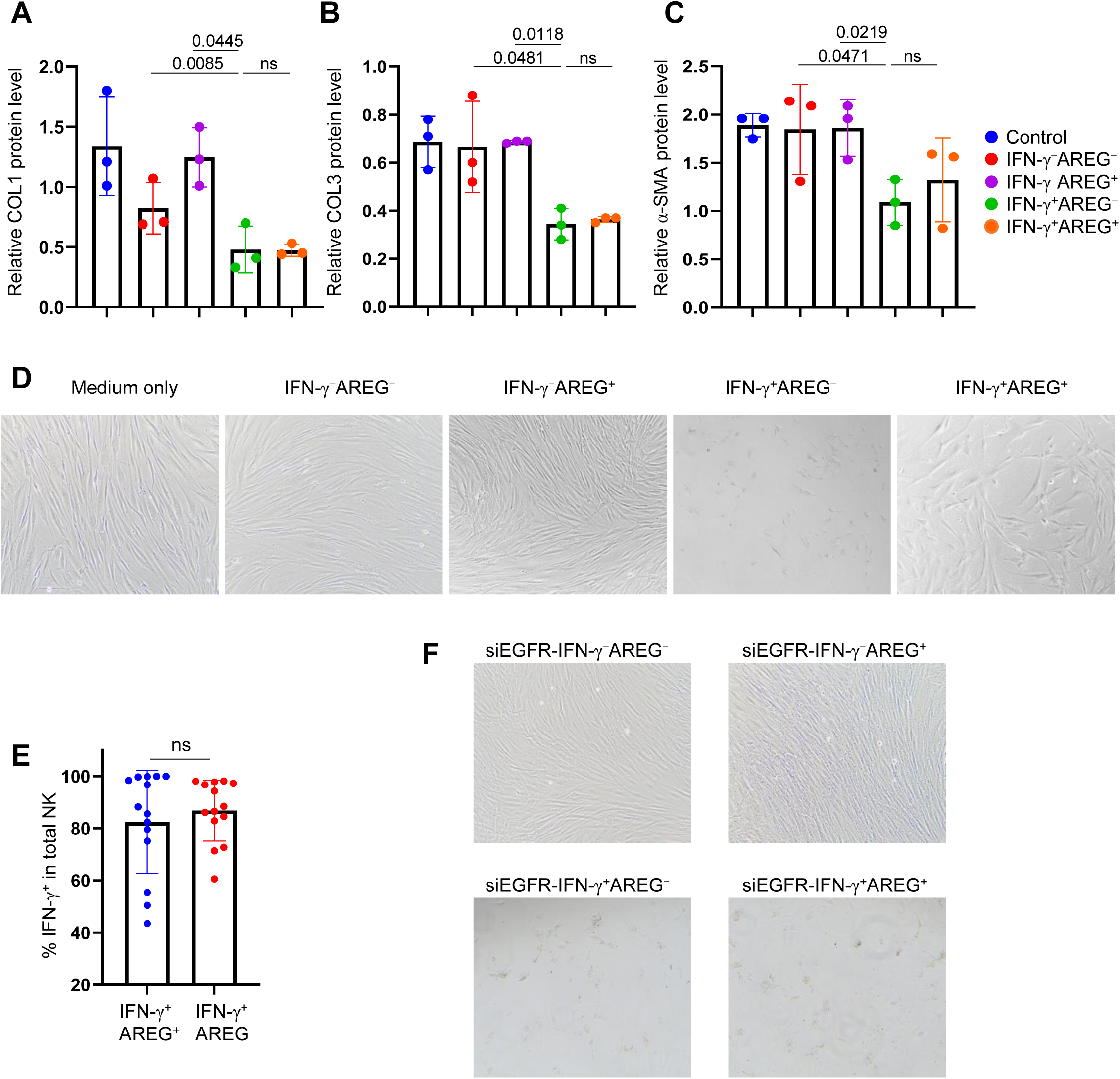
Effects of NK cell derived IFN-γ and AREG on keloid fibroblasts. (A-C) Relative protein levels of COL1 (A), COL3 (B), and α-SMA (C) normalized to GAPDH in keloid lesional fibroblasts cultured with IFN-γ^-^AREG^-^, IFN-γ^+^AREG^-^, IFN-γ^-^AREG^+^ or IFN-γ^+^AREG^+^ NK cell conditioned medium (n=3). (D) Microscopy of keloid lesional fibroblasts under indicated conditions. (E) Percentage of IFN-γ^+^ NK cells from in vitro generated IFN-γ^+^AREG^+^ and IFN-γ^+^AREG^-^ NK cells (n=14). (F) Microscopy images of keloid lesional fibroblasts subjected to EGFR knockdown and treated with conditioned medium as indicated. For (A-C, E), two-tailed paired t-test, data are mean with s.e.m., ns, not significant. Cell culture and stimulation conditions for each panel were detailed in Methods.

**Figure S4.**
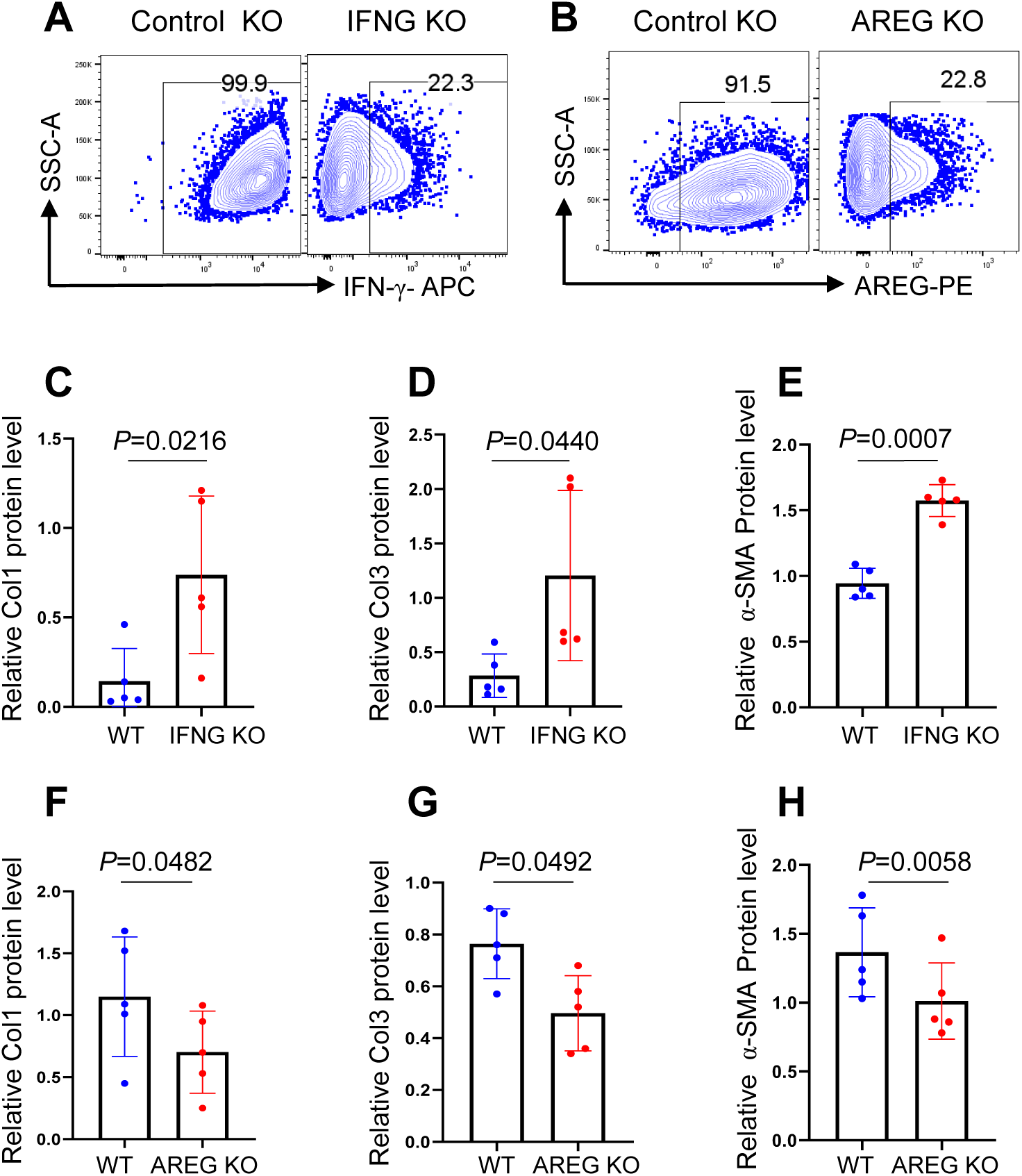
Effect of NK cell IFNG and AREG knockout on keloid progression in mouse xenografts. (A, B) Flow cytometry of IFN-γ (A) and AREG (B) in control, IFNG and AREG knockout NK cells. (C-E) Relative protein levels of COL1 (C), COL3 (D), and α-SMA (E) normalized to GAPDH in keloid tissues injected with wild-type and IFNG KO NK cells (n=5). (F-H) Relative protein levels of COL1 (F), COL3 (G), and α-SMA (H) normalized to GAPDH in keloid tissues injected with wild-type and AREG KO NK cells (n=5). For (C-H), two-tailed paired t-test, data are mean with s.e.m., ns, not significant.

**Figure S5.**
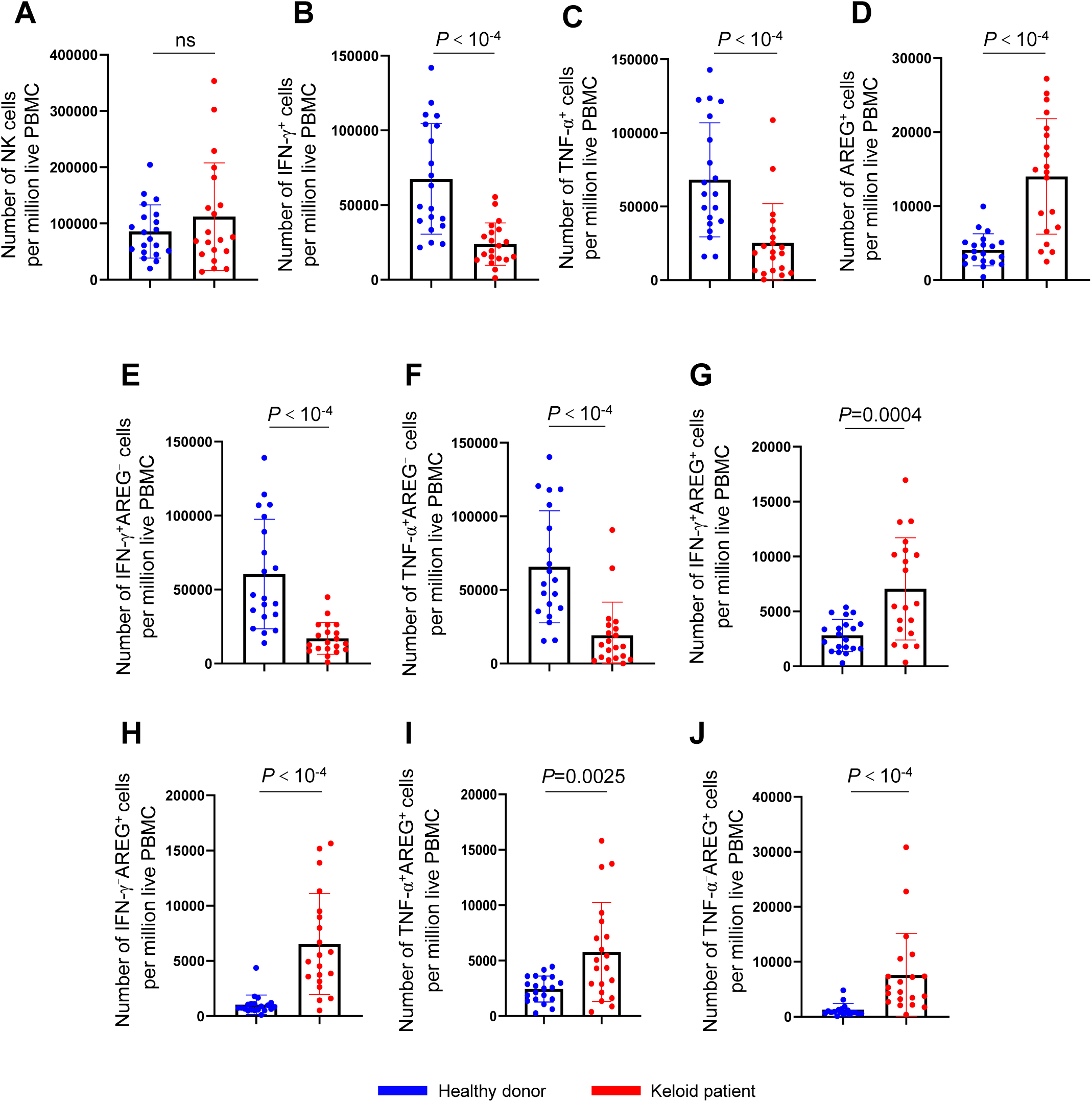
Detection of blood NK cells from keloid patients and healthy donors. (A-J) Number of the total NK cells (A), IFN-γ^+^ (B), TNF-α^+^ (C), AREG^+^ (D), IFN-γ^+^AREG^-^ (E), TNF-α^+^AREG^-^ (F), IFN-γ^+^AREG^+^ (G), IFN-γ^-^AREG^+^ (H), TNF-α^+^AREG^+^ (I) and TNF-α^-^AREG^+^ (J) NK cells in one million live PBMCs from healthy donors and keloid patients (n=20). For (A, C, F, H, J), Mann-Whitney test, for (B, D, E, G, I), two-tailed unpaired t-test, data are mean with s.e.m., ns, not significant.

**Figure S6.**
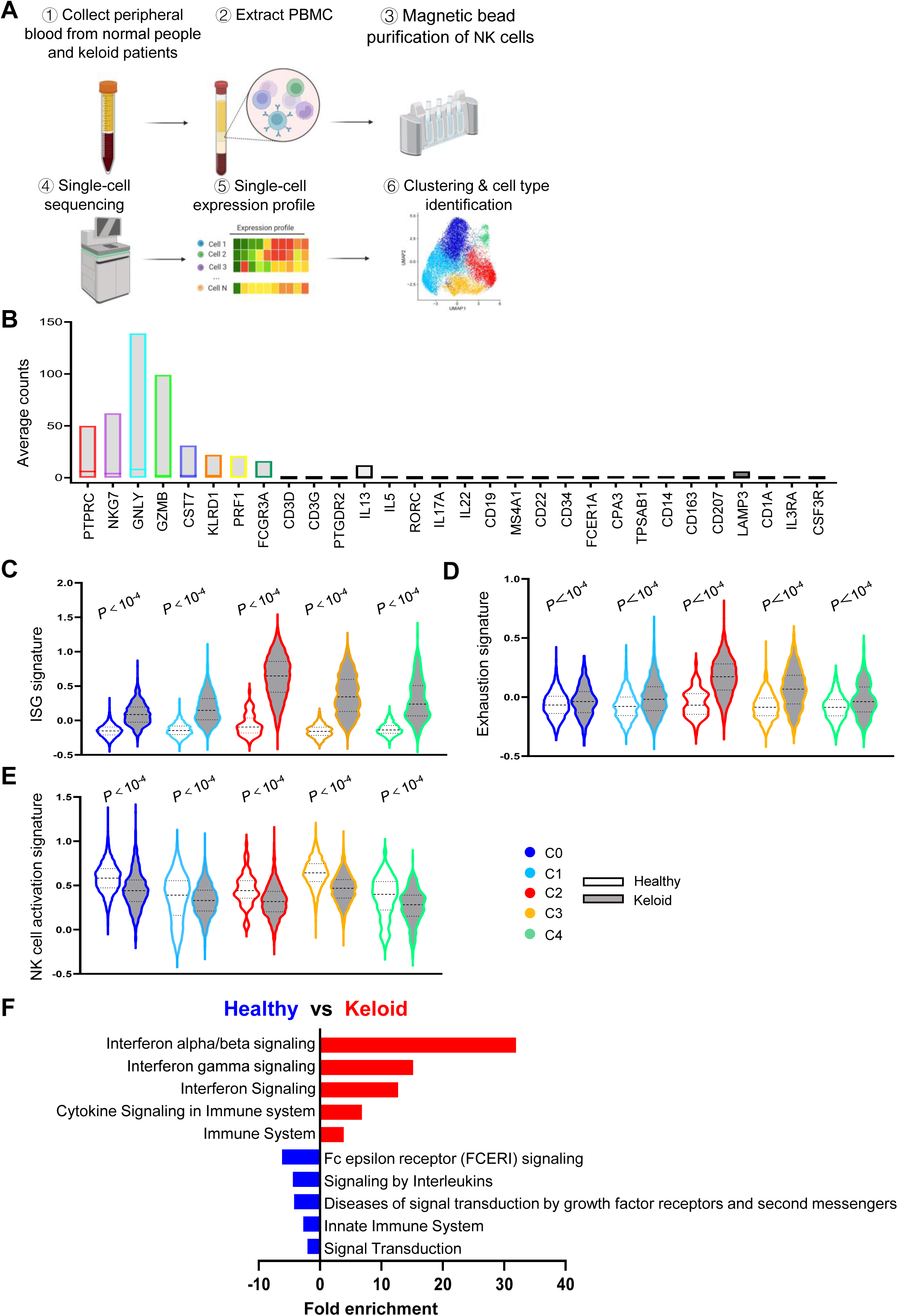
Single-cell sequencing of blood NK cells derived from keloid patients and healthy donors. (A) Experimental design for single-cell sequencing of enriched NK cells isolated from PBMCs of healthy individuals and keloid patients, with each group pooled from 10 individuals. (B) Expression of markers for NK cells and other cell types in the identified NK cell cluster. (C-E) Gene set score analysis of signatures of ISG (C), Exhaustion (D) and NK cell activation (E) of NK clusters in healthy donors and keloid patients. Gene set score was calculated by AddMouduleScore in Seurat, the *P* value was determined by Mann-Whitney test. (F) GO enrichment analysis for commonly upregulated (red) and downregulated (blue) genes in 0-4 NK clusters in keloid.

**Figure S7.**
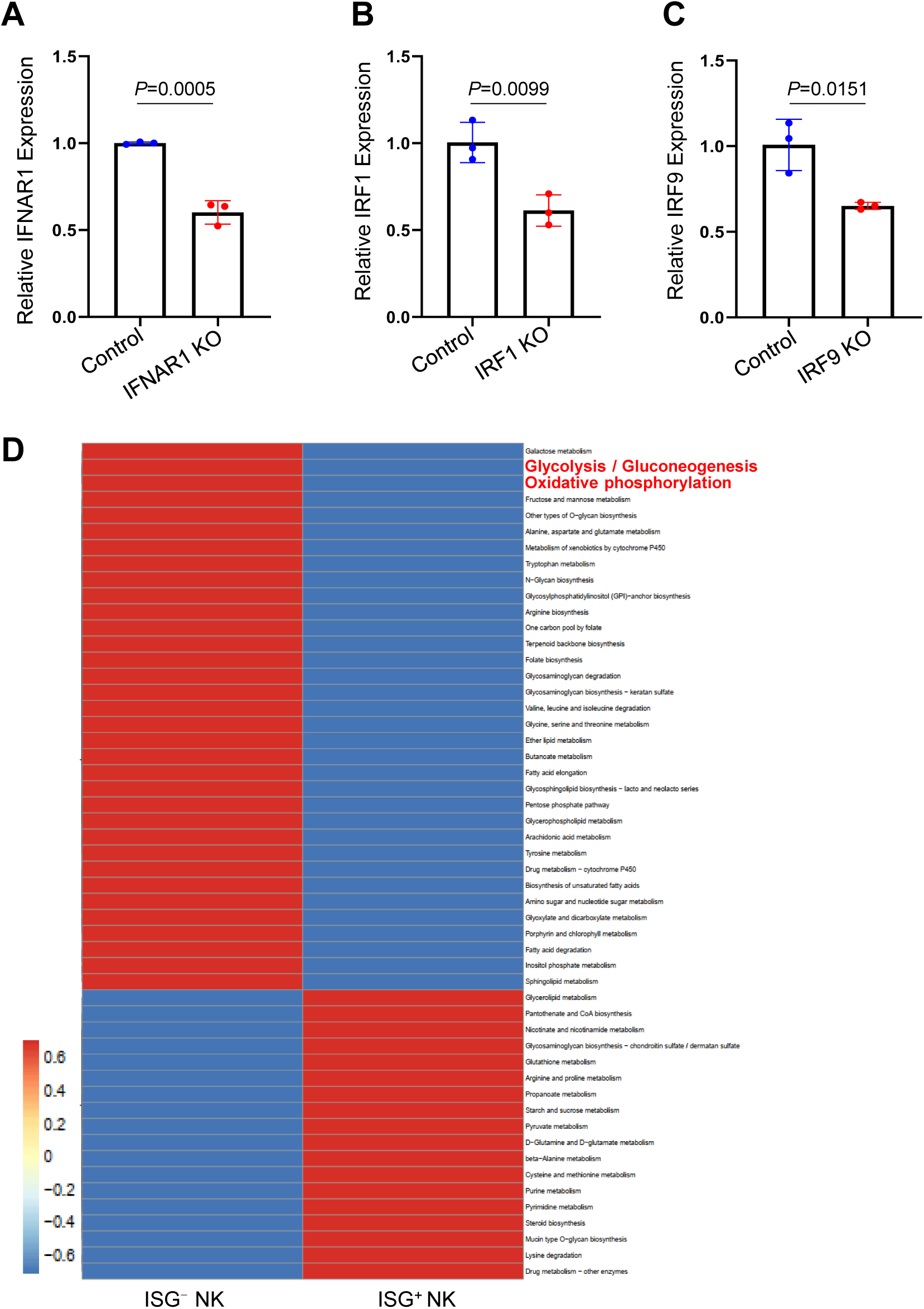
Confirmation of type 1 IFN signaling component knockout and metabolic differences between ISG^+^ and ISG^-^ NK cells. (A-C) Expression of IFNAR1 (A), IRF1 (B), and IRF9 (C) relative to β-actinin in IFNAR1, IRF1, and IRF9 knockout NK cells, measured by quantitative reverse transcription PCR. NK cells electroporated with sgRNA targeting CD19 served as controls (n=3). Metabolic pathway enrichment analysis of ISG^+^NK (C2) versus ISG^-^NK (other NK cell clusters) using scMeta.

## Methods

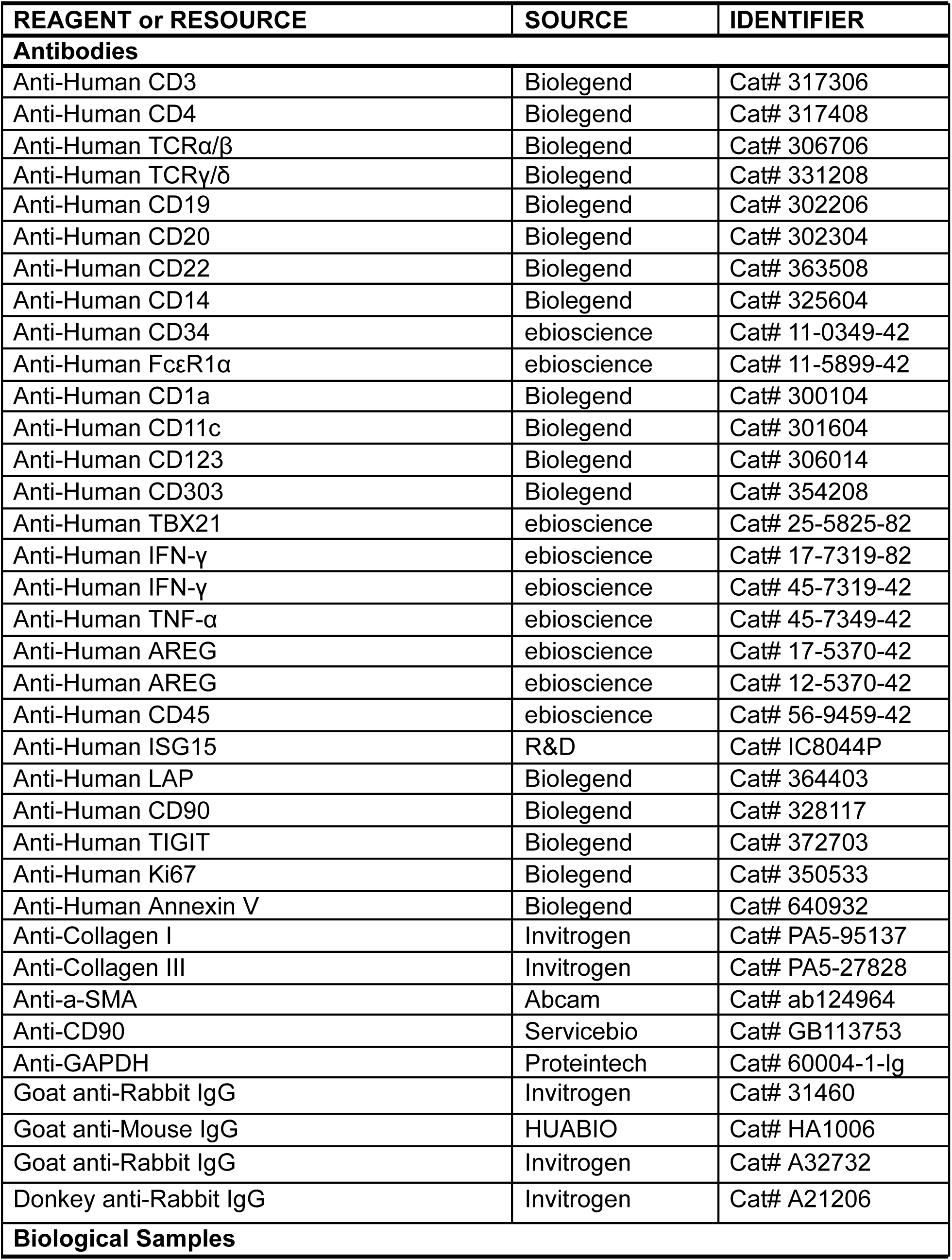
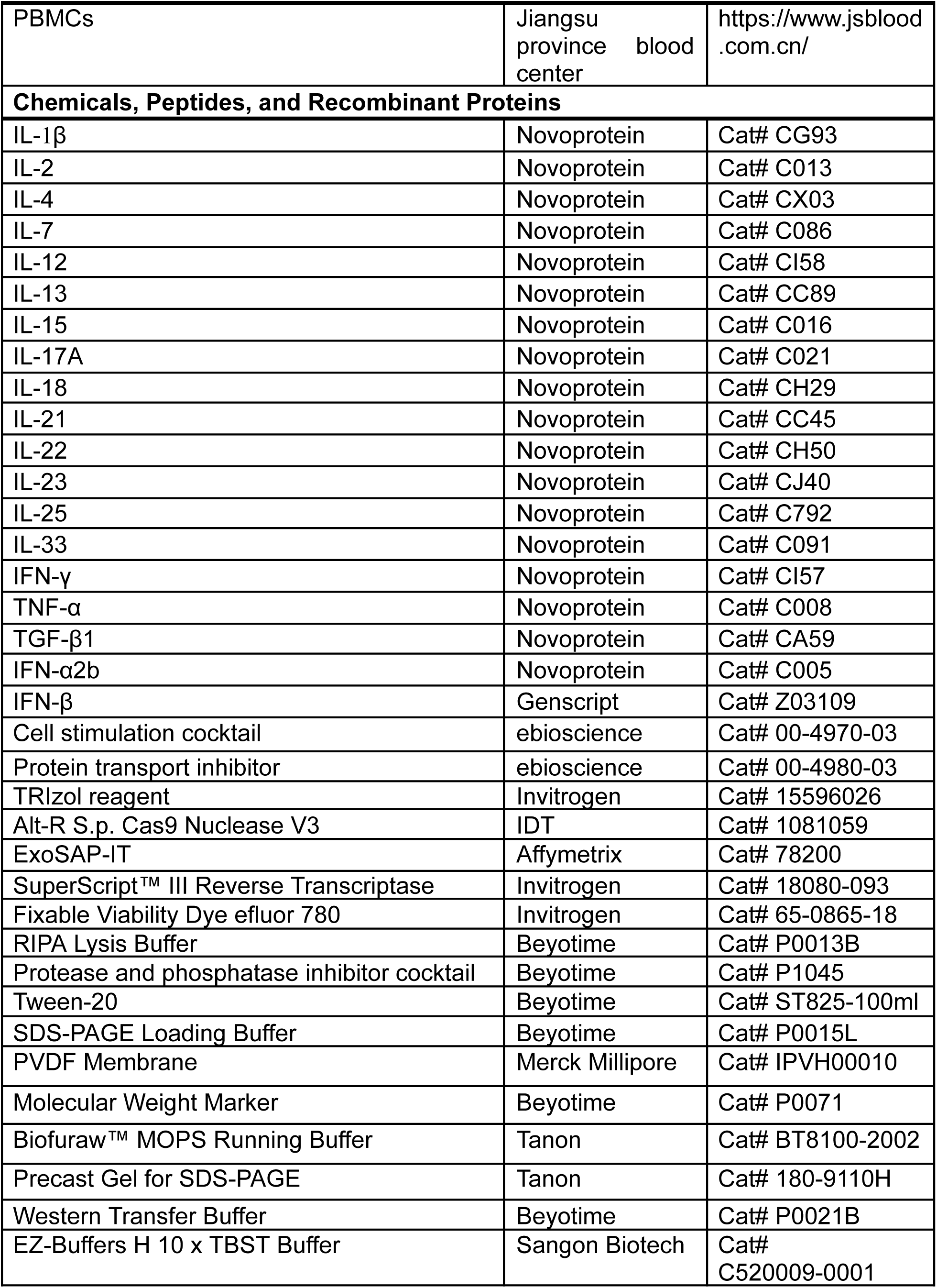
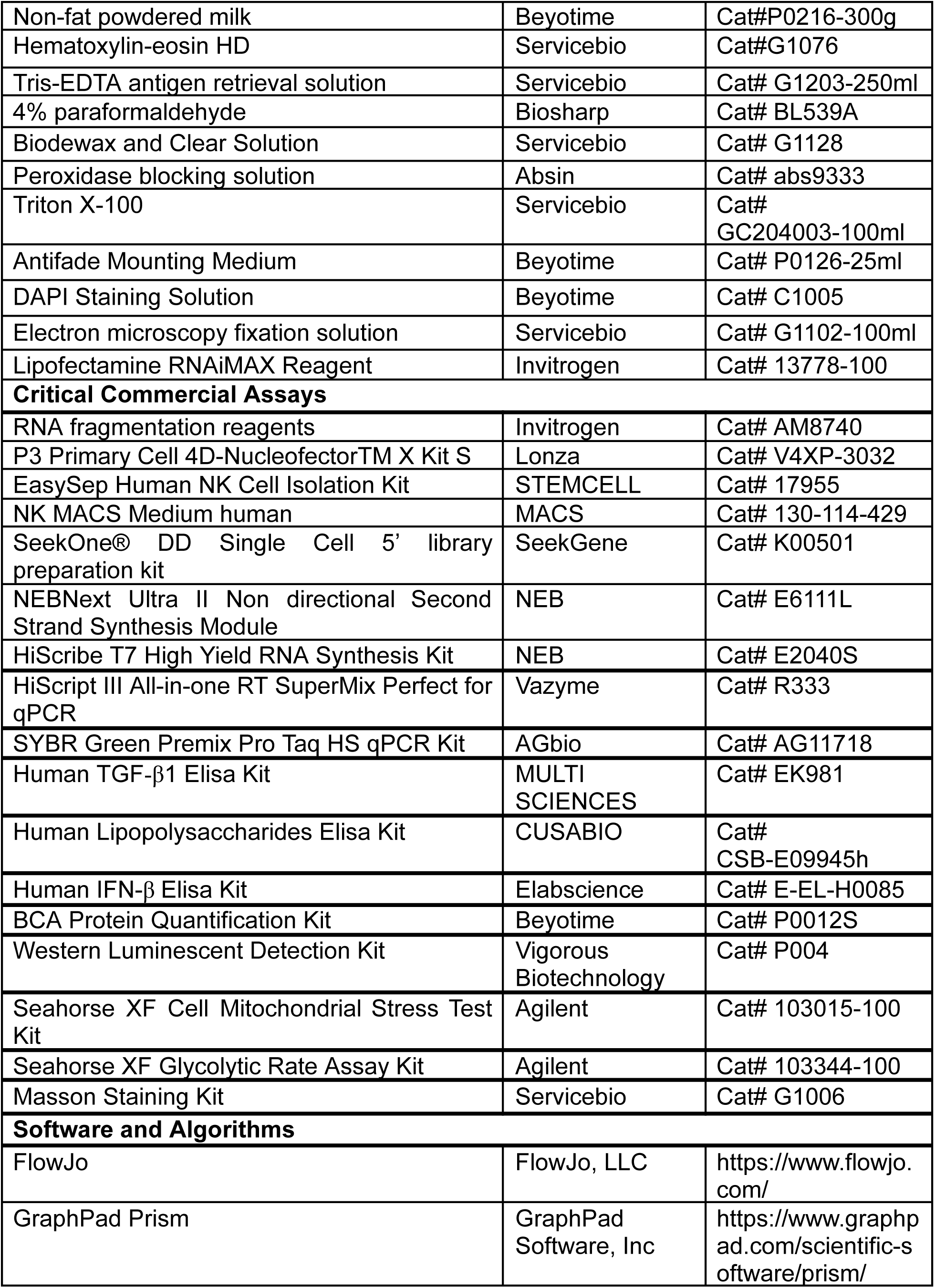
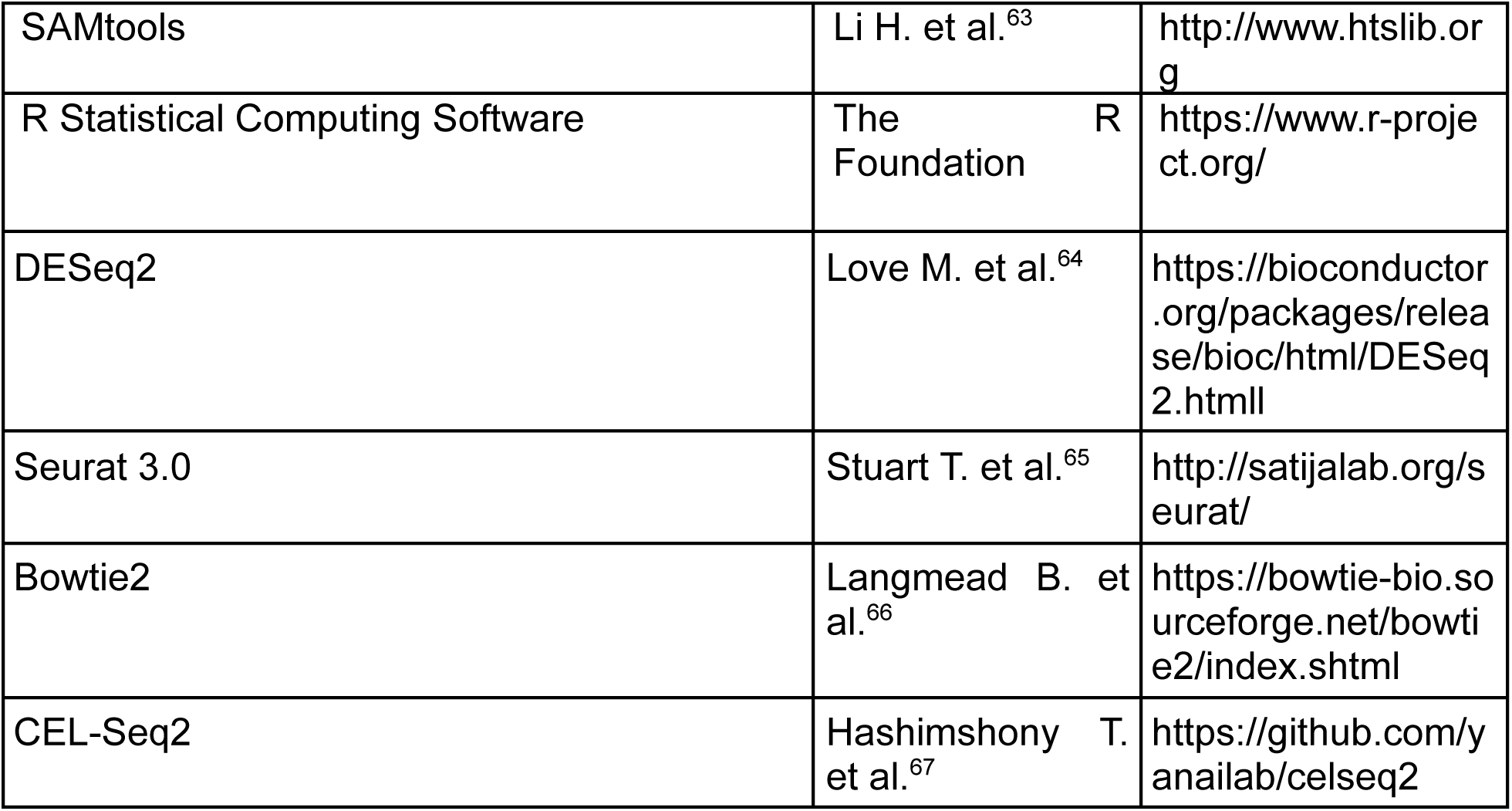

### Resource availability

Further information and requests for resources and reagents should be directed to and will be fulfilled by the lead contact, Yetao Wang.

### Materials availability

This study did not generate new unique reagents.

### Data availability

The datasets produced in this study, including the scRNA-Seq data from Figure 6 and the bulk RNA-Seq data from Figures 3 and 7, are available in the Genome Sequence Archive (GSA) for Humans under accession number: HRA008790, HRA008880 and HRA008788.

### Cell culture and stimulation conditions

All cells were cultured at 37°C containing 5% CO_2_.

**Figures 1A, 1C-1E:** Dermal cells were stimulated with PMA (81 nM) and ionomycin (1.34 lM) (1:500, eBioscience, 00-4970-03) in RPMI 1640 for 2 hrs.

**Figures 2B-2E:** PBMCs or magnetic beads enriched NK cells from PBMCs were stimulated as in Figures 1A-1E.

**Figures 2F and 2G:** Fibroblasts were cultured in DMEM for 16 hrs with Golgi block prior to TGF-β1 detection.

**Figures 2H-2J:** Non-lesional fibroblasts were stimulated with or without indicated cytokines (100 ng/ml each) for 24 hrs. After three washes, co-cultured with PBMCs or magnetic beads enriched NK cells from PBMCs in a transwell. After 48 hrs, the PBMCs or enriched NK cells were collected and stimulated with IL-12 (10 ng/ml), IL-15 (50 ng/ml), and IL-18 (50 ng/ml) for 16 hrs prior to IFN-γ detection.

**Figure 2K:** Non-lesional fibroblasts were stimulated with TGF-β1 (100 ng/ml) for 24 hrs. After three washes, they were cultured in DMEM for 48hrs, the supernatant was collected for ELISA.

**Figure 2L:** Enriched NK cells were stimulated with IL-12 (10 ng/ml) + IL-15 (50 ng/ml) + IL-18 (50 ng/ml) for 16 hrs. After 3 washes, co-cultured with non-lesional and lesional fibroblasts in transwell.

**Figures 3C-3G and 3I:** Enriched NK cells were cultured in the following media as indicated in Figure 3A: basic NK MACS medium, NK MACS medium, basic NK MACS medium supplemented with IL-12 (10 ng/ml), IL-15 (50 ng/ml), and IL-18 (50 ng/ml), and NK MACS medium supplemented with the same cytokines, for 16 hours. After three washes, the NK cells were cultured in DMEM for 48 hours, and the conditioned supernatants were used to culture keloid fibroblasts for 24 (Figure 3C, 3G) or 48 hours (Figure 3D-3F, 3I).

**Figures 5A-5E:** PBMCs or enriched NK cells were stimulated as in Figures 1A-1E.

**Figures 7B-7C:** PBMCs were cultured with or without IFN-β (50 ng/ml) for 16 hours, followed by stimulation with IL-12 (10 ng/ml) + IL-15 (50 ng/ml) + IL-18 (50 ng/ml) for 16 hrs.

**Figures 7K and 7L:** Enriched NK cells were treated with or without IFN-β (10 ng/ml) for 7 days in NK MACS medium.

**Figures 8A-8C:** PBMCs were cultured in RPMI 1640 in the presence or absence of IFN-β (10 ng/ml) for 7 days, then were stimulated with IL-12 (10 ng/ml) + IL-15 (50 ng/ml) + IL-18 (50 ng/ml) for 16 hrs.

**Figures 8D-8I:** Enriched NK cells were cultured in NK MACS medium in the presence or absence of IFN-β (10 ng/ml) for 7 days, then were stimulated with IL-12 (10 ng/ml) + IL-15 (50 ng/ml) + IL-18 (50 ng/ml) for 4 hrs.

**Figures 8J-8K:** Enriched NK cells were cultured in NK MACS medium in the presence of IFN-β (10 ng/ml) for 7 days.

**Figures 8L-8M:** Enriched NK cells were cultured and stimulated as in Figures 8D-8I.

### Clinical samples

Skin keloid lesional and non-lesional samples, along with blood samples from healthy donors and keloid patients, were obtained from the biobank at the Institute of Dermatology, Chinese Academy of Medical Sciences, the Jiangsu Biobank of Clinical Resources, and the Jiangsu Province Blood Center. All participants provided written informed consent for protocols included in the study of immune cell biological functions in skin-related diseases, in accordance with procedures approved by the ethics committee of the Institute of Dermatology, Chinese Academy of Medical Sciences and Peking Union Medical College. There was no blinding of investigators or subjects during the study conduct or analysis, and no power analysis was performed to determine group size. Donor information is provided in Table S1.

### Human skin cell preparation

Isolation of dermal single cells from keloid skin tissues for in vitro culture and flow cytometry: 1. Skin biopsies, after removal of subcutaneous fat, were incubated in 1 mL of dispase (5 U/mL in PBS with 1% penicillin/streptomycin) at 37°C for 2 hours to separate the epidermis from the dermis; 2. The dermis was washed in PBS and then digested in a solution containing collagenase III (3 mg/mL) and DNase (5 µg/mL) in 10% FBS/RPMI 1640 at 37°C for 2 hours, shaking every 30 minutes. 3. The digested dermis was filtered through a 70 µm strainer to collect the flow-through. After centrifugation at 700 x g for 3 minutes, the cells were resuspended in MACS buffer (0.5% BSA and 2 mM EDTA in PBS) and were ready for use.

Culturing primary fibroblasts from human skin: 1. Skin tissues were obtained using sterile techniques and transported in ice-cold PBS after removing adipose tissue; 2. The dermis was washed with PBS and digested overnight at 4°C with Dispase II (5 U/mL in PBS with 1% penicillin/streptomycin). After removing the epidermis, the dermis was washed with PBS (with 1% penicillin/streptomycin) and cut into 1 mm^3^ fragments; 3. Following drainage of excess liquid, the fragments were placed in a 10 cm culture dish to adhere for 20-30 minutes and then cultured in high-glucose DMEM (Gibco, C11995500BT). The fibroblasts were passed on day 14 and subsequently every 3 days thereafter.

### Peripheral blood mononuclear cells (PBMCs) isolation

Blood samples were diluted with an equal volume of serum-free RPMI 1640 (Gibco, 11875093), and leukopaks were washed with 80 ml of serum-free RPMI 1640 before being overlaid on Lymphoprep (STEMCELL, 07851). Both were then centrifuged at 500 x g for 30 minutes at room temperature. After three washes with MACS buffer, the isolated PBMCs were either used immediately or cryopreserved in FBS containing 10% DMSO.

### Keloid xenograft mouse experiment

The keloid xenograft model was established using 7-week-old NCG mice (NOD/ShiLtJGpt-*Prkdc* ^em26Cd52^ Il2rg^em26Cd22^ /Gpt, GemPharmatech, T001475). Human keloid tissues were incubated in dispase II at 4°C for 4 hours to remove the epidermis, after which they were cut into sections measuring approximately 8 × 8 × 10 mm. The anesthetized NCG mice were engrafted with the keloid dermal tissue on their dorsal skin. On days 7 and 14 post-engraftment, the engrafted keloid tissues received injections of wild-type, IFNG knockout, or AREG knockout NK cells, which had been stimulated with IL-12 (10 ng/ml), IL-15 (50 ng/ml), and IL-18 (50 ng/ml) for 16 hours. The engrafted keloid tissues were then dissected on day 21 for downstream analysis. All mice were kept in microisolator cages and given autoclaved food and acidified, autoclaved water within a specific pathogen-free environment. Animal use complied with the guidelines of the Institutional Animal Care and Use Committee at the Hospital for Skin Diseases, Institute of Dermatology, Chinese Academy of Medical Sciences, and Peking Union Medical College. All experimental protocols were reviewed and approved by this committee.

### Flow cytometry

Skin cells or PBMCs were stained with the fixable viability dye eFluor 780 (Invitrogen, 65-0865-18) to assess viability. For surface marker staining, cells were incubated with antibodies in MACS buffer at 4°C for 30 minutes, protected from light. For intracellular staining, cells were fixed and permeabilized using the Foxp3 Staining Kit (eBioscience, 00-5523-00), followed by antibody incubation in the permeabilization buffer at 4°C for 30 minutes, also in the dark. After washing with MACS buffer, the cells were prepared for flow cytometry. Samples with insufficient cell numbers for analysis were excluded from further processing.

### H&E, Masson and immunofluorescence staining

Keloid tissue was harvested from the backs of mice, fixed, embedded in paraffin, and sectioned. The paraffin sections were then deparaffinized through sequential immersion in Dewaxing Transparent Liquid I for 15 minutes, followed by Dewaxing Transparent Liquid II for 15 minutes. The sections were subsequently dehydrated in anhydrous ethanol I for 3 minutes, anhydrous ethanol II for 3 minutes, 95% ethanol for 3 minutes, and 70% ethanol for 3 minutes. The sections were then rinsed with distilled water.

For H&E staining, the sections were first treated with HD constant staining pretreatment solution for 1 minute, then immersed in hematoxylin solution for 3-5 minutes and rinsed with distilled water. The sections were differentiated in a hematoxylin differentiation solution, rinsed with distilled water, and blued in hematoxylin bluing solution, followed by rinsing with tap water. The sections were then dehydrated in 95% ethanol for 1 minute, stained with eosin for 15 seconds, and dehydrated through a series of solutions: absolute ethanol I, II, and III for 2 minutes each, followed by normal butanol I and II for 2 minutes each, and xylene I and II for 2 minutes each. The sections were mounted with neutral gum and examined under a microscope.

For Masson staining, the sections were mordanted in 10% potassium dichromate and 10% trichloroacetic acid for 30 minutes. Nuclei were stained with hematoxylin for 20 minutes, differentiated in hydrochloric acid and ethanol for 15 seconds, and blued in weak ammonia for 15 seconds. The sections were stained with Masson’s solution for 1 minute, rinsed in 1% acetic acid, dehydrated in ethanol, and cleared in xylene I and II for 10 minutes each. The sections were mounted with resin and examined under a microscope.

For immunofluorescence staining, slides were baked at 60°C for 2 hours, deparaffinized, and subjected to antigen retrieval in Tris-EDTA solution at 98°C for 25 minutes. After cooling, the slides were permeabilized with 0.1% Triton X-100 for 10 minutes. Endogenous peroxidase activity was blocked for 15 minutes, followed by incubation with 5% BSA for 1 hour. Primary antibodies—collagen I (1:200, Invitrogen, PA5-95137), collagen III (1:200, Invitrogen, PA5-27828), α-SMA (1:200, Abcam, ab124964), and CD90 (1:200, Servicebio, GB113753)—were applied overnight at 4°C. After three washes with PBST, slides were incubated with species-specific fluorescent secondary antibodies (1:500) for 40 minutes in the dark. Additional washes with PBST were followed by DAPI staining for 10 minutes in the dark. The slides were mounted with anti-fade solution and sealed with nail polish prior to microscopic observation.

### Western blot

Homogenized keloid tissue from mouse dorsal skin and cultured fibroblasts were lysed in RIPA buffer containing 2% protease and phosphatase inhibitors mix (Beyotime, P1045). Protein concentrations were determined, and samples were mixed with loading buffer and denatured at 98°C for 10 minutes. Equal protein amounts were separated by SDS-PAGE and transferred to a PVDF membrane (Merck Millipore, IPVH00010) by wet electroblotting. The membrane was blocked with 5% non-fat milk for 3 hours and incubated overnight at 4°C with primary antibodies against COL1 (1:1000, Invitrogen, PA5-95137), COL3 (1:1000, Invitrogen, PA5-27828), α-SMA (1:1000, Abcam, ab124964), and GAPDH (1:5000, Proteintech, 60004-1-Ig). After washing with TBST, the membrane was incubated with HRP-conjugated secondary antibodies (goat anti-rabbit IgG, 1:2000, Invitrogen, 31460; goat anti-mouse IgG, 1:2000, HUABIO, HA1006) for 3 hours at room temperature. After TBST washing, the membrane was developed using chemiluminescent solution (Vigorous Biotechnology, P004). Imaging was performed with a ChemiDoc Imaging System (Bio-rad), and band intensities were quantified using ImageJ. Target protein levels were calculated as the ratio of target protein to GAPDH.

### Quantitative reverse transcription PCR

RNA was extracted from fibroblasts and NK cells using TRIzol reagent (ThermoFisher, 15596026), and reverse transcribed into cDNA with HiScript III All-in-one RT SuperMix (Vazyme, R333). Quantitative PCR was performed using the following specific primers: EGFR forward: 5’-AGG CAC GAG TAA CAA GCT CAC-3’; EGFR reverse: 5’-ATG AGG ACA TAA CCA GCC ACC-3’; β-actin forward: 5’-AGG CAC CAG GGC GTG ATG GTG-3’; β-actin reverse: 5’-GGT CTC AAA CAT GAT CTG GGT-3’; IFNAR1: forward: 5’-TGT GAG AAA ACA AAA CCA GGA AA-3’; IFNAR1 reverse: 5’-TCT TCA ATG GCT GTT CAG AGA A-3’; IRF1 forward: 5’-ACC CTG GCT AGA GAT GCA GA-3’; IRF1 reverse: 5’-TGC TTT GTA TCG GCC TGT GT-3’; IRF9: forward: 5’-GAA GAG GTG GTT CAG ACT TGG T-3’; IRF9: reverse: 5’-TCT GCT CCA GCA AGT ATC GG-3’.

### Seahorse extracellular flux analysis

Mitochondrial respiration and lactate secretion of NK cells were assessed by measuring oxygen consumption rate (OCR) and glycolytic proton efflux rate (glycoPER), respectively, using an XFe96 extracellular flux analyzer (Agilent). XFe96 microplates (Agilent, 103792-100) were pre-coated with 100 μg/ml polylysine (Sigma, P7280), and 2-3 x 10^5^ NK cells per well were seeded in triplicates in Seahorse XF RPMI medium (Agilent, 103576-100) supplemented with 2 mM L-glutamine (Agilent, 103579-100), 1 mM sodium pyruvate (Agilent, 103578-100), and 10 mM D-glucose (Agilent, 103577-100). After a 1 hour incubation in a CO_2_-free incubator at 37°C, glycolytic and mitochondrial stress tests were performed as per the manufacturer’s protocol.

For glycolysis, basal extracellular acidification rate (ECAR) was measured, followed by the addition of 0.5 μM rotenone (Agilent, 103344-100) and 0.5 μM antimycin A (Agilent, 103344-100) to inhibit mitochondrial complexes I and III, respectively. At the end of the assay, 50 mM 2-DG (Agilent, 103344-100) was added to block glycolysis. Basal glycolysis was defined as glycoPER before rotenone and antimycin A, and compensatory glycolysis as glycoPER after their addition.

For mitochondrial respiration, basal OCR was measured, followed by the addition of 1.5 μM oligomycin (Agilent, 103015-100), 0.5 μM FCCP (Agilent, 103015-100), and 0.5 μM rotenone (Agilent, 103015-100) plus 0.5 μM antimycin A (Agilent, 103015-100). Basal respiration was calculated by subtracting OCR after rotenone and antimycin A from OCR before oligomycin treatment. Maximal respiration was calculated as the difference between OCR after FCCP and after rotenone and antimycin A. Spare respiratory capacity was determined by subtracting basal respiration from maximal respiration. Mitochondrial ATP production was calculated as the difference between OCR before and after oligomycin.

### Transmission electron microscope

Unstimulated NK cells and those stimulated with IFN-β (10 ng/ml) for 7 days were fixed in electron microscopy fixative (Servicebio, G1102-100ml), then dehydrated through a series of graded alcohol solutions. The cells were infiltrated with acetone to facilitate resin infiltration. The samples were embedded in epoxy resin, cured at 60°C, and sectioned into 50–100 nm thick slices using an ultramicrotome. The sections were placed on copper grids, samples were stained with a 2% uranyl acetate solution saturated in alcohol in the dark for 8 minutes, followed by staining with a 2.6% lead citrate solution in a carbon dioxide environment for 8 minutes. Mitochondrial morphology was then examined using a transmission electron microscope (JEOL, JEM-1400) at 100 kV.

### scRNA-Seq library preparation

Single-cell RNA-seq libraries were prepared using the SeekOne DD Single Cell 5’ library preparation kit (SeekGene, K00501). PBMCs (5 x 10^6^) from each of ten healthy donors and ten keloid patients were pooled to form healthy donor and keloid patient groups, respectively. NK cells were enriched from each group using the EasySep Human NK Cell Isolation Kit, adjusted to a concentration of 10^6^ cells/ml with > 90% viability. A total of 20,000 cells for each group were mixed with reverse transcription reagents and loaded into the sample well of the SeekOne® DD Chip S3, with gel beads and partitioning oil dispensed into corresponding wells. After emulsion droplet generation, reverse transcription was performed at 42°C for 90 minutes and then inactivated at 85°C for 5 minutes. The resulting cDNA was purified, amplified by PCR after droplet breakage, and subsequently fragmented, end-repaired, A-tailed, and ligated to sequencing adapters. Indexed PCR was performed, and the sequencing libraries were cleaned with VAHTS DNA Clean Beads (Vazyme, N411-01), then analyzed using Qubit (Thermo Fisher Scientific, Q33226) and Bio-Fragment Analyzer (Bioptic Qsep400). Libraries were sequenced on the Illumina NovaSeq 6000 with a PE150 read length.

### Bulk RNA-Seq library preparation

The bulk RNA-Seq libraries of fibroblasts were prepared using CEL-Seq2^67^. Total RNA was extracted using TRIzol reagent. 100 ng RNA for each library was used for first strand cDNA synthesis using barcoded primers as follows (barcode underlined): IFN-γ^+^AREG^-^ stimulated donor 1 fibroblasts: 5’-GCC GGT AAT ACG ACT CAC TAT AGG GAG TTC TAC AGT CCG ACG ATC NNN NNN <u>AGC TAG</u> TTT TTT TTT TTT TTT TTT TTT TTT V-3’; IFN-γ^+^AREG^-^ stimulated donor 2 fibroblasts: 5’-GCC GGT AAT ACG ACT CAC TAT AGG GAG TTC TAC AGT CCG ACG ATC NNN NNN <u>CAT GCA</u> TTT TTT TTT TTT TTT TTT TTT TTT V-3’; IFN-γ^+^AREG^-^ stimulated donor 3 fibroblasts: 5’-GCC GGT AAT ACG ACT CAC TAT AGG GAG TTC TAC AGT CCG ACG ATC NNN NNN <u>TCA CAG</u> TTT TTT TTT TTT TTT TTT TTT TTT V-3’; IFN-γ^+^AREG^+^ stimulated donor 1 fibroblasts: 5’-GCC GGT AAT ACG ACT CAC TAT AGG GAG TTC TAC AGT CCG ACG ATC NNN NNN <u>AGC TTC</u> TTT TTT TTT TTT TTT TTT TTT TTT V-3’; IFN-γ^+^AREG^+^ stimulated donor 2 fibroblasts: 5’-GCC GGT AAT ACG ACT CAC TAT AGG GAG TTC TAC AGT CCG ACG ATC NNN NNN <u>CAC TAG</u> TTT TTT TTT TTT TTT TTT TTT TTT V-3’; IFN-γ^+^AREG^+^ stimulated 3 donor 3 fibroblasts: 5’-GCC GGT AAT ACG ACT CAC TAT AGG GAG TTC TAC AGT CCG ACG ATC NNN NNN <u>AGT</u> <u>GCA</u> TTT TTT TTT TTT TTT TTT TTT TTT V-3’ (“V” represents any base except T). The second strand was synthesized using the NEBNext second strand synthesis module (NEB, E6111L). The dsDNA was purified using RNAClean XP (Beckman Coulter, A63987) and subsequently subjected to in vitro transcription (IVT) with the HiScribe T7 high yield RNA synthesis kit (NEB, E2040S). Following ExoSAP-IT treatment (Affymetrix, 78200), the IVT RNA was fragmented using RNA fragmentation reagents (Invitrogen, AM8740). The fragmented RNA was then reverse transcribed in the second round using a random hexamer (5’-GCC TTG GCA CCC GAG AAT TCC ANN NNN N-3’). The final library was amplified with indexed primers.: RP1: 5’-AAT GAT ACG GCG ACC ACC GAG ATC TAC ACG TTC AGA GTT CTA CAG TCC GA-3’ and RPI1: 5’-CAA GCA GAA GAC GGC ATA CGA GAT CGT GAT GTG ACT GGA GTT CCT TGG CAC CCG AGA ATT CCA-3’. After quality check, the purified libraries were sequenced by Illumina NovaSeq Xplus.

The bulk RNA-Seq libraries for NK cells were constructed using the VAHTS Universal V8 RNA-seq Library Prep Kit for Illumina (Vazyme, NR605). In brief, total RNA was extracted from NK cells using TRIzol reagent. mRNA was captured with VAHTS mRNA capture beads, and the purified mRNA was fragmented in Frag/Prime buffer at 85°C for 5 minutes, followed by a 2-minute hold at 4°C. Double-stranded cDNA was synthesized using the First and Second Strand Enzyme Mixes, respectively. RNA adaptors were then ligated using a ligation enzyme mix, and size selection was performed in two rounds with VAHTS DNA clean beads (0.6x and 0.1x, respectively). Finally, the libraries were amplified with VAHTS i5 and i7 primers, and paired-end 150-bp reads were generated on the DNBSEQ-T7 platform.

### Cas9 RNP-mediated knockout in NK cells

NK cells were isolated from PBMCs using the EasySep Human NK Cell Isolation Kit (STEMCELL, 17955) and expanded in NK MACS Medium (MACS, 130-114-429) for 10 days to prepare them for electroporation. To form ribonucleoprotein (RNP) complexes, 200 pmol of sgRNA (synthesized by GenScript) was mixed with 100 pmol of Cas9 protein and incubated at room temperature for 20 minutes. One million NK cells were then resuspended in 20 µl of 4D Nucleofector Master Mix (82% P3 solution + 18% Supplement 1; Lonza, V4XP-3032) and mixed with the Cas9 RNP complex for electroporation using the CM137 program. After electroporation, NK cells were cultured in NK MACS Medium for 48 hours prior to downstream experiments. The target genes and sequences for knockout were as follows: CD19: 5’-AGG AAG AAG AGG AGG CGA GG-3’; IFNG: 5’-AAA GAG TGT GGA GAC CAT CA-3’; AREG: 5’-GAG GAC GGT TCA CTA CTA GA-3’; IFNAR1: 5’-AAA CAC TTC TTC ATG GTA TG-3’; IRF1: 5’-CTC TGG TTC TTG GTG AGA GG-3’; IRF9: 5’-CTG GAA ACA TGC AGG CAA GC-3’.

### siRNA-mediated knockdown in fibroblasts

siRNA targeting EGFR was designed using the siDirect (http://sidirect2.rnai.jp/). The antisense strand sequences for the siRNAs were as follows: siRNA1: 5’-UAA AUU CAC UGC UUU GUG GTT-3’; siRNA2: 5’-AUU AUC ACA UCU CCA UCA CTT-3’. Fibroblasts were cultured in 24-well plates. For transfection, 0.25 µl of siRNA or Lipofectamine RNAiMAX (Invitrogen, 13778-100) were each diluted in 25 µl of Opti-MEM (Gibco, 31985070) and incubated at room temperature for 5 minutes. The two components were then gently mixed and incubated at room temperature for an additional 15 minutes. Prior to adding the transfection reagents, the fibroblast culture medium was replaced with 200 µl of serum-free DMEM, and the cells were incubated for 2 hours at 37°C. Following incubation, 50 µl of the transfection reagent mixture was added to the fibroblast cultures. After 6 hours, the medium was replaced with complete DMEM, and the cells were cultured for an additional 24 hours before proceeding with further experiments.

### scRNA-Seq data processing

An average of 34,308 reads, and a median of 1,295 genes per cell were identified using SeesoulTools (v1.0.0). After alignment, cells meeting the following criteria were retained for downstream analysis using Seurat (Version 4.0): (1) feature count between 300 and 6,000; (2) fewer than 20,000 UMIs; (3) less than 20% mitochondrial gene UMIs. Potential doublets and multiplets were filtered out using DoubletFinder, resulting in the integration of 18,595 cells. NK cell identification was based on the expression of NK cell markers (NKG7, GNLY, GZMB, CST7, KLRD1, PRF1, FCGR3A), with exclusion of markers associated with other cell types, including T cells (CD3D, CD3G), B cells (CD19, MS4A1, CD22), ILC3 (RORC, IL17A), ILC2 (IL5, IL13, PTGDR2, HPGD, HPGDS), hematopoietic stem cells (CD34), mast cells (CPA3, TPSAB1, FCER1A), neutrophils (CSF3R), and myeloid cells (CD14, CD163, CD207, LAMP3, IL3RA). A total of 16,558 NK cells were re-clustered into five distinct clusters (resolution = 0.5). Differentially expressed genes within each cluster were identified using the FindAllMarkers function in Seurat. The AddModuleScore function in Seurat was used to calculate gene set scores for NK cell clusters, with the signature genes listed in Table S4^41^.

### Bulk RNA-Seq analysis

The CEL-Seq2 pipeline (https://github.com/yanailab/celseq2)^67^ was used to generate the transcriptome count matrix for all samples, according to the default settings. In brief, Read 2 was assigned to each sample based on the barcode from Read 1, and the data were mapped to the hg19 genome using Bowtie2. UMI counts were then obtained for each sample, and the count matrix was subsequently analyzed with the DESeq2 package in R. Transcript counts were normalized using the variance stabilizing transformation (VST) method in DESeq2. Differential expression analysis was performed with DESeq2, considering genes with |log_2_FC|> 0.5, *P*<0.05 as differentially expressed.

### Statistical analysis

Statistical test was performed using GraphPad Prism 9. Wilcoxon matched-pairs signed rank test, Mann-Whitney test or two tailed paired/unpaired t-test used in this study were specified in the figure legends. Variance was estimated by calculating the mean ± s.e.m. in each group. *P* < 0.05 was considered significant. For gene set score analysis, the *P* value was determined using either a Mann-Whitney test or a two-tailed paired t-test, as indicated in the figure legend. *P* < 0.05 was considered significant.

## Notes

### Competing Interest Statement

The authors have declared no competing interest.

