## Supplementary material for "Natural killer cell dysfunction drives keloid pathogenesis": suppemental tables

| Table S1 \| Donor characteristics and associated experiments | | | | | | |
|---|---|---|---|---|---|---|
| **Sample ID** | **Group** | **Sex** | **Age** | **Distribution of keloid locations among patients** | **Tissues utilized in the experiments** | **Related figures** |
| keloid 1 | keloid | Female | 52 | chest | flow cytometry, skin | figure 1 |
| keloid 2 | keloid | Male | 25 | chest | flow cytometry, skin | figure 1 |
| keloid 3 | keloid | Male | 48 | abdomen | flow cytometry, skin | figure 1 |
| keloid 4 | keloid | Female | 24 | back | flow cytometry, skin | figure 1 |
| keloid 5 | keloid | Male | 25 | chest | flow cytometry, skin | figure 1 |
| keloid 6 | keloid | Male | 42 | chest | flow cytometry, skin, PBMC | figure 1, 5 |
| keloid 7 | keloid | Female | 21 | ear | flow cytometry, skin | figure 1 |
| keloid 8 | keloid | Female | 43 | leg | flow cytometry, skin | figure 1 |
| keloid 9 | keloid | Female | 19 | ear | flow cytometry, skin | figure 1 |
| keloid 10 | keloid | Male | 33 | chest | flow cytometry, skin | figure 1 |
| keloid 11 | keloid | Male | 27 | face | flow cytometry, skin | figure 1 |
| keloid 12 | keloid | Male | 31 | face | flow cytometry, skin | figure 1 |
| keloid 13 | keloid | Female | 53 | abdomen | flow cytometry, PBMC | figure 5 |
| keloid 14 | keloid | Female | 56 | abdomen | flow cytometry, PBMC | figure 5 |
| keloid 15 | keloid | Male | 57 | back | flow cytometry, PBMC | figure 5 |
| keloid 16 | keloid | Male | 28 | face | flow cytometry, PBMC | figure 5 |
| keloid 17 | keloid | Female | 43 | arm | flow cytometry, PBMC | figure 5 |
| keloid 18 | keloid | Male | 25 | face | flow cytometry, PBMC | figure 5 |
| keloid 19 | keloid | Female | 40 | chest | flow cytometry, PBMC | figure 5 |
| keloid 20 | keloid | Male | 28 | face | flow cytometry, PBMC | figure 5 |
| keloid 21 | keloid | Female | 34 | leg | flow cytometry, PBMC | figure 5 |
| keloid 22 | keloid | Male | 26 | face | flow cytometry, PBMC | figure 5 |
| keloid 23 | keloid | Male | 22 | face | flow cytometry, PBMC | figure 5 |
| keloid 24 | keloid | Male | 29 | chest | flow cytometry, PBMC | figure 5 |
| keloid 25 | keloid | Female | 28 | arm | flow cytometry, PBMC; scRNA-seq | figure 5, 6 |
| keloid 26 | keloid | Female | 28 | shoulder | flow cytometry, PBMC; scRNA-seq | figure 5, 6 |
| keloid 27 | keloid | Female | 37 | ear | flow cytometry, PBMC; scRNA-seq | figure 5, 6 |
| keloid 28 | keloid | Male | 24 | face | flow cytometry, PBMC; scRNA-seq | figure 5, 6 |
| keloid 29 | keloid | Female | 22 | chest | flow cytometry, PBMC; scRNA-seq | figure 5, 6 |
| keloid 30 | keloid | Female | 18 | arm | flow cytometry, PBMC; scRNA-seq | figure 5, 6 |
| keloid 31 | keloid | Female | 27 | chest | flow cytometry, PBMC | figure 5 |
| keloid 32 | keloid | Male | 21 | face | xgraft, skin | figure 4 |
| keloid 33 | keloid | Male | 25 | chest | xgraft, skin | figure 4 |
| keloid 34 | keloid | Female | 27 | abdomen | scRNA-seq | figure 6 |
| keloid 35 | keloid | Female | 25 | arm | scRNA-seq | figure 6 |
| keloid 36 | keloid | Male | 31 | face | scRNA-seq | figure 6 |
| keloid 37 | keloid | Male | 22 | chest | scRNA-seq | figure 6 |
| healthy 1 | healthy | Male | 29 | / | flow cytometry, PBMC | figure 5 |
| healthy 2 | healthy | Female | 26 | / | flow cytometry, PBMC | figure 5 |
| healthy 3 | healthy | Male | 35 | / | flow cytometry, PBMC | figure 5 |
| healthy 4 | healthy | Male | 29 | / | flow cytometry, PBMC | figure 5 |
| healthy 5 | healthy | Female | 24 | / | flow cytometry, PBMC | figure 5 |
| healthy 6 | healthy | Female | 33 | / | flow cytometry, PBMC | figure 5 |
| healthy 7 | healthy | Male | 30 | / | flow cytometry, PBMC | figure 5 |
| healthy 8 | healthy | Female | 24 | / | flow cytometry, PBMC | figure 5 |
| healthy 9 | healthy | Male | 23 | / | flow cytometry, PBMC | figure 5 |
| healthy 10 | healthy | Male | 23 | / | flow cytometry, PBMC | figure 5 |
| healthy 11 | healthy | Female | 34 | / | flow cytometry, PBMC | figure 5 |
| healthy 12 | healthy | Female | 44 | / | flow cytometry, PBMC; scRNA-seq | figure 5, 6 |
| healthy 13 | healthy | Male | 25 | / | flow cytometry, PBMC; scRNA-seq | figure 5, 6 |
| healthy 14 | healthy | Male | 27 | / | flow cytometry, PBMC; scRNA-seq | figure 5, 6 |

| | | | | | | |
|---|---|---|---|---|---|---|
| healthy 15 | healthy | Female | 29 | / | flow cytometry, PBMC; scRNA-seq | figure 5, 6 |
| healthy 16 | healthy | Male | 23 | / | flow cytometry, PBMC | figure 5 |
| healthy 17 | healthy | Male | 24 | / | flow cytometry, PBMC | figure 5 |
| healthy 18 | healthy | Female | 24 | / | flow cytometry, PBMC; scRNA-seq | figure 5, 6 |
| healthy 19 | healthy | Female | 21 | / | flow cytometry, PBMC | figure 5 |
| healthy 20 | healthy | Female | 48 | / | flow cytometry, PBMC; scRNA-seq | figure 5, 6 |
| healthy 21 | healthy | Male | 44 | / | scRNA-seq | figure 6 |
| healthy 22 | healthy | Female | 43 | / | scRNA-seq | figure 6 |
| healthy 23 | healthy | Male | 40 | / | scRNA-seq | figure 6 |
| healthy 24 | healthy | Male | 31 | / | scRNA-seq | figure 6 |

| Table S2 \| DEgenes of fibroblasts cultured in IFN-γ+AREG- vs IFN-γ+AREG+ | | |
|---|---|---|
| gene name | log2FC | p value |
| MKLN1-AS2 | 3.733521915 | 0.035989516 |
| TMEM253 | 3.617275215 | 0.004620385 |
| FFAR2 | 2.744915193 | 0.003365164 |
| E2F3-IT1 | 2.546811153 | 0.02189863 |
| APOC1 | 2.536083747 | 0.032591678 |
| EPHB3 | 2.344105013 | 0.020043534 |
| STRA6 | 2.243809433 | 0.037446802 |
| FOSB | 2.146508416 | 0.002387674 |
| MAMSTR | 1.835674334 | 0.035493129 |
| CCNE1 | 1.747794465 | 0.015853621 |
| UPK2 | 1.730184806 | 0.022069672 |
| LFNG | 1.729432855 | 0.024386526 |
| PFKFB4 | 1.692063629 | 4.42612E-06 |
| CD37 | 1.587963029 | 0.013337636 |
| C6orf47-AS1 | 1.44985069 | 0.04746274 |
| ZNF837 | 1.431360625 | 0.024064647 |
| FABP5P3 | 1.391009425 | 0.001713701 |
| SETP14 | 1.360056746 | 0.042851512 |
| TET1 | 1.354809374 | 0.004867014 |
| COL23A1 | 1.322692466 | 0.018516545 |
| CCL28 | 1.3012906 | 0.037796553 |
| ZKSCAN3 | 1.249853785 | 0.019220801 |
| NOP14-AS1 | 1.238782778 | 0.000441956 |
| DHX9P1 | 1.205910393 | 0.031358568 |
| PSENEN | 1.184705874 | 0.046618365 |
| FMO4 | 1.141561242 | 0.046502183 |
| ANKRD37 | 1.127673092 | 6.12838E-05 |
| ZNF395 | 1.118367625 | 6.38897E-05 |
| CTGLF11P | 1.091968425 | 0.026192382 |
| BHLHE41 | 1.039641721 | 0.012673103 |
| PPFIA4 | 1.022190612 | 0.011895975 |
| PSKH1 | 0.97939992 | 0.014900778 |
| ZNF385A | 0.959588476 | 0.009235619 |
| MAPRE3 | 0.956090694 | 0.000809319 |
| PAQR8 | 0.956047182 | 0.000631175 |
| NAGS | 0.943458796 | 0.010898921 |
| PPP1R3C | 0.924643455 | 0.035816546 |
| GPR3 | 0.919433943 | 0.011268349 |
| CPA4 | 0.892296679 | 0.019574019 |
| DDIT4 | 0.888665324 | 0.006709989 |
| CUBN | 0.882830126 | 0.035047136 |
| NFKBID | 0.870958881 | 0.034113366 |
| SAMD10 | 0.868615038 | 0.044899267 |
| ENO2 | 0.863986145 | 0.000701075 |
| ZNF174 | 0.856433986 | 0.045942595 |
| DCAKD | 0.835310924 | 0.017673717 |
| TNFRSF14 | 0.822550396 | 0.016168203 |
| ADAMTS6 | 0.820477895 | 0.01025729 |
| ANGPTL4 | 0.818643943 | 0.013597806 |
| PLTP | 0.811446786 | 0.016193898 |
| NRSN2 | 0.785247605 | 0.001470641 |

| | | |
|---|---|---|
| HOOK2 | 0.7784706 | 0.031577459 |
| FOXP4 | 0.77661964 | 0.013012358 |
| TOR4A | 0.769770279 | 0.014940751 |
| PALM2 | 0.765652405 | 0.016944975 |
| FOXL2 | 0.764538337 | 0.009147606 |
| YPEL3 | 0.764115511 | 0.00833637 |
| PRR24 | 0.761561648 | 0.018943095 |
| SPRYD4 | 0.751492915 | 0.005524783 |
| MIR210HG | 0.750544513 | 0.046031747 |
| NF1 | 0.748081819 | 0.009436313 |
| STC1 | 0.746955934 | 0.000763869 |
| RIN1 | 0.738908402 | 0.023478802 |
| MARCH2 | 0.738476307 | 0.024049238 |
| TCF7L2 | 0.727130845 | 0.007911024 |
| GNL1 | 0.723623993 | 0.002784692 |
| ZP3 | 0.718314017 | 0.034698368 |
| APLP1 | 0.717663009 | 0.011582534 |
| TNFAIP8L3 | 0.716891338 | 0.017303036 |
| STRN4 | 0.713989207 | 0.010747027 |
| INO80B | 0.711504032 | 0.022848398 |
| PAQR7 | 0.709152823 | 0.027720989 |
| SYNPO | 0.709048967 | 0.021705473 |
| HMOX1 | 0.707310724 | 0.000776744 |
| SLC39A7 | 0.706496957 | 0.002125263 |
| RAB4B | 0.705204142 | 0.008438832 |
| SLC39A4 | 0.699824456 | 0.014973394 |
| MT1L | 0.698440377 | 0.024696946 |
| KRT81 | 0.69809217 | 0.027272482 |
| HOXA7 | 0.697181925 | 0.035510084 |
| GDF5OS | 0.69664648 | 0.037244873 |
| RPS6KA5 | 0.696575281 | 0.033716726 |
| C5orf38 | 0.692300936 | 0.032158154 |
| SLC39A11 | 0.691125808 | 0.029226487 |
| C1QTNF6 | 0.68025755 | 0.009095787 |
| GPR153 | 0.679125541 | 0.027805911 |
| SNX27 | 0.66085555 | 0.008643285 |
| C19orf26 | 0.65911057 | 0.023198778 |
| GIGYF1 | 0.659075263 | 0.033691727 |
| SLC29A1 | 0.656470993 | 0.003682318 |
| PEX5 | 0.656401862 | 0.030665862 |
| SOCS2 | 0.654873539 | 0.029866338 |
| NAT14 | 0.654543525 | 0.033023575 |
| AGO2 | 0.652884883 | 0.00925411 |
| C1orf233 | 0.645993954 | 0.017313319 |
| JOSD2 | 0.645272866 | 0.041635152 |
| HSPA1B | 0.643534654 | 0.044909535 |
| ERAP1 | 0.628738689 | 0.031689586 |
| FBXO44 | 0.627712529 | 0.046662081 |
| CXXC5 | 0.62742193 | 0.004337622 |
| LYPLA2 | 0.625254301 | 0.017789239 |
| LINC00963 | 0.620371783 | 0.043923825 |
| MLLT3 | 0.619852915 | 0.045862691 |
| C17orf58 | 0.614853171 | 0.020767879 |
| POLM | 0.614712501 | 0.008792689 |

| | | |
|---|---|---|
| CDC42EP5 | 0.61123703 | 0.026155423 |
| RAB11B | 0.609479164 | 0.041479537 |
| FAM98C | 0.609257642 | 0.013114583 |
| BAX | 0.608504557 | 0.017413239 |
| ITM2C | 0.601112441 | 0.041641103 |
| ADAM17 | 0.598262633 | 0.048289472 |
| THRB-IT1 | 0.59634245 | 0.038283629 |
| GSTM2 | 0.594789952 | 0.0375732 |
| C12orf10 | 0.594731028 | 0.023751902 |
| TAF10 | 0.594299259 | 0.021812871 |
| JUND | 0.593667038 | 0.004373611 |
| TMEM241 | 0.588803352 | 0.049714554 |
| MED24 | 0.588578376 | 0.013582606 |
| TSPYL2 | 0.588306796 | 0.020151567 |
| STMN3 | 0.587369043 | 0.005997457 |
| OR7E38P | 0.586350007 | 0.032708289 |
| SFR1 | 0.582654902 | 0.049944801 |
| CMIP | 0.581773774 | 0.013142701 |
| OTUB1 | 0.581298294 | 0.017661092 |
| IRF3 | 0.580311714 | 0.019913109 |
| HEPN1 | 0.573546588 | 0.049694964 |
| PDK1 | 0.573070597 | 0.046189408 |
| PLAU | 0.572598465 | 0.045246592 |
| DCXR | 0.566286341 | 0.043805516 |
| PYCARD | 0.564463003 | 0.009820533 |
| PTPRS | 0.561991182 | 0.031426157 |
| AAAS | 0.561852182 | 0.043017738 |
| CDCP1 | 0.561771737 | 0.028056872 |
| MDK | 0.561510599 | 0.035534712 |
| GALNT6 | 0.560909858 | 0.047933134 |
| GSTM1 | 0.560259239 | 0.038690494 |
| CYGB | 0.558425792 | 0.009830298 |
| ALYREF | 0.557451886 | 0.028675332 |
| NXN | 0.549239053 | 0.008096101 |
| RILPL2 | 0.549102003 | 0.023454819 |
| OSBPL5 | 0.548970638 | 0.025733749 |
| HSPBP1 | 0.547770181 | 0.032477536 |
| COPRS | 0.546575961 | 0.025907375 |
| SMTN | 0.545061038 | 0.030095569 |
| MARK2 | 0.544026209 | 0.039519201 |
| BRD9 | 0.543627899 | 0.03930486 |
| CCDC124 | 0.541924929 | 0.034327743 |
| PDE6D | 0.540096258 | 0.021789199 |
| PPME1 | 0.534394361 | 0.008815589 |
| QSOX1 | 0.532642035 | 0.032519207 |
| CAPN5 | 0.529702535 | 0.019443917 |
| DOT1L | 0.52834117 | 0.025947477 |
| SELO | 0.527912133 | 0.049134179 |
| BCL9L | 0.526398893 | 0.028186311 |
| FAM210A | 0.525902496 | 0.025265564 |
| NDUFAF4 | 0.520871568 | 0.014908893 |
| SMOX | 0.519000853 | 0.024751334 |
| SCARA3 | 0.518170586 | 0.027606357 |
| DUSP5 | 0.517660029 | 0.038662911 |

| | | |
|---|---|---|
| TCEA2 | 0.514877998 | 0.015247097 |
| GSTM4 | 0.51397783 | 0.040015485 |
| TUBG2 | 0.512262589 | 0.02319761 |
| UBL5 | 0.510834708 | 0.012162346 |
| TRIAP1 | 0.508406043 | 0.049742412 |
| A1BG-AS1 | 0.507374508 | 0.031951162 |
| MLLT11 | 0.506922624 | 0.038014974 |
| KNOP1 | 0.5066562 | 0.044784749 |
| FHOD1 | 0.503373143 | 0.048359977 |
| MLLT10 | 0.501021658 | 0.04304466 |
| CDKN1B | -0.501312582 | 0.034067664 |
| CDK11B | -0.503849455 | 0.028447938 |
| DCTN4 | -0.504416376 | 0.02841375 |
| SCFD1 | -0.51633709 | 0.039838533 |
| MICU2 | -0.517474484 | 0.037255768 |
| TJP2 | -0.52518556 | 0.024990918 |
| CSGALNACT | -0.527741225 | 0.034987882 |
| CHURC1 | -0.529905446 | 0.025170308 |
| TNRC18 | -0.530093352 | 0.046328322 |
| PABPC1P4 | -0.534691177 | 0.034606521 |
| ECT2 | -0.535046723 | 0.037913716 |
| PLK2 | -0.539637114 | 0.041309754 |
| ZNF268 | -0.55937345 | 0.037344034 |
| CEBPD | -0.567165464 | 0.021082071 |
| PSMC3IP | -0.567818628 | 0.040298766 |
| GBF1 | -0.578873903 | 0.011970335 |
| BCAR3 | -0.590454942 | 0.01222362 |
| PABPC3 | -0.596683053 | 0.017186043 |
| NPIPP1 | -0.605067444 | 0.039263294 |
| PTX3 | -0.636353441 | 0.013755719 |
| CYP1B1 | -0.641694045 | 0.01200071 |
| RASSF8 | -0.653407801 | 0.007700228 |
| FAT1 | -0.663879521 | 0.01378059 |
| AHRR | -0.665681981 | 0.009106916 |
| MASTL | -0.67569374 | 0.028141183 |
| ATG2B | -0.680192416 | 0.037713184 |
| RNF6 | -0.693438285 | 0.011947209 |
| SGK3 | -0.700085977 | 0.042143453 |
| MIR22HG | -0.713084755 | 0.011536574 |
| TNFRSF11B | -0.728493383 | 0.005591704 |
| PARP4 | -0.732261185 | 0.007394406 |
| POLQ | -0.742789991 | 0.022711709 |
| GBP1P1 | -0.774356532 | 0.002153516 |
| CDC20P1 | -0.825559235 | 0.024002719 |
| UPF3AP3 | -0.865109245 | 0.023704822 |
| RPL21P131 | -0.874400153 | 0.04139945 |
| CCDC41 | -0.877170589 | 0.017409509 |
| ATP5C1P1 | -0.895376273 | 0.035344298 |
| PHBP3 | -0.937630964 | 0.003128817 |
| FBXL6 | -0.961858136 | 0.044163759 |
| NUPL2 | -0.975150356 | 0.001401581 |
| TFB2M | -1.028695341 | 0.02662576 |
| AHCTF1P1 | -1.091694005 | 0.010692073 |
| ZNF441 | -1.112856703 | 0.044467987 |

| | | |
|---|---|---|
| DOCK11 | -1.277356184 | 0.003049582 |
| MEOX2 | -1.360857906 | 0.046353138 |
| MKI67IPP6 | -1.412555266 | 0.049707627 |
| RN7SL220P | -1.429987719 | 0.02879444 |
| RBMS1P1 | -1.432642793 | 0.045428443 |
| NPM1P27 | -1.444288384 | 0.023665315 |
| HIST1H1A | -1.816702431 | 0.012578806 |
| PPARG | -1.998490031 | 0.031585046 |
| HIST1H2AH | -2.196695044 | 0.042327575 |
| RPL21P127 | -2.213810125 | 0.023609331 |
| RNU6-379P | -2.442655833 | 0.048510766 |
| VIMP1 | -2.536273977 | 0.037925787 |
| NPM1P33 | -2.644199425 | 0.027189521 |
| YBX3P1 | -2.76199938 | 0.039371176 |
| UBE2V1P11 | -2.815036942 | 0.030037245 |
| TMEM213 | -3.216287933 | 0.026203881 |
| RPL21P66 | -3.40858582 | 0.040693572 |
| LINC01004 | -3.437701607 | 0.037924122 |
| ALMS1P | -3.802277009 | 0.023584167 |
| MYCBP2-AS1 | -4.223645447 | 0.012679225 |
| NPM1P18 | -4.958579241 | 0.000733933 |

| Table S3 \| Highly expressed genes for each NK cell cluster | | | | | | | |
|---|---|---|---|---|---|---|---|
| Target cluster | Gene name | Target cluster pct | Other cluster pct | Target cluster mean | Other cluster mean | log2FC | P value |
| 0 | ABHD17A | 0.392 | 0.199 | 2.149 | 1.228 | 0.808 | 2E-133 |
| 0 | ABI3 | 0.162 | 0.106 | 0.726 | 0.466 | 0.349 | 4.5E-21 |
| 0 | ABRACL | 0.2 | 0.148 | 0.897 | 0.798 | 0.169 | 3E-13 |
| 0 | ABT1 | 0.121 | 0.09 | 0.413 | 0.444 | 0.176 | 6.4E-09 |
| 0 | AC007384.1 | 0.174 | 0.087 | 0.824 | 0.492 | 0.744 | 6.4E-47 |
| 0 | AC007942.4 | 0.203 | 0.027 | 1.049 | 0.188 | 2.494 | 0 |
| 0 | AC011446.2 | 0.236 | 0.091 | 1.114 | 0.484 | 0.934 | 1E-124 |
| 0 | AC011476.3 | 0.116 | 0.048 | 0.406 | 0.36 | 0.49 | 3.4E-36 |
| 0 | AC012447.1 | 0.121 | 0.023 | 0.616 | 0.162 | 1.931 | 4E-144 |
| 0 | AC012614.1 | 0.128 | 0.037 | 0.464 | 0.213 | 1.407 | 3E-101 |
| 0 | AC016074.2 | 0.142 | 0.078 | 0.777 | 0.466 | 0.738 | 1.8E-34 |
| 0 | AC018816.1 | 0.104 | 0.044 | 0.48 | 0.244 | 0.918 | 4.5E-44 |
| 0 | AC022217.3 | 0.313 | 0.079 | 1.484 | 0.444 | 1.416 | 2E-299 |
| 0 | AC024171.1 | 0.161 | 0.084 | 0.782 | 0.482 | 0.699 | 3.8E-44 |
| 0 | AC024171.3 | 0.114 | 0.047 | 0.411 | 0.268 | 0.934 | 4.4E-44 |
| 0 | AC026979.2 | 0.141 | 0.044 | 0.637 | 0.247 | 1.368 | 2E-104 |
| 0 | AC044849.1 | 0.204 | 0.047 | 1.034 | 0.338 | 1.612 | 1E-176 |
| 0 | AC087286.2 | 0.217 | 0.044 | 1.128 | 0.338 | 1.738 | 9E-214 |
| 0 | AC092821.3 | 0.428 | 0.389 | 3.281 | 2.694 | 0.284 | 6.5E-41 |
| 0 | AC100778.3 | 0.126 | 0.078 | 0.433 | 0.438 | 0.284 | 3.5E-20 |
| 0 | AC103491.3 | 0.148 | 0.039 | 1.44 | 0.406 | 1.934 | 4E-144 |
| 0 | AC243472.2 | 0.283 | 0.064 | 2.489 | 0.489 | 2.078 | 0 |
| 0 | AC243829.4 | 0.164 | 0.086 | 0.814 | 0.406 | 0.687 | 3.9E-46 |
| 0 | AC244014.3 | 0.149 | 0.024 | 0.998 | 0.174 | 2.42 | 8E-226 |
| 0 | AC244407.2 | 0.12 | 0.074 | 0.434 | 0.403 | 0.406 | 9.2E-20 |
| 0 | ACAA2 | 0.313 | 0.209 | 1.487 | 1.222 | 0.376 | 1.8E-38 |
| 0 | ACAP1 | 0.194 | 0.139 | 0.89 | 0.784 | 0.183 | 1.5E-16 |
| 0 | ACP1 | 0.31 | 0.194 | 1.474 | 1.093 | 0.433 | 4.1E-48 |
| 0 | ACSL3 | 0.27 | 0.172 | 1.298 | 1.018 | 0.341 | 1.8E-38 |
| 0 | ACTB | 0.999 | 0.964 | 70.464 | 39.413 | 0.838 | 0 |
| 0 | ACTG1 | 0.794 | 0.448 | 8.044 | 4.128 | 0.64 | 2E-183 |
| 0 | ACTN4 | 0.272 | 0.199 | 1.323 | 1.213 | 0.124 | 1.5E-18 |
| 0 | ACTR2 | 0.641 | 0.419 | 4.489 | 3.994 | 0.168 | 1.3E-22 |
| 0 | ADD3 | 0.462 | 0.337 | 2.649 | 2.243 | 0.239 | 3.3E-34 |
| 0 | ADGRE4 | 0.449 | 0.318 | 2.783 | 2.336 | 0.243 | 2.2E-42 |
| 0 | ADGRG1 | 0.214 | 0.11 | 1.018 | 0.684 | 0.471 | 8.2E-61 |
| 0 | ADRB2 | 0.196 | 0.104 | 0.941 | 0.494 | 0.662 | 1.7E-42 |
| 0 | AF111167.1 | 0.234 | 0.039 | 1.311 | 0.268 | 2.29 | 0 |
| 0 | AGTPBP1 | 0.684 | 0.442 | 4.137 | 4.49 | 0.163 | 5E-29 |
| 0 | AGTRAP | 0.21 | 0.087 | 0.972 | 0.486 | 1 | 8E-101 |
| 0 | AHNAK | 0.446 | 0.334 | 2.714 | 2.418 | 0.108 | 3.2E-27 |
| 0 | AIMP1 | 0.164 | 0.113 | 0.747 | 0.494 | 0.331 | 1.8E-18 |
| 0 | AKAP13 | 0.684 | 0.472 | 4.344 | 4.912 | 0.122 | 2.9E-17 |
| 0 | AKAP17A | 0.142 | 0.104 | 0.673 | 0.464 | 0.243 | 2.7E-16 |
| 0 | AKIRIN2 | 0.147 | 0.101 | 0.698 | 0.442 | 0.364 | 3.1E-22 |
| 0 | AKNA | 0.428 | 0.301 | 2.497 | 2.146 | 0.212 | 2.4E-37 |
| 0 | AKR1C3 | 0.2 | 0.109 | 0.944 | 0.602 | 0.641 | 2.2E-40 |
| 0 | AL021144.4 | 0.127 | 0.02 | 0.62 | 0.147 | 2.074 | 2E-171 |
| 0 | AL118416.1 | 0.164 | 0.044 | 0.764 | 0.316 | 1.274 | 4E-113 |
| 0 | AL136018.1 | 0.44 | 0.296 | 2.439 | 1.893 | 0.364 | 6.6E-43 |

| | | | | | | | |
|---|---|---|---|---|---|---|---|
| 0 | AL136444.1 | 0.292 | 0.226 | 1.398 | 1.294 | 0.11 | 2.4E-13 |
| 0 | AL360012.1 | 0.124 | 0.02 | 0.633 | 0.146 | 2.121 | 4E-168 |
| 0 | AL390728.4 | 0.177 | 0.132 | 0.802 | 0.701 | 0.194 | 7.1E-12 |
| 0 | AL691403.1 | 0.222 | 0.044 | 1.172 | 0.324 | 1.842 | 2E-261 |
| 0 | ALDH9A1 | 0.108 | 0.074 | 0.464 | 0.399 | 0.219 | 2.8E-11 |
| 0 | ALKBH4 | 0.123 | 0.08 | 0.447 | 0.441 | 0.276 | 6.8E-17 |
| 0 | ALKBH7 | 0.106 | 0.082 | 0.463 | 0.412 | 0.169 | 3.4E-06 |
| 0 | AMD1 | 0.271 | 0.104 | 1.342 | 0.643 | 1.038 | 4E-147 |
| 0 | AMZ2 | 0.167 | 0.094 | 0.77 | 0.426 | 0.441 | 1.2E-37 |
| 0 | ANAPC11 | 0.206 | 0.134 | 0.901 | 0.704 | 0.344 | 4.7E-26 |
| 0 | ANAPC16 | 0.347 | 0.261 | 1.834 | 1.442 | 0.24 | 1.8E-26 |
| 0 | ANAPC4 | 0.306 | 0.237 | 1.492 | 1.374 | 0.118 | 4.8E-14 |
| 0 | ANKRD13A | 0.122 | 0.089 | 0.422 | 0.446 | 0.196 | 2.5E-09 |
| 0 | ANP32B | 0.361 | 0.282 | 1.877 | 1.704 | 0.139 | 1.4E-14 |
| 0 | ANP32E | 0.391 | 0.308 | 2.123 | 1.939 | 0.13 | 2.4E-14 |
| 0 | ANXA11 | 0.262 | 0.196 | 1.242 | 1.123 | 0.144 | 3.7E-16 |
| 0 | ANXA2 | 0.108 | 0.044 | 0.448 | 0.249 | 0.882 | 4.5E-41 |
| 0 | ANXA4 | 0.121 | 0.084 | 0.444 | 0.443 | 0.266 | 7.5E-12 |
| 0 | ANXA6 | 0.264 | 0.144 | 1.278 | 0.862 | 0.469 | 8.4E-67 |
| 0 | AP004329.1 | 0.11 | 0.048 | 0.447 | 0.33 | 0.47 | 2.4E-29 |
| 0 | AP2A1 | 0.144 | 0.109 | 0.684 | 0.604 | 0.176 | 6.2E-14 |
| 0 | AP2S1 | 0.214 | 0.143 | 0.949 | 0.79 | 0.264 | 1.4E-18 |
| 0 | AP3S1 | 0.166 | 0.124 | 0.718 | 0.664 | 0.114 | 4.5E-10 |
| 0 | APEX1 | 0.114 | 0.079 | 0.488 | 0.407 | 0.264 | 1.3E-11 |
| 0 | APH1A | 0.142 | 0.101 | 0.624 | 0.414 | 0.277 | 2E-13 |
| 0 | APLP2 | 0.19 | 0.124 | 0.872 | 0.707 | 0.301 | 2E-24 |
| 0 | APMAP | 0.49 | 0.466 | 3.784 | 3.428 | 0.143 | 4.7E-21 |
| 0 | APOO | 0.169 | 0.114 | 0.744 | 0.604 | 0.302 | 4.9E-20 |
| 0 | APRT | 0.299 | 0.191 | 1.449 | 1.033 | 0.488 | 2.4E-46 |
| 0 | AREG | 0.204 | 0.043 | 1.242 | 0.463 | 1.434 | 8E-190 |
| 0 | ARF1 | 0.433 | 0.416 | 3.17 | 2.807 | 0.174 | 8.4E-24 |
| 0 | ARF4 | 0.343 | 0.171 | 1.904 | 1.042 | 0.847 | 1E-130 |
| 0 | ARF6 | 0.296 | 0.21 | 1.424 | 1.184 | 0.264 | 1.5E-24 |
| 0 | ARHGAP12 | 0.16 | 0.102 | 0.734 | 0.483 | 0.334 | 2.3E-23 |
| 0 | ARHGAP9 | 0.374 | 0.243 | 2.043 | 1.441 | 0.414 | 3E-46 |
| 0 | ARHGDIA | 0.214 | 0.133 | 0.988 | 0.734 | 0.428 | 1.4E-33 |
| 0 | ARHGDIB | 0.807 | 0.494 | 7.629 | 4.26 | 0.437 | 3E-144 |
| 0 | ARID4A | 0.268 | 0.196 | 1.29 | 1.14 | 0.166 | 4E-19 |
| 0 | ARIH1 | 0.406 | 0.289 | 2.247 | 1.946 | 0.214 | 8E-34 |
| 0 | ARIH2 | 0.468 | 0.296 | 2.644 | 1.906 | 0.477 | 2.3E-78 |
| 0 | ARL4C | 0.481 | 0.347 | 3.938 | 2.448 | 0.68 | 4E-144 |
| 0 | ARL6IP1 | 0.444 | 0.329 | 2.438 | 2.023 | 0.269 | 1.1E-31 |
| 0 | ARL6IP4 | 0.272 | 0.217 | 1.313 | 1.222 | 0.104 | 4.7E-10 |
| 0 | ARPC1B | 0.449 | 0.374 | 3.423 | 2.493 | 0.448 | 4.4E-74 |
| 0 | ARPC2 | 0.878 | 0.728 | 9.94 | 7.607 | 0.386 | 9E-106 |
| 0 | ARPC3 | 0.79 | 0.622 | 6.827 | 4.373 | 0.346 | 1.6E-66 |
| 0 | ARPC4 | 0.464 | 0.44 | 3.42 | 3.176 | 0.148 | 2.8E-20 |
| 0 | ARPC4 | 0.372 | 0.192 | 1.908 | 1.107 | 0.784 | 1E-119 |
| 0 | ARPC4L | 0.411 | 0.248 | 2.202 | 1.444 | 0.402 | 4.5E-69 |
| 0 | ARRB2 | 0.283 | 0.193 | 1.384 | 1.104 | 0.324 | 3.9E-30 |
| 0 | ARRDC3 | 0.213 | 0.14 | 0.994 | 0.804 | 0.307 | 2.5E-26 |
| 0 | ARRDC3-AS1 | 0.101 | 0.041 | 0.467 | 0.299 | 0.644 | 1.4E-30 |
| 0 | ASAH1 | 0.147 | 0.098 | 0.648 | 0.431 | 0.286 | 2.2E-17 |
| 0 | ASTE1 | 0.127 | 0.08 | 0.477 | 0.423 | 0.447 | 8.5E-20 |

| | | | | | | | |
|---|---|---|---|---|---|---|---|
| 0 | ATF3 | 0.16 | 0.037 | 0.907 | 0.246 | 1.881 | 1E-165 |
| 0 | ATF4 | 0.139 | 0.094 | 0.614 | 0.412 | 0.26 | 1.1E-14 |
| 0 | ATOX1 | 0.19 | 0.134 | 0.844 | 0.734 | 0.218 | 2.1E-16 |
| 0 | ATP1B3 | 0.143 | 0.074 | 0.634 | 0.437 | 0.437 | 6.8E-40 |
| 0 | ATP4F1D | 0.416 | 0.317 | 2.164 | 1.906 | 0.184 | 1.2E-22 |
| 0 | ATP4F1E | 0.804 | 0.636 | 7.274 | 4.702 | 0.341 | 1.8E-74 |
| 0 | ATP4IF1 | 0.389 | 0.244 | 2.021 | 1.434 | 0.397 | 3.4E-42 |
| 0 | ATP4MC1 | 0.144 | 0.111 | 0.612 | 0.469 | 0.104 | 2.9E-08 |
| 0 | ATP4MC2 | 0.679 | 0.441 | 4.84 | 4.326 | 0.162 | 7.8E-21 |
| 0 | ATP4MC3 | 0.346 | 0.267 | 1.742 | 1.441 | 0.184 | 7.5E-21 |
| 0 | ATP4ME | 0.476 | 0.334 | 2.6 | 2.114 | 0.298 | 3.4E-44 |
| 0 | ATP4MG | 0.432 | 0.408 | 3.138 | 2.701 | 0.216 | 4.2E-30 |
| 0 | ATP4PB | 0.282 | 0.194 | 1.33 | 1.072 | 0.312 | 2.1E-28 |
| 0 | ATP4PF | 0.394 | 0.301 | 2.019 | 1.774 | 0.186 | 1.9E-20 |
| 0 | ATP4PO | 0.268 | 0.196 | 1.247 | 1.061 | 0.244 | 3.4E-19 |
| 0 | ATP6AP2 | 0.276 | 0.213 | 1.284 | 1.183 | 0.119 | 3.9E-13 |
| 0 | ATP6V0B | 0.226 | 0.148 | 1.019 | 0.842 | 0.248 | 2.4E-20 |
| 0 | ATP6V0C | 0.348 | 0.238 | 1.834 | 1.373 | 0.419 | 1.9E-46 |
| 0 | ATP6V0E1 | 0.389 | 0.302 | 2.026 | 1.801 | 0.17 | 1.8E-17 |
| 0 | ATP6V1G1 | 0.424 | 0.327 | 2.239 | 1.941 | 0.199 | 4.4E-21 |
| 0 | ATP8B4 | 0.162 | 0.104 | 0.747 | 0.697 | 0.118 | 1.8E-20 |
| 0 | ATXN7L3B | 0.426 | 0.28 | 2.342 | 1.73 | 0.443 | 4.5E-61 |
| 0 | AUP1 | 0.136 | 0.093 | 0.481 | 0.489 | 0.248 | 2.3E-14 |
| 0 | AURKAIP1 | 0.179 | 0.137 | 0.778 | 0.711 | 0.129 | 4.2E-10 |
| 0 | AXIN1 | 0.208 | 0.104 | 0.966 | 0.704 | 0.446 | 6.1E-62 |
| 0 | AZIN1 | 0.339 | 0.203 | 1.743 | 1.233 | 0.499 | 2.3E-64 |
| 0 | B2M | 1 | 0.997 | 217.622 | 200.23 | 0.12 | 3.4E-42 |
| 0 | B4GALT4 | 0.144 | 0.077 | 0.643 | 0.444 | 0.431 | 4.4E-37 |
| 0 | BANF1 | 0.231 | 0.14 | 1.064 | 0.814 | 0.386 | 3.1E-31 |
| 0 | BAX | 0.213 | 0.16 | 1.018 | 0.881 | 0.209 | 2.1E-13 |
| 0 | BCAP31 | 0.228 | 0.147 | 1.019 | 0.829 | 0.298 | 3.4E-23 |
| 0 | BCAS2 | 0.283 | 0.144 | 1.39 | 0.889 | 0.644 | 4.4E-72 |
| 0 | BCCIP | 0.143 | 0.107 | 0.62 | 0.462 | 0.141 | 1.2E-09 |
| 0 | BCL2 | 0.148 | 0.076 | 0.714 | 0.491 | 0.442 | 3.2E-42 |
| 0 | BCL6 | 0.132 | 0.102 | 0.613 | 0.471 | 0.102 | 1.1E-07 |
| 0 | BCL7B | 0.113 | 0.08 | 0.484 | 0.428 | 0.178 | 6.5E-10 |
| 0 | BCLAF1 | 0.464 | 0.388 | 3.667 | 2.741 | 0.42 | 1.1E-71 |
| 0 | BCR | 0.101 | 0.042 | 0.444 | 0.316 | 0.427 | 1.8E-44 |
| 0 | BEX4 | 0.108 | 0.062 | 0.463 | 0.316 | 0.441 | 1.8E-23 |
| 0 | BHLHE40 | 0.406 | 0.286 | 2.404 | 1.926 | 0.32 | 1.9E-38 |
| 0 | BHLHE40-AS1 | 0.14 | 0.064 | 0.643 | 0.363 | 0.822 | 3.2E-44 |
| 0 | BICRAL | 0.184 | 0.129 | 0.879 | 0.744 | 0.24 | 2.8E-17 |
| 0 | BID | 0.122 | 0.077 | 0.422 | 0.408 | 0.344 | 4.5E-19 |
| 0 | BIN1 | 0.177 | 0.084 | 0.773 | 0.461 | 0.744 | 2.7E-60 |
| 0 | BIN2 | 0.644 | 0.448 | 4.444 | 3.443 | 0.399 | 4.2E-76 |
| 0 | BLOC1S2 | 0.176 | 0.134 | 0.784 | 0.727 | 0.108 | 1E-09 |
| 0 | BLOC1S6 | 0.234 | 0.164 | 1.103 | 0.938 | 0.234 | 3.4E-21 |
| 0 | BNC2 | 0.234 | 0.147 | 1.204 | 1.016 | 0.246 | 2.9E-34 |
| 0 | BORCS7 | 0.107 | 0.083 | 0.48 | 0.444 | 0.114 | 4.9E-06 |
| 0 | BRD2 | 0.291 | 0.218 | 1.484 | 1.267 | 0.229 | 8.7E-19 |
| 0 | BRIX1 | 0.107 | 0.084 | 0.473 | 0.437 | 0.116 | 1.8E-05 |
| 0 | BRK1 | 0.244 | 0.176 | 1.143 | 0.931 | 0.296 | 2.8E-24 |
| 0 | BSG | 0.319 | 0.184 | 1.448 | 1.024 | 0.604 | 9.8E-71 |
| 0 | BTF3 | 0.632 | 0.497 | 4.314 | 3.741 | 0.202 | 3.3E-30 |

| | | | | | | | |
|---|---|---|---|---|---|---|---|
| 0 | BTG1 | 0.9 | 0.798 | 11.844 | 10.874 | 0.124 | 2.1E-15 |
| 0 | BTG2 | 0.899 | 0.617 | 14.219 | 6.988 | 1.123 | 0 |
| 0 | BTG3 | 0.144 | 0.097 | 0.691 | 0.444 | 0.346 | 1.4E-22 |
| 0 | BUD23 | 0.109 | 0.073 | 0.462 | 0.376 | 0.298 | 4.7E-13 |
| 0 | BUD31 | 0.263 | 0.162 | 1.234 | 0.884 | 0.483 | 8.8E-44 |
| 0 | BZW1 | 0.74 | 0.463 | 6.039 | 4.67 | 0.371 | 1.2E-74 |
| 0 | C12orf47 | 0.436 | 0.197 | 2.924 | 1.243 | 1.222 | 6E-216 |
| 0 | C12orf74 | 0.409 | 0.367 | 3.111 | 2.461 | 0.281 | 6.3E-41 |
| 0 | C16orf87 | 0.164 | 0.088 | 0.776 | 0.487 | 0.674 | 3.4E-44 |
| 0 | C18orf32 | 0.228 | 0.143 | 1.039 | 0.811 | 0.347 | 8.4E-27 |
| 0 | C19orf12 | 0.119 | 0.078 | 0.416 | 0.413 | 0.32 | 3.9E-16 |
| 0 | C19orf43 | 0.213 | 0.146 | 0.963 | 0.828 | 0.218 | 9.4E-16 |
| 0 | C1D | 0.231 | 0.173 | 1.069 | 0.946 | 0.176 | 2.4E-14 |
| 0 | C1orf162 | 0.317 | 0.096 | 1.683 | 0.484 | 1.424 | 9E-261 |
| 0 | C1orf174 | 0.113 | 0.078 | 0.494 | 0.414 | 0.243 | 4.1E-12 |
| 0 | C1orf43 | 0.144 | 0.096 | 0.634 | 0.482 | 0.396 | 1.1E-17 |
| 0 | C1QBP | 0.164 | 0.122 | 0.729 | 0.62 | 0.233 | 1.1E-11 |
| 0 | C4orf33 | 0.102 | 0.044 | 0.439 | 0.294 | 0.479 | 7.7E-26 |
| 0 | C9orf16 | 0.384 | 0.242 | 1.947 | 1.427 | 0.448 | 1.7E-42 |
| 0 | C9orf78 | 0.399 | 0.284 | 2.074 | 1.764 | 0.234 | 1.5E-32 |
| 0 | CA4B | 0.239 | 0.122 | 1.124 | 0.743 | 0.499 | 1E-69 |
| 0 | CACYBP | 0.202 | 0.148 | 0.94 | 0.874 | 0.12 | 3.2E-09 |
| 0 | CALCOCO2 | 0.237 | 0.179 | 1.093 | 0.996 | 0.133 | 3.1E-13 |
| 0 | CALM1 | 0.949 | 0.718 | 16.819 | 8.823 | 0.931 | 0 |
| 0 | CALM2 | 0.447 | 0.336 | 2.436 | 2.112 | 0.206 | 1.7E-26 |
| 0 | CALM3 | 0.197 | 0.136 | 0.89 | 0.749 | 0.249 | 4.1E-20 |
| 0 | CALR | 0.23 | 0.16 | 1.094 | 0.931 | 0.233 | 1.9E-21 |
| 0 | CAMLG | 0.163 | 0.118 | 0.724 | 0.634 | 0.191 | 4.9E-13 |
| 0 | CAND2 | 0.187 | 0.129 | 0.846 | 0.709 | 0.27 | 9.4E-19 |
| 0 | CANX | 0.431 | 0.338 | 2.412 | 2.2 | 0.133 | 6.4E-17 |
| 0 | CAP1 | 0.412 | 0.273 | 2.269 | 1.673 | 0.44 | 1.5E-44 |
| 0 | CAPNS1 | 0.193 | 0.103 | 0.864 | 0.448 | 0.649 | 1.4E-41 |
| 0 | CAPZB | 0.661 | 0.483 | 4.647 | 3.422 | 0.403 | 4.3E-72 |
| 0 | CAST | 0.817 | 0.662 | 7.924 | 6.612 | 0.261 | 1.6E-40 |
| 0 | CBLL1 | 0.19 | 0.134 | 0.868 | 0.746 | 0.198 | 4.9E-17 |
| 0 | CBX1 | 0.108 | 0.083 | 0.47 | 0.438 | 0.102 | 2E-06 |
| 0 | CBX4 | 0.17 | 0.129 | 0.781 | 0.727 | 0.102 | 1.8E-09 |
| 0 | CCDC107 | 0.113 | 0.074 | 0.478 | 0.371 | 0.364 | 4.5E-14 |
| 0 | CCDC124 | 0.144 | 0.084 | 0.617 | 0.442 | 0.481 | 9.8E-28 |
| 0 | CCDC173 | 0.144 | 0.031 | 0.697 | 0.204 | 1.776 | 4E-148 |
| 0 | CCDC200 | 0.239 | 0.048 | 1.846 | 0.436 | 1.791 | 4E-242 |
| 0 | CCDC49 | 0.166 | 0.113 | 0.741 | 0.632 | 0.247 | 4.5E-18 |
| 0 | CCDC69 | 0.413 | 0.372 | 2.994 | 2.484 | 0.27 | 4.8E-41 |
| 0 | CCDC84B | 0.288 | 0.204 | 1.367 | 1.143 | 0.249 | 2.2E-24 |
| 0 | CCL3 | 0.31 | 0.112 | 2.844 | 0.974 | 1.442 | 6E-206 |
| 0 | CCL4 | 0.734 | 0.464 | 13.634 | 6.713 | 1.022 | 7E-238 |
| 0 | CCL4L2 | 0.22 | 0.118 | 2.099 | 1.26 | 0.736 | 1.1E-48 |
| 0 | CCND2 | 0.397 | 0.264 | 2.248 | 1.779 | 0.337 | 1.5E-48 |
| 0 | CCND3 | 0.474 | 0.284 | 2.84 | 1.913 | 0.474 | 6E-103 |
| 0 | CCNG1 | 0.167 | 0.112 | 0.744 | 0.624 | 0.274 | 6E-19 |
| 0 | CCNI | 0.448 | 0.341 | 2.444 | 2.27 | 0.164 | 2.3E-19 |
| 0 | CCNK | 0.321 | 0.236 | 1.602 | 1.412 | 0.182 | 8.4E-21 |
| 0 | CCNL1 | 0.448 | 0.299 | 3.718 | 2.126 | 0.806 | 7E-184 |
| 0 | CCNT1 | 0.194 | 0.132 | 0.881 | 0.743 | 0.224 | 4E-21 |

| | | | | | | | |
|---|---|---|---|---|---|---|---|
| 0 | CCT3 | 0.19 | 0.14 | 0.844 | 0.714 | 0.241 | 4.4E-14 |
| 0 | CCT4 | 0.27 | 0.201 | 1.28 | 1.128 | 0.182 | 2.7E-17 |
| 0 | CCT4 | 0.169 | 0.13 | 0.73 | 0.676 | 0.111 | 3.9E-09 |
| 0 | CCT7 | 0.167 | 0.131 | 0.734 | 0.683 | 0.103 | 5E-08 |
| 0 | CD160 | 0.323 | 0.061 | 1.949 | 0.428 | 2.187 | 0 |
| 0 | CD226 | 0.333 | 0.204 | 1.794 | 1.344 | 0.416 | 3E-46 |
| 0 | CD244 | 0.149 | 0.111 | 0.717 | 0.611 | 0.23 | 2.8E-14 |
| 0 | CD300A | 0.248 | 0.148 | 1.272 | 0.928 | 0.444 | 2.7E-44 |
| 0 | CD37 | 0.479 | 0.364 | 2.721 | 2.386 | 0.19 | 2.4E-24 |
| 0 | CD42 | 0.401 | 0.314 | 3.284 | 3.002 | 0.129 | 1.8E-16 |
| 0 | CD43 | 0.692 | 0.421 | 4.197 | 4.119 | 0.334 | 6.4E-64 |
| 0 | CD44 | 0.311 | 0.138 | 1.672 | 0.902 | 0.891 | 4E-134 |
| 0 | CD48 | 0.498 | 0.386 | 2.883 | 2.484 | 0.148 | 1.4E-22 |
| 0 | CD63 | 0.346 | 0.264 | 1.846 | 1.604 | 0.21 | 4.3E-22 |
| 0 | CD69 | 0.793 | 0.422 | 13.064 | 4.198 | 1.638 | 0 |
| 0 | CD7 | 0.667 | 0.433 | 4.036 | 3.313 | 0.604 | 1E-141 |
| 0 | CD83 | 0.223 | 0.086 | 1.182 | 0.467 | 1.061 | 9E-124 |
| 0 | CD8A | 0.12 | 0.093 | 0.613 | 0.448 | 0.134 | 1.1E-06 |
| 0 | CD99 | 0.807 | 0.482 | 7.41 | 4.163 | 0.44 | 1E-148 |
| 0 | CDC37 | 0.368 | 0.287 | 1.864 | 1.736 | 0.103 | 1.1E-14 |
| 0 | CDC42 | 0.906 | 0.694 | 11.286 | 7.133 | 0.662 | 9E-280 |
| 0 | CDC42SE1 | 0.747 | 0.462 | 6.179 | 4.701 | 0.394 | 1.4E-83 |
| 0 | CDK2AP2 | 0.214 | 0.147 | 1.009 | 0.807 | 0.322 | 2.1E-23 |
| 0 | CDK6 | 0.162 | 0.108 | 0.769 | 0.646 | 0.23 | 4.7E-19 |
| 0 | CDKN2AIP | 0.182 | 0.134 | 0.84 | 0.712 | 0.238 | 2.8E-13 |
| 0 | CDKN2D | 0.306 | 0.146 | 1.464 | 0.914 | 0.774 | 4E-96 |
| 0 | CDV3 | 0.313 | 0.188 | 1.446 | 1.084 | 0.411 | 6.2E-49 |
| 0 | CEBPB | 0.169 | 0.08 | 0.801 | 0.419 | 0.936 | 9.9E-62 |
| 0 | CEBPZ | 0.332 | 0.243 | 1.713 | 1.443 | 0.141 | 9.2E-18 |
| 0 | CEMIP2 | 0.479 | 0.342 | 4.321 | 2.988 | 0.432 | 4E-129 |
| 0 | CEP78 | 0.489 | 0.262 | 3.096 | 1.801 | 0.782 | 1E-147 |
| 0 | CFL1 | 0.766 | 0.41 | 6.484 | 4.068 | 0.694 | 3E-204 |
| 0 | CHCHD2 | 0.49 | 0.341 | 2.762 | 2.144 | 0.366 | 1.3E-43 |
| 0 | CHCHD4 | 0.108 | 0.08 | 0.461 | 0.412 | 0.162 | 4.5E-08 |
| 0 | CHD1 | 0.446 | 0.389 | 3.707 | 2.914 | 0.347 | 4.3E-64 |
| 0 | CHD3 | 0.286 | 0.208 | 1.36 | 1.243 | 0.13 | 7E-19 |
| 0 | CHMP2A | 0.18 | 0.129 | 0.794 | 0.677 | 0.229 | 1.4E-14 |
| 0 | CHRAC1 | 0.18 | 0.12 | 0.808 | 0.649 | 0.316 | 1.9E-21 |
| 0 | CHST2 | 0.122 | 0.047 | 0.428 | 0.269 | 0.976 | 1.7E-64 |
| 0 | CIAO2B | 0.228 | 0.178 | 1.046 | 0.944 | 0.149 | 7.7E-11 |
| 0 | CIB1 | 0.279 | 0.204 | 1.324 | 1.144 | 0.21 | 6.5E-20 |
| 0 | CIRBP | 0.482 | 0.302 | 2.724 | 1.911 | 0.411 | 6.3E-84 |
| 0 | CISD2 | 0.164 | 0.127 | 0.774 | 0.683 | 0.181 | 1.4E-09 |
| 0 | CISD3 | 0.179 | 0.082 | 0.779 | 0.434 | 0.84 | 8.8E-70 |
| 0 | CITED2 | 0.186 | 0.094 | 0.924 | 0.431 | 0.799 | 4.6E-47 |
| 0 | CLIC1 | 0.733 | 0.472 | 4.816 | 4.743 | 0.291 | 6.3E-40 |
| 0 | CLK1 | 0.374 | 0.247 | 2.004 | 1.49 | 0.333 | 9.3E-47 |
| 0 | CLTB | 0.178 | 0.123 | 0.793 | 0.664 | 0.244 | 9.4E-18 |
| 0 | CMKLR1 | 0.127 | 0.044 | 0.496 | 0.247 | 1.214 | 6.3E-80 |
| 0 | CMPK1 | 0.374 | 0.284 | 2.009 | 1.708 | 0.234 | 1.4E-22 |
| 0 | CNBP | 0.491 | 0.39 | 2.747 | 2.428 | 0.124 | 7.6E-18 |
| 0 | CNN2 | 0.181 | 0.096 | 0.834 | 0.443 | 0.618 | 3.4E-49 |
| 0 | CNOT1 | 0.274 | 0.196 | 1.284 | 1.149 | 0.148 | 4E-21 |
| 0 | CNOT10 | 0.161 | 0.119 | 0.736 | 0.672 | 0.131 | 1.8E-11 |

| | | | | | | | |
|---|---|---|---|---|---|---|---|
| 0 | CNOT7 | 0.144 | 0.122 | 0.693 | 0.637 | 0.12 | 1.4E-07 |
| 0 | CNOT8 | 0.201 | 0.142 | 0.894 | 0.804 | 0.144 | 4.2E-17 |
| 0 | CNPPD1 | 0.114 | 0.081 | 0.412 | 0.421 | 0.28 | 1.3E-10 |
| 0 | CNPY3 | 0.14 | 0.103 | 0.649 | 0.446 | 0.243 | 1.5E-14 |
| 0 | COG3 | 0.146 | 0.117 | 0.688 | 0.633 | 0.12 | 2.7E-10 |
| 0 | COMMD4 | 0.116 | 0.067 | 0.499 | 0.349 | 0.474 | 1.7E-23 |
| 0 | COMMD6 | 0.633 | 0.476 | 4.193 | 3.438 | 0.244 | 8.8E-41 |
| 0 | COMMD7 | 0.233 | 0.16 | 1.041 | 0.873 | 0.268 | 2.3E-23 |
| 0 | COMT | 0.11 | 0.077 | 0.443 | 0.388 | 0.224 | 2.5E-10 |
| 0 | COPS2 | 0.203 | 0.144 | 0.922 | 0.811 | 0.184 | 7.1E-17 |
| 0 | COPS8 | 0.146 | 0.102 | 0.676 | 0.446 | 0.283 | 1.1E-19 |
| 0 | COPS9 | 0.308 | 0.223 | 1.449 | 1.23 | 0.246 | 7.6E-24 |
| 0 | COQ10B | 0.308 | 0.211 | 1.446 | 1.248 | 0.307 | 2E-31 |
| 0 | CORO1A | 0.443 | 0.37 | 3.489 | 2.416 | 0.472 | 1.4E-81 |
| 0 | COX16 | 0.166 | 0.113 | 0.724 | 0.494 | 0.286 | 3.5E-18 |
| 0 | COX19 | 0.104 | 0.08 | 0.446 | 0.418 | 0.122 | 1.3E-06 |
| 0 | COX20 | 0.171 | 0.121 | 0.769 | 0.671 | 0.197 | 1E-14 |
| 0 | COX4A | 0.314 | 0.23 | 1.408 | 1.27 | 0.247 | 2E-23 |
| 0 | COX4B | 0.396 | 0.269 | 2.033 | 1.414 | 0.424 | 2.7E-48 |
| 0 | COX6A1 | 0.467 | 0.443 | 3.427 | 2.974 | 0.204 | 1.1E-24 |
| 0 | COX6B1 | 0.474 | 0.342 | 2.603 | 2.074 | 0.327 | 3.3E-41 |
| 0 | COX6C | 0.46 | 0.391 | 3.347 | 2.43 | 0.408 | 6.7E-68 |
| 0 | COX7A2 | 0.441 | 0.347 | 2.381 | 2.188 | 0.122 | 4E-16 |
| 0 | COX7A2L | 0.198 | 0.141 | 0.894 | 0.789 | 0.182 | 1.2E-11 |
| 0 | COX7B | 0.284 | 0.206 | 1.321 | 1.101 | 0.264 | 1.3E-22 |
| 0 | COX8A | 0.487 | 0.342 | 2.77 | 2.198 | 0.334 | 3.4E-44 |
| 0 | CPB2-AS1 | 0.32 | 0.241 | 1.492 | 1.424 | 0.16 | 2.8E-18 |
| 0 | CPD | 0.181 | 0.116 | 0.837 | 0.694 | 0.269 | 1.2E-24 |
| 0 | CPQ | 0.27 | 0.186 | 1.382 | 1.162 | 0.24 | 1.3E-26 |
| 0 | CRBN | 0.244 | 0.197 | 1.224 | 1.132 | 0.113 | 1.8E-12 |
| 0 | CRCP | 0.273 | 0.17 | 1.324 | 1.014 | 0.384 | 1.1E-43 |
| 0 | CREM | 0.264 | 0.14 | 1.376 | 0.831 | 0.727 | 9.6E-74 |
| 0 | CRIP1 | 0.484 | 0.393 | 4.411 | 3.279 | 0.428 | 2.5E-82 |
| 0 | CRLF1 | 0.12 | 0.087 | 0.416 | 0.467 | 0.142 | 2.7E-09 |
| 0 | CROCC | 0.422 | 0.122 | 3.207 | 0.884 | 1.848 | 0 |
| 0 | CRY1 | 0.309 | 0.149 | 1.487 | 1.067 | 0.473 | 4E-103 |
| 0 | CS | 0.12 | 0.084 | 0.426 | 0.447 | 0.204 | 1.6E-11 |
| 0 | CSK | 0.227 | 0.168 | 1.047 | 0.921 | 0.184 | 3.6E-14 |
| 0 | CSNK2B | 0.411 | 0.284 | 2.167 | 1.677 | 0.369 | 3.1E-44 |
| 0 | CSRNP1 | 0.16 | 0.046 | 0.732 | 0.33 | 1.148 | 8E-100 |
| 0 | CST7 | 0.944 | 0.794 | 14.491 | 11.344 | 0.448 | 2E-164 |
| 0 | CSTB | 0.221 | 0.177 | 1.042 | 0.968 | 0.107 | 1E-08 |
| 0 | CTBP1 | 0.144 | 0.076 | 0.634 | 0.43 | 0.46 | 2.4E-39 |
| 0 | CTBP2 | 0.243 | 0.169 | 1.17 | 1.034 | 0.177 | 4.3E-22 |
| 0 | CTSC | 0.614 | 0.443 | 4.26 | 3.384 | 0.332 | 3.4E-44 |
| 0 | CTSD | 0.333 | 0.212 | 1.649 | 1.219 | 0.436 | 2.9E-40 |
| 0 | CTSW | 0.913 | 0.631 | 13.782 | 7.477 | 0.882 | 0 |
| 0 | CUTA | 0.332 | 0.188 | 1.648 | 1.074 | 0.626 | 6.8E-78 |
| 0 | CX3CR1 | 0.434 | 0.194 | 2.766 | 1.284 | 1.104 | 1E-203 |
| 0 | CXXC4 | 0.247 | 0.149 | 1.212 | 0.843 | 0.424 | 1.1E-44 |
| 0 | CYB461D2 | 0.118 | 0.073 | 0.409 | 0.383 | 0.41 | 2.4E-19 |
| 0 | CYB4B | 0.214 | 0.164 | 0.994 | 0.907 | 0.131 | 3.7E-11 |
| 0 | CYBA | 0.82 | 0.644 | 8.141 | 6.444 | 0.317 | 1.3E-61 |
| 0 | CYC1 | 0.137 | 0.104 | 0.492 | 0.44 | 0.133 | 3.1E-08 |

| | | | | | | | |
|---|---|---|---|---|---|---|---|
| 0 | CYCS | 0.39 | 0.206 | 2.149 | 1.194 | 0.847 | 1E-123 |
| 0 | CYFIP2 | 0.34 | 0.243 | 1.813 | 1.494 | 0.186 | 6.8E-24 |
| 0 | CYP41A1 | 0.194 | 0.081 | 0.897 | 0.466 | 0.946 | 3.4E-92 |
| 0 | CYTIP | 0.421 | 0.336 | 2.342 | 2.179 | 0.104 | 2.7E-14 |
| 0 | CYTOR | 0.246 | 0.194 | 1.338 | 1.176 | 0.186 | 4.3E-14 |
| 0 | DAD1 | 0.294 | 0.224 | 1.404 | 1.24 | 0.167 | 9.7E-16 |
| 0 | DAZAP1 | 0.136 | 0.088 | 0.607 | 0.481 | 0.334 | 7.1E-19 |
| 0 | DAZAP2 | 0.429 | 0.379 | 3.169 | 2.428 | 0.326 | 4.1E-42 |
| 0 | DBF4 | 0.284 | 0.148 | 1.423 | 0.94 | 0.498 | 6.1E-80 |
| 0 | DBI | 0.498 | 0.274 | 2.908 | 1.646 | 0.821 | 1E-146 |
| 0 | DBNL | 0.142 | 0.087 | 0.622 | 0.462 | 0.428 | 2.5E-24 |
| 0 | DCTN3 | 0.149 | 0.102 | 0.636 | 0.43 | 0.263 | 2.1E-16 |
| 0 | DDIT3 | 0.149 | 0.069 | 0.672 | 0.383 | 0.812 | 2E-46 |
| 0 | DDIT4 | 0.388 | 0.166 | 2.283 | 1.116 | 1.033 | 2E-192 |
| 0 | DDOST | 0.211 | 0.132 | 0.941 | 0.698 | 0.431 | 7.5E-34 |
| 0 | DDT | 0.183 | 0.143 | 0.818 | 0.744 | 0.118 | 7.2E-09 |
| 0 | DDX21 | 0.333 | 0.204 | 1.717 | 1.277 | 0.427 | 4.2E-47 |
| 0 | DDX27 | 0.234 | 0.167 | 1.107 | 0.949 | 0.207 | 6.7E-19 |
| 0 | DDX39A | 0.217 | 0.101 | 1.01 | 0.646 | 0.644 | 2.3E-79 |
| 0 | DDX3Y | 0.203 | 0.092 | 0.988 | 0.434 | 0.888 | 1E-80 |
| 0 | DDX4 | 0.746 | 0.478 | 6.186 | 4.01 | 0.304 | 1.5E-48 |
| 0 | DENND1C | 0.147 | 0.097 | 0.64 | 0.47 | 0.189 | 4.2E-18 |
| 0 | DENND2D | 0.204 | 0.129 | 0.948 | 0.731 | 0.39 | 1.4E-30 |
| 0 | DENND3 | 0.114 | 0.067 | 0.4 | 0.404 | 0.308 | 9.9E-23 |
| 0 | DENR | 0.247 | 0.199 | 1.21 | 1.117 | 0.114 | 3.4E-12 |
| 0 | DGKD | 0.404 | 0.296 | 2.149 | 1.964 | 0.129 | 8.1E-24 |
| 0 | DGKZ | 0.18 | 0.118 | 0.801 | 0.642 | 0.298 | 1.4E-22 |
| 0 | DHRS3 | 0.148 | 0.098 | 0.676 | 0.464 | 0.249 | 4E-18 |
| 0 | DHRS7 | 0.419 | 0.402 | 3.094 | 2.696 | 0.199 | 1.7E-24 |
| 0 | DHX40 | 0.172 | 0.112 | 0.766 | 0.636 | 0.268 | 2.5E-22 |
| 0 | DHX9 | 0.344 | 0.243 | 1.809 | 1.438 | 0.234 | 7.4E-28 |
| 0 | DIAPH1 | 0.436 | 0.394 | 3.344 | 2.844 | 0.229 | 2.4E-39 |
| 0 | DIP2A | 0.466 | 0.377 | 3.714 | 2.837 | 0.389 | 9.4E-83 |
| 0 | DIPK2A | 0.123 | 0.092 | 0.448 | 0.407 | 0.111 | 1.4E-08 |
| 0 | DNAJA1 | 0.411 | 0.369 | 3.188 | 2.643 | 0.264 | 2.5E-41 |
| 0 | DNAJB1 | 0.314 | 0.133 | 1.746 | 0.844 | 1.048 | 1E-144 |
| 0 | DNAJB11 | 0.148 | 0.103 | 0.697 | 0.44 | 0.342 | 2.3E-21 |
| 0 | DNAJB9 | 0.129 | 0.062 | 0.447 | 0.338 | 0.694 | 1.6E-44 |
| 0 | DNAJC3 | 0.343 | 0.262 | 1.79 | 1.636 | 0.13 | 1.1E-20 |
| 0 | DNAJC8 | 0.362 | 0.281 | 1.843 | 1.709 | 0.117 | 1.5E-14 |
| 0 | DNM1L | 0.246 | 0.193 | 1.233 | 1.139 | 0.114 | 2.2E-14 |
| 0 | DNTTIP2 | 0.274 | 0.194 | 1.34 | 1.124 | 0.243 | 4.9E-23 |
| 0 | DOCK4 | 0.204 | 0.14 | 1.078 | 0.99 | 0.123 | 1.4E-20 |
| 0 | DOK2 | 0.3 | 0.122 | 1.41 | 0.716 | 1.074 | 2E-146 |
| 0 | DPH3 | 0.149 | 0.106 | 0.718 | 0.46 | 0.348 | 1.5E-19 |
| 0 | DSTN | 0.488 | 0.28 | 2.949 | 1.908 | 0.634 | 2E-119 |
| 0 | DUSP1 | 0.604 | 0.142 | 4.448 | 1.417 | 1.969 | 0 |
| 0 | DUSP2 | 0.473 | 0.306 | 4.421 | 2.822 | 0.968 | 2E-218 |
| 0 | DYNLL2 | 0.112 | 0.046 | 0.478 | 0.284 | 0.747 | 2.9E-34 |
| 0 | ECD | 0.304 | 0.172 | 1.429 | 1.06 | 0.429 | 1.9E-69 |
| 0 | EEF1A1 | 0.992 | 0.946 | 29.946 | 24.891 | 0.21 | 4.1E-80 |
| 0 | EEF1B2 | 0.648 | 0.44 | 4.649 | 3.321 | 0.486 | 1.4E-98 |
| 0 | EEF1D | 0.714 | 0.468 | 4.274 | 4.721 | 0.16 | 2.3E-26 |
| 0 | EEF1G | 0.416 | 0.319 | 2.213 | 1.987 | 0.146 | 4.3E-20 |

| | | | | | | | |
|---|---|---|---|---|---|---|---|
| 0 | EFCAB2 | 0.102 | 0.048 | 0.444 | 0.286 | 0.667 | 2.5E-37 |
| 0 | EFHD2 | 0.767 | 0.461 | 6.486 | 4.821 | 0.44 | 9E-108 |
| 0 | EGR1 | 0.106 | 0.027 | 0.614 | 0.206 | 1.474 | 6.1E-97 |
| 0 | EID1 | 0.319 | 0.23 | 1.46 | 1.329 | 0.232 | 2.8E-24 |
| 0 | EIF1 | 0.949 | 0.749 | 18.373 | 9.993 | 0.879 | 0 |
| 0 | EIF1AX | 0.276 | 0.212 | 1.394 | 1.242 | 0.166 | 1.4E-14 |
| 0 | EIF2A | 0.192 | 0.134 | 0.873 | 0.733 | 0.242 | 2E-18 |
| 0 | EIF2AK1 | 0.247 | 0.167 | 1.163 | 0.934 | 0.314 | 1.4E-27 |
| 0 | EIF2AK3 | 0.138 | 0.08 | 0.623 | 0.474 | 0.392 | 4.9E-28 |
| 0 | EIF2D | 0.116 | 0.082 | 0.412 | 0.444 | 0.206 | 1.4E-10 |
| 0 | EIF3D | 0.14 | 0.108 | 0.499 | 0.447 | 0.104 | 1.2E-07 |
| 0 | EIF3F | 0.24 | 0.184 | 1.112 | 1.012 | 0.136 | 9.9E-13 |
| 0 | EIF3G | 0.374 | 0.223 | 2.148 | 1.293 | 0.732 | 4.7E-84 |
| 0 | EIF3H | 0.393 | 0.304 | 2.078 | 1.842 | 0.166 | 7.1E-18 |
| 0 | EIF3I | 0.192 | 0.114 | 0.848 | 0.481 | 0.444 | 4.2E-36 |
| 0 | EIF3K | 0.438 | 0.391 | 3.127 | 2.464 | 0.286 | 1.9E-42 |
| 0 | EIF3L | 0.143 | 0.097 | 0.617 | 0.404 | 0.288 | 2.3E-16 |
| 0 | EIF4A | 0.371 | 0.228 | 1.933 | 1.341 | 0.427 | 3.7E-68 |
| 0 | EIF4A1 | 0.302 | 0.184 | 1.476 | 1.042 | 0.488 | 4E-44 |
| 0 | EIF4A2 | 0.284 | 0.196 | 1.364 | 1.132 | 0.27 | 1E-28 |
| 0 | EIF4A3 | 0.163 | 0.094 | 0.739 | 0.431 | 0.477 | 1.2E-33 |
| 0 | EIF4EBP2 | 0.147 | 0.104 | 0.684 | 0.494 | 0.206 | 1.7E-18 |
| 0 | EIF4G2 | 0.471 | 0.374 | 2.694 | 2.468 | 0.127 | 3.2E-16 |
| 0 | EIF4G3 | 0.394 | 0.278 | 2.179 | 1.94 | 0.168 | 9.8E-33 |
| 0 | EIF6 | 0.164 | 0.122 | 0.724 | 0.622 | 0.221 | 9.3E-12 |
| 0 | ELL2 | 0.242 | 0.103 | 1.244 | 0.763 | 0.704 | 6E-107 |
| 0 | ELOB | 0.404 | 0.384 | 2.822 | 2.473 | 0.19 | 4.7E-26 |
| 0 | ELOC | 0.207 | 0.149 | 0.937 | 0.846 | 0.147 | 2.7E-11 |
| 0 | ELOVL1 | 0.102 | 0.064 | 0.43 | 0.323 | 0.414 | 4.7E-14 |
| 0 | EMC10 | 0.149 | 0.096 | 0.642 | 0.497 | 0.369 | 6.7E-21 |
| 0 | EMC3 | 0.184 | 0.143 | 0.84 | 0.764 | 0.138 | 1.8E-09 |
| 0 | EMC4 | 0.176 | 0.122 | 0.766 | 0.638 | 0.264 | 2.7E-17 |
| 0 | EMC7 | 0.134 | 0.106 | 0.493 | 0.441 | 0.104 | 4.6E-07 |
| 0 | EMD | 0.161 | 0.112 | 0.711 | 0.486 | 0.279 | 4.4E-16 |
| 0 | EMP3 | 0.628 | 0.441 | 4.406 | 3.432 | 0.393 | 2.1E-74 |
| 0 | ENO1 | 0.41 | 0.31 | 2.237 | 1.903 | 0.234 | 4.1E-24 |
| 0 | ENO3 | 0.161 | 0.086 | 0.694 | 0.491 | 0.401 | 1.3E-40 |
| 0 | ENOPH1 | 0.107 | 0.087 | 0.474 | 0.442 | 0.1 | 0.00012 |
| 0 | ENSA | 0.447 | 0.294 | 2.416 | 1.898 | 0.407 | 3.4E-62 |
| 0 | ENY2 | 0.203 | 0.141 | 0.942 | 0.762 | 0.321 | 5E-21 |
| 0 | EPC1 | 0.444 | 0.449 | 3.403 | 3.264 | 0.102 | 1.5E-16 |
| 0 | EPS14L1 | 0.136 | 0.092 | 0.494 | 0.401 | 0.246 | 3.5E-15 |
| 0 | ERCC1 | 0.168 | 0.064 | 0.796 | 0.369 | 1.11 | 2.4E-92 |
| 0 | ERVK3-1 | 0.162 | 0.119 | 0.708 | 0.643 | 0.139 | 1.8E-11 |
| 0 | ESYT2 | 0.346 | 0.141 | 1.906 | 1.216 | 0.649 | 3E-147 |
| 0 | ETF1 | 0.327 | 0.194 | 1.666 | 1.192 | 0.482 | 3.5E-62 |
| 0 | ETV3 | 0.201 | 0.122 | 0.913 | 0.678 | 0.429 | 9.3E-34 |
| 0 | EZH2 | 0.16 | 0.082 | 0.738 | 0.437 | 0.46 | 1.6E-44 |
| 0 | FAAP20 | 0.137 | 0.084 | 0.492 | 0.441 | 0.424 | 1.8E-22 |
| 0 | FAM133B | 0.29 | 0.207 | 1.444 | 1.247 | 0.211 | 8.2E-23 |
| 0 | FAM136A | 0.124 | 0.101 | 0.448 | 0.419 | 0.104 | 4.8E-05 |
| 0 | FAM169A | 0.144 | 0.103 | 0.71 | 0.603 | 0.236 | 4E-18 |
| 0 | FAM174C | 0.104 | 0.064 | 0.447 | 0.322 | 0.474 | 1.7E-17 |
| 0 | FAM177A1 | 0.247 | 0.176 | 1.188 | 0.998 | 0.241 | 1.2E-20 |

| | | | | | | | |
|---|---|---|---|---|---|---|---|
| 0 | FAM199X | 0.147 | 0.099 | 0.68 | 0.42 | 0.386 | 4E-24 |
| 0 | FAM32A | 0.136 | 0.081 | 0.491 | 0.426 | 0.472 | 1E-24 |
| 0 | FAM43C | 0.141 | 0.06 | 0.629 | 0.362 | 0.798 | 8E-61 |
| 0 | FAM49B | 0.779 | 0.661 | 6.964 | 6.311 | 0.142 | 2.3E-24 |
| 0 | FAR1 | 0.278 | 0.201 | 1.371 | 1.232 | 0.144 | 1.4E-20 |
| 0 | FAU | 0.987 | 0.906 | 21.073 | 18.682 | 0.174 | 1.8E-44 |
| 0 | FBL | 0.127 | 0.086 | 0.447 | 0.448 | 0.314 | 1.2E-14 |
| 0 | FBXL3 | 0.144 | 0.1 | 0.646 | 0.444 | 0.221 | 2.7E-14 |
| 0 | FBXO33 | 0.238 | 0.147 | 1.166 | 0.949 | 0.297 | 6.2E-38 |
| 0 | FBXW4 | 0.103 | 0.067 | 0.434 | 0.343 | 0.346 | 2.8E-14 |
| 0 | FCER1G | 0.792 | 0.418 | 8.374 | 3.664 | 1.192 | 0 |
| 0 | FCGR3A | 0.779 | 0.404 | 7.938 | 3.469 | 1.143 | 0 |
| 0 | FCMR | 0.148 | 0.07 | 0.643 | 0.39 | 0.746 | 4.9E-43 |
| 0 | FCRL3 | 0.203 | 0.111 | 0.99 | 0.643 | 0.499 | 1.2E-40 |
| 0 | FCRL6 | 0.343 | 0.134 | 1.793 | 0.844 | 1.07 | 2E-190 |
| 0 | FDFT1 | 0.192 | 0.141 | 0.883 | 0.762 | 0.213 | 2.4E-14 |
| 0 | FERMT3 | 0.228 | 0.143 | 1.043 | 0.779 | 0.421 | 9.9E-36 |
| 0 | FEZ1 | 0.146 | 0.067 | 0.734 | 0.379 | 0.942 | 1.3E-69 |
| 0 | FGFBP2 | 0.722 | 0.327 | 6.846 | 2.848 | 1.266 | 0 |
| 0 | FGR | 0.189 | 0.061 | 0.878 | 0.39 | 1.172 | 8E-133 |
| 0 | FIS1 | 0.266 | 0.187 | 1.282 | 1.017 | 0.333 | 1.3E-24 |
| 0 | FKBP11 | 0.24 | 0.16 | 1.214 | 0.914 | 0.411 | 1.5E-34 |
| 0 | FKBP1A | 0.143 | 0.093 | 0.619 | 0.468 | 0.404 | 9.7E-20 |
| 0 | FLNA | 0.292 | 0.148 | 1.442 | 1.044 | 0.462 | 1.2E-74 |
| 0 | FOS | 0.816 | 0.274 | 36.939 | 8.622 | 2.099 | 0 |
| 0 | FOSB | 0.104 | 0.026 | 0.404 | 0.174 | 1.423 | 9E-100 |
| 0 | FOSL2 | 0.21 | 0.11 | 0.993 | 0.664 | 0.479 | 4.9E-48 |
| 0 | FRG1 | 0.193 | 0.14 | 0.878 | 0.778 | 0.174 | 1.4E-14 |
| 0 | FSD1L | 0.137 | 0.098 | 0.601 | 0.431 | 0.177 | 8E-12 |
| 0 | FTL | 0.733 | 0.444 | 6.119 | 4.47 | 0.421 | 1.4E-87 |
| 0 | FUNDC2 | 0.143 | 0.077 | 0.617 | 0.409 | 0.492 | 2.9E-36 |
| 0 | FUS | 0.421 | 0.394 | 3.048 | 2.706 | 0.172 | 4.1E-28 |
| 0 | FUT11 | 0.114 | 0.067 | 0.412 | 0.373 | 0.444 | 1.6E-22 |
| 0 | FXYD4 | 0.646 | 0.474 | 4.644 | 3.848 | 0.267 | 1.2E-42 |
| 0 | G3BP2 | 0.404 | 0.307 | 2.134 | 1.871 | 0.191 | 1.6E-21 |
| 0 | GADD44B | 0.644 | 0.368 | 6.368 | 2.896 | 1.137 | 4E-270 |
| 0 | GADD44GIP1 | 0.118 | 0.084 | 0.496 | 0.424 | 0.227 | 3.9E-10 |
| 0 | GALC | 0.174 | 0.113 | 0.782 | 0.634 | 0.303 | 1.2E-22 |
| 0 | GAPDH | 0.841 | 0.644 | 8.986 | 6.768 | 0.409 | 3E-104 |
| 0 | GARS | 0.173 | 0.13 | 0.766 | 0.713 | 0.103 | 2.4E-10 |
| 0 | GAS7 | 0.241 | 0.164 | 1.329 | 1.029 | 0.37 | 2.2E-33 |
| 0 | GGNBP2 | 0.349 | 0.261 | 1.833 | 1.603 | 0.193 | 8.9E-24 |
| 0 | GHITM | 0.336 | 0.269 | 1.674 | 1.448 | 0.103 | 7.8E-12 |
| 0 | GK4 | 0.376 | 0.278 | 2.047 | 1.844 | 0.14 | 1.4E-22 |
| 0 | GLIPR2 | 0.461 | 0.346 | 2.634 | 2.342 | 0.163 | 1.2E-20 |
| 0 | GLRX | 0.232 | 0.177 | 1.081 | 1.002 | 0.11 | 4.7E-12 |
| 0 | GLS | 0.46 | 0.436 | 3.618 | 3.233 | 0.162 | 3.9E-26 |
| 0 | GLTP | 0.182 | 0.106 | 0.836 | 0.488 | 0.407 | 4.5E-37 |
| 0 | GLUD1 | 0.24 | 0.163 | 1.124 | 0.917 | 0.294 | 3.9E-26 |
| 0 | GMFB | 0.103 | 0.072 | 0.43 | 0.382 | 0.173 | 1.4E-10 |
| 0 | GNAI2 | 0.304 | 0.183 | 1.413 | 1.048 | 0.429 | 4.4E-48 |
| 0 | GNAL | 0.177 | 0.106 | 0.902 | 0.698 | 0.37 | 8.3E-31 |
| 0 | GNAS | 0.422 | 0.32 | 2.378 | 2.119 | 0.166 | 8.3E-22 |
| 0 | GNB2 | 0.248 | 0.17 | 1.209 | 0.948 | 0.34 | 1.2E-31 |

| | | | | | | | |
|---|---|---|---|---|---|---|---|
| 0 | GNG2 | 0.902 | 0.724 | 12.314 | 8.741 | 0.494 | 8E-172 |
| 0 | GNL2 | 0.124 | 0.087 | 0.446 | 0.479 | 0.191 | 2.2E-11 |
| 0 | GNL3L | 0.104 | 0.074 | 0.438 | 0.387 | 0.176 | 1.8E-09 |
| 0 | GNLY | 0.973 | 0.937 | 63.037 | 46.691 | 0.433 | 1E-114 |
| 0 | GPR64 | 0.423 | 0.244 | 2.473 | 1.447 | 0.668 | 4E-101 |
| 0 | GPS2 | 0.34 | 0.233 | 1.631 | 1.341 | 0.283 | 4.4E-34 |
| 0 | GPSM3 | 0.373 | 0.228 | 1.946 | 1.321 | 0.466 | 1.3E-71 |
| 0 | GRASP | 0.104 | 0.024 | 0.449 | 0.172 | 1.414 | 9E-103 |
| 0 | GRPEL1 | 0.133 | 0.087 | 0.478 | 0.444 | 0.379 | 1.2E-17 |
| 0 | GSAP | 0.474 | 0.248 | 2.819 | 1.689 | 0.739 | 3E-144 |
| 0 | GSTP1 | 0.649 | 0.444 | 4.682 | 3.249 | 0.427 | 4E-104 |
| 0 | GTF2B | 0.324 | 0.203 | 1.681 | 1.201 | 0.486 | 7.5E-44 |
| 0 | GTF3A | 0.472 | 0.283 | 2.73 | 1.779 | 0.618 | 8E-104 |
| 0 | GTF3C1 | 0.28 | 0.193 | 1.329 | 1.126 | 0.239 | 2.8E-27 |
| 0 | GTF3C6 | 0.114 | 0.084 | 0.402 | 0.432 | 0.219 | 4.3E-09 |
| 0 | GUK1 | 0.406 | 0.289 | 2.162 | 1.728 | 0.323 | 4.7E-37 |
| 0 | GYPC | 0.341 | 0.241 | 1.729 | 1.396 | 0.309 | 9.7E-31 |
| 0 | GZMA | 0.791 | 0.449 | 8.201 | 4.274 | 0.94 | 0 |
| 0 | GZMB | 0.946 | 0.784 | 20.118 | 12.822 | 0.64 | 2E-284 |
| 0 | GZMH | 0.644 | 0.399 | 4.838 | 3.691 | 0.662 | 2E-142 |
| 0 | GZMM | 0.494 | 0.31 | 4.004 | 2.107 | 0.926 | 2E-236 |
| 0 | H1FX | 0.349 | 0.218 | 2.948 | 1.649 | 0.834 | 3.7E-78 |
| 0 | H2AFJ | 0.144 | 0.076 | 0.724 | 0.43 | 0.742 | 1.4E-41 |
| 0 | H2AFV | 0.396 | 0.3 | 2.114 | 1.888 | 0.164 | 2.7E-21 |
| 0 | H2AFZ | 0.249 | 0.184 | 1.262 | 1.024 | 0.299 | 3.1E-22 |
| 0 | H3F3A | 0.397 | 0.273 | 2.094 | 1.667 | 0.329 | 3E-42 |
| 0 | H3F3B | 0.968 | 0.848 | 23.47 | 14.346 | 0.716 | 0 |
| 0 | HACD4 | 0.202 | 0.131 | 0.937 | 0.733 | 0.343 | 2E-27 |
| 0 | HAUS2 | 0.129 | 0.083 | 0.444 | 0.443 | 0.289 | 6.5E-18 |
| 0 | HAVCR2 | 0.171 | 0.104 | 0.816 | 0.612 | 0.414 | 1.5E-29 |
| 0 | HCST | 0.764 | 0.484 | 7.064 | 3.989 | 0.824 | 2E-248 |
| 0 | HERPUD1 | 0.281 | 0.194 | 1.42 | 1.172 | 0.277 | 1.2E-27 |
| 0 | HIPK1 | 0.3 | 0.189 | 1.481 | 1.147 | 0.369 | 3.1E-46 |
| 0 | HIST1H1B | 0.144 | 0.043 | 0.899 | 0.364 | 1.302 | 2E-100 |
| 0 | HIST1H1C | 0.468 | 0.324 | 4.408 | 2.376 | 0.924 | 2E-187 |
| 0 | HIST1H1D | 0.79 | 0.414 | 11.909 | 6.33 | 0.912 | 1E-264 |
| 0 | HIST1H1E | 0.901 | 0.768 | 14.403 | 11.297 | 0.34 | 3.9E-71 |
| 0 | HIST1H2AJ | 0.137 | 0.043 | 0.749 | 0.273 | 1.477 | 7E-103 |
| 0 | HIST1H2AM | 0.144 | 0.044 | 0.708 | 0.288 | 1.297 | 9.4E-94 |
| 0 | HIST1H4C | 0.734 | 0.446 | 6.487 | 4.014 | 0.394 | 1E-71 |
| 0 | HIST2H2AB | 0.102 | 0.047 | 0.498 | 0.36 | 0.469 | 4.9E-24 |
| 0 | HLA-E | 0.976 | 0.897 | 19.88 | 16.88 | 0.236 | 1.1E-69 |
| 0 | HMGB2 | 0.401 | 0.301 | 2.263 | 1.972 | 0.199 | 4.9E-23 |
| 0 | HMGCS1 | 0.238 | 0.12 | 1.174 | 0.692 | 0.763 | 3.8E-76 |
| 0 | HMGN1 | 0.494 | 0.404 | 2.944 | 2.746 | 0.1 | 3.2E-13 |
| 0 | HMGN2 | 0.42 | 0.39 | 3.108 | 2.638 | 0.237 | 5E-33 |
| 0 | HMGXB3 | 0.13 | 0.094 | 0.467 | 0.407 | 0.161 | 4.9E-10 |
| 0 | HMOX2 | 0.184 | 0.116 | 0.844 | 0.646 | 0.363 | 2.2E-28 |
| 0 | HNRNPA0 | 0.399 | 0.244 | 2.121 | 1.491 | 0.409 | 8.6E-64 |
| 0 | HNRNPA2B1 | 0.901 | 0.749 | 11.264 | 8.441 | 0.416 | 4E-130 |
| 0 | HNRNPA3 | 0.438 | 0.408 | 3.333 | 2.844 | 0.223 | 3.4E-32 |
| 0 | HNRNPAB | 0.321 | 0.238 | 1.608 | 1.416 | 0.184 | 3.5E-20 |
| 0 | HNRNPC | 0.443 | 0.34 | 2.483 | 2.289 | 0.117 | 4.9E-21 |
| 0 | HNRNPDL | 0.886 | 0.664 | 10.344 | 6.763 | 0.613 | 6E-230 |

| | | | | | | | |
|---|---|---|---|---|---|---|---|
| 0 | HNRNPH1 | 0.344 | 0.24 | 1.821 | 1.484 | 0.201 | 1.5E-29 |
| 0 | HNRNPH2 | 0.184 | 0.129 | 0.822 | 0.668 | 0.299 | 1.2E-18 |
| 0 | HNRNPL | 0.296 | 0.177 | 1.486 | 1.064 | 0.48 | 1.1E-44 |
| 0 | HNRNPM | 0.418 | 0.406 | 3.127 | 2.814 | 0.142 | 2.3E-21 |
| 0 | HNRNPR | 0.346 | 0.283 | 1.83 | 1.702 | 0.104 | 2.9E-12 |
| 0 | HNRNPUL1 | 0.274 | 0.182 | 1.306 | 1.049 | 0.303 | 2E-32 |
| 0 | HOOK2 | 0.241 | 0.097 | 1.184 | 0.617 | 0.942 | 9E-123 |
| 0 | HOPX | 0.67 | 0.372 | 4.347 | 2.903 | 0.884 | 7E-244 |
| 0 | HPCAL1 | 0.112 | 0.068 | 0.484 | 0.374 | 0.371 | 4.5E-20 |
| 0 | HSF2 | 0.107 | 0.064 | 0.468 | 0.334 | 0.487 | 4.6E-19 |
| 0 | HSP90AA1 | 0.641 | 0.474 | 4.739 | 3.486 | 0.402 | 4.1E-64 |
| 0 | HSP90AB1 | 0.774 | 0.498 | 7.332 | 4.619 | 0.667 | 2E-212 |
| 0 | HSP90B1 | 0.724 | 0.41 | 6.129 | 4.417 | 0.472 | 4E-111 |
| 0 | HSPA4 | 0.77 | 0.494 | 8.344 | 4.412 | 0.887 | 5E-264 |
| 0 | HSPA9 | 0.281 | 0.203 | 1.34 | 1.188 | 0.184 | 1.4E-20 |
| 0 | HSPB1 | 0.104 | 0.049 | 0.441 | 0.311 | 0.437 | 2.4E-24 |
| 0 | HSPD1 | 0.288 | 0.19 | 1.44 | 1.148 | 0.337 | 1.9E-34 |
| 0 | HSPE1 | 0.214 | 0.118 | 1.032 | 0.668 | 0.627 | 2.3E-43 |
| 0 | HSPH1 | 0.184 | 0.104 | 0.879 | 0.618 | 0.408 | 6.1E-40 |
| 0 | ICAM2 | 0.174 | 0.103 | 0.78 | 0.463 | 0.47 | 1.3E-32 |
| 0 | ICAM3 | 0.194 | 0.107 | 0.883 | 0.471 | 0.628 | 8.8E-48 |
| 0 | ID2 | 0.498 | 0.236 | 3.244 | 1.6 | 1.02 | 1E-221 |
| 0 | IDI1 | 0.304 | 0.174 | 1.481 | 1.026 | 0.624 | 7.7E-69 |
| 0 | IDS | 0.426 | 0.321 | 2.341 | 2.044 | 0.189 | 1.4E-23 |
| 0 | IER2 | 0.881 | 0.447 | 18.839 | 6.98 | 1.432 | 0 |
| 0 | IER4 | 0.101 | 0.039 | 0.44 | 0.218 | 1.044 | 1.1E-44 |
| 0 | IFI27L2 | 0.187 | 0.126 | 0.823 | 0.663 | 0.311 | 1.4E-20 |
| 0 | IFITM2 | 0.89 | 0.737 | 11.987 | 9.71 | 0.304 | 1E-69 |
| 0 | IFNG | 0.181 | 0.043 | 1.194 | 0.342 | 1.764 | 1E-184 |
| 0 | IFNGR1 | 0.37 | 0.201 | 2.143 | 1.314 | 0.704 | 2E-101 |
| 0 | IFRD1 | 0.379 | 0.177 | 2.337 | 1.292 | 0.844 | 1E-143 |
| 0 | IGF2R | 0.288 | 0.187 | 1.431 | 1.198 | 0.246 | 1.2E-36 |
| 0 | IGFBP7 | 0.394 | 0.169 | 2.317 | 1.084 | 1.094 | 1E-198 |
| 0 | IL27RA | 0.168 | 0.12 | 0.777 | 0.67 | 0.216 | 2E-14 |
| 0 | IL2RB | 0.289 | 0.194 | 1.44 | 1.173 | 0.296 | 7.4E-32 |
| 0 | ILF2 | 0.228 | 0.164 | 1.048 | 0.872 | 0.266 | 7.3E-19 |
| 0 | ILF3-DT | 0.161 | 0.093 | 0.726 | 0.498 | 0.444 | 2E-34 |
| 0 | ILKAP | 0.213 | 0.143 | 0.991 | 0.897 | 0.144 | 3.1E-16 |
| 0 | ING3 | 0.109 | 0.079 | 0.481 | 0.419 | 0.198 | 1.3E-09 |
| 0 | INSIG1 | 0.203 | 0.113 | 0.97 | 0.636 | 0.608 | 3.4E-48 |
| 0 | IQCB1 | 0.178 | 0.137 | 0.831 | 0.773 | 0.104 | 1.9E-09 |
| 0 | IQCG | 0.147 | 0.113 | 0.681 | 0.609 | 0.162 | 7.8E-13 |
| 0 | IQGAP1 | 0.781 | 0.644 | 6.639 | 4.933 | 0.162 | 1.2E-24 |
| 0 | IRF2BP2 | 0.142 | 0.098 | 0.688 | 0.447 | 0.306 | 2.1E-20 |
| 0 | IRS2 | 0.107 | 0.043 | 0.488 | 0.289 | 0.747 | 1.8E-42 |
| 0 | ISCA1 | 0.244 | 0.144 | 1.193 | 0.899 | 0.409 | 1.4E-42 |
| 0 | ISG20L2 | 0.126 | 0.084 | 0.444 | 0.439 | 0.337 | 5E-14 |
| 0 | ISY1 | 0.142 | 0.109 | 0.661 | 0.489 | 0.167 | 1.4E-12 |
| 0 | ITGAM | 0.148 | 0.048 | 0.649 | 0.306 | 1.083 | 3E-102 |
| 0 | ITGAX | 0.197 | 0.064 | 0.978 | 0.433 | 1.174 | 5E-137 |
| 0 | ITGB2 | 0.727 | 0.329 | 6.366 | 2.774 | 1.198 | 0 |
| 0 | ITGB7 | 0.169 | 0.071 | 0.746 | 0.392 | 0.929 | 7.8E-78 |
| 0 | ITM2A | 0.144 | 0.107 | 0.641 | 0.483 | 0.148 | 1.3E-10 |
| 0 | ITPRIP | 0.109 | 0.046 | 0.477 | 0.284 | 0.74 | 6.1E-48 |

| | | | | | | | |
|---|---|---|---|---|---|---|---|
| 0 | IVNS1ABP | 0.242 | 0.169 | 1.173 | 1.011 | 0.214 | 5E-22 |
| 0 | JAK1 | 0.908 | 0.803 | 11.983 | 9.947 | 0.267 | 3.7E-46 |
| 0 | JMJD6 | 0.168 | 0.098 | 0.774 | 0.446 | 0.478 | 2.6E-33 |
| 0 | JMY | 0.128 | 0.087 | 0.442 | 0.403 | 0.136 | 1.8E-13 |
| 0 | JOSD1 | 0.377 | 0.238 | 2.09 | 1.403 | 0.476 | 3E-61 |
| 0 | JPT1 | 0.224 | 0.14 | 1.066 | 0.782 | 0.448 | 3.2E-36 |
| 0 | JTB | 0.247 | 0.193 | 1.142 | 1.064 | 0.114 | 9.9E-11 |
| 0 | JUN | 0.814 | 0.37 | 18.847 | 4.137 | 1.874 | 0 |
| 0 | JUNB | 0.697 | 0.282 | 7.413 | 2.437 | 1.466 | 0 |
| 0 | JUND | 0.302 | 0.098 | 1.613 | 0.643 | 1.304 | 6E-220 |
| 0 | KANSL2 | 0.199 | 0.131 | 0.912 | 0.747 | 0.288 | 1.2E-24 |
| 0 | KAT7 | 0.233 | 0.173 | 1.064 | 0.962 | 0.144 | 6.1E-14 |
| 0 | KDELR1 | 0.189 | 0.14 | 0.849 | 0.784 | 0.131 | 2.5E-08 |
| 0 | KDM6B | 0.143 | 0.049 | 0.646 | 0.281 | 1.199 | 4.3E-94 |
| 0 | KIAA1143 | 0.212 | 0.149 | 1.003 | 0.904 | 0.149 | 4.7E-13 |
| 0 | KIR2DL1 | 0.189 | 0.074 | 1.124 | 0.487 | 1.208 | 5E-99 |
| 0 | KIR2DL3 | 0.168 | 0.097 | 0.913 | 0.6 | 0.607 | 4.4E-36 |
| 0 | KIR3DL1 | 0.18 | 0.094 | 1.048 | 0.497 | 0.826 | 1.4E-41 |
| 0 | KIR3DL2 | 0.126 | 0.086 | 0.614 | 0.491 | 0.323 | 2.6E-14 |
| 0 | KLF10 | 0.247 | 0.134 | 1.322 | 0.799 | 0.727 | 1.2E-66 |
| 0 | KLF2 | 0.283 | 0.074 | 1.478 | 0.447 | 1.434 | 2E-248 |
| 0 | KLF3 | 0.42 | 0.224 | 2.404 | 1.444 | 0.734 | 6E-127 |
| 0 | KLF6 | 0.808 | 0.436 | 11.937 | 4.433 | 1.136 | 0 |
| 0 | KLF9 | 0.214 | 0.107 | 1.016 | 0.604 | 0.74 | 2.9E-69 |
| 0 | KLHDC4 | 0.172 | 0.106 | 0.781 | 0.639 | 0.289 | 1.6E-27 |
| 0 | KLHL28 | 0.174 | 0.132 | 0.79 | 0.734 | 0.107 | 2.7E-10 |
| 0 | KLRB1 | 0.676 | 0.438 | 4.676 | 3.622 | 0.648 | 2E-142 |
| 0 | KLRF1 | 0.648 | 0.434 | 4.773 | 3.422 | 0.48 | 2E-103 |
| 0 | KLRK1 | 0.739 | 0.619 | 6.34 | 4.863 | 0.113 | 3.4E-14 |
| 0 | KMT2E-AS1 | 0.264 | 0.073 | 1.387 | 0.448 | 1.63 | 9E-234 |
| 0 | KRT10 | 0.311 | 0.234 | 1.433 | 1.304 | 0.234 | 2.8E-19 |
| 0 | KRTCAP2 | 0.344 | 0.27 | 1.747 | 1.44 | 0.19 | 2.9E-19 |
| 0 | KXD1 | 0.136 | 0.102 | 0.491 | 0.442 | 0.1 | 4.2E-09 |
| 0 | LAIR1 | 0.129 | 0.072 | 0.466 | 0.382 | 0.466 | 9.3E-30 |
| 0 | LAIR2 | 0.313 | 0.09 | 1.843 | 0.488 | 1.648 | 2E-274 |
| 0 | LAMP1 | 0.23 | 0.171 | 1.063 | 0.973 | 0.127 | 6.4E-14 |
| 0 | LAMTOR1 | 0.243 | 0.176 | 1.118 | 0.949 | 0.222 | 1.8E-18 |
| 0 | LAMTOR4 | 0.366 | 0.273 | 1.819 | 1.634 | 0.144 | 4.9E-21 |
| 0 | LAMTOR4 | 0.174 | 0.121 | 0.771 | 0.628 | 0.294 | 8.3E-18 |
| 0 | LAPTM4 | 0.464 | 0.399 | 3.477 | 2.744 | 0.342 | 4.6E-47 |
| 0 | LAT | 0.204 | 0.124 | 0.924 | 0.666 | 0.473 | 7.1E-36 |
| 0 | LAT2 | 0.166 | 0.124 | 0.744 | 0.649 | 0.174 | 4.2E-11 |
| 0 | LATS2 | 0.231 | 0.146 | 1.066 | 0.881 | 0.274 | 6E-32 |
| 0 | LCK | 0.304 | 0.23 | 1.414 | 1.377 | 0.137 | 3.2E-16 |
| 0 | LCP1 | 0.962 | 0.843 | 17.062 | 13.427 | 0.346 | 4E-111 |
| 0 | LGALS1 | 0.399 | 0.184 | 2.824 | 1.308 | 1.11 | 4E-180 |
| 0 | LIME1 | 0.133 | 0.046 | 0.486 | 0.272 | 1.108 | 1.9E-84 |
| 0 | LINC00299 | 0.389 | 0.236 | 2.082 | 1.434 | 0.44 | 1.1E-69 |
| 0 | LINC00861 | 0.264 | 0.174 | 1.312 | 1.064 | 0.3 | 4.5E-31 |
| 0 | LINC00910 | 0.327 | 0.087 | 2.112 | 0.624 | 1.746 | 0 |
| 0 | LINC01004 | 0.143 | 0.091 | 0.642 | 0.418 | 0.331 | 4.9E-21 |
| 0 | LINC01278 | 0.214 | 0.129 | 1.003 | 0.801 | 0.324 | 9.2E-37 |
| 0 | LINC01404 | 0.161 | 0.096 | 0.843 | 0.719 | 0.247 | 1.4E-28 |
| 0 | LINC01478 | 0.178 | 0.117 | 0.794 | 0.684 | 0.216 | 3.8E-22 |

| | | | | | | | |
|---|---|---|---|---|---|---|---|
| 0 | LINC01934 | 0.141 | 0.108 | 0.744 | 0.7 | 0.107 | 1.4E-12 |
| 0 | LITAF | 0.841 | 0.446 | 9.488 | 4.406 | 0.811 | 0 |
| 0 | LLGL2 | 0.179 | 0.088 | 0.821 | 0.42 | 0.648 | 6.9E-48 |
| 0 | LLPH | 0.136 | 0.097 | 0.617 | 0.424 | 0.231 | 4.5E-12 |
| 0 | LMAN2 | 0.244 | 0.186 | 1.142 | 1.029 | 0.141 | 1.8E-13 |
| 0 | LPCAT1 | 0.341 | 0.262 | 1.872 | 1.674 | 0.16 | 4.4E-21 |
| 0 | LRPAP1 | 0.218 | 0.168 | 1.016 | 0.918 | 0.146 | 2.5E-11 |
| 0 | LRRC48 | 0.101 | 0.076 | 0.436 | 0.39 | 0.164 | 3.6E-07 |
| 0 | LRRC8C | 0.12 | 0.078 | 0.441 | 0.468 | 0.211 | 4.9E-16 |
| 0 | LRRFIP1 | 0.604 | 0.444 | 4.084 | 3.416 | 0.216 | 4.6E-40 |
| 0 | LSM10 | 0.124 | 0.087 | 0.421 | 0.428 | 0.284 | 4.9E-13 |
| 0 | LSM12 | 0.179 | 0.122 | 0.796 | 0.632 | 0.333 | 4.1E-20 |
| 0 | LSM3 | 0.168 | 0.121 | 0.741 | 0.644 | 0.198 | 1.9E-13 |
| 0 | LSM4 | 0.138 | 0.078 | 0.489 | 0.414 | 0.407 | 4.7E-31 |
| 0 | LSM7 | 0.244 | 0.178 | 1.179 | 0.942 | 0.308 | 6.2E-24 |
| 0 | LSP1 | 0.627 | 0.487 | 4.144 | 3.723 | 0.144 | 1.8E-26 |
| 0 | LYN | 0.437 | 0.36 | 3.372 | 2.666 | 0.339 | 1.3E-70 |
| 0 | LYPLA1 | 0.164 | 0.119 | 0.739 | 0.636 | 0.217 | 6.5E-13 |
| 0 | LYSMD3 | 0.23 | 0.176 | 1.092 | 1.01 | 0.112 | 3.4E-12 |
| 0 | M6PR | 0.334 | 0.241 | 1.684 | 1.424 | 0.241 | 9.2E-26 |
| 0 | MAF | 0.138 | 0.087 | 0.64 | 0.406 | 0.34 | 2.9E-21 |
| 0 | MAF1 | 0.116 | 0.08 | 0.402 | 0.422 | 0.24 | 4.3E-12 |
| 0 | MAFF | 0.191 | 0.074 | 0.906 | 0.469 | 0.94 | 7E-101 |
| 0 | MAGOH | 0.266 | 0.187 | 1.242 | 1.034 | 0.276 | 4.4E-24 |
| 0 | MAGT1 | 0.193 | 0.144 | 0.88 | 0.816 | 0.11 | 6.4E-12 |
| 0 | MAN1A1 | 0.327 | 0.234 | 1.681 | 1.488 | 0.176 | 4.1E-24 |
| 0 | MANF | 0.109 | 0.07 | 0.464 | 0.349 | 0.37 | 3.2E-14 |
| 0 | MAP1LC3B | 0.692 | 0.413 | 4.16 | 3.847 | 0.42 | 2.2E-80 |
| 0 | MAP3K8 | 0.649 | 0.334 | 4.788 | 2.822 | 1.036 | 0 |
| 0 | MAPK1IP1L | 0.312 | 0.18 | 1.608 | 1.097 | 0.442 | 2.1E-68 |
| 0 | MAPRE1 | 0.227 | 0.14 | 1.039 | 0.842 | 0.286 | 1.5E-27 |
| 0 | MAPRE2 | 0.473 | 0.374 | 3.964 | 2.787 | 0.408 | 2E-104 |
| 0 | MARCKSL1 | 0.104 | 0.04 | 0.476 | 0.232 | 1.034 | 4E-47 |
| 0 | MARS | 0.162 | 0.103 | 0.727 | 0.463 | 0.368 | 2E-24 |
| 0 | MATK | 0.286 | 0.142 | 1.434 | 0.834 | 0.783 | 1.3E-94 |
| 0 | MCRIP1 | 0.12 | 0.074 | 0.414 | 0.382 | 0.432 | 2.5E-19 |
| 0 | MDH2 | 0.249 | 0.169 | 1.184 | 0.91 | 0.382 | 2.6E-34 |
| 0 | MECOM | 0.112 | 0.071 | 0.477 | 0.348 | 0.412 | 1.5E-17 |
| 0 | MED10 | 0.231 | 0.148 | 1.039 | 0.844 | 0.283 | 1.8E-23 |
| 0 | MED30 | 0.191 | 0.126 | 0.846 | 0.678 | 0.334 | 2.5E-24 |
| 0 | MED4 | 0.223 | 0.172 | 1.046 | 0.964 | 0.117 | 2.6E-11 |
| 0 | METAP1 | 0.137 | 0.097 | 0.604 | 0.432 | 0.184 | 2.4E-12 |
| 0 | METRNL | 0.146 | 0.062 | 0.661 | 0.377 | 0.81 | 1.3E-64 |
| 0 | MEX3C | 0.103 | 0.064 | 0.438 | 0.363 | 0.272 | 7E-17 |
| 0 | MIB2 | 0.144 | 0.048 | 0.692 | 0.323 | 1.098 | 4.2E-86 |
| 0 | MICOS13 | 0.148 | 0.109 | 0.636 | 0.449 | 0.186 | 4.5E-11 |
| 0 | MIDN | 0.134 | 0.069 | 0.61 | 0.384 | 0.67 | 3.7E-40 |
| 0 | MIEN1 | 0.244 | 0.16 | 1.136 | 0.866 | 0.391 | 2.5E-32 |
| 0 | MIF | 0.43 | 0.382 | 3.141 | 2.422 | 0.321 | 2.8E-46 |
| 0 | MIF4GD | 0.124 | 0.099 | 0.448 | 0.406 | 0.114 | 4.2E-06 |
| 0 | MIR181A1HG | 0.146 | 0.09 | 0.694 | 0.444 | 0.34 | 1.7E-24 |
| 0 | MIR4434-2HG | 0.119 | 0.079 | 0.44 | 0.493 | 0.147 | 2.1E-14 |
| 0 | MIR646HG | 0.117 | 0.044 | 0.436 | 0.349 | 0.619 | 1.6E-49 |
| 0 | MIR924HG | 0.278 | 0.194 | 1.328 | 1.116 | 0.241 | 9.7E-24 |

| | | | | | | | |
|---|---|---|---|---|---|---|---|
| 0 | MLH1 | 0.102 | 0.074 | 0.44 | 0.404 | 0.143 | 1.3E-07 |
| 0 | MMGT1 | 0.12 | 0.088 | 0.411 | 0.467 | 0.13 | 3.7E-09 |
| 0 | MOB2 | 0.117 | 0.074 | 0.49 | 0.394 | 0.309 | 3.8E-16 |
| 0 | MOB4 | 0.26 | 0.198 | 1.214 | 1.103 | 0.139 | 2.2E-14 |
| 0 | MORF4L1 | 0.6 | 0.449 | 3.841 | 3.14 | 0.29 | 6.3E-46 |
| 0 | MORF4L2 | 0.266 | 0.133 | 1.301 | 0.789 | 0.722 | 4.4E-87 |
| 0 | MPC2 | 0.148 | 0.116 | 0.698 | 0.603 | 0.21 | 6.6E-12 |
| 0 | MRPL10 | 0.222 | 0.129 | 1.022 | 0.716 | 0.414 | 1.9E-44 |
| 0 | MRPL22 | 0.102 | 0.079 | 0.444 | 0.412 | 0.106 | 7.4E-06 |
| 0 | MRPL41 | 0.147 | 0.119 | 0.692 | 0.612 | 0.178 | 4.5E-10 |
| 0 | MRPL42 | 0.143 | 0.112 | 0.644 | 0.467 | 0.207 | 1.1E-11 |
| 0 | MRPL44 | 0.116 | 0.074 | 0.476 | 0.369 | 0.366 | 2.5E-17 |
| 0 | MRPL44 | 0.102 | 0.062 | 0.424 | 0.317 | 0.42 | 4.8E-18 |
| 0 | MRPL9 | 0.132 | 0.094 | 0.476 | 0.404 | 0.188 | 1.5E-11 |
| 0 | MRPS24 | 0.106 | 0.068 | 0.448 | 0.334 | 0.417 | 7.8E-16 |
| 0 | MSL2 | 0.247 | 0.179 | 1.213 | 1.077 | 0.172 | 4.5E-18 |
| 0 | MTPN | 0.481 | 0.341 | 2.746 | 2.249 | 0.293 | 7.8E-39 |
| 0 | MTRF1L | 0.123 | 0.089 | 0.436 | 0.468 | 0.196 | 2.9E-10 |
| 0 | MVD | 0.14 | 0.069 | 0.613 | 0.387 | 0.663 | 1.4E-44 |
| 0 | MXD1 | 0.147 | 0.094 | 0.647 | 0.42 | 0.339 | 5E-21 |
| 0 | MYADM | 0.179 | 0.067 | 0.892 | 0.443 | 1.009 | 2E-99 |
| 0 | MYBL1 | 0.464 | 0.389 | 3.861 | 2.964 | 0.382 | 6.7E-68 |
| 0 | MYDGF | 0.202 | 0.122 | 0.918 | 0.642 | 0.492 | 1.9E-36 |
| 0 | MYL12A | 0.876 | 0.644 | 10.374 | 6.642 | 0.641 | 4E-237 |
| 0 | MYL6 | 0.862 | 0.7 | 9.212 | 7.218 | 0.342 | 1.1E-80 |
| 0 | MYLIP | 0.141 | 0.064 | 0.691 | 0.411 | 0.749 | 2.4E-67 |
| 0 | MYO1G | 0.144 | 0.047 | 0.683 | 0.34 | 0.964 | 2.7E-87 |
| 0 | MYOM2 | 0.32 | 0.048 | 2.644 | 0.443 | 2.262 | 0 |
| 0 | MZT1 | 0.134 | 0.098 | 0.469 | 0.413 | 0.149 | 3.5E-10 |
| 0 | MZT2B | 0.177 | 0.133 | 0.791 | 0.68 | 0.218 | 1.4E-11 |
| 0 | NAA38 | 0.219 | 0.141 | 0.979 | 0.764 | 0.344 | 4.3E-29 |
| 0 | NAAA | 0.104 | 0.069 | 0.44 | 0.366 | 0.298 | 4.5E-14 |
| 0 | NACA | 0.849 | 0.707 | 8.946 | 7.362 | 0.281 | 3.5E-49 |
| 0 | NAMPT | 0.144 | 0.111 | 0.708 | 0.64 | 0.146 | 2.7E-12 |
| 0 | NASP | 0.189 | 0.14 | 0.874 | 0.781 | 0.164 | 6.3E-13 |
| 0 | NAXE | 0.141 | 0.092 | 0.622 | 0.471 | 0.402 | 1.4E-19 |
| 0 | NCBP2AS2 | 0.149 | 0.101 | 0.708 | 0.432 | 0.412 | 4.4E-24 |
| 0 | NCL | 0.799 | 0.674 | 7.469 | 6.437 | 0.192 | 1.5E-28 |
| 0 | NCR3 | 0.288 | 0.06 | 1.43 | 0.382 | 2.001 | 0 |
| 0 | NDE1 | 0.123 | 0.09 | 0.437 | 0.484 | 0.147 | 9.5E-10 |
| 0 | NDFIP1 | 0.318 | 0.234 | 1.473 | 1.411 | 0.147 | 8.5E-21 |
| 0 | NDUFA1 | 0.417 | 0.39 | 2.903 | 2.464 | 0.236 | 1.1E-31 |
| 0 | NDUFA11 | 0.309 | 0.237 | 1.462 | 1.313 | 0.144 | 4.9E-16 |
| 0 | NDUFA12 | 0.304 | 0.18 | 1.444 | 1.011 | 0.424 | 1E-49 |
| 0 | NDUFA13 | 0.363 | 0.279 | 1.798 | 1.604 | 0.163 | 4.1E-18 |
| 0 | NDUFA2 | 0.21 | 0.146 | 0.934 | 0.822 | 0.184 | 1.4E-13 |
| 0 | NDUFA3 | 0.41 | 0.393 | 2.924 | 2.418 | 0.216 | 2.1E-26 |
| 0 | NDUFA4 | 0.433 | 0.304 | 2.328 | 1.834 | 0.344 | 4.1E-43 |
| 0 | NDUFA7 | 0.213 | 0.169 | 0.964 | 0.884 | 0.126 | 6.1E-09 |
| 0 | NDUFAF8 | 0.119 | 0.079 | 0.407 | 0.418 | 0.279 | 1.8E-14 |
| 0 | NDUFB11 | 0.32 | 0.241 | 1.429 | 1.334 | 0.197 | 1E-18 |
| 0 | NDUFB2 | 0.416 | 0.304 | 3.041 | 1.908 | 0.672 | 1E-126 |
| 0 | NDUFB6 | 0.144 | 0.104 | 0.64 | 0.449 | 0.216 | 1E-11 |
| 0 | NDUFB7 | 0.291 | 0.148 | 1.346 | 0.883 | 0.619 | 2.7E-74 |

| | | | | | | | |
|---|---|---|---|---|---|---|---|
| 0 | NDUFB8 | 0.334 | 0.243 | 1.649 | 1.397 | 0.239 | 8.3E-24 |
| 0 | NDUFB9 | 0.304 | 0.198 | 1.444 | 1.134 | 0.36 | 2.9E-40 |
| 0 | NDUFS7 | 0.126 | 0.084 | 0.43 | 0.422 | 0.331 | 2.2E-14 |
| 0 | NDUFS8 | 0.182 | 0.122 | 0.794 | 0.616 | 0.367 | 4.5E-22 |
| 0 | NDUFV1 | 0.16 | 0.112 | 0.707 | 0.497 | 0.242 | 1.9E-14 |
| 0 | NDUFV3 | 0.136 | 0.098 | 0.492 | 0.41 | 0.218 | 2.9E-11 |
| 0 | NEDD8 | 0.344 | 0.271 | 1.713 | 1.477 | 0.119 | 4E-14 |
| 0 | NFE2L2 | 0.331 | 0.244 | 1.789 | 1.604 | 0.147 | 3.5E-16 |
| 0 | NFKB1 | 0.423 | 0.341 | 3.718 | 2.901 | 0.348 | 2.1E-70 |
| 0 | NFKBIA | 0.9 | 0.646 | 21.872 | 9.473 | 1.207 | 0 |
| 0 | NFKBIB | 0.241 | 0.101 | 1.14 | 0.461 | 1.036 | 5E-114 |
| 0 | NFKBIZ | 0.382 | 0.164 | 2.743 | 1.19 | 1.204 | 6E-194 |
| 0 | NHP2 | 0.124 | 0.072 | 0.434 | 0.361 | 0.47 | 3.5E-24 |
| 0 | NKG7 | 0.993 | 0.874 | 40.394 | 19.424 | 1.046 | 0 |
| 0 | NKIRAS1 | 0.171 | 0.121 | 0.762 | 0.668 | 0.19 | 4.1E-14 |
| 0 | NME2 | 0.324 | 0.181 | 1.644 | 1.078 | 0.618 | 2.4E-77 |
| 0 | NME3 | 0.147 | 0.104 | 0.681 | 0.447 | 0.317 | 1E-18 |
| 0 | NMUR1 | 0.114 | 0.038 | 0.4 | 0.229 | 1.126 | 1.2E-74 |
| 0 | NOCT | 0.163 | 0.04 | 0.729 | 0.243 | 1.427 | 5E-146 |
| 0 | NOP48 | 0.327 | 0.232 | 1.689 | 1.442 | 0.227 | 6E-28 |
| 0 | NPC2 | 0.107 | 0.074 | 0.438 | 0.367 | 0.243 | 7.1E-11 |
| 0 | NPM1 | 0.493 | 0.431 | 3.947 | 3.112 | 0.343 | 1.2E-46 |
| 0 | NR4A2 | 0.374 | 0.103 | 2.146 | 0.787 | 1.444 | 0 |
| 0 | NRAS | 0.126 | 0.083 | 0.442 | 0.444 | 0.313 | 4.6E-16 |
| 0 | NSA2 | 0.317 | 0.241 | 1.6 | 1.493 | 0.1 | 2.6E-12 |
| 0 | NT4C | 0.162 | 0.121 | 0.709 | 0.631 | 0.168 | 3.9E-11 |
| 0 | NUCB2 | 0.244 | 0.17 | 1.192 | 1.012 | 0.236 | 1.1E-22 |
| 0 | NUCKS1 | 0.402 | 0.316 | 2.274 | 2.029 | 0.164 | 1.5E-17 |
| 0 | NUDT1 | 0.103 | 0.071 | 0.43 | 0.371 | 0.216 | 3.9E-11 |
| 0 | NUDT4 | 0.138 | 0.09 | 0.492 | 0.488 | 0.279 | 6.9E-18 |
| 0 | NUFIP2 | 0.347 | 0.242 | 1.884 | 1.486 | 0.24 | 2.8E-32 |
| 0 | NUP48 | 0.184 | 0.124 | 0.837 | 0.711 | 0.236 | 2.4E-19 |
| 0 | NUTF2 | 0.111 | 0.074 | 0.448 | 0.371 | 0.304 | 1.4E-13 |
| 0 | NXT1 | 0.221 | 0.083 | 1.042 | 0.481 | 1.116 | 5E-124 |
| 0 | ODC1 | 0.113 | 0.042 | 0.493 | 0.288 | 0.779 | 3.3E-43 |
| 0 | OGDH | 0.104 | 0.074 | 0.443 | 0.399 | 0.186 | 2.3E-09 |
| 0 | OLA1 | 0.272 | 0.204 | 1.281 | 1.193 | 0.103 | 7.2E-14 |
| 0 | OOEP | 0.411 | 0.28 | 2.204 | 1.72 | 0.348 | 9.5E-47 |
| 0 | ORAI1 | 0.129 | 0.074 | 0.487 | 0.394 | 0.474 | 1E-27 |
| 0 | OSBPL3 | 0.128 | 0.081 | 0.463 | 0.409 | 0.146 | 1.2E-18 |
| 0 | OSBPL4 | 0.143 | 0.084 | 0.696 | 0.404 | 0.466 | 1.9E-36 |
| 0 | OSER1 | 0.208 | 0.108 | 0.949 | 0.611 | 0.64 | 6.4E-60 |
| 0 | OST4 | 0.442 | 0.342 | 2.363 | 2.099 | 0.171 | 3.4E-20 |
| 0 | OSTF1 | 0.466 | 0.444 | 3.44 | 3.138 | 0.174 | 3.7E-24 |
| 0 | OSTM1 | 0.233 | 0.184 | 1.09 | 1.006 | 0.116 | 1.4E-09 |
| 0 | OXCT1 | 0.162 | 0.122 | 0.738 | 0.67 | 0.138 | 3.4E-10 |
| 0 | P2RY8 | 0.26 | 0.168 | 1.331 | 1.121 | 0.247 | 4.1E-34 |
| 0 | P4HA1 | 0.133 | 0.091 | 0.498 | 0.408 | 0.236 | 2.1E-14 |
| 0 | P4HB | 0.267 | 0.167 | 1.263 | 0.936 | 0.432 | 2.4E-42 |
| 0 | PA2G4 | 0.397 | 0.316 | 2.099 | 1.92 | 0.128 | 4.2E-14 |
| 0 | PABPC1 | 0.814 | 0.637 | 7.914 | 6.271 | 0.336 | 1.8E-84 |
| 0 | PABPC4 | 0.117 | 0.07 | 0.42 | 0.38 | 0.441 | 4.3E-22 |
| 0 | PAIP2 | 0.477 | 0.378 | 2.704 | 2.476 | 0.127 | 1.8E-16 |
| 0 | PAM | 0.344 | 0.242 | 1.844 | 1.619 | 0.188 | 9E-30 |

| | | | | | | | |
|---|---|---|---|---|---|---|---|
| 0 | PARK7 | 0.37 | 0.263 | 1.886 | 1.446 | 0.287 | 2.3E-32 |
| 0 | PATL2 | 0.193 | 0.122 | 0.926 | 0.696 | 0.412 | 1.3E-28 |
| 0 | PAXX | 0.387 | 0.284 | 2.044 | 1.784 | 0.196 | 1.1E-24 |
| 0 | PCBP1 | 0.616 | 0.406 | 4.143 | 3.873 | 0.101 | 1.4E-14 |
| 0 | PCBP2 | 0.82 | 0.639 | 7.238 | 4.727 | 0.338 | 7.7E-76 |
| 0 | PCED1B-AS1 | 0.464 | 0.308 | 2.686 | 1.973 | 0.444 | 1.5E-63 |
| 0 | PCIF1 | 0.141 | 0.087 | 0.68 | 0.483 | 0.493 | 6.6E-31 |
| 0 | PCNT | 0.18 | 0.111 | 0.842 | 0.678 | 0.311 | 2.5E-28 |
| 0 | PDCL3 | 0.124 | 0.076 | 0.437 | 0.406 | 0.404 | 3.9E-21 |
| 0 | PDGFD | 0.303 | 0.213 | 1.472 | 1.448 | 0.119 | 6.4E-24 |
| 0 | PDIA3 | 0.867 | 0.726 | 9.804 | 7.872 | 0.317 | 6.4E-66 |
| 0 | PDIA4 | 0.186 | 0.121 | 0.872 | 0.641 | 0.421 | 2.4E-24 |
| 0 | PDIA6 | 0.379 | 0.282 | 2.049 | 1.738 | 0.244 | 3.1E-24 |
| 0 | PDZD8 | 0.234 | 0.144 | 1.129 | 0.948 | 0.241 | 4.4E-29 |
| 0 | PECAM1 | 0.142 | 0.036 | 0.728 | 0.23 | 1.649 | 9E-142 |
| 0 | PELI2 | 0.194 | 0.121 | 0.911 | 0.826 | 0.142 | 4.1E-29 |
| 0 | PER1 | 0.142 | 0.049 | 0.63 | 0.328 | 0.944 | 4.2E-89 |
| 0 | PET100 | 0.284 | 0.219 | 1.328 | 1.222 | 0.12 | 7E-14 |
| 0 | PFDN1 | 0.191 | 0.139 | 0.838 | 0.734 | 0.19 | 4.4E-14 |
| 0 | PFDN2 | 0.203 | 0.143 | 0.928 | 0.768 | 0.274 | 1.3E-18 |
| 0 | PFDN4 | 0.484 | 0.464 | 3.671 | 3.322 | 0.144 | 1.2E-20 |
| 0 | PFDN4 | 0.102 | 0.076 | 0.437 | 0.378 | 0.21 | 8.5E-08 |
| 0 | PFN1 | 0.98 | 0.802 | 23.428 | 12.084 | 0.944 | 0 |
| 0 | PGAM1 | 0.2 | 0.136 | 0.9 | 0.709 | 0.343 | 6.2E-22 |
| 0 | PGK1 | 0.366 | 0.267 | 1.846 | 1.624 | 0.191 | 8.4E-26 |
| 0 | PGLS | 0.182 | 0.134 | 0.808 | 0.724 | 0.146 | 4.5E-12 |
| 0 | PGRMC2 | 0.119 | 0.076 | 0.428 | 0.418 | 0.339 | 3.5E-17 |
| 0 | PHB2 | 0.162 | 0.101 | 0.702 | 0.424 | 0.419 | 4.9E-26 |
| 0 | PHF20 | 0.377 | 0.226 | 2.003 | 1.478 | 0.438 | 3.9E-71 |
| 0 | PHPT1 | 0.121 | 0.08 | 0.431 | 0.416 | 0.343 | 7E-16 |
| 0 | PIGT | 0.109 | 0.064 | 0.467 | 0.34 | 0.416 | 4.4E-20 |
| 0 | PIN1 | 0.219 | 0.141 | 0.982 | 0.748 | 0.373 | 1.2E-30 |
| 0 | PLAAT3 | 0.181 | 0.133 | 0.818 | 0.731 | 0.163 | 6.5E-13 |
| 0 | PLAC8 | 0.499 | 0.281 | 3.071 | 1.834 | 0.743 | 1E-140 |
| 0 | PLCL2 | 0.37 | 0.276 | 1.976 | 1.826 | 0.114 | 4.9E-21 |
| 0 | PLEK | 0.791 | 0.467 | 7.684 | 4.244 | 0.441 | 2E-144 |
| 0 | PLEKHA1 | 0.292 | 0.219 | 1.444 | 1.31 | 0.141 | 1.7E-16 |
| 0 | PLEKHA7 | 0.122 | 0.091 | 0.409 | 0.46 | 0.146 | 1.5E-08 |
| 0 | PLEKHB2 | 0.132 | 0.092 | 0.464 | 0.409 | 0.148 | 3.4E-12 |
| 0 | PLEKHM1 | 0.187 | 0.131 | 0.847 | 0.744 | 0.167 | 3.5E-17 |
| 0 | PMAIP1 | 0.312 | 0.102 | 1.973 | 0.686 | 1.424 | 4E-236 |
| 0 | PNKD | 0.174 | 0.112 | 0.779 | 0.602 | 0.372 | 2E-23 |
| 0 | PNN | 0.408 | 0.279 | 2.244 | 1.796 | 0.328 | 1.5E-44 |
| 0 | PNPLA8 | 0.246 | 0.144 | 1.194 | 0.941 | 0.344 | 4.7E-36 |
| 0 | PNRC1 | 0.934 | 0.779 | 14.049 | 10.099 | 0.477 | 1E-172 |
| 0 | POLB | 0.14 | 0.099 | 0.673 | 0.442 | 0.284 | 1.5E-18 |
| 0 | POLD4 | 0.137 | 0.106 | 0.477 | 0.438 | 0.102 | 1.5E-07 |
| 0 | POLE3 | 0.249 | 0.198 | 1.238 | 1.122 | 0.142 | 3.5E-13 |
| 0 | POLR1D | 0.207 | 0.143 | 0.928 | 0.772 | 0.267 | 2.7E-20 |
| 0 | POLR2A | 0.233 | 0.118 | 1.114 | 0.842 | 0.404 | 2.5E-67 |
| 0 | POLR2E | 0.164 | 0.114 | 0.709 | 0.487 | 0.273 | 1.3E-16 |
| 0 | POLR2F | 0.162 | 0.126 | 0.727 | 0.664 | 0.129 | 3E-08 |
| 0 | POLR2G | 0.212 | 0.14 | 0.961 | 0.744 | 0.369 | 7.4E-27 |
| 0 | POLR2J | 0.144 | 0.113 | 0.664 | 0.482 | 0.194 | 1E-11 |

| | | | | | | | |
|---|---|---|---|---|---|---|---|
| 0 | POLR2K | 0.191 | 0.112 | 0.922 | 0.603 | 0.612 | 6E-40 |
| 0 | POLR2L | 0.447 | 0.291 | 2.487 | 1.734 | 0.419 | 3.2E-76 |
| 0 | POLR3GL | 0.308 | 0.217 | 1.427 | 1.214 | 0.33 | 7.3E-29 |
| 0 | POP4 | 0.104 | 0.077 | 0.449 | 0.396 | 0.181 | 2.9E-08 |
| 0 | PPDPF | 0.444 | 0.338 | 2.641 | 2.223 | 0.244 | 1.1E-30 |
| 0 | PPIA | 0.746 | 0.418 | 6.178 | 4.078 | 0.499 | 4E-149 |
| 0 | PPIB | 0.486 | 0.397 | 3.88 | 2.736 | 0.404 | 2.5E-89 |
| 0 | PPIH | 0.122 | 0.086 | 0.417 | 0.439 | 0.236 | 8.3E-12 |
| 0 | PPIL2 | 0.126 | 0.087 | 0.446 | 0.492 | 0.174 | 1.2E-12 |
| 0 | PPM1G | 0.303 | 0.221 | 1.467 | 1.239 | 0.243 | 1.7E-22 |
| 0 | PPM1L | 0.378 | 0.177 | 2.116 | 1.24 | 0.76 | 4E-144 |
| 0 | PPP1CA | 0.466 | 0.323 | 2.634 | 2 | 0.397 | 1.8E-43 |
| 0 | PPP1R10 | 0.431 | 0.224 | 2.644 | 1.419 | 0.898 | 4E-147 |
| 0 | PPP1R11 | 0.188 | 0.136 | 0.849 | 0.724 | 0.247 | 4.2E-14 |
| 0 | PPP1R14A | 0.486 | 0.183 | 4.826 | 1.744 | 1.732 | 0 |
| 0 | PPP1R14B | 0.21 | 0.099 | 0.981 | 0.482 | 0.743 | 4.1E-76 |
| 0 | PPP1R18 | 0.317 | 0.222 | 1.613 | 1.284 | 0.329 | 4.5E-31 |
| 0 | PPP1R7 | 0.2 | 0.141 | 0.904 | 0.789 | 0.196 | 2.8E-12 |
| 0 | PPP2R1A | 0.16 | 0.108 | 0.701 | 0.487 | 0.246 | 4.1E-18 |
| 0 | PPP2R2D | 0.144 | 0.109 | 0.71 | 0.608 | 0.224 | 3.5E-14 |
| 0 | PQBP1 | 0.106 | 0.074 | 0.444 | 0.387 | 0.201 | 8.5E-10 |
| 0 | PRCP | 0.137 | 0.089 | 0.607 | 0.482 | 0.334 | 9.6E-19 |
| 0 | PRDM1 | 0.474 | 0.277 | 2.999 | 1.918 | 0.644 | 7E-114 |
| 0 | PRDX4 | 0.309 | 0.229 | 1.436 | 1.306 | 0.234 | 1E-19 |
| 0 | PRELID1 | 0.248 | 0.193 | 1.246 | 1.047 | 0.237 | 6.6E-17 |
| 0 | PRF1 | 0.704 | 0.423 | 4.986 | 4.447 | 0.429 | 3.3E-84 |
| 0 | PRKAG1 | 0.117 | 0.084 | 0.412 | 0.447 | 0.164 | 2.4E-10 |
| 0 | PRKAR2A | 0.249 | 0.184 | 1.2 | 1.092 | 0.137 | 9.2E-16 |
| 0 | PRMT1 | 0.136 | 0.102 | 0.49 | 0.42 | 0.182 | 5E-09 |
| 0 | PRMT2 | 0.499 | 0.404 | 3.077 | 2.832 | 0.12 | 9E-16 |
| 0 | PRNP | 0.133 | 0.087 | 0.604 | 0.48 | 0.332 | 9.4E-18 |
| 0 | PRPF4B | 0.414 | 0.403 | 3.032 | 2.74 | 0.141 | 2.7E-22 |
| 0 | PRR13 | 0.694 | 0.443 | 4.887 | 4.198 | 0.219 | 3.2E-34 |
| 0 | PRR4 | 0.129 | 0.077 | 0.444 | 0.441 | 0.298 | 4.7E-23 |
| 0 | PRSS23 | 0.363 | 0.124 | 2.004 | 0.864 | 1.214 | 4E-248 |
| 0 | PRXL2C | 0.137 | 0.071 | 0.629 | 0.38 | 0.729 | 6.7E-40 |
| 0 | PSAP | 0.642 | 0.484 | 4.604 | 3.441 | 0.379 | 4.8E-64 |
| 0 | PSMB6 | 0.244 | 0.177 | 1.164 | 0.941 | 0.308 | 9.1E-24 |
| 0 | PSMB7 | 0.109 | 0.081 | 0.462 | 0.429 | 0.107 | 4.3E-08 |
| 0 | PSMC1 | 0.22 | 0.166 | 0.993 | 0.906 | 0.132 | 1.4E-12 |
| 0 | PSMC4 | 0.166 | 0.13 | 0.722 | 0.671 | 0.106 | 2.6E-08 |
| 0 | PSMC4 | 0.143 | 0.113 | 0.676 | 0.499 | 0.174 | 7.8E-11 |
| 0 | PSMD14 | 0.149 | 0.103 | 0.67 | 0.446 | 0.268 | 3.2E-14 |
| 0 | PSMD8 | 0.277 | 0.194 | 1.29 | 1.067 | 0.274 | 6E-24 |
| 0 | PSMD9 | 0.123 | 0.089 | 0.414 | 0.461 | 0.149 | 9.1E-10 |
| 0 | PSTPIP2 | 0.138 | 0.082 | 0.612 | 0.473 | 0.372 | 2.4E-26 |
| 0 | PTBP1 | 0.119 | 0.086 | 0.413 | 0.448 | 0.163 | 6.2E-10 |
| 0 | PTCH2 | 0.188 | 0.04 | 1.363 | 0.334 | 2.023 | 9E-213 |
| 0 | PTGDR | 0.277 | 0.133 | 1.371 | 0.786 | 0.802 | 8E-99 |
| 0 | PTGDS | 0.366 | 0.106 | 6.827 | 1.293 | 2.401 | 0 |
| 0 | PTGER2 | 0.178 | 0.108 | 0.81 | 0.497 | 0.442 | 7.6E-31 |
| 0 | PTMA | 0.993 | 0.937 | 28.977 | 23.847 | 0.281 | 2E-111 |
| 0 | PTP4A2 | 0.162 | 0.103 | 0.726 | 0.447 | 0.409 | 1E-23 |
| 0 | PTPN12 | 0.417 | 0.348 | 3.207 | 2.442 | 0.329 | 1.9E-44 |

| | | | | | | | |
|---|---|---|---|---|---|---|---|
| 0 | PTPN18 | 0.146 | 0.084 | 0.66 | 0.448 | 0.447 | 3.8E-31 |
| 0 | PTPN2 | 0.419 | 0.32 | 2.346 | 2.102 | 0.148 | 3.9E-21 |
| 0 | PTPN6 | 0.137 | 0.047 | 0.62 | 0.332 | 0.902 | 3.9E-63 |
| 0 | PTS | 0.194 | 0.116 | 0.878 | 0.667 | 0.396 | 4.1E-34 |
| 0 | PTTG1IP | 0.174 | 0.11 | 0.784 | 0.494 | 0.4 | 2.9E-27 |
| 0 | PUF60 | 0.118 | 0.087 | 0.492 | 0.441 | 0.149 | 4.8E-09 |
| 0 | PXN | 0.14 | 0.063 | 0.68 | 0.414 | 0.711 | 3.6E-67 |
| 0 | PYROXD1 | 0.121 | 0.097 | 0.442 | 0.409 | 0.119 | 2.5E-05 |
| 0 | RAB11A | 0.316 | 0.181 | 1.494 | 1.049 | 0.489 | 4.1E-71 |
| 0 | RAB11B | 0.121 | 0.092 | 0.42 | 0.483 | 0.104 | 1.7E-07 |
| 0 | RAB1B | 0.197 | 0.131 | 0.891 | 0.713 | 0.321 | 1.2E-23 |
| 0 | RAB29 | 0.301 | 0.174 | 1.486 | 1.014 | 0.441 | 3.9E-64 |
| 0 | RAB2A | 0.46 | 0.343 | 2.632 | 2.44 | 0.109 | 3.2E-22 |
| 0 | RAB37 | 0.141 | 0.064 | 0.673 | 0.361 | 0.899 | 1.3E-67 |
| 0 | RAB4A | 0.237 | 0.173 | 1.107 | 0.996 | 0.142 | 6.9E-17 |
| 0 | RAB4A | 0.147 | 0.111 | 0.706 | 0.61 | 0.21 | 3.3E-14 |
| 0 | RAB4B | 0.127 | 0.092 | 0.449 | 0.493 | 0.144 | 1.4E-10 |
| 0 | RAB4C | 0.289 | 0.203 | 1.369 | 1.139 | 0.264 | 7.8E-26 |
| 0 | RAB4IF | 0.339 | 0.169 | 1.809 | 0.993 | 0.864 | 2E-117 |
| 0 | RAB7A | 0.29 | 0.217 | 1.401 | 1.271 | 0.141 | 6.3E-17 |
| 0 | RAB8A | 0.297 | 0.21 | 1.422 | 1.144 | 0.3 | 1.4E-26 |
| 0 | RAB8B | 0.649 | 0.421 | 4.744 | 4.238 | 0.166 | 7.1E-30 |
| 0 | RAB9A | 0.143 | 0.104 | 0.624 | 0.439 | 0.211 | 1.8E-10 |
| 0 | RABGEF1 | 0.243 | 0.18 | 1.194 | 1.09 | 0.132 | 1.9E-14 |
| 0 | RABL6 | 0.141 | 0.11 | 0.63 | 0.486 | 0.106 | 2.3E-07 |
| 0 | RAC1 | 0.312 | 0.234 | 1.418 | 1.308 | 0.214 | 3.7E-19 |
| 0 | RAC2 | 0.864 | 0.601 | 9.82 | 4.497 | 0.837 | 0 |
| 0 | RACK1 | 0.744 | 0.473 | 6.103 | 4.796 | 0.348 | 7.2E-72 |
| 0 | RAD23A | 0.204 | 0.142 | 0.936 | 0.84 | 0.147 | 2.2E-13 |
| 0 | RALY | 0.331 | 0.242 | 1.671 | 1.443 | 0.212 | 4.7E-24 |
| 0 | RANBP1 | 0.109 | 0.082 | 0.464 | 0.414 | 0.163 | 1.1E-07 |
| 0 | RAP1B | 0.834 | 0.687 | 8.074 | 6.76 | 0.246 | 3.4E-43 |
| 0 | RAP1GAP2 | 0.376 | 0.138 | 2.179 | 1.346 | 0.684 | 2E-217 |
| 0 | RAP2B | 0.107 | 0.046 | 0.449 | 0.294 | 0.643 | 3.3E-31 |
| 0 | RAP2C | 0.171 | 0.12 | 0.771 | 0.641 | 0.266 | 3.6E-16 |
| 0 | RARA | 0.131 | 0.047 | 0.47 | 0.321 | 0.83 | 1.4E-44 |
| 0 | RASA2 | 0.648 | 0.427 | 4.747 | 4.314 | 0.138 | 4E-22 |
| 0 | RASA3 | 0.406 | 0.343 | 3.127 | 2.69 | 0.217 | 4E-47 |
| 0 | RASGEF1B | 0.244 | 0.146 | 1.308 | 0.931 | 0.491 | 8.9E-44 |
| 0 | RASGRP2 | 0.144 | 0.042 | 0.646 | 0.319 | 1.021 | 7.4E-87 |
| 0 | RASSF1 | 0.238 | 0.11 | 1.123 | 0.643 | 0.781 | 4.1E-89 |
| 0 | RASSF4 | 0.18 | 0.069 | 0.846 | 0.423 | 1.002 | 1.4E-96 |
| 0 | RBBP7 | 0.312 | 0.221 | 1.448 | 1.31 | 0.241 | 4.3E-27 |
| 0 | RBFOX2 | 0.203 | 0.134 | 0.93 | 0.731 | 0.346 | 8.3E-24 |
| 0 | RBM18 | 0.143 | 0.114 | 0.668 | 0.608 | 0.137 | 9E-10 |
| 0 | RBM3 | 0.324 | 0.146 | 1.601 | 0.847 | 0.902 | 3E-137 |
| 0 | RBM39 | 0.841 | 0.649 | 9.014 | 6.349 | 0.406 | 9E-144 |
| 0 | RBM7 | 0.114 | 0.077 | 0.479 | 0.406 | 0.24 | 1.1E-12 |
| 0 | RBM8A | 0.492 | 0.394 | 2.843 | 2.628 | 0.119 | 1.8E-14 |
| 0 | RBMX | 0.232 | 0.169 | 1.084 | 0.962 | 0.173 | 9.3E-17 |
| 0 | RCBTB2 | 0.128 | 0.084 | 0.476 | 0.482 | 0.247 | 1E-15 |
| 0 | REL | 0.444 | 0.404 | 3.438 | 2.962 | 0.214 | 1.3E-37 |
| 0 | REX1BD | 0.261 | 0.204 | 1.241 | 1.146 | 0.113 | 1.3E-11 |
| 0 | RGCC | 0.247 | 0.216 | 1.601 | 1.481 | 0.113 | 4.6E-07 |

| | | | | | | | |
|---|---|---|---|---|---|---|---|
| 0 | RGS1 | 0.113 | 0.06 | 0.498 | 0.336 | 0.469 | 1E-30 |
| 0 | RGS19 | 0.149 | 0.043 | 0.671 | 0.283 | 1.246 | 2.2E-93 |
| 0 | RGS2 | 0.18 | 0.077 | 0.889 | 0.429 | 1.042 | 3E-82 |
| 0 | RGS3 | 0.149 | 0.096 | 0.649 | 0.437 | 0.294 | 3.4E-20 |
| 0 | RHOA | 0.727 | 0.428 | 4.743 | 4.071 | 0.496 | 4E-109 |
| 0 | RHOB | 0.103 | 0.028 | 0.489 | 0.162 | 1.488 | 4.3E-90 |
| 0 | RHOC | 0.441 | 0.19 | 2.401 | 1.144 | 1.128 | 4E-232 |
| 0 | RHOF | 0.21 | 0.131 | 0.947 | 0.742 | 0.341 | 1.9E-32 |
| 0 | RHOG | 0.26 | 0.17 | 1.224 | 0.928 | 0.401 | 1.4E-34 |
| 0 | RHOH | 0.284 | 0.209 | 1.482 | 1.314 | 0.174 | 3.3E-20 |
| 0 | RHOQ | 0.101 | 0.064 | 0.433 | 0.366 | 0.242 | 3.4E-14 |
| 0 | RINL | 0.121 | 0.089 | 0.43 | 0.472 | 0.166 | 4.3E-09 |
| 0 | RIOK3 | 0.388 | 0.294 | 2.088 | 1.84 | 0.174 | 2.7E-20 |
| 0 | RIPK2 | 0.148 | 0.082 | 0.664 | 0.467 | 0.406 | 8.4E-34 |
| 0 | RIT1 | 0.106 | 0.08 | 0.447 | 0.412 | 0.147 | 4.9E-07 |
| 0 | RNASEH2C | 0.144 | 0.12 | 0.67 | 0.618 | 0.114 | 6.2E-08 |
| 0 | RNASET2 | 0.142 | 0.1 | 0.632 | 0.447 | 0.209 | 4.9E-13 |
| 0 | RNF11 | 0.21 | 0.148 | 0.941 | 0.876 | 0.118 | 4.3E-12 |
| 0 | RNF124 | 0.478 | 0.268 | 2.823 | 1.832 | 0.624 | 1E-126 |
| 0 | RNF126 | 0.142 | 0.072 | 0.632 | 0.379 | 0.74 | 1.3E-42 |
| 0 | RNF130 | 0.14 | 0.096 | 0.667 | 0.444 | 0.267 | 3.6E-14 |
| 0 | RNF138 | 0.238 | 0.144 | 1.136 | 0.883 | 0.362 | 4.7E-41 |
| 0 | RNF166 | 0.24 | 0.18 | 1.189 | 1.049 | 0.18 | 6.7E-19 |
| 0 | RNF167 | 0.142 | 0.098 | 0.661 | 0.424 | 0.334 | 1.1E-21 |
| 0 | RNF7 | 0.334 | 0.248 | 1.644 | 1.418 | 0.214 | 2.6E-21 |
| 0 | RNH1 | 0.128 | 0.087 | 0.467 | 0.447 | 0.312 | 3.9E-14 |
| 0 | RNMT | 0.274 | 0.191 | 1.348 | 1.164 | 0.221 | 8.7E-26 |
| 0 | RNPS1 | 0.362 | 0.264 | 1.822 | 1.467 | 0.217 | 2E-24 |
| 0 | ROMO1 | 0.284 | 0.206 | 1.321 | 1.137 | 0.217 | 8.4E-21 |
| 0 | RPF1 | 0.194 | 0.127 | 0.887 | 0.72 | 0.302 | 1.5E-24 |
| 0 | RPIA | 0.111 | 0.076 | 0.477 | 0.409 | 0.222 | 3.7E-12 |
| 0 | RPL10 | 0.998 | 0.949 | 46.477 | 39.163 | 0.247 | 5E-113 |
| 0 | RPL10A | 0.699 | 0.48 | 4.444 | 3.887 | 0.489 | 4E-112 |
| 0 | RPL11 | 0.986 | 0.919 | 23.884 | 20.786 | 0.2 | 2.3E-61 |
| 0 | RPL12 | 0.972 | 0.876 | 19.479 | 16.084 | 0.276 | 1.2E-92 |
| 0 | RPL13 | 0.892 | 0.744 | 11.041 | 9.036 | 0.289 | 4.7E-68 |
| 0 | RPL13A | 0.842 | 0.731 | 9.23 | 8.392 | 0.137 | 7.7E-23 |
| 0 | RPL14 | 0.934 | 0.819 | 14.428 | 12.203 | 0.242 | 1.4E-49 |
| 0 | RPL14 | 0.807 | 0.612 | 7.619 | 4.494 | 0.444 | 7E-114 |
| 0 | RPL17 | 0.8 | 0.627 | 7.364 | 4.83 | 0.337 | 8.9E-69 |
| 0 | RPL18 | 0.974 | 0.884 | 19.347 | 17.142 | 0.174 | 2.5E-44 |
| 0 | RPL18A | 0.872 | 0.748 | 10.044 | 8.9 | 0.176 | 3.8E-29 |
| 0 | RPL19 | 0.976 | 0.891 | 19.866 | 17.491 | 0.184 | 2.2E-47 |
| 0 | RPL21 | 0.849 | 0.722 | 9.173 | 8.033 | 0.192 | 1.8E-34 |
| 0 | RPL22 | 0.883 | 0.716 | 9.799 | 7.867 | 0.317 | 9.7E-84 |
| 0 | RPL22L1 | 0.198 | 0.098 | 0.894 | 0.429 | 0.748 | 8.4E-64 |
| 0 | RPL23 | 0.432 | 0.402 | 3.192 | 2.722 | 0.23 | 1.3E-31 |
| 0 | RPL23A | 0.684 | 0.463 | 4.171 | 4.786 | 0.111 | 4.2E-16 |
| 0 | RPL24 | 0.809 | 0.643 | 7.633 | 6.442 | 0.244 | 4.5E-47 |
| 0 | RPL26 | 0.799 | 0.662 | 7.424 | 6.621 | 0.184 | 1.1E-28 |
| 0 | RPL27 | 0.496 | 0.432 | 3.944 | 2.993 | 0.402 | 4.6E-64 |
| 0 | RPL27A | 0.418 | 0.38 | 3.18 | 2.438 | 0.324 | 1.4E-42 |
| 0 | RPL28 | 0.983 | 0.9 | 21.298 | 19.393 | 0.134 | 4.2E-31 |
| 0 | RPL29 | 0.893 | 0.741 | 10.718 | 8.774 | 0.289 | 3E-69 |

| | | | | | | | |
|---|---|---|---|---|---|---|---|
| 0 | RPL3 | 0.663 | 0.494 | 4.747 | 3.872 | 0.297 | 2.3E-41 |
| 0 | RPL30 | 0.989 | 0.914 | 24.21 | 21.317 | 0.184 | 3.3E-49 |
| 0 | RPL31 | 0.749 | 0.619 | 6.474 | 4.448 | 0.223 | 2.6E-34 |
| 0 | RPL32 | 0.949 | 0.843 | 16.294 | 14.037 | 0.214 | 3.8E-47 |
| 0 | RPL34 | 0.973 | 0.881 | 19.6 | 17.331 | 0.177 | 7.6E-43 |
| 0 | RPL34 | 0.803 | 0.626 | 7.318 | 4.884 | 0.314 | 1.3E-68 |
| 0 | RPL36 | 0.806 | 0.674 | 7.464 | 6.727 | 0.14 | 7.2E-24 |
| 0 | RPL37 | 0.942 | 0.831 | 16.141 | 12.79 | 0.336 | 3E-103 |
| 0 | RPL37A | 0.781 | 0.646 | 7.131 | 6.384 | 0.149 | 7.1E-23 |
| 0 | RPL38 | 0.778 | 0.614 | 6.977 | 4.678 | 0.297 | 2.1E-47 |
| 0 | RPL39 | 0.937 | 0.789 | 13.099 | 10.984 | 0.244 | 1.9E-68 |
| 0 | RPL4 | 0.948 | 0.792 | 14.786 | 11.033 | 0.422 | 1E-161 |
| 0 | RPL4 | 0.448 | 0.318 | 2.404 | 2.023 | 0.308 | 2.2E-41 |
| 0 | RPL41 | 0.992 | 0.924 | 33.267 | 27.476 | 0.276 | 1E-112 |
| 0 | RPL6 | 0.948 | 0.829 | 14.462 | 12.262 | 0.334 | 7E-114 |
| 0 | RPL7 | 0.716 | 0.448 | 4.482 | 4.434 | 0.306 | 9.2E-44 |
| 0 | RPL7A | 0.97 | 0.88 | 18.328 | 14.8 | 0.214 | 8.9E-60 |
| 0 | RPL8 | 0.916 | 0.798 | 11.824 | 10.722 | 0.141 | 1.8E-28 |
| 0 | RPL9 | 0.694 | 0.408 | 4.141 | 3.998 | 0.366 | 6.9E-71 |
| 0 | RPLP2 | 0.991 | 0.931 | 27.147 | 24.683 | 0.138 | 1.1E-34 |
| 0 | RPN1 | 0.242 | 0.197 | 1.186 | 1.104 | 0.103 | 4.9E-11 |
| 0 | RPN2 | 0.244 | 0.167 | 1.119 | 0.921 | 0.281 | 1.4E-24 |
| 0 | RPS11 | 0.693 | 0.442 | 4.026 | 4.303 | 0.224 | 1.8E-34 |
| 0 | RPS13 | 0.982 | 0.894 | 20.391 | 18.131 | 0.17 | 4E-47 |
| 0 | RPS14 | 0.96 | 0.867 | 17.733 | 14.843 | 0.162 | 2.4E-37 |
| 0 | RPS14 | 0.872 | 0.727 | 9.617 | 8.09 | 0.249 | 1.8E-42 |
| 0 | RPS14A | 0.962 | 0.842 | 17.421 | 13.722 | 0.344 | 1E-128 |
| 0 | RPS16 | 0.731 | 0.444 | 4.869 | 4.49 | 0.344 | 2.4E-66 |
| 0 | RPS17 | 0.462 | 0.328 | 2.463 | 2.036 | 0.332 | 6.2E-44 |
| 0 | RPS18 | 0.787 | 0.621 | 7.083 | 4.928 | 0.247 | 6.8E-41 |
| 0 | RPS19BP1 | 0.19 | 0.139 | 0.848 | 0.736 | 0.203 | 4.3E-14 |
| 0 | RPS2 | 0.668 | 0.464 | 4.909 | 3.667 | 0.421 | 6E-88 |
| 0 | RPS20 | 0.406 | 0.314 | 2.166 | 1.939 | 0.16 | 2.3E-18 |
| 0 | RPS21 | 0.93 | 0.79 | 13.034 | 11.094 | 0.232 | 3.8E-48 |
| 0 | RPS23 | 0.903 | 0.784 | 11.719 | 10.444 | 0.141 | 1.2E-24 |
| 0 | RPS24 | 0.996 | 0.941 | 32.609 | 27.243 | 0.249 | 2E-121 |
| 0 | RPS24 | 0.646 | 0.494 | 4.434 | 3.804 | 0.243 | 2.9E-39 |
| 0 | RPS27 | 0.978 | 0.881 | 23.314 | 19.194 | 0.281 | 3.5E-98 |
| 0 | RPS28 | 0.969 | 0.867 | 18.682 | 14.971 | 0.226 | 3.4E-63 |
| 0 | RPS29 | 0.889 | 0.737 | 11.446 | 9.249 | 0.318 | 9.1E-80 |
| 0 | RPS3A | 0.943 | 0.811 | 14.699 | 11.642 | 0.336 | 2E-104 |
| 0 | RPS4 | 0.841 | 0.687 | 8.636 | 7.194 | 0.263 | 3.4E-46 |
| 0 | RPS4Y1 | 0.319 | 0.192 | 1.798 | 1.218 | 0.462 | 6.1E-61 |
| 0 | RPS6 | 0.886 | 0.734 | 10.964 | 8.747 | 0.326 | 2E-84 |
| 0 | RPS6KA1 | 0.106 | 0.063 | 0.446 | 0.344 | 0.371 | 1.3E-19 |
| 0 | RPS7 | 0.973 | 0.872 | 19.279 | 14.844 | 0.282 | 3.4E-96 |
| 0 | RPS8 | 0.994 | 0.94 | 31.299 | 26.694 | 0.23 | 2.3E-94 |
| 0 | RPS9 | 0.492 | 0.334 | 2.83 | 2.187 | 0.371 | 3.9E-49 |
| 0 | RPSA | 0.736 | 0.439 | 6.146 | 4.492 | 0.442 | 5E-101 |
| 0 | RRBP1 | 0.274 | 0.187 | 1.367 | 1.148 | 0.242 | 1.8E-28 |
| 0 | RSL1D1 | 0.21 | 0.142 | 0.977 | 0.838 | 0.222 | 2.3E-16 |
| 0 | RSL24D1 | 0.296 | 0.188 | 1.434 | 1.079 | 0.41 | 3.5E-43 |
| 0 | RSRC2 | 0.313 | 0.239 | 1.492 | 1.429 | 0.147 | 1.3E-16 |
| 0 | RSRP1 | 0.674 | 0.43 | 4.304 | 4.294 | 0.304 | 8.5E-44 |

| | | | | | | |
|---|---|---|---|---|---|---|
| 0 | RSU1 | 0.267 | 0.199 | 1.274 | 1.13 | 0.174 | 3.8E-17 |
| 0 | RTN4 | 0.347 | 0.247 | 1.816 | 1.479 | 0.202 | 3.2E-27 |
| 0 | RUNX3 | 0.394 | 0.264 | 2.169 | 1.799 | 0.27 | 2E-44 |
| 0 | RWDD1 | 0.334 | 0.228 | 1.694 | 1.304 | 0.376 | 3.8E-37 |
| 0 | S100A10 | 0.601 | 0.414 | 4.048 | 3.076 | 0.399 | 4E-71 |
| 0 | S100A4 | 0.923 | 0.466 | 19.947 | 9.031 | 1.143 | 0 |
| 0 | S100A6 | 0.701 | 0.464 | 6.234 | 4.287 | 0.441 | 1E-129 |
| 0 | S1PR4 | 0.444 | 0.329 | 3.414 | 2.164 | 0.699 | 4E-141 |
| 0 | SAE1 | 0.103 | 0.049 | 0.447 | 0.33 | 0.469 | 6.1E-22 |
| 0 | SAMD8 | 0.124 | 0.084 | 0.439 | 0.4 | 0.107 | 1.6E-13 |
| 0 | SAMHD1 | 0.464 | 0.34 | 2.734 | 2.406 | 0.126 | 1.7E-23 |
| 0 | SAP18 | 0.478 | 0.348 | 2.609 | 2.142 | 0.284 | 6E-37 |
| 0 | SAR1B | 0.14 | 0.098 | 0.673 | 0.408 | 0.406 | 1.2E-20 |
| 0 | SARS | 0.204 | 0.144 | 0.942 | 0.846 | 0.139 | 6.8E-12 |
| 0 | SBDS | 0.389 | 0.187 | 2.131 | 1.134 | 0.91 | 1E-140 |
| 0 | SCAND1 | 0.311 | 0.217 | 1.422 | 1.203 | 0.34 | 2.9E-30 |
| 0 | SCP2 | 0.348 | 0.238 | 1.809 | 1.401 | 0.369 | 1.5E-39 |
| 0 | SDCBP | 0.36 | 0.278 | 1.934 | 1.774 | 0.124 | 1.7E-16 |
| 0 | SDF2 | 0.103 | 0.07 | 0.434 | 0.349 | 0.319 | 6.8E-12 |
| 0 | SDF2L1 | 0.149 | 0.074 | 0.672 | 0.387 | 0.794 | 2.4E-47 |
| 0 | SDF4 | 0.261 | 0.179 | 1.234 | 0.992 | 0.314 | 1.4E-27 |
| 0 | SEC11C | 0.207 | 0.141 | 0.963 | 0.808 | 0.243 | 4E-16 |
| 0 | SEC13 | 0.124 | 0.094 | 0.432 | 0.491 | 0.116 | 1E-07 |
| 0 | SEC14L1 | 0.263 | 0.177 | 1.242 | 1.004 | 0.304 | 1.1E-29 |
| 0 | SEC61B | 0.444 | 0.343 | 2.444 | 2.221 | 0.144 | 2.5E-16 |
| 0 | SEC61G | 0.366 | 0.267 | 1.816 | 1.442 | 0.236 | 2.8E-24 |
| 0 | SELENOK | 0.431 | 0.269 | 2.37 | 1.668 | 0.407 | 2.1E-76 |
| 0 | SELENOT | 0.277 | 0.204 | 1.332 | 1.164 | 0.194 | 7.9E-19 |
| 0 | SELENOW | 0.374 | 0.204 | 1.903 | 1.199 | 0.667 | 2E-99 |
| 0 | SELPLG | 0.336 | 0.201 | 1.736 | 1.194 | 0.438 | 6.9E-64 |
| 0 | SEMA4D | 0.263 | 0.176 | 1.262 | 1.049 | 0.243 | 4.7E-30 |
| 0 | SEPTIN1 | 0.247 | 0.187 | 1.227 | 1.071 | 0.197 | 2.6E-18 |
| 0 | SEPTIN11 | 0.146 | 0.119 | 0.706 | 0.649 | 0.121 | 2.8E-09 |
| 0 | SERF2 | 0.893 | 0.737 | 10.28 | 8.042 | 0.342 | 1.9E-96 |
| 0 | SERP1 | 0.406 | 0.304 | 2.146 | 1.828 | 0.238 | 2.8E-26 |
| 0 | SERPINB1 | 0.476 | 0.329 | 2.878 | 2.162 | 0.413 | 4.6E-46 |
| 0 | SERPINB6 | 0.168 | 0.113 | 0.744 | 0.604 | 0.321 | 1.3E-19 |
| 0 | SERTAD1 | 0.314 | 0.106 | 1.637 | 0.637 | 1.361 | 1E-220 |
| 0 | SERTAD2 | 0.139 | 0.084 | 0.611 | 0.477 | 0.349 | 3.9E-24 |
| 0 | SERTAD3 | 0.119 | 0.042 | 0.436 | 0.283 | 0.918 | 1.8E-40 |
| 0 | SET | 0.712 | 0.449 | 4.449 | 4.468 | 0.244 | 1.2E-40 |
| 0 | SETD3 | 0.341 | 0.237 | 1.729 | 1.416 | 0.288 | 4.9E-32 |
| 0 | SF3B4 | 0.187 | 0.126 | 0.84 | 0.646 | 0.378 | 6.4E-22 |
| 0 | SF3B6 | 0.2 | 0.143 | 0.876 | 0.764 | 0.194 | 4.1E-16 |
| 0 | SFPQ | 0.349 | 0.267 | 1.869 | 1.646 | 0.183 | 7.2E-23 |
| 0 | SFT2D1 | 0.246 | 0.141 | 1.216 | 0.804 | 0.497 | 2.2E-44 |
| 0 | SFT2D2 | 0.143 | 0.102 | 0.644 | 0.441 | 0.247 | 9.9E-13 |
| 0 | SH3BGRL3 | 0.932 | 0.78 | 14.406 | 11.769 | 0.398 | 4E-123 |
| 0 | SH3BP1 | 0.146 | 0.104 | 0.698 | 0.464 | 0.304 | 4.8E-18 |
| 0 | SH3BP4 | 0.184 | 0.09 | 0.836 | 0.407 | 0.722 | 1.7E-61 |
| 0 | SH3D19 | 0.177 | 0.114 | 0.79 | 0.632 | 0.322 | 4E-24 |
| 0 | SH3GLB1 | 0.479 | 0.389 | 2.771 | 2.484 | 0.101 | 2.7E-14 |
| 0 | SHARPIN | 0.123 | 0.094 | 0.44 | 0.401 | 0.107 | 4.3E-07 |
| 0 | SHKBP1 | 0.101 | 0.076 | 0.432 | 0.393 | 0.137 | 7.7E-07 |

| | | | | | | | |
|---|---|---|---|---|---|---|---|
| 0 | SHLD2 | 0.144 | 0.101 | 0.644 | 0.6 | 0.128 | 4.8E-14 |
| 0 | SIGIRR | 0.199 | 0.121 | 0.92 | 0.703 | 0.387 | 3.7E-34 |
| 0 | SIRT1 | 0.117 | 0.086 | 0.426 | 0.491 | 0.102 | 1.4E-08 |
| 0 | SIRT2 | 0.286 | 0.136 | 1.437 | 0.814 | 0.819 | 9E-106 |
| 0 | SKAP1 | 0.744 | 0.49 | 6.637 | 4.744 | 0.206 | 6.4E-47 |
| 0 | SKP1 | 0.684 | 0.449 | 4.928 | 4.332 | 0.186 | 6.4E-24 |
| 0 | SLC12A2 | 0.119 | 0.064 | 0.448 | 0.394 | 0.476 | 1.4E-29 |
| 0 | SLC20A1 | 0.3 | 0.212 | 1.402 | 1.294 | 0.214 | 4.3E-24 |
| 0 | SLC24A3 | 0.342 | 0.24 | 1.711 | 1.461 | 0.228 | 1.4E-24 |
| 0 | SLC24A32 | 0.166 | 0.101 | 0.74 | 0.474 | 0.387 | 2.3E-29 |
| 0 | SLC24A36 | 0.241 | 0.172 | 1.129 | 0.996 | 0.18 | 1.9E-19 |
| 0 | SLC24A4 | 0.413 | 0.231 | 2.272 | 1.374 | 0.724 | 1E-111 |
| 0 | SLC24A6 | 0.434 | 0.337 | 3.244 | 2.181 | 0.473 | 5E-103 |
| 0 | SLC2A3 | 0.306 | 0.184 | 1.706 | 1.134 | 0.488 | 4E-60 |
| 0 | SLC34D1 | 0.134 | 0.098 | 0.612 | 0.464 | 0.117 | 6.1E-10 |
| 0 | SLC34F1 | 0.147 | 0.116 | 0.683 | 0.603 | 0.181 | 1.4E-11 |
| 0 | SLC38A2 | 0.418 | 0.333 | 2.427 | 2.231 | 0.121 | 7.4E-16 |
| 0 | SLC44A2 | 0.238 | 0.164 | 1.096 | 0.976 | 0.167 | 4.1E-22 |
| 0 | SLC4A4 | 0.14 | 0.107 | 0.691 | 0.636 | 0.119 | 8.7E-13 |
| 0 | SLC9A3R1 | 0.348 | 0.24 | 1.866 | 1.42 | 0.296 | 3.2E-33 |
| 0 | SLCO3A1 | 0.222 | 0.126 | 1.048 | 0.87 | 0.283 | 3.1E-44 |
| 0 | SLCO4C1 | 0.191 | 0.112 | 0.878 | 0.692 | 0.343 | 2.2E-34 |
| 0 | SMAD7 | 0.312 | 0.227 | 1.607 | 1.34 | 0.242 | 2.7E-24 |
| 0 | SMAP2 | 0.381 | 0.244 | 2.021 | 1.611 | 0.327 | 1.8E-44 |
| 0 | SMARCB1 | 0.14 | 0.089 | 0.614 | 0.468 | 0.392 | 2.4E-21 |
| 0 | SMC4 | 0.314 | 0.237 | 1.489 | 1.447 | 0.124 | 4E-17 |
| 0 | SMIM10L1 | 0.199 | 0.147 | 0.934 | 0.793 | 0.236 | 6.1E-14 |
| 0 | SMNDC1 | 0.224 | 0.147 | 1.047 | 0.869 | 0.268 | 1.3E-20 |
| 0 | SNAP29 | 0.24 | 0.188 | 1.137 | 1.036 | 0.134 | 1.4E-11 |
| 0 | SNHG14 | 0.141 | 0.044 | 0.684 | 0.32 | 1.1 | 6.2E-91 |
| 0 | SNHG24 | 0.148 | 0.067 | 0.682 | 0.376 | 0.848 | 1E-48 |
| 0 | SNHG4 | 0.176 | 0.108 | 0.808 | 0.614 | 0.394 | 1.7E-29 |
| 0 | SNHG8 | 0.173 | 0.103 | 0.802 | 0.46 | 0.418 | 4.8E-33 |
| 0 | SNHG9 | 0.227 | 0.094 | 1.048 | 0.414 | 1.04 | 7E-108 |
| 0 | SNRK | 0.406 | 0.312 | 2.194 | 2.044 | 0.102 | 1.8E-18 |
| 0 | SNRNP27 | 0.139 | 0.102 | 0.601 | 0.44 | 0.13 | 6.1E-10 |
| 0 | SNRPA | 0.124 | 0.077 | 0.444 | 0.399 | 0.474 | 7E-21 |
| 0 | SNRPA1 | 0.14 | 0.098 | 0.647 | 0.429 | 0.311 | 1.2E-19 |
| 0 | SNRPB | 0.301 | 0.19 | 1.479 | 1.04 | 0.494 | 1.3E-47 |
| 0 | SNRPB2 | 0.188 | 0.146 | 0.871 | 0.783 | 0.143 | 8.2E-10 |
| 0 | SNRPC | 0.199 | 0.134 | 0.902 | 0.719 | 0.328 | 7.1E-22 |
| 0 | SNRPD1 | 0.2 | 0.127 | 0.892 | 0.682 | 0.388 | 1.5E-29 |
| 0 | SNRPD2 | 0.442 | 0.436 | 3.33 | 3.003 | 0.149 | 7.8E-21 |
| 0 | SNRPE | 0.131 | 0.093 | 0.466 | 0.477 | 0.249 | 4.4E-12 |
| 0 | SNRPF | 0.126 | 0.089 | 0.436 | 0.442 | 0.28 | 4.8E-12 |
| 0 | SNU13 | 0.226 | 0.147 | 1.041 | 0.871 | 0.27 | 7.9E-22 |
| 0 | SNX32 | 0.148 | 0.08 | 0.644 | 0.434 | 0.467 | 1.6E-37 |
| 0 | SPCS1 | 0.404 | 0.329 | 2.111 | 1.963 | 0.104 | 1.3E-11 |
| 0 | SPCS3 | 0.404 | 0.414 | 3.144 | 2.894 | 0.119 | 6.5E-14 |
| 0 | SPON2 | 0.708 | 0.249 | 6.822 | 2.048 | 1.736 | 0 |
| 0 | SPTAN1 | 0.349 | 0.233 | 1.807 | 1.401 | 0.367 | 3.5E-43 |
| 0 | SPTBN1 | 0.249 | 0.162 | 1.24 | 0.999 | 0.323 | 1.7E-38 |
| 0 | SPTY2D1 | 0.214 | 0.13 | 1.011 | 0.744 | 0.423 | 9.3E-37 |
| 0 | SRA1 | 0.179 | 0.128 | 0.78 | 0.703 | 0.141 | 2.9E-14 |

| | | | | | | | |
|---|---|---|---|---|---|---|---|
| 0 | SRGN | 0.912 | 0.76 | 12.637 | 9.489 | 0.413 | 3E-122 |
| 0 | SRP14 | 0.778 | 0.642 | 6.386 | 4.601 | 0.189 | 1.3E-31 |
| 0 | SRP19 | 0.318 | 0.246 | 1.468 | 1.429 | 0.134 | 4.3E-14 |
| 0 | SRP72 | 0.284 | 0.224 | 1.391 | 1.287 | 0.112 | 3E-12 |
| 0 | SRRM1 | 0.439 | 0.422 | 3.281 | 2.948 | 0.14 | 9.7E-22 |
| 0 | SRSF10 | 0.426 | 0.422 | 3.114 | 2.883 | 0.111 | 4E-16 |
| 0 | SRSF2 | 0.234 | 0.141 | 1.092 | 0.794 | 0.449 | 4.3E-43 |
| 0 | SRSF3 | 0.478 | 0.261 | 2.832 | 1.664 | 0.767 | 8E-144 |
| 0 | SRSF4 | 0.647 | 0.471 | 4.792 | 3.443 | 0.432 | 2.5E-82 |
| 0 | SRSF7 | 0.326 | 0.168 | 1.682 | 1.004 | 0.742 | 1E-101 |
| 0 | SSNA1 | 0.194 | 0.126 | 0.88 | 0.638 | 0.464 | 2.3E-27 |
| 0 | SSR2 | 0.376 | 0.264 | 1.974 | 1.436 | 0.362 | 8.7E-39 |
| 0 | SSR4 | 0.44 | 0.322 | 2.43 | 1.966 | 0.363 | 1.1E-43 |
| 0 | SSU72 | 0.184 | 0.134 | 0.83 | 0.744 | 0.146 | 4.5E-13 |
| 0 | ST13 | 0.371 | 0.284 | 1.943 | 1.718 | 0.184 | 2.9E-19 |
| 0 | ST3GAL1 | 0.129 | 0.081 | 0.467 | 0.466 | 0.281 | 3.5E-20 |
| 0 | STARD3NL | 0.241 | 0.184 | 1.222 | 1.074 | 0.186 | 1.4E-16 |
| 0 | STARD4 | 0.116 | 0.082 | 0.409 | 0.429 | 0.244 | 7.8E-11 |
| 0 | STEAP1B | 0.103 | 0.073 | 0.434 | 0.377 | 0.204 | 9.8E-10 |
| 0 | STK17B | 0.826 | 0.694 | 8.181 | 7.229 | 0.178 | 6.2E-29 |
| 0 | STK26 | 0.421 | 0.273 | 2.33 | 1.69 | 0.463 | 5E-63 |
| 0 | STK38 | 0.488 | 0.46 | 3.871 | 3.448 | 0.163 | 1.4E-24 |
| 0 | STMN1 | 0.164 | 0.093 | 0.846 | 0.441 | 0.619 | 8.2E-37 |
| 0 | STMP1 | 0.286 | 0.202 | 1.393 | 1.13 | 0.302 | 2.4E-26 |
| 0 | STN1 | 0.102 | 0.081 | 0.441 | 0.41 | 0.103 | 2.2E-05 |
| 0 | STRAP | 0.24 | 0.146 | 1.122 | 0.834 | 0.429 | 6.1E-42 |
| 0 | STUB1 | 0.141 | 0.111 | 0.646 | 0.444 | 0.243 | 3.4E-11 |
| 0 | SUB1 | 0.388 | 0.278 | 2.036 | 1.694 | 0.264 | 9.9E-32 |
| 0 | SUCO | 0.344 | 0.264 | 1.841 | 1.691 | 0.123 | 1.1E-19 |
| 0 | SUGP2 | 0.136 | 0.083 | 0.48 | 0.474 | 0.287 | 2.7E-22 |
| 0 | SUMO1 | 0.242 | 0.174 | 1.133 | 0.94 | 0.244 | 4.3E-20 |
| 0 | SUMO2 | 0.44 | 0.344 | 2.369 | 2.194 | 0.111 | 4.7E-13 |
| 0 | SUMO3 | 0.138 | 0.076 | 0.602 | 0.393 | 0.614 | 2.4E-34 |
| 0 | SUPT4H1 | 0.14 | 0.111 | 0.644 | 0.469 | 0.202 | 3.3E-11 |
| 0 | SUPV3L1 | 0.119 | 0.068 | 0.499 | 0.377 | 0.404 | 6.8E-24 |
| 0 | SURF4 | 0.136 | 0.103 | 0.602 | 0.444 | 0.147 | 6.9E-09 |
| 0 | SUSD1 | 0.146 | 0.04 | 0.648 | 0.342 | 0.944 | 8.2E-93 |
| 0 | SYAP1 | 0.388 | 0.261 | 2.14 | 1.666 | 0.368 | 1.2E-44 |
| 0 | SYF2 | 0.264 | 0.208 | 1.247 | 1.168 | 0.107 | 1E-11 |
| 0 | SYNE1 | 0.906 | 0.739 | 12.871 | 9.77 | 0.398 | 3E-144 |
| 0 | SYNGR1 | 0.162 | 0.093 | 0.743 | 0.427 | 0.496 | 2.1E-33 |
| 0 | SYTL1 | 0.2 | 0.092 | 0.896 | 0.449 | 0.706 | 6E-74 |
| 0 | SYTL3 | 0.606 | 0.386 | 4.403 | 3.219 | 0.484 | 2E-117 |
| 0 | SZRD1 | 0.21 | 0.142 | 0.964 | 0.847 | 0.187 | 3.5E-16 |
| 0 | TADA3 | 0.114 | 0.077 | 0.49 | 0.406 | 0.271 | 1.3E-13 |
| 0 | TAF10 | 0.207 | 0.143 | 0.919 | 0.744 | 0.304 | 8.1E-21 |
| 0 | TAF13 | 0.147 | 0.074 | 0.641 | 0.404 | 0.664 | 6.8E-44 |
| 0 | TAGAP | 0.397 | 0.242 | 2.763 | 1.601 | 0.787 | 4.3E-86 |
| 0 | TAGLN2 | 0.442 | 0.339 | 3.432 | 2.303 | 0.474 | 1E-107 |
| 0 | TBC1D10C | 0.243 | 0.142 | 1.181 | 0.847 | 0.464 | 9E-46 |
| 0 | TBCB | 0.263 | 0.184 | 1.244 | 1.007 | 0.304 | 1.9E-24 |
| 0 | TBCC | 0.124 | 0.093 | 0.437 | 0.464 | 0.208 | 4.8E-09 |
| 0 | TBX21 | 0.343 | 0.198 | 1.784 | 1.166 | 0.614 | 1.8E-74 |
| 0 | TBXAS1 | 0.139 | 0.081 | 0.642 | 0.481 | 0.417 | 1.4E-27 |

| | | | | | | | |
|---|---|---|---|---|---|---|---|
| 0 | TCF24 | 0.471 | 0.436 | 3.621 | 3.126 | 0.212 | 1.9E-32 |
| 0 | TECR | 0.169 | 0.114 | 0.744 | 0.608 | 0.311 | 2.4E-19 |
| 0 | TENT4B | 0.194 | 0.126 | 0.903 | 0.801 | 0.173 | 4.5E-24 |
| 0 | TENT4C | 0.236 | 0.121 | 1.142 | 0.736 | 0.633 | 2.1E-68 |
| 0 | TES | 0.446 | 0.312 | 2.614 | 2.046 | 0.347 | 4.4E-41 |
| 0 | TEX14 | 0.283 | 0.047 | 2.046 | 0.422 | 1.971 | 0 |
| 0 | TEX264 | 0.106 | 0.062 | 0.441 | 0.317 | 0.411 | 4.5E-21 |
| 0 | TFAM | 0.134 | 0.103 | 0.498 | 0.441 | 0.144 | 9E-08 |
| 0 | TFDP2 | 0.416 | 0.348 | 3.34 | 2.696 | 0.309 | 1.9E-47 |
| 0 | TG | 0.119 | 0.063 | 0.441 | 0.346 | 0.673 | 1.3E-31 |
| 0 | TGFB1 | 0.664 | 0.436 | 4.938 | 3.84 | 0.363 | 7E-112 |
| 0 | TGOLN2 | 0.444 | 0.346 | 2.41 | 2.288 | 0.134 | 3E-18 |
| 0 | THAP12 | 0.147 | 0.124 | 0.72 | 0.664 | 0.117 | 4.6E-07 |
| 0 | THOC7 | 0.208 | 0.147 | 0.976 | 0.792 | 0.302 | 8.1E-20 |
| 0 | THRA | 0.124 | 0.084 | 0.444 | 0.46 | 0.244 | 1.9E-13 |
| 0 | THUMPD1 | 0.261 | 0.196 | 1.27 | 1.111 | 0.194 | 3.4E-16 |
| 0 | THUMPD3-AS1 | 0.17 | 0.102 | 0.777 | 0.46 | 0.471 | 4.3E-31 |
| 0 | TIMM17B | 0.132 | 0.098 | 0.443 | 0.494 | 0.164 | 3E-09 |
| 0 | TIMM44 | 0.104 | 0.064 | 0.426 | 0.344 | 0.304 | 3.3E-16 |
| 0 | TIMP1 | 0.188 | 0.126 | 0.982 | 0.778 | 0.334 | 4.1E-22 |
| 0 | TIPARP | 0.344 | 0.182 | 1.841 | 1.192 | 0.634 | 3E-97 |
| 0 | TKT | 0.108 | 0.072 | 0.448 | 0.379 | 0.24 | 1.9E-13 |
| 0 | TLE1 | 0.303 | 0.144 | 1.443 | 0.987 | 0.644 | 6.9E-93 |
| 0 | TLE4 | 0.411 | 0.288 | 3.076 | 1.891 | 0.702 | 8E-139 |
| 0 | TLE4 | 0.407 | 0.344 | 3.149 | 2.491 | 0.338 | 1.8E-42 |
| 0 | TLN1 | 0.674 | 0.483 | 4.967 | 3.617 | 0.447 | 6.6E-92 |
| 0 | TM2D3 | 0.101 | 0.072 | 0.426 | 0.376 | 0.179 | 2.3E-09 |
| 0 | TMA7 | 0.73 | 0.482 | 4.609 | 4.761 | 0.236 | 1.4E-37 |
| 0 | TMBIM1 | 0.13 | 0.047 | 0.48 | 0.307 | 0.914 | 4.4E-44 |
| 0 | TMC6 | 0.113 | 0.073 | 0.484 | 0.384 | 0.336 | 4.3E-16 |
| 0 | TMED2 | 0.24 | 0.19 | 1.189 | 1.08 | 0.139 | 6.1E-14 |
| 0 | TMED4 | 0.249 | 0.192 | 1.144 | 1.044 | 0.144 | 2.1E-12 |
| 0 | TMED7 | 0.146 | 0.104 | 0.633 | 0.447 | 0.184 | 3.2E-12 |
| 0 | TMED9 | 0.181 | 0.109 | 0.807 | 0.473 | 0.494 | 1.8E-33 |
| 0 | TMEM107 | 0.127 | 0.041 | 0.497 | 0.282 | 1.082 | 4.5E-62 |
| 0 | TMEM109 | 0.194 | 0.136 | 0.884 | 0.724 | 0.284 | 1.7E-18 |
| 0 | TMEM160 | 0.2 | 0.143 | 0.887 | 0.744 | 0.242 | 2E-16 |
| 0 | TMEM167B | 0.144 | 0.099 | 0.626 | 0.416 | 0.279 | 2.2E-14 |
| 0 | TMEM170A | 0.163 | 0.127 | 0.74 | 0.682 | 0.118 | 4.7E-08 |
| 0 | TMEM183A | 0.191 | 0.138 | 0.844 | 0.741 | 0.188 | 3.3E-14 |
| 0 | TMEM230 | 0.244 | 0.144 | 1.117 | 0.864 | 0.368 | 1.4E-34 |
| 0 | TMEM242 | 0.133 | 0.09 | 0.484 | 0.487 | 0.267 | 9.2E-14 |
| 0 | TMEM243 | 0.112 | 0.084 | 0.484 | 0.449 | 0.107 | 2.1E-07 |
| 0 | TMEM246 | 0.103 | 0.07 | 0.432 | 0.337 | 0.348 | 2E-12 |
| 0 | TMEM248 | 0.28 | 0.222 | 1.304 | 1.206 | 0.113 | 4E-11 |
| 0 | TMEM71 | 0.306 | 0.196 | 1.614 | 1.189 | 0.44 | 1.3E-44 |
| 0 | TMEM9B | 0.217 | 0.169 | 1.011 | 0.936 | 0.112 | 3.4E-10 |
| 0 | TMIGD2 | 0.197 | 0.116 | 0.971 | 0.69 | 0.494 | 8.4E-39 |
| 0 | TMSB10 | 0.974 | 0.822 | 21.906 | 13.478 | 0.69 | 0 |
| 0 | TMX4 | 0.491 | 0.391 | 2.964 | 2.703 | 0.134 | 4E-18 |
| 0 | TNFRSF1B | 0.184 | 0.102 | 0.842 | 0.487 | 0.438 | 3.6E-44 |
| 0 | TOB1 | 0.111 | 0.06 | 0.484 | 0.334 | 0.433 | 2.3E-28 |
| 0 | TOMM20 | 0.233 | 0.144 | 1.082 | 0.828 | 0.386 | 2.8E-30 |
| 0 | TOMM22 | 0.17 | 0.093 | 0.742 | 0.479 | 0.641 | 1.5E-42 |

| | | | | | | | |
|---|---|---|---|---|---|---|---|
| 0 | TOMM4 | 0.264 | 0.177 | 1.246 | 0.947 | 0.38 | 2.4E-32 |
| 0 | TOMM6 | 0.298 | 0.226 | 1.434 | 1.249 | 0.2 | 6.2E-17 |
| 0 | TPD42 | 0.147 | 0.094 | 0.671 | 0.434 | 0.33 | 3.9E-21 |
| 0 | TPI1 | 0.474 | 0.44 | 3.647 | 3.014 | 0.274 | 1.4E-36 |
| 0 | TPM3 | 0.842 | 0.641 | 8.867 | 6.01 | 0.461 | 9E-182 |
| 0 | TPM4 | 0.342 | 0.139 | 1.928 | 0.946 | 1.012 | 1E-174 |
| 0 | TPP2 | 0.386 | 0.289 | 2.039 | 1.871 | 0.124 | 4.1E-22 |
| 0 | TPST2 | 0.38 | 0.173 | 2 | 1.031 | 0.946 | 6E-166 |
| 0 | TRA2A | 0.327 | 0.217 | 1.642 | 1.341 | 0.282 | 1.1E-38 |
| 0 | TRA2B | 0.409 | 0.347 | 3.088 | 2.316 | 0.414 | 4.7E-64 |
| 0 | TRABD2A | 0.168 | 0.101 | 0.728 | 0.474 | 0.344 | 6.8E-30 |
| 0 | TRAPPC1 | 0.291 | 0.209 | 1.393 | 1.147 | 0.268 | 2.1E-23 |
| 0 | TRBC2 | 0.441 | 0.423 | 3.461 | 3.064 | 0.216 | 1.1E-28 |
| 0 | TRDC | 0.383 | 0.29 | 2.19 | 1.901 | 0.204 | 4.9E-23 |
| 0 | TRIM8 | 0.103 | 0.076 | 0.438 | 0.402 | 0.127 | 7.7E-08 |
| 0 | TSC22D3 | 0.406 | 0.144 | 3.714 | 1.29 | 1.426 | 0 |
| 0 | TSN | 0.123 | 0.086 | 0.422 | 0.442 | 0.238 | 4.8E-12 |
| 0 | TSPAN2 | 0.127 | 0.084 | 0.444 | 0.442 | 0.328 | 4.6E-16 |
| 0 | TSPAN32 | 0.132 | 0.047 | 0.474 | 0.334 | 0.784 | 2E-47 |
| 0 | TSPO | 0.204 | 0.14 | 0.936 | 0.804 | 0.219 | 7.3E-14 |
| 0 | TSPYL2 | 0.322 | 0.122 | 2.24 | 0.944 | 1.231 | 5E-192 |
| 0 | TSTD1 | 0.229 | 0.176 | 1.086 | 0.977 | 0.144 | 6.9E-12 |
| 0 | TTC3 | 0.343 | 0.272 | 1.871 | 1.736 | 0.108 | 4.3E-16 |
| 0 | TTC38 | 0.339 | 0.134 | 1.804 | 0.842 | 1.083 | 8E-187 |
| 0 | TUBA1A | 0.406 | 0.207 | 2.476 | 1.314 | 0.97 | 3E-146 |
| 0 | TUBA1B | 0.297 | 0.144 | 1.488 | 0.83 | 0.842 | 1E-106 |
| 0 | TUBA4A | 0.142 | 0.047 | 0.636 | 0.321 | 0.986 | 6.1E-70 |
| 0 | TUBB4B | 0.177 | 0.087 | 0.828 | 0.491 | 0.743 | 7.9E-48 |
| 0 | TWF1 | 0.266 | 0.148 | 1.264 | 0.844 | 0.467 | 2.3E-62 |
| 0 | TWF2 | 0.142 | 0.092 | 0.604 | 0.474 | 0.341 | 6.3E-19 |
| 0 | TWISTNB | 0.103 | 0.048 | 0.448 | 0.271 | 0.726 | 1.2E-38 |
| 0 | TXK | 0.417 | 0.366 | 3.322 | 2.626 | 0.339 | 8.1E-42 |
| 0 | TXN | 0.214 | 0.146 | 0.974 | 0.847 | 0.203 | 4.1E-16 |
| 0 | TXN2 | 0.114 | 0.064 | 0.499 | 0.32 | 0.642 | 4.1E-27 |
| 0 | TXNIP | 0.972 | 0.831 | 32.466 | 17.033 | 0.931 | 0 |
| 0 | TXNL4A | 0.171 | 0.12 | 0.739 | 0.62 | 0.243 | 6.9E-16 |
| 0 | TYROBP | 0.904 | 0.777 | 11.382 | 9.987 | 0.189 | 1.4E-33 |
| 0 | UAP1 | 0.172 | 0.073 | 0.799 | 0.443 | 0.818 | 6.9E-78 |
| 0 | UBA42 | 0.823 | 0.678 | 7.747 | 6.742 | 0.201 | 1.7E-36 |
| 0 | UBB | 0.83 | 0.631 | 7.984 | 4.839 | 0.441 | 2E-117 |
| 0 | UBC | 0.893 | 0.778 | 10.84 | 9.646 | 0.167 | 2.8E-27 |
| 0 | UBE2A | 0.198 | 0.146 | 0.881 | 0.767 | 0.199 | 4.4E-14 |
| 0 | UBE2D3 | 0.464 | 0.437 | 3.444 | 3.11 | 0.142 | 1E-24 |
| 0 | UBE2E3 | 0.333 | 0.24 | 1.678 | 1.492 | 0.169 | 1.5E-19 |
| 0 | UBE2F | 0.34 | 0.227 | 1.818 | 1.374 | 0.403 | 3.8E-49 |
| 0 | UBE2J1 | 0.218 | 0.137 | 1.004 | 0.747 | 0.408 | 1.4E-33 |
| 0 | UBE2L3 | 0.331 | 0.243 | 1.64 | 1.403 | 0.126 | 4.2E-16 |
| 0 | UBE2S | 0.138 | 0.042 | 0.494 | 0.266 | 1.148 | 1E-76 |
| 0 | UBE2V1 | 0.413 | 0.344 | 3.084 | 2.329 | 0.404 | 3.5E-61 |
| 0 | UBE2V2 | 0.166 | 0.124 | 0.739 | 0.674 | 0.134 | 6.1E-11 |
| 0 | UBL4 | 0.388 | 0.296 | 1.969 | 1.73 | 0.187 | 8E-20 |
| 0 | UBLCP1 | 0.232 | 0.184 | 1.097 | 1.019 | 0.107 | 1.6E-09 |
| 0 | UCHL4 | 0.242 | 0.181 | 1.214 | 1.011 | 0.266 | 4E-21 |
| 0 | UCP2 | 0.437 | 0.203 | 2.48 | 1.277 | 0.948 | 1E-186 |

| | | | | | | | |
|---|---|---|---|---|---|---|---|
| 0 | UFC1 | 0.289 | 0.182 | 1.386 | 0.99 | 0.486 | 9E-47 |
| 0 | UFD1 | 0.164 | 0.116 | 0.74 | 0.623 | 0.248 | 8.4E-14 |
| 0 | UFM1 | 0.293 | 0.206 | 1.44 | 1.163 | 0.309 | 1.2E-27 |
| 0 | UPF3A | 0.213 | 0.162 | 1.032 | 0.914 | 0.174 | 3.7E-12 |
| 0 | UQCR11 | 0.48 | 0.444 | 3.491 | 3.114 | 0.206 | 4.3E-26 |
| 0 | UQCRB | 0.42 | 0.339 | 2.314 | 2.114 | 0.13 | 1.3E-13 |
| 0 | UQCRC1 | 0.118 | 0.072 | 0.491 | 0.347 | 0.449 | 3.5E-21 |
| 0 | UQCRFS1 | 0.232 | 0.14 | 1.068 | 0.803 | 0.413 | 1.4E-32 |
| 0 | UQCRH | 0.344 | 0.244 | 1.71 | 1.424 | 0.264 | 1.2E-29 |
| 0 | UQCRQ | 0.318 | 0.237 | 1.436 | 1.324 | 0.214 | 1.8E-20 |
| 0 | USB1 | 0.108 | 0.06 | 0.467 | 0.321 | 0.439 | 2.2E-24 |
| 0 | USP11 | 0.184 | 0.128 | 0.842 | 0.724 | 0.217 | 7.8E-18 |
| 0 | USP12 | 0.161 | 0.116 | 0.716 | 0.634 | 0.174 | 2.3E-13 |
| 0 | USP36 | 0.123 | 0.08 | 0.444 | 0.491 | 0.148 | 8.8E-16 |
| 0 | UXS1 | 0.102 | 0.076 | 0.444 | 0.404 | 0.164 | 1.2E-07 |
| 0 | VAMP2 | 0.484 | 0.448 | 3.696 | 3.229 | 0.194 | 2E-26 |
| 0 | VAMP3 | 0.118 | 0.08 | 0.406 | 0.421 | 0.264 | 1.3E-13 |
| 0 | VAMP7 | 0.144 | 0.103 | 0.606 | 0.433 | 0.184 | 1.5E-12 |
| 0 | VAMP8 | 0.37 | 0.276 | 1.863 | 1.606 | 0.214 | 4.4E-23 |
| 0 | VAPA | 0.32 | 0.243 | 1.487 | 1.462 | 0.119 | 4.3E-17 |
| 0 | VASP | 0.244 | 0.12 | 1.164 | 0.676 | 0.784 | 2.9E-84 |
| 0 | VAV3 | 0.494 | 0.347 | 3.124 | 2.714 | 0.203 | 4.1E-39 |
| 0 | VCL | 0.367 | 0.191 | 2.004 | 1.221 | 0.714 | 3E-110 |
| 0 | VCP | 0.292 | 0.224 | 1.414 | 1.297 | 0.124 | 2.2E-14 |
| 0 | VDAC2 | 0.294 | 0.192 | 1.419 | 1.084 | 0.387 | 2.9E-39 |
| 0 | VMA21 | 0.122 | 0.068 | 0.416 | 0.341 | 0.444 | 2.3E-29 |
| 0 | VPS26A | 0.277 | 0.193 | 1.338 | 1.12 | 0.247 | 1E-24 |
| 0 | VPS26B | 0.102 | 0.067 | 0.433 | 0.36 | 0.268 | 1.2E-13 |
| 0 | VPS37B | 0.322 | 0.2 | 1.612 | 1.31 | 0.299 | 4.9E-49 |
| 0 | VSIR | 0.124 | 0.044 | 0.434 | 0.262 | 1.034 | 3.7E-72 |
| 0 | WARS2 | 0.144 | 0.047 | 0.691 | 0.347 | 0.996 | 1E-74 |
| 0 | WAS | 0.44 | 0.334 | 2.48 | 2.094 | 0.244 | 1E-28 |
| 0 | WBP11 | 0.301 | 0.218 | 1.468 | 1.294 | 0.182 | 1.2E-21 |
| 0 | WDR1 | 0.21 | 0.134 | 0.973 | 0.749 | 0.376 | 1.4E-29 |
| 0 | WDR26 | 0.249 | 0.182 | 1.287 | 1.14 | 0.162 | 1.2E-22 |
| 0 | WDR37 | 0.236 | 0.147 | 1.106 | 0.848 | 0.384 | 1.7E-36 |
| 0 | WDR43 | 0.194 | 0.139 | 0.886 | 0.794 | 0.148 | 1.3E-14 |
| 0 | WDR44 | 0.128 | 0.094 | 0.443 | 0.482 | 0.174 | 3.3E-09 |
| 0 | WDR44B | 0.242 | 0.163 | 1.116 | 0.921 | 0.276 | 8.4E-27 |
| 0 | WDR74 | 0.297 | 0.082 | 1.774 | 0.483 | 1.606 | 5E-271 |
| 0 | XBP1 | 0.313 | 0.147 | 1.474 | 0.874 | 0.848 | 2E-121 |
| 0 | XCL1 | 0.282 | 0.144 | 1.819 | 1.618 | 0.169 | 4E-66 |
| 0 | XCL2 | 0.323 | 0.143 | 2.062 | 1.247 | 0.714 | 8E-122 |
| 0 | XYLT1 | 0.114 | 0.064 | 0.409 | 0.417 | 0.286 | 1.7E-24 |
| 0 | YBX1 | 0.732 | 0.418 | 4.774 | 3.947 | 0.444 | 7E-134 |
| 0 | YES1 | 0.479 | 0.243 | 2.939 | 1.999 | 0.446 | 8E-140 |
| 0 | YME1L1 | 0.414 | 0.264 | 2.318 | 1.687 | 0.449 | 3.4E-64 |
| 0 | YPEL4 | 0.427 | 0.307 | 3.337 | 2.114 | 0.648 | 2E-134 |
| 0 | YTHDF3 | 0.208 | 0.147 | 0.944 | 0.874 | 0.127 | 4E-17 |
| 0 | YWHAB | 0.771 | 0.641 | 6.486 | 4.997 | 0.134 | 2.8E-17 |
| 0 | YWHAQ | 0.406 | 0.361 | 3.007 | 2.449 | 0.296 | 1.3E-44 |
| 0 | YWHAZ | 0.941 | 0.809 | 13.904 | 10.388 | 0.42 | 7E-144 |
| 0 | Z93241.1 | 0.178 | 0.033 | 1.037 | 0.242 | 2.1 | 3E-224 |
| 0 | Z99127.4 | 0.114 | 0.036 | 0.43 | 0.204 | 1.371 | 2.3E-82 |

| | | | | | | | |
|---|---|---|---|---|---|---|---|
| 0 | ZAP70 | 0.314 | 0.227 | 1.448 | 1.418 | 0.126 | 4.2E-23 |
| 0 | ZBTB1 | 0.402 | 0.348 | 3.149 | 2.444 | 0.37 | 4.9E-46 |
| 0 | ZBTB10 | 0.213 | 0.078 | 1.022 | 0.486 | 1.073 | 4E-124 |
| 0 | ZBTB16 | 0.264 | 0.164 | 1.3 | 1.142 | 0.174 | 4.1E-39 |
| 0 | ZBTB21 | 0.114 | 0.063 | 0.494 | 0.361 | 0.444 | 1.7E-26 |
| 0 | ZC3H12A | 0.161 | 0.074 | 0.739 | 0.423 | 0.804 | 2.4E-61 |
| 0 | ZC3H14 | 0.323 | 0.234 | 1.634 | 1.436 | 0.186 | 8.3E-23 |
| 0 | ZCCHC17 | 0.191 | 0.131 | 0.867 | 0.742 | 0.224 | 2.1E-19 |
| 0 | ZCRB1 | 0.113 | 0.088 | 0.4 | 0.461 | 0.117 | 4.1E-06 |
| 0 | ZEB2 | 0.828 | 0.718 | 9.19 | 8.429 | 0.108 | 1.4E-20 |
| 0 | ZFAND2A | 0.116 | 0.074 | 0.498 | 0.393 | 0.343 | 1.4E-17 |
| 0 | ZFAND4 | 0.246 | 0.177 | 1.244 | 1.032 | 0.282 | 1E-23 |
| 0 | ZFAS1 | 0.124 | 0.087 | 0.439 | 0.474 | 0.188 | 8.1E-12 |
| 0 | ZFC3H1 | 0.391 | 0.293 | 2.186 | 1.9 | 0.203 | 6.8E-24 |
| 0 | ZFP36 | 0.747 | 0.329 | 9.481 | 2.906 | 1.706 | 0 |
| 0 | ZFP36L1 | 0.474 | 0.36 | 2.949 | 2.404 | 0.236 | 2E-28 |
| 0 | ZFP36L2 | 0.837 | 0.634 | 9.388 | 6.418 | 0.449 | 6E-140 |
| 0 | ZFP91 | 0.234 | 0.179 | 1.102 | 1.01 | 0.127 | 1.9E-12 |
| 0 | ZHX1 | 0.111 | 0.082 | 0.404 | 0.447 | 0.144 | 2E-08 |
| 0 | ZMAT4 | 0.102 | 0.047 | 0.419 | 0.309 | 0.747 | 1.6E-38 |
| 0 | ZNF240 | 0.132 | 0.06 | 0.487 | 0.349 | 0.749 | 1.2E-49 |
| 0 | ZNF276 | 0.161 | 0.103 | 0.736 | 0.493 | 0.311 | 7.3E-23 |
| 0 | ZNF331 | 0.304 | 0.108 | 1.916 | 0.883 | 1.118 | 3E-193 |
| 0 | ZNF394 | 0.227 | 0.144 | 1.042 | 0.878 | 0.261 | 8.6E-24 |
| 0 | ZNF410 | 0.123 | 0.08 | 0.437 | 0.448 | 0.229 | 1E-16 |
| 0 | ZNF428 | 0.17 | 0.117 | 0.761 | 0.6 | 0.342 | 4.2E-18 |
| 0 | ZNF622 | 0.146 | 0.097 | 0.638 | 0.424 | 0.28 | 1E-17 |
| 0 | ZNF791 | 0.124 | 0.079 | 0.448 | 0.438 | 0.323 | 6.7E-19 |
| 0 | ZNF831 | 0.146 | 0.089 | 0.67 | 0.42 | 0.366 | 1.3E-24 |
| 0 | ZNF92 | 0.136 | 0.089 | 0.609 | 0.48 | 0.344 | 4.7E-18 |
| 0 | ZNHIT1 | 0.163 | 0.103 | 0.727 | 0.438 | 0.434 | 3.5E-24 |
| 0 | ZNRD2 | 0.146 | 0.082 | 0.629 | 0.434 | 0.434 | 8E-33 |
| 1 | 45357 | 0.194 | 0.249 | 1.604 | 1.232 | 0.381 | 4.1E-07 |
| 1 | 45358 | 0.232 | 0.331 | 1.79 | 1.648 | 0.119 | 4.9E-14 |
| 1 | AAK1 | 0.402 | 0.433 | 3.702 | 3.221 | 0.201 | 6.5E-05 |
| 1 | AATF | 0.114 | 0.17 | 0.87 | 0.764 | 0.187 | 9.5E-14 |
| 1 | ABCB1 | 0.247 | 0.336 | 2.207 | 1.82 | 0.278 | 4.4E-06 |
| 1 | ABCD3 | 0.102 | 0.128 | 0.77 | 0.443 | 0.479 | 0.00072 |
| 1 | ABCF1 | 0.162 | 0.238 | 1.231 | 1.132 | 0.121 | 1.4E-16 |
| 1 | ABHD2 | 0.181 | 0.24 | 1.497 | 1.23 | 0.283 | 4.2E-10 |
| 1 | ABI1 | 0.343 | 0.466 | 2.948 | 2.611 | 0.18 | 3.4E-08 |
| 1 | ABLIM1 | 0.186 | 0.246 | 1.424 | 1.294 | 0.236 | 2.3E-10 |
| 1 | AC004687.1 | 0.103 | 0.137 | 0.747 | 0.604 | 0.323 | 2.3E-06 |
| 1 | AC008014.1 | 0.137 | 0.196 | 1.196 | 0.986 | 0.279 | 3.8E-12 |
| 1 | AC014849.1 | 0.402 | 0.436 | 3.699 | 3.306 | 0.162 | 1.5E-06 |
| 1 | AC079793.1 | 0.101 | 0.141 | 0.744 | 0.644 | 0.203 | 7.1E-13 |
| 1 | AC118449.1 | 0.106 | 0.148 | 0.773 | 0.714 | 0.112 | 1.7E-13 |
| 1 | AC244297.3 | 0.33 | 0.434 | 3.002 | 2.484 | 0.216 | 4.2E-06 |
| 1 | ACAP2 | 0.33 | 0.447 | 2.877 | 2.498 | 0.147 | 4E-11 |
| 1 | ACBD3 | 0.104 | 0.144 | 0.764 | 0.69 | 0.148 | 1.3E-13 |
| 1 | ACBD4 | 0.118 | 0.177 | 0.886 | 0.794 | 0.148 | 8.8E-14 |
| 1 | ACBD6 | 0.138 | 0.197 | 1.062 | 0.919 | 0.21 | 5E-12 |
| 1 | ACIN1 | 0.247 | 0.363 | 2.029 | 1.876 | 0.113 | 1.9E-18 |
| 1 | ACTN4 | 0.169 | 0.246 | 1.374 | 1.18 | 0.219 | 1.3E-14 |

| | | | | | | | |
|---|---|---|---|---|---|---|---|
| 1 | ACYP2 | 0.17 | 0.233 | 1.314 | 1.098 | 0.261 | 7.9E-10 |
| 1 | ADCY7 | 0.109 | 0.149 | 0.823 | 0.644 | 0.332 | 2.2E-07 |
| 1 | ADGRE4 | 0.264 | 0.403 | 2.633 | 2.383 | 0.144 | 1.4E-27 |
| 1 | ADGRG1 | 0.104 | 0.149 | 0.847 | 0.74 | 0.174 | 8.9E-14 |
| 1 | ADGRG4 | 0.109 | 0.144 | 0.829 | 0.644 | 0.361 | 4.2E-06 |
| 1 | ADK | 0.181 | 0.233 | 1.478 | 1.144 | 0.346 | 3.5E-05 |
| 1 | AEBP2 | 0.143 | 0.219 | 1.181 | 1.048 | 0.173 | 3.3E-13 |
| 1 | AFF4 | 0.19 | 0.279 | 1.461 | 1.363 | 0.196 | 1.2E-16 |
| 1 | AFTPH | 0.143 | 0.203 | 1.071 | 0.943 | 0.184 | 1.9E-11 |
| 1 | AGO2 | 0.116 | 0.146 | 0.901 | 0.643 | 0.464 | 0.00027 |
| 1 | AGO3 | 0.149 | 0.216 | 1.228 | 1.024 | 0.262 | 1.6E-08 |
| 1 | AGPAT4 | 0.173 | 0.241 | 1.327 | 1.184 | 0.163 | 4.7E-12 |
| 1 | AGPS | 0.106 | 0.17 | 0.809 | 0.744 | 0.1 | 2.5E-19 |
| 1 | AHCTF1 | 0.118 | 0.167 | 0.904 | 0.764 | 0.243 | 1.8E-10 |
| 1 | AHNAK | 0.303 | 0.404 | 2.91 | 2.406 | 0.274 | 4.1E-07 |
| 1 | AKAP13 | 0.404 | 0.646 | 4.33 | 4.892 | 0.124 | 0.0001 |
| 1 | AKAP9 | 0.33 | 0.427 | 2.944 | 2.407 | 0.237 | 0.00011 |
| 1 | AKNA | 0.282 | 0.367 | 2.668 | 2.048 | 0.382 | 0.00013 |
| 1 | AL138963.4 | 0.187 | 0.279 | 1.489 | 1.341 | 0.244 | 1.7E-17 |
| 1 | ALG13 | 0.129 | 0.198 | 1.01 | 0.924 | 0.126 | 4.4E-18 |
| 1 | ANKHD1 | 0.298 | 0.4 | 2.634 | 2.194 | 0.264 | 1.2E-07 |
| 1 | ANKRD10 | 0.146 | 0.216 | 1.183 | 0.997 | 0.247 | 2.5E-10 |
| 1 | ANKRD17 | 0.224 | 0.298 | 1.868 | 1.497 | 0.319 | 2.2E-07 |
| 1 | ANKRD28 | 0.126 | 0.167 | 1.017 | 0.767 | 0.407 | 2.7E-06 |
| 1 | ANKRD36 | 0.193 | 0.244 | 1.603 | 1.163 | 0.463 | 0.00099 |
| 1 | ANKRD36C | 0.217 | 0.283 | 1.804 | 1.416 | 0.349 | 7.3E-06 |
| 1 | ANKRD44 | 0.499 | 0.482 | 4.464 | 3.88 | 0.494 | 8.8E-09 |
| 1 | ANO6 | 0.124 | 0.178 | 0.942 | 0.792 | 0.249 | 8.3E-11 |
| 1 | AOAH | 0.801 | 0.86 | 14.623 | 12.843 | 0.282 | 9.4E-32 |
| 1 | AP1G1 | 0.141 | 0.217 | 1.213 | 0.999 | 0.279 | 2.9E-12 |
| 1 | AP2A2 | 0.111 | 0.141 | 0.861 | 0.664 | 0.373 | 2.2E-07 |
| 1 | AP3B1 | 0.294 | 0.404 | 2.472 | 2.187 | 0.234 | 3.1E-09 |
| 1 | APBB1IP | 0.411 | 0.44 | 3.798 | 3.403 | 0.148 | 2.1E-05 |
| 1 | APC | 0.177 | 0.249 | 1.391 | 1.227 | 0.182 | 2.6E-12 |
| 1 | APMAP | 0.404 | 0.442 | 3.703 | 3.447 | 0.103 | 4.9E-09 |
| 1 | APPBP2 | 0.107 | 0.143 | 0.783 | 0.639 | 0.294 | 6.9E-07 |
| 1 | AQR | 0.141 | 0.198 | 1.074 | 0.904 | 0.249 | 1.3E-10 |
| 1 | ARFGEF1 | 0.273 | 0.361 | 2.314 | 1.91 | 0.278 | 8.6E-07 |
| 1 | ARGLU1 | 0.402 | 0.644 | 4.062 | 4.448 | 0.141 | 0.00317 |
| 1 | ARHGAP14 | 0.476 | 0.704 | 7.197 | 4.619 | 0.347 | 0.00037 |
| 1 | ARHGAP30 | 0.21 | 0.302 | 1.689 | 1.478 | 0.192 | 2E-14 |
| 1 | ARHGAP4 | 0.101 | 0.143 | 0.788 | 0.714 | 0.143 | 1.4E-14 |
| 1 | ARHGEF1 | 0.233 | 0.331 | 1.912 | 1.719 | 0.144 | 4.3E-14 |
| 1 | ARHGEF7 | 0.174 | 0.22 | 1.414 | 1.021 | 0.469 | 0.00037 |
| 1 | ARID1A | 0.114 | 0.146 | 0.899 | 0.69 | 0.381 | 1.2E-07 |
| 1 | ARID4A | 0.166 | 0.214 | 1.307 | 1.034 | 0.336 | 1.4E-05 |
| 1 | ARID4B | 0.443 | 0.489 | 4.474 | 4.006 | 0.192 | 4.9E-06 |
| 1 | ARIH1 | 0.243 | 0.349 | 2.281 | 1.914 | 0.243 | 4.8E-13 |
| 1 | ARMC8 | 0.109 | 0.148 | 0.832 | 0.646 | 0.366 | 3E-07 |
| 1 | ASAP1 | 0.124 | 0.171 | 0.973 | 0.788 | 0.304 | 1.2E-08 |
| 1 | ASB3 | 0.124 | 0.16 | 0.944 | 0.72 | 0.393 | 3.9E-05 |
| 1 | ASCC3 | 0.216 | 0.307 | 1.816 | 1.483 | 0.199 | 2.6E-14 |
| 1 | ASH1L | 0.323 | 0.418 | 2.926 | 2.342 | 0.321 | 0.00084 |
| 1 | ASXL2 | 0.13 | 0.184 | 0.944 | 0.847 | 0.171 | 2E-11 |

| | | | | | | | |
|---|---|---|---|---|---|---|---|
| 1 | ATAD2B | 0.24 | 0.346 | 2.032 | 1.77 | 0.199 | 4.1E-14 |
| 1 | ATE1 | 0.148 | 0.219 | 1.218 | 1.02 | 0.246 | 3.8E-10 |
| 1 | ATF2 | 0.139 | 0.191 | 1.063 | 0.877 | 0.276 | 3.7E-09 |
| 1 | ATF6 | 0.126 | 0.177 | 0.944 | 0.798 | 0.243 | 4.4E-10 |
| 1 | ATF6B | 0.106 | 0.16 | 0.749 | 0.704 | 0.107 | 1.5E-14 |
| 1 | ATF7IP | 0.303 | 0.423 | 2.684 | 2.346 | 0.194 | 6.4E-12 |
| 1 | ATG10 | 0.109 | 0.142 | 0.843 | 0.671 | 0.329 | 6.1E-09 |
| 1 | ATG14 | 0.133 | 0.186 | 0.977 | 0.834 | 0.226 | 2.6E-10 |
| 1 | ATP10A | 0.14 | 0.182 | 1.144 | 0.838 | 0.448 | 3E-05 |
| 1 | ATP11B | 0.338 | 0.448 | 3.042 | 2.444 | 0.262 | 2.3E-05 |
| 1 | ATP2B1 | 0.173 | 0.23 | 1.448 | 1.108 | 0.396 | 8.4E-07 |
| 1 | ATP2B4 | 0.248 | 0.341 | 2.126 | 1.87 | 0.184 | 4.2E-10 |
| 1 | ATP2C1 | 0.139 | 0.191 | 1.043 | 0.86 | 0.291 | 4.8E-09 |
| 1 | ATP8A1 | 0.479 | 0.444 | 4.099 | 3.718 | 0.446 | 1.2E-07 |
| 1 | ATP9B | 0.194 | 0.267 | 1.612 | 1.284 | 0.328 | 1.9E-09 |
| 1 | ATR | 0.144 | 0.229 | 1.273 | 1.112 | 0.194 | 2.3E-14 |
| 1 | ATRX | 0.373 | 0.498 | 3.434 | 3.02 | 0.184 | 2E-06 |
| 1 | ATXN10 | 0.101 | 0.137 | 0.741 | 0.601 | 0.322 | 3.6E-07 |
| 1 | ATXN2 | 0.118 | 0.164 | 0.901 | 0.736 | 0.292 | 2.8E-09 |
| 1 | ATXN7 | 0.119 | 0.168 | 0.927 | 0.76 | 0.286 | 4.4E-10 |
| 1 | ATXN7L1 | 0.108 | 0.14 | 0.823 | 0.629 | 0.386 | 3.3E-05 |
| 1 | AUH | 0.163 | 0.226 | 1.284 | 1.06 | 0.277 | 6.1E-10 |
| 1 | AVL9 | 0.114 | 0.148 | 0.864 | 0.711 | 0.281 | 1.9E-08 |
| 1 | AXIN1 | 0.111 | 0.148 | 0.998 | 0.671 | 0.471 | 4.2E-06 |
| 1 | B4GALT1 | 0.236 | 0.33 | 1.969 | 1.664 | 0.243 | 2.3E-11 |
| 1 | BABAM2 | 0.17 | 0.208 | 1.422 | 0.999 | 0.41 | 0.00749 |
| 1 | BACH1 | 0.173 | 0.24 | 1.36 | 1.14 | 0.243 | 1.3E-10 |
| 1 | BAZ1B | 0.148 | 0.224 | 1.143 | 1.064 | 0.104 | 1.1E-13 |
| 1 | BAZ2A | 0.147 | 0.222 | 1.108 | 1.03 | 0.106 | 1.2E-17 |
| 1 | BAZ2B | 0.128 | 0.179 | 1.037 | 0.864 | 0.261 | 3.1E-10 |
| 1 | BBX | 0.314 | 0.402 | 2.846 | 2.289 | 0.32 | 0.0016 |
| 1 | BCL11B | 0.219 | 0.303 | 1.806 | 1.469 | 0.202 | 2.4E-11 |
| 1 | BDP1 | 0.284 | 0.384 | 2.443 | 2.147 | 0.179 | 1.4E-09 |
| 1 | BICRAL | 0.106 | 0.164 | 0.828 | 0.762 | 0.119 | 4.6E-17 |
| 1 | BMPR2 | 0.124 | 0.18 | 0.996 | 0.849 | 0.213 | 4.5E-12 |
| 1 | BNC2 | 0.144 | 0.187 | 1.337 | 0.936 | 0.414 | 1.2E-05 |
| 1 | BNIP3L | 0.136 | 0.179 | 1.04 | 0.811 | 0.349 | 2.3E-06 |
| 1 | BOD1L1 | 0.184 | 0.244 | 1.449 | 1.27 | 0.19 | 1.2E-10 |
| 1 | BPTF | 0.397 | 0.412 | 3.734 | 3.178 | 0.233 | 0.00641 |
| 1 | BRAF | 0.239 | 0.304 | 2.09 | 1.443 | 0.428 | 0.00111 |
| 1 | BRWD1 | 0.242 | 0.337 | 2.042 | 1.746 | 0.224 | 4.1E-08 |
| 1 | BRWD3 | 0.12 | 0.148 | 0.903 | 0.714 | 0.337 | 2E-06 |
| 1 | BTAF1 | 0.174 | 0.239 | 1.418 | 1.124 | 0.334 | 4.1E-09 |
| 1 | BTBD1 | 0.123 | 0.188 | 0.943 | 0.84 | 0.166 | 1.5E-16 |
| 1 | BTBD9 | 0.203 | 0.266 | 1.84 | 1.371 | 0.432 | 9.7E-06 |
| 1 | BTN3A1 | 0.279 | 0.374 | 2.311 | 2.042 | 0.178 | 1.2E-08 |
| 1 | BTN3A3 | 0.126 | 0.188 | 0.932 | 0.844 | 0.123 | 3.3E-15 |
| 1 | C2CD4 | 0.111 | 0.142 | 0.807 | 0.681 | 0.244 | 6.4E-08 |
| 1 | CAB39 | 0.184 | 0.244 | 1.429 | 1.184 | 0.271 | 1E-07 |
| 1 | CAMK1D | 0.2 | 0.283 | 1.88 | 1.444 | 0.381 | 9.8E-12 |
| 1 | CAMK2D | 0.217 | 0.297 | 1.922 | 1.426 | 0.333 | 4.4E-09 |
| 1 | CAMKMT | 0.148 | 0.203 | 1.321 | 1.028 | 0.362 | 1.5E-05 |
| 1 | CAMTA1 | 0.111 | 0.167 | 0.83 | 0.736 | 0.173 | 1.5E-14 |
| 1 | CAND1 | 0.173 | 0.243 | 1.307 | 1.146 | 0.177 | 6.7E-12 |

| | | | | | | | |
|---|---|---|---|---|---|---|---|
| 1 | CAPN7 | 0.176 | 0.24 | 1.348 | 1.202 | 0.164 | 4.1E-13 |
| 1 | CAPRIN1 | 0.149 | 0.223 | 1.214 | 1.033 | 0.234 | 4.1E-11 |
| 1 | CAPZA2 | 0.174 | 0.263 | 1.347 | 1.222 | 0.141 | 1.4E-18 |
| 1 | CARD11 | 0.219 | 0.293 | 2.13 | 1.412 | 0.494 | 4.3E-07 |
| 1 | CARD8 | 0.19 | 0.262 | 1.439 | 1.267 | 0.184 | 1.7E-10 |
| 1 | CASC3 | 0.113 | 0.148 | 0.844 | 0.644 | 0.384 | 7E-06 |
| 1 | CASP8 | 0.188 | 0.277 | 1.402 | 1.321 | 0.184 | 2.8E-16 |
| 1 | CBFA2T2 | 0.113 | 0.138 | 0.847 | 0.617 | 0.474 | 0.00264 |
| 1 | CBFB | 0.133 | 0.197 | 1.011 | 0.89 | 0.184 | 4.9E-14 |
| 1 | CBL | 0.106 | 0.148 | 0.747 | 0.694 | 0.124 | 1.1E-13 |
| 1 | CBR4 | 0.142 | 0.171 | 1.096 | 0.824 | 0.411 | 0.00881 |
| 1 | CCAR1 | 0.133 | 0.188 | 1.001 | 0.844 | 0.244 | 2.1E-10 |
| 1 | CCDC12 | 0.164 | 0.231 | 1.197 | 1.069 | 0.163 | 1.6E-11 |
| 1 | CCDC18-AS1 | 0.122 | 0.173 | 0.908 | 0.817 | 0.143 | 4.6E-11 |
| 1 | CCDC82 | 0.114 | 0.148 | 0.847 | 0.724 | 0.241 | 1.3E-08 |
| 1 | CCDC88C | 0.213 | 0.284 | 1.723 | 1.413 | 0.287 | 1.4E-07 |
| 1 | CCDC91 | 0.24 | 0.341 | 2.084 | 1.791 | 0.218 | 1.5E-13 |
| 1 | CCDC93 | 0.127 | 0.173 | 1.009 | 0.789 | 0.344 | 2.4E-08 |
| 1 | CCM2 | 0.148 | 0.187 | 1.101 | 0.87 | 0.339 | 0.00013 |
| 1 | CCNDBP1 | 0.117 | 0.163 | 0.806 | 0.719 | 0.166 | 2.1E-09 |
| 1 | CCNH | 0.393 | 0.433 | 3.964 | 3.694 | 0.102 | 1.7E-11 |
| 1 | CCNL2 | 0.124 | 0.163 | 0.919 | 0.744 | 0.304 | 4E-06 |
| 1 | CCNT1 | 0.106 | 0.172 | 0.838 | 0.767 | 0.128 | 2.8E-20 |
| 1 | CCNT2 | 0.143 | 0.224 | 1.109 | 1.019 | 0.122 | 3E-21 |
| 1 | CCNY | 0.116 | 0.144 | 1.014 | 0.702 | 0.432 | 2.4E-06 |
| 1 | CCPG1 | 0.2 | 0.281 | 1.492 | 1.38 | 0.113 | 1.7E-13 |
| 1 | CD247 | 0.816 | 0.896 | 14.348 | 12.004 | 0.346 | 2.2E-24 |
| 1 | CD46 | 0.317 | 0.434 | 2.668 | 2.404 | 0.141 | 1.4E-09 |
| 1 | CD48 | 0.194 | 0.263 | 1.479 | 1.318 | 0.166 | 9.7E-10 |
| 1 | CDC14A | 0.141 | 0.223 | 1.249 | 1.091 | 0.194 | 3.4E-14 |
| 1 | CDC16 | 0.131 | 0.187 | 0.928 | 0.848 | 0.113 | 2.4E-12 |
| 1 | CDC27 | 0.127 | 0.18 | 0.968 | 0.812 | 0.243 | 1.5E-10 |
| 1 | CDC42SE2 | 0.641 | 0.777 | 8.223 | 7.023 | 0.228 | 0.00023 |
| 1 | CDC73 | 0.208 | 0.299 | 1.628 | 1.477 | 0.141 | 4.6E-14 |
| 1 | CDK11A | 0.13 | 0.192 | 1.02 | 0.87 | 0.231 | 1.1E-13 |
| 1 | CDK11B | 0.14 | 0.214 | 1.124 | 0.986 | 0.191 | 1.1E-17 |
| 1 | CDK12 | 0.124 | 0.176 | 0.946 | 0.79 | 0.261 | 2.4E-10 |
| 1 | CDK13 | 0.287 | 0.379 | 2.408 | 2 | 0.326 | 3.9E-05 |
| 1 | CDK17 | 0.419 | 0.449 | 4.343 | 3.406 | 0.312 | 0.00399 |
| 1 | CDK19 | 0.134 | 0.189 | 1.037 | 0.869 | 0.244 | 1.4E-10 |
| 1 | CDK4RAP2 | 0.144 | 0.208 | 1.104 | 0.996 | 0.14 | 1.3E-13 |
| 1 | CDKAL1 | 0.237 | 0.303 | 2.042 | 1.42 | 0.433 | 0.00062 |
| 1 | CDYL | 0.101 | 0.123 | 0.806 | 0.434 | 0.493 | 0.00473 |
| 1 | CELF1 | 0.124 | 0.16 | 0.978 | 0.704 | 0.474 | 8.8E-05 |
| 1 | CEMIP2 | 0.342 | 0.443 | 3.971 | 3.08 | 0.367 | 0.00103 |
| 1 | CENPC | 0.219 | 0.319 | 1.794 | 1.621 | 0.147 | 8.5E-17 |
| 1 | CEP192 | 0.131 | 0.17 | 1.017 | 0.778 | 0.388 | 1.2E-05 |
| 1 | CEP340 | 0.311 | 0.409 | 2.684 | 2.348 | 0.193 | 8.5E-07 |
| 1 | CEP84L | 0.211 | 0.293 | 1.74 | 1.469 | 0.244 | 2.7E-11 |
| 1 | CEP94 | 0.141 | 0.214 | 1.172 | 0.996 | 0.234 | 1.3E-11 |
| 1 | CERT1 | 0.184 | 0.246 | 1.429 | 1.242 | 0.191 | 2.3E-11 |
| 1 | CFDP1 | 0.124 | 0.177 | 0.939 | 0.831 | 0.174 | 3.3E-11 |
| 1 | CHCHD3 | 0.144 | 0.22 | 1.196 | 0.997 | 0.262 | 1.5E-11 |
| 1 | CHD1 | 0.343 | 0.481 | 3.426 | 3.006 | 0.189 | 2.5E-10 |

| | | | | | | | |
|---|---|---|---|---|---|---|---|
| 1 | CHD2 | 0.404 | 0.444 | 3.944 | 3.667 | 0.109 | 2.2E-10 |
| 1 | CHD3 | 0.178 | 0.248 | 1.377 | 1.227 | 0.166 | 6.3E-14 |
| 1 | CHD4 | 0.216 | 0.306 | 1.804 | 1.423 | 0.244 | 1.4E-12 |
| 1 | CHD6 | 0.164 | 0.203 | 1.301 | 0.933 | 0.48 | 0.00399 |
| 1 | CHD8 | 0.142 | 0.199 | 1.091 | 0.914 | 0.243 | 2.4E-10 |
| 1 | CHD9 | 0.206 | 0.284 | 1.606 | 1.417 | 0.18 | 3.1E-11 |
| 1 | CHIC2 | 0.236 | 0.339 | 1.914 | 1.702 | 0.169 | 2E-14 |
| 1 | CIR1 | 0.216 | 0.314 | 1.694 | 1.474 | 0.106 | 1.5E-17 |
| 1 | CKAP4 | 0.119 | 0.143 | 0.88 | 0.684 | 0.364 | 4.6E-05 |
| 1 | CLASP1 | 0.279 | 0.36 | 2.394 | 1.894 | 0.339 | 0.00048 |
| 1 | CLASP2 | 0.208 | 0.264 | 1.761 | 1.306 | 0.431 | 0.00046 |
| 1 | CLINT1 | 0.281 | 0.399 | 2.361 | 2.142 | 0.134 | 4.2E-14 |
| 1 | CLIP1 | 0.171 | 0.24 | 1.318 | 1.229 | 0.1 | 1.7E-16 |
| 1 | CLIP4 | 0.136 | 0.172 | 0.984 | 0.797 | 0.304 | 9.3E-05 |
| 1 | CLK1 | 0.227 | 0.313 | 1.924 | 1.604 | 0.263 | 3.7E-10 |
| 1 | CLK4 | 0.12 | 0.144 | 0.88 | 0.68 | 0.372 | 7E-05 |
| 1 | CLN3 | 0.103 | 0.132 | 0.72 | 0.494 | 0.276 | 0.0001 |
| 1 | CMIP | 0.224 | 0.292 | 2.007 | 1.469 | 0.44 | 6.2E-05 |
| 1 | CNOT1 | 0.149 | 0.24 | 1.264 | 1.16 | 0.124 | 1.9E-22 |
| 1 | CNOT10 | 0.101 | 0.147 | 0.76 | 0.646 | 0.211 | 1.6E-11 |
| 1 | CNOT2 | 0.303 | 0.414 | 2.648 | 2.237 | 0.244 | 4.9E-09 |
| 1 | CNOT4 | 0.244 | 0.344 | 2.038 | 1.762 | 0.21 | 1.2E-12 |
| 1 | CNST | 0.144 | 0.214 | 1.08 | 0.994 | 0.119 | 6.8E-16 |
| 1 | COA1 | 0.134 | 0.181 | 1.016 | 0.834 | 0.286 | 1.4E-07 |
| 1 | COG4 | 0.224 | 0.291 | 1.909 | 1.44 | 0.397 | 7.4E-05 |
| 1 | COMMD1 | 0.118 | 0.182 | 0.941 | 0.842 | 0.176 | 2.4E-17 |
| 1 | COMMD10 | 0.139 | 0.192 | 1.141 | 0.904 | 0.334 | 3.9E-09 |
| 1 | COP1 | 0.261 | 0.33 | 2.197 | 1.71 | 0.361 | 0.00201 |
| 1 | COPA | 0.146 | 0.211 | 1.184 | 0.984 | 0.264 | 4.9E-13 |
| 1 | CPLANE1 | 0.144 | 0.21 | 1.177 | 1.009 | 0.222 | 2.9E-09 |
| 1 | CPQ | 0.148 | 0.237 | 1.294 | 1.192 | 0.12 | 4.8E-18 |
| 1 | CPSF6 | 0.144 | 0.212 | 1.209 | 0.968 | 0.322 | 6.5E-09 |
| 1 | CPT1A | 0.141 | 0.224 | 1.184 | 1.049 | 0.16 | 2.8E-16 |
| 1 | CRACR2A | 0.101 | 0.141 | 0.826 | 0.623 | 0.406 | 2.9E-08 |
| 1 | CRADD | 0.112 | 0.148 | 0.886 | 0.72 | 0.3 | 8.4E-10 |
| 1 | CRCP | 0.144 | 0.229 | 1.163 | 1.077 | 0.111 | 2.3E-23 |
| 1 | CREB1 | 0.164 | 0.228 | 1.24 | 1.119 | 0.16 | 7.8E-11 |
| 1 | CREBBP | 0.217 | 0.284 | 1.848 | 1.372 | 0.438 | 4.8E-05 |
| 1 | CRLF3 | 0.183 | 0.242 | 1.397 | 1.133 | 0.302 | 1.3E-06 |
| 1 | CRTC3 | 0.116 | 0.148 | 0.881 | 0.704 | 0.322 | 1.6E-07 |
| 1 | CRY1 | 0.147 | 0.216 | 1.494 | 1.081 | 0.467 | 1.5E-08 |
| 1 | CSGALNACT2 | 0.233 | 0.326 | 1.87 | 1.667 | 0.166 | 1.4E-12 |
| 1 | CSNK1D | 0.122 | 0.174 | 0.946 | 0.767 | 0.317 | 8.4E-11 |
| 1 | CSNK1G1 | 0.103 | 0.146 | 0.782 | 0.644 | 0.276 | 1.1E-09 |
| 1 | CSNK1G3 | 0.219 | 0.313 | 1.774 | 1.447 | 0.188 | 4.3E-14 |
| 1 | CTBP2 | 0.142 | 0.211 | 1.26 | 0.98 | 0.362 | 1.9E-09 |
| 1 | CTCF | 0.184 | 0.246 | 1.397 | 1.231 | 0.182 | 3E-11 |
| 1 | CTDSP1 | 0.104 | 0.148 | 0.723 | 0.644 | 0.144 | 1.1E-10 |
| 1 | CTDSPL2 | 0.194 | 0.274 | 1.416 | 1.31 | 0.211 | 8.4E-13 |
| 1 | CTNNB1 | 0.182 | 0.268 | 1.464 | 1.276 | 0.199 | 1.5E-16 |
| 1 | CTR9 | 0.18 | 0.248 | 1.343 | 1.222 | 0.137 | 2.6E-14 |
| 1 | CUL1 | 0.114 | 0.167 | 0.844 | 0.749 | 0.17 | 8.3E-12 |
| 1 | CUL2 | 0.129 | 0.186 | 0.936 | 0.844 | 0.149 | 2.1E-12 |
| 1 | CUL3 | 0.208 | 0.27 | 1.783 | 1.311 | 0.444 | 4.1E-05 |

| | | | | | | | |
|---|---|---|---|---|---|---|---|
| 1 | CUX1 | 0.144 | 0.197 | 1.168 | 0.911 | 0.348 | 2E-08 |
| 1 | CWF19L2 | 0.147 | 0.211 | 1.262 | 0.986 | 0.344 | 8E-08 |
| 1 | CYFIP2 | 0.217 | 0.314 | 1.792 | 1.49 | 0.173 | 9.7E-16 |
| 1 | CYLD | 0.381 | 0.4 | 3.437 | 3.247 | 0.119 | 2.9E-07 |
| 1 | CYP20A1 | 0.174 | 0.244 | 1.37 | 1.249 | 0.122 | 1.3E-14 |
| 1 | DAAM1 | 0.124 | 0.161 | 0.944 | 0.778 | 0.28 | 2.4E-05 |
| 1 | DCAF10 | 0.103 | 0.127 | 0.74 | 0.486 | 0.346 | 0.00148 |
| 1 | DCAF4 | 0.199 | 0.274 | 1.471 | 1.336 | 0.233 | 2.4E-10 |
| 1 | DCAF7 | 0.116 | 0.174 | 0.862 | 0.778 | 0.149 | 8.2E-14 |
| 1 | DCP1A | 0.102 | 0.144 | 0.802 | 0.623 | 0.364 | 9.1E-09 |
| 1 | DCTN4 | 0.112 | 0.164 | 0.871 | 0.736 | 0.243 | 2E-12 |
| 1 | DDHD1 | 0.182 | 0.249 | 1.41 | 1.243 | 0.269 | 1.1E-12 |
| 1 | DDX27 | 0.141 | 0.209 | 1.043 | 0.977 | 0.107 | 1.3E-14 |
| 1 | DDX42 | 0.109 | 0.16 | 0.842 | 0.73 | 0.206 | 1.9E-12 |
| 1 | DDX46 | 0.209 | 0.308 | 1.644 | 1.428 | 0.107 | 1.2E-18 |
| 1 | DDX6 | 0.437 | 0.482 | 4.346 | 3.801 | 0.193 | 0.00011 |
| 1 | DEF6 | 0.131 | 0.191 | 0.996 | 0.869 | 0.197 | 1.4E-13 |
| 1 | DENND1A | 0.128 | 0.164 | 1.008 | 0.748 | 0.412 | 0.00011 |
| 1 | DENND1B | 0.197 | 0.281 | 1.642 | 1.396 | 0.234 | 1.2E-13 |
| 1 | DENND4A | 0.272 | 0.364 | 2.496 | 1.983 | 0.332 | 4.8E-07 |
| 1 | DENND4C | 0.224 | 0.287 | 1.847 | 1.396 | 0.404 | 0.00074 |
| 1 | DENND6A | 0.122 | 0.183 | 0.908 | 0.829 | 0.131 | 3E-14 |
| 1 | DGKD | 0.276 | 0.344 | 2.364 | 1.843 | 0.349 | 0.00044 |
| 1 | DHX14 | 0.141 | 0.194 | 1.093 | 0.886 | 0.304 | 6.4E-09 |
| 1 | DHX8 | 0.118 | 0.174 | 0.874 | 0.764 | 0.192 | 2.5E-13 |
| 1 | DIAPH1 | 0.348 | 0.481 | 3.266 | 2.862 | 0.191 | 6E-11 |
| 1 | DIAPH2 | 0.138 | 0.213 | 1.203 | 1.04 | 0.194 | 7.1E-19 |
| 1 | DICER1 | 0.148 | 0.211 | 1.228 | 0.999 | 0.298 | 1.1E-07 |
| 1 | DIDO1 | 0.122 | 0.16 | 0.918 | 0.709 | 0.373 | 4.1E-06 |
| 1 | DIP2A | 0.363 | 0.468 | 3.467 | 2.846 | 0.321 | 0.00268 |
| 1 | DIP2B | 0.123 | 0.168 | 0.946 | 0.74 | 0.34 | 4.7E-08 |
| 1 | DIPK1A | 0.26 | 0.38 | 2.247 | 2.074 | 0.122 | 7.1E-19 |
| 1 | DLEU2 | 0.196 | 0.264 | 1.638 | 1.346 | 0.283 | 1.4E-08 |
| 1 | DLG1 | 0.147 | 0.207 | 1.236 | 0.988 | 0.324 | 4.7E-07 |
| 1 | DMXL1 | 0.149 | 0.234 | 1.204 | 1.12 | 0.104 | 1.9E-16 |
| 1 | DNAJB14 | 0.283 | 0.394 | 2.387 | 2.127 | 0.167 | 1.2E-11 |
| 1 | DNAJB6 | 0.379 | 0.422 | 3.447 | 3.248 | 0.127 | 4.9E-12 |
| 1 | DNAJC1 | 0.314 | 0.416 | 2.72 | 2.32 | 0.229 | 1.2E-06 |
| 1 | DNAJC3 | 0.214 | 0.327 | 1.767 | 1.638 | 0.109 | 7E-22 |
| 1 | DNAJC7 | 0.103 | 0.149 | 0.764 | 0.703 | 0.123 | 9.1E-16 |
| 1 | DNM2 | 0.179 | 0.229 | 1.47 | 1.047 | 0.476 | 0.00013 |
| 1 | DOCK10 | 0.462 | 0.494 | 4.798 | 4.173 | 0.201 | 0.00999 |
| 1 | DOCK11 | 0.243 | 0.344 | 2.146 | 1.78 | 0.269 | 2.6E-09 |
| 1 | DOCK2 | 0.284 | 0.393 | 2.448 | 2.082 | 0.292 | 3.3E-09 |
| 1 | DOCK4 | 0.129 | 0.173 | 1.236 | 0.904 | 0.441 | 2.2E-07 |
| 1 | DOCK8 | 0.467 | 0.61 | 4.884 | 4.434 | 0.14 | 1.9E-05 |
| 1 | DPYD | 0.471 | 0.44 | 4.108 | 3.647 | 0.486 | 1.7E-07 |
| 1 | DTD1 | 0.104 | 0.163 | 0.812 | 0.734 | 0.146 | 1.6E-17 |
| 1 | DTHD1 | 0.212 | 0.284 | 1.814 | 1.614 | 0.167 | 3.4E-10 |
| 1 | DUSP10 | 0.104 | 0.16 | 0.808 | 0.724 | 0.148 | 9.9E-16 |
| 1 | DYNC1H1 | 0.242 | 0.344 | 2.168 | 1.74 | 0.318 | 4.5E-08 |
| 1 | DYRK1A | 0.246 | 0.34 | 2.296 | 1.782 | 0.364 | 4.7E-08 |
| 1 | EHBP1 | 0.104 | 0.138 | 0.81 | 0.644 | 0.328 | 8.1E-06 |
| 1 | EHMT1 | 0.248 | 0.341 | 2.076 | 1.697 | 0.291 | 1.4E-09 |

| | | | | | | | |
|---|---|---|---|---|---|---|---|
| 1 | EIF2S3 | 0.123 | 0.193 | 0.934 | 0.864 | 0.113 | 4.2E-19 |
| 1 | EIF4G3 | 0.247 | 0.339 | 2.393 | 1.816 | 0.398 | 8.4E-06 |
| 1 | ELF2 | 0.27 | 0.372 | 2.243 | 1.944 | 0.206 | 7.6E-10 |
| 1 | ELK4 | 0.104 | 0.146 | 0.771 | 0.696 | 0.148 | 9.2E-14 |
| 1 | ELL2 | 0.106 | 0.163 | 1.071 | 0.82 | 0.384 | 1.5E-14 |
| 1 | ELMO1 | 0.214 | 0.277 | 2.037 | 1.424 | 0.416 | 0.00016 |
| 1 | ELOVL4 | 0.123 | 0.181 | 0.967 | 0.824 | 0.23 | 9.3E-14 |
| 1 | ELP4 | 0.129 | 0.172 | 1.042 | 0.839 | 0.312 | 2.7E-07 |
| 1 | EML4 | 0.411 | 0.422 | 3.994 | 3.448 | 0.208 | 0.00772 |
| 1 | EMSY | 0.103 | 0.141 | 0.769 | 0.626 | 0.297 | 1.2E-07 |
| 1 | ENTPD4 | 0.119 | 0.169 | 0.907 | 0.749 | 0.276 | 4.4E-10 |
| 1 | EP300 | 0.188 | 0.264 | 1.433 | 1.266 | 0.276 | 4.1E-11 |
| 1 | EPB41 | 0.428 | 0.642 | 4.76 | 4.644 | 0.307 | 0.00888 |
| 1 | EPC2 | 0.164 | 0.231 | 1.221 | 1.08 | 0.177 | 6.1E-12 |
| 1 | EPS14 | 0.323 | 0.449 | 2.83 | 2.461 | 0.144 | 3E-12 |
| 1 | ERAP2 | 0.211 | 0.294 | 1.677 | 1.446 | 0.117 | 2.1E-13 |
| 1 | ERBIN | 0.33 | 0.446 | 2.893 | 2.499 | 0.211 | 7.9E-08 |
| 1 | ERC1 | 0.144 | 0.199 | 1.187 | 0.947 | 0.326 | 6.1E-09 |
| 1 | ERCC4 | 0.161 | 0.231 | 1.207 | 1.116 | 0.113 | 3.7E-14 |
| 1 | ERCC6L2 | 0.114 | 0.144 | 0.901 | 0.712 | 0.341 | 4.5E-07 |
| 1 | ERGIC3 | 0.106 | 0.163 | 0.742 | 0.702 | 0.1 | 2.4E-15 |
| 1 | ERN1 | 0.143 | 0.2 | 1.16 | 0.927 | 0.323 | 7.4E-06 |
| 1 | ERO1B | 0.138 | 0.203 | 1.048 | 0.949 | 0.147 | 2E-14 |
| 1 | ESCO1 | 0.127 | 0.173 | 0.969 | 0.797 | 0.282 | 4.5E-08 |
| 1 | ESYT2 | 0.161 | 0.237 | 1.74 | 1.246 | 0.471 | 1.3E-13 |
| 1 | ETV6 | 0.102 | 0.142 | 0.837 | 0.647 | 0.371 | 3.8E-08 |
| 1 | EXOC2 | 0.142 | 0.224 | 1.143 | 1.064 | 0.102 | 3.7E-16 |
| 1 | EXOC6B | 0.132 | 0.164 | 1.071 | 0.764 | 0.486 | 0.00119 |
| 1 | EZH1 | 0.149 | 0.182 | 1.092 | 0.827 | 0.4 | 0.00274 |
| 1 | F2R | 0.316 | 0.411 | 2.842 | 2.449 | 0.147 | 1.2E-06 |
| 1 | FAF1 | 0.211 | 0.264 | 1.904 | 1.311 | 0.439 | 0.00403 |
| 1 | FAF2 | 0.102 | 0.141 | 0.74 | 0.604 | 0.314 | 4.9E-08 |
| 1 | FAM107B | 0.243 | 0.348 | 2.047 | 1.844 | 0.147 | 2.9E-14 |
| 1 | FAM120B | 0.143 | 0.204 | 1.078 | 0.93 | 0.212 | 2.3E-12 |
| 1 | FAM13B | 0.206 | 0.284 | 1.627 | 1.404 | 0.213 | 3E-11 |
| 1 | FAM160B1 | 0.114 | 0.171 | 0.833 | 0.746 | 0.14 | 9.2E-14 |
| 1 | FAM214A | 0.194 | 0.248 | 1.449 | 1.274 | 0.281 | 1.8E-07 |
| 1 | FAM91A1 | 0.132 | 0.184 | 1.043 | 0.839 | 0.328 | 1.8E-09 |
| 1 | FAR1 | 0.174 | 0.248 | 1.434 | 1.191 | 0.269 | 4.3E-12 |
| 1 | FBXL17 | 0.244 | 0.319 | 2.329 | 1.712 | 0.444 | 0.00489 |
| 1 | FBXO11 | 0.219 | 0.306 | 1.884 | 1.449 | 0.274 | 4.2E-11 |
| 1 | FBXO21 | 0.107 | 0.149 | 0.779 | 0.698 | 0.148 | 2.5E-13 |
| 1 | FBXO33 | 0.137 | 0.192 | 1.214 | 0.911 | 0.414 | 8.4E-10 |
| 1 | FBXW11 | 0.198 | 0.278 | 1.44 | 1.338 | 0.203 | 4.4E-12 |
| 1 | FBXW7 | 0.277 | 0.373 | 2.423 | 2.011 | 0.328 | 3.8E-07 |
| 1 | FCHO2 | 0.104 | 0.132 | 0.776 | 0.603 | 0.363 | 0.0004 |
| 1 | FCHSD2 | 0.164 | 0.221 | 1.338 | 1.084 | 0.302 | 1.4E-08 |
| 1 | FGD3 | 0.239 | 0.301 | 1.908 | 1.49 | 0.346 | 0.00343 |
| 1 | FIP1L1 | 0.177 | 0.264 | 1.364 | 1.246 | 0.119 | 1.4E-18 |
| 1 | FKBP8 | 0.114 | 0.171 | 0.81 | 0.743 | 0.104 | 2E-14 |
| 1 | FLI1 | 0.182 | 0.244 | 1.44 | 1.219 | 0.241 | 3.1E-11 |
| 1 | FLNA | 0.142 | 0.224 | 1.247 | 1.162 | 0.102 | 9.4E-23 |
| 1 | FMNL1 | 0.121 | 0.174 | 0.911 | 0.79 | 0.204 | 7.1E-12 |
| 1 | FMR1 | 0.103 | 0.144 | 0.797 | 0.689 | 0.21 | 3.7E-13 |

| | | | | | | | |
|---|---|---|---|---|---|---|---|
| 1 | FNBP1 | 0.387 | 0.434 | 3.662 | 3.28 | 0.149 | 4.2E-10 |
| 1 | FNBP4 | 0.261 | 0.372 | 2.136 | 1.917 | 0.146 | 1.2E-14 |
| 1 | FNDC3A | 0.194 | 0.29 | 1.602 | 1.463 | 0.131 | 3.1E-19 |
| 1 | FNDC3B | 0.297 | 0.404 | 3.006 | 2.298 | 0.387 | 1.7E-07 |
| 1 | FNIP1 | 0.174 | 0.233 | 1.396 | 1.102 | 0.341 | 2E-07 |
| 1 | FOSL2 | 0.102 | 0.148 | 0.817 | 0.733 | 0.146 | 1.5E-16 |
| 1 | FOXJ3 | 0.193 | 0.273 | 1.493 | 1.311 | 0.188 | 4.9E-13 |
| 1 | FOXN2 | 0.147 | 0.237 | 1.189 | 1.11 | 0.1 | 8E-19 |
| 1 | FOXO3 | 0.114 | 0.147 | 0.878 | 0.641 | 0.431 | 8.8E-05 |
| 1 | FOXP1 | 0.443 | 0.428 | 4.648 | 3.498 | 0.413 | 0.00048 |
| 1 | FP236383.3 | 0.61 | 0.492 | 12.42 | 4.431 | 1.179 | 4E-100 |
| 1 | FRYL | 0.324 | 0.424 | 2.967 | 2.342 | 0.334 | 0.00036 |
| 1 | FUBP1 | 0.209 | 0.307 | 1.717 | 1.424 | 0.171 | 9.8E-17 |
| 1 | FYN | 0.707 | 0.814 | 10.648 | 8.434 | 0.336 | 4.6E-14 |
| 1 | G2E3 | 0.108 | 0.148 | 0.811 | 0.712 | 0.189 | 4.4E-12 |
| 1 | GAB3 | 0.134 | 0.178 | 1.046 | 0.812 | 0.364 | 2.2E-06 |
| 1 | GABARAPL1 | 0.177 | 0.26 | 1.374 | 1.26 | 0.124 | 3.3E-17 |
| 1 | GAK | 0.101 | 0.141 | 0.801 | 0.614 | 0.382 | 2E-08 |
| 1 | GALNT1 | 0.122 | 0.172 | 0.967 | 0.772 | 0.324 | 2.8E-10 |
| 1 | GAPVD1 | 0.112 | 0.144 | 0.833 | 0.681 | 0.291 | 2.4E-08 |
| 1 | GARS-DT | 0.106 | 0.141 | 0.833 | 0.682 | 0.289 | 8E-10 |
| 1 | GATAD2A | 0.149 | 0.207 | 1.24 | 0.934 | 0.419 | 2.4E-05 |
| 1 | GATAD2B | 0.22 | 0.281 | 1.818 | 1.368 | 0.41 | 0.00033 |
| 1 | GBF1 | 0.128 | 0.182 | 1.014 | 0.828 | 0.294 | 4.4E-11 |
| 1 | GCC2 | 0.304 | 0.404 | 2.687 | 2.316 | 0.214 | 1.7E-07 |
| 1 | GCNT1 | 0.137 | 0.168 | 1.101 | 0.849 | 0.376 | 0.0015 |
| 1 | GFOD1 | 0.104 | 0.14 | 0.844 | 0.638 | 0.422 | 1.5E-06 |
| 1 | GFPT1 | 0.102 | 0.142 | 0.74 | 0.683 | 0.136 | 4.8E-13 |
| 1 | GGA2 | 0.197 | 0.242 | 1.412 | 1.266 | 0.246 | 1.7E-05 |
| 1 | GIGYF2 | 0.137 | 0.183 | 1.044 | 0.83 | 0.331 | 3.4E-07 |
| 1 | GIMAP4 | 0.197 | 0.249 | 1.428 | 1.399 | 0.127 | 1.1E-08 |
| 1 | GIT2 | 0.138 | 0.187 | 1.024 | 0.861 | 0.24 | 2.8E-08 |
| 1 | GK4 | 0.247 | 0.332 | 2.204 | 1.766 | 0.32 | 0.00013 |
| 1 | GLG1 | 0.26 | 0.361 | 2.169 | 1.86 | 0.222 | 8E-11 |
| 1 | GMCL1 | 0.118 | 0.171 | 0.87 | 0.766 | 0.183 | 3.3E-12 |
| 1 | GMDS | 0.123 | 0.178 | 1.036 | 0.827 | 0.324 | 3.4E-12 |
| 1 | GMDS-DT | 0.169 | 0.217 | 1.389 | 1.113 | 0.32 | 1.6E-05 |
| 1 | GNA13 | 0.167 | 0.243 | 1.384 | 1.144 | 0.261 | 4.3E-14 |
| 1 | GNB1 | 0.139 | 0.181 | 1.168 | 0.807 | 0.434 | 2.4E-05 |
| 1 | GOLGA4 | 0.334 | 0.444 | 3.064 | 2.661 | 0.203 | 3.4E-08 |
| 1 | GOLGA8A | 0.104 | 0.141 | 0.802 | 0.614 | 0.383 | 1.4E-06 |
| 1 | GOLGA8B | 0.149 | 0.204 | 1.122 | 0.94 | 0.24 | 1.1E-09 |
| 1 | GOLPH3 | 0.101 | 0.147 | 0.77 | 0.636 | 0.277 | 4.1E-11 |
| 1 | GON4L | 0.131 | 0.187 | 0.999 | 0.84 | 0.232 | 2.7E-11 |
| 1 | GOSR1 | 0.11 | 0.164 | 0.82 | 0.717 | 0.194 | 2.7E-13 |
| 1 | GPATCH2L | 0.137 | 0.198 | 1 | 0.917 | 0.124 | 1.4E-13 |
| 1 | GPATCH8 | 0.298 | 0.412 | 2.644 | 2.286 | 0.216 | 7E-10 |
| 1 | GPBP1L1 | 0.128 | 0.171 | 0.981 | 0.766 | 0.347 | 3.4E-07 |
| 1 | GPCPD1 | 0.306 | 0.413 | 2.649 | 2.246 | 0.237 | 1.2E-07 |
| 1 | GPR174 | 0.139 | 0.2 | 1.048 | 0.919 | 0.191 | 1.1E-12 |
| 1 | GPRIN3 | 0.143 | 0.23 | 1.296 | 1.132 | 0.196 | 1.8E-17 |
| 1 | GRAP2 | 0.107 | 0.143 | 0.804 | 0.712 | 0.174 | 4.4E-11 |
| 1 | GRIPAP1 | 0.107 | 0.161 | 0.776 | 0.714 | 0.118 | 1.9E-14 |
| 1 | GRK4 | 0.116 | 0.161 | 0.969 | 0.724 | 0.419 | 7.9E-09 |

| | | | | | | | |
|---|---|---|---|---|---|---|---|
| 1 | GSAP | 0.233 | 0.346 | 2.12 | 1.971 | 0.104 | 4.6E-24 |
| 1 | GSE1 | 0.102 | 0.14 | 0.803 | 0.627 | 0.347 | 4.7E-08 |
| 1 | GSK3B | 0.36 | 0.493 | 3.262 | 2.824 | 0.208 | 7.3E-08 |
| 1 | GTDC1 | 0.162 | 0.208 | 1.29 | 1.009 | 0.344 | 2.7E-05 |
| 1 | GTF2E2 | 0.142 | 0.232 | 1.143 | 1.066 | 0.102 | 2.5E-19 |
| 1 | GTF2I | 0.238 | 0.332 | 1.917 | 1.726 | 0.141 | 8.9E-13 |
| 1 | H2AFV | 0.241 | 0.367 | 2.046 | 1.903 | 0.112 | 7.5E-18 |
| 1 | HBP1 | 0.167 | 0.236 | 1.314 | 1.097 | 0.261 | 3.5E-11 |
| 1 | HEATR4B | 0.139 | 0.193 | 1.022 | 0.886 | 0.206 | 4.3E-10 |
| 1 | HECTD1 | 0.203 | 0.277 | 1.484 | 1.347 | 0.234 | 2.5E-09 |
| 1 | HECTD4 | 0.127 | 0.167 | 0.972 | 0.749 | 0.347 | 2.1E-06 |
| 1 | HELZ | 0.3 | 0.388 | 2.831 | 2.126 | 0.413 | 0.00264 |
| 1 | HERC2 | 0.244 | 0.306 | 1.983 | 1.419 | 0.384 | 0.00485 |
| 1 | HERC4 | 0.188 | 0.247 | 1.476 | 1.262 | 0.226 | 4.4E-10 |
| 1 | HEXA | 0.109 | 0.149 | 0.832 | 0.712 | 0.224 | 6.7E-12 |
| 1 | HIPK1 | 0.162 | 0.241 | 1.334 | 1.2 | 0.144 | 1.4E-21 |
| 1 | HIPK2 | 0.24 | 0.32 | 2.187 | 1.644 | 0.411 | 0.00067 |
| 1 | HIPK3 | 0.148 | 0.227 | 1.184 | 1.06 | 0.149 | 1.2E-13 |
| 1 | HIVEP2 | 0.148 | 0.207 | 1.296 | 0.989 | 0.39 | 7.3E-06 |
| 1 | HIVEP3 | 0.2 | 0.264 | 1.824 | 1.337 | 0.449 | 1.5E-06 |
| 1 | HNRNPH1 | 0.214 | 0.314 | 1.796 | 1.482 | 0.183 | 1.1E-16 |
| 1 | HNRNPH3 | 0.121 | 0.184 | 0.9 | 0.826 | 0.124 | 2.4E-16 |
| 1 | HPS3 | 0.141 | 0.21 | 1.107 | 0.982 | 0.173 | 4.9E-11 |
| 1 | HTT | 0.141 | 0.224 | 1.164 | 1.028 | 0.179 | 1.4E-14 |
| 1 | HUWE1 | 0.178 | 0.274 | 1.44 | 1.324 | 0.121 | 9.9E-23 |
| 1 | IFNAR2 | 0.187 | 0.243 | 1.431 | 1.219 | 0.232 | 4.6E-09 |
| 1 | IGF2R | 0.171 | 0.24 | 1.492 | 1.142 | 0.372 | 9.2E-11 |
| 1 | IKZF2 | 0.11 | 0.161 | 0.947 | 0.77 | 0.299 | 4.1E-12 |
| 1 | IL18RAP | 0.139 | 0.201 | 1.124 | 0.944 | 0.236 | 1.8E-12 |
| 1 | ILF3 | 0.194 | 0.288 | 1.493 | 1.4 | 0.187 | 8.7E-18 |
| 1 | ILK | 0.112 | 0.162 | 0.778 | 0.706 | 0.14 | 6.1E-12 |
| 1 | ILKAP | 0.134 | 0.189 | 1.046 | 0.863 | 0.277 | 1.7E-10 |
| 1 | ILRUN | 0.133 | 0.198 | 0.98 | 0.893 | 0.133 | 7.7E-14 |
| 1 | IMMP2L | 0.138 | 0.191 | 1.148 | 0.991 | 0.224 | 3.9E-10 |
| 1 | INO80D | 0.184 | 0.264 | 1.403 | 1.284 | 0.226 | 4E-13 |
| 1 | INPP4A | 0.197 | 0.249 | 1.642 | 1.182 | 0.474 | 0.00197 |
| 1 | INPP4D | 0.307 | 0.397 | 2.8 | 2.174 | 0.364 | 0.00089 |
| 1 | INTS6 | 0.132 | 0.183 | 1.003 | 0.84 | 0.244 | 1.8E-09 |
| 1 | INVS | 0.139 | 0.177 | 1.163 | 0.837 | 0.474 | 0.00018 |
| 1 | IP6K2 | 0.114 | 0.171 | 0.876 | 0.767 | 0.191 | 3.3E-14 |
| 1 | IPO7 | 0.108 | 0.149 | 0.796 | 0.667 | 0.244 | 4E-08 |
| 1 | IQGAP2 | 0.464 | 0.622 | 4.88 | 4.382 | 0.144 | 9.3E-08 |
| 1 | IREB2 | 0.126 | 0.183 | 0.942 | 0.81 | 0.218 | 1.4E-12 |
| 1 | IRF3 | 0.109 | 0.144 | 0.799 | 0.646 | 0.306 | 1.4E-06 |
| 1 | IST1 | 0.143 | 0.201 | 1.088 | 0.911 | 0.247 | 1.5E-10 |
| 1 | ITCH | 0.341 | 0.439 | 3.118 | 2.447 | 0.344 | 0.00775 |
| 1 | ITFG1 | 0.224 | 0.297 | 1.932 | 1.436 | 0.332 | 2.1E-06 |
| 1 | ITGAE | 0.181 | 0.223 | 1.404 | 1.133 | 0.31 | 0.00109 |
| 1 | ITGAL | 0.323 | 0.432 | 2.966 | 2.442 | 0.28 | 1.1E-06 |
| 1 | ITK | 0.23 | 0.292 | 1.89 | 1.488 | 0.344 | 0.00032 |
| 1 | ITSN2 | 0.308 | 0.434 | 2.72 | 2.462 | 0.144 | 2.1E-13 |
| 1 | IVNS1ABP | 0.143 | 0.214 | 1.131 | 1.022 | 0.146 | 4.3E-17 |
| 1 | JAKMIP2 | 0.117 | 0.169 | 0.904 | 0.794 | 0.187 | 4.4E-12 |
| 1 | JARID2 | 0.228 | 0.289 | 1.966 | 1.41 | 0.48 | 0.00323 |

| | | | | | | | |
|---|---|---|---|---|---|---|---|
| 1 | JAZF1 | 0.429 | 0.626 | 4.849 | 4.64 | 0.334 | 0.00027 |
| 1 | JMJD1C | 0.349 | 0.484 | 3.277 | 2.962 | 0.146 | 4.4E-12 |
| 1 | JPX | 0.122 | 0.173 | 0.938 | 0.803 | 0.223 | 1.1E-10 |
| 1 | KANSL2 | 0.108 | 0.173 | 0.834 | 0.774 | 0.108 | 6.4E-19 |
| 1 | KAT2B | 0.161 | 0.199 | 1.339 | 0.926 | 0.432 | 0.00363 |
| 1 | KAT6B | 0.332 | 0.429 | 3.091 | 2.463 | 0.327 | 0.00124 |
| 1 | KCTD20 | 0.144 | 0.224 | 1.164 | 1.047 | 0.144 | 3.8E-14 |
| 1 | KDM2A | 0.212 | 0.286 | 1.804 | 1.417 | 0.349 | 1E-07 |
| 1 | KDM3A | 0.11 | 0.14 | 0.849 | 0.629 | 0.433 | 0.00015 |
| 1 | KDM3B | 0.109 | 0.14 | 0.816 | 0.674 | 0.273 | 2.2E-08 |
| 1 | KDM4A | 0.289 | 0.386 | 2.467 | 2.089 | 0.24 | 4.1E-07 |
| 1 | KDM4B | 0.149 | 0.203 | 1.197 | 0.961 | 0.316 | 4.2E-05 |
| 1 | KDM4C | 0.243 | 0.319 | 2.12 | 1.486 | 0.419 | 0.0001 |
| 1 | KDM6A | 0.262 | 0.364 | 2.489 | 1.947 | 0.347 | 2.1E-10 |
| 1 | KDM7A | 0.181 | 0.246 | 1.413 | 1.216 | 0.217 | 1.7E-09 |
| 1 | KIAA0232 | 0.139 | 0.2 | 1.02 | 0.924 | 0.142 | 3.7E-13 |
| 1 | KIAA0486 | 0.11 | 0.141 | 0.821 | 0.694 | 0.241 | 1.9E-08 |
| 1 | KIAA1109 | 0.342 | 0.447 | 3.17 | 2.64 | 0.264 | 6.8E-06 |
| 1 | KIAA2026 | 0.221 | 0.286 | 1.787 | 1.423 | 0.329 | 2.4E-05 |
| 1 | KIDINS220 | 0.164 | 0.24 | 1.244 | 1.126 | 0.146 | 4.4E-14 |
| 1 | KIF13B | 0.164 | 0.246 | 1.303 | 1.146 | 0.172 | 2.4E-17 |
| 1 | KIFAP3 | 0.184 | 0.268 | 1.411 | 1.288 | 0.132 | 1.8E-14 |
| 1 | KIN | 0.114 | 0.161 | 0.79 | 0.718 | 0.138 | 2.1E-10 |
| 1 | KLC1 | 0.136 | 0.192 | 1.044 | 0.864 | 0.274 | 7.4E-11 |
| 1 | KLF13 | 0.141 | 0.204 | 1.094 | 0.914 | 0.249 | 1.2E-12 |
| 1 | KLHL2 | 0.114 | 0.146 | 0.873 | 0.649 | 0.404 | 0.00013 |
| 1 | KLHL20 | 0.109 | 0.144 | 0.844 | 0.702 | 0.284 | 1.9E-09 |
| 1 | KMT2A | 0.263 | 0.347 | 2.229 | 1.871 | 0.243 | 6.7E-07 |
| 1 | KMT4B | 0.169 | 0.223 | 1.278 | 1.068 | 0.248 | 1.2E-06 |
| 1 | KPNA3 | 0.146 | 0.213 | 1.131 | 0.972 | 0.219 | 2.2E-13 |
| 1 | KPNA4 | 0.208 | 0.313 | 1.64 | 1.434 | 0.104 | 8E-21 |
| 1 | KRAS | 0.244 | 0.336 | 1.93 | 1.733 | 0.144 | 8.1E-11 |
| 1 | KRCC1 | 0.112 | 0.146 | 0.769 | 0.709 | 0.118 | 1.4E-09 |
| 1 | KRIT1 | 0.124 | 0.179 | 0.916 | 0.82 | 0.16 | 5E-12 |
| 1 | L3MBTL4 | 0.141 | 0.179 | 1.167 | 0.867 | 0.43 | 0.00014 |
| 1 | LARP7 | 0.112 | 0.17 | 0.877 | 0.8 | 0.132 | 1.4E-14 |
| 1 | LASP1 | 0.137 | 0.184 | 1.063 | 0.83 | 0.347 | 1.7E-07 |
| 1 | LATS2 | 0.137 | 0.188 | 1.067 | 0.868 | 0.297 | 1.1E-08 |
| 1 | LBR | 0.347 | 0.492 | 3.213 | 2.937 | 0.129 | 4.9E-11 |
| 1 | LCOR | 0.218 | 0.298 | 1.798 | 1.468 | 0.292 | 4.8E-09 |
| 1 | LCORL | 0.14 | 0.203 | 1.184 | 0.974 | 0.282 | 2.9E-08 |
| 1 | LCP2 | 0.283 | 0.382 | 2.407 | 2.078 | 0.212 | 7.2E-09 |
| 1 | LDLRAD4 | 0.14 | 0.171 | 1.099 | 0.846 | 0.377 | 0.00122 |
| 1 | LINC00299 | 0.208 | 0.317 | 1.784 | 1.64 | 0.113 | 9.4E-23 |
| 1 | LINC00623 | 0.267 | 0.348 | 2.204 | 1.949 | 0.17 | 3.4E-09 |
| 1 | LINC00861 | 0.143 | 0.226 | 1.224 | 1.094 | 0.161 | 4.2E-16 |
| 1 | LINC01138 | 0.389 | 0.416 | 3.649 | 3.306 | 0.142 | 2.2E-07 |
| 1 | LINC01278 | 0.118 | 0.172 | 0.989 | 0.794 | 0.314 | 2.3E-12 |
| 1 | LINC01478 | 0.104 | 0.14 | 0.824 | 0.661 | 0.319 | 3E-10 |
| 1 | LINC01619 | 0.308 | 0.394 | 3.013 | 2.307 | 0.384 | 0.00344 |
| 1 | LINC02246 | 0.214 | 0.308 | 1.72 | 1.469 | 0.132 | 1.5E-14 |
| 1 | LINC02328 | 0.113 | 0.142 | 0.891 | 0.726 | 0.294 | 1.8E-07 |
| 1 | LINGO2 | 0.334 | 0.343 | 4.04 | 2.81 | 0.424 | 0.0004 |
| 1 | LMBR1 | 0.168 | 0.232 | 1.363 | 1.101 | 0.308 | 8.1E-10 |

| | | | | | | | |
|---|---|---|---|---|---|---|---|
| 1 | LMBRD1 | 0.208 | 0.284 | 1.612 | 1.387 | 0.216 | 1.2E-09 |
| 1 | LONP2 | 0.161 | 0.244 | 1.23 | 1.138 | 0.112 | 1.4E-19 |
| 1 | LPCAT1 | 0.223 | 0.321 | 1.87 | 1.664 | 0.168 | 2.5E-14 |
| 1 | LPGAT1 | 0.274 | 0.366 | 2.248 | 1.944 | 0.201 | 1E-07 |
| 1 | LPP | 0.301 | 0.419 | 3.086 | 2.363 | 0.384 | 1.2E-09 |
| 1 | LPXN | 0.146 | 0.227 | 1.141 | 1.044 | 0.113 | 3.4E-14 |
| 1 | LRBA | 0.432 | 0.446 | 4.444 | 3.717 | 0.248 | 0.00792 |
| 1 | LRCH1 | 0.106 | 0.144 | 0.802 | 0.689 | 0.22 | 9.2E-12 |
| 1 | LRCH3 | 0.164 | 0.207 | 1.27 | 0.966 | 0.394 | 0.00084 |
| 1 | LRMP | 0.167 | 0.233 | 1.267 | 1.096 | 0.209 | 6.3E-11 |
| 1 | LRPPRC | 0.13 | 0.191 | 0.974 | 0.842 | 0.193 | 1E-13 |
| 1 | LRRFIP2 | 0.131 | 0.198 | 1.009 | 0.894 | 0.173 | 1.6E-14 |
| 1 | LSM14A | 0.363 | 0.404 | 3.184 | 2.946 | 0.113 | 3.4E-11 |
| 1 | LUC7L | 0.163 | 0.214 | 1.26 | 0.974 | 0.371 | 2.2E-06 |
| 1 | LUC7L2 | 0.292 | 0.408 | 2.446 | 2.179 | 0.23 | 8.6E-11 |
| 1 | LUC7L3 | 0.288 | 0.364 | 2.473 | 1.912 | 0.371 | 0.00731 |
| 1 | LUZP1 | 0.134 | 0.17 | 1.009 | 0.802 | 0.331 | 0.0001 |
| 1 | LYN | 0.332 | 0.442 | 3.242 | 2.686 | 0.271 | 9.4E-09 |
| 1 | MACO1 | 0.104 | 0.147 | 0.811 | 0.716 | 0.18 | 2.3E-13 |
| 1 | MALAT1 | 1 | 1 | 776.904 | 697.45 | 0.146 | 1E-47 |
| 1 | MALT1 | 0.112 | 0.141 | 0.844 | 0.667 | 0.347 | 3E-07 |
| 1 | MAN1A1 | 0.2 | 0.293 | 1.667 | 1.482 | 0.17 | 6.4E-17 |
| 1 | MAN1A2 | 0.247 | 0.343 | 2.168 | 1.843 | 0.234 | 4.2E-10 |
| 1 | MAN2A1 | 0.142 | 0.194 | 1.139 | 0.906 | 0.33 | 1.3E-08 |
| 1 | MANBA | 0.108 | 0.138 | 0.788 | 0.614 | 0.36 | 7.2E-05 |
| 1 | MAP2K4 | 0.149 | 0.211 | 1.231 | 0.973 | 0.339 | 8.1E-07 |
| 1 | MAP3K2 | 0.282 | 0.384 | 2.383 | 2.068 | 0.204 | 1.9E-09 |
| 1 | MAP3K4 | 0.137 | 0.171 | 1.024 | 0.784 | 0.384 | 0.00043 |
| 1 | MAP4K4 | 0.197 | 0.271 | 1.474 | 1.331 | 0.242 | 4.8E-10 |
| 1 | MAP4K4 | 0.104 | 0.134 | 0.818 | 0.609 | 0.424 | 8.5E-05 |
| 1 | MAPK1 | 0.378 | 0.487 | 3.402 | 2.871 | 0.244 | 0.00991 |
| 1 | MAPK14 | 0.144 | 0.183 | 1.119 | 0.838 | 0.418 | 0.00013 |
| 1 | MAPK8 | 0.144 | 0.189 | 1.222 | 0.904 | 0.434 | 4.1E-06 |
| 1 | MAPKAP1 | 0.161 | 0.233 | 1.241 | 1.087 | 0.203 | 4.8E-14 |
| 1 | MAPRE2 | 0.336 | 0.481 | 3.308 | 3.044 | 0.12 | 2.8E-19 |
| 1 | MARF1 | 0.12 | 0.141 | 0.873 | 0.669 | 0.384 | 0.00026 |
| 1 | MARK3 | 0.187 | 0.244 | 1.407 | 1.208 | 0.319 | 1E-08 |
| 1 | MAX | 0.144 | 0.227 | 1.142 | 1.043 | 0.117 | 2.3E-21 |
| 1 | MBD4 | 0.332 | 0.43 | 3.141 | 2.497 | 0.274 | 0.00014 |
| 1 | MBNL2 | 0.178 | 0.242 | 1.387 | 1.141 | 0.268 | 3.4E-09 |
| 1 | MBNL3 | 0.114 | 0.164 | 0.864 | 0.749 | 0.19 | 1.4E-11 |
| 1 | MCM9 | 0.13 | 0.177 | 1.018 | 0.81 | 0.329 | 8.7E-08 |
| 1 | MCPH1 | 0.148 | 0.228 | 1.241 | 1.094 | 0.182 | 4.2E-14 |
| 1 | MCTP2 | 0.484 | 0.473 | 4.228 | 3.76 | 0.474 | 7.6E-06 |
| 1 | MDM2 | 0.141 | 0.194 | 1.107 | 0.89 | 0.314 | 4.4E-09 |
| 1 | MDM4 | 0.147 | 0.233 | 1.2 | 1.093 | 0.134 | 4.4E-17 |
| 1 | MECP2 | 0.191 | 0.244 | 1.472 | 1.231 | 0.342 | 4.5E-07 |
| 1 | MED13 | 0.176 | 0.234 | 1.368 | 1.096 | 0.32 | 3.7E-07 |
| 1 | MED14 | 0.146 | 0.234 | 1.207 | 1.107 | 0.124 | 4.8E-18 |
| 1 | MED14 | 0.106 | 0.141 | 0.794 | 0.676 | 0.234 | 2.8E-10 |
| 1 | MEGF9 | 0.124 | 0.174 | 0.922 | 0.819 | 0.172 | 2.8E-10 |
| 1 | METAP2 | 0.121 | 0.172 | 0.914 | 0.776 | 0.238 | 6.4E-11 |
| 1 | METTL14 | 0.117 | 0.144 | 0.909 | 0.699 | 0.379 | 8.7E-06 |
| 1 | MFSD6 | 0.104 | 0.144 | 0.764 | 0.679 | 0.173 | 1.1E-12 |

| | | | | | | | |
|---|---|---|---|---|---|---|---|
| 1 | MGA | 0.111 | 0.146 | 0.817 | 0.66 | 0.308 | 2.1E-06 |
| 1 | MGMT | 0.114 | 0.161 | 0.861 | 0.718 | 0.26 | 2.8E-09 |
| 1 | MIA3 | 0.183 | 0.268 | 1.444 | 1.321 | 0.129 | 2.4E-16 |
| 1 | MICU2 | 0.187 | 0.269 | 1.438 | 1.327 | 0.213 | 8.2E-14 |
| 1 | MIER1 | 0.298 | 0.433 | 2.449 | 2.362 | 0.11 | 2.4E-17 |
| 1 | MIGA1 | 0.123 | 0.174 | 0.91 | 0.777 | 0.228 | 2.8E-10 |
| 1 | MINDY2 | 0.136 | 0.183 | 1.033 | 0.84 | 0.299 | 1.6E-07 |
| 1 | MINDY3 | 0.104 | 0.144 | 0.79 | 0.679 | 0.219 | 3.1E-12 |
| 1 | MIR181A2HG | 0.149 | 0.186 | 1.142 | 0.973 | 0.244 | 0.00013 |
| 1 | MITD1 | 0.12 | 0.169 | 0.867 | 0.76 | 0.189 | 7.4E-10 |
| 1 | MKLN1 | 0.243 | 0.346 | 2.129 | 1.768 | 0.268 | 2.8E-09 |
| 1 | MNAT1 | 0.184 | 0.242 | 1.484 | 1.209 | 0.296 | 4.9E-09 |
| 1 | MON2 | 0.241 | 0.339 | 2.141 | 1.719 | 0.324 | 2.5E-07 |
| 1 | MPHOSPH8 | 0.249 | 0.34 | 2.116 | 1.833 | 0.207 | 4.9E-09 |
| 1 | MPHOSPH9 | 0.113 | 0.147 | 0.848 | 0.698 | 0.282 | 7.9E-09 |
| 1 | MPP7 | 0.134 | 0.177 | 1.024 | 0.809 | 0.341 | 1E-06 |
| 1 | MRE11 | 0.108 | 0.141 | 0.786 | 0.664 | 0.244 | 2.9E-09 |
| 1 | MRPS4 | 0.117 | 0.164 | 0.809 | 0.728 | 0.143 | 4.4E-10 |
| 1 | MSH3 | 0.112 | 0.166 | 0.869 | 0.77 | 0.174 | 1.2E-13 |
| 1 | MSI2 | 0.187 | 0.236 | 1.403 | 1.124 | 0.418 | 0.00023 |
| 1 | MSL2 | 0.148 | 0.224 | 1.187 | 1.081 | 0.134 | 1.3E-18 |
| 1 | MT-ATP6 | 0.729 | 0.849 | 11.43 | 9.423 | 0.279 | 3.8E-07 |
| 1 | MT-ATP8 | 0.438 | 0.471 | 4.682 | 3.882 | 0.27 | 0.00275 |
| 1 | MT-CO1 | 0.998 | 0.999 | 69.214 | 44.974 | 0.332 | 5E-160 |
| 1 | MT-CO2 | 0.863 | 0.944 | 18.829 | 14.137 | 0.314 | 4.1E-21 |
| 1 | MT-CO3 | 0.837 | 0.932 | 17.614 | 14.332 | 0.298 | 1.8E-16 |
| 1 | MT-CYB | 0.676 | 0.781 | 9.476 | 7.399 | 0.347 | 4E-14 |
| 1 | MTDH | 0.263 | 0.378 | 2.171 | 1.979 | 0.134 | 9.8E-16 |
| 1 | MTG1 | 0.112 | 0.169 | 0.814 | 0.743 | 0.132 | 2.6E-14 |
| 1 | MTMR3 | 0.118 | 0.148 | 0.938 | 0.647 | 0.436 | 0.00084 |
| 1 | MT-ND2 | 0.744 | 0.846 | 12.18 | 9.923 | 0.296 | 3.3E-08 |
| 1 | MT-ND3 | 0.418 | 0.444 | 4.318 | 3.493 | 0.264 | 0.00224 |
| 1 | MT-ND4 | 0.881 | 0.948 | 20.443 | 14.901 | 0.362 | 7.5E-42 |
| 1 | MT-ND4 | 0.691 | 0.804 | 9.866 | 7.811 | 0.337 | 2.4E-11 |
| 1 | MT-ND4L | 0.923 | 0.947 | 27.704 | 19.244 | 0.424 | 2E-109 |
| 1 | MTREX | 0.14 | 0.201 | 1.073 | 0.918 | 0.224 | 3.4E-12 |
| 1 | MTSS1 | 0.248 | 0.339 | 2.19 | 1.926 | 0.186 | 2.9E-11 |
| 1 | MYH9 | 0.293 | 0.372 | 2.477 | 2.02 | 0.341 | 0.00244 |
| 1 | MYO4A | 0.122 | 0.167 | 0.992 | 0.744 | 0.394 | 6.1E-08 |
| 1 | MYO9A | 0.144 | 0.184 | 1.143 | 0.877 | 0.394 | 4.2E-05 |
| 1 | MYO9B | 0.141 | 0.182 | 1.204 | 0.816 | 0.463 | 0.00011 |
| 1 | N4BP2L2 | 0.33 | 0.436 | 2.923 | 2.473 | 0.241 | 9.8E-06 |
| 1 | NAA34 | 0.107 | 0.141 | 0.81 | 0.668 | 0.277 | 1.4E-09 |
| 1 | NAA40 | 0.267 | 0.366 | 2.186 | 2.004 | 0.124 | 4.5E-12 |
| 1 | NAE1 | 0.103 | 0.14 | 0.739 | 0.672 | 0.137 | 7.7E-12 |
| 1 | NARF | 0.139 | 0.199 | 1.093 | 0.924 | 0.242 | 4.2E-12 |
| 1 | NBAS | 0.117 | 0.173 | 0.904 | 0.792 | 0.192 | 4.7E-14 |
| 1 | NBPF19 | 0.146 | 0.198 | 1.147 | 0.931 | 0.313 | 6.3E-05 |
| 1 | NBPF26 | 0.128 | 0.177 | 0.923 | 0.788 | 0.227 | 6.4E-09 |
| 1 | NCALD | 0.307 | 0.408 | 3.231 | 2.369 | 0.447 | 3.6E-05 |
| 1 | NCBP3 | 0.264 | 0.338 | 2.208 | 1.829 | 0.271 | 9.3E-05 |
| 1 | NCKAP1L | 0.163 | 0.22 | 1.26 | 1.013 | 0.314 | 4.4E-08 |
| 1 | NCOA2 | 0.314 | 0.411 | 2.828 | 2.363 | 0.249 | 1.4E-05 |
| 1 | NCOA3 | 0.177 | 0.24 | 1.408 | 1.274 | 0.144 | 1.8E-13 |

| | | | | | | | |
|---|---|---|---|---|---|---|---|
| 1 | NCOA6 | 0.147 | 0.213 | 1.068 | 0.972 | 0.136 | 1.1E-13 |
| 1 | NCOA7 | 0.183 | 0.264 | 1.449 | 1.324 | 0.139 | 4.2E-14 |
| 1 | NDRG3 | 0.173 | 0.241 | 1.246 | 1.114 | 0.161 | 3.7E-11 |
| 1 | NDUFV2 | 0.211 | 0.302 | 1.477 | 1.471 | 0.1 | 7E-15 |
| 1 | NEAT1 | 0.246 | 0.336 | 2.448 | 1.801 | 0.449 | 0.00008 |
| 1 | NEDD9 | 0.144 | 0.203 | 1.24 | 0.947 | 0.401 | 1.9E-06 |
| 1 | NEK7 | 0.268 | 0.369 | 2.264 | 1.986 | 0.19 | 1.7E-10 |
| 1 | NEMF | 0.209 | 0.294 | 1.642 | 1.474 | 0.163 | 2.5E-13 |
| 1 | NF1 | 0.219 | 0.294 | 1.812 | 1.443 | 0.223 | 4.1E-09 |
| 1 | NFATC2 | 0.263 | 0.34 | 2.327 | 1.776 | 0.39 | 0.00031 |
| 1 | NFKB1 | 0.308 | 0.449 | 3.396 | 3.011 | 0.173 | 6.4E-21 |
| 1 | NGLY1 | 0.171 | 0.248 | 1.374 | 1.224 | 0.168 | 4.7E-19 |
| 1 | NIBAN1 | 0.172 | 0.243 | 1.374 | 1.264 | 0.121 | 7.7E-17 |
| 1 | NIN | 0.197 | 0.289 | 1.629 | 1.474 | 0.144 | 3E-17 |
| 1 | NKTR | 0.307 | 0.407 | 2.62 | 2.208 | 0.247 | 2.9E-06 |
| 1 | NLRC4 | 0.171 | 0.234 | 1.366 | 1.161 | 0.234 | 7.7E-10 |
| 1 | NLRP1 | 0.194 | 0.242 | 1.662 | 1.173 | 0.402 | 0.00446 |
| 1 | NMT2 | 0.117 | 0.144 | 0.832 | 0.686 | 0.278 | 3.3E-06 |
| 1 | NNT | 0.112 | 0.144 | 0.836 | 0.701 | 0.244 | 2.3E-08 |
| 1 | NOTCH2 | 0.133 | 0.172 | 0.96 | 0.78 | 0.3 | 1.4E-05 |
| 1 | NPEPPS | 0.304 | 0.414 | 2.623 | 2.241 | 0.22 | 1.3E-08 |
| 1 | NPTN | 0.104 | 0.14 | 0.741 | 0.646 | 0.197 | 1.1E-10 |
| 1 | NR2C2 | 0.144 | 0.194 | 1.112 | 0.911 | 0.288 | 4.6E-08 |
| 1 | NRDC | 0.248 | 0.366 | 2.106 | 1.949 | 0.111 | 4.3E-14 |
| 1 | NRF1 | 0.109 | 0.144 | 0.807 | 0.64 | 0.312 | 1.4E-06 |
| 1 | NRIP1 | 0.14 | 0.214 | 1.194 | 1.019 | 0.23 | 4.4E-12 |
| 1 | NSD1 | 0.243 | 0.343 | 2.107 | 1.831 | 0.203 | 4.1E-11 |
| 1 | NSF | 0.274 | 0.342 | 2.364 | 1.911 | 0.307 | 0.00019 |
| 1 | NSMCE2 | 0.168 | 0.228 | 1.414 | 1.108 | 0.342 | 2.8E-08 |
| 1 | NSRP1 | 0.141 | 0.203 | 1.086 | 0.963 | 0.172 | 7.4E-13 |
| 1 | NT4C2 | 0.137 | 0.174 | 1.047 | 0.791 | 0.404 | 0.00021 |
| 1 | NUFIP2 | 0.207 | 0.321 | 1.746 | 1.632 | 0.106 | 8.8E-24 |
| 1 | NUMA1 | 0.129 | 0.178 | 0.973 | 0.81 | 0.264 | 2.7E-09 |
| 1 | NUMB | 0.118 | 0.147 | 0.88 | 0.699 | 0.333 | 1.8E-06 |
| 1 | NUP143 | 0.16 | 0.239 | 1.311 | 1.126 | 0.219 | 1.3E-16 |
| 1 | NUP214 | 0.123 | 0.171 | 0.904 | 0.743 | 0.263 | 2.2E-09 |
| 1 | NUP44 | 0.107 | 0.164 | 0.818 | 0.743 | 0.138 | 1E-14 |
| 1 | NUP48 | 0.104 | 0.162 | 0.798 | 0.723 | 0.142 | 2.3E-16 |
| 1 | NUP98 | 0.166 | 0.239 | 1.29 | 1.113 | 0.212 | 2.2E-13 |
| 1 | NUTM2A-AS1 | 0.164 | 0.228 | 1.293 | 1.072 | 0.271 | 1E-09 |
| 1 | NUTM2B-AS1 | 0.14 | 0.209 | 1.103 | 0.974 | 0.179 | 6.2E-11 |
| 1 | OCIAD2 | 0.148 | 0.207 | 1.048 | 0.948 | 0.13 | 1.4E-11 |
| 1 | OGT | 0.218 | 0.28 | 1.799 | 1.344 | 0.409 | 0.00044 |
| 1 | OPA1 | 0.16 | 0.239 | 1.264 | 1.133 | 0.149 | 2.5E-17 |
| 1 | ORC2 | 0.116 | 0.141 | 0.834 | 0.684 | 0.289 | 8.4E-06 |
| 1 | OSBPL9 | 0.162 | 0.228 | 1.293 | 1.074 | 0.266 | 8E-11 |
| 1 | OTULIN | 0.292 | 0.399 | 2.422 | 2.302 | 0.131 | 4.2E-11 |
| 1 | P2RY8 | 0.148 | 0.214 | 1.427 | 1.049 | 0.43 | 4.7E-08 |
| 1 | PACSIN2 | 0.108 | 0.139 | 0.786 | 0.6 | 0.39 | 4.9E-05 |
| 1 | PAFAH1B1 | 0.348 | 0.473 | 3.069 | 2.679 | 0.196 | 2.7E-08 |
| 1 | PAM | 0.218 | 0.299 | 1.912 | 1.47 | 0.284 | 3E-10 |
| 1 | PAPOLG | 0.101 | 0.143 | 0.733 | 0.629 | 0.221 | 6.3E-10 |
| 1 | PARN | 0.136 | 0.204 | 1.043 | 0.941 | 0.149 | 1.1E-16 |
| 1 | PARP14 | 0.214 | 0.279 | 1.68 | 1.373 | 0.292 | 4.7E-06 |

| | | | | | | | |
|---|---|---|---|---|---|---|---|
| 1 | PAXBP1 | 0.146 | 0.204 | 1.092 | 0.949 | 0.203 | 4.1E-11 |
| 1 | PCBP1-AS1 | 0.131 | 0.196 | 0.994 | 0.9 | 0.144 | 1.5E-14 |
| 1 | PCCA | 0.122 | 0.142 | 0.989 | 0.681 | 0.439 | 0.00138 |
| 1 | PCED1B | 0.269 | 0.338 | 2.281 | 1.844 | 0.306 | 0.0008 |
| 1 | PCF11 | 0.134 | 0.187 | 1.026 | 0.881 | 0.219 | 2.8E-10 |
| 1 | PCID2 | 0.101 | 0.134 | 0.71 | 0.604 | 0.233 | 8.2E-07 |
| 1 | PCM1 | 0.374 | 0.493 | 3.396 | 2.988 | 0.184 | 9.1E-06 |
| 1 | PCMTD1 | 0.239 | 0.34 | 1.911 | 1.771 | 0.11 | 2.1E-14 |
| 1 | PCMTD2 | 0.129 | 0.187 | 0.932 | 0.849 | 0.117 | 1.3E-13 |
| 1 | PCNT | 0.101 | 0.147 | 0.817 | 0.68 | 0.264 | 3E-11 |
| 1 | PCNX4 | 0.199 | 0.246 | 1.477 | 1.264 | 0.318 | 3.8E-05 |
| 1 | PDCD4 | 0.398 | 0.433 | 3.862 | 3.414 | 0.136 | 4.3E-08 |
| 1 | PDE4D | 0.432 | 0.46 | 4.279 | 3.772 | 0.182 | 0.00031 |
| 1 | PDE7A | 0.241 | 0.303 | 2.11 | 1.476 | 0.421 | 0.00148 |
| 1 | PDGFD | 0.194 | 0.262 | 1.771 | 1.339 | 0.403 | 1.9E-07 |
| 1 | PDK3 | 0.123 | 0.16 | 0.903 | 0.716 | 0.334 | 8.9E-06 |
| 1 | PDS4A | 0.2 | 0.272 | 1.614 | 1.294 | 0.32 | 2.4E-08 |
| 1 | PDS4B | 0.192 | 0.246 | 1.611 | 1.246 | 0.36 | 4.8E-07 |
| 1 | PDSS2 | 0.107 | 0.139 | 0.837 | 0.628 | 0.414 | 4E-05 |
| 1 | PDXDC1 | 0.171 | 0.229 | 1.347 | 1.062 | 0.343 | 1.5E-07 |
| 1 | PDZD8 | 0.132 | 0.201 | 1.094 | 0.944 | 0.198 | 6.4E-17 |
| 1 | PHACTR2 | 0.147 | 0.203 | 1.308 | 1.007 | 0.377 | 1.2E-05 |
| 1 | PHACTR4 | 0.148 | 0.226 | 1.21 | 1.044 | 0.198 | 8.7E-13 |
| 1 | PHC3 | 0.207 | 0.281 | 1.64 | 1.373 | 0.247 | 1.3E-08 |
| 1 | PHF14 | 0.123 | 0.164 | 0.934 | 0.744 | 0.309 | 7.1E-07 |
| 1 | PHF20 | 0.198 | 0.307 | 1.74 | 1.472 | 0.144 | 4.4E-24 |
| 1 | PHF20L1 | 0.22 | 0.304 | 1.73 | 1.443 | 0.146 | 4E-11 |
| 1 | PHF3 | 0.339 | 0.46 | 2.976 | 2.706 | 0.137 | 1.6E-09 |
| 1 | PHIP | 0.26 | 0.361 | 2.202 | 1.948 | 0.169 | 5E-12 |
| 1 | PHKB | 0.126 | 0.191 | 0.978 | 0.864 | 0.177 | 4.6E-16 |
| 1 | PI4KA | 0.118 | 0.144 | 0.934 | 0.643 | 0.438 | 0.00485 |
| 1 | PIAS1 | 0.238 | 0.328 | 1.941 | 1.649 | 0.234 | 1.7E-10 |
| 1 | PIBF1 | 0.172 | 0.234 | 1.393 | 1.174 | 0.244 | 4.1E-10 |
| 1 | PICALM | 0.344 | 0.441 | 3.101 | 2.416 | 0.302 | 0.00412 |
| 1 | PIK3C2A | 0.148 | 0.204 | 1.294 | 0.947 | 0.436 | 3.8E-05 |
| 1 | PIK3C3 | 0.11 | 0.147 | 0.814 | 0.69 | 0.24 | 1.1E-10 |
| 1 | PIK3CA | 0.149 | 0.217 | 1.163 | 1.029 | 0.176 | 7.9E-14 |
| 1 | PIK3CG | 0.143 | 0.207 | 1.14 | 0.981 | 0.216 | 1.8E-08 |
| 1 | PIK3IP1 | 0.171 | 0.232 | 1.266 | 1.084 | 0.223 | 4.3E-09 |
| 1 | PIK3R1 | 0.413 | 0.644 | 4.864 | 4.303 | 0.144 | 0.00712 |
| 1 | PIK3R4 | 0.193 | 0.246 | 1.669 | 1.284 | 0.378 | 4.3E-07 |
| 1 | PIKFYVE | 0.113 | 0.14 | 0.819 | 0.662 | 0.307 | 2.1E-06 |
| 1 | PITPNC1 | 0.446 | 0.626 | 6.809 | 4.726 | 0.427 | 2E-18 |
| 1 | PJA2 | 0.342 | 0.447 | 2.926 | 2.493 | 0.174 | 2.8E-07 |
| 1 | PKN2 | 0.238 | 0.343 | 1.911 | 1.717 | 0.144 | 2.8E-15 |
| 1 | PLCG2 | 0.212 | 0.276 | 1.824 | 1.368 | 0.417 | 4.7E-05 |
| 1 | PLCL2 | 0.233 | 0.34 | 2.078 | 1.764 | 0.236 | 2.3E-16 |
| 1 | PLEKHA1 | 0.19 | 0.264 | 1.444 | 1.3 | 0.162 | 1.1E-11 |
| 1 | PLEKHA2 | 0.361 | 0.464 | 3.3 | 2.704 | 0.287 | 0.00272 |
| 1 | PLEKHM1 | 0.114 | 0.164 | 0.887 | 0.728 | 0.284 | 6.8E-11 |
| 1 | PMPCB | 0.102 | 0.142 | 0.76 | 0.662 | 0.199 | 4.1E-13 |
| 1 | PMS1 | 0.102 | 0.142 | 0.733 | 0.678 | 0.111 | 8.7E-13 |
| 1 | PNISR | 0.314 | 0.434 | 2.708 | 2.434 | 0.144 | 1.2E-11 |
| 1 | PNPLA8 | 0.134 | 0.204 | 1.094 | 0.974 | 0.167 | 2.1E-17 |

| | | | | | | | |
|---|---|---|---|---|---|---|---|
| 1 | POGZ | 0.132 | 0.177 | 0.971 | 0.81 | 0.261 | 9.2E-08 |
| 1 | POLR2A | 0.121 | 0.167 | 1.188 | 0.787 | 0.494 | 3.5E-08 |
| 1 | POLR2B | 0.161 | 0.213 | 1.209 | 0.994 | 0.282 | 4.5E-07 |
| 1 | POU2F1 | 0.124 | 0.148 | 0.964 | 0.724 | 0.414 | 0.00011 |
| 1 | PPHLN1 | 0.171 | 0.234 | 1.313 | 1.092 | 0.266 | 2.6E-09 |
| 1 | PPIG | 0.162 | 0.248 | 1.27 | 1.183 | 0.103 | 4.5E-21 |
| 1 | PPM1A | 0.148 | 0.196 | 1.131 | 0.884 | 0.344 | 1.4E-06 |
| 1 | PPM1B | 0.14 | 0.184 | 1.196 | 0.838 | 0.414 | 0.00336 |
| 1 | PPM1L | 0.172 | 0.269 | 1.603 | 1.444 | 0.14 | 2.9E-24 |
| 1 | PPP1CB | 0.174 | 0.266 | 1.377 | 1.28 | 0.104 | 8.6E-21 |
| 1 | PPP1R12A | 0.382 | 0.416 | 3.466 | 3.21 | 0.142 | 1.3E-07 |
| 1 | PPP1R21 | 0.116 | 0.148 | 0.844 | 0.707 | 0.274 | 8.5E-08 |
| 1 | PPP2R2A | 0.21 | 0.316 | 1.744 | 1.601 | 0.124 | 1.4E-20 |
| 1 | PPP2R2B | 0.242 | 0.34 | 2.328 | 1.809 | 0.364 | 1.9E-07 |
| 1 | PPP2R3C | 0.129 | 0.184 | 0.99 | 0.844 | 0.214 | 3.9E-12 |
| 1 | PPP2R4A | 0.208 | 0.296 | 1.636 | 1.484 | 0.14 | 4.8E-14 |
| 1 | PPP2R4C | 0.49 | 0.639 | 4.3 | 4.708 | 0.171 | 0.0003 |
| 1 | PPP2R4E | 0.21 | 0.288 | 1.649 | 1.41 | 0.234 | 8E-10 |
| 1 | PPP3CB | 0.134 | 0.197 | 0.989 | 0.908 | 0.124 | 4.1E-14 |
| 1 | PPP3CC | 0.247 | 0.384 | 2.244 | 2.061 | 0.129 | 1.1E-21 |
| 1 | PPP3R1 | 0.219 | 0.309 | 1.8 | 1.428 | 0.236 | 7.1E-12 |
| 1 | PPP4R3A | 0.197 | 0.273 | 1.479 | 1.333 | 0.244 | 2.2E-10 |
| 1 | PPP6C | 0.141 | 0.202 | 1.046 | 0.908 | 0.204 | 3.4E-12 |
| 1 | PPP6R2 | 0.187 | 0.249 | 1.444 | 1.239 | 0.318 | 4.2E-10 |
| 1 | PPP6R3 | 0.221 | 0.292 | 1.828 | 1.433 | 0.341 | 1.9E-06 |
| 1 | PRDM2 | 0.249 | 0.338 | 2.148 | 1.747 | 0.29 | 1.9E-08 |
| 1 | PREX1 | 0.174 | 0.23 | 1.471 | 1.088 | 0.434 | 3.8E-06 |
| 1 | PRKAA1 | 0.11 | 0.144 | 0.784 | 0.683 | 0.2 | 1.2E-09 |
| 1 | PRKACB | 0.443 | 0.474 | 4.408 | 3.891 | 0.18 | 0.00071 |
| 1 | PRKAG2 | 0.104 | 0.141 | 0.806 | 0.689 | 0.227 | 7.6E-12 |
| 1 | PRKAR2A | 0.146 | 0.228 | 1.231 | 1.069 | 0.202 | 4.8E-14 |
| 1 | PRKCB | 0.214 | 0.282 | 2.131 | 1.441 | 0.464 | 3.4E-05 |
| 1 | PRKCH | 0.634 | 0.738 | 9.421 | 6.411 | 0.433 | 1.9E-13 |
| 1 | PRKCQ | 0.244 | 0.332 | 2.21 | 1.736 | 0.349 | 6.4E-05 |
| 1 | PRKD3 | 0.183 | 0.244 | 1.404 | 1.248 | 0.169 | 3.4E-11 |
| 1 | PRKDC | 0.21 | 0.308 | 1.644 | 1.414 | 0.128 | 1.5E-16 |
| 1 | PRKRIP1 | 0.101 | 0.148 | 0.741 | 0.64 | 0.188 | 7.2E-12 |
| 1 | PRKX | 0.197 | 0.271 | 1.684 | 1.349 | 0.32 | 1.8E-09 |
| 1 | PRKY | 0.108 | 0.14 | 0.818 | 0.641 | 0.341 | 1.4E-05 |
| 1 | PRMT2 | 0.347 | 0.476 | 3.044 | 2.827 | 0.112 | 3.3E-11 |
| 1 | PRPF38B | 0.276 | 0.386 | 2.242 | 2.062 | 0.121 | 3.5E-13 |
| 1 | PRPF40A | 0.244 | 0.364 | 2.047 | 1.899 | 0.109 | 3.3E-16 |
| 1 | PRPF4B | 0.344 | 0.482 | 3.033 | 2.731 | 0.142 | 1E-12 |
| 1 | PRPF6 | 0.142 | 0.214 | 1.047 | 0.984 | 0.102 | 1.5E-17 |
| 1 | PRPF8 | 0.114 | 0.14 | 0.848 | 0.644 | 0.391 | 4.8E-06 |
| 1 | PRR14L | 0.141 | 0.221 | 1.126 | 1.024 | 0.137 | 4.8E-15 |
| 1 | PRR4L | 0.142 | 0.196 | 1.149 | 0.906 | 0.344 | 1.6E-05 |
| 1 | PRRC2B | 0.23 | 0.297 | 1.902 | 1.467 | 0.374 | 0.00012 |
| 1 | PRRC2C | 0.468 | 0.61 | 4.813 | 4.192 | 0.199 | 0.0028 |
| 1 | PSME4 | 0.261 | 0.342 | 2.34 | 1.774 | 0.398 | 4.7E-05 |
| 1 | PSPC1 | 0.147 | 0.21 | 1.243 | 0.968 | 0.361 | 2.8E-07 |
| 1 | PTAR1 | 0.139 | 0.206 | 1.068 | 0.941 | 0.182 | 7.1E-14 |
| 1 | PTBP2 | 0.134 | 0.176 | 1.074 | 0.803 | 0.421 | 4.3E-06 |
| 1 | PTEN | 0.187 | 0.278 | 1.401 | 1.366 | 0.136 | 4.9E-18 |

| | | | | | | | |
|---|---|---|---|---|---|---|---|
| 1 | PTK2B | 0.197 | 0.24 | 1.48 | 1.227 | 0.364 | 0.00014 |
| 1 | PTPN12 | 0.334 | 0.44 | 3.062 | 2.484 | 0.244 | 3.5E-06 |
| 1 | PTPN22 | 0.326 | 0.426 | 2.893 | 2.444 | 0.179 | 2.5E-06 |
| 1 | PTPN4 | 0.342 | 0.49 | 3.204 | 2.944 | 0.118 | 4.1E-12 |
| 1 | PTPRA | 0.346 | 0.483 | 3.237 | 2.813 | 0.202 | 9.8E-08 |
| 1 | PTPRE | 0.266 | 0.38 | 2.237 | 2.049 | 0.126 | 4.1E-16 |
| 1 | PUM1 | 0.176 | 0.244 | 1.404 | 1.134 | 0.309 | 6.4E-10 |
| 1 | PUM2 | 0.297 | 0.381 | 2.442 | 2.008 | 0.346 | 0.00277 |
| 1 | PXK | 0.114 | 0.143 | 0.824 | 0.691 | 0.243 | 3.9E-07 |
| 1 | R3HDM1 | 0.174 | 0.238 | 1.349 | 1.144 | 0.224 | 2E-09 |
| 1 | R3HDM2 | 0.274 | 0.382 | 2.264 | 2.034 | 0.144 | 2.5E-12 |
| 1 | RAB10 | 0.168 | 0.24 | 1.274 | 1.176 | 0.116 | 1.8E-17 |
| 1 | RAB11FIP1 | 0.229 | 0.312 | 1.837 | 1.448 | 0.247 | 3.3E-09 |
| 1 | RAB18 | 0.124 | 0.174 | 0.961 | 0.768 | 0.323 | 2.9E-09 |
| 1 | RAB1A | 0.141 | 0.203 | 1.212 | 0.942 | 0.364 | 8.9E-08 |
| 1 | RAB21 | 0.127 | 0.196 | 0.943 | 0.886 | 0.104 | 1E-18 |
| 1 | RAB27A | 0.224 | 0.311 | 1.818 | 1.486 | 0.197 | 1.5E-11 |
| 1 | RAB27B | 0.132 | 0.197 | 1.016 | 0.937 | 0.116 | 1.5E-14 |
| 1 | RAB2A | 0.301 | 0.427 | 2.8 | 2.343 | 0.247 | 9.4E-13 |
| 1 | RAB3GAP2 | 0.136 | 0.202 | 1.044 | 0.933 | 0.164 | 1.4E-14 |
| 1 | RAB6A | 0.194 | 0.276 | 1.412 | 1.317 | 0.199 | 1.2E-12 |
| 1 | RAB8B | 0.462 | 0.611 | 4.809 | 4.178 | 0.203 | 0.00023 |
| 1 | RABEP1 | 0.16 | 0.211 | 1.226 | 0.992 | 0.306 | 7.5E-07 |
| 1 | RABGAP1 | 0.116 | 0.163 | 0.89 | 0.739 | 0.27 | 1.1E-09 |
| 1 | RABGAP1L | 0.644 | 0.76 | 8.988 | 7.144 | 0.331 | 1.1E-06 |
| 1 | RABGEF1 | 0.16 | 0.218 | 1.293 | 1.034 | 0.322 | 1.7E-08 |
| 1 | RAD40 | 0.106 | 0.144 | 0.817 | 0.724 | 0.172 | 3.6E-12 |
| 1 | RAD41B | 0.184 | 0.249 | 1.621 | 1.326 | 0.289 | 2.4E-09 |
| 1 | RALA | 0.177 | 0.236 | 1.381 | 1.268 | 0.123 | 2.1E-09 |
| 1 | RALGAPA1 | 0.314 | 0.418 | 2.819 | 2.398 | 0.234 | 1E-06 |
| 1 | RALGAPA2 | 0.14 | 0.181 | 1.074 | 0.838 | 0.36 | 1E-05 |
| 1 | RALGAPB | 0.121 | 0.16 | 0.908 | 0.714 | 0.346 | 8.5E-07 |
| 1 | RANBP9 | 0.109 | 0.138 | 0.864 | 0.612 | 0.497 | 0.00037 |
| 1 | RAP1A | 0.383 | 0.498 | 3.498 | 3.027 | 0.209 | 0.00141 |
| 1 | RAP1GAP2 | 0.174 | 0.224 | 2.21 | 1.287 | 0.78 | 0.00014 |
| 1 | RAP1GDS1 | 0.176 | 0.247 | 1.441 | 1.241 | 0.226 | 3.5E-14 |
| 1 | RAPGEF2 | 0.104 | 0.133 | 0.843 | 0.612 | 0.48 | 0.00016 |
| 1 | RAPGEF6 | 0.177 | 0.248 | 1.42 | 1.264 | 0.167 | 8E-16 |
| 1 | RARS2 | 0.114 | 0.164 | 0.884 | 0.728 | 0.28 | 4.4E-11 |
| 1 | RASA1 | 0.188 | 0.239 | 1.496 | 1.172 | 0.343 | 9.5E-05 |
| 1 | RASA2 | 0.448 | 0.614 | 4.749 | 4.284 | 0.148 | 4.8E-07 |
| 1 | RASAL3 | 0.134 | 0.19 | 0.992 | 0.844 | 0.214 | 4.7E-11 |
| 1 | RASGRP1 | 0.182 | 0.242 | 1.467 | 1.231 | 0.243 | 6.2E-11 |
| 1 | RASSF3 | 0.144 | 0.202 | 1.1 | 0.922 | 0.244 | 8.3E-11 |
| 1 | RASSF4 | 0.224 | 0.333 | 1.816 | 1.664 | 0.124 | 9.7E-20 |
| 1 | RB1 | 0.206 | 0.293 | 1.702 | 1.469 | 0.213 | 2.1E-13 |
| 1 | RB1CC1 | 0.248 | 0.344 | 2.212 | 1.864 | 0.247 | 1.4E-09 |
| 1 | RBL2 | 0.449 | 0.607 | 4.474 | 4.166 | 0.103 | 1.2E-07 |
| 1 | RBM26 | 0.184 | 0.268 | 1.436 | 1.314 | 0.127 | 1.4E-16 |
| 1 | RBM27 | 0.184 | 0.271 | 1.426 | 1.316 | 0.117 | 1.3E-17 |
| 1 | RBM33 | 0.263 | 0.364 | 2.209 | 1.874 | 0.237 | 3.5E-10 |
| 1 | RBM4 | 0.134 | 0.204 | 0.977 | 0.907 | 0.108 | 1.9E-17 |
| 1 | RBM6 | 0.218 | 0.288 | 1.816 | 1.424 | 0.34 | 3.8E-06 |
| 1 | RBMS1 | 0.189 | 0.276 | 1.414 | 1.368 | 0.147 | 3.9E-16 |

| | | | | | | | |
|---|---|---|---|---|---|---|---|
| 1 | RC3H1 | 0.332 | 0.467 | 2.92 | 2.64 | 0.144 | 6.4E-13 |
| 1 | RC3H2 | 0.134 | 0.187 | 0.984 | 0.844 | 0.204 | 3E-10 |
| 1 | RCOR3 | 0.163 | 0.222 | 1.234 | 1.046 | 0.24 | 2.1E-09 |
| 1 | RECQL | 0.16 | 0.234 | 1.231 | 1.123 | 0.133 | 4.3E-16 |
| 1 | REL | 0.344 | 0.492 | 3.331 | 2.984 | 0.148 | 7.4E-12 |
| 1 | RELB | 0.111 | 0.166 | 0.899 | 0.748 | 0.266 | 1.9E-13 |
| 1 | RELCH | 0.16 | 0.229 | 1.22 | 1.076 | 0.182 | 7.2E-13 |
| 1 | RERE | 0.174 | 0.241 | 1.417 | 1.142 | 0.398 | 9.5E-10 |
| 1 | REV3L | 0.134 | 0.184 | 1.043 | 0.86 | 0.279 | 3.2E-09 |
| 1 | RFC1 | 0.166 | 0.244 | 1.338 | 1.168 | 0.196 | 9.1E-16 |
| 1 | RFFL | 0.14 | 0.206 | 1.19 | 0.944 | 0.334 | 9.1E-09 |
| 1 | RFTN1 | 0.282 | 0.367 | 2.388 | 1.931 | 0.307 | 0.0001 |
| 1 | RFX3 | 0.2 | 0.248 | 1.673 | 1.296 | 0.367 | 2.5E-05 |
| 1 | RFX3-AS1 | 0.113 | 0.147 | 0.897 | 0.719 | 0.319 | 1.1E-08 |
| 1 | RFX7 | 0.146 | 0.186 | 1.171 | 0.869 | 0.431 | 0.0001 |
| 1 | RHOBTB3 | 0.146 | 0.19 | 1.044 | 0.898 | 0.231 | 1.4E-06 |
| 1 | RIC1 | 0.113 | 0.161 | 0.903 | 0.737 | 0.293 | 2.4E-10 |
| 1 | RICTOR | 0.291 | 0.4 | 2.443 | 2.179 | 0.229 | 1.3E-10 |
| 1 | RIF1 | 0.197 | 0.28 | 1.443 | 1.421 | 0.128 | 1.2E-14 |
| 1 | RIN3 | 0.2 | 0.242 | 1.77 | 1.233 | 0.422 | 0.00194 |
| 1 | RIPK1 | 0.142 | 0.234 | 1.184 | 1.094 | 0.114 | 4.9E-20 |
| 1 | RLF | 0.144 | 0.204 | 1.204 | 0.944 | 0.34 | 6.8E-11 |
| 1 | RMND4A | 0.112 | 0.144 | 0.849 | 0.689 | 0.32 | 4.5E-08 |
| 1 | RNF111 | 0.146 | 0.219 | 1.188 | 1.012 | 0.231 | 3.1E-11 |
| 1 | RNF114 | 0.228 | 0.314 | 1.849 | 1.494 | 0.221 | 7.7E-11 |
| 1 | RNF124 | 0.249 | 0.37 | 2.237 | 2.064 | 0.114 | 1.4E-21 |
| 1 | RNF13 | 0.141 | 0.204 | 1.062 | 0.947 | 0.164 | 8.2E-14 |
| 1 | RNF138 | 0.119 | 0.198 | 1.014 | 0.928 | 0.128 | 3E-24 |
| 1 | RNF144 | 0.121 | 0.168 | 0.889 | 0.742 | 0.261 | 6.8E-09 |
| 1 | RNF166 | 0.144 | 0.224 | 1.174 | 1.047 | 0.164 | 4.5E-14 |
| 1 | RNF216 | 0.206 | 0.278 | 1.686 | 1.321 | 0.342 | 4.1E-08 |
| 1 | RNF38 | 0.147 | 0.203 | 1.306 | 0.944 | 0.469 | 3.5E-05 |
| 1 | RNGTT | 0.161 | 0.236 | 1.311 | 1.114 | 0.233 | 4.7E-14 |
| 1 | RNMT | 0.16 | 0.244 | 1.302 | 1.181 | 0.14 | 8.8E-20 |
| 1 | RNPC3 | 0.148 | 0.21 | 1.08 | 0.963 | 0.164 | 9.6E-12 |
| 1 | ROCK2 | 0.108 | 0.137 | 0.788 | 0.607 | 0.378 | 0.00016 |
| 1 | RORA | 0.419 | 0.616 | 4.634 | 4.449 | 0.306 | 0.00128 |
| 1 | RPAP2 | 0.269 | 0.363 | 2.244 | 1.964 | 0.191 | 8.9E-09 |
| 1 | RPRD2 | 0.147 | 0.214 | 1.097 | 1.018 | 0.108 | 1.2E-14 |
| 1 | RPS6KA3 | 0.237 | 0.33 | 1.96 | 1.719 | 0.189 | 4.8E-12 |
| 1 | RPS6KA4 | 0.131 | 0.202 | 1.09 | 0.941 | 0.197 | 4.9E-18 |
| 1 | RPS6KC1 | 0.164 | 0.238 | 1.309 | 1.161 | 0.173 | 3.6E-14 |
| 1 | RRAS2 | 0.123 | 0.182 | 1.04 | 0.829 | 0.327 | 4.1E-13 |
| 1 | RSBN1 | 0.103 | 0.14 | 0.748 | 0.669 | 0.181 | 8.2E-12 |
| 1 | RSBN1L | 0.217 | 0.303 | 1.714 | 1.436 | 0.148 | 4.7E-12 |
| 1 | RSF1 | 0.292 | 0.396 | 2.46 | 2.198 | 0.162 | 4.6E-10 |
| 1 | RSRC1 | 0.142 | 0.196 | 1.122 | 0.921 | 0.284 | 1.4E-09 |
| 1 | RTF1 | 0.144 | 0.208 | 1.123 | 0.97 | 0.211 | 6.2E-13 |
| 1 | RTN4 | 0.214 | 0.323 | 1.726 | 1.61 | 0.1 | 1.9E-21 |
| 1 | RUNX3 | 0.229 | 0.339 | 2.108 | 1.806 | 0.224 | 4.4E-18 |
| 1 | S100PBP | 0.177 | 0.228 | 1.324 | 1.098 | 0.271 | 1.3E-05 |
| 1 | SACM1L | 0.194 | 0.272 | 1.418 | 1.313 | 0.209 | 1.2E-11 |
| 1 | SAFB | 0.204 | 0.28 | 1.449 | 1.37 | 0.187 | 4.8E-10 |
| 1 | SAFB2 | 0.133 | 0.193 | 0.987 | 0.878 | 0.169 | 2E-12 |

| | | | | | | | |
|---|---|---|---|---|---|---|---|
| 1 | SAP30BP | 0.104 | 0.144 | 0.772 | 0.679 | 0.187 | 2.7E-12 |
| 1 | SASH3 | 0.126 | 0.191 | 0.941 | 0.868 | 0.132 | 1.1E-16 |
| 1 | SATB1-AS1 | 0.114 | 0.144 | 0.884 | 0.777 | 0.184 | 3.2E-08 |
| 1 | SBF2 | 0.131 | 0.174 | 1.04 | 0.83 | 0.324 | 1.8E-07 |
| 1 | SBNO1 | 0.136 | 0.189 | 1.007 | 0.877 | 0.199 | 3.9E-10 |
| 1 | SCAF4 | 0.162 | 0.233 | 1.282 | 1.066 | 0.266 | 6.8E-13 |
| 1 | SCAF8 | 0.204 | 0.266 | 1.686 | 1.268 | 0.41 | 2.7E-05 |
| 1 | SCAPER | 0.199 | 0.263 | 1.603 | 1.324 | 0.274 | 2.1E-07 |
| 1 | SCFD1 | 0.144 | 0.214 | 1.142 | 1.004 | 0.198 | 8.8E-16 |
| 1 | SCYL2 | 0.144 | 0.206 | 1.087 | 0.943 | 0.189 | 7.9E-12 |
| 1 | SEC24B | 0.141 | 0.219 | 1.238 | 1.017 | 0.283 | 4.1E-13 |
| 1 | SEC63 | 0.17 | 0.243 | 1.376 | 1.147 | 0.263 | 1E-12 |
| 1 | SECISBP2 | 0.149 | 0.21 | 1.121 | 0.963 | 0.22 | 2.1E-11 |
| 1 | SECISBP2L | 0.224 | 0.322 | 1.797 | 1.642 | 0.13 | 1.3E-15 |
| 1 | SEMA4D | 0.143 | 0.226 | 1.226 | 1.064 | 0.204 | 1.7E-14 |
| 1 | SENP2 | 0.107 | 0.144 | 0.787 | 0.69 | 0.191 | 4.4E-11 |
| 1 | SENP4 | 0.188 | 0.263 | 1.408 | 1.274 | 0.243 | 4.4E-11 |
| 1 | SENP6 | 0.297 | 0.411 | 2.408 | 2.319 | 0.113 | 1.5E-12 |
| 1 | SENP7 | 0.238 | 0.32 | 1.932 | 1.626 | 0.248 | 2.2E-08 |
| 1 | SEPTIN9 | 0.144 | 0.217 | 1.234 | 1.017 | 0.279 | 6.7E-11 |
| 1 | SERINC3 | 0.164 | 0.24 | 1.24 | 1.124 | 0.141 | 3.2E-14 |
| 1 | SESN1 | 0.164 | 0.228 | 1.262 | 1.106 | 0.191 | 4.4E-11 |
| 1 | SETD3 | 0.211 | 0.296 | 1.614 | 1.444 | 0.141 | 1.8E-12 |
| 1 | SETD4 | 0.176 | 0.246 | 1.334 | 1.178 | 0.181 | 9E-12 |
| 1 | SFMBT2 | 0.111 | 0.141 | 0.968 | 0.692 | 0.484 | 2.1E-07 |
| 1 | SFSWAP | 0.138 | 0.2 | 1.018 | 0.92 | 0.146 | 1.8E-13 |
| 1 | SGCD | 0.114 | 0.16 | 1.149 | 1.037 | 0.16 | 8.9E-11 |
| 1 | SGK3 | 0.136 | 0.187 | 1.03 | 0.897 | 0.199 | 1.2E-09 |
| 1 | SH3KBP1 | 0.436 | 0.472 | 4.331 | 3.739 | 0.212 | 0.00028 |
| 1 | SHPRH | 0.137 | 0.18 | 1.018 | 0.84 | 0.277 | 1E-06 |
| 1 | SIDT1 | 0.222 | 0.303 | 1.92 | 1.464 | 0.294 | 8.5E-09 |
| 1 | SIK3 | 0.283 | 0.378 | 2.831 | 2.094 | 0.434 | 1.9E-06 |
| 1 | SIN3A | 0.137 | 0.194 | 1.083 | 0.916 | 0.241 | 1.4E-11 |
| 1 | SIPA1L1 | 0.222 | 0.326 | 1.978 | 1.813 | 0.126 | 2E-19 |
| 1 | SLA2 | 0.18 | 0.224 | 1.469 | 1.106 | 0.41 | 0.00043 |
| 1 | SLC14A4 | 0.182 | 0.244 | 1.434 | 1.249 | 0.201 | 3.4E-11 |
| 1 | SLC16A7 | 0.139 | 0.201 | 1.116 | 0.94 | 0.232 | 6.9E-13 |
| 1 | SLC30A9 | 0.104 | 0.147 | 0.743 | 0.64 | 0.193 | 2.7E-09 |
| 1 | SLC34A3 | 0.117 | 0.169 | 0.874 | 0.763 | 0.196 | 1.2E-11 |
| 1 | SLC38A1 | 0.442 | 0.492 | 4.812 | 4.034 | 0.244 | 0.00346 |
| 1 | SLC38A2 | 0.276 | 0.399 | 2.414 | 2.224 | 0.117 | 1.7E-17 |
| 1 | SLC44A2 | 0.137 | 0.211 | 1.081 | 0.976 | 0.148 | 9E-19 |
| 1 | SLC4A7 | 0.164 | 0.224 | 1.297 | 1.123 | 0.208 | 6.7E-10 |
| 1 | SLC9A9 | 0.146 | 0.208 | 1.193 | 0.987 | 0.273 | 4.8E-12 |
| 1 | SLCO3A1 | 0.129 | 0.167 | 1.184 | 0.793 | 0.478 | 7.5E-05 |
| 1 | SLCO4C1 | 0.101 | 0.142 | 0.824 | 0.708 | 0.219 | 1.1E-13 |
| 1 | SLF2 | 0.137 | 0.179 | 1.031 | 0.848 | 0.283 | 3E-06 |
| 1 | SLFN13 | 0.147 | 0.221 | 1.221 | 1.049 | 0.206 | 9.1E-12 |
| 1 | SLK | 0.133 | 0.184 | 1.02 | 0.844 | 0.244 | 1E-09 |
| 1 | SLMAP | 0.166 | 0.224 | 1.299 | 1.047 | 0.311 | 4.1E-08 |
| 1 | SLTM | 0.274 | 0.4 | 2.368 | 2.174 | 0.123 | 7.1E-18 |
| 1 | SLU7 | 0.102 | 0.143 | 0.729 | 0.664 | 0.137 | 2E-13 |
| 1 | SMAD2 | 0.176 | 0.26 | 1.38 | 1.243 | 0.141 | 1.8E-16 |
| 1 | SMAD4 | 0.118 | 0.161 | 0.817 | 0.726 | 0.171 | 9.3E-09 |

| | | | | | | | |
|---|---|---|---|---|---|---|---|
| 1 | SMAP1 | 0.219 | 0.304 | 1.776 | 1.401 | 0.243 | 8.4E-11 |
| 1 | SMARCA2 | 0.242 | 0.343 | 2.064 | 1.783 | 0.212 | 1.3E-13 |
| 1 | SMARCC1 | 0.2 | 0.294 | 1.642 | 1.444 | 0.183 | 1.4E-16 |
| 1 | SMC3 | 0.141 | 0.22 | 1.131 | 1.034 | 0.128 | 1.7E-14 |
| 1 | SMC4 | 0.2 | 0.291 | 1.496 | 1.444 | 0.144 | 1.6E-16 |
| 1 | SMC4 | 0.101 | 0.149 | 0.764 | 0.692 | 0.144 | 1.9E-12 |
| 1 | SMG1 | 0.241 | 0.342 | 2.124 | 1.744 | 0.276 | 7.2E-09 |
| 1 | SMG6 | 0.143 | 0.199 | 1.244 | 0.914 | 0.447 | 2.1E-05 |
| 1 | SMG7 | 0.112 | 0.142 | 0.836 | 0.683 | 0.293 | 1.1E-07 |
| 1 | SMURF1 | 0.101 | 0.128 | 0.818 | 0.482 | 0.49 | 0.00046 |
| 1 | SNAP23 | 0.141 | 0.204 | 1.044 | 0.941 | 0.163 | 1E-13 |
| 1 | SND1 | 0.104 | 0.138 | 0.824 | 0.497 | 0.464 | 9.1E-06 |
| 1 | SNRK | 0.269 | 0.376 | 2.279 | 1.993 | 0.194 | 3.1E-12 |
| 1 | SNRNP200 | 0.141 | 0.206 | 1.062 | 0.939 | 0.177 | 2.1E-14 |
| 1 | SNRNP70 | 0.102 | 0.148 | 0.771 | 0.696 | 0.149 | 6.1E-16 |
| 1 | SNTB1 | 0.103 | 0.16 | 0.906 | 0.744 | 0.263 | 1.2E-14 |
| 1 | SNW1 | 0.17 | 0.249 | 1.313 | 1.198 | 0.131 | 7.4E-16 |
| 1 | SNX1 | 0.122 | 0.171 | 0.941 | 0.744 | 0.319 | 1.4E-09 |
| 1 | SNX13 | 0.114 | 0.143 | 0.876 | 0.676 | 0.374 | 3.8E-06 |
| 1 | SNX14 | 0.179 | 0.243 | 1.387 | 1.162 | 0.246 | 4.1E-09 |
| 1 | SNX4 | 0.123 | 0.146 | 0.902 | 0.684 | 0.396 | 0.00026 |
| 1 | SOAT1 | 0.114 | 0.147 | 0.871 | 0.71 | 0.294 | 3.6E-08 |
| 1 | SOS1 | 0.276 | 0.343 | 2.378 | 1.96 | 0.279 | 0.00027 |
| 1 | SP3 | 0.243 | 0.339 | 2.018 | 1.706 | 0.242 | 4.3E-11 |
| 1 | SP4 | 0.103 | 0.161 | 0.774 | 0.713 | 0.119 | 1.8E-16 |
| 1 | SPAG9 | 0.204 | 0.283 | 1.649 | 1.376 | 0.27 | 4.1E-10 |
| 1 | SPART | 0.112 | 0.164 | 0.818 | 0.749 | 0.108 | 6.9E-13 |
| 1 | SPATA13 | 0.226 | 0.324 | 1.802 | 1.66 | 0.119 | 1.4E-16 |
| 1 | SPECC1L | 0.161 | 0.21 | 1.249 | 0.993 | 0.33 | 4.9E-06 |
| 1 | SPEN | 0.177 | 0.244 | 1.373 | 1.241 | 0.146 | 5E-14 |
| 1 | SPG11 | 0.174 | 0.242 | 1.343 | 1.173 | 0.194 | 9.1E-11 |
| 1 | SPG7 | 0.106 | 0.149 | 0.783 | 0.664 | 0.239 | 1.6E-09 |
| 1 | SPIDR | 0.271 | 0.342 | 2.494 | 1.834 | 0.443 | 0.00446 |
| 1 | SPOCK2 | 0.17 | 0.244 | 1.467 | 1.31 | 0.163 | 1.8E-14 |
| 1 | SPSB3 | 0.201 | 0.277 | 1.411 | 1.321 | 0.193 | 1.8E-10 |
| 1 | SPTBN1 | 0.137 | 0.217 | 1.126 | 1.046 | 0.104 | 8.4E-22 |
| 1 | SQSTM1 | 0.344 | 0.443 | 3.042 | 2.736 | 0.148 | 1E-06 |
| 1 | SRBD1 | 0.129 | 0.176 | 1.02 | 0.833 | 0.293 | 2E-08 |
| 1 | SREK1 | 0.148 | 0.213 | 1.162 | 1.001 | 0.214 | 2.2E-13 |
| 1 | SRSF11 | 0.488 | 0.64 | 4.944 | 4.494 | 0.141 | 4.2E-05 |
| 1 | SRSF4 | 0.212 | 0.3 | 1.666 | 1.487 | 0.163 | 4.1E-13 |
| 1 | SS18 | 0.137 | 0.18 | 1.032 | 0.823 | 0.326 | 1.7E-06 |
| 1 | SSBP3 | 0.12 | 0.14 | 0.943 | 0.669 | 0.411 | 0.00094 |
| 1 | SSH1 | 0.114 | 0.139 | 0.877 | 0.608 | 0.428 | 0.00927 |
| 1 | ST6GAL1 | 0.149 | 0.216 | 1.284 | 1.038 | 0.307 | 8.4E-13 |
| 1 | ST8SIA4 | 0.211 | 0.297 | 1.719 | 1.433 | 0.166 | 1.9E-13 |
| 1 | ST8SIA6 | 0.144 | 0.174 | 1.122 | 0.842 | 0.413 | 0.00949 |
| 1 | STAG2 | 0.36 | 0.489 | 3.247 | 2.946 | 0.14 | 4.2E-09 |
| 1 | STAM | 0.112 | 0.144 | 0.84 | 0.698 | 0.266 | 1.4E-08 |
| 1 | STAM2 | 0.11 | 0.167 | 0.824 | 0.746 | 0.124 | 2.5E-14 |
| 1 | STAT4 | 0.647 | 0.769 | 8.704 | 7.216 | 0.271 | 1.1E-08 |
| 1 | STAT4B | 0.142 | 0.194 | 1.096 | 0.913 | 0.264 | 7.5E-09 |
| 1 | STAU1 | 0.188 | 0.27 | 1.449 | 1.317 | 0.147 | 8E-15 |
| 1 | STAU2 | 0.16 | 0.209 | 1.271 | 1.009 | 0.333 | 3E-06 |

| | | | | | | | |
|---|---|---|---|---|---|---|---|
| 1 | STIM2 | 0.227 | 0.323 | 1.936 | 1.632 | 0.246 | 4.3E-13 |
| 1 | STK38 | 0.4 | 0.447 | 3.741 | 3.493 | 0.103 | 4.3E-10 |
| 1 | STK4 | 0.434 | 0.701 | 4.66 | 4.249 | 0.109 | 0.00044 |
| 1 | STRN | 0.147 | 0.191 | 1.121 | 0.884 | 0.341 | 6.7E-06 |
| 1 | STRN3 | 0.113 | 0.149 | 0.88 | 0.649 | 0.416 | 4.8E-06 |
| 1 | STT3B | 0.201 | 0.29 | 1.648 | 1.428 | 0.214 | 1.1E-14 |
| 1 | STX7 | 0.109 | 0.148 | 0.803 | 0.712 | 0.174 | 2.1E-11 |
| 1 | STX8 | 0.164 | 0.216 | 1.384 | 1.008 | 0.448 | 6.8E-06 |
| 1 | STXBP4 | 0.143 | 0.194 | 1.124 | 0.927 | 0.279 | 3.4E-08 |
| 1 | SUCLG2 | 0.178 | 0.24 | 1.447 | 1.147 | 0.334 | 3.2E-08 |
| 1 | SUCO | 0.222 | 0.326 | 1.882 | 1.66 | 0.181 | 1.2E-17 |
| 1 | SUDS3 | 0.134 | 0.194 | 0.941 | 0.872 | 0.126 | 2.5E-13 |
| 1 | SUMF1 | 0.148 | 0.192 | 1.247 | 0.923 | 0.434 | 1.5E-05 |
| 1 | SUN2 | 0.144 | 0.223 | 1.2 | 1.039 | 0.208 | 1.4E-13 |
| 1 | SUPT6H | 0.144 | 0.207 | 1.081 | 0.938 | 0.204 | 7E-12 |
| 1 | SUSD6 | 0.127 | 0.148 | 1.069 | 0.7 | 0.611 | 0.00196 |
| 1 | SYK | 0.111 | 0.14 | 0.814 | 0.664 | 0.294 | 3.8E-07 |
| 1 | SYTL2 | 0.181 | 0.239 | 1.48 | 1.222 | 0.277 | 6.3E-07 |
| 1 | TAB2 | 0.127 | 0.176 | 0.974 | 0.798 | 0.289 | 1.4E-09 |
| 1 | TACC1 | 0.227 | 0.29 | 1.84 | 1.437 | 0.364 | 0.00033 |
| 1 | TAF1 | 0.114 | 0.142 | 0.898 | 0.686 | 0.388 | 7E-07 |
| 1 | TAF14 | 0.21 | 0.288 | 1.619 | 1.429 | 0.18 | 1.4E-10 |
| 1 | TAF1D | 0.147 | 0.22 | 1.114 | 1.032 | 0.112 | 1.2E-16 |
| 1 | TAF3 | 0.144 | 0.187 | 1.142 | 0.87 | 0.406 | 2.7E-05 |
| 1 | TAOK1 | 0.137 | 0.196 | 1.049 | 0.908 | 0.222 | 1E-11 |
| 1 | TAOK3 | 0.377 | 0.404 | 3.621 | 3.008 | 0.268 | 3.9E-05 |
| 1 | TASOR2 | 0.162 | 0.232 | 1.242 | 1.101 | 0.173 | 1.9E-13 |
| 1 | TAX1BP1 | 0.373 | 0.417 | 3.338 | 3.104 | 0.104 | 3.9E-13 |
| 1 | TBC1D14 | 0.114 | 0.14 | 0.862 | 0.681 | 0.341 | 8.4E-06 |
| 1 | TBC1D31 | 0.128 | 0.172 | 0.934 | 0.784 | 0.243 | 8.7E-08 |
| 1 | TBC1D4 | 0.344 | 0.463 | 3.432 | 2.717 | 0.337 | 0.00162 |
| 1 | TBCK | 0.134 | 0.188 | 1.013 | 0.888 | 0.19 | 1E-10 |
| 1 | TBK1 | 0.167 | 0.246 | 1.281 | 1.178 | 0.12 | 6.2E-17 |
| 1 | TBRG1 | 0.192 | 0.276 | 1.444 | 1.343 | 0.106 | 2.3E-14 |
| 1 | TC2N | 0.109 | 0.169 | 0.941 | 0.836 | 0.17 | 8E-17 |
| 1 | TCAF2 | 0.121 | 0.18 | 0.916 | 0.83 | 0.141 | 1.2E-14 |
| 1 | TCF24 | 0.377 | 0.424 | 3.434 | 3.189 | 0.107 | 2.8E-12 |
| 1 | TENT2 | 0.267 | 0.374 | 2.24 | 2.021 | 0.149 | 2.3E-13 |
| 1 | TENT4B | 0.12 | 0.149 | 1.026 | 0.732 | 0.487 | 6.4E-06 |
| 1 | TERF1 | 0.116 | 0.172 | 0.869 | 0.787 | 0.144 | 3.2E-14 |
| 1 | TES | 0.271 | 0.397 | 2.324 | 2.167 | 0.1 | 4.4E-20 |
| 1 | TET2 | 0.19 | 0.284 | 1.468 | 1.407 | 0.146 | 9.2E-19 |
| 1 | TFDP2 | 0.344 | 0.434 | 3.37 | 2.639 | 0.342 | 0.00724 |
| 1 | TGFB1 | 0.41 | 0.441 | 4.818 | 3.83 | 0.331 | 2.4E-07 |
| 1 | TGFBR1 | 0.123 | 0.184 | 0.936 | 0.83 | 0.174 | 8E-14 |
| 1 | TGFBR2 | 0.26 | 0.343 | 2.13 | 1.798 | 0.244 | 4.4E-07 |
| 1 | THADA | 0.122 | 0.148 | 0.964 | 0.723 | 0.414 | 2.4E-05 |
| 1 | THOC1 | 0.122 | 0.178 | 0.969 | 0.822 | 0.238 | 6.7E-13 |
| 1 | THOC2 | 0.193 | 0.284 | 1.427 | 1.42 | 0.104 | 4.3E-18 |
| 1 | TLE1 | 0.148 | 0.218 | 1.296 | 1.081 | 0.262 | 7.9E-10 |
| 1 | TLK1 | 0.217 | 0.304 | 1.673 | 1.409 | 0.149 | 6.4E-13 |
| 1 | TLK2 | 0.134 | 0.177 | 1.048 | 0.794 | 0.414 | 6.1E-06 |
| 1 | TM9SF3 | 0.274 | 0.4 | 2.276 | 2.108 | 0.11 | 2.5E-18 |
| 1 | TMEM131 | 0.178 | 0.24 | 1.448 | 1.189 | 0.294 | 4.4E-11 |

| | | | | | | | |
|---|---|---|---|---|---|---|---|
| 1 | TMEM131L | 0.24 | 0.333 | 1.987 | 1.684 | 0.238 | 1.2E-10 |
| 1 | TMEM134 | 0.104 | 0.138 | 0.777 | 0.618 | 0.329 | 2.3E-06 |
| 1 | TMEM164 | 0.134 | 0.187 | 0.983 | 0.846 | 0.217 | 1.3E-09 |
| 1 | TMEM181 | 0.149 | 0.212 | 1.266 | 1.004 | 0.334 | 3E-11 |
| 1 | TMEM244 | 0.113 | 0.147 | 0.841 | 0.647 | 0.374 | 3E-05 |
| 1 | TMEM41B | 0.184 | 0.228 | 1.394 | 1.084 | 0.364 | 0.00322 |
| 1 | TMEM43 | 0.103 | 0.141 | 0.717 | 0.623 | 0.203 | 1E-07 |
| 1 | TMEM87A | 0.199 | 0.29 | 1.622 | 1.444 | 0.167 | 1.6E-14 |
| 1 | TMX3 | 0.117 | 0.174 | 0.844 | 0.779 | 0.118 | 2.8E-14 |
| 1 | TNFAIP8 | 0.183 | 0.267 | 1.411 | 1.338 | 0.176 | 1.4E-14 |
| 1 | TNIP1 | 0.106 | 0.14 | 0.797 | 0.664 | 0.264 | 7.2E-10 |
| 1 | TNKS | 0.206 | 0.283 | 1.692 | 1.387 | 0.286 | 1E-09 |
| 1 | TNKS2 | 0.213 | 0.301 | 1.694 | 1.49 | 0.184 | 1.4E-12 |
| 1 | TNPO1 | 0.143 | 0.217 | 1.198 | 1.002 | 0.249 | 2.2E-11 |
| 1 | TNRC6A | 0.169 | 0.224 | 1.377 | 1.049 | 0.379 | 4.3E-07 |
| 1 | TOP2B | 0.283 | 0.383 | 2.423 | 2.106 | 0.202 | 9E-09 |
| 1 | TOPBP1 | 0.104 | 0.147 | 0.793 | 0.694 | 0.19 | 2.1E-13 |
| 1 | TPP2 | 0.241 | 0.341 | 2.144 | 1.807 | 0.247 | 7.7E-12 |
| 1 | TPR | 0.309 | 0.426 | 2.646 | 2.442 | 0.116 | 9.7E-12 |
| 1 | TRA2A | 0.18 | 0.284 | 1.427 | 1.39 | 0.136 | 4.1E-24 |
| 1 | TRAF3 | 0.107 | 0.133 | 0.868 | 0.491 | 0.446 | 0.00181 |
| 1 | TRAPPC10 | 0.331 | 0.437 | 2.914 | 2.439 | 0.247 | 1.8E-05 |
| 1 | TRAPPC8 | 0.142 | 0.199 | 1.063 | 0.917 | 0.213 | 9E-11 |
| 1 | TRIM33 | 0.198 | 0.248 | 1.619 | 1.241 | 0.372 | 9.8E-06 |
| 1 | TRIM44 | 0.126 | 0.179 | 0.991 | 0.822 | 0.27 | 2.9E-11 |
| 1 | TRIP11 | 0.136 | 0.189 | 1.04 | 0.891 | 0.223 | 2.7E-10 |
| 1 | TRIP12 | 0.311 | 0.41 | 2.674 | 2.269 | 0.237 | 3.9E-06 |
| 1 | TRPC4AP | 0.132 | 0.18 | 1.041 | 0.822 | 0.344 | 4.5E-08 |
| 1 | TRPM7 | 0.232 | 0.317 | 1.908 | 1.491 | 0.263 | 3.2E-09 |
| 1 | TSEN44 | 0.109 | 0.168 | 0.806 | 0.731 | 0.141 | 1.5E-14 |
| 1 | TSG101 | 0.141 | 0.187 | 1.06 | 0.841 | 0.317 | 1.5E-06 |
| 1 | TSPOAP1-AS1 | 0.204 | 0.247 | 1.642 | 1.288 | 0.341 | 0.00019 |
| 1 | TTC14 | 0.198 | 0.28 | 1.446 | 1.372 | 0.172 | 9.5E-13 |
| 1 | TTC17 | 0.304 | 0.392 | 2.644 | 2.19 | 0.272 | 0.00021 |
| 1 | TTC39C | 0.262 | 0.344 | 2.194 | 2.017 | 0.122 | 8.1E-11 |
| 1 | TULP4 | 0.148 | 0.219 | 1.107 | 1.02 | 0.117 | 1.1E-14 |
| 1 | TUT7 | 0.291 | 0.413 | 2.41 | 2.339 | 0.102 | 3.3E-14 |
| 1 | TYW1 | 0.107 | 0.147 | 0.826 | 0.703 | 0.233 | 1.1E-11 |
| 1 | UBA2 | 0.164 | 0.237 | 1.241 | 1.106 | 0.164 | 2.1E-13 |
| 1 | UBAC2 | 0.166 | 0.234 | 1.34 | 1.103 | 0.281 | 2.8E-11 |
| 1 | UBAP1 | 0.124 | 0.171 | 0.978 | 0.76 | 0.362 | 2.7E-08 |
| 1 | UBE2B | 0.199 | 0.283 | 1.497 | 1.343 | 0.147 | 2.4E-13 |
| 1 | UBE2E1 | 0.121 | 0.148 | 0.942 | 0.714 | 0.401 | 8.1E-06 |
| 1 | UBE2G1 | 0.144 | 0.214 | 1.238 | 0.988 | 0.324 | 1.1E-09 |
| 1 | UBE2H | 0.172 | 0.234 | 1.34 | 1.122 | 0.246 | 1.2E-09 |
| 1 | UBE2I | 0.168 | 0.247 | 1.223 | 1.127 | 0.118 | 4.8E-16 |
| 1 | UBE2K | 0.164 | 0.24 | 1.31 | 1.124 | 0.222 | 6E-14 |
| 1 | UBE2Q2 | 0.141 | 0.217 | 1.167 | 1.003 | 0.218 | 4.2E-13 |
| 1 | UBE2R2 | 0.113 | 0.174 | 0.836 | 0.774 | 0.109 | 1.1E-16 |
| 1 | UBE2W | 0.106 | 0.16 | 0.763 | 0.707 | 0.109 | 9.7E-14 |
| 1 | UBE3A | 0.193 | 0.268 | 1.424 | 1.274 | 0.249 | 1.9E-10 |
| 1 | UBE3C | 0.128 | 0.174 | 0.946 | 0.782 | 0.29 | 3.6E-08 |
| 1 | UBQLN1 | 0.182 | 0.266 | 1.43 | 1.267 | 0.174 | 4.3E-14 |
| 1 | UBR1 | 0.173 | 0.234 | 1.426 | 1.1 | 0.374 | 4.2E-08 |

| | | | | | | | |
|---|---|---|---|---|---|---|---|
| 1 | UBR2 | 0.24 | 0.32 | 1.941 | 1.673 | 0.222 | 2.7E-08 |
| 1 | UBR3 | 0.143 | 0.181 | 1.111 | 0.829 | 0.423 | 0.00013 |
| 1 | UBR4 | 0.226 | 0.284 | 1.869 | 1.387 | 0.43 | 0.00199 |
| 1 | UBR4 | 0.109 | 0.14 | 0.868 | 0.648 | 0.4 | 8.7E-08 |
| 1 | UBXN7 | 0.114 | 0.143 | 0.898 | 0.673 | 0.417 | 3E-06 |
| 1 | UHRF1BP1L | 0.133 | 0.191 | 1.006 | 0.883 | 0.188 | 6.4E-12 |
| 1 | UIMC1 | 0.13 | 0.179 | 0.98 | 0.82 | 0.246 | 2.3E-09 |
| 1 | UMAD1 | 0.144 | 0.202 | 1.213 | 0.971 | 0.32 | 1.2E-06 |
| 1 | UPF2 | 0.206 | 0.29 | 1.647 | 1.44 | 0.193 | 1.3E-12 |
| 1 | USF3 | 0.111 | 0.162 | 0.814 | 0.739 | 0.141 | 3E-12 |
| 1 | USP16 | 0.149 | 0.236 | 1.24 | 1.134 | 0.127 | 2.3E-16 |
| 1 | USP24 | 0.228 | 0.312 | 1.942 | 1.489 | 0.297 | 3.1E-09 |
| 1 | USP24 | 0.198 | 0.277 | 1.63 | 1.34 | 0.271 | 3.5E-11 |
| 1 | USP33 | 0.248 | 0.343 | 2.037 | 1.863 | 0.129 | 7.9E-14 |
| 1 | USP34 | 0.428 | 0.461 | 4.112 | 3.43 | 0.22 | 0.00232 |
| 1 | USP48 | 0.164 | 0.238 | 1.329 | 1.127 | 0.237 | 1.4E-13 |
| 1 | USP8 | 0.177 | 0.244 | 1.34 | 1.169 | 0.198 | 9.1E-11 |
| 1 | USP9X | 0.238 | 0.309 | 2.044 | 1.442 | 0.414 | 9.7E-05 |
| 1 | USP9Y | 0.124 | 0.144 | 0.972 | 0.734 | 0.404 | 0.00032 |
| 1 | UVRAG | 0.216 | 0.319 | 1.803 | 1.646 | 0.132 | 4.8E-19 |
| 1 | VAV3 | 0.32 | 0.437 | 3.224 | 2.637 | 0.29 | 6.4E-09 |
| 1 | VDAC3 | 0.107 | 0.164 | 0.782 | 0.728 | 0.102 | 2.2E-14 |
| 1 | VMP1 | 0.134 | 0.2 | 1.049 | 0.94 | 0.172 | 4.1E-15 |
| 1 | VOPP1 | 0.176 | 0.264 | 1.423 | 1.308 | 0.121 | 4.7E-19 |
| 1 | VPS13A | 0.22 | 0.307 | 1.823 | 1.444 | 0.24 | 4.5E-11 |
| 1 | VPS13D | 0.144 | 0.18 | 1.173 | 0.818 | 0.419 | 0.00218 |
| 1 | VPS36 | 0.137 | 0.202 | 0.983 | 0.908 | 0.114 | 8.7E-14 |
| 1 | VPS37B | 0.179 | 0.264 | 1.603 | 1.296 | 0.307 | 2.3E-14 |
| 1 | VPS41 | 0.106 | 0.16 | 0.801 | 0.718 | 0.149 | 2E-14 |
| 1 | VPS43 | 0.137 | 0.204 | 1.081 | 0.937 | 0.206 | 9.1E-14 |
| 1 | VPS8 | 0.141 | 0.213 | 1.19 | 1.008 | 0.24 | 2.3E-11 |
| 1 | VTI1A | 0.262 | 0.344 | 2.234 | 1.802 | 0.311 | 1.1E-05 |
| 1 | VWA8 | 0.101 | 0.141 | 0.818 | 0.649 | 0.334 | 1.7E-08 |
| 1 | WAC | 0.266 | 0.349 | 2.247 | 1.867 | 0.274 | 3.6E-08 |
| 1 | WAKMAR2 | 0.122 | 0.17 | 0.961 | 0.826 | 0.219 | 1.3E-09 |
| 1 | WASHC4 | 0.16 | 0.214 | 1.211 | 1.013 | 0.248 | 4.1E-08 |
| 1 | WDFY1 | 0.133 | 0.204 | 1.04 | 0.934 | 0.168 | 1.5E-17 |
| 1 | WDPCP | 0.14 | 0.183 | 1.224 | 0.887 | 0.464 | 0.00318 |
| 1 | WDR11 | 0.144 | 0.2 | 1.077 | 0.932 | 0.208 | 7.2E-10 |
| 1 | WDR26 | 0.163 | 0.226 | 1.416 | 1.076 | 0.396 | 2E-09 |
| 1 | WDR37 | 0.127 | 0.197 | 0.987 | 0.892 | 0.146 | 1.2E-18 |
| 1 | WDR43 | 0.11 | 0.178 | 0.866 | 0.798 | 0.118 | 1.9E-20 |
| 1 | WDR48 | 0.141 | 0.198 | 1.111 | 0.891 | 0.319 | 1.8E-10 |
| 1 | WDR7 | 0.14 | 0.179 | 1.118 | 0.818 | 0.44 | 0.00019 |
| 1 | WDR70 | 0.184 | 0.231 | 1.437 | 1.064 | 0.431 | 0.0022 |
| 1 | WHAMM | 0.138 | 0.219 | 1.108 | 1.029 | 0.106 | 1.4E-22 |
| 1 | WNK1 | 0.214 | 0.293 | 1.764 | 1.446 | 0.288 | 2.2E-09 |
| 1 | WSB1 | 0.178 | 0.248 | 1.368 | 1.204 | 0.184 | 1.4E-11 |
| 1 | WWP1 | 0.134 | 0.184 | 0.996 | 0.824 | 0.274 | 8.4E-09 |
| 1 | XIST | 0.126 | 0.183 | 1.182 | 0.897 | 0.399 | 4.3E-12 |
| 1 | XKR6 | 0.104 | 0.128 | 0.788 | 0.474 | 0.449 | 0.00264 |
| 1 | XPC | 0.187 | 0.244 | 1.423 | 1.23 | 0.21 | 2.5E-09 |
| 1 | XPO1 | 0.227 | 0.32 | 1.894 | 1.629 | 0.218 | 1E-12 |
| 1 | XPO4 | 0.121 | 0.161 | 0.896 | 0.72 | 0.316 | 1.2E-06 |

| | | | | | | | |
|---|---|---|---|---|---|---|---|
| 1 | XPR1 | 0.131 | 0.174 | 1.06 | 0.783 | 0.436 | 1.6E-06 |
| 1 | YAF2 | 0.149 | 0.204 | 1.207 | 0.94 | 0.36 | 1.3E-08 |
| 1 | YARS | 0.219 | 0.301 | 1.744 | 1.478 | 0.146 | 4E-12 |
| 1 | YES1 | 0.243 | 0.343 | 2.729 | 2.044 | 0.416 | 3.5E-10 |
| 1 | YIPF4 | 0.107 | 0.14 | 0.77 | 0.618 | 0.316 | 1.3E-05 |
| 1 | YPEL1 | 0.116 | 0.148 | 0.843 | 0.663 | 0.363 | 7.4E-05 |
| 1 | YPEL3 | 0.249 | 0.344 | 2.079 | 1.884 | 0.141 | 2.8E-11 |
| 1 | YTHDC1 | 0.362 | 0.499 | 3.273 | 2.968 | 0.141 | 3.3E-10 |
| 1 | YWHAE | 0.147 | 0.241 | 1.22 | 1.112 | 0.133 | 9.2E-20 |
| 1 | YY1 | 0.148 | 0.234 | 1.201 | 1.101 | 0.126 | 1.9E-16 |
| 1 | ZAP70 | 0.201 | 0.279 | 1.704 | 1.331 | 0.347 | 1.3E-09 |
| 1 | ZBTB11 | 0.119 | 0.147 | 0.906 | 0.716 | 0.34 | 2.6E-06 |
| 1 | ZBTB20 | 0.49 | 0.643 | 8.182 | 6.179 | 0.404 | 4.5E-14 |
| 1 | ZBTB24 | 0.114 | 0.168 | 0.844 | 0.77 | 0.134 | 6.4E-13 |
| 1 | ZBTB44 | 0.146 | 0.204 | 1.1 | 0.94 | 0.211 | 3.5E-11 |
| 1 | ZC3H13 | 0.194 | 0.268 | 1.432 | 1.34 | 0.193 | 3.1E-11 |
| 1 | ZC3H18 | 0.109 | 0.164 | 0.836 | 0.731 | 0.194 | 4E-14 |
| 1 | ZC3H6 | 0.102 | 0.138 | 0.72 | 0.648 | 0.142 | 8.9E-08 |
| 1 | ZC3H7A | 0.187 | 0.249 | 1.429 | 1.267 | 0.174 | 1.7E-11 |
| 1 | ZCCHC17 | 0.114 | 0.166 | 0.849 | 0.743 | 0.194 | 4.9E-12 |
| 1 | ZCCHC7 | 0.224 | 0.296 | 1.892 | 1.401 | 0.334 | 4.9E-06 |
| 1 | ZDHHC17 | 0.103 | 0.14 | 0.764 | 0.674 | 0.181 | 1.1E-11 |
| 1 | ZDHHC20 | 0.284 | 0.401 | 2.447 | 2.172 | 0.173 | 7.4E-13 |
| 1 | ZFAND6 | 0.234 | 0.314 | 1.949 | 1.444 | 0.334 | 6E-07 |
| 1 | ZFC3H1 | 0.246 | 0.361 | 2.089 | 1.931 | 0.114 | 1.5E-19 |
| 1 | ZFR | 0.244 | 0.363 | 2.073 | 1.871 | 0.148 | 3.5E-14 |
| 1 | ZFX | 0.143 | 0.214 | 1.104 | 1.021 | 0.112 | 3.3E-17 |
| 1 | ZFYVE16 | 0.111 | 0.162 | 0.811 | 0.749 | 0.114 | 1.8E-12 |
| 1 | ZFYVE28 | 0.139 | 0.168 | 1.08 | 0.794 | 0.443 | 0.0075 |
| 1 | ZMYM2 | 0.192 | 0.263 | 1.43 | 1.27 | 0.269 | 1.7E-09 |
| 1 | ZMYM4 | 0.102 | 0.149 | 0.746 | 0.647 | 0.203 | 9.6E-12 |
| 1 | ZMYM6 | 0.128 | 0.179 | 0.968 | 0.814 | 0.24 | 2E-09 |
| 1 | ZMYND11 | 0.199 | 0.27 | 1.462 | 1.298 | 0.267 | 4.5E-09 |
| 1 | ZMYND8 | 0.143 | 0.188 | 1.124 | 0.864 | 0.38 | 4.2E-06 |
| 1 | ZNF106 | 0.141 | 0.216 | 1.14 | 1 | 0.201 | 1.3E-12 |
| 1 | ZNF131 | 0.163 | 0.24 | 1.304 | 1.136 | 0.2 | 1.2E-14 |
| 1 | ZNF148 | 0.233 | 0.314 | 1.881 | 1.46 | 0.27 | 1.5E-08 |
| 1 | ZNF292 | 0.283 | 0.367 | 2.463 | 1.931 | 0.341 | 0.00032 |
| 1 | ZNF331 | 0.124 | 0.187 | 1.293 | 1.132 | 0.191 | 9.9E-16 |
| 1 | ZNF407 | 0.14 | 0.178 | 1.214 | 0.822 | 0.462 | 0.00024 |
| 1 | ZNF418A | 0.104 | 0.147 | 0.744 | 0.678 | 0.133 | 6.4E-10 |
| 1 | ZNF441 | 0.169 | 0.234 | 1.303 | 1.11 | 0.231 | 1.2E-10 |
| 1 | ZNF600 | 0.218 | 0.299 | 1.742 | 1.464 | 0.144 | 3.8E-11 |
| 1 | ZNF609 | 0.116 | 0.142 | 0.924 | 0.629 | 0.447 | 0.00444 |
| 1 | ZNF611 | 0.117 | 0.144 | 0.842 | 0.682 | 0.303 | 2.9E-06 |
| 1 | ZNF638 | 0.263 | 0.363 | 2.144 | 1.964 | 0.133 | 4.9E-12 |
| 1 | ZNF642 | 0.162 | 0.238 | 1.249 | 1.141 | 0.142 | 6.4E-16 |
| 1 | ZNF644 | 0.279 | 0.38 | 2.432 | 2.012 | 0.274 | 1.9E-08 |
| 1 | ZNF644 | 0.122 | 0.161 | 0.918 | 0.737 | 0.316 | 1.3E-06 |
| 1 | ZNRF2 | 0.128 | 0.177 | 1.003 | 0.808 | 0.311 | 8.4E-09 |
| 1 | ZRANB2 | 0.307 | 0.427 | 2.479 | 2.394 | 0.107 | 1.4E-12 |
| 1 | ZSWIM6 | 0.138 | 0.194 | 1.178 | 0.922 | 0.344 | 4.8E-11 |
| 1 | ZZEF1 | 0.141 | 0.183 | 1.09 | 0.832 | 0.39 | 2.2E-05 |
| 2 | 45354 | 0.263 | 0.104 | 1.348 | 0.644 | 1.066 | 2E-107 |

| | | | | | | | |
|---|---|---|---|---|---|---|---|
| 2 | 45359 | 0.177 | 0.106 | 0.78 | 0.482 | 0.422 | 7.6E-24 |
| 2 | AASDHPPT | 0.191 | 0.107 | 0.826 | 0.444 | 0.476 | 4.9E-33 |
| 2 | ABCB1 | 0.414 | 0.287 | 2.284 | 1.879 | 0.282 | 2.4E-28 |
| 2 | ABCE1 | 0.116 | 0.08 | 0.487 | 0.396 | 0.297 | 2.3E-09 |
| 2 | ABCF1 | 0.27 | 0.2 | 1.273 | 1.142 | 0.147 | 1.8E-12 |
| 2 | ABHD17B | 0.103 | 0.067 | 0.431 | 0.348 | 0.306 | 1.2E-10 |
| 2 | ABHD18 | 0.143 | 0.087 | 0.648 | 0.473 | 0.444 | 1.4E-24 |
| 2 | ABHD2 | 0.337 | 0.203 | 1.702 | 1.237 | 0.46 | 2.2E-44 |
| 2 | ABI2 | 0.104 | 0.069 | 0.419 | 0.366 | 0.194 | 4.4E-10 |
| 2 | ABLIM1 | 0.299 | 0.218 | 1.423 | 1.338 | 0.186 | 1.9E-14 |
| 2 | ABRAXAS2 | 0.104 | 0.048 | 0.448 | 0.301 | 0.476 | 2.8E-19 |
| 2 | AC004441.1 | 0.207 | 0.079 | 0.914 | 0.449 | 1.024 | 2.4E-89 |
| 2 | AC004462.1 | 0.114 | 0.074 | 0.492 | 0.384 | 0.36 | 1.3E-12 |
| 2 | AC004687.1 | 0.177 | 0.114 | 0.748 | 0.636 | 0.234 | 1.5E-16 |
| 2 | AC008014.1 | 0.227 | 0.164 | 1.137 | 1.04 | 0.128 | 4.3E-12 |
| 2 | AC008104.3 | 0.119 | 0.041 | 0.496 | 0.278 | 0.834 | 2.3E-40 |
| 2 | AC008874.3 | 0.374 | 0.247 | 2.034 | 1.434 | 0.407 | 1.7E-34 |
| 2 | AC009093.2 | 0.148 | 0.101 | 0.648 | 0.494 | 0.148 | 1.1E-15 |
| 2 | AC009126.1 | 0.126 | 0.064 | 0.433 | 0.344 | 0.631 | 1.9E-27 |
| 2 | AC010894.3 | 0.377 | 0.231 | 1.828 | 1.319 | 0.471 | 4E-47 |
| 2 | AC014849.1 | 0.601 | 0.467 | 3.804 | 3.349 | 0.18 | 4E-20 |
| 2 | AC016831.7 | 0.133 | 0.076 | 0.446 | 0.427 | 0.344 | 3.2E-21 |
| 2 | AC068487.4 | 0.102 | 0.073 | 0.448 | 0.402 | 0.146 | 7.5E-07 |
| 2 | AC099063.1 | 0.213 | 0.094 | 0.929 | 0.421 | 0.834 | 2.4E-68 |
| 2 | AC118449.1 | 0.193 | 0.13 | 0.872 | 0.704 | 0.306 | 2.3E-16 |
| 2 | AC136474.1 | 0.174 | 0.107 | 0.731 | 0.449 | 0.389 | 4.9E-21 |
| 2 | AC139887.2 | 0.127 | 0.076 | 0.411 | 0.406 | 0.332 | 1.8E-17 |
| 2 | AC242426.2 | 0.102 | 0.049 | 0.422 | 0.317 | 0.414 | 4.2E-16 |
| 2 | AC243829.1 | 0.138 | 0.093 | 0.602 | 0.404 | 0.242 | 9.2E-12 |
| 2 | AC244297.3 | 0.473 | 0.361 | 3.741 | 2.404 | 0.483 | 2.4E-86 |
| 2 | AC244402.1 | 0.223 | 0.148 | 0.978 | 0.787 | 0.314 | 1.8E-19 |
| 2 | ACADM | 0.241 | 0.134 | 1.141 | 0.737 | 0.63 | 1.1E-48 |
| 2 | ACADSB | 0.101 | 0.072 | 0.419 | 0.376 | 0.144 | 8.4E-07 |
| 2 | ACADVL | 0.164 | 0.11 | 0.687 | 0.482 | 0.237 | 4.4E-14 |
| 2 | ACIN1 | 0.429 | 0.302 | 2.239 | 1.861 | 0.267 | 1.1E-26 |
| 2 | ACLY | 0.114 | 0.077 | 0.449 | 0.403 | 0.143 | 6.1E-10 |
| 2 | ACSL4 | 0.16 | 0.104 | 0.682 | 0.476 | 0.244 | 9.2E-15 |
| 2 | ACSL4 | 0.118 | 0.074 | 0.496 | 0.394 | 0.329 | 6.1E-13 |
| 2 | ACTR10 | 0.244 | 0.178 | 1.08 | 0.972 | 0.141 | 4.3E-12 |
| 2 | ACTR3 | 0.689 | 0.448 | 4.784 | 4.448 | 0.104 | 9.1E-12 |
| 2 | ACTR6 | 0.144 | 0.094 | 0.604 | 0.48 | 0.332 | 1.1E-14 |
| 2 | ACYP2 | 0.294 | 0.194 | 1.341 | 1.132 | 0.244 | 4.2E-24 |
| 2 | ADAM10 | 0.76 | 0.476 | 6.692 | 4.934 | 0.439 | 2.4E-66 |
| 2 | ADAM28 | 0.214 | 0.067 | 1.103 | 0.398 | 1.472 | 2E-133 |
| 2 | ADAM8 | 0.164 | 0.119 | 0.704 | 0.632 | 0.146 | 3.3E-09 |
| 2 | ADAMTSL4-AS | 0.11 | 0.062 | 0.484 | 0.384 | 0.334 | 2E-18 |
| 2 | ADAR | 0.713 | 0.34 | 4.368 | 2.447 | 1.133 | 2E-291 |
| 2 | ADARB2 | 0.14 | 0.041 | 0.609 | 0.298 | 1.031 | 3.4E-64 |
| 2 | ADCY7 | 0.182 | 0.124 | 0.781 | 0.696 | 0.164 | 7.8E-13 |
| 2 | ADH4 | 0.224 | 0.163 | 0.991 | 0.844 | 0.216 | 2.1E-12 |
| 2 | ADPGK | 0.138 | 0.082 | 0.464 | 0.433 | 0.383 | 3.8E-19 |
| 2 | ADRM1 | 0.174 | 0.123 | 0.703 | 0.642 | 0.13 | 3.2E-11 |
| 2 | ADSS | 0.332 | 0.244 | 1.617 | 1.462 | 0.146 | 2.2E-14 |
| 2 | AF164147.1 | 0.487 | 0.241 | 4.832 | 1.792 | 1.431 | 4E-284 |

| | | | | | | | |
|---|---|---|---|---|---|---|---|
| 2 | AFF1 | 0.309 | 0.173 | 1.427 | 1.011 | 0.496 | 1.1E-44 |
| 2 | AFTPH | 0.244 | 0.17 | 1.096 | 0.962 | 0.187 | 3.5E-14 |
| 2 | AGAP2 | 0.138 | 0.104 | 0.468 | 0.427 | 0.109 | 1.4E-06 |
| 2 | AGGF1 | 0.198 | 0.12 | 0.848 | 0.644 | 0.392 | 3.5E-24 |
| 2 | AGK | 0.214 | 0.132 | 0.983 | 0.721 | 0.447 | 7.3E-27 |
| 2 | AGL | 0.126 | 0.093 | 0.443 | 0.494 | 0.134 | 6.8E-07 |
| 2 | AGO2 | 0.208 | 0.121 | 0.894 | 0.702 | 0.34 | 7.3E-30 |
| 2 | AGO4 | 0.134 | 0.088 | 0.46 | 0.498 | 0.17 | 4.2E-12 |
| 2 | AHCTF1 | 0.194 | 0.141 | 0.86 | 0.801 | 0.102 | 3.2E-10 |
| 2 | AHCYL2 | 0.141 | 0.08 | 0.494 | 0.47 | 0.336 | 3.9E-22 |
| 2 | AHI1 | 0.244 | 0.16 | 1.378 | 1.076 | 0.347 | 1.4E-28 |
| 2 | AIDA | 0.181 | 0.109 | 0.801 | 0.469 | 0.493 | 1.3E-24 |
| 2 | AK3 | 0.134 | 0.086 | 0.441 | 0.442 | 0.284 | 1.9E-14 |
| 2 | AK9 | 0.148 | 0.084 | 0.646 | 0.464 | 0.474 | 8.2E-24 |
| 2 | AKAP10 | 0.19 | 0.12 | 0.813 | 0.64 | 0.322 | 1.2E-20 |
| 2 | AKAP11 | 0.172 | 0.12 | 0.763 | 0.643 | 0.248 | 1.4E-11 |
| 2 | AKAP4 | 0.27 | 0.174 | 1.429 | 1.081 | 0.403 | 3.3E-27 |
| 2 | AKAP7 | 0.119 | 0.072 | 0.416 | 0.382 | 0.434 | 2.4E-16 |
| 2 | AKAP9 | 0.484 | 0.374 | 2.872 | 2.612 | 0.137 | 1.8E-14 |
| 2 | AKR1B1 | 0.129 | 0.094 | 0.427 | 0.474 | 0.149 | 1.7E-07 |
| 2 | AL137846.1 | 0.103 | 0.042 | 0.424 | 0.269 | 0.662 | 3E-24 |
| 2 | AL644468.1 | 0.291 | 0.16 | 1.376 | 0.946 | 0.441 | 3.1E-41 |
| 2 | ALG2 | 0.11 | 0.076 | 0.464 | 0.382 | 0.282 | 1.1E-08 |
| 2 | ALG6 | 0.102 | 0.071 | 0.411 | 0.382 | 0.106 | 6.1E-08 |
| 2 | ALOX4AP | 0.761 | 0.412 | 8.474 | 3.469 | 1.289 | 0 |
| 2 | ANAPC4 | 0.148 | 0.107 | 0.637 | 0.488 | 0.114 | 1.2E-08 |
| 2 | ANAPC7 | 0.121 | 0.072 | 0.498 | 0.379 | 0.394 | 4.3E-17 |
| 2 | ANGEL2 | 0.167 | 0.12 | 0.72 | 0.633 | 0.184 | 8.9E-10 |
| 2 | ANKAR | 0.164 | 0.104 | 0.77 | 0.492 | 0.38 | 3.4E-17 |
| 2 | ANKFY1 | 0.284 | 0.112 | 1.344 | 0.624 | 1.114 | 8E-121 |
| 2 | ANKH | 0.144 | 0.083 | 0.607 | 0.466 | 0.383 | 3.9E-23 |
| 2 | ANKIB1 | 0.427 | 0.24 | 2.422 | 1.477 | 0.619 | 4E-69 |
| 2 | ANKRD12 | 0.732 | 0.482 | 4.714 | 4.083 | 0.169 | 2.7E-22 |
| 2 | ANKRD17 | 0.366 | 0.244 | 1.824 | 1.479 | 0.209 | 1.7E-23 |
| 2 | ANXA1 | 0.794 | 0.646 | 8.687 | 7.176 | 0.276 | 9.7E-34 |
| 2 | AOAH | 0.961 | 0.814 | 19.964 | 12.447 | 0.681 | 7E-249 |
| 2 | AP001011.1 | 0.244 | 0.142 | 1.104 | 0.829 | 0.414 | 8.7E-29 |
| 2 | AP3M1 | 0.136 | 0.083 | 0.474 | 0.434 | 0.404 | 4.7E-18 |
| 2 | AP3S1 | 0.172 | 0.13 | 0.724 | 0.67 | 0.113 | 7.1E-08 |
| 2 | AP4M1 | 0.131 | 0.084 | 0.422 | 0.446 | 0.226 | 6.7E-13 |
| 2 | APBB1IP | 0.611 | 0.472 | 3.924 | 3.443 | 0.184 | 1.3E-20 |
| 2 | APC | 0.297 | 0.209 | 1.396 | 1.248 | 0.141 | 4.2E-17 |
| 2 | API4 | 0.24 | 0.168 | 1.08 | 0.898 | 0.267 | 2.1E-16 |
| 2 | APIP | 0.109 | 0.047 | 0.434 | 0.291 | 0.476 | 4.4E-23 |
| 2 | APOBEC3C | 0.291 | 0.187 | 1.366 | 1.032 | 0.404 | 3.8E-30 |
| 2 | APOBEC3D | 0.129 | 0.061 | 0.44 | 0.324 | 0.739 | 4.7E-36 |
| 2 | APOBEC3F | 0.134 | 0.044 | 0.447 | 0.291 | 0.912 | 2.1E-40 |
| 2 | APOBEC3G | 0.649 | 0.367 | 4.14 | 2.741 | 0.904 | 2E-172 |
| 2 | APOL1 | 0.227 | 0.073 | 1.04 | 0.386 | 1.444 | 5E-134 |
| 2 | APOL2 | 0.141 | 0.06 | 0.646 | 0.314 | 1.043 | 4.8E-60 |
| 2 | APOL3 | 0.186 | 0.083 | 0.814 | 0.467 | 0.801 | 2.1E-48 |
| 2 | APOL6 | 0.437 | 0.271 | 3.643 | 1.7 | 1.103 | 1E-173 |
| 2 | APPBP2 | 0.192 | 0.118 | 0.838 | 0.644 | 0.344 | 2.2E-23 |
| 2 | APPL2 | 0.214 | 0.132 | 0.936 | 0.744 | 0.311 | 9.2E-24 |

| | | | | | | | |
|---|---|---|---|---|---|---|---|
| 2 | ARAP2 | 0.828 | 0.628 | 7.973 | 4.847 | 0.447 | 1E-84 |
| 2 | ARCN1 | 0.241 | 0.16 | 1.092 | 0.847 | 0.349 | 7.7E-26 |
| 2 | ARFGAP2 | 0.173 | 0.119 | 0.724 | 0.633 | 0.193 | 9E-13 |
| 2 | ARFGAP3 | 0.142 | 0.094 | 0.6 | 0.407 | 0.244 | 4.4E-12 |
| 2 | ARFIP1 | 0.192 | 0.114 | 0.829 | 0.618 | 0.423 | 1.7E-26 |
| 2 | ARHGAP17 | 0.144 | 0.1 | 0.621 | 0.441 | 0.172 | 8.7E-11 |
| 2 | ARHGAP18 | 0.139 | 0.094 | 0.603 | 0.409 | 0.243 | 4.1E-11 |
| 2 | ARHGAP24 | 0.314 | 0.227 | 1.412 | 1.344 | 0.17 | 1.4E-14 |
| 2 | ARHGAP34 | 0.149 | 0.09 | 0.684 | 0.4 | 0.442 | 4.8E-26 |
| 2 | ARHGEF1 | 0.383 | 0.28 | 1.916 | 1.744 | 0.126 | 1.5E-16 |
| 2 | ARHGEF12 | 0.138 | 0.096 | 0.483 | 0.413 | 0.184 | 1.1E-09 |
| 2 | ARHGEF3 | 0.719 | 0.403 | 4.878 | 3.014 | 0.963 | 3E-218 |
| 2 | ARHGEF6 | 0.281 | 0.187 | 1.299 | 1.036 | 0.326 | 8.7E-24 |
| 2 | ARID2 | 0.409 | 0.292 | 2.097 | 1.898 | 0.144 | 9.8E-21 |
| 2 | ARID4A | 0.284 | 0.203 | 1.313 | 1.164 | 0.172 | 7.3E-16 |
| 2 | ARL6IP6 | 0.163 | 0.094 | 0.719 | 0.494 | 0.437 | 4.4E-24 |
| 2 | ARL8B | 0.223 | 0.167 | 0.973 | 0.903 | 0.108 | 1.8E-09 |
| 2 | ARMC1 | 0.142 | 0.096 | 0.644 | 0.412 | 0.347 | 1.5E-16 |
| 2 | ARMH4 | 0.11 | 0.074 | 0.463 | 0.398 | 0.219 | 2.9E-09 |
| 2 | ARNT | 0.136 | 0.084 | 0.464 | 0.44 | 0.348 | 1.7E-14 |
| 2 | ARNTL | 0.396 | 0.284 | 2.026 | 1.783 | 0.184 | 4.8E-21 |
| 2 | ASAP1 | 0.203 | 0.144 | 0.912 | 0.837 | 0.124 | 1.3E-11 |
| 2 | ASB8 | 0.124 | 0.09 | 0.418 | 0.442 | 0.194 | 1E-07 |
| 2 | ASCC3 | 0.349 | 0.249 | 1.894 | 1.611 | 0.234 | 4.4E-20 |
| 2 | ASF1A | 0.12 | 0.072 | 0.498 | 0.376 | 0.406 | 1.4E-16 |
| 2 | ASH1L | 0.481 | 0.366 | 2.712 | 2.401 | 0.117 | 7.8E-16 |
| 2 | ASXL2 | 0.248 | 0.149 | 1.096 | 0.837 | 0.389 | 4.3E-32 |
| 2 | ATF1 | 0.17 | 0.117 | 0.747 | 0.631 | 0.262 | 4.5E-13 |
| 2 | ATF7 | 0.126 | 0.091 | 0.433 | 0.48 | 0.142 | 8.8E-08 |
| 2 | ATF7IP2 | 0.134 | 0.092 | 0.461 | 0.416 | 0.119 | 1.2E-10 |
| 2 | ATG2B | 0.138 | 0.091 | 0.447 | 0.479 | 0.218 | 9.5E-13 |
| 2 | ATG3 | 0.168 | 0.103 | 0.726 | 0.439 | 0.429 | 1.3E-20 |
| 2 | ATG7 | 0.182 | 0.116 | 0.792 | 0.674 | 0.233 | 1.9E-18 |
| 2 | ATL3 | 0.214 | 0.146 | 0.948 | 0.778 | 0.284 | 2.2E-16 |
| 2 | ATP10A | 0.26 | 0.148 | 1.209 | 0.884 | 0.442 | 6.7E-42 |
| 2 | ATP11C | 0.114 | 0.072 | 0.441 | 0.394 | 0.193 | 3E-12 |
| 2 | ATP4MPL | 0.363 | 0.274 | 1.734 | 1.433 | 0.178 | 1.2E-13 |
| 2 | ATP4PD | 0.238 | 0.177 | 1.048 | 0.944 | 0.147 | 9.5E-11 |
| 2 | ATP6V0D1 | 0.104 | 0.077 | 0.421 | 0.386 | 0.126 | 3.9E-06 |
| 2 | ATP6V1A | 0.13 | 0.081 | 0.441 | 0.44 | 0.291 | 7.1E-16 |
| 2 | ATP8A1 | 0.669 | 0.4 | 4.844 | 4.038 | 0.264 | 3E-37 |
| 2 | ATPAF1 | 0.11 | 0.076 | 0.442 | 0.381 | 0.249 | 6.4E-09 |
| 2 | ATRN | 0.124 | 0.079 | 0.423 | 0.432 | 0.274 | 6.2E-14 |
| 2 | ATXN1 | 0.436 | 0.297 | 2.411 | 2.033 | 0.246 | 2.4E-31 |
| 2 | ATXN10 | 0.169 | 0.114 | 0.724 | 0.636 | 0.186 | 4.5E-13 |
| 2 | ATXN2 | 0.204 | 0.137 | 0.892 | 0.77 | 0.212 | 2.5E-16 |
| 2 | ATXN3 | 0.187 | 0.112 | 0.84 | 0.627 | 0.423 | 1.3E-24 |
| 2 | AUTS2 | 0.437 | 0.266 | 2.373 | 1.834 | 0.371 | 1.3E-44 |
| 2 | AVL9 | 0.187 | 0.134 | 0.81 | 0.742 | 0.107 | 1.4E-10 |
| 2 | AZI2 | 0.234 | 0.104 | 1.076 | 0.471 | 0.914 | 2.1E-74 |
| 2 | B2M | 1 | 0.997 | 226.66 | 200.75 | 0.174 | 1.2E-60 |
| 2 | B3GALT4 | 0.104 | 0.064 | 0.414 | 0.326 | 0.343 | 2.8E-13 |
| 2 | B3GNT2 | 0.308 | 0.24 | 1.404 | 1.383 | 0.12 | 1E-09 |
| 2 | B4GALT4 | 0.109 | 0.046 | 0.448 | 0.31 | 0.462 | 6.2E-24 |

| | | | | | | | |
|---|---|---|---|---|---|---|---|
| 2 | BACH1 | 0.287 | 0.203 | 1.383 | 1.186 | 0.222 | 1.3E-17 |
| 2 | BACH2 | 0.264 | 0.14 | 1.43 | 0.961 | 0.473 | 4.1E-44 |
| 2 | BAG1 | 0.186 | 0.101 | 0.824 | 0.428 | 0.644 | 4.9E-36 |
| 2 | BAZ1A | 0.361 | 0.272 | 1.904 | 1.641 | 0.206 | 4.2E-14 |
| 2 | BAZ2A | 0.282 | 0.179 | 1.284 | 1.007 | 0.343 | 6.3E-30 |
| 2 | BBS9 | 0.24 | 0.162 | 1.094 | 0.99 | 0.144 | 2.4E-17 |
| 2 | BBX | 0.481 | 0.34 | 2.822 | 2.406 | 0.23 | 1.1E-24 |
| 2 | BCAP29 | 0.148 | 0.102 | 0.624 | 0.426 | 0.248 | 2.4E-11 |
| 2 | BCL2L11 | 0.118 | 0.084 | 0.498 | 0.462 | 0.109 | 7.1E-08 |
| 2 | BCL2L13 | 0.213 | 0.144 | 0.927 | 0.807 | 0.2 | 6.5E-12 |
| 2 | BCOR | 0.102 | 0.064 | 0.426 | 0.327 | 0.381 | 9.1E-12 |
| 2 | BICD1 | 0.143 | 0.073 | 0.689 | 0.414 | 0.734 | 1.7E-40 |
| 2 | BIRC2 | 0.313 | 0.214 | 1.487 | 1.229 | 0.274 | 1.7E-22 |
| 2 | BIRC3 | 0.273 | 0.14 | 1.398 | 0.993 | 0.493 | 1.2E-47 |
| 2 | BLMH | 0.184 | 0.092 | 0.8 | 0.477 | 0.744 | 1.3E-44 |
| 2 | BLOC1S4 | 0.114 | 0.078 | 0.472 | 0.41 | 0.203 | 3.2E-09 |
| 2 | BLZF1 | 0.136 | 0.067 | 0.488 | 0.344 | 0.768 | 1.9E-34 |
| 2 | BMPR2 | 0.263 | 0.14 | 1.284 | 0.823 | 0.641 | 3.4E-43 |
| 2 | BMT2 | 0.171 | 0.11 | 0.714 | 0.499 | 0.244 | 6.8E-17 |
| 2 | BNIP2 | 0.218 | 0.163 | 0.946 | 0.874 | 0.128 | 1.8E-09 |
| 2 | BOD1L1 | 0.33 | 0.21 | 1.644 | 1.263 | 0.38 | 1.2E-33 |
| 2 | BPGM | 0.147 | 0.09 | 0.66 | 0.461 | 0.417 | 1.9E-19 |
| 2 | BPTF | 0.464 | 0.443 | 3.499 | 3.314 | 0.119 | 2.8E-12 |
| 2 | BRCC3 | 0.108 | 0.068 | 0.423 | 0.343 | 0.304 | 4.1E-12 |
| 2 | BRD3 | 0.244 | 0.143 | 1.134 | 0.836 | 0.44 | 6.5E-28 |
| 2 | BRD7 | 0.308 | 0.221 | 1.608 | 1.321 | 0.284 | 4.4E-19 |
| 2 | BRD8 | 0.197 | 0.122 | 0.843 | 0.644 | 0.404 | 2.8E-23 |
| 2 | BROX | 0.278 | 0.192 | 1.281 | 1.078 | 0.249 | 4.4E-19 |
| 2 | BRWD3 | 0.207 | 0.132 | 0.909 | 0.74 | 0.276 | 7.4E-21 |
| 2 | BST2 | 0.789 | 0.344 | 7.042 | 2.478 | 1.409 | 0 |
| 2 | BTBD10 | 0.174 | 0.112 | 0.737 | 0.61 | 0.273 | 3.4E-17 |
| 2 | BTBD7 | 0.136 | 0.086 | 0.444 | 0.47 | 0.236 | 1.5E-14 |
| 2 | BTN2A1 | 0.14 | 0.087 | 0.6 | 0.469 | 0.346 | 2.4E-16 |
| 2 | BTN3A1 | 0.497 | 0.309 | 2.781 | 1.993 | 0.481 | 1.6E-63 |
| 2 | BTN3A2 | 0.467 | 0.284 | 2.444 | 1.746 | 0.449 | 3.3E-64 |
| 2 | BTN3A3 | 0.267 | 0.144 | 1.211 | 0.81 | 0.481 | 3.7E-49 |
| 2 | BTRC | 0.138 | 0.093 | 0.472 | 0.416 | 0.148 | 1.5E-11 |
| 2 | C11orf44 | 0.144 | 0.092 | 0.66 | 0.404 | 0.384 | 2.4E-20 |
| 2 | C11orf49 | 0.103 | 0.06 | 0.411 | 0.324 | 0.34 | 2.1E-14 |
| 2 | C12orf64 | 0.144 | 0.082 | 0.674 | 0.446 | 0.498 | 2.1E-30 |
| 2 | C14orf119 | 0.129 | 0.066 | 0.498 | 0.332 | 0.483 | 2E-28 |
| 2 | C14orf40 | 0.184 | 0.107 | 0.806 | 0.469 | 0.402 | 2.7E-27 |
| 2 | C17orf49 | 0.376 | 0.247 | 1.977 | 1.413 | 0.386 | 1.5E-34 |
| 2 | C17orf80 | 0.108 | 0.046 | 0.449 | 0.299 | 0.489 | 1.3E-23 |
| 2 | C18orf21 | 0.108 | 0.074 | 0.444 | 0.384 | 0.242 | 1.2E-08 |
| 2 | C19orf24 | 0.106 | 0.074 | 0.416 | 0.379 | 0.136 | 1.4E-07 |
| 2 | C1GALT1 | 0.414 | 0.218 | 2.14 | 1.287 | 0.74 | 2.1E-93 |
| 2 | C1orf131 | 0.101 | 0.044 | 0.424 | 0.284 | 0.484 | 1.5E-19 |
| 2 | C21orf91 | 0.298 | 0.174 | 1.496 | 0.941 | 0.643 | 1.1E-48 |
| 2 | C2CD4 | 0.192 | 0.127 | 0.832 | 0.7 | 0.249 | 4.7E-17 |
| 2 | C3orf38 | 0.211 | 0.142 | 0.944 | 0.813 | 0.216 | 5E-12 |
| 2 | C4orf22 | 0.111 | 0.066 | 0.44 | 0.334 | 0.427 | 3.4E-16 |
| 2 | C4orf3 | 0.424 | 0.28 | 2.221 | 1.611 | 0.463 | 4.7E-42 |
| 2 | C4orf41 | 0.171 | 0.103 | 0.766 | 0.449 | 0.444 | 4.3E-23 |

| | | | | | | | |
|---|---|---|---|---|---|---|---|
| 2 | C4orf46 | 0.673 | 0.421 | 4.791 | 3.004 | 0.674 | 2E-120 |
| 2 | C6orf62 | 0.347 | 0.217 | 1.811 | 1.246 | 0.428 | 4.4E-49 |
| 2 | C6orf89 | 0.162 | 0.112 | 0.674 | 0.474 | 0.232 | 6.7E-12 |
| 2 | C8orf33 | 0.122 | 0.092 | 0.412 | 0.467 | 0.132 | 7.3E-06 |
| 2 | C8orf76 | 0.104 | 0.07 | 0.417 | 0.367 | 0.184 | 2.8E-09 |
| 2 | C9orf84 | 0.109 | 0.066 | 0.463 | 0.346 | 0.419 | 6.4E-14 |
| 2 | CAB39 | 0.293 | 0.21 | 1.381 | 1.242 | 0.143 | 2E-14 |
| 2 | CACYBP | 0.22 | 0.161 | 0.974 | 0.879 | 0.147 | 3.8E-11 |
| 2 | CALCOCO1 | 0.113 | 0.068 | 0.447 | 0.363 | 0.333 | 2E-14 |
| 2 | CALCOCO2 | 0.246 | 0.183 | 1.18 | 0.991 | 0.242 | 1.3E-14 |
| 2 | CAMK1D | 0.341 | 0.236 | 1.748 | 1.444 | 0.178 | 9.2E-23 |
| 2 | CAND1 | 0.296 | 0.203 | 1.369 | 1.172 | 0.224 | 9.8E-21 |
| 2 | CANX | 0.442 | 0.346 | 2.444 | 2.221 | 0.144 | 1.3E-14 |
| 2 | CAPN2 | 0.4 | 0.284 | 2.174 | 1.729 | 0.331 | 6.4E-26 |
| 2 | CAPN7 | 0.303 | 0.208 | 1.418 | 1.214 | 0.222 | 1.1E-20 |
| 2 | CAPRIN1 | 0.262 | 0.189 | 1.207 | 1.069 | 0.174 | 2.8E-14 |
| 2 | CAPZA1 | 0.463 | 0.341 | 2.401 | 2.143 | 0.164 | 4E-20 |
| 2 | CAPZA2 | 0.304 | 0.218 | 1.39 | 1.237 | 0.169 | 2E-16 |
| 2 | CARD16 | 0.446 | 0.296 | 3.632 | 1.847 | 0.974 | 1E-142 |
| 2 | CARD19 | 0.116 | 0.072 | 0.478 | 0.379 | 0.334 | 1.7E-14 |
| 2 | CARD8 | 0.349 | 0.214 | 1.722 | 1.239 | 0.474 | 3.4E-44 |
| 2 | CARD8-AS1 | 0.104 | 0.044 | 0.461 | 0.292 | 0.66 | 9.9E-22 |
| 2 | CARS | 0.126 | 0.069 | 0.42 | 0.364 | 0.413 | 1.7E-23 |
| 2 | CASC4 | 0.308 | 0.224 | 1.464 | 1.33 | 0.138 | 1.2E-14 |
| 2 | CASD1 | 0.164 | 0.109 | 0.712 | 0.492 | 0.264 | 1.2E-14 |
| 2 | CASK | 0.174 | 0.084 | 0.838 | 0.474 | 0.819 | 1.9E-47 |
| 2 | CASP1 | 0.242 | 0.109 | 1.199 | 0.493 | 1.016 | 1.3E-88 |
| 2 | CASP10 | 0.146 | 0.079 | 0.674 | 0.422 | 0.674 | 4E-36 |
| 2 | CASP3 | 0.263 | 0.146 | 1.23 | 0.803 | 0.614 | 4E-47 |
| 2 | CASP4 | 0.34 | 0.196 | 1.709 | 1.118 | 0.612 | 2.3E-62 |
| 2 | CASP8AP2 | 0.178 | 0.114 | 0.804 | 0.62 | 0.377 | 4.7E-19 |
| 2 | CASTOR3 | 0.109 | 0.071 | 0.436 | 0.391 | 0.147 | 1.3E-10 |
| 2 | CBFA2T2 | 0.2 | 0.114 | 0.894 | 0.644 | 0.448 | 6E-32 |
| 2 | CBR4 | 0.249 | 0.142 | 1.187 | 0.848 | 0.468 | 3.3E-39 |
| 2 | CBWD2 | 0.14 | 0.06 | 0.666 | 0.319 | 1.063 | 4.1E-61 |
| 2 | CBX3 | 0.409 | 0.313 | 2.096 | 1.914 | 0.131 | 1.4E-13 |
| 2 | CCAR1 | 0.218 | 0.149 | 0.972 | 0.882 | 0.14 | 3.4E-11 |
| 2 | CCDC117 | 0.114 | 0.074 | 0.471 | 0.384 | 0.29 | 2.7E-12 |
| 2 | CCDC130 | 0.144 | 0.103 | 0.6 | 0.441 | 0.124 | 2.7E-09 |
| 2 | CCDC146 | 0.161 | 0.1 | 0.72 | 0.44 | 0.389 | 1.9E-19 |
| 2 | CCDC18 | 0.104 | 0.073 | 0.447 | 0.388 | 0.204 | 2.2E-08 |
| 2 | CCDC18-AS1 | 0.216 | 0.143 | 0.982 | 0.818 | 0.263 | 1.1E-18 |
| 2 | CCDC28A | 0.139 | 0.094 | 0.493 | 0.404 | 0.234 | 8.9E-11 |
| 2 | CCDC32 | 0.142 | 0.098 | 0.486 | 0.416 | 0.183 | 3.9E-10 |
| 2 | CCDC47 | 0.173 | 0.121 | 0.748 | 0.646 | 0.211 | 8.5E-12 |
| 2 | CCDC66 | 0.147 | 0.093 | 0.633 | 0.401 | 0.338 | 1.2E-16 |
| 2 | CCDC82 | 0.201 | 0.131 | 0.922 | 0.737 | 0.323 | 1.3E-19 |
| 2 | CCDC84B | 0.291 | 0.216 | 1.316 | 1.184 | 0.141 | 7.8E-13 |
| 2 | CCDC90B | 0.141 | 0.104 | 0.637 | 0.443 | 0.232 | 2E-11 |
| 2 | CCL4 | 0.949 | 0.912 | 47.323 | 42.644 | 0.123 | 8.9E-07 |
| 2 | CCL4L2 | 0.121 | 0.144 | 2.178 | 1.363 | 0.676 | 1.3E-06 |
| 2 | CCM2 | 0.234 | 0.16 | 1.073 | 0.921 | 0.221 | 2.1E-17 |
| 2 | CCNDBP1 | 0.204 | 0.134 | 0.863 | 0.723 | 0.244 | 3.2E-17 |
| 2 | CCNG2 | 0.146 | 0.084 | 0.64 | 0.436 | 0.444 | 3.4E-23 |

| | | | | | | | |
|---|---|---|---|---|---|---|---|
| 2 | CCNL2 | 0.203 | 0.139 | 0.922 | 0.778 | 0.246 | 8.5E-16 |
| 2 | CCPG1 | 0.326 | 0.238 | 1.478 | 1.384 | 0.19 | 1.7E-14 |
| 2 | CCSER1 | 0.111 | 0.074 | 0.441 | 0.38 | 0.247 | 6.1E-10 |
| 2 | CCSER2 | 0.801 | 0.492 | 7.744 | 4.414 | 0.418 | 8E-102 |
| 2 | CCT3 | 0.197 | 0.146 | 0.848 | 0.731 | 0.231 | 2.5E-10 |
| 2 | CCT4 | 0.184 | 0.132 | 0.773 | 0.674 | 0.196 | 2.3E-11 |
| 2 | CCT6A | 0.309 | 0.2 | 1.431 | 1.094 | 0.386 | 4.5E-30 |
| 2 | CCT7 | 0.18 | 0.133 | 0.768 | 0.683 | 0.168 | 3.9E-09 |
| 2 | CCT8 | 0.26 | 0.177 | 1.203 | 0.947 | 0.344 | 1.8E-20 |
| 2 | CD164 | 0.747 | 0.434 | 4.826 | 4.044 | 0.427 | 1.1E-84 |
| 2 | CD2AP | 0.303 | 0.146 | 1.498 | 0.899 | 0.736 | 2.6E-69 |
| 2 | CD2BP2 | 0.126 | 0.079 | 0.404 | 0.393 | 0.36 | 2.1E-14 |
| 2 | CD38 | 0.742 | 0.283 | 8.161 | 2.07 | 1.979 | 0 |
| 2 | CD44 | 0.627 | 0.409 | 4.697 | 3.264 | 0.424 | 3E-80 |
| 2 | CD46 | 0.402 | 0.372 | 2.784 | 2.43 | 0.197 | 4.2E-22 |
| 2 | CD47 | 0.439 | 0.421 | 3.143 | 2.844 | 0.144 | 9.7E-14 |
| 2 | CD48 | 0.428 | 0.394 | 3.144 | 2.468 | 0.297 | 2.1E-24 |
| 2 | CD48 | 0.348 | 0.214 | 1.84 | 1.272 | 0.433 | 3.4E-41 |
| 2 | CD68 | 0.169 | 0.021 | 0.879 | 0.116 | 2.918 | 2E-248 |
| 2 | CD74 | 0.902 | 0.668 | 12.269 | 7.402 | 0.71 | 2E-193 |
| 2 | CD82 | 0.208 | 0.073 | 0.941 | 0.448 | 1.087 | 5E-104 |
| 2 | CD8A | 0.131 | 0.094 | 0.634 | 0.462 | 0.174 | 4.3E-08 |
| 2 | CD96 | 0.884 | 0.494 | 11.281 | 6.319 | 0.836 | 7E-260 |
| 2 | CDC14A | 0.267 | 0.184 | 1.277 | 1.116 | 0.194 | 1E-18 |
| 2 | CDC16 | 0.233 | 0.144 | 1.044 | 0.847 | 0.301 | 7.5E-20 |
| 2 | CDC26 | 0.142 | 0.11 | 0.643 | 0.444 | 0.213 | 4E-09 |
| 2 | CDC27 | 0.21 | 0.142 | 0.924 | 0.842 | 0.117 | 2.9E-11 |
| 2 | CDC37L1 | 0.112 | 0.076 | 0.447 | 0.399 | 0.196 | 1E-09 |
| 2 | CDC40 | 0.247 | 0.181 | 1.173 | 1.009 | 0.217 | 3.7E-16 |
| 2 | CDC42EP3 | 0.446 | 0.334 | 2.414 | 2.178 | 0.208 | 2.1E-19 |
| 2 | CDC42SE2 | 0.876 | 0.7 | 9.247 | 7.034 | 0.396 | 4.4E-78 |
| 2 | CDC4L | 0.213 | 0.163 | 0.979 | 0.901 | 0.12 | 4.4E-08 |
| 2 | CDK19 | 0.228 | 0.148 | 1.013 | 0.907 | 0.149 | 1.9E-14 |
| 2 | CDK4RAP2 | 0.242 | 0.174 | 1.136 | 1.01 | 0.169 | 7.3E-13 |
| 2 | CDK8 | 0.146 | 0.092 | 0.629 | 0.414 | 0.292 | 1.3E-16 |
| 2 | CDKN1B | 0.441 | 0.347 | 2.413 | 2.317 | 0.117 | 1.6E-10 |
| 2 | CEBPD | 0.46 | 0.207 | 3.074 | 1.333 | 1.206 | 2E-177 |
| 2 | CEP104 | 0.163 | 0.098 | 0.701 | 0.421 | 0.43 | 1.4E-21 |
| 2 | CEP134 | 0.141 | 0.1 | 0.668 | 0.461 | 0.243 | 6.5E-14 |
| 2 | CEP162 | 0.118 | 0.063 | 0.404 | 0.344 | 0.443 | 4.2E-24 |
| 2 | CEP290 | 0.16 | 0.107 | 0.736 | 0.498 | 0.299 | 7.3E-14 |
| 2 | CEP340 | 0.488 | 0.342 | 2.804 | 2.387 | 0.232 | 4.1E-26 |
| 2 | CEP44 | 0.108 | 0.071 | 0.434 | 0.382 | 0.189 | 4.5E-10 |
| 2 | CEP47 | 0.242 | 0.189 | 1.141 | 1.043 | 0.13 | 4.5E-08 |
| 2 | CEP63 | 0.117 | 0.074 | 0.477 | 0.387 | 0.302 | 4.7E-13 |
| 2 | CEPT1 | 0.124 | 0.074 | 0.41 | 0.401 | 0.347 | 1.1E-17 |
| 2 | CERK | 0.164 | 0.097 | 0.708 | 0.426 | 0.429 | 1.4E-22 |
| 2 | CERS6 | 0.146 | 0.101 | 0.636 | 0.48 | 0.133 | 4.2E-11 |
| 2 | CERT1 | 0.337 | 0.209 | 1.648 | 1.237 | 0.423 | 7.4E-39 |
| 2 | CFAP298 | 0.127 | 0.093 | 0.422 | 0.486 | 0.106 | 3.3E-07 |
| 2 | CFLAR | 0.646 | 0.491 | 4.484 | 3.639 | 0.302 | 3.7E-36 |
| 2 | CGGBP1 | 0.473 | 0.339 | 2.686 | 2.128 | 0.336 | 2.5E-30 |
| 2 | CHD8 | 0.238 | 0.167 | 1.084 | 0.94 | 0.19 | 4.1E-14 |
| 2 | CHIC2 | 0.386 | 0.287 | 1.924 | 1.741 | 0.144 | 4E-16 |

| | | | | | | | |
|---|---|---|---|---|---|---|---|
| 2 | CHKB | 0.183 | 0.096 | 0.763 | 0.499 | 0.612 | 6.2E-38 |
| 2 | CHM | 0.132 | 0.073 | 0.444 | 0.383 | 0.408 | 9.8E-24 |
| 2 | CHMP2A | 0.178 | 0.137 | 0.747 | 0.702 | 0.11 | 4.6E-07 |
| 2 | CHMP4 | 0.294 | 0.101 | 1.36 | 0.44 | 1.332 | 3E-160 |
| 2 | CHMP4A | 0.247 | 0.184 | 1.18 | 1.013 | 0.221 | 4.6E-14 |
| 2 | CHORDC1 | 0.213 | 0.142 | 0.946 | 0.828 | 0.208 | 1.4E-12 |
| 2 | CHST11 | 0.469 | 0.324 | 2.66 | 2.301 | 0.209 | 7E-32 |
| 2 | CHST12 | 0.434 | 0.231 | 2.306 | 1.346 | 0.777 | 2.1E-98 |
| 2 | CHTOP | 0.102 | 0.069 | 0.394 | 0.362 | 0.126 | 1.3E-08 |
| 2 | CHURC1 | 0.243 | 0.184 | 1.146 | 1.003 | 0.204 | 7E-14 |
| 2 | CIAO2A | 0.117 | 0.073 | 0.486 | 0.36 | 0.434 | 2.9E-14 |
| 2 | CKAP2 | 0.176 | 0.104 | 0.782 | 0.48 | 0.432 | 4.2E-24 |
| 2 | CKAP4 | 0.18 | 0.133 | 0.8 | 0.739 | 0.114 | 1.1E-08 |
| 2 | CKLF | 0.444 | 0.29 | 2.431 | 1.834 | 0.406 | 3.5E-44 |
| 2 | CLASP1 | 0.437 | 0.31 | 2.279 | 2.016 | 0.177 | 2.1E-23 |
| 2 | CLCN3 | 0.188 | 0.114 | 0.828 | 0.619 | 0.419 | 1.5E-23 |
| 2 | CLDND1 | 0.487 | 0.236 | 3.317 | 1.433 | 1.114 | 2E-146 |
| 2 | CLEC2B | 0.661 | 0.448 | 4.666 | 3.369 | 0.47 | 9.7E-71 |
| 2 | CLEC2D | 0.447 | 0.346 | 3.724 | 2.644 | 0.488 | 8.8E-61 |
| 2 | CLIC3 | 0.747 | 0.41 | 7.046 | 4.494 | 0.648 | 1E-117 |
| 2 | CLINT1 | 0.447 | 0.341 | 2.444 | 2.174 | 0.169 | 3.2E-16 |
| 2 | CLIP1 | 0.286 | 0.209 | 1.364 | 1.236 | 0.143 | 2.8E-13 |
| 2 | CLIP4 | 0.243 | 0.142 | 1.114 | 0.804 | 0.47 | 9.2E-34 |
| 2 | CLN3 | 0.22 | 0.101 | 0.997 | 0.449 | 0.834 | 1.3E-64 |
| 2 | CLNS1A | 0.207 | 0.143 | 0.902 | 0.802 | 0.17 | 2.1E-10 |
| 2 | CLPX | 0.167 | 0.118 | 0.694 | 0.624 | 0.146 | 1.5E-10 |
| 2 | CLSTN3 | 0.119 | 0.067 | 0.486 | 0.347 | 0.487 | 4.2E-20 |
| 2 | CLTA | 0.284 | 0.203 | 1.279 | 1.167 | 0.132 | 4.3E-14 |
| 2 | CLTC | 0.408 | 0.343 | 2.762 | 2.21 | 0.322 | 3.5E-36 |
| 2 | CMPK2 | 0.427 | 0.07 | 2.444 | 0.44 | 2.403 | 0 |
| 2 | CMTM6 | 0.309 | 0.229 | 1.382 | 1.276 | 0.116 | 9.1E-13 |
| 2 | CMTR1 | 0.199 | 0.094 | 0.884 | 0.418 | 0.772 | 1.2E-44 |
| 2 | CMTR2 | 0.117 | 0.064 | 0.484 | 0.338 | 0.417 | 1.1E-21 |
| 2 | CNDP2 | 0.141 | 0.084 | 0.648 | 0.439 | 0.461 | 1.4E-26 |
| 2 | CNOT4 | 0.419 | 0.288 | 2.122 | 1.797 | 0.24 | 6.8E-28 |
| 2 | CNOT6 | 0.104 | 0.066 | 0.432 | 0.342 | 0.336 | 9.5E-12 |
| 2 | CNOT9 | 0.142 | 0.109 | 0.64 | 0.481 | 0.141 | 1.7E-09 |
| 2 | CNTRL | 0.278 | 0.191 | 1.308 | 1.136 | 0.203 | 2E-18 |
| 2 | COA1 | 0.224 | 0.143 | 0.99 | 0.874 | 0.179 | 1.1E-14 |
| 2 | COA3 | 0.132 | 0.071 | 0.421 | 0.342 | 0.467 | 2.9E-24 |
| 2 | COA4 | 0.14 | 0.092 | 0.489 | 0.488 | 0.271 | 4.1E-13 |
| 2 | COA6 | 0.104 | 0.047 | 0.424 | 0.298 | 0.41 | 7.4E-20 |
| 2 | COG2 | 0.111 | 0.068 | 0.446 | 0.341 | 0.378 | 7.1E-15 |
| 2 | COG6 | 0.162 | 0.103 | 0.684 | 0.467 | 0.271 | 4.3E-17 |
| 2 | COLGALT2 | 0.162 | 0.081 | 0.709 | 0.446 | 0.67 | 2.5E-38 |
| 2 | COMMD2 | 0.14 | 0.11 | 0.619 | 0.463 | 0.136 | 4.5E-08 |
| 2 | COMMD3 | 0.108 | 0.074 | 0.439 | 0.374 | 0.228 | 2E-08 |
| 2 | COMMD8 | 0.141 | 0.101 | 0.489 | 0.438 | 0.128 | 9.2E-09 |
| 2 | COMT | 0.109 | 0.082 | 0.446 | 0.399 | 0.161 | 1.3E-05 |
| 2 | COPB1 | 0.447 | 0.268 | 2.382 | 1.633 | 0.444 | 1.3E-64 |
| 2 | COPB2 | 0.26 | 0.14 | 1.144 | 0.811 | 0.409 | 1.9E-40 |
| 2 | COPS4 | 0.149 | 0.104 | 0.488 | 0.429 | 0.144 | 3.8E-10 |
| 2 | COPS6 | 0.147 | 0.119 | 0.643 | 0.608 | 0.102 | 4.4E-07 |
| 2 | COX17 | 0.308 | 0.202 | 1.404 | 1.122 | 0.323 | 3.2E-27 |

| | | | | | | | |
|---|---|---|---|---|---|---|---|
| 2 | COX7B | 0.281 | 0.218 | 1.234 | 1.141 | 0.1 | 9.4E-09 |
| 2 | CPEB3 | 0.106 | 0.043 | 0.432 | 0.294 | 0.446 | 2.5E-24 |
| 2 | CPLANE1 | 0.289 | 0.17 | 1.411 | 0.991 | 0.41 | 1.8E-42 |
| 2 | CPNE3 | 0.274 | 0.2 | 1.246 | 1.118 | 0.167 | 8.1E-14 |
| 2 | CPSF2 | 0.147 | 0.084 | 0.644 | 0.46 | 0.484 | 7.8E-23 |
| 2 | CPSF3 | 0.101 | 0.063 | 0.434 | 0.324 | 0.418 | 1.5E-12 |
| 2 | CRADD | 0.194 | 0.131 | 0.842 | 0.761 | 0.144 | 4.4E-14 |
| 2 | CREB1 | 0.317 | 0.183 | 1.46 | 1.077 | 0.434 | 4.3E-49 |
| 2 | CREB3L2 | 0.118 | 0.083 | 0.494 | 0.428 | 0.209 | 2.1E-08 |
| 2 | CREBRF | 0.329 | 0.24 | 1.497 | 1.398 | 0.192 | 1.2E-16 |
| 2 | CRLF3 | 0.32 | 0.201 | 1.428 | 1.146 | 0.403 | 1.2E-34 |
| 2 | CRTC3 | 0.208 | 0.13 | 0.888 | 0.738 | 0.267 | 1.2E-22 |
| 2 | CRYZL1 | 0.122 | 0.079 | 0.417 | 0.414 | 0.322 | 2.2E-12 |
| 2 | CSGALNACT2 | 0.376 | 0.277 | 1.894 | 1.701 | 0.146 | 2.8E-16 |
| 2 | CSK | 0.233 | 0.174 | 1.036 | 0.942 | 0.137 | 8.5E-10 |
| 2 | CSNK1A1 | 0.391 | 0.286 | 1.944 | 1.713 | 0.19 | 2.7E-17 |
| 2 | CSNK1G1 | 0.178 | 0.122 | 0.744 | 0.678 | 0.144 | 3.3E-13 |
| 2 | CSNK1G2 | 0.146 | 0.098 | 0.628 | 0.437 | 0.224 | 8.9E-13 |
| 2 | CSNK2A1 | 0.232 | 0.176 | 1.013 | 0.943 | 0.103 | 4.7E-09 |
| 2 | CTBS | 0.204 | 0.114 | 0.932 | 0.636 | 0.442 | 4.7E-34 |
| 2 | CTCF | 0.303 | 0.217 | 1.434 | 1.244 | 0.192 | 1.8E-16 |
| 2 | CTNNA1 | 0.133 | 0.094 | 0.47 | 0.408 | 0.164 | 2.1E-08 |
| 2 | CTR9 | 0.299 | 0.218 | 1.373 | 1.239 | 0.148 | 2.7E-14 |
| 2 | CTSO | 0.103 | 0.064 | 0.431 | 0.333 | 0.369 | 1.8E-12 |
| 2 | CTSS | 0.443 | 0.284 | 2.432 | 1.694 | 0.479 | 1.2E-44 |
| 2 | CUL2 | 0.216 | 0.146 | 0.946 | 0.848 | 0.147 | 1.1E-11 |
| 2 | CUL4 | 0.324 | 0.202 | 1.416 | 1.143 | 0.407 | 6.3E-37 |
| 2 | CUL4A | 0.317 | 0.23 | 1.49 | 1.338 | 0.146 | 1.7E-14 |
| 2 | CUL4B | 0.187 | 0.104 | 0.84 | 0.441 | 0.608 | 4.7E-33 |
| 2 | CUTC | 0.121 | 0.086 | 0.401 | 0.441 | 0.184 | 2.5E-08 |
| 2 | CWC22 | 0.19 | 0.11 | 0.839 | 0.494 | 0.497 | 1.2E-28 |
| 2 | CWC27 | 0.144 | 0.112 | 0.697 | 0.622 | 0.164 | 1.4E-08 |
| 2 | CXCR3 | 0.218 | 0.082 | 1.083 | 0.486 | 1.146 | 1E-99 |
| 2 | CXCR4 | 0.44 | 0.263 | 2.422 | 1.629 | 0.63 | 6.1E-68 |
| 2 | CXorf38 | 0.207 | 0.143 | 0.949 | 0.78 | 0.282 | 2.7E-14 |
| 2 | CYB4B | 0.229 | 0.168 | 1.014 | 0.914 | 0.14 | 4.5E-11 |
| 2 | CYBC1 | 0.104 | 0.074 | 0.426 | 0.374 | 0.184 | 1.6E-07 |
| 2 | CYC1 | 0.149 | 0.107 | 0.499 | 0.446 | 0.134 | 4.7E-09 |
| 2 | CYLD | 0.662 | 0.417 | 4.739 | 3.043 | 0.634 | 1E-108 |
| 2 | CYP2R1 | 0.113 | 0.073 | 0.477 | 0.392 | 0.284 | 3.4E-12 |
| 2 | CYTH3 | 0.186 | 0.088 | 0.841 | 0.491 | 0.774 | 2E-41 |
| 2 | DAAM1 | 0.214 | 0.134 | 0.994 | 0.8 | 0.314 | 4.6E-23 |
| 2 | DAB1 | 0.144 | 0.102 | 0.61 | 0.442 | 0.144 | 1.6E-09 |
| 2 | DARS | 0.18 | 0.118 | 0.761 | 0.628 | 0.278 | 1.4E-16 |
| 2 | DAXX | 0.137 | 0.088 | 0.462 | 0.446 | 0.301 | 9.9E-14 |
| 2 | DCAF10 | 0.172 | 0.108 | 0.744 | 0.616 | 0.294 | 4.7E-19 |
| 2 | DCAF11 | 0.104 | 0.074 | 0.419 | 0.384 | 0.128 | 4.1E-07 |
| 2 | DCAF6 | 0.141 | 0.102 | 0.63 | 0.462 | 0.164 | 2.9E-12 |
| 2 | DCAF7 | 0.223 | 0.14 | 0.991 | 0.767 | 0.371 | 1.9E-24 |
| 2 | DCAF8 | 0.334 | 0.227 | 1.48 | 1.31 | 0.27 | 1.7E-24 |
| 2 | DCLRE1C | 0.111 | 0.064 | 0.44 | 0.343 | 0.389 | 1.4E-17 |
| 2 | DCP2 | 0.308 | 0.216 | 1.444 | 1.227 | 0.244 | 2.8E-19 |
| 2 | DCPS | 0.134 | 0.08 | 0.467 | 0.403 | 0.492 | 4.1E-19 |
| 2 | DCTN1 | 0.143 | 0.101 | 0.603 | 0.428 | 0.191 | 9.5E-10 |

| | | | | | | | |
|---|---|---|---|---|---|---|---|
| 2 | DCTN2 | 0.146 | 0.101 | 0.604 | 0.4 | 0.272 | 1.9E-11 |
| 2 | DCUN1D1 | 0.149 | 0.108 | 0.628 | 0.482 | 0.11 | 2.4E-08 |
| 2 | DDI2 | 0.134 | 0.064 | 0.491 | 0.344 | 0.737 | 3.9E-34 |
| 2 | DDRGK1 | 0.112 | 0.076 | 0.444 | 0.384 | 0.237 | 1.5E-09 |
| 2 | DDT | 0.187 | 0.147 | 0.824 | 0.762 | 0.111 | 4.9E-06 |
| 2 | DDX1 | 0.179 | 0.132 | 0.749 | 0.707 | 0.102 | 1.1E-08 |
| 2 | DDX10 | 0.104 | 0.073 | 0.436 | 0.398 | 0.13 | 4.9E-08 |
| 2 | DDX17 | 0.444 | 0.414 | 3.287 | 2.864 | 0.198 | 9.2E-23 |
| 2 | DDX23 | 0.13 | 0.084 | 0.441 | 0.44 | 0.264 | 2.1E-12 |
| 2 | DDX3X | 0.476 | 0.444 | 3.714 | 3.319 | 0.162 | 1.5E-16 |
| 2 | DDX42 | 0.143 | 0.106 | 0.629 | 0.448 | 0.198 | 4.4E-11 |
| 2 | DDX44 | 0.129 | 0.099 | 0.468 | 0.417 | 0.134 | 1E-05 |
| 2 | DDX48 | 0.68 | 0.211 | 6.694 | 1.491 | 2.167 | 0 |
| 2 | DDX49 | 0.203 | 0.131 | 0.9 | 0.709 | 0.344 | 1.1E-19 |
| 2 | DDX60 | 0.648 | 0.233 | 4.913 | 1.62 | 1.601 | 0 |
| 2 | DDX60L | 0.736 | 0.264 | 6.942 | 2.009 | 1.789 | 0 |
| 2 | DECR1 | 0.342 | 0.222 | 1.789 | 1.298 | 0.463 | 1.3E-40 |
| 2 | DEF6 | 0.223 | 0.16 | 0.99 | 0.894 | 0.147 | 2.2E-12 |
| 2 | DEGS1 | 0.139 | 0.098 | 0.487 | 0.414 | 0.189 | 1.8E-09 |
| 2 | DEK | 0.449 | 0.342 | 2.424 | 2.201 | 0.198 | 3.1E-19 |
| 2 | DENND1A | 0.214 | 0.138 | 0.99 | 0.811 | 0.288 | 3.5E-21 |
| 2 | DERA | 0.127 | 0.088 | 0.418 | 0.469 | 0.142 | 1.9E-09 |
| 2 | DERL1 | 0.377 | 0.233 | 1.869 | 1.34 | 0.48 | 2.4E-47 |
| 2 | DESI2 | 0.164 | 0.114 | 0.67 | 0.604 | 0.141 | 4.4E-11 |
| 2 | DGCR6L | 0.121 | 0.084 | 0.497 | 0.428 | 0.214 | 1.8E-08 |
| 2 | DGKA | 0.114 | 0.074 | 0.462 | 0.387 | 0.244 | 9.4E-12 |
| 2 | DGLUCY | 0.193 | 0.104 | 0.868 | 0.486 | 0.466 | 1.7E-37 |
| 2 | DHX29 | 0.209 | 0.134 | 0.942 | 0.766 | 0.313 | 8.6E-21 |
| 2 | DHX30 | 0.193 | 0.132 | 0.83 | 0.684 | 0.279 | 6E-14 |
| 2 | DHX36 | 0.34 | 0.243 | 1.71 | 1.494 | 0.194 | 1.5E-14 |
| 2 | DHX48 | 0.104 | 0.023 | 0.444 | 0.117 | 1.922 | 4.1E-98 |
| 2 | DIMT1 | 0.111 | 0.066 | 0.448 | 0.349 | 0.363 | 1.2E-14 |
| 2 | DIPK1A | 0.492 | 0.307 | 2.848 | 1.98 | 0.43 | 3.4E-66 |
| 2 | DIS3 | 0.146 | 0.099 | 0.643 | 0.431 | 0.3 | 1E-16 |
| 2 | DIS3L | 0.102 | 0.068 | 0.424 | 0.344 | 0.246 | 7.9E-10 |
| 2 | DLEU2 | 0.366 | 0.214 | 1.883 | 1.34 | 0.48 | 1.4E-44 |
| 2 | DLGAP1-AS1 | 0.147 | 0.094 | 0.667 | 0.406 | 0.399 | 2.4E-21 |
| 2 | DMAC2L | 0.149 | 0.104 | 0.696 | 0.471 | 0.286 | 9.3E-14 |
| 2 | DMTF1 | 0.331 | 0.243 | 1.606 | 1.436 | 0.162 | 1.8E-14 |
| 2 | DMXL1 | 0.264 | 0.197 | 1.216 | 1.133 | 0.101 | 4.5E-11 |
| 2 | DNAJA1 | 0.409 | 0.389 | 3.141 | 2.739 | 0.198 | 3.3E-18 |
| 2 | DNAJB6 | 0.602 | 0.446 | 3.946 | 3.22 | 0.293 | 1.3E-33 |
| 2 | DNAJC10 | 0.234 | 0.173 | 1.043 | 0.961 | 0.118 | 2.3E-10 |
| 2 | DNAJC13 | 0.194 | 0.117 | 0.879 | 0.644 | 0.447 | 4.6E-26 |
| 2 | DNAJC19 | 0.172 | 0.128 | 0.742 | 0.676 | 0.134 | 2.4E-08 |
| 2 | DNAJC24 | 0.143 | 0.092 | 0.491 | 0.401 | 0.24 | 2.2E-14 |
| 2 | DNAJC4 | 0.12 | 0.082 | 0.492 | 0.444 | 0.14 | 2.8E-09 |
| 2 | DNAJC7 | 0.184 | 0.131 | 0.794 | 0.708 | 0.164 | 2.1E-11 |
| 2 | DNM2 | 0.282 | 0.197 | 1.283 | 1.177 | 0.124 | 4.6E-17 |
| 2 | DNMT1 | 0.244 | 0.177 | 1.147 | 1.004 | 0.192 | 5E-13 |
| 2 | DNPEP | 0.117 | 0.06 | 0.493 | 0.298 | 0.726 | 3.2E-27 |
| 2 | DNPH1 | 0.206 | 0.061 | 0.932 | 0.341 | 1.441 | 4E-134 |
| 2 | DOCK10 | 0.712 | 0.414 | 4.494 | 4.144 | 0.407 | 1.5E-49 |
| 2 | DOCK8 | 0.743 | 0.421 | 6.242 | 4.227 | 0.464 | 1E-108 |

| | | | | | | |
|---|---|---|---|---|---|---|
| 2 | DOCK9 | 0.149 | 0.066 | 0.674 | 0.374 | 0.849 | 4.2E-48 |
| 2 | DOP1A | 0.306 | 0.134 | 1.644 | 0.793 | 1.043 | 3E-106 |
| 2 | DPF2 | 0.104 | 0.06 | 0.414 | 0.309 | 0.423 | 2.8E-17 |
| 2 | DPP8 | 0.224 | 0.148 | 1.02 | 0.868 | 0.233 | 3.8E-14 |
| 2 | DPY30 | 0.134 | 0.092 | 0.481 | 0.494 | 0.232 | 1E-10 |
| 2 | DPYD | 0.642 | 0.49 | 4.717 | 4.014 | 0.233 | 3E-31 |
| 2 | DRAP1 | 0.76 | 0.442 | 6.444 | 3.126 | 1.046 | 2E-247 |
| 2 | DTHD1 | 0.341 | 0.243 | 1.942 | 1.626 | 0.246 | 2.8E-20 |
| 2 | DTX3L | 0.769 | 0.31 | 6.836 | 2.236 | 1.613 | 0 |
| 2 | DUSP11 | 0.233 | 0.174 | 1.014 | 0.946 | 0.101 | 1.9E-09 |
| 2 | DUSP16 | 0.208 | 0.074 | 1.002 | 0.437 | 1.198 | 1E-101 |
| 2 | DUSP22 | 0.131 | 0.082 | 0.442 | 0.426 | 0.373 | 1.5E-14 |
| 2 | DYNC1I2 | 0.174 | 0.111 | 0.741 | 0.496 | 0.313 | 4.7E-18 |
| 2 | DYNLT1 | 0.402 | 0.268 | 3.148 | 1.612 | 0.966 | 1E-134 |
| 2 | DYNLT3 | 0.17 | 0.124 | 0.743 | 0.646 | 0.201 | 1.4E-09 |
| 2 | E2F3 | 0.261 | 0.103 | 1.221 | 0.49 | 1.048 | 6E-108 |
| 2 | ECE1 | 0.161 | 0.064 | 0.749 | 0.343 | 1.104 | 1.6E-66 |
| 2 | ECH1 | 0.292 | 0.224 | 1.334 | 1.239 | 0.107 | 1.5E-09 |
| 2 | ECHDC1 | 0.222 | 0.137 | 0.973 | 0.746 | 0.383 | 8.3E-26 |
| 2 | ECHS1 | 0.131 | 0.078 | 0.446 | 0.381 | 0.419 | 4.1E-19 |
| 2 | ECPAS | 0.241 | 0.141 | 1.092 | 0.788 | 0.47 | 6.5E-34 |
| 2 | EDEM3 | 0.117 | 0.076 | 0.404 | 0.414 | 0.284 | 9.4E-12 |
| 2 | EDF1 | 0.397 | 0.293 | 2.02 | 1.682 | 0.264 | 8.7E-19 |
| 2 | EEF1AKMT2 | 0.122 | 0.083 | 0.494 | 0.449 | 0.139 | 8.8E-10 |
| 2 | EFCAB14 | 0.366 | 0.277 | 1.838 | 1.629 | 0.174 | 3.4E-14 |
| 2 | EFL1 | 0.113 | 0.074 | 0.464 | 0.388 | 0.249 | 1.5E-11 |
| 2 | EFR3A | 0.387 | 0.244 | 1.977 | 1.416 | 0.383 | 1E-36 |
| 2 | EGLN1 | 0.122 | 0.084 | 0.491 | 0.424 | 0.206 | 2.4E-09 |
| 2 | EHBP1 | 0.164 | 0.119 | 0.744 | 0.691 | 0.106 | 1.9E-09 |
| 2 | EHD4 | 0.144 | 0.089 | 0.703 | 0.483 | 0.441 | 1.3E-24 |
| 2 | EIF2AK2 | 0.849 | 0.338 | 10.613 | 2.964 | 1.84 | 0 |
| 2 | EIF2AK4 | 0.124 | 0.092 | 0.428 | 0.491 | 0.104 | 7.4E-07 |
| 2 | EIF2S1 | 0.149 | 0.113 | 0.673 | 0.481 | 0.213 | 2.2E-10 |
| 2 | EIF2S2 | 0.289 | 0.201 | 1.361 | 1.143 | 0.241 | 3.4E-19 |
| 2 | EIF2S3 | 0.216 | 0.149 | 0.973 | 0.87 | 0.161 | 4.7E-11 |
| 2 | EIF3E | 0.219 | 0.163 | 0.987 | 0.873 | 0.177 | 1.9E-10 |
| 2 | EIF3M | 0.246 | 0.172 | 1.104 | 0.904 | 0.289 | 6.5E-17 |
| 2 | EIF4B | 0.263 | 0.19 | 1.224 | 1.111 | 0.14 | 9E-14 |
| 2 | EIF4E | 0.131 | 0.083 | 0.431 | 0.413 | 0.364 | 1.5E-14 |
| 2 | EIF4E2 | 0.12 | 0.079 | 0.402 | 0.408 | 0.301 | 1.9E-11 |
| 2 | EIF4E3 | 0.177 | 0.084 | 0.776 | 0.464 | 0.737 | 2.3E-48 |
| 2 | EIF4EBP1 | 0.14 | 0.048 | 0.468 | 0.303 | 0.904 | 1.3E-40 |
| 2 | EIF4G1 | 0.198 | 0.137 | 0.837 | 0.734 | 0.187 | 1.1E-13 |
| 2 | ELAVL1 | 0.138 | 0.094 | 0.474 | 0.479 | 0.261 | 1.5E-11 |
| 2 | ELF1 | 0.722 | 0.404 | 4.643 | 3.838 | 0.446 | 1.7E-84 |
| 2 | ELK3 | 0.249 | 0.143 | 1.243 | 0.814 | 0.61 | 1.9E-46 |
| 2 | ELK4 | 0.217 | 0.122 | 0.992 | 0.662 | 0.483 | 1.3E-37 |
| 2 | ELL | 0.131 | 0.072 | 0.447 | 0.391 | 0.484 | 1.1E-23 |
| 2 | ELMOD2 | 0.101 | 0.062 | 0.4 | 0.327 | 0.292 | 7.9E-13 |
| 2 | ELP4 | 0.211 | 0.147 | 0.992 | 0.889 | 0.148 | 4.8E-14 |
| 2 | EMC2 | 0.114 | 0.062 | 0.466 | 0.326 | 0.414 | 1.1E-21 |
| 2 | EMC6 | 0.127 | 0.093 | 0.41 | 0.449 | 0.184 | 2.2E-07 |
| 2 | EML4 | 0.624 | 0.444 | 4.43 | 3.468 | 0.343 | 8.5E-44 |
| 2 | EMSY | 0.184 | 0.116 | 0.776 | 0.642 | 0.242 | 7.5E-20 |

| | | | | | | | |
|---|---|---|---|---|---|---|---|
| 2 | ENDOD1 | 0.179 | 0.076 | 0.836 | 0.396 | 1.079 | 4.2E-64 |
| 2 | ENOSF1 | 0.138 | 0.081 | 0.484 | 0.436 | 0.422 | 2.4E-20 |
| 2 | ENOX2 | 0.131 | 0.081 | 0.444 | 0.438 | 0.312 | 6.9E-16 |
| 2 | ENPP4 | 0.298 | 0.209 | 1.407 | 1.201 | 0.229 | 2.4E-18 |
| 2 | ENTPD1-AS1 | 0.142 | 0.099 | 0.693 | 0.444 | 0.321 | 3.7E-14 |
| 2 | EOMES | 0.199 | 0.116 | 0.877 | 0.634 | 0.466 | 6.7E-29 |
| 2 | EPB41 | 0.732 | 0.476 | 4.721 | 4.878 | 0.23 | 3.1E-31 |
| 2 | EPC2 | 0.264 | 0.197 | 1.194 | 1.113 | 0.103 | 4.8E-11 |
| 2 | EPDR1 | 0.144 | 0.104 | 0.731 | 0.607 | 0.268 | 2.4E-12 |
| 2 | EPRS | 0.209 | 0.143 | 0.934 | 0.849 | 0.122 | 3.3E-10 |
| 2 | EPS14 | 0.426 | 0.381 | 3.037 | 2.469 | 0.242 | 2.3E-28 |
| 2 | EPSTI1 | 0.87 | 0.332 | 11.863 | 3.008 | 1.98 | 0 |
| 2 | ERAP1 | 0.391 | 0.231 | 1.991 | 1.329 | 0.484 | 4.3E-60 |
| 2 | ERAP2 | 0.407 | 0.237 | 2.194 | 1.46 | 0.488 | 1E-66 |
| 2 | ERC1 | 0.243 | 0.167 | 1.147 | 1.002 | 0.194 | 3.9E-17 |
| 2 | ERCC4 | 0.26 | 0.197 | 1.229 | 1.129 | 0.122 | 7.2E-10 |
| 2 | ERGIC2 | 0.144 | 0.109 | 0.64 | 0.482 | 0.137 | 4.7E-10 |
| 2 | ERI1 | 0.101 | 0.063 | 0.413 | 0.328 | 0.332 | 3.3E-12 |
| 2 | ERICH3 | 0.101 | 0.011 | 0.469 | 0.072 | 2.99 | 2E-149 |
| 2 | ERLEC1 | 0.301 | 0.214 | 1.436 | 1.214 | 0.241 | 2.5E-17 |
| 2 | ERMP1 | 0.111 | 0.071 | 0.46 | 0.38 | 0.274 | 3.7E-12 |
| 2 | ERP44 | 0.242 | 0.17 | 1.131 | 0.928 | 0.284 | 4.5E-20 |
| 2 | ESD | 0.113 | 0.048 | 0.466 | 0.296 | 0.642 | 8E-24 |
| 2 | ETFA | 0.216 | 0.142 | 0.944 | 0.743 | 0.344 | 1.5E-20 |
| 2 | ETFB | 0.132 | 0.09 | 0.416 | 0.48 | 0.106 | 2.7E-10 |
| 2 | ETNK1 | 0.413 | 0.26 | 2.133 | 1.602 | 0.413 | 1.7E-46 |
| 2 | ETV6 | 0.222 | 0.109 | 1.037 | 0.641 | 0.696 | 1.1E-44 |
| 2 | EVI2A | 0.273 | 0.173 | 1.313 | 0.993 | 0.403 | 2.7E-29 |
| 2 | EVI2B | 0.339 | 0.249 | 1.689 | 1.429 | 0.143 | 8.4E-12 |
| 2 | EVL | 0.634 | 0.473 | 4.348 | 3.429 | 0.346 | 1.6E-38 |
| 2 | EWSR1 | 0.33 | 0.234 | 1.433 | 1.344 | 0.18 | 1.9E-18 |
| 2 | EXOC1 | 0.193 | 0.114 | 0.838 | 0.62 | 0.434 | 1.7E-26 |
| 2 | EXOC2 | 0.314 | 0.176 | 1.411 | 1.001 | 0.494 | 3.8E-44 |
| 2 | EXOC3 | 0.112 | 0.068 | 0.443 | 0.343 | 0.363 | 3.7E-14 |
| 2 | EXOC4 | 0.18 | 0.106 | 0.774 | 0.46 | 0.468 | 3.1E-24 |
| 2 | EXOSC10 | 0.149 | 0.109 | 0.669 | 0.486 | 0.193 | 8.4E-12 |
| 2 | EXOSC8 | 0.178 | 0.136 | 0.792 | 0.724 | 0.129 | 1.3E-07 |
| 2 | EXOSC9 | 0.141 | 0.094 | 0.499 | 0.494 | 0.276 | 4.5E-12 |
| 2 | EXT1 | 0.103 | 0.063 | 0.427 | 0.394 | 0.113 | 8.6E-13 |
| 2 | EYA3 | 0.14 | 0.088 | 0.628 | 0.479 | 0.393 | 9.9E-22 |
| 2 | EZH1 | 0.222 | 0.16 | 0.979 | 0.902 | 0.117 | 1.2E-11 |
| 2 | F2R | 0.402 | 0.343 | 3.074 | 2.444 | 0.267 | 5E-31 |
| 2 | FAF2 | 0.168 | 0.12 | 0.704 | 0.642 | 0.133 | 3.4E-10 |
| 2 | FAM102A | 0.126 | 0.047 | 0.431 | 0.308 | 0.784 | 1.4E-38 |
| 2 | FAM102B | 0.146 | 0.102 | 0.684 | 0.472 | 0.246 | 9E-14 |
| 2 | FAM104A | 0.129 | 0.081 | 0.422 | 0.414 | 0.331 | 4.4E-15 |
| 2 | FAM111A | 0.268 | 0.196 | 1.244 | 1.118 | 0.166 | 2.7E-13 |
| 2 | FAM118A | 0.166 | 0.104 | 0.749 | 0.471 | 0.392 | 1.4E-19 |
| 2 | FAM120AOS | 0.176 | 0.12 | 0.767 | 0.637 | 0.268 | 1.2E-13 |
| 2 | FAM120B | 0.242 | 0.172 | 1.072 | 0.96 | 0.149 | 8E-14 |
| 2 | FAM136A | 0.137 | 0.102 | 0.466 | 0.423 | 0.114 | 8E-07 |
| 2 | FAM13B | 0.333 | 0.242 | 1.487 | 1.446 | 0.124 | 1.2E-14 |
| 2 | FAM160B1 | 0.199 | 0.141 | 0.843 | 0.766 | 0.144 | 7.2E-12 |
| 2 | FAM192A | 0.142 | 0.108 | 0.647 | 0.473 | 0.174 | 7.8E-10 |

| | | | | | | | |
|---|---|---|---|---|---|---|---|
| 2 | FAM193A | 0.128 | 0.089 | 0.424 | 0.488 | 0.102 | 4.2E-09 |
| 2 | FAM214A | 0.312 | 0.22 | 1.43 | 1.332 | 0.2 | 2.5E-18 |
| 2 | FAM217B | 0.149 | 0.104 | 0.62 | 0.464 | 0.134 | 7.1E-10 |
| 2 | FAM228B | 0.164 | 0.123 | 0.714 | 0.66 | 0.116 | 6.4E-08 |
| 2 | FAM3C | 0.127 | 0.084 | 0.44 | 0.434 | 0.34 | 7.5E-12 |
| 2 | FAM40A | 0.124 | 0.093 | 0.417 | 0.48 | 0.107 | 4E-06 |
| 2 | FAM76B | 0.184 | 0.14 | 0.824 | 0.744 | 0.129 | 4.9E-08 |
| 2 | FAM98B | 0.188 | 0.124 | 0.781 | 0.669 | 0.224 | 8.2E-16 |
| 2 | FANCA | 0.109 | 0.034 | 0.474 | 0.188 | 1.34 | 1.4E-62 |
| 2 | FAS | 0.134 | 0.06 | 0.464 | 0.313 | 0.842 | 1.4E-43 |
| 2 | FASLG | 0.374 | 0.123 | 1.937 | 0.7 | 1.469 | 8E-227 |
| 2 | FBXL4 | 0.119 | 0.076 | 0.469 | 0.426 | 0.14 | 1.3E-12 |
| 2 | FBXO28 | 0.106 | 0.068 | 0.444 | 0.342 | 0.367 | 1.4E-11 |
| 2 | FBXO34 | 0.392 | 0.244 | 2.173 | 1.466 | 0.473 | 3.3E-48 |
| 2 | FBXO38 | 0.171 | 0.102 | 0.734 | 0.447 | 0.399 | 4.9E-23 |
| 2 | FBXO6 | 0.214 | 0.088 | 0.998 | 0.447 | 1.147 | 7.2E-86 |
| 2 | FBXO9 | 0.244 | 0.184 | 1.098 | 1.016 | 0.113 | 2.4E-12 |
| 2 | FBXW11 | 0.323 | 0.236 | 1.414 | 1.382 | 0.131 | 1E-14 |
| 2 | FCHO2 | 0.149 | 0.114 | 0.709 | 0.641 | 0.124 | 3.7E-09 |
| 2 | FCHSD2 | 0.274 | 0.186 | 1.347 | 1.132 | 0.24 | 7.5E-21 |
| 2 | FDPS | 0.132 | 0.091 | 0.422 | 0.464 | 0.17 | 7.8E-10 |
| 2 | FDX1 | 0.132 | 0.099 | 0.442 | 0.494 | 0.129 | 1.5E-06 |
| 2 | FECH | 0.123 | 0.043 | 0.428 | 0.27 | 0.967 | 1.9E-41 |
| 2 | FEM1B | 0.101 | 0.076 | 0.424 | 0.386 | 0.136 | 3.7E-05 |
| 2 | FER | 0.148 | 0.113 | 0.719 | 0.641 | 0.144 | 1.4E-09 |
| 2 | FEZ2 | 0.138 | 0.084 | 0.462 | 0.442 | 0.314 | 1.7E-17 |
| 2 | FGD3 | 0.373 | 0.26 | 1.844 | 1.483 | 0.229 | 2E-23 |
| 2 | FGFR1OP2 | 0.367 | 0.243 | 1.826 | 1.441 | 0.331 | 4.7E-32 |
| 2 | FHL3 | 0.129 | 0.066 | 0.444 | 0.329 | 0.743 | 1.1E-29 |
| 2 | FIBP | 0.134 | 0.088 | 0.47 | 0.438 | 0.379 | 2.5E-13 |
| 2 | FKBP14 | 0.107 | 0.067 | 0.44 | 0.34 | 0.362 | 7.7E-13 |
| 2 | FKBP3 | 0.133 | 0.094 | 0.44 | 0.489 | 0.17 | 8.9E-09 |
| 2 | FKBP4 | 0.214 | 0.106 | 0.963 | 0.491 | 0.704 | 4.5E-43 |
| 2 | FLII | 0.101 | 0.069 | 0.406 | 0.362 | 0.166 | 1.4E-08 |
| 2 | FLOT1 | 0.112 | 0.074 | 0.448 | 0.388 | 0.206 | 4.1E-10 |
| 2 | FMR1 | 0.207 | 0.122 | 0.912 | 0.684 | 0.412 | 1.6E-29 |
| 2 | FNDC3A | 0.376 | 0.232 | 1.922 | 1.421 | 0.434 | 4.4E-44 |
| 2 | FNTA | 0.19 | 0.11 | 0.807 | 0.49 | 0.443 | 9.1E-29 |
| 2 | FOCAD | 0.143 | 0.081 | 0.648 | 0.447 | 0.424 | 2E-30 |
| 2 | FOPNL | 0.132 | 0.094 | 0.44 | 0.488 | 0.174 | 4E-08 |
| 2 | FOXJ3 | 0.308 | 0.233 | 1.46 | 1.343 | 0.109 | 8.9E-12 |
| 2 | FOXN2 | 0.3 | 0.19 | 1.434 | 1.072 | 0.421 | 1.7E-32 |
| 2 | FOXP1 | 0.678 | 0.466 | 4.844 | 3.681 | 0.399 | 4.9E-68 |
| 2 | FRA10AC1 | 0.127 | 0.091 | 0.424 | 0.474 | 0.142 | 4.6E-08 |
| 2 | FRYL | 0.486 | 0.371 | 2.746 | 2.419 | 0.124 | 1.3E-16 |
| 2 | FTH1 | 0.974 | 0.914 | 26.334 | 20 | 0.397 | 3.3E-11 |
| 2 | FTX | 0.349 | 0.217 | 1.814 | 1.363 | 0.412 | 1.4E-44 |
| 2 | FUBP3 | 0.134 | 0.089 | 0.463 | 0.47 | 0.249 | 8.3E-13 |
| 2 | FYB1 | 0.748 | 0.464 | 6.391 | 4.646 | 0.46 | 4.4E-73 |
| 2 | FYN | 0.888 | 0.744 | 10.919 | 8.806 | 0.31 | 2.4E-64 |
| 2 | FYTTD1 | 0.369 | 0.227 | 1.836 | 1.308 | 0.489 | 8.4E-46 |
| 2 | GABARAPL1 | 0.309 | 0.214 | 1.481 | 1.249 | 0.233 | 3.9E-20 |
| 2 | GABARAPL2 | 0.321 | 0.218 | 1.464 | 1.2 | 0.287 | 3.6E-24 |
| 2 | GABPA | 0.144 | 0.096 | 0.668 | 0.403 | 0.41 | 9.5E-19 |

| | | | | | | | |
|---|---|---|---|---|---|---|---|
| 2 | GALNT10 | 0.164 | 0.108 | 0.687 | 0.617 | 0.144 | 2.4E-14 |
| 2 | GALNT2 | 0.124 | 0.084 | 0.434 | 0.447 | 0.226 | 1.7E-10 |
| 2 | GALNT3 | 0.213 | 0.146 | 0.943 | 0.804 | 0.244 | 6.4E-16 |
| 2 | GAPVD1 | 0.184 | 0.13 | 0.783 | 0.721 | 0.12 | 3.7E-11 |
| 2 | GARS-DT | 0.182 | 0.127 | 0.807 | 0.717 | 0.171 | 1.5E-12 |
| 2 | GART | 0.138 | 0.092 | 0.484 | 0.473 | 0.306 | 1.5E-12 |
| 2 | GATA3 | 0.277 | 0.184 | 1.406 | 1.117 | 0.332 | 4.8E-24 |
| 2 | GATAD1 | 0.124 | 0.088 | 0.423 | 0.466 | 0.164 | 3.5E-08 |
| 2 | GATAD2B | 0.341 | 0.243 | 1.626 | 1.496 | 0.12 | 4.4E-18 |
| 2 | GBP1 | 0.404 | 0.138 | 4.661 | 0.974 | 2.249 | 0 |
| 2 | GBP2 | 0.644 | 0.331 | 4.836 | 2.322 | 1.048 | 8E-216 |
| 2 | GBP3 | 0.27 | 0.112 | 1.278 | 0.614 | 1.046 | 2E-102 |
| 2 | GBP4 | 0.888 | 0.406 | 18.347 | 4.276 | 1.798 | 0 |
| 2 | GBP4 | 0.742 | 0.294 | 7.647 | 2.144 | 1.834 | 0 |
| 2 | GCC2 | 0.464 | 0.34 | 2.682 | 2.389 | 0.167 | 1.8E-18 |
| 2 | GCH1 | 0.167 | 0.107 | 0.776 | 0.481 | 0.419 | 3.1E-18 |
| 2 | GCHFR | 0.124 | 0.082 | 0.471 | 0.444 | 0.349 | 3.1E-12 |
| 2 | GCNT1 | 0.232 | 0.141 | 1.164 | 0.884 | 0.397 | 4.2E-29 |
| 2 | GDI2 | 0.426 | 0.278 | 2.123 | 1.487 | 0.419 | 9.9E-42 |
| 2 | GFM1 | 0.123 | 0.084 | 0.403 | 0.433 | 0.218 | 6.1E-10 |
| 2 | GFOD1 | 0.194 | 0.114 | 0.916 | 0.667 | 0.447 | 8.9E-29 |
| 2 | GGA2 | 0.392 | 0.2 | 2.031 | 1.202 | 0.747 | 3.5E-94 |
| 2 | GHITM | 0.342 | 0.274 | 1.706 | 1.468 | 0.122 | 4E-10 |
| 2 | GIGYF2 | 0.233 | 0.144 | 1.019 | 0.876 | 0.218 | 2.2E-19 |
| 2 | GIMAP1 | 0.33 | 0.116 | 1.687 | 0.674 | 1.324 | 6E-174 |
| 2 | GIMAP2 | 0.249 | 0.101 | 1.212 | 0.431 | 1.191 | 3E-113 |
| 2 | GIMAP4 | 0.471 | 0.188 | 2.669 | 1.179 | 1.179 | 2E-213 |
| 2 | GIMAP4 | 0.394 | 0.187 | 2.2 | 1.116 | 0.98 | 4E-123 |
| 2 | GIMAP6 | 0.174 | 0.066 | 0.789 | 0.341 | 1.168 | 6.9E-79 |
| 2 | GIMAP7 | 0.834 | 0.483 | 9.194 | 4.246 | 1.114 | 0 |
| 2 | GIMAP8 | 0.162 | 0.044 | 0.72 | 0.284 | 1.338 | 7.9E-88 |
| 2 | GIT2 | 0.228 | 0.148 | 1.018 | 0.894 | 0.188 | 3E-14 |
| 2 | GKAP1 | 0.103 | 0.06 | 0.446 | 0.314 | 0.404 | 1.5E-16 |
| 2 | GLCCI1 | 0.343 | 0.23 | 1.946 | 1.442 | 0.336 | 3.4E-33 |
| 2 | GLIPR2 | 0.49 | 0.364 | 2.808 | 2.344 | 0.244 | 2.2E-22 |
| 2 | GLO1 | 0.231 | 0.166 | 1.01 | 0.88 | 0.198 | 4.3E-13 |
| 2 | GLOD4 | 0.142 | 0.111 | 0.619 | 0.464 | 0.13 | 4.3E-08 |
| 2 | GLRX | 0.249 | 0.181 | 1.201 | 0.988 | 0.282 | 4.3E-14 |
| 2 | GLT8D1 | 0.108 | 0.069 | 0.442 | 0.34 | 0.334 | 6.8E-12 |
| 2 | GMCL1 | 0.197 | 0.144 | 0.872 | 0.784 | 0.141 | 2.7E-10 |
| 2 | GMDS-DT | 0.309 | 0.178 | 1.492 | 1.123 | 0.404 | 3.7E-48 |
| 2 | GNG4 | 0.486 | 0.326 | 2.726 | 1.944 | 0.487 | 1.5E-48 |
| 2 | GNPAT | 0.11 | 0.076 | 0.443 | 0.401 | 0.173 | 2.1E-08 |
| 2 | GOLGA1 | 0.129 | 0.084 | 0.441 | 0.448 | 0.271 | 1.4E-12 |
| 2 | GOLGA2 | 0.103 | 0.072 | 0.414 | 0.373 | 0.141 | 7.3E-08 |
| 2 | GOLGA4 | 0.118 | 0.08 | 0.497 | 0.408 | 0.284 | 2.1E-10 |
| 2 | GOLGA7 | 0.284 | 0.213 | 1.31 | 1.184 | 0.144 | 4.8E-12 |
| 2 | GOLIM4 | 0.113 | 0.084 | 0.414 | 0.46 | 0.162 | 2.4E-06 |
| 2 | GOPC | 0.243 | 0.142 | 1.044 | 0.79 | 0.403 | 6.7E-34 |
| 2 | GPALPP1 | 0.122 | 0.079 | 0.41 | 0.412 | 0.307 | 1.4E-12 |
| 2 | GPATCH2 | 0.119 | 0.083 | 0.493 | 0.446 | 0.143 | 8E-09 |
| 2 | GPATCH2L | 0.24 | 0.162 | 1.134 | 0.904 | 0.327 | 9.7E-24 |
| 2 | GPBP1 | 0.642 | 0.398 | 4.337 | 2.692 | 0.688 | 3E-114 |
| 2 | GPR144 | 0.246 | 0.068 | 1.384 | 0.396 | 1.806 | 9E-200 |

| | | | | | | | |
|---|---|---|---|---|---|---|---|
| 2 | GPR171 | 0.184 | 0.041 | 0.871 | 0.282 | 1.628 | 3E-136 |
| 2 | GPR174 | 0.26 | 0.162 | 1.194 | 0.912 | 0.39 | 1E-29 |
| 2 | GPR180 | 0.118 | 0.038 | 0.434 | 0.203 | 1.393 | 1.2E-67 |
| 2 | GPR89B | 0.133 | 0.089 | 0.444 | 0.472 | 0.232 | 8.6E-12 |
| 2 | GRIPAP1 | 0.213 | 0.128 | 0.947 | 0.69 | 0.448 | 2.9E-28 |
| 2 | GRSF1 | 0.191 | 0.129 | 0.822 | 0.669 | 0.296 | 2.9E-14 |
| 2 | GSDMD | 0.119 | 0.063 | 0.407 | 0.318 | 0.674 | 4.4E-24 |
| 2 | GSTO1 | 0.181 | 0.114 | 0.749 | 0.607 | 0.322 | 2.3E-19 |
| 2 | GTF2A1 | 0.198 | 0.121 | 0.843 | 0.641 | 0.373 | 3.8E-24 |
| 2 | GTF2F1 | 0.187 | 0.126 | 0.801 | 0.644 | 0.291 | 7E-16 |
| 2 | GTF2F2 | 0.144 | 0.088 | 0.62 | 0.474 | 0.384 | 1.1E-18 |
| 2 | GTF2H1 | 0.179 | 0.11 | 0.778 | 0.481 | 0.42 | 8.7E-22 |
| 2 | GTF2I | 0.414 | 0.274 | 2.207 | 1.701 | 0.376 | 3.1E-37 |
| 2 | GTF2IRD2B | 0.104 | 0.069 | 0.428 | 0.343 | 0.279 | 1.2E-09 |
| 2 | GTF3C3 | 0.112 | 0.068 | 0.477 | 0.349 | 0.443 | 2.4E-14 |
| 2 | GXYLT1 | 0.124 | 0.087 | 0.412 | 0.448 | 0.163 | 8.6E-09 |
| 2 | GYG1 | 0.199 | 0.133 | 0.862 | 0.7 | 0.3 | 4.9E-17 |
| 2 | HACD3 | 0.11 | 0.072 | 0.433 | 0.383 | 0.174 | 9.2E-11 |
| 2 | HADHB | 0.221 | 0.136 | 0.966 | 0.739 | 0.388 | 3.5E-26 |
| 2 | HAGH | 0.107 | 0.067 | 0.429 | 0.33 | 0.378 | 6.9E-13 |
| 2 | HAT1 | 0.141 | 0.094 | 0.484 | 0.493 | 0.246 | 1E-11 |
| 2 | HAVCR2 | 0.169 | 0.114 | 0.819 | 0.641 | 0.344 | 8.7E-14 |
| 2 | HCP4 | 0.129 | 0.068 | 0.407 | 0.369 | 0.46 | 3.8E-26 |
| 2 | HDAC1 | 0.303 | 0.223 | 1.417 | 1.263 | 0.164 | 6E-14 |
| 2 | HDAC2 | 0.217 | 0.162 | 0.974 | 0.872 | 0.16 | 4.5E-10 |
| 2 | HDAC4 | 0.123 | 0.079 | 0.418 | 0.44 | 0.201 | 3.1E-13 |
| 2 | HDAC8 | 0.17 | 0.104 | 0.772 | 0.494 | 0.378 | 2.5E-20 |
| 2 | HDLBP | 0.196 | 0.136 | 0.867 | 0.732 | 0.244 | 4E-14 |
| 2 | HEATR1 | 0.144 | 0.094 | 0.631 | 0.41 | 0.306 | 2.4E-14 |
| 2 | HECTD4 | 0.212 | 0.141 | 0.942 | 0.807 | 0.224 | 1.3E-17 |
| 2 | HEG1 | 0.116 | 0.061 | 0.479 | 0.337 | 0.406 | 1.1E-23 |
| 2 | HELB | 0.249 | 0.102 | 1.301 | 0.446 | 1.243 | 4E-111 |
| 2 | HELQ | 0.117 | 0.077 | 0.49 | 0.397 | 0.302 | 3E-11 |
| 2 | HELZ2 | 0.194 | 0.048 | 0.941 | 0.249 | 1.863 | 1E-162 |
| 2 | HENMT1 | 0.114 | 0.077 | 0.477 | 0.391 | 0.287 | 1.9E-10 |
| 2 | HERC3 | 0.138 | 0.097 | 0.486 | 0.442 | 0.113 | 1.4E-09 |
| 2 | HERC4 | 0.739 | 0.193 | 7.94 | 1.463 | 2.344 | 0 |
| 2 | HERC4 | 0.343 | 0.208 | 1.714 | 1.242 | 0.444 | 2.2E-40 |
| 2 | HERC6 | 0.447 | 0.142 | 3.672 | 0.942 | 1.948 | 0 |
| 2 | HERPUD2 | 0.344 | 0.249 | 1.767 | 1.494 | 0.242 | 2.8E-19 |
| 2 | HEXD | 0.193 | 0.09 | 0.849 | 0.482 | 0.834 | 3.1E-44 |
| 2 | HIBADH | 0.144 | 0.091 | 0.669 | 0.498 | 0.426 | 8.2E-22 |
| 2 | HIBCH | 0.147 | 0.11 | 0.684 | 0.613 | 0.16 | 2.4E-10 |
| 2 | HIF1A | 0.848 | 0.433 | 9.477 | 4.711 | 1.008 | 6E-284 |
| 2 | HIF1A-AS3 | 0.144 | 0.076 | 0.601 | 0.41 | 0.441 | 2.8E-30 |
| 2 | HINT1 | 0.449 | 0.432 | 3.43 | 2.932 | 0.226 | 2.5E-20 |
| 2 | HINT2 | 0.142 | 0.112 | 0.632 | 0.484 | 0.114 | 1.1E-07 |
| 2 | HIP1 | 0.184 | 0.118 | 0.807 | 0.691 | 0.222 | 4.7E-18 |
| 2 | HIPK3 | 0.263 | 0.191 | 1.188 | 1.083 | 0.132 | 4E-13 |
| 2 | HIST2H2AC | 0.284 | 0.197 | 1.311 | 1.102 | 0.24 | 4.7E-19 |
| 2 | HIVEP1 | 0.223 | 0.102 | 1.024 | 0.48 | 0.822 | 1.2E-66 |
| 2 | HIVEP2 | 0.307 | 0.166 | 1.449 | 1.014 | 0.426 | 4E-47 |
| 2 | HK1 | 0.148 | 0.093 | 0.634 | 0.498 | 0.34 | 3.5E-17 |
| 2 | HLA-A | 0.996 | 0.933 | 34.724 | 24.482 | 0.444 | 5E-283 |

| | | | | | | | |
|---|---|---|---|---|---|---|---|
| 2 | HLA-B | 0.998 | 0.963 | 40.431 | 32.902 | 0.301 | 2.2E-92 |
| 2 | HLA-C | 0.48 | 0.448 | 3.624 | 3.141 | 0.202 | 4.9E-20 |
| 2 | HLA-E | 0.978 | 0.908 | 19.477 | 17.393 | 0.163 | 4.8E-22 |
| 2 | HLA-F | 0.694 | 0.406 | 4.048 | 3.776 | 0.419 | 3E-44 |
| 2 | HLTF | 0.168 | 0.106 | 0.741 | 0.49 | 0.348 | 1E-18 |
| 2 | HM13 | 0.187 | 0.101 | 0.864 | 0.43 | 0.706 | 1.3E-37 |
| 2 | HMBOX1 | 0.143 | 0.097 | 0.644 | 0.464 | 0.214 | 1.1E-14 |
| 2 | HMG20A | 0.123 | 0.07 | 0.413 | 0.362 | 0.403 | 1.1E-20 |
| 2 | HMGA1 | 0.119 | 0.064 | 0.483 | 0.323 | 0.48 | 1E-22 |
| 2 | HMGB1 | 0.868 | 0.769 | 9.181 | 8.499 | 0.111 | 3.8E-11 |
| 2 | HMGN3 | 0.414 | 0.314 | 2.189 | 1.899 | 0.204 | 8.5E-16 |
| 2 | HMGN4 | 0.2 | 0.142 | 0.846 | 0.748 | 0.176 | 1.6E-12 |
| 2 | HNRNPD | 0.443 | 0.344 | 2.417 | 2.244 | 0.107 | 2.4E-12 |
| 2 | HNRNPF | 0.414 | 0.302 | 2.988 | 1.841 | 0.698 | 9.2E-93 |
| 2 | HNRNPH2 | 0.187 | 0.136 | 0.789 | 0.697 | 0.18 | 4.5E-10 |
| 2 | HNRNPK | 0.747 | 0.608 | 4.891 | 4.19 | 0.183 | 7.4E-21 |
| 2 | HNRNPUL2 | 0.164 | 0.11 | 0.691 | 0.476 | 0.263 | 4.4E-14 |
| 2 | HOXB2 | 0.116 | 0.069 | 0.483 | 0.374 | 0.371 | 1E-16 |
| 2 | HPS3 | 0.243 | 0.177 | 1.144 | 0.996 | 0.212 | 6.4E-16 |
| 2 | HS2ST1 | 0.163 | 0.098 | 0.731 | 0.446 | 0.394 | 1.7E-21 |
| 2 | HSBP1 | 0.147 | 0.088 | 0.619 | 0.444 | 0.476 | 4E-20 |
| 2 | HSD17B11 | 0.162 | 0.106 | 0.672 | 0.476 | 0.223 | 6.9E-14 |
| 2 | HSH2D | 0.444 | 0.243 | 2.441 | 1.486 | 0.774 | 4.8E-94 |
| 2 | HSPA4 | 0.273 | 0.166 | 1.203 | 0.928 | 0.374 | 7.1E-33 |
| 2 | HSPA8 | 0.878 | 0.646 | 12.832 | 6.443 | 0.992 | 9E-232 |
| 2 | HSPB11 | 0.174 | 0.12 | 0.742 | 0.634 | 0.247 | 5E-13 |
| 2 | HTATIP2 | 0.173 | 0.094 | 0.741 | 0.483 | 0.617 | 7.7E-32 |
| 2 | IAH1 | 0.104 | 0.07 | 0.407 | 0.344 | 0.197 | 2.5E-09 |
| 2 | IARS2 | 0.224 | 0.172 | 1 | 0.93 | 0.106 | 3.4E-08 |
| 2 | ICAM2 | 0.172 | 0.114 | 0.788 | 0.492 | 0.414 | 1.8E-16 |
| 2 | ICE1 | 0.304 | 0.201 | 1.411 | 1.177 | 0.36 | 2.3E-27 |
| 2 | ICE2 | 0.137 | 0.092 | 0.461 | 0.482 | 0.219 | 2.2E-11 |
| 2 | IDH2 | 0.344 | 0.193 | 1.698 | 1.071 | 0.664 | 2.3E-63 |
| 2 | IER3IP1 | 0.238 | 0.17 | 1.03 | 0.89 | 0.211 | 1.3E-13 |
| 2 | IFI16 | 0.888 | 0.494 | 10.871 | 4.384 | 1.31 | 0 |
| 2 | IFI34 | 0.449 | 0.114 | 2.643 | 0.644 | 2.036 | 0 |
| 2 | IFI44 | 0.69 | 0.174 | 6.02 | 1.339 | 2.168 | 0 |
| 2 | IFI44L | 0.781 | 0.187 | 9.83 | 1.771 | 2.473 | 0 |
| 2 | IFI6 | 0.733 | 0.177 | 8.026 | 1.427 | 2.491 | 0 |
| 2 | IFIH1 | 0.444 | 0.148 | 3.944 | 0.937 | 2.074 | 0 |
| 2 | IFIT1 | 0.732 | 0.146 | 9.149 | 1.368 | 2.743 | 0 |
| 2 | IFIT2 | 0.499 | 0.174 | 4.96 | 1.396 | 2.094 | 0 |
| 2 | IFIT3 | 0.832 | 0.238 | 17.242 | 2.823 | 2.612 | 0 |
| 2 | IFIT4 | 0.471 | 0.144 | 2.718 | 0.911 | 1.477 | 0 |
| 2 | IFITM2 | 0.909 | 0.744 | 12.494 | 9.924 | 0.332 | 2.4E-44 |
| 2 | IFITM3 | 0.943 | 0.712 | 27.871 | 9.26 | 1.49 | 0 |
| 2 | IFNAR1 | 0.198 | 0.134 | 0.874 | 0.726 | 0.269 | 3.8E-14 |
| 2 | IGF1R | 0.193 | 0.121 | 0.838 | 0.73 | 0.198 | 1.4E-20 |
| 2 | IGFLR1 | 0.176 | 0.113 | 0.74 | 0.477 | 0.349 | 1.5E-18 |
| 2 | IK | 0.346 | 0.274 | 1.774 | 1.626 | 0.126 | 1.9E-11 |
| 2 | IKBIP | 0.102 | 0.062 | 0.419 | 0.32 | 0.388 | 6.3E-14 |
| 2 | IKBKB | 0.113 | 0.07 | 0.444 | 0.381 | 0.244 | 9E-14 |
| 2 | IKZF2 | 0.213 | 0.129 | 1.031 | 0.786 | 0.391 | 7.4E-28 |
| 2 | IKZF3 | 0.468 | 0.444 | 3.743 | 3.371 | 0.144 | 8.4E-14 |

| | | | | | | | |
|---|---|---|---|---|---|---|---|
| 2 | IL10RA | 0.443 | 0.431 | 3.398 | 2.996 | 0.182 | 4.2E-16 |
| 2 | IL12RB2 | 0.162 | 0.09 | 0.813 | 0.473 | 0.404 | 4.7E-29 |
| 2 | IL14 | 0.124 | 0.043 | 0.464 | 0.24 | 1.237 | 1.5E-64 |
| 2 | IL14RA | 0.142 | 0.037 | 0.631 | 0.198 | 1.672 | 1E-109 |
| 2 | IL16 | 0.414 | 0.314 | 2.929 | 1.976 | 0.467 | 4.8E-73 |
| 2 | IL18RAP | 0.246 | 0.166 | 1.191 | 0.974 | 0.29 | 7.1E-20 |
| 2 | IL2RG | 0.894 | 0.788 | 10.413 | 9.636 | 0.126 | 4.3E-13 |
| 2 | IL6ST | 0.142 | 0.102 | 0.669 | 0.467 | 0.239 | 7.1E-13 |
| 2 | ILK | 0.218 | 0.129 | 0.929 | 0.687 | 0.434 | 1.1E-29 |
| 2 | IMMP2L | 0.239 | 0.149 | 1.172 | 1.021 | 0.199 | 8.4E-20 |
| 2 | IMMT | 0.276 | 0.19 | 1.304 | 1.048 | 0.316 | 4.2E-20 |
| 2 | IMP3 | 0.2 | 0.134 | 0.844 | 0.697 | 0.294 | 3.7E-16 |
| 2 | IMPA1 | 0.16 | 0.104 | 0.664 | 0.46 | 0.248 | 6.7E-14 |
| 2 | INPP4D | 0.467 | 0.344 | 2.646 | 2.326 | 0.191 | 3.7E-21 |
| 2 | INTS12 | 0.104 | 0.073 | 0.431 | 0.4 | 0.107 | 1.4E-07 |
| 2 | INTS6L | 0.114 | 0.077 | 0.471 | 0.418 | 0.172 | 6.2E-10 |
| 2 | IP6K1 | 0.111 | 0.076 | 0.462 | 0.417 | 0.147 | 9.4E-09 |
| 2 | IPCEF1 | 0.364 | 0.13 | 1.961 | 0.741 | 1.384 | 3E-194 |
| 2 | IPO8 | 0.12 | 0.063 | 0.476 | 0.332 | 0.42 | 1.3E-24 |
| 2 | IQGAP2 | 0.667 | 0.449 | 4.838 | 4.487 | 0.108 | 2.2E-11 |
| 2 | IRAK2 | 0.11 | 0.049 | 0.449 | 0.274 | 0.706 | 2.1E-34 |
| 2 | IRAK4 | 0.284 | 0.192 | 1.317 | 1.082 | 0.284 | 1.4E-22 |
| 2 | IREB2 | 0.226 | 0.141 | 0.944 | 0.833 | 0.198 | 3.1E-18 |
| 2 | IRF1 | 0.919 | 0.801 | 14.489 | 11.449 | 0.436 | 6.4E-68 |
| 2 | IRF2 | 0.488 | 0.234 | 2.83 | 1.404 | 1.01 | 5E-144 |
| 2 | IRF3 | 0.177 | 0.123 | 0.766 | 0.683 | 0.167 | 4.1E-12 |
| 2 | IRF4 | 0.111 | 0.03 | 0.423 | 0.171 | 1.614 | 4.4E-81 |
| 2 | IRF7 | 0.633 | 0.177 | 4.642 | 1.164 | 1.998 | 0 |
| 2 | IRF9 | 0.472 | 0.213 | 2.486 | 1.304 | 0.988 | 2E-163 |
| 2 | ISCU | 0.337 | 0.24 | 1.644 | 1.468 | 0.173 | 7.8E-14 |
| 2 | ISG14 | 0.777 | 0.222 | 9.484 | 1.808 | 2.406 | 0 |
| 2 | ISG20 | 0.861 | 0.322 | 12.443 | 2.494 | 2.274 | 0 |
| 2 | ITGA1 | 0.204 | 0.07 | 0.988 | 0.424 | 1.218 | 4E-108 |
| 2 | ITGAE | 0.364 | 0.176 | 1.984 | 1.061 | 0.903 | 5E-103 |
| 2 | ITGB3BP | 0.129 | 0.08 | 0.429 | 0.447 | 0.244 | 8.2E-14 |
| 2 | ITM2B | 0.941 | 0.783 | 14.486 | 9.449 | 0.611 | 9E-187 |
| 2 | ITM2C | 0.149 | 0.063 | 0.799 | 0.384 | 1.049 | 2.7E-63 |
| 2 | ITPR1 | 0.239 | 0.14 | 1.104 | 0.789 | 0.484 | 1.3E-34 |
| 2 | ITPR2 | 0.404 | 0.349 | 2.961 | 2.421 | 0.291 | 6.3E-36 |
| 2 | IVD | 0.126 | 0.088 | 0.426 | 0.46 | 0.193 | 6.1E-09 |
| 2 | IWS1 | 0.217 | 0.137 | 0.98 | 0.742 | 0.402 | 1.7E-24 |
| 2 | JADE2 | 0.162 | 0.1 | 0.664 | 0.441 | 0.298 | 4.5E-19 |
| 2 | JAK2 | 0.407 | 0.143 | 2.271 | 0.892 | 1.347 | 5E-206 |
| 2 | JAK3 | 0.172 | 0.089 | 0.724 | 0.477 | 0.603 | 1.8E-36 |
| 2 | JKAMP | 0.168 | 0.086 | 0.678 | 0.442 | 0.484 | 2.3E-36 |
| 2 | JMJD1C | 0.441 | 0.414 | 3.478 | 2.979 | 0.223 | 1.1E-24 |
| 2 | JPX | 0.222 | 0.141 | 1.021 | 0.811 | 0.332 | 8.9E-23 |
| 2 | KANSL1L | 0.109 | 0.049 | 0.44 | 0.263 | 0.774 | 1.2E-33 |
| 2 | KANSL3 | 0.164 | 0.106 | 0.692 | 0.484 | 0.241 | 1.8E-16 |
| 2 | KARS | 0.138 | 0.101 | 0.474 | 0.428 | 0.123 | 1.1E-07 |
| 2 | KAT6A | 0.409 | 0.374 | 2.971 | 2.476 | 0.206 | 1.5E-24 |
| 2 | KATNA1 | 0.103 | 0.07 | 0.424 | 0.374 | 0.178 | 8.8E-09 |
| 2 | KATNBL1 | 0.148 | 0.098 | 0.62 | 0.412 | 0.277 | 1.3E-13 |
| 2 | KDELR1 | 0.199 | 0.143 | 0.873 | 0.792 | 0.141 | 1.2E-07 |

| | | | | | | | |
|---|---|---|---|---|---|---|---|
| 2 | KDM3B | 0.194 | 0.124 | 0.844 | 0.696 | 0.279 | 4E-20 |
| 2 | KDM4B | 0.284 | 0.168 | 1.343 | 0.974 | 0.461 | 8.5E-40 |
| 2 | KDM7A | 0.304 | 0.207 | 1.494 | 1.236 | 0.274 | 2.4E-22 |
| 2 | KEAP1 | 0.163 | 0.077 | 0.707 | 0.407 | 0.798 | 4.5E-44 |
| 2 | KHDRBS1 | 0.418 | 0.317 | 2.127 | 1.921 | 0.147 | 7.9E-14 |
| 2 | KIAA0040 | 0.129 | 0.07 | 0.444 | 0.347 | 0.673 | 6.2E-24 |
| 2 | KIAA0319L | 0.171 | 0.093 | 0.744 | 0.412 | 0.439 | 1.8E-32 |
| 2 | KIAA0486 | 0.177 | 0.129 | 0.786 | 0.727 | 0.113 | 2E-09 |
| 2 | KIAA2026 | 0.344 | 0.247 | 1.691 | 1.414 | 0.16 | 4.5E-18 |
| 2 | KIF13A | 0.144 | 0.073 | 0.674 | 0.424 | 0.666 | 2.4E-41 |
| 2 | KIF21A | 0.184 | 0.126 | 0.847 | 0.709 | 0.274 | 2.4E-14 |
| 2 | KIN | 0.183 | 0.137 | 0.79 | 0.731 | 0.111 | 3.4E-08 |
| 2 | KLHDC2 | 0.168 | 0.12 | 0.707 | 0.642 | 0.138 | 8E-10 |
| 2 | KLHL20 | 0.184 | 0.129 | 0.82 | 0.739 | 0.149 | 1.5E-12 |
| 2 | KLHL4 | 0.19 | 0.088 | 0.837 | 0.488 | 0.777 | 4.3E-44 |
| 2 | KLHL6 | 0.229 | 0.071 | 1.026 | 0.404 | 1.343 | 8E-143 |
| 2 | KLHL7 | 0.11 | 0.066 | 0.44 | 0.347 | 0.376 | 1.3E-14 |
| 2 | KLRC1 | 0.363 | 0.247 | 2.644 | 2.133 | 0.316 | 6.7E-28 |
| 2 | KLRD1 | 0.892 | 0.791 | 11.434 | 10.104 | 0.191 | 2.9E-19 |
| 2 | KMT2A | 0.429 | 0.294 | 2.344 | 1.913 | 0.3 | 3.4E-31 |
| 2 | KMT4B | 0.276 | 0.189 | 1.34 | 1.096 | 0.29 | 3.8E-20 |
| 2 | KPNA1 | 0.473 | 0.292 | 2.74 | 1.921 | 0.417 | 8.1E-64 |
| 2 | KPNA4 | 0.26 | 0.143 | 1.244 | 0.802 | 0.644 | 4.5E-48 |
| 2 | KPNB1 | 0.423 | 0.276 | 3.097 | 1.72 | 0.848 | 5E-134 |
| 2 | KRAS | 0.398 | 0.284 | 2.002 | 1.746 | 0.189 | 1.5E-20 |
| 2 | KRCC1 | 0.213 | 0.126 | 0.949 | 0.68 | 0.497 | 6.2E-30 |
| 2 | KRIT1 | 0.221 | 0.148 | 0.989 | 0.823 | 0.264 | 4.7E-18 |
| 2 | KTN1 | 0.462 | 0.419 | 3.442 | 2.9 | 0.247 | 3.9E-26 |
| 2 | KYAT3 | 0.129 | 0.069 | 0.444 | 0.36 | 0.494 | 1.1E-24 |
| 2 | L3MBTL3 | 0.134 | 0.077 | 0.618 | 0.42 | 0.447 | 7.8E-23 |
| 2 | L3MBTL4 | 0.286 | 0.14 | 1.413 | 0.872 | 0.696 | 1.6E-72 |
| 2 | LACTB | 0.123 | 0.06 | 0.421 | 0.319 | 0.708 | 4.4E-31 |
| 2 | LAGE3 | 0.104 | 0.072 | 0.404 | 0.346 | 0.186 | 1.8E-08 |
| 2 | LAP3 | 0.418 | 0.172 | 2.373 | 0.991 | 1.26 | 3E-186 |
| 2 | LARP4B | 0.144 | 0.103 | 0.499 | 0.449 | 0.101 | 1.1E-08 |
| 2 | LARP7 | 0.2 | 0.139 | 0.896 | 0.811 | 0.144 | 3.4E-13 |
| 2 | LBH | 0.742 | 0.402 | 6.941 | 4.38 | 0.666 | 3E-127 |
| 2 | LBR | 0.448 | 0.423 | 3.436 | 2.943 | 0.224 | 4.8E-23 |
| 2 | LCMT1 | 0.102 | 0.069 | 0.409 | 0.34 | 0.224 | 1E-08 |
| 2 | LCORL | 0.263 | 0.168 | 1.272 | 0.996 | 0.344 | 4.1E-27 |
| 2 | LCP2 | 0.442 | 0.327 | 2.444 | 2.112 | 0.268 | 3.1E-24 |
| 2 | LDB2 | 0.128 | 0.048 | 0.641 | 0.284 | 1.177 | 7.2E-48 |
| 2 | LDHA | 0.414 | 0.424 | 3.143 | 2.93 | 0.106 | 3.1E-08 |
| 2 | LDLRAD4 | 0.268 | 0.138 | 1.362 | 0.839 | 0.699 | 1.1E-49 |
| 2 | LEMD2 | 0.101 | 0.064 | 0.413 | 0.329 | 0.328 | 4.3E-11 |
| 2 | LEMD3 | 0.167 | 0.12 | 0.69 | 0.639 | 0.109 | 8.3E-10 |
| 2 | LEO1 | 0.139 | 0.083 | 0.497 | 0.442 | 0.401 | 4.1E-19 |
| 2 | LEPROTL1 | 0.39 | 0.28 | 1.963 | 1.667 | 0.236 | 1.1E-21 |
| 2 | LGALS8 | 0.204 | 0.112 | 0.887 | 0.61 | 0.439 | 9.4E-37 |
| 2 | LGALS9 | 0.244 | 0.046 | 1.282 | 0.286 | 2.166 | 7E-241 |
| 2 | LINC00278 | 0.137 | 0.088 | 0.649 | 0.423 | 0.334 | 1.6E-14 |
| 2 | LINC00476 | 0.109 | 0.071 | 0.449 | 0.408 | 0.172 | 1.5E-10 |
| 2 | LINC00623 | 0.442 | 0.3 | 2.414 | 1.94 | 0.374 | 4.5E-41 |
| 2 | LINC00667 | 0.101 | 0.067 | 0.414 | 0.337 | 0.3 | 3.8E-10 |

| | | | | | | | |
|---|---|---|---|---|---|---|---|
| 2 | LINC00996 | 0.339 | 0.096 | 2.404 | 0.782 | 1.68 | 7E-240 |
| 2 | LINC01138 | 0.641 | 0.434 | 4.414 | 3.187 | 0.403 | 1.9E-78 |
| 2 | LINC01684 | 0.114 | 0.049 | 0.484 | 0.264 | 0.874 | 8.5E-39 |
| 2 | LINC02100 | 0.144 | 0.046 | 0.644 | 0.324 | 0.994 | 9.2E-62 |
| 2 | LINC02328 | 0.207 | 0.124 | 0.966 | 0.742 | 0.381 | 4.9E-27 |
| 2 | LINC02384 | 0.192 | 0.129 | 0.974 | 0.764 | 0.349 | 4.1E-17 |
| 2 | LINGO2 | 0.418 | 0.31 | 4.024 | 2.836 | 0.824 | 1E-100 |
| 2 | LINS1 | 0.106 | 0.072 | 0.419 | 0.374 | 0.16 | 3.9E-09 |
| 2 | LIPA | 0.409 | 0.28 | 2.869 | 1.744 | 0.718 | 1E-106 |
| 2 | LMAN1 | 0.278 | 0.199 | 1.24 | 1.09 | 0.187 | 1.5E-14 |
| 2 | LMBR1 | 0.274 | 0.197 | 1.299 | 1.166 | 0.146 | 2E-14 |
| 2 | LMNB1 | 0.144 | 0.086 | 0.619 | 0.442 | 0.443 | 4.5E-20 |
| 2 | LMO4 | 0.176 | 0.114 | 0.793 | 0.614 | 0.368 | 3.3E-17 |
| 2 | LNPEP | 0.762 | 0.424 | 6.261 | 4.174 | 0.484 | 5E-116 |
| 2 | LPGAT1 | 0.447 | 0.312 | 2.47 | 1.964 | 0.331 | 7.9E-32 |
| 2 | LPIN2 | 0.44 | 0.206 | 2.476 | 1.232 | 1.064 | 1E-146 |
| 2 | LPXN | 0.248 | 0.191 | 1.173 | 1.064 | 0.141 | 1.4E-11 |
| 2 | LRBA | 0.632 | 0.489 | 4.41 | 3.864 | 0.19 | 8.2E-24 |
| 2 | LRCH1 | 0.214 | 0.122 | 0.944 | 0.678 | 0.492 | 1.3E-34 |
| 2 | LRCH3 | 0.294 | 0.171 | 1.374 | 1.002 | 0.446 | 2.5E-42 |
| 2 | LRIF1 | 0.227 | 0.131 | 1.072 | 0.718 | 0.479 | 3.9E-34 |
| 2 | LRP10 | 0.147 | 0.11 | 0.647 | 0.471 | 0.202 | 1.4E-10 |
| 2 | LSM10 | 0.126 | 0.092 | 0.404 | 0.444 | 0.179 | 2.4E-07 |
| 2 | LSM2 | 0.178 | 0.124 | 0.724 | 0.632 | 0.197 | 4.5E-12 |
| 2 | LSM6 | 0.149 | 0.092 | 0.631 | 0.463 | 0.446 | 4E-18 |
| 2 | LUZP1 | 0.244 | 0.139 | 1.194 | 0.803 | 0.474 | 2.7E-40 |
| 2 | LY6E | 0.896 | 0.488 | 12.468 | 4.214 | 1.464 | 0 |
| 2 | LYPLAL1 | 0.127 | 0.079 | 0.422 | 0.416 | 0.328 | 3.4E-14 |
| 2 | LYSMD2 | 0.329 | 0.108 | 1.462 | 0.481 | 1.427 | 1E-194 |
| 2 | LYST | 0.848 | 0.443 | 11.449 | 4.884 | 0.96 | 4E-276 |
| 2 | MAGED1 | 0.123 | 0.089 | 0.437 | 0.487 | 0.14 | 1.4E-07 |
| 2 | MAGED2 | 0.118 | 0.066 | 0.492 | 0.33 | 0.478 | 1.4E-20 |
| 2 | MAML2 | 0.368 | 0.184 | 2.098 | 1.224 | 0.774 | 7.8E-92 |
| 2 | MAP2K1 | 0.144 | 0.094 | 0.627 | 0.423 | 0.262 | 3.7E-18 |
| 2 | MAP3K4 | 0.389 | 0.226 | 2.016 | 1.418 | 0.408 | 6.4E-49 |
| 2 | MAP4 | 0.39 | 0.273 | 1.942 | 1.646 | 0.246 | 2.7E-24 |
| 2 | MAP4K3 | 0.111 | 0.066 | 0.449 | 0.344 | 0.372 | 1.1E-14 |
| 2 | MAP7D1 | 0.129 | 0.094 | 0.449 | 0.49 | 0.163 | 1.8E-07 |
| 2 | MAPK1 | 0.464 | 0.426 | 3.479 | 2.947 | 0.234 | 2.5E-24 |
| 2 | MAPKAP1 | 0.266 | 0.197 | 1.21 | 1.127 | 0.103 | 1.1E-11 |
| 2 | MARF1 | 0.212 | 0.124 | 0.94 | 0.694 | 0.437 | 7.8E-30 |
| 2 | MARK2 | 0.13 | 0.093 | 0.432 | 0.489 | 0.12 | 4.3E-08 |
| 2 | MAT2B | 0.434 | 0.324 | 3.136 | 2.023 | 0.632 | 4.2E-84 |
| 2 | MBD4 | 0.462 | 0.362 | 3.637 | 2.496 | 0.487 | 8.4E-69 |
| 2 | MBNL1 | 0.933 | 0.841 | 12.446 | 11.318 | 0.137 | 3E-18 |
| 2 | MBTD1 | 0.144 | 0.099 | 0.608 | 0.447 | 0.144 | 1.1E-10 |
| 2 | MCCC2 | 0.107 | 0.069 | 0.432 | 0.348 | 0.271 | 2.1E-11 |
| 2 | MCFD2 | 0.122 | 0.087 | 0.407 | 0.443 | 0.164 | 4E-08 |
| 2 | MCL1 | 0.403 | 0.324 | 2.982 | 2.138 | 0.48 | 2E-47 |
| 2 | MCM3 | 0.116 | 0.064 | 0.47 | 0.324 | 0.431 | 4E-20 |
| 2 | MCM9 | 0.206 | 0.141 | 0.934 | 0.869 | 0.104 | 2.5E-10 |
| 2 | MCMBP | 0.109 | 0.077 | 0.432 | 0.39 | 0.14 | 9E-08 |
| 2 | MCPH1 | 0.248 | 0.193 | 1.23 | 1.124 | 0.129 | 6.3E-11 |
| 2 | MCRS1 | 0.103 | 0.07 | 0.406 | 0.344 | 0.193 | 7.7E-09 |

| | | | | | | | |
|---|---|---|---|---|---|---|---|
| 2 | MCUB | 0.384 | 0.168 | 1.989 | 1.003 | 0.988 | 1E-134 |
| 2 | MDH1 | 0.234 | 0.144 | 1.004 | 0.827 | 0.281 | 1.2E-20 |
| 2 | ME2 | 0.341 | 0.191 | 1.707 | 1.068 | 0.676 | 2E-62 |
| 2 | MEA1 | 0.16 | 0.124 | 0.678 | 0.632 | 0.101 | 8.8E-06 |
| 2 | MED1 | 0.132 | 0.098 | 0.467 | 0.41 | 0.142 | 4.7E-07 |
| 2 | MED13 | 0.309 | 0.194 | 1.424 | 1.136 | 0.327 | 1E-32 |
| 2 | MED13L | 0.427 | 0.316 | 2.314 | 2.104 | 0.137 | 1.5E-18 |
| 2 | MED6 | 0.111 | 0.078 | 0.436 | 0.406 | 0.104 | 1.3E-07 |
| 2 | MEF2A | 0.272 | 0.204 | 1.323 | 1.203 | 0.137 | 3.6E-11 |
| 2 | MEGF9 | 0.197 | 0.149 | 0.924 | 0.838 | 0.142 | 1.2E-08 |
| 2 | MESD | 0.261 | 0.179 | 1.178 | 0.991 | 0.249 | 2.5E-18 |
| 2 | METTL14 | 0.192 | 0.131 | 0.843 | 0.742 | 0.182 | 8.1E-14 |
| 2 | METTL14 | 0.137 | 0.088 | 0.466 | 0.464 | 0.287 | 4.3E-14 |
| 2 | METTL23 | 0.124 | 0.09 | 0.4 | 0.444 | 0.17 | 8.1E-08 |
| 2 | METTL24 | 0.101 | 0.064 | 0.403 | 0.342 | 0.194 | 9.2E-11 |
| 2 | METTL4 | 0.144 | 0.101 | 0.493 | 0.431 | 0.16 | 4.3E-10 |
| 2 | METTL4 | 0.108 | 0.04 | 0.449 | 0.26 | 0.82 | 1.5E-30 |
| 2 | MFAP1 | 0.117 | 0.049 | 0.483 | 0.308 | 0.646 | 4.4E-27 |
| 2 | MFSD1 | 0.121 | 0.081 | 0.488 | 0.414 | 0.236 | 7E-11 |
| 2 | MFSD6 | 0.177 | 0.128 | 0.741 | 0.698 | 0.104 | 9.4E-10 |
| 2 | MFSD8 | 0.111 | 0.063 | 0.446 | 0.334 | 0.441 | 1.3E-18 |
| 2 | MGA | 0.182 | 0.124 | 0.82 | 0.689 | 0.241 | 1.8E-14 |
| 2 | MGAT1 | 0.167 | 0.101 | 0.688 | 0.432 | 0.372 | 3.5E-21 |
| 2 | MIA3 | 0.327 | 0.221 | 1.621 | 1.307 | 0.311 | 4.2E-26 |
| 2 | MIB1 | 0.16 | 0.094 | 0.676 | 0.422 | 0.374 | 3.1E-22 |
| 2 | MICOS13 | 0.141 | 0.114 | 0.623 | 0.473 | 0.121 | 1E-06 |
| 2 | MINDY2 | 0.248 | 0.14 | 1.112 | 0.86 | 0.37 | 3.3E-31 |
| 2 | MIR181A2HG | 0.324 | 0.141 | 1.773 | 0.874 | 1.02 | 2E-114 |
| 2 | MITD1 | 0.208 | 0.141 | 0.911 | 0.771 | 0.24 | 7.3E-16 |
| 2 | MKRN1 | 0.188 | 0.129 | 0.849 | 0.683 | 0.314 | 1.2E-14 |
| 2 | MLEC | 0.178 | 0.124 | 0.748 | 0.634 | 0.237 | 7.2E-12 |
| 2 | MLKL | 0.134 | 0.048 | 0.494 | 0.308 | 0.947 | 8.6E-48 |
| 2 | MLLT3 | 0.418 | 0.244 | 3.377 | 1.786 | 0.919 | 9E-149 |
| 2 | MMADHC | 0.274 | 0.186 | 1.243 | 1.011 | 0.309 | 1.4E-21 |
| 2 | MNAT1 | 0.304 | 0.214 | 1.492 | 1.261 | 0.243 | 9.1E-20 |
| 2 | MOB1A | 0.646 | 0.418 | 4.187 | 3.844 | 0.124 | 4.1E-13 |
| 2 | MOB1B | 0.141 | 0.087 | 0.602 | 0.441 | 0.414 | 4.2E-17 |
| 2 | MOCS2 | 0.102 | 0.069 | 0.426 | 0.34 | 0.281 | 6E-09 |
| 2 | MORC3 | 0.416 | 0.239 | 2.247 | 1.437 | 0.644 | 9.1E-72 |
| 2 | MOSPD2 | 0.13 | 0.07 | 0.449 | 0.348 | 0.683 | 1.9E-26 |
| 2 | MOV10 | 0.113 | 0.041 | 0.468 | 0.224 | 1.046 | 4.1E-43 |
| 2 | MPG | 0.114 | 0.08 | 0.444 | 0.41 | 0.114 | 8.9E-08 |
| 2 | MPHOSPH10 | 0.124 | 0.087 | 0.438 | 0.44 | 0.248 | 1.7E-09 |
| 2 | MPHOSPH9 | 0.192 | 0.132 | 0.813 | 0.734 | 0.147 | 1.2E-13 |
| 2 | MPLKIP | 0.121 | 0.082 | 0.487 | 0.411 | 0.244 | 4.9E-10 |
| 2 | MPP7 | 0.22 | 0.14 | 1.013 | 0.843 | 0.247 | 3.1E-17 |
| 2 | MR1 | 0.133 | 0.08 | 0.474 | 0.422 | 0.444 | 3.2E-18 |
| 2 | MRFAP1 | 0.246 | 0.136 | 1.161 | 0.71 | 0.709 | 1.4E-43 |
| 2 | MRFAP1L1 | 0.134 | 0.074 | 0.464 | 0.382 | 0.464 | 9.4E-24 |
| 2 | MRPL13 | 0.131 | 0.092 | 0.434 | 0.47 | 0.189 | 2.2E-09 |
| 2 | MRPL16 | 0.174 | 0.111 | 0.737 | 0.486 | 0.33 | 1.8E-18 |
| 2 | MRPL19 | 0.111 | 0.078 | 0.464 | 0.394 | 0.236 | 6.4E-08 |
| 2 | MRPL20 | 0.191 | 0.134 | 0.827 | 0.698 | 0.244 | 5E-13 |
| 2 | MRPL34 | 0.141 | 0.106 | 0.603 | 0.43 | 0.187 | 6.1E-10 |

| | | | | | | | |
|---|---|---|---|---|---|---|---|
| 2 | MRPL34 | 0.11 | 0.048 | 0.434 | 0.294 | 0.462 | 9.1E-23 |
| 2 | MRPL41 | 0.146 | 0.1 | 0.494 | 0.403 | 0.24 | 2.6E-11 |
| 2 | MRPL42 | 0.117 | 0.089 | 0.496 | 0.439 | 0.177 | 1.2E-05 |
| 2 | MRPL43 | 0.174 | 0.128 | 0.71 | 0.648 | 0.131 | 1E-08 |
| 2 | MRPL47 | 0.214 | 0.149 | 0.903 | 0.838 | 0.109 | 2.4E-09 |
| 2 | MRPS10 | 0.143 | 0.103 | 0.494 | 0.423 | 0.184 | 4.5E-12 |
| 2 | MRPS18B | 0.102 | 0.061 | 0.404 | 0.299 | 0.432 | 3.4E-14 |
| 2 | MRPS18C | 0.214 | 0.12 | 0.934 | 0.634 | 0.447 | 4.8E-37 |
| 2 | MRPS21 | 0.277 | 0.207 | 1.263 | 1.126 | 0.166 | 6.6E-12 |
| 2 | MRPS23 | 0.11 | 0.067 | 0.428 | 0.332 | 0.364 | 8.6E-14 |
| 2 | MRPS27 | 0.127 | 0.086 | 0.432 | 0.439 | 0.278 | 4.6E-11 |
| 2 | MRPS28 | 0.124 | 0.079 | 0.433 | 0.409 | 0.381 | 1E-13 |
| 2 | MRPS31 | 0.208 | 0.126 | 0.886 | 0.669 | 0.404 | 6.6E-26 |
| 2 | MRPS4 | 0.204 | 0.136 | 0.873 | 0.73 | 0.248 | 1.8E-17 |
| 2 | MRPS6 | 0.431 | 0.247 | 2.366 | 1.494 | 0.47 | 2.3E-64 |
| 2 | MRPS9 | 0.142 | 0.11 | 0.679 | 0.486 | 0.211 | 6.5E-09 |
| 2 | MSH2 | 0.129 | 0.078 | 0.424 | 0.416 | 0.334 | 1.3E-16 |
| 2 | MSL3 | 0.219 | 0.109 | 1.003 | 0.487 | 0.774 | 8.6E-44 |
| 2 | MSMO1 | 0.129 | 0.088 | 0.444 | 0.449 | 0.246 | 1.5E-10 |
| 2 | MSN | 0.83 | 0.643 | 7.896 | 6.067 | 0.38 | 1.2E-61 |
| 2 | MT2A | 0.868 | 0.429 | 19.344 | 4.482 | 2.078 | 0 |
| 2 | MTERF1 | 0.14 | 0.091 | 0.494 | 0.406 | 0.233 | 1.4E-13 |
| 2 | MTF2 | 0.162 | 0.111 | 0.687 | 0.604 | 0.184 | 2.2E-12 |
| 2 | MTG1 | 0.246 | 0.129 | 1.084 | 0.699 | 0.633 | 5E-41 |
| 2 | MTHFD2 | 0.178 | 0.082 | 0.834 | 0.426 | 0.968 | 1E-44 |
| 2 | MTHFS | 0.103 | 0.062 | 0.429 | 0.31 | 0.469 | 4.5E-14 |
| 2 | MTIF2 | 0.101 | 0.068 | 0.437 | 0.363 | 0.27 | 3.7E-09 |
| 2 | MTIF3 | 0.21 | 0.139 | 0.909 | 0.743 | 0.272 | 4.3E-18 |
| 2 | MTLN | 0.119 | 0.082 | 0.482 | 0.407 | 0.242 | 2.7E-09 |
| 2 | MTOR | 0.113 | 0.084 | 0.494 | 0.441 | 0.133 | 1.9E-06 |
| 2 | MVP | 0.108 | 0.068 | 0.44 | 0.344 | 0.314 | 3.9E-12 |
| 2 | MX1 | 0.908 | 0.282 | 22.396 | 3.496 | 2.679 | 0 |
| 2 | MX2 | 0.646 | 0.144 | 4.732 | 1.144 | 2.324 | 0 |
| 2 | MYCBP | 0.149 | 0.081 | 0.623 | 0.399 | 0.644 | 4.8E-29 |
| 2 | MYCBP2 | 0.698 | 0.494 | 4.294 | 3.783 | 0.484 | 2.8E-71 |
| 2 | MYD88 | 0.138 | 0.042 | 0.611 | 0.241 | 1.281 | 1.3E-63 |
| 2 | MZT2B | 0.178 | 0.139 | 0.746 | 0.704 | 0.104 | 2.3E-06 |
| 2 | N4BP1 | 0.413 | 0.232 | 3.004 | 1.49 | 1.012 | 9E-184 |
| 2 | N4BP2L1 | 0.234 | 0.134 | 1.047 | 0.733 | 0.427 | 1E-34 |
| 2 | N4BP2L2 | 0.428 | 0.372 | 3.086 | 2.424 | 0.289 | 1.1E-34 |
| 2 | NAA20 | 0.138 | 0.084 | 0.441 | 0.423 | 0.381 | 2.6E-17 |
| 2 | NAA40 | 0.47 | 0.303 | 2.482 | 1.944 | 0.402 | 1.2E-48 |
| 2 | NADK | 0.114 | 0.069 | 0.402 | 0.348 | 0.426 | 1.3E-16 |
| 2 | NAE1 | 0.173 | 0.126 | 0.74 | 0.684 | 0.113 | 1.8E-09 |
| 2 | NAP1L4 | 0.46 | 0.396 | 3.34 | 2.64 | 0.334 | 1.1E-37 |
| 2 | NAPA | 0.238 | 0.146 | 1.074 | 0.861 | 0.318 | 2.9E-22 |
| 2 | NARS | 0.22 | 0.14 | 0.971 | 0.82 | 0.244 | 2.2E-16 |
| 2 | NBN | 0.392 | 0.217 | 2.046 | 1.264 | 0.694 | 2.2E-76 |
| 2 | NBPF14 | 0.169 | 0.124 | 0.747 | 0.674 | 0.147 | 3E-08 |
| 2 | NBPF19 | 0.234 | 0.173 | 1.082 | 0.991 | 0.127 | 9.1E-11 |
| 2 | NBPF26 | 0.213 | 0.149 | 0.912 | 0.817 | 0.149 | 3.1E-13 |
| 2 | NBR1 | 0.268 | 0.19 | 1.217 | 1.039 | 0.228 | 4.7E-16 |
| 2 | NCAM1 | 0.311 | 0.229 | 1.786 | 1.42 | 0.232 | 3.5E-16 |
| 2 | NCBP2 | 0.13 | 0.091 | 0.419 | 0.461 | 0.172 | 4.9E-09 |

| | | | | | | | |
|---|---|---|---|---|---|---|---|
| 2 | NCBP3 | 0.444 | 0.283 | 2.472 | 1.824 | 0.494 | 2.1E-47 |
| 2 | NCK1 | 0.331 | 0.217 | 1.447 | 1.242 | 0.317 | 8.1E-30 |
| 2 | NCOA2 | 0.441 | 0.343 | 3.277 | 2.347 | 0.474 | 1.3E-64 |
| 2 | NCOA3 | 0.39 | 0.19 | 2.046 | 1.161 | 0.824 | 8E-107 |
| 2 | NCOA4 | 0.271 | 0.18 | 1.217 | 0.974 | 0.32 | 1.9E-23 |
| 2 | NCOA7 | 0.291 | 0.224 | 1.476 | 1.347 | 0.131 | 2.5E-10 |
| 2 | NCOR1 | 0.478 | 0.44 | 3.491 | 3.221 | 0.147 | 3.7E-17 |
| 2 | NCR1 | 0.398 | 0.276 | 2.081 | 1.643 | 0.332 | 1.4E-28 |
| 2 | NDRG3 | 0.272 | 0.207 | 1.227 | 1.144 | 0.101 | 7.1E-10 |
| 2 | NDUFA7 | 0.232 | 0.171 | 0.986 | 0.89 | 0.148 | 4.4E-11 |
| 2 | NDUFA9 | 0.148 | 0.092 | 0.611 | 0.489 | 0.322 | 4.7E-17 |
| 2 | NDUFAB1 | 0.203 | 0.147 | 0.834 | 0.744 | 0.143 | 7.2E-11 |
| 2 | NDUFAF3 | 0.197 | 0.134 | 0.826 | 0.694 | 0.241 | 4.7E-14 |
| 2 | NDUFB3 | 0.118 | 0.084 | 0.464 | 0.41 | 0.181 | 2E-07 |
| 2 | NDUFB4 | 0.241 | 0.18 | 1.087 | 0.944 | 0.203 | 6.2E-14 |
| 2 | NDUFB4 | 0.181 | 0.114 | 0.783 | 0.481 | 0.43 | 1.2E-19 |
| 2 | NDUFC1 | 0.237 | 0.16 | 1.047 | 0.846 | 0.291 | 1.4E-18 |
| 2 | NDUFS1 | 0.224 | 0.149 | 1.017 | 0.801 | 0.344 | 4.1E-20 |
| 2 | NDUFS2 | 0.148 | 0.099 | 0.637 | 0.416 | 0.303 | 7.7E-18 |
| 2 | NDUFS4 | 0.133 | 0.086 | 0.444 | 0.447 | 0.243 | 1.3E-13 |
| 2 | NDUFV2 | 0.384 | 0.247 | 1.843 | 1.432 | 0.372 | 2.9E-38 |
| 2 | NECAP2 | 0.124 | 0.093 | 0.414 | 0.474 | 0.119 | 1.4E-06 |
| 2 | NEK1 | 0.264 | 0.172 | 1.244 | 0.983 | 0.339 | 3.6E-24 |
| 2 | NEK7 | 0.482 | 0.303 | 2.66 | 1.946 | 0.444 | 1.1E-44 |
| 2 | NEK9 | 0.124 | 0.084 | 0.42 | 0.438 | 0.247 | 3.9E-11 |
| 2 | NEO1 | 0.107 | 0.076 | 0.47 | 0.402 | 0.227 | 1.5E-07 |
| 2 | NEPRO | 0.132 | 0.083 | 0.446 | 0.437 | 0.323 | 1.8E-14 |
| 2 | NF1 | 0.409 | 0.239 | 2.247 | 1.407 | 0.482 | 8.9E-66 |
| 2 | NFATC2IP | 0.179 | 0.12 | 0.747 | 0.648 | 0.204 | 1E-14 |
| 2 | NFE2L1 | 0.128 | 0.084 | 0.431 | 0.433 | 0.292 | 8.2E-12 |
| 2 | NFYB | 0.101 | 0.064 | 0.414 | 0.337 | 0.301 | 4.3E-11 |
| 2 | NFYC | 0.111 | 0.073 | 0.442 | 0.392 | 0.206 | 3E-10 |
| 2 | NGDN | 0.107 | 0.077 | 0.434 | 0.394 | 0.137 | 3.4E-07 |
| 2 | NHLRC2 | 0.113 | 0.066 | 0.484 | 0.349 | 0.43 | 1.1E-17 |
| 2 | NHLRC3 | 0.116 | 0.074 | 0.49 | 0.381 | 0.362 | 1E-12 |
| 2 | NIFK | 0.11 | 0.08 | 0.441 | 0.408 | 0.144 | 1.6E-06 |
| 2 | NIN | 0.329 | 0.243 | 1.71 | 1.487 | 0.202 | 3.5E-16 |
| 2 | NIPAL3 | 0.107 | 0.048 | 0.446 | 0.303 | 0.447 | 3.9E-21 |
| 2 | NIPBL | 0.644 | 0.403 | 4.323 | 3.911 | 0.144 | 2.7E-20 |
| 2 | NLK | 0.124 | 0.074 | 0.413 | 0.418 | 0.297 | 4.3E-17 |
| 2 | NLRC3 | 0.292 | 0.198 | 1.364 | 1.162 | 0.231 | 2.8E-21 |
| 2 | NLRC4 | 0.378 | 0.177 | 1.914 | 1.083 | 0.821 | 2E-110 |
| 2 | NME7 | 0.107 | 0.043 | 0.444 | 0.284 | 0.682 | 3E-26 |
| 2 | NMI | 0.412 | 0.128 | 2.28 | 0.748 | 1.607 | 1E-280 |
| 2 | NMRK1 | 0.321 | 0.168 | 1.477 | 0.962 | 0.713 | 6.6E-70 |
| 2 | NMT1 | 0.161 | 0.117 | 0.672 | 0.607 | 0.144 | 2.8E-09 |
| 2 | NMT2 | 0.213 | 0.126 | 0.942 | 0.689 | 0.466 | 9.9E-30 |
| 2 | NOD1 | 0.102 | 0.043 | 0.433 | 0.273 | 0.664 | 3.4E-23 |
| 2 | NOL11 | 0.102 | 0.066 | 0.403 | 0.343 | 0.232 | 7.1E-11 |
| 2 | NOL7 | 0.223 | 0.16 | 1.026 | 0.861 | 0.243 | 4.3E-13 |
| 2 | NOL8 | 0.16 | 0.103 | 0.703 | 0.462 | 0.322 | 1.5E-16 |
| 2 | NOP10 | 0.308 | 0.222 | 1.421 | 1.229 | 0.209 | 1.7E-16 |
| 2 | NOP14 | 0.131 | 0.094 | 0.466 | 0.499 | 0.183 | 4.9E-08 |
| 2 | NOSIP | 0.144 | 0.096 | 0.493 | 0.482 | 0.301 | 3.5E-13 |

| | | | | | | | |
|---|---|---|---|---|---|---|---|
| 2 | NOTCH2 | 0.238 | 0.142 | 1.081 | 0.789 | 0.444 | 1E-31 |
| 2 | NPAT | 0.318 | 0.226 | 1.444 | 1.34 | 0.214 | 2.5E-18 |
| 2 | NPC1 | 0.41 | 0.284 | 2.184 | 1.849 | 0.24 | 4.8E-26 |
| 2 | NPC2 | 0.101 | 0.08 | 0.411 | 0.383 | 0.102 | 0.00081 |
| 2 | NPHP3 | 0.118 | 0.068 | 0.483 | 0.346 | 0.439 | 6.7E-19 |
| 2 | NR2C1 | 0.147 | 0.103 | 0.493 | 0.444 | 0.124 | 4.2E-10 |
| 2 | NR2C2 | 0.247 | 0.163 | 1.117 | 0.949 | 0.236 | 2.6E-21 |
| 2 | NR3C1 | 0.83 | 0.486 | 8.802 | 4.493 | 0.644 | 5E-147 |
| 2 | NRBF2 | 0.164 | 0.109 | 0.708 | 0.472 | 0.308 | 9.3E-15 |
| 2 | NRDC | 0.462 | 0.302 | 2.447 | 1.883 | 0.442 | 1.3E-47 |
| 2 | NRIP1 | 0.27 | 0.176 | 1.302 | 1.03 | 0.339 | 1.4E-24 |
| 2 | NSL1 | 0.147 | 0.106 | 0.649 | 0.447 | 0.221 | 1.1E-12 |
| 2 | NSMAF | 0.144 | 0.107 | 0.689 | 0.492 | 0.221 | 6.3E-11 |
| 2 | NSMCE2 | 0.264 | 0.194 | 1.294 | 1.193 | 0.117 | 2.9E-12 |
| 2 | NSMCE4A | 0.126 | 0.079 | 0.427 | 0.413 | 0.342 | 6.5E-14 |
| 2 | NSRP1 | 0.244 | 0.168 | 1.143 | 0.974 | 0.23 | 3.5E-17 |
| 2 | NSUN2 | 0.174 | 0.12 | 0.744 | 0.66 | 0.194 | 1.5E-12 |
| 2 | NT4C3A | 0.484 | 0.246 | 4.169 | 1.479 | 1.401 | 3E-283 |
| 2 | NT4DC1 | 0.178 | 0.104 | 0.768 | 0.483 | 0.398 | 2.3E-24 |
| 2 | NUB1 | 0.698 | 0.369 | 4.649 | 2.421 | 1.166 | 1E-244 |
| 2 | NUCB1 | 0.149 | 0.114 | 0.664 | 0.616 | 0.108 | 1.8E-06 |
| 2 | NUDC | 0.179 | 0.121 | 0.744 | 0.616 | 0.292 | 4.5E-14 |
| 2 | NUDT21 | 0.138 | 0.101 | 0.449 | 0.421 | 0.103 | 1.8E-07 |
| 2 | NUDT4 | 0.124 | 0.078 | 0.419 | 0.404 | 0.347 | 2.7E-14 |
| 2 | NUMA1 | 0.206 | 0.142 | 0.916 | 0.844 | 0.101 | 4.4E-10 |
| 2 | NUP107 | 0.142 | 0.106 | 0.661 | 0.49 | 0.163 | 2.1E-10 |
| 2 | NUP40 | 0.164 | 0.106 | 0.701 | 0.464 | 0.31 | 4.8E-17 |
| 2 | NUTM2A-AS1 | 0.273 | 0.193 | 1.268 | 1.12 | 0.179 | 3.9E-16 |
| 2 | OARD1 | 0.144 | 0.104 | 0.606 | 0.436 | 0.177 | 3.5E-09 |
| 2 | OAS1 | 0.614 | 0.122 | 4.694 | 0.871 | 2.43 | 0 |
| 2 | OAS2 | 0.733 | 0.214 | 6.076 | 1.411 | 2.007 | 0 |
| 2 | OAS3 | 0.367 | 0.048 | 2.076 | 0.36 | 2.427 | 0 |
| 2 | OASL | 0.747 | 0.272 | 7.411 | 2.334 | 1.684 | 0 |
| 2 | OAZ1 | 0.482 | 0.462 | 3.479 | 3.188 | 0.167 | 1.7E-14 |
| 2 | OBI1 | 0.137 | 0.071 | 0.494 | 0.362 | 0.714 | 4.4E-30 |
| 2 | OCIAD1 | 0.423 | 0.318 | 2.184 | 1.933 | 0.176 | 1.9E-16 |
| 2 | OCIAD2 | 0.29 | 0.164 | 1.363 | 0.907 | 0.488 | 1.4E-47 |
| 2 | ODF2 | 0.114 | 0.083 | 0.476 | 0.431 | 0.143 | 4.1E-07 |
| 2 | ODF2L | 0.414 | 0.244 | 3.444 | 1.447 | 1.144 | 7E-183 |
| 2 | ODF3B | 0.141 | 0.028 | 0.638 | 0.148 | 2.108 | 6E-144 |
| 2 | OGA | 0.448 | 0.439 | 3.43 | 3.162 | 0.117 | 4.3E-14 |
| 2 | OGFR | 0.232 | 0.104 | 1.087 | 0.444 | 0.972 | 2E-74 |
| 2 | OGT | 0.334 | 0.243 | 1.601 | 1.483 | 0.11 | 6.8E-16 |
| 2 | OIP4-AS1 | 0.218 | 0.16 | 0.967 | 0.861 | 0.168 | 4.5E-11 |
| 2 | OPTN | 0.494 | 0.393 | 3.802 | 2.681 | 0.404 | 6.7E-68 |
| 2 | ORAI2 | 0.109 | 0.048 | 0.47 | 0.307 | 0.611 | 4.9E-22 |
| 2 | ORC2 | 0.203 | 0.126 | 0.922 | 0.694 | 0.409 | 3.6E-24 |
| 2 | ORC3 | 0.111 | 0.078 | 0.449 | 0.404 | 0.18 | 4.3E-08 |
| 2 | ORC4 | 0.142 | 0.091 | 0.48 | 0.481 | 0.268 | 2.1E-14 |
| 2 | ORC4 | 0.123 | 0.074 | 0.417 | 0.414 | 0.318 | 1.2E-14 |
| 2 | ORMDL1 | 0.347 | 0.27 | 1.762 | 1.467 | 0.169 | 6.1E-14 |
| 2 | OS9 | 0.204 | 0.144 | 0.881 | 0.772 | 0.19 | 1.3E-12 |
| 2 | OSTC | 0.22 | 0.143 | 0.942 | 0.801 | 0.249 | 3.5E-14 |
| 2 | OTUB1 | 0.179 | 0.129 | 0.737 | 0.644 | 0.172 | 2.8E-10 |

| | | | | | | | |
|---|---|---|---|---|---|---|---|
| 2 | OTUD4 | 0.203 | 0.13 | 0.901 | 0.704 | 0.347 | 4.7E-21 |
| 2 | OTUD7B | 0.101 | 0.049 | 0.431 | 0.329 | 0.393 | 3.7E-16 |
| 2 | OTULIN | 0.434 | 0.326 | 3.289 | 2.18 | 0.493 | 8.3E-82 |
| 2 | OTULINL | 0.266 | 0.146 | 1.223 | 0.948 | 0.367 | 4.4E-38 |
| 2 | P2RY10 | 0.163 | 0.063 | 0.692 | 0.338 | 1.032 | 1.1E-68 |
| 2 | PACS1 | 0.424 | 0.304 | 2.323 | 2.017 | 0.204 | 9.7E-23 |
| 2 | PAFAH1B2 | 0.121 | 0.084 | 0.498 | 0.44 | 0.144 | 1.2E-08 |
| 2 | PAG1 | 0.18 | 0.101 | 0.783 | 0.474 | 0.444 | 3.8E-29 |
| 2 | PAIP1 | 0.149 | 0.092 | 0.621 | 0.474 | 0.387 | 6.2E-18 |
| 2 | PAK2 | 0.46 | 0.446 | 3.396 | 3.149 | 0.104 | 4.3E-10 |
| 2 | PALM2-AKAP2 | 0.272 | 0.042 | 1.412 | 0.294 | 2.363 | 4E-306 |
| 2 | PAPOLG | 0.184 | 0.117 | 0.778 | 0.639 | 0.283 | 1.2E-19 |
| 2 | PARG | 0.169 | 0.108 | 0.702 | 0.49 | 0.249 | 6.3E-17 |
| 2 | PARP1 | 0.184 | 0.137 | 0.792 | 0.719 | 0.139 | 4.4E-09 |
| 2 | PARP10 | 0.129 | 0.04 | 0.447 | 0.263 | 1.046 | 1.2E-44 |
| 2 | PARP11 | 0.333 | 0.127 | 1.737 | 0.733 | 1.244 | 4E-146 |
| 2 | PARP12 | 0.313 | 0.103 | 1.447 | 0.494 | 1.378 | 2E-182 |
| 2 | PARP14 | 0.801 | 0.32 | 7.866 | 2.449 | 1.677 | 0 |
| 2 | PARP4 | 0.41 | 0.273 | 3.012 | 1.72 | 0.808 | 9E-120 |
| 2 | PARP8 | 0.83 | 0.644 | 8.014 | 6.291 | 0.349 | 1.3E-49 |
| 2 | PARP9 | 0.677 | 0.228 | 4.982 | 1.629 | 1.613 | 0 |
| 2 | PARVG | 0.224 | 0.141 | 1.011 | 0.811 | 0.317 | 3.8E-19 |
| 2 | PBDC1 | 0.132 | 0.084 | 0.469 | 0.438 | 0.377 | 9.8E-14 |
| 2 | PBXIP1 | 0.164 | 0.111 | 0.73 | 0.609 | 0.263 | 1.4E-13 |
| 2 | PCAT1 | 0.103 | 0.032 | 0.403 | 0.194 | 1.376 | 6.1E-63 |
| 2 | PCBP1 | 0.683 | 0.407 | 4.846 | 3.761 | 0.368 | 3.4E-44 |
| 2 | PCBP1-AS1 | 0.234 | 0.161 | 1.073 | 0.901 | 0.242 | 2.3E-17 |
| 2 | PCED1B | 0.423 | 0.292 | 2.343 | 1.914 | 0.298 | 3.4E-31 |
| 2 | PCF11 | 0.23 | 0.147 | 1.062 | 0.902 | 0.236 | 2.4E-17 |
| 2 | PCGF4 | 0.61 | 0.34 | 4.219 | 2.213 | 0.931 | 6E-163 |
| 2 | PCID2 | 0.189 | 0.109 | 0.827 | 0.6 | 0.463 | 1.7E-28 |
| 2 | PCM1 | 0.447 | 0.431 | 3.403 | 3.044 | 0.203 | 1.8E-20 |
| 2 | PCMTD1 | 0.443 | 0.277 | 2.342 | 1.704 | 0.448 | 4.1E-44 |
| 2 | PCMTD2 | 0.234 | 0.143 | 1.078 | 0.842 | 0.348 | 2.5E-22 |
| 2 | PCNX4 | 0.362 | 0.21 | 1.832 | 1.271 | 0.428 | 4.7E-44 |
| 2 | PCYT1A | 0.188 | 0.11 | 0.806 | 0.6 | 0.427 | 1.2E-26 |
| 2 | PDCD10 | 0.244 | 0.174 | 1.072 | 0.964 | 0.143 | 3.7E-13 |
| 2 | PDCD4 | 0.63 | 0.447 | 4.421 | 3.44 | 0.394 | 8.6E-47 |
| 2 | PDCD6 | 0.284 | 0.204 | 1.249 | 1.114 | 0.174 | 7.1E-14 |
| 2 | PDCD6IP | 0.389 | 0.272 | 1.927 | 1.641 | 0.232 | 4.1E-23 |
| 2 | PDE3B | 0.608 | 0.383 | 4.874 | 3.346 | 0.439 | 3.4E-88 |
| 2 | PDE4B | 0.123 | 0.04 | 0.492 | 0.226 | 1.386 | 3.2E-69 |
| 2 | PDE4D | 0.634 | 0.492 | 4.378 | 3.849 | 0.186 | 8.5E-23 |
| 2 | PDE4DIP | 0.148 | 0.084 | 0.63 | 0.448 | 0.491 | 6.4E-23 |
| 2 | PDE6D | 0.166 | 0.108 | 0.688 | 0.446 | 0.308 | 1.4E-14 |
| 2 | PDE7B | 0.111 | 0.038 | 0.444 | 0.278 | 0.966 | 4E-47 |
| 2 | PDHB | 0.166 | 0.107 | 0.716 | 0.447 | 0.361 | 1.5E-16 |
| 2 | PDHX | 0.107 | 0.063 | 0.441 | 0.333 | 0.408 | 1.1E-15 |
| 2 | PDIA3 | 0.861 | 0.747 | 9.144 | 8.287 | 0.144 | 1.5E-11 |
| 2 | PDIA4 | 0.187 | 0.13 | 0.818 | 0.694 | 0.234 | 3.2E-13 |
| 2 | PDIA6 | 0.382 | 0.294 | 2.016 | 1.793 | 0.169 | 2.6E-13 |
| 2 | PDK3 | 0.216 | 0.133 | 0.947 | 0.74 | 0.372 | 8.5E-26 |
| 2 | PDSS2 | 0.176 | 0.118 | 0.773 | 0.682 | 0.181 | 1E-14 |
| 2 | PEAK1 | 0.23 | 0.163 | 1.071 | 0.994 | 0.106 | 1.7E-13 |

| | | | | | | | |
|---|---|---|---|---|---|---|---|
| 2 | PEX1 | 0.131 | 0.096 | 0.464 | 0.406 | 0.16 | 2.6E-07 |
| 2 | PEX3 | 0.11 | 0.067 | 0.448 | 0.36 | 0.314 | 2.7E-14 |
| 2 | PHACTR2 | 0.311 | 0.161 | 1.492 | 1.004 | 0.664 | 3.8E-69 |
| 2 | PHAX | 0.146 | 0.112 | 0.672 | 0.49 | 0.187 | 3E-09 |
| 2 | PHC3 | 0.33 | 0.24 | 1.461 | 1.442 | 0.114 | 1.4E-14 |
| 2 | PHF11 | 0.447 | 0.246 | 3.444 | 1.63 | 1.121 | 2E-210 |
| 2 | PHF14 | 0.198 | 0.14 | 0.898 | 0.798 | 0.17 | 1.7E-12 |
| 2 | PHF21A | 0.394 | 0.284 | 2.064 | 1.837 | 0.169 | 2.3E-19 |
| 2 | PHF3 | 0.424 | 0.397 | 3.131 | 2.724 | 0.2 | 4E-21 |
| 2 | PHF6 | 0.127 | 0.08 | 0.438 | 0.411 | 0.389 | 4.4E-14 |
| 2 | PHIP | 0.409 | 0.31 | 2.274 | 1.99 | 0.194 | 9.2E-17 |
| 2 | PHKB | 0.223 | 0.147 | 0.981 | 0.886 | 0.147 | 1.1E-13 |
| 2 | PHPT1 | 0.118 | 0.086 | 0.482 | 0.442 | 0.124 | 9.7E-07 |
| 2 | PHTF1 | 0.142 | 0.099 | 0.636 | 0.426 | 0.274 | 1.9E-14 |
| 2 | PHYKPL | 0.16 | 0.088 | 0.647 | 0.441 | 0.42 | 2E-28 |
| 2 | PI4K2B | 0.288 | 0.108 | 1.403 | 0.604 | 1.216 | 3E-134 |
| 2 | PIBF1 | 0.28 | 0.2 | 1.377 | 1.221 | 0.174 | 4.3E-16 |
| 2 | PICALM | 0.416 | 0.386 | 3.027 | 2.646 | 0.194 | 4.1E-22 |
| 2 | PIGC | 0.176 | 0.124 | 0.747 | 0.644 | 0.189 | 4.8E-11 |
| 2 | PIGF | 0.118 | 0.072 | 0.472 | 0.369 | 0.346 | 1.4E-14 |
| 2 | PIGL | 0.109 | 0.068 | 0.448 | 0.371 | 0.271 | 8.2E-13 |
| 2 | PIGN | 0.12 | 0.079 | 0.414 | 0.444 | 0.207 | 2.1E-11 |
| 2 | PIK3AP1 | 0.343 | 0.217 | 1.921 | 1.291 | 0.473 | 2.2E-46 |
| 2 | PIK3C3 | 0.184 | 0.131 | 0.794 | 0.719 | 0.143 | 1.8E-11 |
| 2 | PIK3CB | 0.14 | 0.071 | 0.484 | 0.389 | 0.484 | 8.7E-32 |
| 2 | PIK3CD | 0.239 | 0.147 | 1.026 | 0.869 | 0.24 | 2.9E-26 |
| 2 | PIK3CG | 0.311 | 0.162 | 1.499 | 0.934 | 0.682 | 2.9E-66 |
| 2 | PIK3IP1 | 0.276 | 0.197 | 1.284 | 1.116 | 0.203 | 9.6E-16 |
| 2 | PIK3R1 | 0.712 | 0.484 | 6.328 | 4.312 | 0.243 | 4E-24 |
| 2 | PIKFYVE | 0.18 | 0.129 | 0.773 | 0.703 | 0.136 | 9.4E-11 |
| 2 | PILRB | 0.12 | 0.077 | 0.401 | 0.416 | 0.27 | 7.9E-13 |
| 2 | PITPNB | 0.277 | 0.199 | 1.248 | 1.114 | 0.174 | 4.7E-14 |
| 2 | PITPNC1 | 0.73 | 0.476 | 6.27 | 4.244 | 0.247 | 4.8E-39 |
| 2 | PJA2 | 0.434 | 0.394 | 3.049 | 2.631 | 0.213 | 1.4E-24 |
| 2 | PLAAT4 | 0.882 | 0.473 | 10.709 | 4.173 | 1.04 | 0 |
| 2 | PLCG2 | 0.324 | 0.24 | 1.636 | 1.497 | 0.128 | 2.9E-14 |
| 2 | PLCL1 | 0.117 | 0.07 | 0.449 | 0.442 | 0.28 | 7.3E-16 |
| 2 | PLCXD2 | 0.139 | 0.074 | 0.601 | 0.404 | 0.473 | 7.1E-28 |
| 2 | PLEKHA2 | 0.461 | 0.402 | 3.347 | 2.808 | 0.248 | 4.7E-34 |
| 2 | PLEKHA3 | 0.119 | 0.083 | 0.486 | 0.422 | 0.204 | 1.7E-08 |
| 2 | PLEKHA4 | 0.107 | 0.047 | 0.403 | 0.268 | 0.907 | 2.7E-34 |
| 2 | PLEKHO1 | 0.148 | 0.079 | 0.688 | 0.444 | 0.628 | 9.4E-37 |
| 2 | PLP2 | 0.144 | 0.101 | 0.684 | 0.464 | 0.279 | 2E-14 |
| 2 | PLRG1 | 0.144 | 0.101 | 0.607 | 0.423 | 0.214 | 9.8E-10 |
| 2 | PLSCR1 | 0.83 | 0.27 | 11.907 | 2.41 | 2.304 | 0 |
| 2 | PMEPA1 | 0.144 | 0.043 | 0.644 | 0.247 | 1.33 | 5E-90 |
| 2 | PML | 0.13 | 0.042 | 0.468 | 0.269 | 1.08 | 4.8E-42 |
| 2 | PMS1 | 0.181 | 0.124 | 0.791 | 0.676 | 0.226 | 6.9E-13 |
| 2 | PMS2 | 0.147 | 0.102 | 0.48 | 0.434 | 0.121 | 2.4E-10 |
| 2 | PNPT1 | 0.429 | 0.114 | 2.643 | 0.686 | 1.944 | 0 |
| 2 | POLD4 | 0.147 | 0.108 | 0.609 | 0.437 | 0.181 | 1.1E-07 |
| 2 | POLK | 0.144 | 0.097 | 0.634 | 0.43 | 0.262 | 3.1E-12 |
| 2 | POLR2B | 0.269 | 0.18 | 1.224 | 1.033 | 0.244 | 1.9E-21 |
| 2 | POMP | 0.343 | 0.26 | 1.646 | 1.403 | 0.14 | 7.9E-13 |

| | | | | | | | |
|---|---|---|---|---|---|---|---|
| 2 | POU2F1 | 0.192 | 0.137 | 0.86 | 0.793 | 0.118 | 2.4E-11 |
| 2 | POU2F2 | 0.107 | 0.073 | 0.443 | 0.404 | 0.164 | 1.2E-08 |
| 2 | PPARA | 0.106 | 0.048 | 0.441 | 0.3 | 0.443 | 4.8E-20 |
| 2 | PPCS | 0.241 | 0.173 | 1.092 | 0.948 | 0.189 | 2E-13 |
| 2 | PPID | 0.202 | 0.137 | 0.899 | 0.723 | 0.314 | 8.2E-16 |
| 2 | PPM1K | 0.44 | 0.222 | 3.713 | 1.423 | 1.383 | 2E-279 |
| 2 | PPP1CC | 0.261 | 0.169 | 1.181 | 0.914 | 0.368 | 1.2E-24 |
| 2 | PPP1R12A | 0.476 | 0.448 | 3.611 | 3.269 | 0.144 | 1.5E-17 |
| 2 | PPP1R2 | 0.479 | 0.294 | 2.486 | 1.827 | 0.401 | 4.3E-62 |
| 2 | PPP1R21 | 0.19 | 0.134 | 0.817 | 0.744 | 0.134 | 1.4E-11 |
| 2 | PPP2R2A | 0.392 | 0.246 | 2.029 | 1.468 | 0.372 | 2.3E-37 |
| 2 | PPP3CB | 0.239 | 0.162 | 1.049 | 0.908 | 0.221 | 4.1E-18 |
| 2 | PPP4R2 | 0.166 | 0.109 | 0.706 | 0.474 | 0.296 | 2.3E-14 |
| 2 | PPP4R3B | 0.434 | 0.296 | 2.229 | 1.843 | 0.274 | 8.2E-32 |
| 2 | PPTC7 | 0.147 | 0.103 | 0.643 | 0.44 | 0.248 | 2.2E-14 |
| 2 | PPWD1 | 0.238 | 0.142 | 1.016 | 0.839 | 0.276 | 7.3E-23 |
| 2 | PRDX3 | 0.142 | 0.064 | 0.474 | 0.319 | 0.849 | 3.1E-43 |
| 2 | PRDX4 | 0.308 | 0.241 | 1.444 | 1.346 | 0.1 | 3.5E-09 |
| 2 | PRDX6 | 0.349 | 0.246 | 1.748 | 1.412 | 0.308 | 1.4E-26 |
| 2 | PRH1 | 0.104 | 0.062 | 0.441 | 0.343 | 0.341 | 8.5E-14 |
| 2 | PRIM2 | 0.172 | 0.116 | 0.741 | 0.677 | 0.149 | 1.5E-13 |
| 2 | PRIMPOL | 0.108 | 0.06 | 0.44 | 0.316 | 0.412 | 1.8E-19 |
| 2 | PRKAA1 | 0.182 | 0.131 | 0.787 | 0.702 | 0.164 | 1.4E-10 |
| 2 | PRKACB | 0.644 | 0.404 | 4.714 | 3.924 | 0.264 | 2.3E-31 |
| 2 | PRKAG2 | 0.191 | 0.123 | 0.863 | 0.699 | 0.303 | 1.4E-19 |
| 2 | PRKAR1A | 0.184 | 0.104 | 0.769 | 0.447 | 0.467 | 3.5E-29 |
| 2 | PRKCE | 0.173 | 0.103 | 0.782 | 0.617 | 0.342 | 8E-24 |
| 2 | PRKCQ | 0.42 | 0.282 | 2.264 | 1.816 | 0.318 | 2E-34 |
| 2 | PRKCQ-AS1 | 0.187 | 0.094 | 0.844 | 0.402 | 0.74 | 7.4E-44 |
| 2 | PRKD2 | 0.14 | 0.072 | 0.674 | 0.402 | 0.748 | 1.9E-39 |
| 2 | PRKD3 | 0.304 | 0.214 | 1.449 | 1.267 | 0.204 | 6.7E-18 |
| 2 | PRMT7 | 0.144 | 0.073 | 0.647 | 0.38 | 0.769 | 1.9E-42 |
| 2 | PRPF38B | 0.434 | 0.33 | 2.264 | 2.092 | 0.114 | 1.3E-13 |
| 2 | PRPF6 | 0.242 | 0.179 | 1.087 | 0.993 | 0.13 | 2.6E-11 |
| 2 | PRPS2 | 0.104 | 0.04 | 0.426 | 0.243 | 0.812 | 1.7E-29 |
| 2 | PRR14L | 0.271 | 0.182 | 1.241 | 1.019 | 0.284 | 9.4E-22 |
| 2 | PRR4L | 0.263 | 0.164 | 1.197 | 0.943 | 0.344 | 3.1E-29 |
| 2 | PRRC1 | 0.119 | 0.087 | 0.489 | 0.441 | 0.118 | 4E-07 |
| 2 | PSENEN | 0.174 | 0.122 | 0.726 | 0.613 | 0.244 | 4.3E-12 |
| 2 | PSMA1 | 0.291 | 0.214 | 1.411 | 1.243 | 0.183 | 1.9E-13 |
| 2 | PSMA2 | 0.233 | 0.13 | 1.044 | 0.671 | 0.639 | 1.4E-41 |
| 2 | PSMA3 | 0.247 | 0.134 | 1.186 | 0.734 | 0.692 | 9E-44 |
| 2 | PSMA4 | 0.434 | 0.249 | 2.304 | 1.443 | 0.674 | 3.5E-79 |
| 2 | PSMA4 | 0.284 | 0.181 | 1.342 | 0.967 | 0.473 | 2.3E-30 |
| 2 | PSMA6 | 0.418 | 0.274 | 2.227 | 1.498 | 0.479 | 4.5E-44 |
| 2 | PSMB10 | 0.637 | 0.43 | 4.188 | 2.913 | 0.424 | 2.5E-74 |
| 2 | PSMB2 | 0.234 | 0.161 | 1.002 | 0.837 | 0.249 | 3.4E-17 |
| 2 | PSMB3 | 0.293 | 0.207 | 1.292 | 1.102 | 0.229 | 2.4E-17 |
| 2 | PSMB8 | 0.492 | 0.308 | 3.788 | 1.882 | 1.009 | 2E-181 |
| 2 | PSMB9 | 0.896 | 0.481 | 10.496 | 4.193 | 1.029 | 0 |
| 2 | PSMC2 | 0.137 | 0.1 | 0.463 | 0.408 | 0.148 | 8.4E-08 |
| 2 | PSMC4 | 0.149 | 0.117 | 0.648 | 0.614 | 0.1 | 2.8E-08 |
| 2 | PSMC6 | 0.197 | 0.134 | 0.84 | 0.709 | 0.262 | 9.1E-14 |
| 2 | PSMD11 | 0.124 | 0.078 | 0.404 | 0.407 | 0.312 | 2.6E-14 |

| | | | | | | | |
|---|---|---|---|---|---|---|---|
| 2 | PSMD12 | 0.146 | 0.113 | 0.662 | 0.613 | 0.112 | 9.4E-09 |
| 2 | PSMD13 | 0.138 | 0.102 | 0.464 | 0.414 | 0.137 | 1.3E-07 |
| 2 | PSMD2 | 0.167 | 0.11 | 0.684 | 0.464 | 0.276 | 1.6E-14 |
| 2 | PSMD4 | 0.274 | 0.193 | 1.21 | 1.017 | 0.24 | 2.1E-17 |
| 2 | PSMD4 | 0.101 | 0.067 | 0.416 | 0.363 | 0.196 | 2.3E-09 |
| 2 | PSME1 | 0.863 | 0.696 | 8.727 | 6.872 | 0.344 | 1.8E-44 |
| 2 | PSME2 | 0.683 | 0.426 | 4.046 | 3.007 | 0.746 | 4E-129 |
| 2 | PSMG2 | 0.264 | 0.184 | 1.211 | 0.999 | 0.277 | 1.2E-18 |
| 2 | PTGER4 | 0.36 | 0.266 | 1.906 | 1.718 | 0.14 | 7E-16 |
| 2 | PTK2B | 0.349 | 0.204 | 1.736 | 1.262 | 0.46 | 6.4E-44 |
| 2 | PTPN11 | 0.282 | 0.218 | 1.348 | 1.23 | 0.132 | 6.2E-10 |
| 2 | PTPN22 | 0.419 | 0.364 | 3.282 | 2.437 | 0.372 | 3.3E-40 |
| 2 | PTPRC | 0.997 | 0.977 | 33.891 | 31.444 | 0.104 | 2.6E-18 |
| 2 | PTPRJ | 0.412 | 0.342 | 3.082 | 2.403 | 0.3 | 6.6E-39 |
| 2 | PURA | 0.177 | 0.114 | 0.767 | 0.619 | 0.309 | 4.2E-17 |
| 2 | PUS7L | 0.121 | 0.087 | 0.416 | 0.446 | 0.179 | 1.4E-07 |
| 2 | PWP1 | 0.143 | 0.099 | 0.467 | 0.409 | 0.146 | 3.4E-10 |
| 2 | PWWP2A | 0.144 | 0.094 | 0.662 | 0.417 | 0.344 | 1.3E-18 |
| 2 | PWWP3A | 0.148 | 0.096 | 0.674 | 0.408 | 0.407 | 2.3E-20 |
| 2 | PXK | 0.198 | 0.128 | 0.843 | 0.71 | 0.264 | 4.8E-19 |
| 2 | PYCARD | 0.198 | 0.13 | 0.86 | 0.666 | 0.369 | 1E-18 |
| 2 | PYHIN1 | 0.833 | 0.64 | 9.016 | 6.794 | 0.408 | 3.8E-69 |
| 2 | PYURF | 0.362 | 0.24 | 1.789 | 1.41 | 0.343 | 1.4E-26 |
| 2 | QKI | 0.369 | 0.267 | 1.843 | 1.641 | 0.166 | 1.4E-18 |
| 2 | QSER1 | 0.129 | 0.042 | 0.482 | 0.217 | 1.424 | 6.3E-74 |
| 2 | R3HDM1 | 0.306 | 0.197 | 1.404 | 1.149 | 0.374 | 1.5E-30 |
| 2 | RAB10 | 0.313 | 0.203 | 1.447 | 1.146 | 0.334 | 3.6E-29 |
| 2 | RAB11FIP1 | 0.38 | 0.263 | 1.884 | 1.493 | 0.243 | 4.5E-24 |
| 2 | RAB21 | 0.233 | 0.16 | 1.006 | 0.888 | 0.181 | 2E-16 |
| 2 | RAB27A | 0.424 | 0.241 | 2.212 | 1.446 | 0.417 | 1.5E-62 |
| 2 | RAB27B | 0.241 | 0.161 | 1.108 | 0.933 | 0.249 | 9.4E-20 |
| 2 | RAB28 | 0.141 | 0.091 | 0.622 | 0.409 | 0.288 | 4.3E-19 |
| 2 | RAB2B | 0.129 | 0.097 | 0.441 | 0.494 | 0.129 | 2.1E-06 |
| 2 | RAB30-DT | 0.112 | 0.048 | 0.468 | 0.298 | 0.642 | 2.3E-24 |
| 2 | RAB3GAP1 | 0.224 | 0.101 | 1.084 | 0.471 | 0.924 | 6.1E-71 |
| 2 | RAB3GAP2 | 0.231 | 0.169 | 1.031 | 0.947 | 0.108 | 1.2E-11 |
| 2 | RABAC1 | 0.484 | 0.329 | 2.616 | 2.024 | 0.37 | 2.6E-40 |
| 2 | RABEP1 | 0.249 | 0.18 | 1.201 | 1.042 | 0.204 | 9.3E-17 |
| 2 | RABGAP1 | 0.203 | 0.134 | 0.874 | 0.771 | 0.181 | 2.2E-16 |
| 2 | RABGAP1L | 0.827 | 0.699 | 8.434 | 7.621 | 0.146 | 9.3E-24 |
| 2 | RAD21 | 0.426 | 0.316 | 2.171 | 1.942 | 0.16 | 2.4E-17 |
| 2 | RAD41B | 0.327 | 0.206 | 1.74 | 1.348 | 0.348 | 1.2E-34 |
| 2 | RAD9A | 0.101 | 0.063 | 0.414 | 0.328 | 0.338 | 4E-12 |
| 2 | RAF1 | 0.164 | 0.117 | 0.687 | 0.633 | 0.117 | 8.9E-10 |
| 2 | RALA | 0.414 | 0.174 | 2.348 | 1.08 | 1.127 | 1E-166 |
| 2 | RALB | 0.214 | 0.104 | 0.987 | 0.46 | 0.818 | 7E-46 |
| 2 | RALBP1 | 0.211 | 0.144 | 0.921 | 0.846 | 0.122 | 4.9E-10 |
| 2 | RALGAPA2 | 0.219 | 0.146 | 0.994 | 0.901 | 0.141 | 1.1E-12 |
| 2 | RALGAPB | 0.194 | 0.137 | 0.841 | 0.764 | 0.144 | 1.5E-12 |
| 2 | RALGPS2 | 0.116 | 0.064 | 0.496 | 0.348 | 0.468 | 2.2E-20 |
| 2 | RAN | 0.312 | 0.23 | 1.426 | 1.276 | 0.16 | 4.3E-14 |
| 2 | RAP1A | 0.474 | 0.434 | 3.442 | 3.108 | 0.188 | 1.7E-21 |
| 2 | RAPGEF6 | 0.347 | 0.204 | 1.806 | 1.212 | 0.474 | 7E-49 |
| 2 | RASA1 | 0.366 | 0.191 | 1.834 | 1.162 | 0.649 | 5E-79 |

| | | | | | | | |
|---|---|---|---|---|---|---|---|
| 2 | RASAL3 | 0.229 | 0.149 | 0.987 | 0.883 | 0.16 | 8.4E-14 |
| 2 | RASGRP1 | 0.296 | 0.214 | 1.438 | 1.283 | 0.164 | 1.5E-14 |
| 2 | RBBP4 | 0.441 | 0.327 | 2.328 | 1.991 | 0.226 | 4E-23 |
| 2 | RBBP6 | 0.302 | 0.217 | 1.499 | 1.264 | 0.246 | 6.7E-17 |
| 2 | RBCK1 | 0.232 | 0.113 | 1.049 | 0.6 | 0.804 | 9.7E-61 |
| 2 | RBM17 | 0.376 | 0.288 | 1.867 | 1.684 | 0.149 | 1.4E-12 |
| 2 | RBM23 | 0.326 | 0.221 | 1.441 | 1.249 | 0.292 | 2.4E-24 |
| 2 | RBM26 | 0.317 | 0.223 | 1.477 | 1.308 | 0.27 | 1.4E-20 |
| 2 | RBM28 | 0.121 | 0.088 | 0.417 | 0.46 | 0.169 | 2.5E-07 |
| 2 | RBM4 | 0.164 | 0.114 | 0.704 | 0.624 | 0.173 | 1.5E-11 |
| 2 | RBM41 | 0.124 | 0.073 | 0.426 | 0.384 | 0.444 | 2.5E-18 |
| 2 | RBM42 | 0.131 | 0.087 | 0.441 | 0.444 | 0.311 | 2.1E-12 |
| 2 | RBM43 | 0.16 | 0.061 | 0.734 | 0.326 | 1.173 | 1.2E-71 |
| 2 | RBM48 | 0.194 | 0.142 | 0.838 | 0.762 | 0.138 | 1.4E-09 |
| 2 | RBMS1 | 0.348 | 0.223 | 1.778 | 1.339 | 0.409 | 2.1E-41 |
| 2 | RBPJ | 0.436 | 0.19 | 2.441 | 1.179 | 1.044 | 5E-149 |
| 2 | RBX1 | 0.211 | 0.139 | 0.931 | 0.743 | 0.306 | 1E-18 |
| 2 | RC3H2 | 0.23 | 0.146 | 1.028 | 0.87 | 0.24 | 8.7E-18 |
| 2 | RCC2 | 0.103 | 0.072 | 0.42 | 0.371 | 0.178 | 1.8E-07 |
| 2 | RCOR3 | 0.247 | 0.19 | 1.177 | 1.094 | 0.104 | 8E-12 |
| 2 | RCSD1 | 0.368 | 0.269 | 1.866 | 1.627 | 0.198 | 1.4E-17 |
| 2 | RDH11 | 0.124 | 0.082 | 0.497 | 0.432 | 0.203 | 3.7E-11 |
| 2 | RDH14 | 0.112 | 0.048 | 0.466 | 0.283 | 0.717 | 3.5E-24 |
| 2 | RDX | 0.111 | 0.07 | 0.448 | 0.374 | 0.294 | 8E-13 |
| 2 | REEP3 | 0.11 | 0.049 | 0.467 | 0.303 | 0.623 | 1.1E-21 |
| 2 | REEP4 | 0.377 | 0.284 | 1.849 | 1.64 | 0.173 | 8.1E-14 |
| 2 | RELCH | 0.281 | 0.19 | 1.299 | 1.087 | 0.247 | 3.3E-21 |
| 2 | RER1 | 0.17 | 0.119 | 0.687 | 0.618 | 0.141 | 2.1E-11 |
| 2 | RESF1 | 0.791 | 0.469 | 7.916 | 4.404 | 0.424 | 2.1E-96 |
| 2 | REV3L | 0.212 | 0.148 | 0.993 | 0.906 | 0.132 | 6.3E-10 |
| 2 | REXO2 | 0.104 | 0.076 | 0.429 | 0.38 | 0.176 | 2.1E-06 |
| 2 | REXO4 | 0.114 | 0.084 | 0.492 | 0.443 | 0.121 | 7.2E-06 |
| 2 | RFFL | 0.261 | 0.172 | 1.17 | 0.996 | 0.232 | 2.8E-22 |
| 2 | RFX3 | 0.334 | 0.218 | 1.671 | 1.37 | 0.286 | 2.3E-29 |
| 2 | RFX7 | 0.247 | 0.144 | 1.187 | 0.924 | 0.361 | 4.6E-33 |
| 2 | RGS1 | 0.111 | 0.068 | 0.408 | 0.347 | 0.409 | 8.4E-14 |
| 2 | RHBDD1 | 0.103 | 0.067 | 0.432 | 0.364 | 0.246 | 4.5E-11 |
| 2 | RHBDF2 | 0.106 | 0.061 | 0.434 | 0.328 | 0.404 | 3.4E-17 |
| 2 | RHOBTB3 | 0.219 | 0.166 | 1.04 | 0.931 | 0.148 | 6.3E-09 |
| 2 | RHOT1 | 0.196 | 0.129 | 0.841 | 0.698 | 0.284 | 4.8E-17 |
| 2 | RICTOR | 0.448 | 0.343 | 2.477 | 2.268 | 0.127 | 1.5E-17 |
| 2 | RIF1 | 0.373 | 0.226 | 1.901 | 1.372 | 0.471 | 7.8E-48 |
| 2 | RIPK1 | 0.288 | 0.189 | 1.346 | 1.074 | 0.334 | 6.4E-26 |
| 2 | RNASEH2B | 0.296 | 0.196 | 1.418 | 1.126 | 0.332 | 1.1E-24 |
| 2 | RNASEH2C | 0.174 | 0.121 | 0.708 | 0.618 | 0.196 | 4.2E-12 |
| 2 | RNF111 | 0.267 | 0.183 | 1.214 | 1.041 | 0.222 | 1.3E-18 |
| 2 | RNF113A | 0.123 | 0.09 | 0.496 | 0.46 | 0.109 | 7.1E-07 |
| 2 | RNF13 | 0.261 | 0.167 | 1.188 | 0.942 | 0.334 | 3.6E-26 |
| 2 | RNF134 | 0.134 | 0.071 | 0.468 | 0.367 | 0.631 | 1.2E-28 |
| 2 | RNF14 | 0.107 | 0.07 | 0.43 | 0.349 | 0.262 | 3E-10 |
| 2 | RNF146 | 0.144 | 0.09 | 0.646 | 0.473 | 0.472 | 1.9E-23 |
| 2 | RNF149 | 0.447 | 0.369 | 3.467 | 2.396 | 0.433 | 3.2E-64 |
| 2 | RNF169 | 0.122 | 0.087 | 0.417 | 0.474 | 0.123 | 3.7E-08 |
| 2 | RNF181 | 0.188 | 0.137 | 0.797 | 0.707 | 0.173 | 3.5E-10 |

| | | | | | | | |
|---|---|---|---|---|---|---|---|
| 2 | RNF19A | 0.494 | 0.444 | 4.047 | 3.246 | 0.318 | 1E-32 |
| 2 | RNF213 | 0.976 | 0.827 | 22.477 | 11.441 | 0.968 | 0 |
| 2 | RNF213-AS1 | 0.169 | 0.094 | 0.719 | 0.498 | 0.431 | 1.3E-28 |
| 2 | RNF24 | 0.106 | 0.069 | 0.442 | 0.368 | 0.297 | 4.2E-11 |
| 2 | RNF34 | 0.194 | 0.129 | 0.809 | 0.709 | 0.19 | 4.3E-16 |
| 2 | RNFT1 | 0.166 | 0.107 | 0.692 | 0.463 | 0.297 | 1.9E-16 |
| 2 | RO60 | 0.414 | 0.311 | 2.164 | 1.944 | 0.144 | 4.2E-16 |
| 2 | ROCK2 | 0.184 | 0.114 | 0.784 | 0.643 | 0.286 | 3.1E-20 |
| 2 | RORA | 0.698 | 0.449 | 4.441 | 4.807 | 0.181 | 2E-20 |
| 2 | RORA-AS1 | 0.232 | 0.173 | 1.047 | 0.946 | 0.144 | 1.4E-10 |
| 2 | RPA2 | 0.163 | 0.12 | 0.701 | 0.623 | 0.17 | 2.3E-08 |
| 2 | RPA3 | 0.117 | 0.078 | 0.476 | 0.389 | 0.291 | 5E-11 |
| 2 | RPAP2 | 0.424 | 0.312 | 2.294 | 2.008 | 0.193 | 1.8E-19 |
| 2 | RPAP3 | 0.141 | 0.084 | 0.632 | 0.443 | 0.483 | 8.4E-20 |
| 2 | RPL36AL | 0.662 | 0.434 | 4.407 | 4.041 | 0.144 | 3.2E-14 |
| 2 | RPL7L1 | 0.179 | 0.119 | 0.774 | 0.62 | 0.319 | 4.9E-16 |
| 2 | RPLP1 | 0.992 | 0.944 | 39.772 | 36.704 | 0.116 | 1.8E-17 |
| 2 | RPP38 | 0.129 | 0.078 | 0.43 | 0.419 | 0.342 | 2E-16 |
| 2 | RPRD2 | 0.249 | 0.179 | 1.143 | 1.024 | 0.148 | 2.5E-13 |
| 2 | RPS6KA3 | 0.399 | 0.277 | 2.086 | 1.739 | 0.263 | 1.2E-26 |
| 2 | RPS6KA4 | 0.243 | 0.164 | 1.16 | 0.964 | 0.267 | 1.1E-19 |
| 2 | RPS6KB1 | 0.147 | 0.106 | 0.662 | 0.479 | 0.194 | 1.3E-12 |
| 2 | RPS6KC1 | 0.331 | 0.188 | 1.64 | 1.119 | 0.442 | 1.2E-44 |
| 2 | RSAD2 | 0.644 | 0.13 | 4.683 | 1.044 | 2.444 | 0 |
| 2 | RSBN1L | 0.401 | 0.247 | 2.062 | 1.494 | 0.464 | 6.9E-49 |
| 2 | RSF1 | 0.449 | 0.34 | 2.421 | 2.236 | 0.173 | 1.4E-19 |
| 2 | RTN3 | 0.216 | 0.163 | 0.944 | 0.884 | 0.111 | 8.1E-09 |
| 2 | RTP4 | 0.224 | 0.039 | 1.007 | 0.209 | 2.268 | 2E-269 |
| 2 | RTRAF | 0.221 | 0.16 | 0.937 | 0.84 | 0.147 | 8.4E-12 |
| 2 | RUBCN | 0.114 | 0.066 | 0.498 | 0.343 | 0.499 | 4E-19 |
| 2 | RUFY3 | 0.123 | 0.066 | 0.413 | 0.346 | 0.427 | 5E-24 |
| 2 | RWDD3 | 0.142 | 0.096 | 0.497 | 0.496 | 0.268 | 1.1E-11 |
| 2 | S100A11 | 0.381 | 0.26 | 2.024 | 1.646 | 0.299 | 7.8E-28 |
| 2 | S100PBP | 0.317 | 0.188 | 1.414 | 1.101 | 0.46 | 1.4E-42 |
| 2 | S1PR1 | 0.144 | 0.101 | 0.627 | 0.448 | 0.194 | 8.8E-10 |
| 2 | SACM1L | 0.343 | 0.224 | 1.664 | 1.321 | 0.334 | 4.9E-31 |
| 2 | SAFB | 0.324 | 0.24 | 1.61 | 1.396 | 0.206 | 7.1E-14 |
| 2 | SAMD3 | 0.664 | 0.42 | 4.814 | 4.144 | 0.213 | 4.6E-24 |
| 2 | SAMD9 | 0.841 | 0.38 | 10.27 | 3.143 | 1.703 | 0 |
| 2 | SAMD9L | 0.841 | 0.324 | 9.912 | 2.811 | 1.818 | 0 |
| 2 | SAMHD1 | 0.47 | 0.366 | 2.969 | 2.488 | 0.244 | 8.2E-18 |
| 2 | SAMSN1 | 0.403 | 0.237 | 3.244 | 1.646 | 0.979 | 1E-163 |
| 2 | SART3 | 0.101 | 0.064 | 0.402 | 0.331 | 0.28 | 1.8E-11 |
| 2 | SASH3 | 0.223 | 0.148 | 0.991 | 0.874 | 0.179 | 1.1E-13 |
| 2 | SAT1 | 0.294 | 0.144 | 1.647 | 0.879 | 0.904 | 1.8E-69 |
| 2 | SATB1-AS1 | 0.272 | 0.113 | 1.396 | 0.688 | 1.022 | 2E-101 |
| 2 | SBF2 | 0.21 | 0.14 | 1.002 | 0.879 | 0.189 | 3.1E-13 |
| 2 | SCAF11 | 0.741 | 0.464 | 4.744 | 4.49 | 0.324 | 1.3E-46 |
| 2 | SCAMP1 | 0.183 | 0.118 | 0.794 | 0.634 | 0.326 | 4.8E-18 |
| 2 | SCAMP2 | 0.148 | 0.104 | 0.608 | 0.434 | 0.184 | 1.4E-09 |
| 2 | SCARB2 | 0.134 | 0.049 | 0.489 | 0.324 | 0.863 | 4.2E-44 |
| 2 | SCLT1 | 0.484 | 0.347 | 3.04 | 2.373 | 0.348 | 1.4E-33 |
| 2 | SCML4 | 0.336 | 0.2 | 1.779 | 1.232 | 0.43 | 1.6E-49 |
| 2 | SCOC | 0.117 | 0.072 | 0.474 | 0.364 | 0.376 | 8.1E-14 |

| | | | | | | | |
|---|---|---|---|---|---|---|---|
| 2 | SCYL2 | 0.241 | 0.172 | 1.142 | 0.967 | 0.24 | 8.8E-18 |
| 2 | SDCCAG8 | 0.132 | 0.077 | 0.464 | 0.416 | 0.44 | 4.2E-20 |
| 2 | SDHA | 0.206 | 0.143 | 0.911 | 0.788 | 0.209 | 9.1E-14 |
| 2 | SDHAF2 | 0.246 | 0.172 | 1.048 | 0.918 | 0.19 | 2.1E-14 |
| 2 | SDHB | 0.198 | 0.136 | 0.834 | 0.724 | 0.204 | 4.8E-14 |
| 2 | SDHC | 0.19 | 0.134 | 0.8 | 0.698 | 0.196 | 1.3E-11 |
| 2 | SDHD | 0.243 | 0.174 | 1.049 | 0.918 | 0.193 | 1.4E-13 |
| 2 | SEC11A | 0.312 | 0.234 | 1.446 | 1.333 | 0.127 | 1.5E-12 |
| 2 | SEC22B | 0.174 | 0.122 | 0.731 | 0.67 | 0.126 | 1.6E-11 |
| 2 | SEC23IP | 0.129 | 0.072 | 0.434 | 0.383 | 0.48 | 1.9E-22 |
| 2 | SEC24D | 0.129 | 0.087 | 0.439 | 0.472 | 0.191 | 4.4E-11 |
| 2 | SEC62 | 0.473 | 0.349 | 2.448 | 2.322 | 0.14 | 9.2E-16 |
| 2 | SECISBP2L | 0.374 | 0.271 | 1.847 | 1.66 | 0.162 | 3.5E-18 |
| 2 | SEL1L | 0.347 | 0.219 | 1.714 | 1.241 | 0.466 | 2.4E-44 |
| 2 | SELENOH | 0.294 | 0.228 | 1.383 | 1.236 | 0.162 | 1.4E-09 |
| 2 | SEM1 | 0.341 | 0.232 | 1.611 | 1.293 | 0.318 | 1.9E-26 |
| 2 | SENP1 | 0.131 | 0.091 | 0.446 | 0.492 | 0.177 | 1.5E-09 |
| 2 | SENP6 | 0.486 | 0.349 | 2.742 | 2.303 | 0.247 | 2.4E-27 |
| 2 | SENP7 | 0.38 | 0.273 | 1.923 | 1.687 | 0.189 | 4.2E-20 |
| 2 | SEPHS2 | 0.141 | 0.094 | 0.624 | 0.48 | 0.377 | 1.6E-17 |
| 2 | SEPTIN2 | 0.417 | 0.306 | 2.174 | 1.879 | 0.211 | 2E-19 |
| 2 | SEPTIN6 | 0.407 | 0.378 | 3.014 | 2.414 | 0.261 | 1.8E-24 |
| 2 | SEPTIN7 | 0.92 | 0.81 | 11.673 | 10.029 | 0.219 | 9.7E-36 |
| 2 | SERBP1 | 0.44 | 0.421 | 3.174 | 2.906 | 0.127 | 1.2E-14 |
| 2 | SERINC1 | 0.449 | 0.338 | 2.336 | 2.064 | 0.178 | 6.5E-17 |
| 2 | SERINC3 | 0.284 | 0.2 | 1.286 | 1.136 | 0.179 | 4.6E-17 |
| 2 | SESN1 | 0.296 | 0.188 | 1.423 | 1.101 | 0.37 | 2.2E-31 |
| 2 | SESN3 | 0.143 | 0.094 | 0.642 | 0.44 | 0.273 | 6.9E-18 |
| 2 | SETD2 | 0.619 | 0.448 | 4.071 | 3.346 | 0.279 | 7.1E-34 |
| 2 | SETD4 | 0.314 | 0.203 | 1.413 | 1.17 | 0.371 | 4.2E-31 |
| 2 | SETDB2 | 0.201 | 0.111 | 0.9 | 0.498 | 0.491 | 1E-36 |
| 2 | SETX | 0.648 | 0.424 | 4.784 | 3.043 | 0.643 | 4E-101 |
| 2 | SF3A1 | 0.13 | 0.092 | 0.436 | 0.474 | 0.176 | 8.1E-09 |
| 2 | SF3A3 | 0.141 | 0.11 | 0.62 | 0.464 | 0.134 | 6.4E-06 |
| 2 | SF3B4 | 0.244 | 0.16 | 1.064 | 0.83 | 0.349 | 3.3E-22 |
| 2 | SFI1 | 0.104 | 0.068 | 0.433 | 0.377 | 0.2 | 9E-11 |
| 2 | SGK3 | 0.224 | 0.149 | 1.069 | 0.914 | 0.224 | 4.8E-14 |
| 2 | SGMS1 | 0.308 | 0.196 | 1.471 | 1.19 | 0.306 | 1.7E-31 |
| 2 | SH2D1B | 0.604 | 0.379 | 4.122 | 2.69 | 0.616 | 8.3E-94 |
| 2 | SH2D2A | 0.627 | 0.379 | 4.449 | 2.724 | 0.743 | 1E-121 |
| 2 | SH3BGRL | 0.494 | 0.386 | 2.779 | 2.492 | 0.147 | 1.4E-14 |
| 2 | SHFL | 0.419 | 0.176 | 2.327 | 1.034 | 1.169 | 3E-172 |
| 2 | SHISA4 | 0.244 | 0.144 | 1.177 | 0.826 | 0.412 | 1.1E-34 |
| 2 | SHPRH | 0.241 | 0.149 | 1.104 | 0.846 | 0.368 | 2.7E-27 |
| 2 | SIDT1 | 0.43 | 0.242 | 2.338 | 1.444 | 0.499 | 4.3E-77 |
| 2 | SIPA1L1 | 0.441 | 0.247 | 2.739 | 1.681 | 0.704 | 1.5E-84 |
| 2 | SKIL | 0.438 | 0.286 | 3.304 | 1.924 | 0.78 | 3E-127 |
| 2 | SLA | 0.342 | 0.241 | 1.816 | 1.444 | 0.234 | 2.1E-20 |
| 2 | SLA2 | 0.316 | 0.187 | 1.48 | 1.143 | 0.444 | 5E-44 |
| 2 | SLAMF6 | 0.164 | 0.112 | 0.729 | 0.494 | 0.294 | 1.8E-13 |
| 2 | SLAMF7 | 0.377 | 0.239 | 1.946 | 1.411 | 0.464 | 4.1E-42 |
| 2 | SLBP | 0.166 | 0.118 | 0.719 | 0.624 | 0.202 | 1.5E-10 |
| 2 | SLC12A6 | 0.147 | 0.098 | 0.61 | 0.442 | 0.171 | 1.5E-12 |
| 2 | SLC20A2 | 0.166 | 0.106 | 0.748 | 0.473 | 0.384 | 6.4E-18 |

| | | | | | | | |
|---|---|---|---|---|---|---|---|
| 2 | SLC23A2 | 0.111 | 0.068 | 0.471 | 0.372 | 0.342 | 3.2E-14 |
| 2 | SLC24A40 | 0.109 | 0.077 | 0.438 | 0.4 | 0.13 | 1.4E-07 |
| 2 | SLC24A46 | 0.136 | 0.093 | 0.447 | 0.492 | 0.181 | 8.2E-11 |
| 2 | SLC30A4 | 0.142 | 0.108 | 0.642 | 0.48 | 0.167 | 7.8E-10 |
| 2 | SLC30A6 | 0.104 | 0.064 | 0.429 | 0.348 | 0.301 | 2.5E-12 |
| 2 | SLC30A7 | 0.192 | 0.12 | 0.82 | 0.649 | 0.314 | 1.9E-21 |
| 2 | SLC30A9 | 0.174 | 0.124 | 0.734 | 0.67 | 0.134 | 3.1E-10 |
| 2 | SLC38A1 | 0.662 | 0.42 | 4.636 | 4.223 | 0.134 | 2.2E-19 |
| 2 | SLC38A9 | 0.134 | 0.083 | 0.6 | 0.484 | 0.311 | 1E-15 |
| 2 | SLC39A7 | 0.11 | 0.066 | 0.447 | 0.336 | 0.414 | 4.4E-14 |
| 2 | SLC40A1 | 0.128 | 0.086 | 0.43 | 0.433 | 0.291 | 1.3E-11 |
| 2 | SLC4A1AP | 0.124 | 0.093 | 0.442 | 0.496 | 0.126 | 1.3E-06 |
| 2 | SLC66A3 | 0.13 | 0.088 | 0.43 | 0.44 | 0.237 | 1.1E-10 |
| 2 | SLC6A6 | 0.133 | 0.067 | 0.46 | 0.374 | 0.477 | 8.9E-31 |
| 2 | SLC9A8 | 0.164 | 0.114 | 0.678 | 0.617 | 0.137 | 8.4E-12 |
| 2 | SLF2 | 0.239 | 0.149 | 1.108 | 0.867 | 0.344 | 3.1E-27 |
| 2 | SLFN12 | 0.111 | 0.044 | 0.481 | 0.233 | 1.048 | 1.3E-44 |
| 2 | SLFN12L | 0.744 | 0.446 | 6.388 | 3.714 | 0.783 | 3E-178 |
| 2 | SLFN13 | 0.288 | 0.181 | 1.393 | 1.043 | 0.403 | 1E-32 |
| 2 | SLFN4 | 0.811 | 0.446 | 8.421 | 4.109 | 1.042 | 1E-300 |
| 2 | SLIRP | 0.149 | 0.11 | 0.62 | 0.472 | 0.114 | 1.1E-07 |
| 2 | SLK | 0.221 | 0.146 | 0.992 | 0.893 | 0.143 | 1.7E-13 |
| 2 | SLMAP | 0.263 | 0.193 | 1.203 | 1.117 | 0.108 | 4.4E-12 |
| 2 | SMAD3 | 0.14 | 0.092 | 0.613 | 0.437 | 0.191 | 1.2E-17 |
| 2 | SMAD4 | 0.224 | 0.129 | 1.004 | 0.703 | 0.413 | 8.5E-34 |
| 2 | SMARCA4 | 0.443 | 0.308 | 2.376 | 1.917 | 0.31 | 1.9E-30 |
| 2 | SMARCAD1 | 0.189 | 0.13 | 0.846 | 0.707 | 0.277 | 8.4E-14 |
| 2 | SMC1A | 0.248 | 0.183 | 1.204 | 1.017 | 0.243 | 4.9E-16 |
| 2 | SMC3 | 0.286 | 0.177 | 1.36 | 1.004 | 0.438 | 1E-33 |
| 2 | SMC6 | 0.16 | 0.111 | 0.704 | 0.494 | 0.244 | 1.1E-11 |
| 2 | SMCHD1 | 0.876 | 0.627 | 10.923 | 4.926 | 0.882 | 1E-226 |
| 2 | SMG7 | 0.206 | 0.124 | 0.924 | 0.693 | 0.413 | 1.4E-27 |
| 2 | SMIM14 | 0.222 | 0.136 | 0.993 | 0.746 | 0.413 | 8E-27 |
| 2 | SMIM14 | 0.118 | 0.078 | 0.473 | 0.403 | 0.229 | 6.4E-11 |
| 2 | SMIM19 | 0.166 | 0.101 | 0.68 | 0.428 | 0.363 | 3.1E-21 |
| 2 | SMIM26 | 0.127 | 0.094 | 0.42 | 0.478 | 0.121 | 7.4E-07 |
| 2 | SMIM29 | 0.116 | 0.074 | 0.47 | 0.367 | 0.347 | 3.7E-13 |
| 2 | SMIM7 | 0.161 | 0.109 | 0.647 | 0.486 | 0.166 | 2.4E-12 |
| 2 | SMURF2 | 0.243 | 0.149 | 1.084 | 0.944 | 0.2 | 8.6E-21 |
| 2 | SNAP23 | 0.249 | 0.169 | 1.114 | 0.94 | 0.232 | 9.5E-19 |
| 2 | SNAP29 | 0.261 | 0.191 | 1.174 | 1.043 | 0.172 | 4.7E-13 |
| 2 | SNAPC4 | 0.118 | 0.074 | 0.479 | 0.374 | 0.344 | 2E-14 |
| 2 | SNAPIN | 0.104 | 0.049 | 0.446 | 0.3 | 0.474 | 4.3E-18 |
| 2 | SNRNP40 | 0.147 | 0.094 | 0.612 | 0.488 | 0.324 | 1.1E-14 |
| 2 | SNRPE | 0.13 | 0.099 | 0.438 | 0.494 | 0.12 | 4.6E-06 |
| 2 | SNRPG | 0.296 | 0.207 | 1.346 | 1.113 | 0.284 | 1.4E-19 |
| 2 | SNTB1 | 0.209 | 0.126 | 0.998 | 0.764 | 0.384 | 2.4E-27 |
| 2 | SNX1 | 0.206 | 0.144 | 0.886 | 0.802 | 0.144 | 1.2E-13 |
| 2 | SNX10 | 0.141 | 0.09 | 0.607 | 0.468 | 0.374 | 3.1E-14 |
| 2 | SNX14 | 0.304 | 0.204 | 1.424 | 1.198 | 0.24 | 4.9E-24 |
| 2 | SNX2 | 0.166 | 0.109 | 0.693 | 0.477 | 0.264 | 4.4E-14 |
| 2 | SNX20 | 0.196 | 0.094 | 0.846 | 0.408 | 0.744 | 1.5E-42 |
| 2 | SNX3 | 0.343 | 0.267 | 1.678 | 1.443 | 0.121 | 2.7E-12 |
| 2 | SNX6 | 0.401 | 0.336 | 2.841 | 2.079 | 0.446 | 1.6E-48 |

| | | | | | | | |
|---|---|---|---|---|---|---|---|
| 2 | SOCS1 | 0.132 | 0.08 | 0.638 | 0.423 | 0.494 | 2.4E-18 |
| 2 | SOS1 | 0.403 | 0.29 | 2.931 | 1.922 | 0.609 | 1.5E-88 |
| 2 | SOS2 | 0.311 | 0.23 | 1.443 | 1.399 | 0.141 | 6.4E-14 |
| 2 | SP1 | 0.104 | 0.063 | 0.444 | 0.349 | 0.348 | 1.3E-14 |
| 2 | SP100 | 0.934 | 0.638 | 13.816 | 6.494 | 1.089 | 0 |
| 2 | SP110 | 0.798 | 0.399 | 7.792 | 2.998 | 1.378 | 0 |
| 2 | SP140 | 0.481 | 0.23 | 2.873 | 1.433 | 1.004 | 7E-146 |
| 2 | SP140L | 0.34 | 0.238 | 1.686 | 1.36 | 0.311 | 3.8E-23 |
| 2 | SP2 | 0.13 | 0.081 | 0.422 | 0.401 | 0.38 | 2.1E-14 |
| 2 | SP4 | 0.181 | 0.133 | 0.784 | 0.723 | 0.118 | 4.7E-09 |
| 2 | SPAG1 | 0.108 | 0.073 | 0.438 | 0.398 | 0.137 | 4.3E-09 |
| 2 | SPART | 0.237 | 0.127 | 1.078 | 0.714 | 0.494 | 2.7E-46 |
| 2 | SPAST | 0.167 | 0.116 | 0.736 | 0.639 | 0.204 | 1.2E-11 |
| 2 | SPATA13 | 0.398 | 0.269 | 2.044 | 1.634 | 0.322 | 3E-32 |
| 2 | SPATA4 | 0.149 | 0.113 | 0.707 | 0.646 | 0.13 | 4.9E-10 |
| 2 | SPATS2L | 0.248 | 0.046 | 1.288 | 0.266 | 2.277 | 8E-281 |
| 2 | SPG11 | 0.331 | 0.196 | 1.622 | 1.146 | 0.402 | 7E-48 |
| 2 | SPIDR | 0.418 | 0.297 | 2.249 | 2.014 | 0.149 | 2.5E-23 |
| 2 | SPIN1 | 0.134 | 0.088 | 0.446 | 0.467 | 0.241 | 2.8E-13 |
| 2 | SPOP | 0.179 | 0.11 | 0.748 | 0.496 | 0.328 | 3.1E-21 |
| 2 | SPPL2A | 0.619 | 0.37 | 4.031 | 2.468 | 0.708 | 2E-118 |
| 2 | SPTLC2 | 0.37 | 0.204 | 1.922 | 1.149 | 0.729 | 1.8E-73 |
| 2 | SQOR | 0.178 | 0.08 | 0.784 | 0.419 | 0.904 | 4.3E-46 |
| 2 | SQSTM1 | 0.491 | 0.379 | 3.841 | 2.626 | 0.442 | 1.3E-80 |
| 2 | SRBD1 | 0.244 | 0.143 | 1.138 | 0.844 | 0.431 | 1E-34 |
| 2 | SRGAP2 | 0.197 | 0.079 | 0.902 | 0.434 | 1.044 | 2.7E-79 |
| 2 | SRGAP2C | 0.134 | 0.044 | 0.619 | 0.244 | 1.342 | 4.5E-73 |
| 2 | SRGN | 0.884 | 0.787 | 11.084 | 10.27 | 0.11 | 3.5E-10 |
| 2 | SRP44 | 0.248 | 0.16 | 1.202 | 0.889 | 0.434 | 7.4E-31 |
| 2 | SRP9 | 0.34 | 0.248 | 1.476 | 1.446 | 0.114 | 8.2E-12 |
| 2 | SRSF4 | 0.339 | 0.246 | 1.647 | 1.426 | 0.11 | 4E-12 |
| 2 | SRSF9 | 0.286 | 0.214 | 1.304 | 1.162 | 0.166 | 1.8E-12 |
| 2 | SS18 | 0.237 | 0.14 | 1.074 | 0.844 | 0.332 | 4.8E-24 |
| 2 | SS18L2 | 0.149 | 0.109 | 0.614 | 0.461 | 0.13 | 7.2E-08 |
| 2 | SSBP1 | 0.216 | 0.146 | 0.927 | 0.823 | 0.172 | 1E-11 |
| 2 | SSR3 | 0.267 | 0.184 | 1.194 | 1.008 | 0.244 | 2.6E-18 |
| 2 | ST3GAL4 | 0.129 | 0.09 | 0.444 | 0.467 | 0.221 | 1.9E-09 |
| 2 | STAG2 | 0.616 | 0.408 | 3.912 | 2.874 | 0.444 | 4.4E-67 |
| 2 | STAM2 | 0.212 | 0.134 | 0.964 | 0.739 | 0.384 | 2.9E-23 |
| 2 | STAMBP | 0.114 | 0.084 | 0.487 | 0.442 | 0.108 | 3.7E-06 |
| 2 | STAMBPL1 | 0.114 | 0.07 | 0.467 | 0.378 | 0.304 | 2.1E-14 |
| 2 | STARD7 | 0.206 | 0.141 | 0.906 | 0.761 | 0.241 | 1.1E-14 |
| 2 | STAT1 | 0.927 | 0.4 | 14.872 | 4.226 | 1.909 | 0 |
| 2 | STAT2 | 0.434 | 0.149 | 3.742 | 0.971 | 1.947 | 0 |
| 2 | STAT3 | 0.448 | 0.337 | 2.434 | 2.142 | 0.178 | 7.8E-18 |
| 2 | STAT4 | 0.844 | 0.707 | 8.723 | 7.401 | 0.218 | 1E-30 |
| 2 | STAT4B | 0.248 | 0.161 | 1.124 | 0.942 | 0.244 | 9.2E-23 |
| 2 | STAT6 | 0.211 | 0.144 | 0.934 | 0.846 | 0.142 | 1.4E-10 |
| 2 | STAU2 | 0.279 | 0.174 | 1.319 | 1.049 | 0.33 | 4.9E-31 |
| 2 | STIP1 | 0.197 | 0.11 | 0.882 | 0.474 | 0.621 | 9.1E-34 |
| 2 | STK17A | 0.869 | 0.683 | 9.793 | 7.008 | 0.483 | 7E-103 |
| 2 | STOM | 0.499 | 0.289 | 2.949 | 1.746 | 0.742 | 1.4E-96 |
| 2 | STRADA | 0.124 | 0.087 | 0.43 | 0.474 | 0.147 | 1.4E-08 |
| 2 | STRN | 0.228 | 0.164 | 1.036 | 0.949 | 0.126 | 9E-12 |

| | | | | | | | |
|---|---|---|---|---|---|---|---|
| 2 | STRN3 | 0.181 | 0.128 | 0.781 | 0.723 | 0.111 | 2.7E-11 |
| 2 | STX16 | 0.326 | 0.222 | 1.606 | 1.302 | 0.303 | 1.3E-24 |
| 2 | STX17 | 0.241 | 0.119 | 1.17 | 0.646 | 0.834 | 3.6E-70 |
| 2 | STX7 | 0.202 | 0.128 | 0.914 | 0.706 | 0.372 | 4.2E-22 |
| 2 | STXBP3 | 0.238 | 0.168 | 1.073 | 0.936 | 0.196 | 4.5E-14 |
| 2 | STXBP4 | 0.232 | 0.164 | 1.064 | 0.978 | 0.122 | 6.4E-13 |
| 2 | STYX | 0.168 | 0.1 | 0.707 | 0.428 | 0.421 | 9.3E-23 |
| 2 | SUCLA2 | 0.144 | 0.104 | 0.619 | 0.447 | 0.14 | 4.7E-09 |
| 2 | SUDS3 | 0.249 | 0.148 | 1.103 | 0.844 | 0.369 | 1.9E-26 |
| 2 | SUGT1 | 0.228 | 0.174 | 1.009 | 0.939 | 0.103 | 1.9E-08 |
| 2 | SUMO2 | 0.464 | 0.362 | 2.406 | 2.211 | 0.122 | 4.5E-13 |
| 2 | SUN1 | 0.111 | 0.076 | 0.446 | 0.39 | 0.194 | 6.5E-09 |
| 2 | SUPT16H | 0.266 | 0.188 | 1.18 | 1.048 | 0.171 | 7.3E-16 |
| 2 | SUPT20H | 0.269 | 0.18 | 1.216 | 0.991 | 0.296 | 7E-22 |
| 2 | SUPT3H | 0.29 | 0.146 | 1.476 | 0.86 | 0.779 | 2.6E-71 |
| 2 | SUPT6H | 0.243 | 0.174 | 1.08 | 0.964 | 0.162 | 1E-13 |
| 2 | SUZ12 | 0.328 | 0.24 | 1.482 | 1.412 | 0.163 | 4.1E-14 |
| 2 | SYNE2 | 0.914 | 0.829 | 13.674 | 12.674 | 0.11 | 7.2E-08 |
| 2 | SYNGR2 | 0.198 | 0.111 | 0.84 | 0.477 | 0.442 | 2.9E-33 |
| 2 | SYNRG | 0.43 | 0.311 | 2.287 | 1.92 | 0.243 | 7E-23 |
| 2 | TAB2 | 0.209 | 0.149 | 0.947 | 0.838 | 0.176 | 1.2E-12 |
| 2 | TAF12 | 0.137 | 0.097 | 0.472 | 0.41 | 0.164 | 4.4E-09 |
| 2 | TAF14 | 0.348 | 0.242 | 1.807 | 1.426 | 0.342 | 3.8E-28 |
| 2 | TAF1B | 0.114 | 0.079 | 0.463 | 0.404 | 0.193 | 9.2E-09 |
| 2 | TAF2 | 0.163 | 0.117 | 0.7 | 0.64 | 0.106 | 1.5E-09 |
| 2 | TALDO1 | 0.187 | 0.14 | 0.784 | 0.724 | 0.114 | 1.6E-08 |
| 2 | TANC2 | 0.148 | 0.11 | 0.707 | 0.644 | 0.111 | 2.1E-10 |
| 2 | TANK | 0.394 | 0.239 | 2.138 | 1.494 | 0.417 | 9E-44 |
| 2 | TAOK1 | 0.231 | 0.164 | 1.04 | 0.941 | 0.144 | 3.8E-13 |
| 2 | TAP1 | 0.434 | 0.239 | 3.203 | 1.443 | 1.141 | 4E-214 |
| 2 | TAP2 | 0.491 | 0.226 | 2.787 | 1.384 | 1.009 | 4E-172 |
| 2 | TAPBP | 0.687 | 0.466 | 4.904 | 3.391 | 0.433 | 6.1E-86 |
| 2 | TAPBPL | 0.141 | 0.092 | 0.467 | 0.468 | 0.276 | 1.4E-13 |
| 2 | TARS | 0.211 | 0.124 | 0.984 | 0.686 | 0.422 | 1.5E-30 |
| 2 | TASOR | 0.341 | 0.242 | 1.773 | 1.423 | 0.219 | 3.3E-19 |
| 2 | TASOR2 | 0.287 | 0.192 | 1.371 | 1.101 | 0.317 | 1.8E-24 |
| 2 | TATDN1 | 0.114 | 0.079 | 0.487 | 0.404 | 0.27 | 3.4E-09 |
| 2 | TBC1D1 | 0.318 | 0.164 | 1.609 | 0.94 | 0.761 | 2.7E-72 |
| 2 | TBC1D22A | 0.464 | 0.338 | 2.444 | 2.234 | 0.187 | 1.1E-22 |
| 2 | TBC1D2B | 0.16 | 0.09 | 0.689 | 0.4 | 0.461 | 9E-27 |
| 2 | TBC1D32 | 0.106 | 0.06 | 0.434 | 0.338 | 0.361 | 1.4E-17 |
| 2 | TBCC | 0.129 | 0.096 | 0.417 | 0.48 | 0.107 | 1.9E-06 |
| 2 | TBCD | 0.176 | 0.112 | 0.744 | 0.628 | 0.266 | 2E-18 |
| 2 | TBCK | 0.23 | 0.148 | 1.063 | 0.902 | 0.238 | 4.3E-17 |
| 2 | TBK1 | 0.316 | 0.199 | 1.434 | 1.144 | 0.424 | 4.6E-34 |
| 2 | TBL1X | 0.261 | 0.168 | 1.169 | 1.044 | 0.161 | 1.2E-24 |
| 2 | TBL1XR1 | 0.439 | 0.269 | 2.418 | 1.772 | 0.449 | 3.4E-44 |
| 2 | TBPL1 | 0.168 | 0.123 | 0.723 | 0.646 | 0.161 | 4.7E-09 |
| 2 | TCAF2 | 0.224 | 0.147 | 0.969 | 0.834 | 0.214 | 2.6E-19 |
| 2 | TCAIM | 0.103 | 0.047 | 0.449 | 0.303 | 0.466 | 1.6E-19 |
| 2 | TCEANC2 | 0.108 | 0.061 | 0.443 | 0.319 | 0.409 | 2.1E-18 |
| 2 | TCF12 | 0.411 | 0.39 | 2.996 | 2.771 | 0.112 | 1.5E-16 |
| 2 | TDP2 | 0.168 | 0.101 | 0.714 | 0.44 | 0.403 | 1E-21 |
| 2 | TDRD3 | 0.174 | 0.117 | 0.772 | 0.646 | 0.234 | 8.4E-14 |

| | | | | | | | |
|---|---|---|---|---|---|---|---|
| 2 | TDRD7 | 0.16 | 0.062 | 0.712 | 0.344 | 1.002 | 1.7E-67 |
| 2 | TENT2 | 0.424 | 0.32 | 2.372 | 2.034 | 0.221 | 7.6E-18 |
| 2 | TENT4A | 0.249 | 0.132 | 1.243 | 0.749 | 0.713 | 1.4E-42 |
| 2 | TEP1 | 0.109 | 0.069 | 0.439 | 0.374 | 0.231 | 4.3E-12 |
| 2 | TERF2 | 0.144 | 0.104 | 0.603 | 0.442 | 0.129 | 8.9E-09 |
| 2 | TERF2IP | 0.378 | 0.274 | 1.926 | 1.616 | 0.243 | 4.8E-21 |
| 2 | TESPA1 | 0.13 | 0.066 | 0.468 | 0.344 | 0.718 | 7.7E-30 |
| 2 | TFB1M | 0.101 | 0.049 | 0.427 | 0.313 | 0.447 | 4.1E-16 |
| 2 | TFCP2 | 0.144 | 0.082 | 0.669 | 0.444 | 0.489 | 1.2E-31 |
| 2 | TFG | 0.202 | 0.123 | 0.862 | 0.642 | 0.426 | 4.4E-26 |
| 2 | TFIP11 | 0.107 | 0.077 | 0.426 | 0.379 | 0.168 | 3.3E-07 |
| 2 | TGFBR2 | 0.449 | 0.284 | 2.478 | 1.787 | 0.471 | 3.1E-46 |
| 2 | TGIF1 | 0.123 | 0.084 | 0.421 | 0.439 | 0.249 | 1.5E-09 |
| 2 | TGS1 | 0.181 | 0.128 | 0.812 | 0.692 | 0.231 | 9.2E-12 |
| 2 | THAP1 | 0.114 | 0.082 | 0.497 | 0.43 | 0.209 | 4.3E-08 |
| 2 | THAP6 | 0.141 | 0.104 | 0.641 | 0.461 | 0.191 | 8.4E-11 |
| 2 | THEMIS2 | 0.16 | 0.121 | 0.728 | 0.642 | 0.149 | 2.7E-07 |
| 2 | THRAP3 | 0.4 | 0.366 | 2.836 | 2.369 | 0.26 | 4.9E-26 |
| 2 | TIAM2 | 0.101 | 0.047 | 0.409 | 0.244 | 0.684 | 3.2E-29 |
| 2 | TIGIT | 0.478 | 0.266 | 3.44 | 1.9 | 0.902 | 4E-108 |
| 2 | TIMM17B | 0.138 | 0.101 | 0.468 | 0.499 | 0.186 | 1.2E-07 |
| 2 | TIMM8B | 0.11 | 0.076 | 0.44 | 0.374 | 0.267 | 7.1E-09 |
| 2 | TINF2 | 0.167 | 0.103 | 0.708 | 0.444 | 0.382 | 4.3E-20 |
| 2 | TIPRL | 0.237 | 0.141 | 1 | 0.804 | 0.314 | 4E-24 |
| 2 | TLR1 | 0.102 | 0.044 | 0.428 | 0.29 | 0.463 | 7.2E-21 |
| 2 | TM2D1 | 0.141 | 0.101 | 0.649 | 0.444 | 0.242 | 4.5E-13 |
| 2 | TM7SF3 | 0.137 | 0.084 | 0.491 | 0.447 | 0.402 | 7.6E-17 |
| 2 | TM9SF2 | 0.418 | 0.244 | 2.119 | 1.492 | 0.406 | 1.3E-44 |
| 2 | TMBIM4 | 0.38 | 0.239 | 1.821 | 1.383 | 0.397 | 1.9E-42 |
| 2 | TMBIM6 | 0.899 | 0.769 | 9.444 | 8.46 | 0.148 | 4.1E-20 |
| 2 | TMCC1 | 0.143 | 0.107 | 0.669 | 0.499 | 0.149 | 1.5E-10 |
| 2 | TMCO1 | 0.289 | 0.223 | 1.349 | 1.243 | 0.129 | 8.7E-10 |
| 2 | TMED4 | 0.334 | 0.243 | 1.63 | 1.44 | 0.169 | 3.6E-16 |
| 2 | TMEM106B | 0.261 | 0.146 | 1.184 | 0.843 | 0.492 | 6.2E-36 |
| 2 | TMEM123 | 0.648 | 0.329 | 4.49 | 2.384 | 1.229 | 1E-249 |
| 2 | TMEM126B | 0.161 | 0.099 | 0.684 | 0.407 | 0.434 | 9.3E-20 |
| 2 | TMEM131L | 0.403 | 0.28 | 2.064 | 1.727 | 0.247 | 2.4E-27 |
| 2 | TMEM134 | 0.113 | 0.087 | 0.478 | 0.444 | 0.106 | 6.7E-05 |
| 2 | TMEM140 | 0.274 | 0.107 | 1.338 | 0.619 | 1.113 | 3E-119 |
| 2 | TMEM141 | 0.12 | 0.078 | 0.479 | 0.389 | 0.3 | 4.2E-12 |
| 2 | TMEM146 | 0.122 | 0.077 | 0.426 | 0.422 | 0.318 | 1.7E-13 |
| 2 | TMEM14B | 0.224 | 0.149 | 0.991 | 0.897 | 0.144 | 2.9E-13 |
| 2 | TMEM164 | 0.23 | 0.147 | 1.014 | 0.866 | 0.228 | 4.2E-17 |
| 2 | TMEM167A | 0.143 | 0.112 | 0.648 | 0.494 | 0.124 | 4.4E-08 |
| 2 | TMEM168 | 0.144 | 0.08 | 0.648 | 0.428 | 0.619 | 2.3E-33 |
| 2 | TMEM170A | 0.174 | 0.13 | 0.742 | 0.688 | 0.128 | 1.7E-08 |
| 2 | TMEM179B | 0.273 | 0.183 | 1.174 | 0.987 | 0.24 | 3.2E-21 |
| 2 | TMEM244 | 0.194 | 0.123 | 0.849 | 0.693 | 0.31 | 4.4E-21 |
| 2 | TMEM248 | 0.3 | 0.226 | 1.334 | 1.214 | 0.136 | 1.4E-11 |
| 2 | TMEM260 | 0.148 | 0.074 | 0.609 | 0.419 | 0.44 | 1.6E-32 |
| 2 | TMEM263 | 0.227 | 0.146 | 1.012 | 0.84 | 0.269 | 3.3E-16 |
| 2 | TMEM33 | 0.144 | 0.091 | 0.489 | 0.483 | 0.286 | 9.4E-16 |
| 2 | TMEM34B | 0.164 | 0.098 | 0.694 | 0.407 | 0.443 | 1.3E-22 |
| 2 | TMEM40A | 0.44 | 0.422 | 3.062 | 2.789 | 0.134 | 2.4E-14 |

| | | | | | | | |
|---|---|---|---|---|---|---|---|
| 2 | TMEM41B | 0.304 | 0.194 | 1.438 | 1.134 | 0.341 | 1.5E-30 |
| 2 | TMEM43 | 0.189 | 0.114 | 0.816 | 0.62 | 0.397 | 6.7E-24 |
| 2 | TMEM49 | 0.366 | 0.272 | 1.803 | 1.488 | 0.183 | 9.3E-16 |
| 2 | TMEM62 | 0.181 | 0.093 | 0.79 | 0.412 | 0.624 | 6E-40 |
| 2 | TMIGD2 | 0.198 | 0.127 | 0.939 | 0.737 | 0.34 | 2.4E-20 |
| 2 | TMOD3 | 0.462 | 0.431 | 3.288 | 2.921 | 0.171 | 1.7E-19 |
| 2 | TMPO | 0.344 | 0.236 | 1.749 | 1.394 | 0.326 | 3.6E-26 |
| 2 | TMSB4X | 0.994 | 0.934 | 36.424 | 26.618 | 0.443 | 2E-190 |
| 2 | TMX1 | 0.283 | 0.148 | 1.299 | 0.848 | 0.616 | 1.6E-40 |
| 2 | TMX3 | 0.201 | 0.144 | 0.861 | 0.788 | 0.128 | 7.2E-11 |
| 2 | TNFAIP3 | 0.707 | 0.446 | 4.913 | 4.893 | 0.273 | 1.4E-36 |
| 2 | TNFRSF14 | 0.162 | 0.119 | 0.664 | 0.617 | 0.109 | 1.7E-08 |
| 2 | TNFRSF18 | 0.111 | 0.048 | 0.411 | 0.378 | 0.433 | 4.1E-24 |
| 2 | TNFSF10 | 0.428 | 0.128 | 4.449 | 0.948 | 2.411 | 0 |
| 2 | TNFSF13B | 0.141 | 0.034 | 0.701 | 0.189 | 1.889 | 8E-120 |
| 2 | TNIP1 | 0.198 | 0.122 | 0.871 | 0.674 | 0.371 | 4.1E-24 |
| 2 | TNK2 | 0.104 | 0.042 | 0.446 | 0.236 | 0.917 | 2E-40 |
| 2 | TNPO3 | 0.166 | 0.119 | 0.736 | 0.669 | 0.139 | 8.9E-10 |
| 2 | TOMM70 | 0.131 | 0.082 | 0.441 | 0.418 | 0.374 | 1.7E-14 |
| 2 | TOP1 | 0.372 | 0.244 | 1.924 | 1.443 | 0.309 | 1.3E-27 |
| 2 | TOP2B | 0.486 | 0.32 | 2.801 | 2.086 | 0.426 | 1.7E-48 |
| 2 | TOR1AIP1 | 0.27 | 0.176 | 1.194 | 0.973 | 0.296 | 1.1E-24 |
| 2 | TOX | 0.478 | 0.361 | 2.904 | 2.614 | 0.142 | 3.2E-18 |
| 2 | TPR | 0.484 | 0.366 | 2.8 | 2.448 | 0.194 | 1.4E-18 |
| 2 | TRAF1 | 0.113 | 0.07 | 0.474 | 0.386 | 0.301 | 3.9E-14 |
| 2 | TRAF3 | 0.167 | 0.114 | 0.73 | 0.674 | 0.114 | 3.6E-12 |
| 2 | TRAFD1 | 0.293 | 0.099 | 1.414 | 0.446 | 1.444 | 1E-164 |
| 2 | TRAK2 | 0.104 | 0.07 | 0.429 | 0.344 | 0.273 | 1.1E-09 |
| 2 | TRAM1 | 0.468 | 0.368 | 2.6 | 2.328 | 0.149 | 8.4E-14 |
| 2 | TRANK1 | 0.63 | 0.279 | 4.471 | 1.89 | 1.242 | 2E-282 |
| 2 | TRAPPC11 | 0.114 | 0.076 | 0.468 | 0.417 | 0.167 | 9E-11 |
| 2 | TRAPPC3 | 0.108 | 0.076 | 0.424 | 0.378 | 0.17 | 1.1E-07 |
| 2 | TRAPPC4 | 0.192 | 0.134 | 0.81 | 0.681 | 0.24 | 2.6E-13 |
| 2 | TRAPPC4 | 0.109 | 0.072 | 0.422 | 0.346 | 0.246 | 1.3E-10 |
| 2 | TRAPPC6B | 0.139 | 0.087 | 0.491 | 0.446 | 0.373 | 3.3E-16 |
| 2 | TRAPPC8 | 0.244 | 0.164 | 1.164 | 0.924 | 0.334 | 4.5E-24 |
| 2 | TRDMT1 | 0.116 | 0.072 | 0.488 | 0.392 | 0.316 | 6.8E-14 |
| 2 | TRG-AS1 | 0.344 | 0.238 | 1.973 | 1.704 | 0.212 | 3.7E-23 |
| 2 | TRGV7 | 0.312 | 0.194 | 1.449 | 1.131 | 0.462 | 9E-37 |
| 2 | TRGV9 | 0.138 | 0.083 | 0.604 | 0.423 | 0.413 | 7.5E-20 |
| 2 | TRIM14 | 0.343 | 0.134 | 1.767 | 0.801 | 1.141 | 2E-149 |
| 2 | TRIM21 | 0.214 | 0.061 | 0.992 | 0.323 | 1.618 | 1E-143 |
| 2 | TRIM22 | 0.87 | 0.378 | 11.067 | 3.306 | 1.743 | 0 |
| 2 | TRIM23 | 0.117 | 0.078 | 0.467 | 0.414 | 0.174 | 1.1E-10 |
| 2 | TRIM24 | 0.296 | 0.094 | 1.494 | 0.431 | 1.494 | 2E-182 |
| 2 | TRIM24 | 0.123 | 0.091 | 0.414 | 0.476 | 0.114 | 9.3E-07 |
| 2 | TRIM27 | 0.192 | 0.099 | 0.818 | 0.412 | 0.677 | 1.7E-42 |
| 2 | TRIM34 | 0.121 | 0.044 | 0.416 | 0.232 | 1.14 | 1.9E-44 |
| 2 | TRIM37 | 0.119 | 0.083 | 0.499 | 0.442 | 0.141 | 1.1E-08 |
| 2 | TRIM38 | 0.413 | 0.19 | 2.228 | 1.114 | 0.999 | 7E-138 |
| 2 | TRIM4 | 0.389 | 0.164 | 1.972 | 0.934 | 1.078 | 2E-141 |
| 2 | TRIM4 | 0.134 | 0.086 | 0.441 | 0.442 | 0.26 | 2E-14 |
| 2 | TRIM44 | 0.244 | 0.143 | 1.126 | 0.824 | 0.448 | 3.3E-34 |
| 2 | TRIM46 | 0.267 | 0.117 | 1.264 | 0.666 | 0.924 | 6.7E-90 |

| | | | | | | | |
|---|---|---|---|---|---|---|---|
| 2 | TRIP11 | 0.232 | 0.148 | 1.07 | 0.913 | 0.228 | 2.4E-17 |
| 2 | TRIP12 | 0.414 | 0.347 | 2.943 | 2.287 | 0.369 | 1.8E-44 |
| 2 | TRIP4 | 0.11 | 0.07 | 0.439 | 0.386 | 0.184 | 1.7E-11 |
| 2 | TRIR | 0.276 | 0.197 | 1.234 | 1.046 | 0.224 | 9.4E-16 |
| 2 | TRMT10B | 0.127 | 0.077 | 0.427 | 0.404 | 0.384 | 9.8E-17 |
| 2 | TRMT11 | 0.106 | 0.062 | 0.446 | 0.328 | 0.443 | 9.1E-16 |
| 2 | TRMT112 | 0.313 | 0.216 | 1.431 | 1.182 | 0.276 | 2.1E-21 |
| 2 | TRNT1 | 0.14 | 0.09 | 0.637 | 0.487 | 0.388 | 2.1E-20 |
| 2 | TRPC4AP | 0.229 | 0.14 | 1.023 | 0.872 | 0.231 | 3E-20 |
| 2 | TRRAP | 0.167 | 0.112 | 0.714 | 0.616 | 0.214 | 2.1E-14 |
| 2 | TSC22D1 | 0.17 | 0.103 | 0.729 | 0.473 | 0.347 | 1.8E-21 |
| 2 | TSC22D4 | 0.178 | 0.126 | 0.764 | 0.664 | 0.203 | 1.4E-11 |
| 2 | TSEN14 | 0.212 | 0.149 | 0.928 | 0.784 | 0.244 | 4.4E-14 |
| 2 | TSG101 | 0.223 | 0.161 | 0.98 | 0.908 | 0.11 | 1.1E-11 |
| 2 | TSNAX | 0.224 | 0.14 | 0.99 | 0.811 | 0.288 | 4.5E-19 |
| 2 | TSPAN3 | 0.106 | 0.062 | 0.433 | 0.323 | 0.423 | 1.8E-16 |
| 2 | TSPAN4 | 0.141 | 0.082 | 0.678 | 0.481 | 0.496 | 4.1E-28 |
| 2 | TSPOAP1-AS1 | 0.382 | 0.208 | 1.944 | 1.291 | 0.491 | 1.1E-73 |
| 2 | TSR2 | 0.147 | 0.112 | 0.644 | 0.471 | 0.198 | 4.1E-10 |
| 2 | TTBK2 | 0.134 | 0.088 | 0.47 | 0.4 | 0.189 | 1.6E-12 |
| 2 | TTC1 | 0.24 | 0.184 | 1.084 | 0.999 | 0.118 | 2.2E-11 |
| 2 | TTC17 | 0.441 | 0.343 | 2.436 | 2.301 | 0.141 | 4.5E-16 |
| 2 | TTC33 | 0.141 | 0.087 | 0.624 | 0.46 | 0.443 | 2.5E-23 |
| 2 | TTC37 | 0.24 | 0.179 | 1.112 | 1.017 | 0.128 | 1.4E-10 |
| 2 | TTC39B | 0.391 | 0.182 | 2.134 | 1.067 | 0.999 | 5E-126 |
| 2 | TTC39C | 0.497 | 0.287 | 2.944 | 1.89 | 0.639 | 1.6E-87 |
| 2 | TTF1 | 0.107 | 0.071 | 0.48 | 0.378 | 0.343 | 3E-10 |
| 2 | TUBB | 0.193 | 0.141 | 0.827 | 0.761 | 0.12 | 1.1E-09 |
| 2 | TULP4 | 0.327 | 0.166 | 1.467 | 0.938 | 0.74 | 4.3E-77 |
| 2 | TUT4 | 0.463 | 0.441 | 3.477 | 3.178 | 0.171 | 2E-17 |
| 2 | TUT7 | 0.489 | 0.347 | 2.842 | 2.301 | 0.304 | 2.4E-31 |
| 2 | TVP23B | 0.174 | 0.096 | 0.719 | 0.492 | 0.448 | 6.8E-31 |
| 2 | TXNDC14 | 0.107 | 0.066 | 0.43 | 0.342 | 0.332 | 3.7E-13 |
| 2 | TXNDC9 | 0.17 | 0.094 | 0.738 | 0.487 | 0.499 | 9.7E-30 |
| 2 | TYMP | 0.279 | 0.063 | 1.383 | 0.347 | 1.993 | 8E-264 |
| 2 | TYROBP | 0.907 | 0.794 | 11.207 | 10.223 | 0.133 | 1.5E-12 |
| 2 | U2SURP | 0.424 | 0.326 | 2.204 | 2.028 | 0.121 | 2.4E-13 |
| 2 | UBA3 | 0.186 | 0.122 | 0.818 | 0.649 | 0.334 | 4.2E-17 |
| 2 | UBA4 | 0.114 | 0.082 | 0.446 | 0.424 | 0.104 | 3.6E-07 |
| 2 | UBAP1 | 0.204 | 0.144 | 0.886 | 0.822 | 0.107 | 4.9E-12 |
| 2 | UBC | 0.906 | 0.792 | 10.838 | 9.824 | 0.142 | 1.2E-12 |
| 2 | UBE2B | 0.33 | 0.239 | 1.424 | 1.367 | 0.148 | 7.8E-16 |
| 2 | UBE2D1 | 0.114 | 0.082 | 0.461 | 0.422 | 0.127 | 1.2E-07 |
| 2 | UBE2E1 | 0.196 | 0.134 | 0.849 | 0.776 | 0.148 | 1E-13 |
| 2 | UBE2I | 0.284 | 0.207 | 1.264 | 1.137 | 0.144 | 3.6E-14 |
| 2 | UBE2L6 | 0.449 | 0.226 | 3.448 | 1.334 | 1.37 | 2E-271 |
| 2 | UBE2W | 0.188 | 0.132 | 0.804 | 0.709 | 0.182 | 2.9E-12 |
| 2 | UBE3A | 0.306 | 0.229 | 1.44 | 1.341 | 0.103 | 2.5E-12 |
| 2 | UBE4A | 0.219 | 0.169 | 1.002 | 0.934 | 0.102 | 4.6E-08 |
| 2 | UBE4B | 0.142 | 0.099 | 0.641 | 0.461 | 0.193 | 6E-14 |
| 2 | UBN2 | 0.113 | 0.069 | 0.461 | 0.38 | 0.278 | 1.9E-14 |
| 2 | UBR2 | 0.436 | 0.262 | 2.346 | 1.642 | 0.414 | 4.4E-62 |
| 2 | UBXN1 | 0.32 | 0.244 | 1.402 | 1.366 | 0.137 | 3.7E-11 |
| 2 | UBXN2A | 0.212 | 0.124 | 0.982 | 0.692 | 0.404 | 2.1E-30 |

| | | | | | | | |
|---|---|---|---|---|---|---|---|
| 2 | UBXN4 | 0.433 | 0.399 | 3.132 | 2.683 | 0.223 | 8.8E-24 |
| 2 | UCHL3 | 0.122 | 0.076 | 0.498 | 0.41 | 0.278 | 3.5E-14 |
| 2 | UEVLD | 0.136 | 0.086 | 0.49 | 0.443 | 0.382 | 4E-14 |
| 2 | UFL1 | 0.244 | 0.168 | 1.16 | 0.924 | 0.329 | 1.5E-22 |
| 2 | UGP2 | 0.271 | 0.192 | 1.239 | 1.062 | 0.222 | 3.1E-16 |
| 2 | UMAD1 | 0.247 | 0.173 | 1.149 | 1.032 | 0.144 | 1.1E-14 |
| 2 | UNC40 | 0.168 | 0.092 | 0.704 | 0.488 | 0.431 | 1E-29 |
| 2 | UPF2 | 0.343 | 0.242 | 1.746 | 1.468 | 0.248 | 1.8E-24 |
| 2 | UPF3B | 0.126 | 0.09 | 0.42 | 0.478 | 0.121 | 9.7E-08 |
| 2 | UPP1 | 0.297 | 0.214 | 1.448 | 1.244 | 0.217 | 7.5E-16 |
| 2 | UQCR10 | 0.329 | 0.243 | 1.412 | 1.388 | 0.123 | 4.5E-11 |
| 2 | UQCR11 | 0.484 | 0.472 | 3.449 | 3.212 | 0.102 | 3E-10 |
| 2 | URI1 | 0.176 | 0.13 | 0.762 | 0.71 | 0.1 | 7.5E-09 |
| 2 | USO1 | 0.279 | 0.141 | 1.26 | 0.846 | 0.474 | 4.8E-42 |
| 2 | USP1 | 0.192 | 0.134 | 0.834 | 0.744 | 0.164 | 3.1E-12 |
| 2 | USP14 | 0.809 | 0.448 | 7.486 | 4.493 | 0.744 | 3E-179 |
| 2 | USP18 | 0.368 | 0.063 | 1.942 | 0.366 | 2.406 | 0 |
| 2 | USP24 | 0.426 | 0.243 | 2.224 | 1.601 | 0.474 | 2.9E-61 |
| 2 | USP28 | 0.397 | 0.272 | 2.06 | 1.641 | 0.319 | 5E-29 |
| 2 | USP32 | 0.14 | 0.1 | 0.626 | 0.462 | 0.144 | 6.4E-13 |
| 2 | USP33 | 0.448 | 0.29 | 2.427 | 1.813 | 0.421 | 1.3E-46 |
| 2 | USP37 | 0.143 | 0.099 | 0.619 | 0.439 | 0.201 | 1.3E-10 |
| 2 | USP42 | 0.172 | 0.104 | 0.747 | 0.471 | 0.389 | 4.3E-22 |
| 2 | USP43 | 0.123 | 0.081 | 0.431 | 0.436 | 0.286 | 1.8E-11 |
| 2 | USP8 | 0.284 | 0.208 | 1.324 | 1.204 | 0.137 | 1.5E-13 |
| 2 | USPL1 | 0.184 | 0.114 | 0.801 | 0.624 | 0.36 | 9.4E-22 |
| 2 | UTP11 | 0.109 | 0.074 | 0.441 | 0.392 | 0.2 | 8.5E-09 |
| 2 | UTP6 | 0.128 | 0.08 | 0.427 | 0.422 | 0.319 | 3.4E-14 |
| 2 | UTRN | 0.961 | 0.849 | 16.281 | 12.048 | 0.434 | 1E-122 |
| 2 | UVRAG | 0.393 | 0.261 | 2.121 | 1.608 | 0.4 | 1.3E-37 |
| 2 | UXT | 0.261 | 0.197 | 1.147 | 1.046 | 0.132 | 3.1E-10 |
| 2 | VAMP4 | 0.213 | 0.113 | 0.944 | 0.473 | 0.723 | 4.3E-44 |
| 2 | VCPIP1 | 0.241 | 0.143 | 1.103 | 0.779 | 0.401 | 3E-34 |
| 2 | VDAC1 | 0.292 | 0.192 | 1.321 | 1.027 | 0.363 | 1E-26 |
| 2 | VEZF1 | 0.326 | 0.221 | 1.441 | 1.264 | 0.284 | 1.3E-24 |
| 2 | VEZT | 0.113 | 0.082 | 0.477 | 0.438 | 0.123 | 3.8E-07 |
| 2 | VHL | 0.204 | 0.143 | 0.897 | 0.806 | 0.146 | 2.4E-09 |
| 2 | VIRMA | 0.26 | 0.184 | 1.174 | 1.01 | 0.217 | 3.4E-14 |
| 2 | VOPP1 | 0.299 | 0.221 | 1.474 | 1.319 | 0.16 | 4.1E-14 |
| 2 | VPS13C | 0.727 | 0.478 | 4.404 | 4.773 | 0.206 | 3E-24 |
| 2 | VPS26C | 0.119 | 0.08 | 0.484 | 0.426 | 0.183 | 4.3E-10 |
| 2 | VPS28 | 0.363 | 0.262 | 1.71 | 1.494 | 0.194 | 4.2E-18 |
| 2 | VPS34 | 0.207 | 0.12 | 0.876 | 0.624 | 0.488 | 4.8E-32 |
| 2 | VPS36 | 0.243 | 0.167 | 1.074 | 0.903 | 0.249 | 1.7E-17 |
| 2 | VPS37A | 0.128 | 0.079 | 0.419 | 0.413 | 0.329 | 1.2E-14 |
| 2 | VPS39 | 0.164 | 0.101 | 0.702 | 0.442 | 0.373 | 1.8E-20 |
| 2 | VPS40 | 0.148 | 0.106 | 0.642 | 0.604 | 0.108 | 1.1E-12 |
| 2 | VPS41 | 0.198 | 0.13 | 0.844 | 0.724 | 0.22 | 3.7E-17 |
| 2 | VPS44 | 0.128 | 0.074 | 0.416 | 0.391 | 0.398 | 1.1E-19 |
| 2 | VPS44 | 0.122 | 0.074 | 0.413 | 0.424 | 0.269 | 4.3E-14 |
| 2 | VPS4B | 0.361 | 0.246 | 1.7 | 1.486 | 0.194 | 2E-20 |
| 2 | VRK1 | 0.117 | 0.083 | 0.489 | 0.442 | 0.112 | 6.5E-08 |
| 2 | VRK2 | 0.229 | 0.147 | 1.046 | 0.894 | 0.224 | 1.7E-16 |
| 2 | VTA1 | 0.168 | 0.1 | 0.697 | 0.434 | 0.38 | 2.2E-23 |

| | | | | | | | |
|---|---|---|---|---|---|---|---|
| 2 | VTI1A | 0.408 | 0.297 | 2.17 | 1.9 | 0.192 | 9.7E-21 |
| 2 | VTI1B | 0.213 | 0.14 | 0.924 | 0.829 | 0.147 | 4.4E-13 |
| 2 | WAKMAR2 | 0.22 | 0.139 | 1.077 | 0.827 | 0.381 | 4.8E-24 |
| 2 | WAPL | 0.301 | 0.211 | 1.398 | 1.219 | 0.197 | 2.4E-18 |
| 2 | WARS | 0.108 | 0.043 | 0.478 | 0.292 | 0.986 | 2.6E-28 |
| 2 | WASHC1 | 0.211 | 0.161 | 0.944 | 0.883 | 0.112 | 3.8E-08 |
| 2 | WASHC3 | 0.162 | 0.108 | 0.687 | 0.448 | 0.299 | 7E-14 |
| 2 | WASHC4 | 0.307 | 0.173 | 1.468 | 0.996 | 0.46 | 1.7E-42 |
| 2 | WDR11 | 0.264 | 0.164 | 1.234 | 0.927 | 0.413 | 1.6E-30 |
| 2 | WDR33 | 0.243 | 0.191 | 1.148 | 1.074 | 0.108 | 4.5E-10 |
| 2 | WDR36 | 0.131 | 0.089 | 0.442 | 0.472 | 0.224 | 7.9E-11 |
| 2 | WDR44 | 0.138 | 0.09 | 0.472 | 0.491 | 0.22 | 4.8E-13 |
| 2 | WDR44 | 0.129 | 0.1 | 0.429 | 0.493 | 0.101 | 1.7E-05 |
| 2 | WDR7 | 0.229 | 0.142 | 1.047 | 0.89 | 0.249 | 1.3E-19 |
| 2 | WDR83OS | 0.31 | 0.236 | 1.403 | 1.3 | 0.111 | 1.6E-10 |
| 2 | WIPF1 | 0.971 | 0.814 | 16.998 | 10.947 | 0.633 | 2E-230 |
| 2 | WRNIP1 | 0.102 | 0.066 | 0.41 | 0.334 | 0.293 | 1.8E-10 |
| 2 | XAF1 | 0.872 | 0.372 | 9.962 | 3.099 | 1.684 | 0 |
| 2 | XIAP | 0.326 | 0.188 | 1.629 | 1.069 | 0.608 | 3.5E-43 |
| 2 | XKR6 | 0.174 | 0.108 | 0.774 | 0.618 | 0.327 | 4.5E-21 |
| 2 | XPC | 0.32 | 0.212 | 1.44 | 1.241 | 0.321 | 1.7E-26 |
| 2 | XPO1 | 0.381 | 0.269 | 1.943 | 1.67 | 0.219 | 1.5E-22 |
| 2 | XRCC4 | 0.484 | 0.442 | 3.419 | 3.118 | 0.174 | 2.4E-17 |
| 2 | XRCC4 | 0.101 | 0.07 | 0.429 | 0.391 | 0.134 | 1E-07 |
| 2 | XRCC6 | 0.267 | 0.198 | 1.2 | 1.081 | 0.141 | 9.6E-12 |
| 2 | XRN1 | 0.834 | 0.414 | 9.393 | 4.306 | 1.124 | 0 |
| 2 | YARS | 0.427 | 0.241 | 2.249 | 1.4 | 0.491 | 1.4E-74 |
| 2 | YEATS2 | 0.179 | 0.08 | 0.819 | 0.436 | 0.908 | 1E-47 |
| 2 | YIPF3 | 0.107 | 0.062 | 0.434 | 0.317 | 0.447 | 2.2E-16 |
| 2 | YIPF4 | 0.172 | 0.119 | 0.733 | 0.644 | 0.161 | 4.5E-12 |
| 2 | YIPF4 | 0.134 | 0.084 | 0.437 | 0.443 | 0.244 | 2.3E-14 |
| 2 | YPEL1 | 0.189 | 0.126 | 0.839 | 0.703 | 0.246 | 2.1E-16 |
| 2 | YPEL3 | 0.446 | 0.293 | 2.46 | 1.841 | 0.418 | 1.1E-47 |
| 2 | YWHAG | 0.106 | 0.07 | 0.424 | 0.376 | 0.174 | 2.4E-10 |
| 2 | ZBED4 | 0.127 | 0.077 | 0.437 | 0.42 | 0.344 | 3.5E-16 |
| 2 | ZBP1 | 0.49 | 0.2 | 3.242 | 1.231 | 1.398 | 7E-238 |
| 2 | ZBTB20 | 0.846 | 0.486 | 9.311 | 6.324 | 0.448 | 3E-146 |
| 2 | ZBTB24 | 0.192 | 0.141 | 0.869 | 0.779 | 0.147 | 7.3E-10 |
| 2 | ZBTB33 | 0.124 | 0.083 | 0.404 | 0.427 | 0.242 | 3.3E-11 |
| 2 | ZBTB40 | 0.104 | 0.066 | 0.414 | 0.342 | 0.278 | 1.3E-11 |
| 2 | ZBTB41 | 0.101 | 0.064 | 0.417 | 0.336 | 0.31 | 3E-11 |
| 2 | ZBTB43 | 0.109 | 0.079 | 0.473 | 0.423 | 0.163 | 8.9E-07 |
| 2 | ZBTB8OS | 0.178 | 0.127 | 0.749 | 0.642 | 0.2 | 2.8E-11 |
| 2 | ZC3H13 | 0.326 | 0.224 | 1.632 | 1.344 | 0.268 | 1.4E-22 |
| 2 | ZC3H14 | 0.17 | 0.106 | 0.726 | 0.472 | 0.344 | 2.9E-19 |
| 2 | ZC3H6 | 0.171 | 0.116 | 0.782 | 0.648 | 0.27 | 1.1E-13 |
| 2 | ZC3H7A | 0.329 | 0.214 | 1.468 | 1.269 | 0.304 | 2.2E-29 |
| 2 | ZC3HAV1 | 0.487 | 0.401 | 3.868 | 2.887 | 0.422 | 6.2E-46 |
| 2 | ZCCHC2 | 0.27 | 0.123 | 1.289 | 0.713 | 0.844 | 1.1E-83 |
| 2 | ZDHHC13 | 0.106 | 0.06 | 0.448 | 0.324 | 0.46 | 6.2E-18 |
| 2 | ZDHHC14 | 0.239 | 0.067 | 1.186 | 0.41 | 1.434 | 1E-169 |
| 2 | ZDHHC17 | 0.18 | 0.124 | 0.784 | 0.687 | 0.192 | 1.3E-12 |
| 2 | ZDHHC2 | 0.123 | 0.082 | 0.41 | 0.421 | 0.276 | 2.7E-11 |
| 2 | ZDHHC4 | 0.174 | 0.129 | 0.748 | 0.678 | 0.143 | 1.8E-08 |

| | | | | | | | |
|---|---|---|---|---|---|---|---|
| 2 | ZEB1 | 0.173 | 0.104 | 0.788 | 0.64 | 0.302 | 1.9E-22 |
| 2 | ZFP14 | 0.119 | 0.067 | 0.494 | 0.349 | 0.463 | 4.5E-20 |
| 2 | ZFR | 0.414 | 0.307 | 2.12 | 1.9 | 0.148 | 7.4E-18 |
| 2 | ZFX | 0.247 | 0.177 | 1.207 | 1.014 | 0.24 | 1.9E-18 |
| 2 | ZFYVE16 | 0.221 | 0.128 | 1.03 | 0.714 | 0.428 | 8.7E-34 |
| 2 | ZKSCAN1 | 0.144 | 0.1 | 0.644 | 0.436 | 0.264 | 2.1E-14 |
| 2 | ZMAT2 | 0.142 | 0.097 | 0.483 | 0.494 | 0.234 | 3.8E-11 |
| 2 | ZMPSTE24 | 0.112 | 0.084 | 0.449 | 0.428 | 0.103 | 8.2E-06 |
| 2 | ZMYM4 | 0.229 | 0.16 | 1.068 | 0.884 | 0.272 | 2.3E-14 |
| 2 | ZMYM6 | 0.214 | 0.141 | 0.931 | 0.842 | 0.127 | 6.2E-13 |
| 2 | ZMYND11 | 0.316 | 0.231 | 1.426 | 1.347 | 0.169 | 4.3E-14 |
| 2 | ZMYND8 | 0.234 | 0.149 | 1.062 | 0.929 | 0.192 | 4.9E-17 |
| 2 | ZNF106 | 0.247 | 0.183 | 1.127 | 1.034 | 0.123 | 4E-11 |
| 2 | ZNF107 | 0.147 | 0.099 | 0.713 | 0.433 | 0.42 | 3.1E-18 |
| 2 | ZNF131 | 0.282 | 0.199 | 1.311 | 1.167 | 0.167 | 1.9E-16 |
| 2 | ZNF141 | 0.116 | 0.078 | 0.484 | 0.414 | 0.226 | 2.6E-10 |
| 2 | ZNF146 | 0.148 | 0.107 | 0.648 | 0.461 | 0.208 | 1.7E-12 |
| 2 | ZNF148 | 0.374 | 0.268 | 1.836 | 1.632 | 0.17 | 1.5E-19 |
| 2 | ZNF207 | 0.499 | 0.369 | 2.723 | 2.376 | 0.197 | 2.5E-21 |
| 2 | ZNF217 | 0.38 | 0.219 | 1.938 | 1.304 | 0.472 | 1.1E-61 |
| 2 | ZNF22 | 0.19 | 0.144 | 0.824 | 0.766 | 0.106 | 1.4E-07 |
| 2 | ZNF224 | 0.142 | 0.073 | 0.631 | 0.392 | 0.688 | 4.7E-39 |
| 2 | ZNF24 | 0.2 | 0.148 | 0.867 | 0.776 | 0.161 | 9.7E-10 |
| 2 | ZNF267 | 0.262 | 0.12 | 1.266 | 0.641 | 0.949 | 2.4E-81 |
| 2 | ZNF277 | 0.143 | 0.087 | 0.613 | 0.464 | 0.399 | 1.4E-18 |
| 2 | ZNF28 | 0.117 | 0.084 | 0.496 | 0.443 | 0.164 | 4.3E-07 |
| 2 | ZNF280D | 0.173 | 0.121 | 0.727 | 0.66 | 0.141 | 1.1E-11 |
| 2 | ZNF281 | 0.11 | 0.079 | 0.466 | 0.406 | 0.197 | 3.4E-07 |
| 2 | ZNF302 | 0.194 | 0.106 | 0.861 | 0.464 | 0.608 | 4.3E-36 |
| 2 | ZNF322 | 0.101 | 0.06 | 0.384 | 0.306 | 0.329 | 1.8E-14 |
| 2 | ZNF337 | 0.112 | 0.064 | 0.448 | 0.331 | 0.436 | 3.5E-17 |
| 2 | ZNF33A | 0.213 | 0.126 | 0.943 | 0.692 | 0.461 | 1.3E-29 |
| 2 | ZNF37A | 0.141 | 0.087 | 0.477 | 0.486 | 0.249 | 6.4E-17 |
| 2 | ZNF418B | 0.102 | 0.06 | 0.442 | 0.322 | 0.446 | 3.9E-14 |
| 2 | ZNF429 | 0.103 | 0.062 | 0.43 | 0.318 | 0.434 | 1.5E-14 |
| 2 | ZNF467 | 0.106 | 0.074 | 0.441 | 0.39 | 0.176 | 3.2E-08 |
| 2 | ZNF600 | 0.347 | 0.246 | 1.786 | 1.489 | 0.169 | 4.5E-16 |
| 2 | ZNF611 | 0.202 | 0.128 | 0.837 | 0.714 | 0.228 | 8.9E-20 |
| 2 | ZNF626 | 0.109 | 0.078 | 0.442 | 0.408 | 0.147 | 3.9E-07 |
| 2 | ZNF638 | 0.464 | 0.3 | 2.444 | 1.918 | 0.408 | 9.1E-48 |
| 2 | ZNF642 | 0.274 | 0.199 | 1.271 | 1.161 | 0.13 | 1.2E-13 |
| 2 | ZNF644 | 0.243 | 0.174 | 1.097 | 0.978 | 0.166 | 2.7E-13 |
| 2 | ZNF644 | 0.199 | 0.137 | 0.889 | 0.778 | 0.192 | 2E-14 |
| 2 | ZNF674 | 0.117 | 0.082 | 0.472 | 0.431 | 0.131 | 2.9E-08 |
| 2 | ZNF701 | 0.162 | 0.094 | 0.649 | 0.484 | 0.446 | 4.1E-24 |
| 2 | ZNF706 | 0.239 | 0.148 | 1.04 | 0.849 | 0.306 | 1.2E-20 |
| 2 | ZNF718 | 0.131 | 0.086 | 0.444 | 0.443 | 0.266 | 1.2E-12 |
| 2 | ZNF720 | 0.147 | 0.084 | 0.637 | 0.442 | 0.427 | 3.8E-23 |
| 2 | ZNF721 | 0.226 | 0.168 | 1.02 | 0.932 | 0.129 | 2.8E-10 |
| 2 | ZNF74D | 0.118 | 0.061 | 0.484 | 0.339 | 0.414 | 2.5E-24 |
| 2 | ZNF766 | 0.103 | 0.073 | 0.417 | 0.374 | 0.141 | 3.4E-07 |
| 2 | ZNF770 | 0.179 | 0.124 | 0.746 | 0.644 | 0.21 | 2E-12 |
| 2 | ZNF800 | 0.346 | 0.27 | 1.794 | 1.616 | 0.141 | 4.4E-13 |
| 2 | ZNF827 | 0.146 | 0.098 | 0.603 | 0.443 | 0.124 | 3.4E-12 |

| | | | | | | | |
|---|---|---|---|---|---|---|---|
| 2 | ZNF83 | 0.247 | 0.173 | 1.166 | 0.998 | 0.224 | 6.2E-20 |
| 2 | ZNF91 | 0.274 | 0.193 | 1.264 | 1.084 | 0.222 | 3.1E-17 |
| 2 | ZNFX1 | 0.386 | 0.141 | 2.124 | 0.889 | 1.247 | 8E-178 |
| 2 | ZNRD1 | 0.143 | 0.104 | 0.626 | 0.433 | 0.231 | 7.1E-12 |
| 2 | ZNRF2 | 0.219 | 0.148 | 0.99 | 0.849 | 0.222 | 1.8E-17 |
| 2 | ZRANB2 | 0.476 | 0.367 | 2.682 | 2.409 | 0.144 | 8.8E-16 |
| 2 | ZRSR2 | 0.192 | 0.124 | 0.864 | 0.672 | 0.363 | 9E-19 |
| 3 | 45358 | 0.338 | 0.291 | 1.81 | 1.674 | 0.111 | 3.7E-05 |
| 3 | AASDHPPT | 0.149 | 0.117 | 0.699 | 0.486 | 0.244 | 2E-05 |
| 3 | ABCB1 | 0.348 | 0.301 | 2.084 | 1.927 | 0.113 | 2E-06 |
| 3 | ABHD13 | 0.118 | 0.096 | 0.438 | 0.494 | 0.123 | 0.00126 |
| 3 | ABHD18 | 0.116 | 0.096 | 0.442 | 0.496 | 0.144 | 0.00264 |
| 3 | ABHD3 | 0.139 | 0.114 | 0.683 | 0.618 | 0.144 | 0.00041 |
| 3 | AC004441.1 | 0.122 | 0.098 | 0.487 | 0.422 | 0.169 | 0.00038 |
| 3 | AC006369.1 | 0.161 | 0.09 | 0.874 | 0.401 | 0.803 | 1.4E-27 |
| 3 | AC008874.3 | 0.334 | 0.248 | 2.024 | 1.442 | 0.384 | 4.8E-16 |
| 3 | AC010894.3 | 0.287 | 0.241 | 1.499 | 1.393 | 0.104 | 0.00027 |
| 3 | AC016831.7 | 0.113 | 0.081 | 0.427 | 0.434 | 0.28 | 3.4E-07 |
| 3 | AC022074.1 | 0.199 | 0.164 | 0.964 | 0.871 | 0.148 | 0.00012 |
| 3 | AC099063.1 | 0.132 | 0.112 | 0.634 | 0.486 | 0.113 | 0.00468 |
| 3 | AC136474.1 | 0.147 | 0.114 | 0.689 | 0.472 | 0.269 | 4.3E-06 |
| 3 | AC243829.1 | 0.138 | 0.094 | 0.693 | 0.492 | 0.494 | 1.4E-11 |
| 3 | AC243829.2 | 0.138 | 0.03 | 0.814 | 0.164 | 2.303 | 9E-128 |
| 3 | ACADM | 0.188 | 0.149 | 0.942 | 0.784 | 0.262 | 1.5E-06 |
| 3 | ACSL4 | 0.149 | 0.109 | 0.716 | 0.473 | 0.322 | 9.8E-09 |
| 3 | ADAM10 | 0.662 | 0.499 | 4.819 | 4.144 | 0.178 | 7.4E-10 |
| 3 | ADAR | 0.46 | 0.407 | 3.283 | 2.907 | 0.176 | 1.2E-06 |
| 3 | ADIPOR2 | 0.236 | 0.197 | 1.18 | 1.044 | 0.163 | 1.3E-05 |
| 3 | ADRB2 | 0.161 | 0.126 | 0.804 | 0.677 | 0.248 | 1.7E-06 |
| 3 | AFTPH | 0.21 | 0.178 | 1.062 | 0.973 | 0.127 | 0.00042 |
| 3 | AGGF1 | 0.173 | 0.127 | 0.84 | 0.663 | 0.34 | 1.3E-09 |
| 3 | AGO3 | 0.232 | 0.191 | 1.2 | 1.074 | 0.16 | 9E-06 |
| 3 | AHCTF1 | 0.184 | 0.144 | 0.904 | 0.794 | 0.186 | 8E-07 |
| 3 | AHNAK | 0.468 | 0.343 | 3.09 | 2.484 | 0.314 | 1.4E-23 |
| 3 | AK3 | 0.114 | 0.091 | 0.443 | 0.444 | 0.283 | 0.00024 |
| 3 | AK9 | 0.129 | 0.09 | 0.616 | 0.476 | 0.372 | 1.7E-09 |
| 3 | AKAP11 | 0.149 | 0.126 | 0.734 | 0.642 | 0.174 | 0.00241 |
| 3 | AKAP4 | 0.248 | 0.18 | 1.489 | 1.081 | 0.461 | 2.9E-19 |
| 3 | AL136444.1 | 0.284 | 0.238 | 1.473 | 1.3 | 0.18 | 2.4E-06 |
| 3 | AMBRA1 | 0.606 | 0.432 | 4.396 | 4.048 | 0.119 | 8.1E-08 |
| 3 | ANAPC4 | 0.134 | 0.11 | 0.637 | 0.49 | 0.111 | 0.00073 |
| 3 | ANGEL2 | 0.147 | 0.123 | 0.746 | 0.632 | 0.239 | 6.2E-06 |
| 3 | ANKFY1 | 0.167 | 0.138 | 0.814 | 0.744 | 0.133 | 0.0002 |
| 3 | ANKRD12 | 0.668 | 0.498 | 4.62 | 4.12 | 0.134 | 9.5E-10 |
| 3 | ANKRD13C | 0.142 | 0.119 | 0.703 | 0.631 | 0.146 | 0.00124 |
| 3 | ANKRD49 | 0.133 | 0.113 | 0.641 | 0.492 | 0.114 | 0.00414 |
| 3 | ANP32E | 0.382 | 0.324 | 2.183 | 1.96 | 0.146 | 1.6E-07 |
| 3 | ANXA1 | 0.784 | 0.662 | 8.629 | 7.233 | 0.244 | 3.5E-26 |
| 3 | ANXA4 | 0.171 | 0.048 | 0.902 | 0.308 | 1.449 | 3E-86 |
| 3 | AP001011.1 | 0.191 | 0.164 | 0.949 | 0.864 | 0.134 | 0.00142 |
| 3 | AP2B1 | 0.276 | 0.233 | 1.431 | 1.301 | 0.137 | 2.3E-05 |
| 3 | AP3S1 | 0.144 | 0.134 | 0.744 | 0.669 | 0.144 | 0.00481 |
| 3 | AP4M1 | 0.108 | 0.09 | 0.494 | 0.443 | 0.126 | 0.00444 |
| 3 | APAF1 | 0.141 | 0.114 | 0.663 | 0.486 | 0.178 | 0.00014 |

| | | | | | | | |
|---|---|---|---|---|---|---|---|
| 3 | APBB1IP | 0.468 | 0.484 | 3.901 | 3.471 | 0.168 | 4.5E-11 |
| 3 | API4 | 0.204 | 0.176 | 0.997 | 0.918 | 0.118 | 0.00102 |
| 3 | APOBEC3G | 0.424 | 0.398 | 3.814 | 3.062 | 0.317 | 2.2E-26 |
| 3 | ARF4 | 0.249 | 0.224 | 1.304 | 1.183 | 0.142 | 0.00032 |
| 3 | ARHGAP10 | 0.167 | 0.093 | 0.841 | 0.432 | 0.661 | 1.4E-27 |
| 3 | ARHGAP17 | 0.126 | 0.104 | 0.604 | 0.446 | 0.122 | 0.00215 |
| 3 | ARHGAP24 | 0.304 | 0.231 | 1.676 | 1.32 | 0.344 | 2.4E-14 |
| 3 | ARHGAP34 | 0.129 | 0.097 | 0.611 | 0.419 | 0.236 | 2.3E-06 |
| 3 | ARHGEF12 | 0.124 | 0.1 | 0.607 | 0.411 | 0.249 | 0.00014 |
| 3 | ARHGEF3 | 0.42 | 0.448 | 3.82 | 3.469 | 0.139 | 7.8E-08 |
| 3 | ARID4B | 0.214 | 0.148 | 1.181 | 0.874 | 0.434 | 1.2E-16 |
| 3 | ARL6IP4 | 0.634 | 0.43 | 4.774 | 4.044 | 0.239 | 3.4E-19 |
| 3 | ARL6IP6 | 0.131 | 0.102 | 0.619 | 0.42 | 0.243 | 3.1E-05 |
| 3 | ARMC1 | 0.127 | 0.103 | 0.611 | 0.424 | 0.223 | 0.00034 |
| 3 | ARNT | 0.11 | 0.092 | 0.4 | 0.446 | 0.134 | 0.00684 |
| 3 | ARPP19 | 0.279 | 0.224 | 1.446 | 1.232 | 0.241 | 1.2E-08 |
| 3 | ASXL2 | 0.194 | 0.162 | 0.968 | 0.868 | 0.147 | 8.4E-05 |
| 3 | ATAD2 | 0.126 | 0.094 | 0.619 | 0.403 | 0.301 | 1.9E-06 |
| 3 | ATF1 | 0.149 | 0.12 | 0.744 | 0.637 | 0.224 | 3.1E-07 |
| 3 | ATL3 | 0.194 | 0.142 | 0.92 | 0.789 | 0.222 | 2E-07 |
| 3 | ATP1A1 | 0.22 | 0.146 | 1.082 | 0.827 | 0.389 | 1.4E-14 |
| 3 | ATP2B4 | 0.396 | 0.306 | 2.33 | 1.89 | 0.302 | 8.3E-17 |
| 3 | ATP4MD | 0.367 | 0.314 | 1.974 | 1.776 | 0.143 | 1.4E-06 |
| 3 | ATP4PD | 0.21 | 0.184 | 1.039 | 0.962 | 0.112 | 0.00373 |
| 3 | ATP6AP2 | 0.247 | 0.227 | 1.304 | 1.196 | 0.126 | 0.00134 |
| 3 | ATP6V1A | 0.114 | 0.084 | 0.461 | 0.442 | 0.313 | 1.2E-06 |
| 3 | ATXN1 | 0.379 | 0.311 | 2.28 | 2.068 | 0.141 | 4.5E-09 |
| 3 | ATXN7L1 | 0.146 | 0.124 | 0.761 | 0.683 | 0.146 | 4.4E-05 |
| 3 | AUH | 0.238 | 0.199 | 1.214 | 1.121 | 0.114 | 3.1E-05 |
| 3 | AZI2 | 0.144 | 0.123 | 0.761 | 0.642 | 0.243 | 2.4E-05 |
| 3 | B2M | 1 | 0.998 | 223.431 | 202.14 | 0.144 | 1E-43 |
| 3 | BABAM1 | 0.148 | 0.136 | 0.737 | 0.671 | 0.136 | 0.00348 |
| 3 | BAG4 | 0.108 | 0.088 | 0.408 | 0.446 | 0.188 | 0.00117 |
| 3 | BAZ1B | 0.244 | 0.194 | 1.283 | 1.047 | 0.28 | 9.5E-09 |
| 3 | BBS9 | 0.209 | 0.17 | 1.077 | 0.997 | 0.111 | 1.4E-05 |
| 3 | BCL11B | 0.364 | 0.249 | 2.094 | 1.47 | 0.416 | 4.7E-26 |
| 3 | BCL2L11 | 0.132 | 0.083 | 0.669 | 0.433 | 0.629 | 3E-14 |
| 3 | BECN1 | 0.143 | 0.117 | 0.693 | 0.603 | 0.2 | 0.00035 |
| 3 | BICD1 | 0.127 | 0.08 | 0.684 | 0.424 | 0.69 | 1.1E-14 |
| 3 | BIRC3 | 0.244 | 0.149 | 1.464 | 0.994 | 0.449 | 4.5E-26 |
| 3 | BLOC1S1 | 0.339 | 0.268 | 1.814 | 1.484 | 0.289 | 1.3E-12 |
| 3 | BMI1 | 0.177 | 0.124 | 0.844 | 0.648 | 0.398 | 2.7E-12 |
| 3 | BMPR1A | 0.103 | 0.047 | 0.406 | 0.278 | 0.866 | 4.5E-18 |
| 3 | BMPR2 | 0.201 | 0.144 | 1.037 | 0.881 | 0.234 | 2.5E-08 |
| 3 | BRD8 | 0.168 | 0.129 | 0.772 | 0.664 | 0.217 | 7.3E-07 |
| 3 | BTBD7 | 0.112 | 0.092 | 0.44 | 0.474 | 0.184 | 0.00194 |
| 3 | BTD | 0.102 | 0.084 | 0.417 | 0.428 | 0.271 | 0.00261 |
| 3 | BTF3 | 0.499 | 0.426 | 4.196 | 3.867 | 0.118 | 2.4E-07 |
| 3 | BTG3 | 0.132 | 0.111 | 0.636 | 0.478 | 0.137 | 0.00268 |
| 3 | BTN3A1 | 0.417 | 0.329 | 2.434 | 2.062 | 0.298 | 3.5E-16 |
| 3 | BTN3A2 | 0.362 | 0.309 | 2.04 | 1.862 | 0.132 | 4.2E-06 |
| 3 | C11orf48 | 0.489 | 0.434 | 3.03 | 2.816 | 0.106 | 2.4E-05 |
| 3 | C12orf64 | 0.114 | 0.092 | 0.439 | 0.477 | 0.177 | 0.00088 |
| 3 | C12orf74 | 0.421 | 0.389 | 3.486 | 2.487 | 0.43 | 4.4E-33 |

| | | | | | | | |
|---|---|---|---|---|---|---|---|
| 3 | C14orf40 | 0.14 | 0.118 | 0.702 | 0.494 | 0.239 | 0.00128 |
| 3 | C18orf32 | 0.194 | 0.171 | 0.96 | 0.864 | 0.142 | 0.00418 |
| 3 | C1orf21 | 0.608 | 0.409 | 4.447 | 3.917 | 0.218 | 1.4E-14 |
| 3 | C21orf91 | 0.244 | 0.187 | 1.24 | 1.013 | 0.292 | 1E-10 |
| 3 | C3orf38 | 0.184 | 0.148 | 0.932 | 0.82 | 0.184 | 0.00144 |
| 3 | C4orf3 | 0.337 | 0.3 | 1.843 | 1.697 | 0.119 | 0.0004 |
| 3 | C4orf46 | 0.433 | 0.444 | 3.662 | 3.249 | 0.168 | 2.7E-10 |
| 3 | C6orf89 | 0.143 | 0.117 | 0.664 | 0.48 | 0.194 | 0.00043 |
| 3 | C7orf40 | 0.114 | 0.084 | 0.407 | 0.424 | 0.244 | 1.3E-06 |
| 3 | CAB39 | 0.271 | 0.216 | 1.414 | 1.241 | 0.19 | 1.4E-08 |
| 3 | CADM1 | 0.139 | 0.03 | 0.71 | 0.162 | 2.137 | 1E-126 |
| 3 | CALM2 | 0.41 | 0.361 | 2.343 | 2.183 | 0.102 | 7.2E-05 |
| 3 | CAMKMT | 0.217 | 0.183 | 1.214 | 1.111 | 0.13 | 9.5E-05 |
| 3 | CAMLG | 0.146 | 0.127 | 0.738 | 0.646 | 0.192 | 0.00014 |
| 3 | CAND1 | 0.249 | 0.214 | 1.306 | 1.189 | 0.134 | 0.00031 |
| 3 | CAPN2 | 0.378 | 0.292 | 2.192 | 1.739 | 0.334 | 8.6E-17 |
| 3 | CARD19 | 0.106 | 0.074 | 0.414 | 0.376 | 0.449 | 2.2E-07 |
| 3 | CARNMT1 | 0.126 | 0.099 | 0.604 | 0.499 | 0.276 | 6.3E-05 |
| 3 | CASP1 | 0.197 | 0.123 | 1.031 | 0.642 | 0.684 | 1.2E-23 |
| 3 | CAST | 0.774 | 0.694 | 7.621 | 6.887 | 0.146 | 2.2E-10 |
| 3 | CBL | 0.163 | 0.137 | 0.786 | 0.703 | 0.16 | 0.00071 |
| 3 | CBR4 | 0.196 | 0.144 | 1.044 | 0.892 | 0.242 | 9.4E-07 |
| 3 | CCDC14 | 0.12 | 0.098 | 0.48 | 0.492 | 0.239 | 0.00045 |
| 3 | CCDC146 | 0.132 | 0.107 | 0.644 | 0.468 | 0.182 | 0.00037 |
| 3 | CCDC18-AS1 | 0.179 | 0.142 | 0.916 | 0.834 | 0.134 | 0.00118 |
| 3 | CCDC24 | 0.127 | 0.098 | 0.622 | 0.494 | 0.328 | 1.1E-05 |
| 3 | CCDC66 | 0.124 | 0.098 | 0.491 | 0.412 | 0.207 | 8.7E-05 |
| 3 | CCDC88C | 0.324 | 0.248 | 1.822 | 1.463 | 0.316 | 2.3E-14 |
| 3 | CCDC91 | 0.343 | 0.3 | 2.063 | 1.849 | 0.14 | 4.3E-05 |
| 3 | CCDC92 | 0.104 | 0.083 | 0.497 | 0.414 | 0.263 | 0.00035 |
| 3 | CCL4 | 0.996 | 0.904 | 81.043 | 48.491 | 0.738 | 0 |
| 3 | CCNDBP1 | 0.167 | 0.144 | 0.812 | 0.737 | 0.14 | 0.00415 |
| 3 | CCNH | 0.468 | 0.471 | 4.462 | 3.664 | 0.284 | 9.8E-19 |
| 3 | CCSER2 | 0.714 | 0.614 | 6.649 | 4.683 | 0.226 | 4.9E-18 |
| 3 | CD2 | 0.391 | 0.166 | 2.644 | 1.069 | 1.306 | 1E-140 |
| 3 | CD37 | 0.448 | 0.39 | 2.714 | 2.444 | 0.142 | 1.7E-06 |
| 3 | CD3E | 0.386 | 0.146 | 3.234 | 1.018 | 1.668 | 4E-191 |
| 3 | CD42 | 0.481 | 0.298 | 4.928 | 2.482 | 1.199 | 2E-183 |
| 3 | CD46 | 0.446 | 0.386 | 2.714 | 2.443 | 0.146 | 1E-06 |
| 3 | CD48 | 0.464 | 0.411 | 2.849 | 2.638 | 0.116 | 2.9E-05 |
| 3 | CD48 | 0.291 | 0.23 | 1.413 | 1.347 | 0.168 | 9.8E-09 |
| 3 | CD6 | 0.147 | 0.074 | 0.734 | 0.399 | 0.88 | 9E-40 |
| 3 | CD74 | 0.744 | 0.701 | 9.177 | 8.198 | 0.163 | 0.00035 |
| 3 | CD99 | 0.741 | 0.631 | 6.649 | 4.712 | 0.219 | 7.8E-19 |
| 3 | CDC26 | 0.144 | 0.113 | 0.664 | 0.444 | 0.264 | 1.2E-05 |
| 3 | CDC42SE2 | 0.804 | 0.718 | 7.894 | 7.344 | 0.104 | 3E-10 |
| 3 | CDC73 | 0.308 | 0.261 | 1.629 | 1.41 | 0.109 | 2.7E-05 |
| 3 | CDK4RAP2 | 0.22 | 0.181 | 1.127 | 1.016 | 0.14 | 1.7E-05 |
| 3 | CDK8 | 0.127 | 0.097 | 0.497 | 0.423 | 0.191 | 9.9E-06 |
| 3 | CDKN1B | 0.444 | 0.361 | 2.724 | 2.286 | 0.243 | 6.2E-14 |
| 3 | CDKN2AIP | 0.166 | 0.144 | 0.794 | 0.741 | 0.102 | 0.00848 |
| 3 | CEACAM21 | 0.124 | 0.1 | 0.601 | 0.428 | 0.188 | 0.00024 |
| 3 | CEP104 | 0.129 | 0.106 | 0.618 | 0.441 | 0.193 | 0.00137 |
| 3 | CEP120 | 0.142 | 0.114 | 0.711 | 0.609 | 0.224 | 0.00016 |

| | | | | | | | |
|---|---|---|---|---|---|---|---|
| 3 | CEP290 | 0.142 | 0.112 | 0.698 | 0.609 | 0.196 | 3.7E-05 |
| 3 | CEP340 | 0.427 | 0.367 | 2.668 | 2.424 | 0.138 | 6.4E-07 |
| 3 | CEP47 | 0.236 | 0.192 | 1.162 | 1.044 | 0.141 | 8.7E-06 |
| 3 | CERK | 0.134 | 0.104 | 0.626 | 0.446 | 0.198 | 4.1E-05 |
| 3 | CHD7 | 0.14 | 0.109 | 0.698 | 0.611 | 0.191 | 1.9E-05 |
| 3 | CHST12 | 0.299 | 0.261 | 1.647 | 1.493 | 0.142 | 0.00015 |
| 3 | CIB1 | 0.264 | 0.22 | 1.37 | 1.168 | 0.231 | 6E-07 |
| 3 | CKAP2 | 0.173 | 0.108 | 0.838 | 0.476 | 0.441 | 2E-19 |
| 3 | CLASP1 | 0.383 | 0.324 | 2.198 | 2.038 | 0.109 | 1E-06 |
| 3 | CLEC2D | 0.473 | 0.348 | 4.613 | 2.431 | 0.866 | 2E-106 |
| 3 | CLIP1 | 0.263 | 0.216 | 1.429 | 1.229 | 0.218 | 3.8E-07 |
| 3 | CLPX | 0.143 | 0.122 | 0.691 | 0.627 | 0.14 | 4.9E-05 |
| 3 | CLSTN1 | 0.178 | 0.141 | 0.843 | 0.774 | 0.139 | 0.00077 |
| 3 | CMPK1 | 0.346 | 0.303 | 1.948 | 1.77 | 0.138 | 3.9E-06 |
| 3 | CMPK2 | 0.164 | 0.128 | 1.001 | 0.791 | 0.34 | 4.4E-07 |
| 3 | CNBP | 0.481 | 0.409 | 2.888 | 2.444 | 0.183 | 2E-09 |
| 3 | CNIH4 | 0.111 | 0.087 | 0.428 | 0.436 | 0.276 | 0.00014 |
| 3 | CNTRL | 0.242 | 0.201 | 1.272 | 1.148 | 0.148 | 1.9E-05 |
| 3 | COG6 | 0.136 | 0.109 | 0.642 | 0.476 | 0.18 | 0.00013 |
| 3 | COMMD2 | 0.137 | 0.113 | 0.646 | 0.448 | 0.233 | 0.00045 |
| 3 | COMMD6 | 0.468 | 0.414 | 3.991 | 3.684 | 0.114 | 4.6E-05 |
| 3 | COPS9 | 0.282 | 0.242 | 1.42 | 1.274 | 0.144 | 6.4E-05 |
| 3 | COX6A1 | 0.434 | 0.47 | 3.318 | 3.071 | 0.111 | 5E-06 |
| 3 | COX7A2 | 0.424 | 0.377 | 2.384 | 2.22 | 0.104 | 9.8E-05 |
| 3 | COX7C | 0.644 | 0.446 | 4.604 | 4.077 | 0.176 | 1.6E-12 |
| 3 | CPSF2 | 0.116 | 0.092 | 0.444 | 0.481 | 0.207 | 0.0002 |
| 3 | CRADD | 0.181 | 0.134 | 0.903 | 0.743 | 0.262 | 9.1E-09 |
| 3 | CREB1 | 0.236 | 0.202 | 1.241 | 1.147 | 0.124 | 0.00037 |
| 3 | CREBRF | 0.283 | 0.24 | 1.429 | 1.416 | 0.11 | 0.00114 |
| 3 | CREBZF | 0.134 | 0.114 | 0.646 | 0.492 | 0.148 | 0.00604 |
| 3 | CRIP1 | 0.629 | 0.417 | 4.433 | 3.289 | 0.724 | 1E-92 |
| 3 | CST7 | 0.923 | 0.824 | 14.447 | 12.219 | 0.243 | 3.2E-34 |
| 3 | CTNNA1 | 0.134 | 0.096 | 0.633 | 0.499 | 0.343 | 4.6E-09 |
| 3 | CTSS | 0.344 | 0.304 | 2.049 | 1.807 | 0.181 | 1.3E-06 |
| 3 | CUL4 | 0.246 | 0.218 | 1.318 | 1.189 | 0.149 | 8.5E-05 |
| 3 | CUL4A | 0.291 | 0.237 | 1.409 | 1.339 | 0.173 | 1.7E-07 |
| 3 | CWC14 | 0.167 | 0.146 | 0.836 | 0.746 | 0.166 | 0.00614 |
| 3 | CWC27 | 0.143 | 0.114 | 0.69 | 0.626 | 0.141 | 0.00028 |
| 3 | CX3CR1 | 0.314 | 0.247 | 1.842 | 1.697 | 0.126 | 2.1E-07 |
| 3 | CXorf38 | 0.18 | 0.14 | 0.878 | 0.798 | 0.137 | 0.00019 |
| 3 | CYBA | 0.774 | 0.691 | 7.488 | 6.914 | 0.134 | 9E-09 |
| 3 | CYLD | 0.403 | 0.442 | 3.644 | 3.299 | 0.143 | 8.1E-06 |
| 3 | CYTOR | 0.249 | 0.207 | 1.33 | 1.204 | 0.144 | 4.5E-06 |
| 3 | DAAM1 | 0.21 | 0.138 | 1.14 | 0.778 | 0.463 | 1.7E-20 |
| 3 | DAPK2 | 0.126 | 0.034 | 0.637 | 0.178 | 1.839 | 1.4E-88 |
| 3 | DCAF10 | 0.14 | 0.116 | 0.704 | 0.63 | 0.16 | 0.00099 |
| 3 | DCAF8 | 0.283 | 0.24 | 1.498 | 1.333 | 0.168 | 8.9E-06 |
| 3 | DDT | 0.178 | 0.14 | 0.877 | 0.744 | 0.216 | 0.00024 |
| 3 | DDX24 | 0.448 | 0.391 | 2.776 | 2.447 | 0.182 | 2.2E-09 |
| 3 | DDX42 | 0.13 | 0.111 | 0.613 | 0.444 | 0.148 | 0.00929 |
| 3 | DDX48 | 0.367 | 0.281 | 2.749 | 2.34 | 0.232 | 3.1E-14 |
| 3 | DDX6 | 0.497 | 0.422 | 4.284 | 3.931 | 0.124 | 4.2E-08 |
| 3 | DDX60 | 0.397 | 0.29 | 2.749 | 2.104 | 0.391 | 4.9E-24 |
| 3 | DDX60L | 0.448 | 0.33 | 3.636 | 2.748 | 0.404 | 4.5E-28 |

| | | | | | | | |
|---|---|---|---|---|---|---|---|
| 3 | DEK | 0.409 | 0.344 | 2.477 | 2.22 | 0.148 | 1.6E-06 |
| 3 | DENND10 | 0.103 | 0.086 | 0.496 | 0.434 | 0.192 | 0.00644 |
| 3 | DIAPH2 | 0.227 | 0.18 | 1.226 | 1.08 | 0.183 | 3.5E-07 |
| 3 | DMTF1 | 0.304 | 0.24 | 1.649 | 1.432 | 0.212 | 4.2E-08 |
| 3 | DNAJC19 | 0.166 | 0.13 | 0.774 | 0.672 | 0.206 | 4.2E-06 |
| 3 | DNAJC24 | 0.123 | 0.097 | 0.497 | 0.402 | 0.248 | 9.8E-05 |
| 3 | DNAJC4 | 0.116 | 0.084 | 0.436 | 0.437 | 0.293 | 6.9E-07 |
| 3 | DOCK8 | 0.634 | 0.449 | 4.024 | 4.408 | 0.147 | 2.4E-12 |
| 3 | DPP8 | 0.193 | 0.166 | 0.98 | 0.88 | 0.146 | 0.00177 |
| 3 | DR1 | 0.29 | 0.246 | 1.434 | 1.43 | 0.102 | 0.00107 |
| 3 | DSTN | 0.437 | 0.324 | 2.83 | 2.108 | 0.424 | 1.2E-26 |
| 3 | DTD1 | 0.164 | 0.139 | 0.813 | 0.741 | 0.114 | 0.00127 |
| 3 | DTX3L | 0.481 | 0.374 | 3.627 | 2.948 | 0.299 | 1.8E-19 |
| 3 | DUT | 0.204 | 0.149 | 1.001 | 0.784 | 0.34 | 2.2E-11 |
| 3 | DYNLT1 | 0.36 | 0.3 | 2.119 | 1.842 | 0.202 | 3.2E-08 |
| 3 | DYNLT3 | 0.143 | 0.128 | 0.734 | 0.641 | 0.174 | 0.00066 |
| 3 | ECHDC1 | 0.173 | 0.148 | 0.842 | 0.774 | 0.137 | 0.00212 |
| 3 | EEF1D | 0.68 | 0.499 | 4.282 | 4.813 | 0.134 | 1.4E-09 |
| 3 | EFHD2 | 0.7 | 0.607 | 4.849 | 4.248 | 0.149 | 4.2E-12 |
| 3 | EIF1B | 0.221 | 0.191 | 1.103 | 0.986 | 0.162 | 0.00068 |
| 3 | EIF2AK2 | 0.422 | 0.414 | 4.218 | 4.148 | 0.328 | 4.3E-22 |
| 3 | EIF3A | 0.346 | 0.291 | 1.917 | 1.749 | 0.124 | 1.2E-06 |
| 3 | EIF3F | 0.228 | 0.196 | 1.122 | 1.027 | 0.128 | 0.00061 |
| 3 | EIF3M | 0.22 | 0.179 | 1.068 | 0.917 | 0.22 | 2.2E-06 |
| 3 | EIF4E3 | 0.122 | 0.097 | 0.494 | 0.407 | 0.231 | 0.00014 |
| 3 | ELK3 | 0.191 | 0.149 | 1.014 | 0.869 | 0.223 | 4.4E-05 |
| 3 | EMB | 0.498 | 0.417 | 3.324 | 2.84 | 0.222 | 3.8E-12 |
| 3 | EMC7 | 0.132 | 0.111 | 0.601 | 0.446 | 0.112 | 0.00349 |
| 3 | EMSY | 0.146 | 0.124 | 0.744 | 0.66 | 0.192 | 2.2E-05 |
| 3 | ENDOD1 | 0.11 | 0.092 | 0.43 | 0.464 | 0.193 | 0.00436 |
| 3 | ENPP4 | 0.267 | 0.217 | 1.396 | 1.209 | 0.207 | 3.1E-07 |
| 3 | EOMES | 0.161 | 0.124 | 0.801 | 0.646 | 0.288 | 1.2E-06 |
| 3 | EPB41L4A | 0.134 | 0.062 | 0.664 | 0.343 | 0.944 | 1.7E-37 |
| 3 | EPDR1 | 0.168 | 0.104 | 0.894 | 0.482 | 0.622 | 2.1E-20 |
| 3 | EPS14 | 0.474 | 0.394 | 3.086 | 2.474 | 0.261 | 1.4E-13 |
| 3 | EPSTI1 | 0.438 | 0.407 | 4.627 | 4.389 | 0.349 | 3.5E-31 |
| 3 | ERCC4 | 0.244 | 0.201 | 1.248 | 1.127 | 0.149 | 8.9E-06 |
| 3 | ERGIC2 | 0.134 | 0.114 | 0.642 | 0.483 | 0.139 | 0.004 |
| 3 | ERP29 | 0.249 | 0.186 | 1.222 | 0.972 | 0.331 | 1.3E-12 |
| 3 | ERVK3-1 | 0.143 | 0.128 | 0.702 | 0.644 | 0.102 | 0.00143 |
| 3 | EVI2B | 0.344 | 0.249 | 1.926 | 1.492 | 0.369 | 4.1E-20 |
| 3 | EXOC1 | 0.149 | 0.122 | 0.746 | 0.641 | 0.239 | 1E-06 |
| 3 | EXOC2 | 0.236 | 0.194 | 1.217 | 1.069 | 0.187 | 5E-06 |
| 3 | EXOC6B | 0.199 | 0.144 | 1.003 | 0.844 | 0.249 | 4.4E-11 |
| 3 | EXOSC10 | 0.139 | 0.114 | 0.642 | 0.493 | 0.113 | 0.00074 |
| 3 | EXOSC8 | 0.166 | 0.14 | 0.793 | 0.727 | 0.127 | 0.00121 |
| 3 | EYA3 | 0.118 | 0.096 | 0.441 | 0.499 | 0.119 | 0.00111 |
| 3 | EZR | 0.432 | 0.346 | 2.666 | 2.2 | 0.278 | 1.6E-14 |
| 3 | F2R | 0.449 | 0.349 | 3.973 | 2.412 | 0.72 | 6.5E-84 |
| 3 | FAM102A | 0.108 | 0.062 | 0.414 | 0.318 | 0.698 | 1.6E-16 |
| 3 | FAM104A | 0.106 | 0.087 | 0.493 | 0.423 | 0.218 | 0.00241 |
| 3 | FAM111A | 0.239 | 0.203 | 1.216 | 1.129 | 0.107 | 0.00014 |
| 3 | FAM117A | 0.104 | 0.082 | 0.489 | 0.418 | 0.224 | 0.00029 |
| 3 | FAM118A | 0.163 | 0.106 | 0.793 | 0.469 | 0.48 | 1.4E-14 |

| | | | | | | | |
|---|---|---|---|---|---|---|---|
| 3 | FAM120AOS | 0.149 | 0.127 | 0.723 | 0.649 | 0.144 | 0.00263 |
| 3 | FAM160B1 | 0.2 | 0.143 | 0.974 | 0.747 | 0.381 | 1.1E-12 |
| 3 | FAM162A | 0.119 | 0.092 | 0.436 | 0.446 | 0.232 | 3.6E-05 |
| 3 | FAM169A | 0.134 | 0.116 | 0.679 | 0.627 | 0.114 | 0.00907 |
| 3 | FAM217B | 0.13 | 0.11 | 0.626 | 0.466 | 0.147 | 0.00322 |
| 3 | FAM228B | 0.148 | 0.124 | 0.78 | 0.64 | 0.261 | 1E-05 |
| 3 | FAM76A | 0.113 | 0.08 | 0.422 | 0.408 | 0.346 | 4.4E-08 |
| 3 | FAM76B | 0.173 | 0.144 | 0.846 | 0.741 | 0.188 | 0.00019 |
| 3 | FAS | 0.102 | 0.068 | 0.498 | 0.332 | 0.486 | 1E-09 |
| 3 | FAU | 0.976 | 0.922 | 21.687 | 18.977 | 0.193 | 1.8E-29 |
| 3 | FBXL4 | 0.347 | 0.304 | 1.913 | 1.781 | 0.103 | 7.8E-06 |
| 3 | FBXO32 | 0.122 | 0.084 | 0.476 | 0.442 | 0.382 | 4E-09 |
| 3 | FBXO42 | 0.102 | 0.083 | 0.482 | 0.429 | 0.17 | 0.00223 |
| 3 | FBXO9 | 0.218 | 0.194 | 1.12 | 1.014 | 0.143 | 0.00461 |
| 3 | FBXW11 | 0.299 | 0.243 | 1.447 | 1.381 | 0.164 | 4.1E-08 |
| 3 | FCGR3A | 0.491 | 0.402 | 4.331 | 4.768 | 0.161 | 9.7E-10 |
| 3 | FCRL6 | 0.262 | 0.183 | 1.464 | 1.071 | 0.442 | 2.5E-19 |
| 3 | FDX1 | 0.124 | 0.101 | 0.478 | 0.491 | 0.236 | 0.00034 |
| 3 | FEM1B | 0.102 | 0.077 | 0.462 | 0.38 | 0.283 | 2.1E-05 |
| 3 | FGFBP2 | 0.444 | 0.424 | 4.762 | 3.892 | 0.291 | 3.1E-22 |
| 3 | FGL2 | 0.116 | 0.049 | 0.447 | 0.31 | 0.848 | 1.1E-24 |
| 3 | FKBP8 | 0.184 | 0.146 | 0.891 | 0.741 | 0.246 | 3.3E-06 |
| 3 | FTO | 0.147 | 0.121 | 0.726 | 0.664 | 0.126 | 0.00041 |
| 3 | FTX | 0.291 | 0.233 | 1.611 | 1.413 | 0.189 | 1.2E-08 |
| 3 | GAB3 | 0.21 | 0.144 | 1.076 | 0.848 | 0.328 | 7.2E-11 |
| 3 | GABARAP | 0.422 | 0.446 | 3.114 | 2.906 | 0.1 | 1.9E-07 |
| 3 | GABARAPL1 | 0.271 | 0.224 | 1.41 | 1.279 | 0.141 | 3.4E-06 |
| 3 | GABPA | 0.128 | 0.103 | 0.606 | 0.419 | 0.224 | 0.00014 |
| 3 | GAS4 | 0.174 | 0.147 | 0.846 | 0.744 | 0.164 | 0.00031 |
| 3 | GBP1 | 0.289 | 0.188 | 2.021 | 1.447 | 0.376 | 4E-28 |
| 3 | GBP2 | 0.447 | 0.376 | 3.09 | 2.71 | 0.189 | 3.2E-10 |
| 3 | GBP3 | 0.17 | 0.134 | 0.817 | 0.717 | 0.188 | 1E-05 |
| 3 | GBP4 | 0.647 | 0.461 | 8.166 | 7.487 | 0.124 | 2.5E-13 |
| 3 | GCLM | 0.127 | 0.104 | 0.624 | 0.434 | 0.226 | 0.0004 |
| 3 | GCNT1 | 0.204 | 0.149 | 1.138 | 0.898 | 0.343 | 9E-12 |
| 3 | GIMAP4 | 0.263 | 0.217 | 1.392 | 1.293 | 0.107 | 1.1E-05 |
| 3 | GLIPR1 | 0.31 | 0.26 | 1.646 | 1.408 | 0.126 | 6.6E-06 |
| 3 | GLO1 | 0.198 | 0.173 | 0.964 | 0.893 | 0.111 | 0.00442 |
| 3 | GLOD4 | 0.134 | 0.114 | 0.628 | 0.464 | 0.142 | 0.00682 |
| 3 | GLUL | 0.124 | 0.096 | 0.497 | 0.499 | 0.249 | 1.7E-05 |
| 3 | GNB2 | 0.217 | 0.192 | 1.099 | 1.012 | 0.119 | 0.00449 |
| 3 | GNLY | 0.986 | 0.941 | 44.787 | 40.734 | 0.137 | 6.7E-20 |
| 3 | GNPTAB | 0.633 | 0.4 | 4.906 | 3.816 | 0.362 | 8.5E-34 |
| 3 | GOLGA1 | 0.112 | 0.088 | 0.424 | 0.444 | 0.208 | 0.00027 |
| 3 | GOLGA4 | 0.101 | 0.084 | 0.471 | 0.416 | 0.182 | 0.00664 |
| 3 | GPATCH2L | 0.203 | 0.173 | 1.023 | 0.931 | 0.136 | 0.00044 |
| 3 | GPHN | 0.164 | 0.109 | 0.897 | 0.601 | 0.477 | 1.9E-14 |
| 3 | GPR144 | 0.132 | 0.096 | 0.699 | 0.448 | 0.341 | 6.3E-08 |
| 3 | GPR174 | 0.21 | 0.174 | 1.061 | 0.944 | 0.167 | 3.9E-05 |
| 3 | GPR89B | 0.12 | 0.093 | 0.484 | 0.47 | 0.312 | 3.5E-05 |
| 3 | GRAP2 | 0.244 | 0.119 | 1.276 | 0.648 | 0.978 | 2.6E-63 |
| 3 | GSTK1 | 0.307 | 0.264 | 1.636 | 1.481 | 0.144 | 4.1E-05 |
| 3 | GTDC1 | 0.234 | 0.184 | 1.244 | 1.079 | 0.207 | 9.5E-08 |
| 3 | GUK1 | 0.372 | 0.314 | 2.046 | 1.822 | 0.168 | 2.2E-07 |

| | | | | | | | |
|---|---|---|---|---|---|---|---|
| 3 | GXYLT1 | 0.114 | 0.09 | 0.446 | 0.443 | 0.269 | 0.00019 |
| 3 | GZMB | 0.916 | 0.82 | 14.914 | 14.799 | 0.104 | 1.4E-11 |
| 3 | GZMH | 0.69 | 0.433 | 7.023 | 3.843 | 0.87 | 1E-143 |
| 3 | HDDC2 | 0.239 | 0.144 | 1.181 | 0.809 | 0.447 | 1.4E-24 |
| 3 | HEATR1 | 0.122 | 0.1 | 0.467 | 0.424 | 0.11 | 0.00203 |
| 3 | HELZ2 | 0.101 | 0.069 | 0.473 | 0.363 | 0.383 | 4.9E-08 |
| 3 | HERC1 | 0.477 | 0.364 | 3.104 | 2.464 | 0.276 | 4.4E-22 |
| 3 | HERC4 | 0.391 | 0.272 | 3.644 | 2.422 | 0.431 | 2.4E-32 |
| 3 | HERC4 | 0.268 | 0.227 | 1.447 | 1.312 | 0.141 | 3E-05 |
| 3 | HERC6 | 0.272 | 0.203 | 1.682 | 1.39 | 0.274 | 4.7E-13 |
| 3 | HERPUD2 | 0.332 | 0.244 | 1.861 | 1.486 | 0.324 | 8.4E-14 |
| 3 | HEXB | 0.104 | 0.088 | 0.49 | 0.433 | 0.18 | 0.00477 |
| 3 | HIBADH | 0.124 | 0.098 | 0.607 | 0.414 | 0.241 | 7.3E-05 |
| 3 | HIVEP3 | 0.292 | 0.234 | 1.497 | 1.483 | 0.107 | 2.6E-08 |
| 3 | HLA-A | 0.988 | 0.936 | 28.987 | 26.026 | 0.144 | 1.1E-23 |
| 3 | HLA-DPA1 | 0.477 | 0.244 | 2.961 | 1.398 | 1.083 | 1E-134 |
| 3 | HLA-DPB1 | 0.426 | 0.21 | 2.472 | 1.182 | 1.122 | 4E-124 |
| 3 | HLA-DRB1 | 0.346 | 0.068 | 2.376 | 0.348 | 2.731 | 0 |
| 3 | HLA-DRB4 | 0.274 | 0.027 | 1.713 | 0.12 | 3.836 | 0 |
| 3 | HMGB2 | 0.411 | 0.316 | 2.442 | 1.989 | 0.296 | 1.5E-17 |
| 3 | HPS3 | 0.236 | 0.182 | 1.207 | 0.992 | 0.284 | 6.3E-10 |
| 3 | HSPA4 | 0.211 | 0.181 | 1.047 | 0.963 | 0.134 | 0.00048 |
| 3 | HTATSF1 | 0.196 | 0.167 | 1.013 | 0.871 | 0.218 | 0.00033 |
| 3 | IBTK | 0.143 | 0.129 | 0.731 | 0.68 | 0.103 | 0.00263 |
| 3 | ICE2 | 0.119 | 0.096 | 0.442 | 0.486 | 0.186 | 0.00062 |
| 3 | IDH2 | 0.263 | 0.212 | 1.389 | 1.144 | 0.278 | 2.5E-08 |
| 3 | IFI16 | 0.647 | 0.449 | 6.384 | 4.382 | 0.246 | 1.7E-16 |
| 3 | IFI27L2 | 0.173 | 0.139 | 0.791 | 0.696 | 0.184 | 2.4E-05 |
| 3 | IFI44 | 0.309 | 0.248 | 2.386 | 2.129 | 0.164 | 2.2E-06 |
| 3 | IFI44L | 0.361 | 0.28 | 3.664 | 3.114 | 0.234 | 7.9E-14 |
| 3 | IFI6 | 0.374 | 0.248 | 3.246 | 2.48 | 0.388 | 4.7E-28 |
| 3 | IFIT1 | 0.36 | 0.23 | 3.473 | 2.6 | 0.448 | 7.4E-37 |
| 3 | IFIT2 | 0.31 | 0.239 | 2.431 | 2.163 | 0.169 | 9.4E-12 |
| 3 | IFIT3 | 0.444 | 0.323 | 6.803 | 4.124 | 0.409 | 4.5E-34 |
| 3 | IFIT4 | 0.266 | 0.2 | 1.486 | 1.184 | 0.326 | 9.4E-13 |
| 3 | IKZF3 | 0.466 | 0.449 | 3.974 | 3.344 | 0.249 | 2.1E-18 |
| 3 | IKZF4 | 0.111 | 0.086 | 0.419 | 0.444 | 0.224 | 9.4E-05 |
| 3 | IL10RA | 0.407 | 0.442 | 3.399 | 3.009 | 0.176 | 4.3E-09 |
| 3 | IL32 | 0.777 | 0.447 | 11.832 | 6.117 | 0.942 | 1E-210 |
| 3 | INO80 | 0.143 | 0.127 | 0.734 | 0.671 | 0.129 | 0.00049 |
| 3 | INO80E | 0.104 | 0.084 | 0.462 | 0.422 | 0.132 | 0.00372 |
| 3 | INTS10 | 0.144 | 0.117 | 0.727 | 0.492 | 0.294 | 7.4E-05 |
| 3 | INTS8 | 0.114 | 0.094 | 0.441 | 0.49 | 0.143 | 0.00217 |
| 3 | IRAK4 | 0.24 | 0.202 | 1.21 | 1.108 | 0.127 | 8.7E-05 |
| 3 | IRF2 | 0.331 | 0.271 | 1.891 | 1.616 | 0.227 | 7.1E-09 |
| 3 | IRF3 | 0.164 | 0.127 | 0.807 | 0.678 | 0.242 | 1.3E-06 |
| 3 | IRF7 | 0.303 | 0.249 | 1.966 | 1.749 | 0.169 | 1.4E-07 |
| 3 | IRF9 | 0.314 | 0.249 | 1.844 | 1.474 | 0.323 | 1.3E-11 |
| 3 | ISCU | 0.333 | 0.243 | 1.862 | 1.437 | 0.374 | 1.5E-16 |
| 3 | ISG14 | 0.401 | 0.306 | 3.484 | 3.113 | 0.203 | 1.1E-16 |
| 3 | ITGA1 | 0.13 | 0.088 | 0.646 | 0.403 | 0.363 | 1E-10 |
| 3 | ITGB1 | 0.683 | 0.444 | 6.091 | 4.246 | 0.417 | 4E-42 |
| 3 | IWS1 | 0.181 | 0.146 | 0.88 | 0.767 | 0.199 | 1.1E-05 |
| 3 | JADE2 | 0.131 | 0.107 | 0.614 | 0.444 | 0.148 | 0.00065 |

| | | | | | | | |
|---|---|---|---|---|---|---|---|
| 3 | JAK3 | 0.129 | 0.099 | 0.488 | 0.409 | 0.209 | 1.1E-05 |
| 3 | JAKMIP2 | 0.201 | 0.143 | 1.048 | 0.793 | 0.403 | 1.4E-13 |
| 3 | JAZF1 | 0.661 | 0.481 | 4.444 | 4.977 | 0.13 | 8.2E-10 |
| 3 | KDM7A | 0.282 | 0.214 | 1.438 | 1.237 | 0.314 | 5E-13 |
| 3 | KEAP1 | 0.119 | 0.088 | 0.462 | 0.442 | 0.349 | 1.4E-06 |
| 3 | KIAA0040 | 0.104 | 0.076 | 0.482 | 0.367 | 0.394 | 2.1E-06 |
| 3 | KIAA0232 | 0.211 | 0.174 | 1.073 | 0.937 | 0.194 | 2.2E-05 |
| 3 | KIAA1109 | 0.404 | 0.403 | 3.23 | 2.746 | 0.234 | 6.3E-18 |
| 3 | KIAA1328 | 0.121 | 0.096 | 0.487 | 0.436 | 0.13 | 0.00022 |
| 3 | KIF13B | 0.242 | 0.212 | 1.281 | 1.192 | 0.103 | 3.8E-05 |
| 3 | KIF21A | 0.224 | 0.12 | 1.18 | 0.646 | 0.847 | 3.5E-44 |
| 3 | KIF4B | 0.293 | 0.247 | 1.444 | 1.434 | 0.117 | 0.00049 |
| 3 | KIN | 0.164 | 0.141 | 0.792 | 0.733 | 0.112 | 0.00484 |
| 3 | KIR2DL3 | 0.194 | 0.104 | 1.078 | 0.623 | 0.79 | 1.3E-36 |
| 3 | KIR3DL2 | 0.141 | 0.09 | 0.712 | 0.494 | 0.424 | 9.2E-14 |
| 3 | KLRC2 | 0.876 | 0.393 | 14.473 | 3.462 | 2.128 | 0 |
| 3 | KLRC3 | 0.843 | 0.402 | 11.261 | 3.493 | 1.648 | 0 |
| 3 | KLRC4 | 0.491 | 0.141 | 3.411 | 0.929 | 1.917 | 0 |
| 3 | KLRK1 | 0.724 | 0.642 | 6.412 | 4.913 | 0.139 | 3.1E-10 |
| 3 | KMT2A | 0.388 | 0.306 | 2.264 | 1.943 | 0.221 | 2.8E-13 |
| 3 | KPNA4 | 0.192 | 0.149 | 0.961 | 0.868 | 0.148 | 0.00011 |
| 3 | KPNB1 | 0.369 | 0.311 | 2.124 | 1.936 | 0.134 | 6.4E-07 |
| 3 | KRAS | 0.379 | 0.292 | 2.173 | 1.734 | 0.326 | 7.1E-17 |
| 3 | KRTCAP2 | 0.344 | 0.286 | 1.784 | 1.472 | 0.183 | 6.7E-08 |
| 3 | L3MBTL3 | 0.116 | 0.082 | 0.443 | 0.44 | 0.304 | 7.4E-08 |
| 3 | LAG3 | 0.171 | 0.017 | 0.998 | 0.091 | 3.447 | 2E-290 |
| 3 | LAP3 | 0.242 | 0.211 | 1.348 | 1.216 | 0.149 | 0.00107 |
| 3 | LAT | 0.173 | 0.143 | 0.827 | 0.727 | 0.187 | 0.00014 |
| 3 | LATS1 | 0.143 | 0.129 | 0.74 | 0.677 | 0.128 | 0.00199 |
| 3 | LCK | 0.316 | 0.24 | 1.729 | 1.362 | 0.344 | 4.2E-14 |
| 3 | LGALS8 | 0.146 | 0.124 | 0.72 | 0.648 | 0.14 | 0.00344 |
| 3 | LIME1 | 0.104 | 0.064 | 0.443 | 0.333 | 0.706 | 4E-13 |
| 3 | LIN44 | 0.132 | 0.103 | 0.621 | 0.422 | 0.249 | 2E-05 |
| 3 | LINC00278 | 0.129 | 0.091 | 0.641 | 0.43 | 0.273 | 1.1E-08 |
| 3 | LINC00342 | 0.164 | 0.136 | 0.847 | 0.787 | 0.106 | 0.00038 |
| 3 | LINC00944 | 0.162 | 0.036 | 0.833 | 0.209 | 1.996 | 2E-141 |
| 3 | LINC01871 | 0.244 | 0.071 | 1.404 | 0.378 | 1.894 | 3E-177 |
| 3 | LINC02328 | 0.161 | 0.134 | 0.83 | 0.773 | 0.103 | 0.00108 |
| 3 | LINC02384 | 0.202 | 0.129 | 1.132 | 0.743 | 0.608 | 1.5E-21 |
| 3 | LIPA | 0.38 | 0.31 | 2.236 | 1.891 | 0.241 | 3.2E-11 |
| 3 | LMAN1 | 0.239 | 0.208 | 1.217 | 1.099 | 0.148 | 0.00041 |
| 3 | LNPK | 0.119 | 0.097 | 0.462 | 0.49 | 0.198 | 0.00061 |
| 3 | LPAR6 | 0.16 | 0.111 | 0.834 | 0.629 | 0.406 | 3.4E-12 |
| 3 | LPIN1 | 0.142 | 0.117 | 0.677 | 0.604 | 0.163 | 0.00073 |
| 3 | LPIN2 | 0.283 | 0.241 | 1.611 | 1.444 | 0.148 | 2.6E-05 |
| 3 | LRBA | 0.622 | 0.494 | 4.701 | 3.831 | 0.294 | 3.1E-28 |
| 3 | LRPAP1 | 0.203 | 0.18 | 1.014 | 0.934 | 0.12 | 0.00474 |
| 3 | LRRC40 | 0.128 | 0.107 | 0.601 | 0.446 | 0.111 | 0.00277 |
| 3 | LSM6 | 0.123 | 0.099 | 0.473 | 0.478 | 0.261 | 0.0003 |
| 3 | LSP1 | 0.488 | 0.417 | 4.202 | 3.784 | 0.141 | 1.4E-08 |
| 3 | LY74 | 0.24 | 0.194 | 1.243 | 1.094 | 0.194 | 1E-06 |
| 3 | LYAR | 0.388 | 0.288 | 2.464 | 1.881 | 0.389 | 2.3E-22 |
| 3 | MAML2 | 0.249 | 0.21 | 1.449 | 1.348 | 0.21 | 1.4E-07 |
| 3 | MAP3K4 | 0.189 | 0.144 | 0.943 | 0.841 | 0.149 | 4.9E-05 |

| | | | | | | | |
|---|---|---|---|---|---|---|---|
| 3 | MAP4 | 0.34 | 0.286 | 1.848 | 1.674 | 0.142 | 8.4E-07 |
| 3 | MAP4K4 | 0.276 | 0.241 | 1.406 | 1.397 | 0.108 | 0.00043 |
| 3 | MBD4 | 0.184 | 0.142 | 0.884 | 0.802 | 0.143 | 8.3E-05 |
| 3 | MBNL2 | 0.243 | 0.214 | 1.327 | 1.213 | 0.129 | 0.00013 |
| 3 | MBP | 0.834 | 0.747 | 8.944 | 8.184 | 0.129 | 4.1E-11 |
| 3 | MCFD2 | 0.107 | 0.091 | 0.41 | 0.444 | 0.167 | 0.00983 |
| 3 | MCOLN2 | 0.174 | 0.033 | 0.976 | 0.19 | 2.361 | 1E-179 |
| 3 | MDFIC | 0.162 | 0.119 | 0.823 | 0.642 | 0.336 | 2.9E-09 |
| 3 | MEA1 | 0.162 | 0.126 | 0.742 | 0.62 | 0.277 | 1.7E-06 |
| 3 | MED10 | 0.201 | 0.176 | 0.994 | 0.893 | 0.144 | 0.00247 |
| 3 | MED14 | 0.262 | 0.199 | 1.366 | 1.101 | 0.311 | 6.9E-12 |
| 3 | MED17 | 0.14 | 0.112 | 0.669 | 0.477 | 0.213 | 7.8E-05 |
| 3 | MEGF9 | 0.189 | 0.142 | 0.94 | 0.836 | 0.183 | 9.1E-06 |
| 3 | MGAT4A | 0.443 | 0.374 | 2.739 | 2.378 | 0.204 | 7E-11 |
| 3 | MGST3 | 0.164 | 0.114 | 0.794 | 0.474 | 0.47 | 6.8E-13 |
| 3 | MIB1 | 0.122 | 0.104 | 0.491 | 0.442 | 0.126 | 0.00495 |
| 3 | MINDY2 | 0.204 | 0.161 | 1.022 | 0.884 | 0.209 | 7.4E-07 |
| 3 | MIR4434-2HG | 0.11 | 0.088 | 0.467 | 0.4 | 0.181 | 0.00034 |
| 3 | MIR924HG | 0.262 | 0.211 | 1.322 | 1.143 | 0.197 | 1.2E-07 |
| 3 | MITD1 | 0.18 | 0.148 | 0.848 | 0.787 | 0.108 | 0.00014 |
| 3 | MLLT10 | 0.174 | 0.124 | 0.861 | 0.646 | 0.393 | 3.7E-11 |
| 3 | MLLT6 | 0.174 | 0.148 | 0.841 | 0.787 | 0.114 | 0.00079 |
| 3 | MMGT1 | 0.113 | 0.094 | 0.446 | 0.468 | 0.22 | 0.00345 |
| 3 | MORC3 | 0.314 | 0.263 | 1.784 | 1.444 | 0.208 | 2.1E-07 |
| 3 | MPC1 | 0.166 | 0.137 | 0.793 | 0.717 | 0.146 | 0.00031 |
| 3 | MRFAP1L1 | 0.108 | 0.081 | 0.417 | 0.396 | 0.386 | 6.4E-06 |
| 3 | MRPL13 | 0.117 | 0.096 | 0.432 | 0.473 | 0.17 | 0.0011 |
| 3 | MRPL20 | 0.164 | 0.141 | 0.781 | 0.71 | 0.138 | 0.00325 |
| 3 | MRPL33 | 0.141 | 0.118 | 0.683 | 0.601 | 0.183 | 0.00147 |
| 3 | MRPL41 | 0.143 | 0.126 | 0.674 | 0.629 | 0.102 | 0.00048 |
| 3 | MRPL41 | 0.127 | 0.104 | 0.466 | 0.411 | 0.147 | 0.0023 |
| 3 | MRPL47 | 0.19 | 0.164 | 0.92 | 0.837 | 0.137 | 0.00284 |
| 3 | MRPS21 | 0.247 | 0.213 | 1.3 | 1.124 | 0.21 | 1.4E-06 |
| 3 | MRPS28 | 0.104 | 0.084 | 0.484 | 0.422 | 0.197 | 0.00105 |
| 3 | MRPS30 | 0.124 | 0.1 | 0.474 | 0.497 | 0.208 | 0.00014 |
| 3 | MRPS31 | 0.16 | 0.137 | 0.774 | 0.696 | 0.144 | 0.00294 |
| 3 | MRPS34 | 0.187 | 0.147 | 0.939 | 0.812 | 0.211 | 0.00014 |
| 3 | MRPS34 | 0.129 | 0.106 | 0.612 | 0.427 | 0.217 | 0.00098 |
| 3 | MSH6 | 0.103 | 0.084 | 0.478 | 0.428 | 0.16 | 0.00306 |
| 3 | MSI2 | 0.249 | 0.213 | 1.348 | 1.233 | 0.139 | 1.2E-06 |
| 3 | MSL3 | 0.161 | 0.123 | 0.767 | 0.641 | 0.249 | 4E-07 |
| 3 | MT1X | 0.104 | 0.063 | 0.489 | 0.303 | 0.692 | 1E-14 |
| 3 | MT2A | 0.613 | 0.488 | 8.22 | 7.013 | 0.229 | 7.3E-24 |
| 3 | MTIF3 | 0.178 | 0.147 | 0.883 | 0.763 | 0.211 | 0.00011 |
| 3 | MTLN | 0.109 | 0.084 | 0.49 | 0.408 | 0.263 | 0.00014 |
| 3 | MTPN | 0.436 | 0.381 | 2.448 | 2.37 | 0.11 | 2.7E-05 |
| 3 | MX1 | 0.484 | 0.376 | 7.429 | 6.736 | 0.141 | 3.4E-18 |
| 3 | MX2 | 0.306 | 0.232 | 2.266 | 1.901 | 0.243 | 1.4E-13 |
| 3 | MXD4 | 0.114 | 0.086 | 0.434 | 0.433 | 0.306 | 4.3E-06 |
| 3 | MYCBP | 0.113 | 0.089 | 0.41 | 0.426 | 0.262 | 0.00023 |
| 3 | MYH9 | 0.412 | 0.333 | 2.462 | 2.163 | 0.187 | 1.2E-11 |
| 3 | MYL12B | 0.72 | 0.616 | 4.804 | 4.004 | 0.214 | 4.4E-18 |
| 3 | MYO1F | 0.482 | 0.408 | 3.116 | 2.819 | 0.144 | 4.6E-09 |
| 3 | MYO4A | 0.19 | 0.144 | 0.931 | 0.817 | 0.188 | 4.2E-08 |

| | | | | | | | |
|---|---|---|---|---|---|---|---|
| 3 | MYO6 | 0.23 | 0.072 | 1.242 | 0.404 | 1.619 | 4E-134 |
| 3 | MYO9A | 0.198 | 0.166 | 1.041 | 0.946 | 0.137 | 0.00017 |
| 3 | N4BP1 | 0.332 | 0.273 | 1.993 | 1.716 | 0.216 | 1.5E-08 |
| 3 | NAA38 | 0.184 | 0.161 | 0.883 | 0.818 | 0.11 | 0.00475 |
| 3 | NAA40 | 0.436 | 0.314 | 2.671 | 1.949 | 0.448 | 6.6E-31 |
| 3 | NACA | 0.807 | 0.742 | 8.303 | 7.743 | 0.101 | 4.3E-06 |
| 3 | NBPF12 | 0.101 | 0.081 | 0.448 | 0.427 | 0.103 | 0.00182 |
| 3 | NBPF14 | 0.144 | 0.129 | 0.738 | 0.678 | 0.123 | 0.00135 |
| 3 | NCBP2 | 0.114 | 0.094 | 0.44 | 0.447 | 0.264 | 0.00146 |
| 3 | NCK1 | 0.276 | 0.231 | 1.446 | 1.267 | 0.2 | 3.9E-06 |
| 3 | NCOR1 | 0.424 | 0.463 | 3.441 | 3.242 | 0.128 | 1.2E-06 |
| 3 | NDUFA1 | 0.478 | 0.418 | 2.816 | 2.444 | 0.141 | 2.7E-06 |
| 3 | NDUFB10 | 0.248 | 0.214 | 1.287 | 1.13 | 0.188 | 4.8E-06 |
| 3 | NDUFB3 | 0.106 | 0.088 | 0.49 | 0.407 | 0.268 | 0.00281 |
| 3 | NDUFB4 | 0.226 | 0.187 | 1.107 | 0.944 | 0.229 | 6.9E-06 |
| 3 | NDUFC1 | 0.218 | 0.166 | 1.034 | 0.864 | 0.261 | 1.8E-09 |
| 3 | NDUFS1 | 0.194 | 0.147 | 0.924 | 0.824 | 0.167 | 1.2E-05 |
| 3 | NDUFS4 | 0.243 | 0.21 | 1.234 | 1.106 | 0.149 | 6.3E-06 |
| 3 | NDUFV2 | 0.309 | 0.264 | 1.628 | 1.484 | 0.133 | 3.8E-05 |
| 3 | NEDD1 | 0.14 | 0.111 | 0.663 | 0.474 | 0.204 | 4.4E-05 |
| 3 | NEK1 | 0.214 | 0.184 | 1.109 | 1.014 | 0.128 | 0.00072 |
| 3 | NEK9 | 0.114 | 0.087 | 0.433 | 0.438 | 0.283 | 2.3E-05 |
| 3 | NEO1 | 0.116 | 0.076 | 0.434 | 0.393 | 0.447 | 3.3E-11 |
| 3 | NF1 | 0.312 | 0.262 | 1.742 | 1.62 | 0.112 | 6.9E-06 |
| 3 | NFATC2IP | 0.149 | 0.127 | 0.722 | 0.646 | 0.139 | 0.00273 |
| 3 | NFX1 | 0.116 | 0.09 | 0.43 | 0.464 | 0.191 | 4E-05 |
| 3 | NIBAN1 | 0.274 | 0.218 | 1.463 | 1.273 | 0.201 | 3.7E-09 |
| 3 | NKTR | 0.418 | 0.364 | 2.414 | 2.317 | 0.118 | 4.8E-06 |
| 3 | NLRC4 | 0.244 | 0.207 | 1.34 | 1.211 | 0.146 | 4E-05 |
| 3 | NMI | 0.204 | 0.174 | 1.104 | 1.004 | 0.138 | 0.00063 |
| 3 | NMRK1 | 0.236 | 0.188 | 1.214 | 1.044 | 0.216 | 1.2E-07 |
| 3 | NMT1 | 0.141 | 0.12 | 0.696 | 0.604 | 0.203 | 2.7E-05 |
| 3 | NNT | 0.164 | 0.136 | 0.796 | 0.738 | 0.109 | 0.00079 |
| 3 | NOL7 | 0.206 | 0.164 | 0.994 | 0.872 | 0.189 | 1.8E-06 |
| 3 | NPC1 | 0.347 | 0.299 | 2.069 | 1.879 | 0.139 | 3.2E-07 |
| 3 | NPTN | 0.164 | 0.129 | 0.766 | 0.674 | 0.184 | 4.8E-06 |
| 3 | NR1D2 | 0.218 | 0.164 | 1.114 | 0.916 | 0.281 | 1.4E-10 |
| 3 | NR2C2 | 0.203 | 0.174 | 1.047 | 0.966 | 0.114 | 0.00126 |
| 3 | NRF1 | 0.148 | 0.128 | 0.748 | 0.693 | 0.13 | 0.00018 |
| 3 | NSD3 | 0.389 | 0.314 | 2.241 | 1.896 | 0.241 | 2.1E-12 |
| 3 | NSF | 0.39 | 0.314 | 2.396 | 2.004 | 0.247 | 4.4E-13 |
| 3 | NSL1 | 0.143 | 0.11 | 0.662 | 0.447 | 0.248 | 4.4E-06 |
| 3 | NSMCE4A | 0.103 | 0.084 | 0.474 | 0.426 | 0.144 | 0.00307 |
| 3 | NUB1 | 0.497 | 0.414 | 3.462 | 3.008 | 0.203 | 3.2E-12 |
| 3 | NUP107 | 0.132 | 0.111 | 0.648 | 0.493 | 0.149 | 0.00262 |
| 3 | NUP40 | 0.132 | 0.113 | 0.627 | 0.482 | 0.107 | 0.00697 |
| 3 | NUTM2B-AS1 | 0.22 | 0.184 | 1.109 | 1.002 | 0.147 | 8.9E-05 |
| 3 | OAS1 | 0.268 | 0.199 | 1.824 | 1.499 | 0.284 | 4.2E-13 |
| 3 | OAS2 | 0.382 | 0.293 | 2.702 | 2.241 | 0.263 | 1.5E-14 |
| 3 | OASL | 0.432 | 0.328 | 4.192 | 2.907 | 0.837 | 1.7E-91 |
| 3 | OAZ1 | 0.442 | 0.473 | 3.691 | 3.181 | 0.214 | 3.2E-10 |
| 3 | ODF2L | 0.334 | 0.284 | 2.16 | 1.844 | 0.229 | 8E-07 |
| 3 | OGA | 0.413 | 0.441 | 3.408 | 3.174 | 0.102 | 7.7E-07 |
| 3 | OGFR | 0.144 | 0.121 | 0.746 | 0.631 | 0.243 | 1.1E-05 |

| | | | | | | | |
|---|---|---|---|---|---|---|---|
| 3 | OOEP | 0.367 | 0.31 | 2.104 | 1.82 | 0.209 | 4.7E-08 |
| 3 | ORC3 | 0.104 | 0.08 | 0.494 | 0.4 | 0.304 | 3.9E-05 |
| 3 | ORC4 | 0.102 | 0.081 | 0.487 | 0.423 | 0.204 | 0.00069 |
| 3 | ORMDL1 | 0.344 | 0.274 | 1.811 | 1.464 | 0.211 | 6.5E-11 |
| 3 | OST4 | 0.419 | 0.363 | 2.36 | 2.144 | 0.139 | 1.3E-06 |
| 3 | OTUB1 | 0.147 | 0.134 | 0.724 | 0.649 | 0.136 | 0.00429 |
| 3 | OTUD4 | 0.166 | 0.139 | 0.811 | 0.726 | 0.161 | 0.00048 |
| 3 | OTUD6B-AS1 | 0.144 | 0.134 | 0.766 | 0.712 | 0.106 | 0.00741 |
| 3 | OXCT1 | 0.142 | 0.131 | 0.732 | 0.683 | 0.101 | 0.00712 |
| 3 | PALLD | 0.106 | 0.048 | 0.403 | 0.293 | 0.78 | 2.4E-19 |
| 3 | PALM2-AKAP2 | 0.106 | 0.088 | 0.444 | 0.401 | 0.147 | 0.00462 |
| 3 | PANK2 | 0.139 | 0.1 | 0.648 | 0.416 | 0.341 | 1.9E-08 |
| 3 | PANK3 | 0.104 | 0.087 | 0.486 | 0.444 | 0.129 | 0.00738 |
| 3 | PARP11 | 0.184 | 0.16 | 0.968 | 0.9 | 0.106 | 0.00481 |
| 3 | PARP12 | 0.184 | 0.132 | 0.944 | 0.729 | 0.389 | 4.6E-12 |
| 3 | PARP14 | 0.483 | 0.391 | 3.893 | 3.332 | 0.224 | 3.1E-14 |
| 3 | PARP14 | 0.303 | 0.249 | 1.614 | 1.441 | 0.144 | 7.4E-07 |
| 3 | PARP9 | 0.399 | 0.291 | 2.977 | 2.089 | 0.411 | 2.7E-27 |
| 3 | PATL2 | 0.182 | 0.136 | 0.921 | 0.736 | 0.323 | 3.4E-09 |
| 3 | PAXX | 0.379 | 0.303 | 2.208 | 1.8 | 0.294 | 2E-13 |
| 3 | PCED1B | 0.373 | 0.304 | 2.267 | 1.943 | 0.222 | 4.1E-10 |
| 3 | PCGF4 | 0.449 | 0.377 | 2.766 | 2.433 | 0.127 | 2.4E-08 |
| 3 | PCID2 | 0.143 | 0.12 | 0.696 | 0.63 | 0.144 | 0.00129 |
| 3 | PCM1 | 0.41 | 0.443 | 3.324 | 3.09 | 0.106 | 8.3E-07 |
| 3 | PCNX2 | 0.149 | 0.092 | 0.817 | 0.487 | 0.747 | 9.5E-24 |
| 3 | PCYT1A | 0.142 | 0.119 | 0.744 | 0.617 | 0.269 | 4.5E-06 |
| 3 | PDCD10 | 0.217 | 0.182 | 1.071 | 0.968 | 0.147 | 0.0001 |
| 3 | PDCD4 | 0.141 | 0.128 | 0.704 | 0.641 | 0.112 | 0.0028 |
| 3 | PDE3B | 0.436 | 0.403 | 4.444 | 3.479 | 0.343 | 7.8E-31 |
| 3 | PDE4DIP | 0.117 | 0.093 | 0.443 | 0.469 | 0.21 | 0.00022 |
| 3 | PDHB | 0.143 | 0.113 | 0.649 | 0.473 | 0.202 | 2.9E-05 |
| 3 | PDLIM4 | 0.129 | 0.109 | 0.614 | 0.467 | 0.116 | 0.00417 |
| 3 | PEBP1 | 0.172 | 0.147 | 0.843 | 0.719 | 0.229 | 0.00118 |
| 3 | PEX1 | 0.129 | 0.098 | 0.604 | 0.4 | 0.272 | 3.3E-06 |
| 3 | PFDN4 | 0.449 | 0.488 | 3.684 | 3.378 | 0.124 | 2.4E-07 |
| 3 | PGAM1 | 0.194 | 0.148 | 0.923 | 0.737 | 0.323 | 6.9E-09 |
| 3 | PHC3 | 0.296 | 0.249 | 1.488 | 1.441 | 0.141 | 4.7E-06 |
| 3 | PHF11 | 0.391 | 0.294 | 2.441 | 1.883 | 0.38 | 6.3E-21 |
| 3 | PHF3 | 0.482 | 0.409 | 2.996 | 2.762 | 0.118 | 6.8E-08 |
| 3 | PHIP | 0.387 | 0.317 | 2.27 | 2 | 0.183 | 1.4E-09 |
| 3 | PI4K2B | 0.187 | 0.131 | 0.989 | 0.702 | 0.494 | 7.8E-14 |
| 3 | PICALM | 0.471 | 0.398 | 2.961 | 2.669 | 0.14 | 1.5E-09 |
| 3 | PITPNB | 0.24 | 0.206 | 1.279 | 1.114 | 0.198 | 3.1E-06 |
| 3 | PKM | 0.181 | 0.143 | 0.9 | 0.764 | 0.237 | 1.8E-06 |
| 3 | PLAAT3 | 0.189 | 0.14 | 0.917 | 0.728 | 0.332 | 6.2E-10 |
| 3 | PLAAT4 | 0.713 | 0.613 | 6.691 | 6.048 | 0.143 | 7.9E-13 |
| 3 | PLCL1 | 0.124 | 0.071 | 0.704 | 0.428 | 0.719 | 3.2E-19 |
| 3 | PLCXD2 | 0.104 | 0.082 | 0.47 | 0.433 | 0.118 | 0.00044 |
| 3 | PLEKHA7 | 0.116 | 0.097 | 0.424 | 0.464 | 0.174 | 0.00386 |
| 3 | PLEKHF1 | 0.141 | 0.114 | 0.679 | 0.482 | 0.223 | 0.00018 |
| 3 | PLEKHF2 | 0.172 | 0.134 | 0.847 | 0.687 | 0.302 | 1.1E-06 |
| 3 | PLP2 | 0.139 | 0.106 | 0.684 | 0.469 | 0.264 | 1.7E-06 |
| 3 | PMS1 | 0.161 | 0.13 | 0.774 | 0.683 | 0.182 | 9.2E-05 |
| 3 | PNPT1 | 0.221 | 0.16 | 1.3 | 0.984 | 0.4 | 3.4E-13 |

| | | | | | | | |
|---|---|---|---|---|---|---|---|
| 3 | POLK | 0.132 | 0.101 | 0.641 | 0.432 | 0.27 | 3.8E-06 |
| 3 | POLR2I | 0.103 | 0.081 | 0.463 | 0.379 | 0.287 | 0.00031 |
| 3 | PPDPF | 0.446 | 0.347 | 2.789 | 2.27 | 0.297 | 1E-17 |
| 3 | PPM1K | 0.348 | 0.266 | 2.331 | 1.74 | 0.422 | 3E-20 |
| 3 | PPP1CC | 0.24 | 0.174 | 1.247 | 0.91 | 0.467 | 6.4E-14 |
| 3 | PPP1R21 | 0.17 | 0.14 | 0.821 | 0.744 | 0.14 | 0.00019 |
| 3 | PPP2R2B | 0.37 | 0.3 | 2.134 | 1.947 | 0.126 | 6.8E-10 |
| 3 | PPP2R4C | 0.686 | 0.472 | 4.849 | 4.74 | 0.303 | 2.4E-28 |
| 3 | PPP3CB | 0.208 | 0.17 | 1.061 | 0.913 | 0.217 | 4.4E-06 |
| 3 | PPP3CC | 0.401 | 0.33 | 2.349 | 2.086 | 0.171 | 7.7E-10 |
| 3 | PPP4R3B | 0.36 | 0.314 | 2.03 | 1.89 | 0.103 | 9.3E-05 |
| 3 | PPWD1 | 0.191 | 0.163 | 0.961 | 0.844 | 0.169 | 0.00103 |
| 3 | PRDM1 | 0.421 | 0.32 | 2.704 | 2.143 | 0.329 | 2.5E-20 |
| 3 | PRELID1 | 0.244 | 0.206 | 1.22 | 1.093 | 0.148 | 7.8E-05 |
| 3 | PREPL | 0.132 | 0.111 | 0.644 | 0.484 | 0.164 | 0.00201 |
| 3 | PRKAG2 | 0.162 | 0.13 | 0.796 | 0.716 | 0.142 | 4.2E-05 |
| 3 | PRKCI | 0.101 | 0.071 | 0.444 | 0.379 | 0.264 | 4.1E-07 |
| 3 | PRKY | 0.163 | 0.123 | 0.829 | 0.678 | 0.291 | 4.5E-08 |
| 3 | PRPF3 | 0.132 | 0.104 | 0.623 | 0.446 | 0.192 | 8.4E-05 |
| 3 | PRPF39 | 0.143 | 0.122 | 0.699 | 0.634 | 0.14 | 0.00371 |
| 3 | PRPSAP1 | 0.111 | 0.083 | 0.417 | 0.407 | 0.344 | 8.2E-06 |
| 3 | PRSS23 | 0.231 | 0.189 | 1.324 | 1.177 | 0.171 | 6.8E-06 |
| 3 | PSIP1 | 0.328 | 0.273 | 1.79 | 1.472 | 0.188 | 1.7E-07 |
| 3 | PSMA2 | 0.171 | 0.144 | 0.804 | 0.724 | 0.14 | 0.00066 |
| 3 | PSMA3 | 0.174 | 0.143 | 0.876 | 0.803 | 0.126 | 0.00802 |
| 3 | PSMB3 | 0.248 | 0.218 | 1.222 | 1.121 | 0.124 | 0.0022 |
| 3 | PSMB9 | 0.734 | 0.62 | 6.971 | 6.003 | 0.216 | 7.5E-20 |
| 3 | PSMC1 | 0.204 | 0.178 | 1 | 0.919 | 0.121 | 0.00285 |
| 3 | PSMC2 | 0.126 | 0.103 | 0.476 | 0.407 | 0.184 | 0.00097 |
| 3 | PTBP3 | 0.478 | 0.496 | 3.946 | 3.467 | 0.19 | 3.3E-12 |
| 3 | PTGER4 | 0.341 | 0.272 | 2.044 | 1.7 | 0.266 | 1.8E-11 |
| 3 | PTGES3 | 0.414 | 0.341 | 2.343 | 2.002 | 0.233 | 9.6E-12 |
| 3 | PTMS | 0.144 | 0.022 | 0.737 | 0.118 | 2.644 | 2E-181 |
| 3 | PTP4A2 | 0.144 | 0.116 | 0.678 | 0.486 | 0.211 | 4.4E-05 |
| 3 | PTPN22 | 0.462 | 0.38 | 3.017 | 2.607 | 0.211 | 1.5E-11 |
| 3 | PTPRC | 0.992 | 0.979 | 34.218 | 31.383 | 0.166 | 1.7E-34 |
| 3 | PTPRE | 0.377 | 0.334 | 2.291 | 2.08 | 0.139 | 9E-05 |
| 3 | PTPRJ | 0.44 | 0.37 | 2.804 | 2.47 | 0.126 | 9.4E-09 |
| 3 | PTPRM | 0.103 | 0.044 | 0.447 | 0.326 | 0.748 | 8.1E-21 |
| 3 | PURA | 0.167 | 0.119 | 0.84 | 0.609 | 0.481 | 3.1E-11 |
| 3 | PUS7L | 0.108 | 0.091 | 0.401 | 0.46 | 0.124 | 0.00904 |
| 3 | PWWP2A | 0.122 | 0.103 | 0.481 | 0.436 | 0.116 | 0.00414 |
| 3 | PYHIN1 | 0.772 | 0.666 | 8.166 | 7.014 | 0.219 | 2.2E-21 |
| 3 | PYURF | 0.306 | 0.264 | 1.487 | 1.448 | 0.122 | 7.9E-05 |
| 3 | R3HCC1L | 0.191 | 0.148 | 0.934 | 0.774 | 0.271 | 8.9E-08 |
| 3 | RAB10 | 0.243 | 0.217 | 1.289 | 1.194 | 0.11 | 0.00021 |
| 3 | RAB14 | 0.348 | 0.316 | 1.964 | 1.817 | 0.112 | 0.00025 |
| 3 | RAB22A | 0.187 | 0.144 | 0.89 | 0.74 | 0.246 | 9.4E-08 |
| 3 | RAB28 | 0.122 | 0.098 | 0.486 | 0.419 | 0.176 | 0.00037 |
| 3 | RAB9A | 0.14 | 0.112 | 0.646 | 0.449 | 0.234 | 7.4E-05 |
| 3 | RABAC1 | 0.413 | 0.347 | 2.376 | 2.084 | 0.188 | 2.5E-08 |
| 3 | RABGAP1L | 0.804 | 0.707 | 8.463 | 7.641 | 0.147 | 6.7E-17 |
| 3 | RAD23B | 0.342 | 0.286 | 1.791 | 1.662 | 0.108 | 1.4E-06 |
| 3 | RAD40 | 0.149 | 0.134 | 0.806 | 0.747 | 0.11 | 0.00142 |

| | | | | | | | |
|---|---|---|---|---|---|---|---|
| 3 | RAD41B | 0.279 | 0.218 | 1.619 | 1.391 | 0.219 | 3.8E-10 |
| 3 | RALGAPB | 0.181 | 0.141 | 0.843 | 0.768 | 0.133 | 2E-06 |
| 3 | RALGDS | 0.144 | 0.113 | 0.671 | 0.601 | 0.148 | 2.6E-05 |
| 3 | RAPGEF1 | 0.279 | 0.213 | 1.424 | 1.277 | 0.147 | 7.5E-11 |
| 3 | RBFOX2 | 0.178 | 0.141 | 0.839 | 0.781 | 0.103 | 0.001 |
| 3 | RBM48 | 0.181 | 0.146 | 0.884 | 0.746 | 0.227 | 1.6E-05 |
| 3 | RBPJ | 0.268 | 0.228 | 1.432 | 1.382 | 0.149 | 4.1E-05 |
| 3 | RBX1 | 0.177 | 0.148 | 0.898 | 0.764 | 0.233 | 0.00024 |
| 3 | RCBTB2 | 0.129 | 0.092 | 0.644 | 0.484 | 0.433 | 1E-08 |
| 3 | RECQL | 0.241 | 0.202 | 1.306 | 1.133 | 0.204 | 1.1E-07 |
| 3 | REEP4 | 0.347 | 0.291 | 1.9 | 1.638 | 0.214 | 4.9E-10 |
| 3 | RELCH | 0.241 | 0.2 | 1.243 | 1.102 | 0.186 | 7.5E-06 |
| 3 | RER1 | 0.147 | 0.123 | 0.74 | 0.611 | 0.277 | 5E-06 |
| 3 | RESF1 | 0.702 | 0.492 | 7.086 | 4.726 | 0.308 | 6.1E-24 |
| 3 | RFX3-AS1 | 0.164 | 0.138 | 0.836 | 0.768 | 0.122 | 0.00071 |
| 3 | RGS10 | 0.104 | 0.083 | 0.421 | 0.431 | 0.273 | 0.00063 |
| 3 | RGS9 | 0.178 | 0.083 | 0.861 | 0.446 | 0.94 | 1.1E-47 |
| 3 | RHOT1 | 0.163 | 0.137 | 0.793 | 0.713 | 0.143 | 0.00128 |
| 3 | RMDN1 | 0.143 | 0.121 | 0.644 | 0.608 | 0.107 | 0.00446 |
| 3 | RNASEH2B | 0.272 | 0.203 | 1.436 | 1.132 | 0.343 | 5E-14 |
| 3 | RNF114 | 0.126 | 0.101 | 0.493 | 0.402 | 0.24 | 0.00024 |
| 3 | RNF13 | 0.216 | 0.178 | 1.071 | 0.97 | 0.143 | 2.7E-05 |
| 3 | RNF134 | 0.101 | 0.079 | 0.466 | 0.391 | 0.242 | 0.00028 |
| 3 | RNF149 | 0.474 | 0.389 | 2.937 | 2.423 | 0.219 | 3.4E-13 |
| 3 | RNF167 | 0.136 | 0.11 | 0.61 | 0.446 | 0.132 | 0.00035 |
| 3 | RNF168 | 0.244 | 0.203 | 1.333 | 1.124 | 0.246 | 3E-08 |
| 3 | RNF169 | 0.108 | 0.09 | 0.421 | 0.474 | 0.133 | 0.00845 |
| 3 | RNF19A | 0.448 | 0.446 | 3.933 | 3.292 | 0.247 | 4.5E-17 |
| 3 | RNF213 | 0.91 | 0.844 | 14.072 | 13.214 | 0.19 | 4.5E-19 |
| 3 | RNF213-AS1 | 0.126 | 0.104 | 0.489 | 0.428 | 0.149 | 0.00239 |
| 3 | RNF34 | 0.164 | 0.137 | 0.786 | 0.716 | 0.134 | 0.00047 |
| 3 | RNF6 | 0.126 | 0.103 | 0.602 | 0.424 | 0.199 | 0.00065 |
| 3 | RNF7 | 0.3 | 0.268 | 1.604 | 1.464 | 0.132 | 0.00122 |
| 3 | RNPC3 | 0.226 | 0.182 | 1.131 | 0.98 | 0.206 | 1.2E-06 |
| 3 | RORA | 0.648 | 0.47 | 4.476 | 4.804 | 0.214 | 1.4E-14 |
| 3 | RPL11 | 0.978 | 0.932 | 23.362 | 21.402 | 0.126 | 2.5E-14 |
| 3 | RPL14 | 0.92 | 0.842 | 14.232 | 12.648 | 0.17 | 6.1E-18 |
| 3 | RPL18A | 0.849 | 0.771 | 10.133 | 9.081 | 0.148 | 4.2E-13 |
| 3 | RPL19 | 0.963 | 0.907 | 19.771 | 17.909 | 0.143 | 3.2E-17 |
| 3 | RPL21 | 0.83 | 0.74 | 9.244 | 8.211 | 0.173 | 1.5E-14 |
| 3 | RPL23A | 0.686 | 0.483 | 4.472 | 4.78 | 0.221 | 2E-17 |
| 3 | RPL26 | 0.768 | 0.69 | 7.644 | 6.742 | 0.179 | 3.2E-12 |
| 3 | RPL27A | 0.403 | 0.406 | 3.211 | 2.641 | 0.282 | 9.4E-18 |
| 3 | RPL28 | 0.972 | 0.916 | 22.068 | 19.478 | 0.173 | 2.1E-21 |
| 3 | RPL29 | 0.846 | 0.774 | 10.014 | 9.227 | 0.118 | 1.9E-07 |
| 3 | RPL30 | 0.978 | 0.93 | 24.771 | 21.707 | 0.191 | 4E-29 |
| 3 | RPL32 | 0.941 | 0.874 | 16.121 | 14.449 | 0.148 | 8.7E-20 |
| 3 | RPL34 | 0.96 | 0.899 | 20.011 | 17.641 | 0.182 | 2.8E-23 |
| 3 | RPL34 | 0.741 | 0.664 | 6.843 | 6.208 | 0.143 | 4.4E-10 |
| 3 | RPL34A | 0.937 | 0.861 | 16.048 | 13.498 | 0.239 | 2.5E-34 |
| 3 | RPL36 | 0.788 | 0.701 | 7.416 | 6.843 | 0.134 | 4.3E-09 |
| 3 | RPL36A | 0.743 | 0.622 | 6.864 | 4.449 | 0.331 | 1.3E-30 |
| 3 | RPL36AL | 0.624 | 0.444 | 4.404 | 4.083 | 0.109 | 1.1E-07 |
| 3 | RPL37A | 0.747 | 0.681 | 7.184 | 6.401 | 0.144 | 1.7E-09 |

| | | | | | | | |
|---|---|---|---|---|---|---|---|
| 3 | RPL38 | 0.742 | 0.647 | 7.047 | 4.884 | 0.26 | 6.1E-22 |
| 3 | RPL39 | 0.914 | 0.818 | 13.397 | 11.289 | 0.247 | 1.5E-32 |
| 3 | RPL41 | 0.988 | 0.937 | 32.91 | 28.417 | 0.207 | 1.1E-34 |
| 3 | RPL7A | 0.949 | 0.897 | 17.937 | 16.296 | 0.138 | 6.4E-16 |
| 3 | RPLP0 | 0.849 | 0.743 | 9.446 | 8.734 | 0.13 | 2E-11 |
| 3 | RPLP1 | 0.994 | 0.944 | 40.249 | 36.716 | 0.133 | 1.4E-20 |
| 3 | RPLP2 | 0.987 | 0.942 | 28.122 | 24.93 | 0.174 | 3.5E-28 |
| 3 | RPRD2 | 0.226 | 0.184 | 1.214 | 1.014 | 0.248 | 2E-06 |
| 3 | RPS11 | 0.644 | 0.474 | 4.988 | 4.432 | 0.171 | 3.7E-11 |
| 3 | RPS12 | 0.988 | 0.937 | 30.449 | 24.894 | 0.296 | 6.3E-67 |
| 3 | RPS14 | 0.942 | 0.884 | 19.418 | 14.872 | 0.291 | 1.4E-41 |
| 3 | RPS14 | 0.846 | 0.746 | 9.441 | 8.378 | 0.174 | 9E-16 |
| 3 | RPS14A | 0.93 | 0.868 | 14.928 | 14.611 | 0.124 | 1E-10 |
| 3 | RPS16 | 0.679 | 0.493 | 4.392 | 4.89 | 0.141 | 2.1E-09 |
| 3 | RPS18 | 0.747 | 0.646 | 7.019 | 6.134 | 0.194 | 1.9E-14 |
| 3 | RPS19 | 0.872 | 0.779 | 11.484 | 9.894 | 0.228 | 2E-26 |
| 3 | RPS2 | 0.482 | 0.414 | 4.397 | 3.967 | 0.149 | 2.9E-07 |
| 3 | RPS20 | 0.381 | 0.334 | 2.128 | 1.984 | 0.101 | 6.5E-05 |
| 3 | RPS21 | 0.906 | 0.817 | 13.193 | 11.394 | 0.211 | 4.1E-23 |
| 3 | RPS23 | 0.88 | 0.809 | 11.693 | 10.746 | 0.12 | 2E-10 |
| 3 | RPS24 | 0.499 | 0.429 | 4.293 | 3.971 | 0.113 | 2.4E-06 |
| 3 | RPS26 | 0.891 | 0.834 | 18.497 | 16.414 | 0.171 | 6.2E-11 |
| 3 | RPS27 | 0.966 | 0.899 | 24.22 | 19.73 | 0.296 | 1.2E-41 |
| 3 | RPS27A | 0.996 | 0.944 | 33.969 | 29.417 | 0.203 | 4.2E-41 |
| 3 | RPS27L | 0.234 | 0.191 | 1.144 | 1.063 | 0.119 | 4.2E-06 |
| 3 | RPS28 | 0.946 | 0.887 | 18.268 | 16.402 | 0.147 | 6.6E-17 |
| 3 | RPS29 | 0.868 | 0.766 | 11.734 | 9.612 | 0.288 | 2.9E-34 |
| 3 | RPS3 | 0.978 | 0.91 | 23.348 | 18.324 | 0.34 | 7.1E-79 |
| 3 | RPS4X | 0.881 | 0.784 | 11.063 | 9.448 | 0.211 | 1.4E-20 |
| 3 | RPS4Y1 | 0.276 | 0.221 | 1.483 | 1.344 | 0.224 | 1.7E-08 |
| 3 | RPS7 | 0.949 | 0.892 | 17.972 | 16.664 | 0.109 | 1.5E-09 |
| 3 | RPS8 | 0.989 | 0.94 | 30.609 | 27.494 | 0.14 | 2.4E-23 |
| 3 | RSAD2 | 0.296 | 0.208 | 2.368 | 1.774 | 0.414 | 1.2E-20 |
| 3 | RSRC1 | 0.217 | 0.171 | 1.148 | 0.96 | 0.248 | 4.1E-08 |
| 3 | RTTN | 0.104 | 0.076 | 0.401 | 0.412 | 0.282 | 2E-06 |
| 3 | RUBCN | 0.104 | 0.069 | 0.474 | 0.361 | 0.397 | 7.2E-10 |
| 3 | RUFY2 | 0.124 | 0.104 | 0.494 | 0.442 | 0.133 | 0.0072 |
| 3 | RUNX1 | 0.398 | 0.329 | 2.41 | 2.194 | 0.134 | 2.1E-09 |
| 3 | S100A4 | 0.787 | 0.641 | 14.918 | 11.764 | 0.343 | 4.1E-30 |
| 3 | S100A6 | 0.693 | 0.406 | 6.777 | 4.42 | 0.484 | 3.2E-73 |
| 3 | S1PR1 | 0.13 | 0.104 | 0.614 | 0.443 | 0.14 | 0.00043 |
| 3 | SAMD9 | 0.444 | 0.444 | 4.286 | 4.248 | 0.312 | 4.4E-22 |
| 3 | SAMD9L | 0.424 | 0.396 | 4.334 | 3.844 | 0.473 | 9.1E-33 |
| 3 | SAMSN1 | 0.388 | 0.266 | 2.408 | 1.826 | 0.448 | 7.1E-32 |
| 3 | SCAF11 | 0.649 | 0.484 | 4.241 | 4.714 | 0.146 | 2.9E-09 |
| 3 | SCAMP2 | 0.14 | 0.108 | 0.639 | 0.432 | 0.264 | 9.1E-06 |
| 3 | SCP2 | 0.316 | 0.262 | 1.647 | 1.496 | 0.148 | 8.1E-07 |
| 3 | SDCCAG8 | 0.107 | 0.084 | 0.488 | 0.434 | 0.167 | 0.00025 |
| 3 | SDHA | 0.181 | 0.149 | 0.864 | 0.8 | 0.111 | 0.00012 |
| 3 | SDHC | 0.166 | 0.141 | 0.792 | 0.703 | 0.172 | 0.00163 |
| 3 | SEC62 | 0.418 | 0.372 | 2.428 | 2.334 | 0.114 | 7.6E-05 |
| 3 | SECISBP2L | 0.334 | 0.282 | 1.844 | 1.668 | 0.146 | 1.5E-06 |
| 3 | SEL1L | 0.303 | 0.233 | 1.468 | 1.281 | 0.292 | 8E-13 |
| 3 | SELENOF | 0.203 | 0.176 | 0.993 | 0.904 | 0.133 | 0.00212 |

| | | | | | | | |
|---|---|---|---|---|---|---|---|
| 3 | SELENOS | 0.204 | 0.169 | 0.989 | 0.884 | 0.16 | 6.9E-05 |
| 3 | SELENOT | 0.247 | 0.22 | 1.303 | 1.198 | 0.121 | 0.00019 |
| 3 | SEM1 | 0.29 | 0.244 | 1.488 | 1.324 | 0.168 | 1.4E-05 |
| 3 | SENP6 | 0.434 | 0.362 | 2.724 | 2.322 | 0.231 | 6.1E-11 |
| 3 | SERINC1 | 0.407 | 0.348 | 2.292 | 2.081 | 0.14 | 4.2E-07 |
| 3 | SERPINB9 | 0.161 | 0.109 | 0.743 | 0.48 | 0.377 | 6.5E-13 |
| 3 | SESN3 | 0.204 | 0.087 | 1.044 | 0.472 | 1.149 | 6.7E-69 |
| 3 | SETBP1 | 0.138 | 0.077 | 0.669 | 0.437 | 0.613 | 4.8E-23 |
| 3 | SETDB2 | 0.169 | 0.119 | 0.836 | 0.618 | 0.434 | 6E-12 |
| 3 | SF3B2 | 0.462 | 0.403 | 2.842 | 2.443 | 0.144 | 1.6E-07 |
| 3 | SFXN1 | 0.141 | 0.112 | 0.667 | 0.478 | 0.206 | 9.8E-05 |
| 3 | SGCD | 0.377 | 0.103 | 2.932 | 0.74 | 1.967 | 1E-287 |
| 3 | SH3BGRL | 0.442 | 0.397 | 2.7 | 2.414 | 0.102 | 2.7E-05 |
| 3 | SH3BGRL3 | 0.921 | 0.808 | 14.113 | 12.47 | 0.277 | 1.2E-39 |
| 3 | SHFL | 0.261 | 0.211 | 1.471 | 1.226 | 0.262 | 1.1E-07 |
| 3 | SHISA4 | 0.2 | 0.167 | 0.987 | 0.87 | 0.182 | 0.00012 |
| 3 | SIK2 | 0.122 | 0.094 | 0.492 | 0.404 | 0.229 | 6.4E-05 |
| 3 | SKIL | 0.388 | 0.321 | 2.478 | 2.114 | 0.229 | 3E-10 |
| 3 | SLAMF6 | 0.141 | 0.118 | 0.673 | 0.608 | 0.146 | 0.00171 |
| 3 | SLC14A4 | 0.266 | 0.223 | 1.442 | 1.288 | 0.162 | 1.4E-05 |
| 3 | SLC20A1 | 0.288 | 0.229 | 1.49 | 1.313 | 0.277 | 7.5E-10 |
| 3 | SLC24A24 | 0.132 | 0.074 | 0.646 | 0.384 | 0.74 | 3.2E-21 |
| 3 | SLC30A7 | 0.141 | 0.13 | 0.729 | 0.68 | 0.101 | 0.00604 |
| 3 | SLC34E2B | 0.119 | 0.097 | 0.462 | 0.494 | 0.183 | 0.00099 |
| 3 | SLC34F1 | 0.147 | 0.123 | 0.731 | 0.608 | 0.266 | 3.8E-06 |
| 3 | SLC3A2 | 0.234 | 0.192 | 1.172 | 1.038 | 0.174 | 3E-06 |
| 3 | SLC4A7 | 0.303 | 0.187 | 1.743 | 1.081 | 0.697 | 1.3E-40 |
| 3 | SLC66A3 | 0.112 | 0.092 | 0.417 | 0.444 | 0.186 | 0.00189 |
| 3 | SLC6A6 | 0.106 | 0.074 | 0.422 | 0.387 | 0.43 | 1E-07 |
| 3 | SLC9A9 | 0.224 | 0.18 | 1.144 | 1.039 | 0.143 | 4.8E-07 |
| 3 | SLF1 | 0.108 | 0.086 | 0.437 | 0.446 | 0.234 | 0.00063 |
| 3 | SLFN12L | 0.601 | 0.492 | 4.133 | 4.02 | 0.343 | 4E-26 |
| 3 | SLFN4 | 0.491 | 0.406 | 4.708 | 4.744 | 0.266 | 4.9E-16 |
| 3 | SMARCA4 | 0.377 | 0.324 | 2.143 | 1.972 | 0.119 | 6.1E-06 |
| 3 | SMARCAD1 | 0.166 | 0.136 | 0.833 | 0.714 | 0.219 | 0.00012 |
| 3 | SMARCE1 | 0.274 | 0.24 | 1.434 | 1.337 | 0.103 | 0.00046 |
| 3 | SMCHD1 | 0.74 | 0.647 | 7.774 | 6.64 | 0.228 | 1.4E-18 |
| 3 | SMIM7 | 0.134 | 0.116 | 0.639 | 0.491 | 0.113 | 0.00789 |
| 3 | SNAP23 | 0.208 | 0.179 | 1.04 | 0.966 | 0.119 | 0.00104 |
| 3 | SNHG14 | 0.137 | 0.101 | 0.641 | 0.437 | 0.278 | 1.9E-07 |
| 3 | SNHG6 | 0.248 | 0.201 | 1.266 | 1.073 | 0.239 | 4.4E-07 |
| 3 | SNRNP200 | 0.214 | 0.179 | 1.044 | 0.967 | 0.126 | 6.3E-05 |
| 3 | SNRPA | 0.11 | 0.088 | 0.499 | 0.434 | 0.198 | 0.00072 |
| 3 | SNX17 | 0.124 | 0.106 | 0.466 | 0.416 | 0.132 | 0.00903 |
| 3 | SNX20 | 0.136 | 0.108 | 0.66 | 0.443 | 0.244 | 4.4E-05 |
| 3 | SNX4 | 0.143 | 0.124 | 0.734 | 0.643 | 0.193 | 0.00017 |
| 3 | SOD1 | 0.264 | 0.219 | 1.337 | 1.171 | 0.191 | 1.7E-06 |
| 3 | SOS1 | 0.371 | 0.32 | 2.231 | 2.078 | 0.103 | 1.4E-05 |
| 3 | SP100 | 0.776 | 0.674 | 9.276 | 7.427 | 0.301 | 6.1E-26 |
| 3 | SP110 | 0.444 | 0.446 | 4.407 | 3.747 | 0.234 | 1.1E-13 |
| 3 | SP140 | 0.344 | 0.262 | 2.041 | 1.624 | 0.329 | 1.3E-14 |
| 3 | SP4 | 0.162 | 0.138 | 0.804 | 0.721 | 0.149 | 0.0014 |
| 3 | SPATA13 | 0.338 | 0.283 | 1.873 | 1.678 | 0.149 | 3.3E-07 |
| 3 | SPCS1 | 0.384 | 0.344 | 2.134 | 1.984 | 0.106 | 0.00025 |

| | | | | | | | |
|---|---|---|---|---|---|---|---|
| 3 | SPG11 | 0.24 | 0.214 | 1.333 | 1.212 | 0.138 | 0.00024 |
| 3 | SPOCK2 | 0.289 | 0.207 | 1.796 | 1.286 | 0.482 | 6.2E-20 |
| 3 | SPSB3 | 0.294 | 0.244 | 1.498 | 1.364 | 0.134 | 1.8E-06 |
| 3 | SRGAP2 | 0.14 | 0.093 | 0.672 | 0.49 | 0.446 | 2.2E-12 |
| 3 | SRPRA | 0.336 | 0.282 | 1.784 | 1.446 | 0.198 | 2.4E-07 |
| 3 | SSB | 0.223 | 0.184 | 1.116 | 1.001 | 0.146 | 1.8E-05 |
| 3 | SSR1 | 0.197 | 0.169 | 0.983 | 0.897 | 0.132 | 0.001 |
| 3 | ST3GAL4 | 0.124 | 0.092 | 0.474 | 0.464 | 0.311 | 1.3E-06 |
| 3 | ST8SIA4 | 0.32 | 0.249 | 1.813 | 1.447 | 0.22 | 2.4E-09 |
| 3 | ST8SIA6 | 0.189 | 0.16 | 1.01 | 0.923 | 0.13 | 0.00079 |
| 3 | STAMBP | 0.114 | 0.084 | 0.426 | 0.446 | 0.239 | 7.3E-06 |
| 3 | STARD7 | 0.184 | 0.147 | 0.902 | 0.767 | 0.234 | 1.3E-06 |
| 3 | STAT1 | 0.473 | 0.479 | 6.773 | 6.201 | 0.127 | 1.6E-13 |
| 3 | STAT2 | 0.27 | 0.218 | 1.46 | 1.443 | 0.112 | 6.8E-07 |
| 3 | STAT3 | 0.41 | 0.347 | 2.409 | 2.164 | 0.144 | 7.4E-08 |
| 3 | STAT4B | 0.238 | 0.166 | 1.244 | 0.926 | 0.427 | 2.1E-17 |
| 3 | STK38 | 0.464 | 0.486 | 3.872 | 3.428 | 0.134 | 9E-09 |
| 3 | STUB1 | 0.14 | 0.12 | 0.648 | 0.473 | 0.179 | 0.00471 |
| 3 | STX17 | 0.161 | 0.139 | 0.794 | 0.738 | 0.107 | 0.0062 |
| 3 | STXBP3 | 0.204 | 0.176 | 1.019 | 0.94 | 0.1 | 0.00138 |
| 3 | STXBP4 | 0.207 | 0.171 | 1.084 | 0.977 | 0.142 | 4.1E-05 |
| 3 | SUCLA2 | 0.129 | 0.108 | 0.603 | 0.462 | 0.102 | 0.00386 |
| 3 | SUDS3 | 0.208 | 0.168 | 1.009 | 0.879 | 0.199 | 4.2E-06 |
| 3 | SUN2 | 0.232 | 0.194 | 1.167 | 1.08 | 0.112 | 4.3E-05 |
| 3 | SUPT20H | 0.224 | 0.19 | 1.116 | 1.016 | 0.136 | 0.00025 |
| 3 | SUPT4H1 | 0.142 | 0.119 | 0.672 | 0.48 | 0.212 | 0.00131 |
| 3 | SURF4 | 0.137 | 0.108 | 0.636 | 0.448 | 0.216 | 3.9E-05 |
| 3 | SVIP | 0.12 | 0.092 | 0.448 | 0.473 | 0.239 | 2.6E-05 |
| 3 | SYF2 | 0.247 | 0.218 | 1.274 | 1.18 | 0.111 | 0.00014 |
| 3 | SYNE1 | 0.862 | 0.774 | 11.404 | 10.443 | 0.112 | 8.7E-14 |
| 3 | SYNE2 | 0.929 | 0.829 | 14.492 | 12.383 | 0.323 | 7.1E-43 |
| 3 | SYNGR1 | 0.143 | 0.108 | 0.717 | 0.468 | 0.336 | 4.1E-07 |
| 3 | SYNJ1 | 0.113 | 0.089 | 0.427 | 0.446 | 0.208 | 0.00024 |
| 3 | SYT11 | 0.301 | 0.197 | 1.624 | 1.046 | 0.634 | 4.7E-32 |
| 3 | SYTL2 | 0.263 | 0.212 | 1.472 | 1.28 | 0.202 | 4.3E-07 |
| 3 | TAF1B | 0.104 | 0.082 | 0.491 | 0.402 | 0.291 | 0.00041 |
| 3 | TAF7 | 0.284 | 0.244 | 1.406 | 1.368 | 0.139 | 7.4E-05 |
| 3 | TANK | 0.317 | 0.247 | 1.849 | 1.464 | 0.24 | 9.4E-09 |
| 3 | TAOK1 | 0.204 | 0.171 | 1.043 | 0.942 | 0.161 | 0.00011 |
| 3 | TAP1 | 0.346 | 0.281 | 2.014 | 1.718 | 0.23 | 7E-10 |
| 3 | TAP2 | 0.311 | 0.266 | 1.804 | 1.603 | 0.171 | 9.4E-06 |
| 3 | TBC1D23 | 0.18 | 0.146 | 0.882 | 0.816 | 0.112 | 0.0034 |
| 3 | TBC1D2B | 0.16 | 0.092 | 0.799 | 0.487 | 0.714 | 2.7E-24 |
| 3 | TBCA | 0.31 | 0.267 | 1.647 | 1.404 | 0.131 | 3E-05 |
| 3 | TBCD | 0.174 | 0.114 | 0.89 | 0.608 | 0.44 | 3E-17 |
| 3 | TBK1 | 0.247 | 0.213 | 1.32 | 1.194 | 0.144 | 6.3E-06 |
| 3 | TBL1XR1 | 0.348 | 0.291 | 2.004 | 1.864 | 0.104 | 1.2E-06 |
| 3 | TCAF2 | 0.179 | 0.147 | 0.923 | 0.848 | 0.123 | 0.0077 |
| 3 | TCEA1 | 0.418 | 0.363 | 2.402 | 2.203 | 0.124 | 1.1E-05 |
| 3 | TCTEX1D2 | 0.101 | 0.067 | 0.487 | 0.346 | 0.494 | 3.3E-09 |
| 3 | TDP2 | 0.136 | 0.109 | 0.641 | 0.448 | 0.2 | 0.00011 |
| 3 | TDRD3 | 0.182 | 0.118 | 0.937 | 0.63 | 0.473 | 1.5E-18 |
| 3 | TECR | 0.161 | 0.124 | 0.746 | 0.633 | 0.246 | 2.4E-06 |
| 3 | TENT4A | 0.182 | 0.147 | 0.944 | 0.827 | 0.191 | 1.4E-05 |

| | | | | | | | |
|---|---|---|---|---|---|---|---|
| 3 | TERF1 | 0.173 | 0.14 | 0.868 | 0.804 | 0.108 | 0.00364 |
| 3 | TGFBR2 | 0.37 | 0.304 | 2.066 | 1.882 | 0.134 | 1E-08 |
| 3 | TGIF1 | 0.11 | 0.089 | 0.403 | 0.444 | 0.18 | 0.00093 |
| 3 | THAP12 | 0.167 | 0.128 | 0.801 | 0.649 | 0.28 | 4.9E-07 |
| 3 | THRA | 0.13 | 0.09 | 0.496 | 0.464 | 0.347 | 2E-09 |
| 3 | TIGIT | 0.377 | 0.29 | 2.431 | 2.149 | 0.178 | 1.4E-14 |
| 3 | TM9SF2 | 0.334 | 0.274 | 1.812 | 1.466 | 0.21 | 9.7E-09 |
| 3 | TMBIM6 | 0.861 | 0.78 | 9.424 | 8.614 | 0.13 | 1.5E-10 |
| 3 | TMCC3 | 0.141 | 0.094 | 0.701 | 0.413 | 0.44 | 1.7E-12 |
| 3 | TMCO1 | 0.274 | 0.228 | 1.392 | 1.24 | 0.167 | 2.5E-06 |
| 3 | TMED4 | 0.232 | 0.204 | 1.184 | 1.047 | 0.163 | 0.00164 |
| 3 | TMEM106B | 0.196 | 0.17 | 1 | 0.887 | 0.173 | 0.00193 |
| 3 | TMEM109 | 0.174 | 0.14 | 0.847 | 0.749 | 0.16 | 0.00143 |
| 3 | TMEM117 | 0.128 | 0.097 | 0.641 | 0.461 | 0.192 | 4.2E-06 |
| 3 | TMEM14B | 0.239 | 0.149 | 1.274 | 0.849 | 0.487 | 4.7E-23 |
| 3 | TMEM170A | 0.146 | 0.134 | 0.749 | 0.69 | 0.119 | 0.0047 |
| 3 | TMEM248 | 0.14 | 0.118 | 0.647 | 0.494 | 0.144 | 0.0023 |
| 3 | TMEM30A | 0.371 | 0.303 | 2.088 | 1.748 | 0.248 | 1E-10 |
| 3 | TMEM34B | 0.13 | 0.106 | 0.627 | 0.424 | 0.247 | 0.00033 |
| 3 | TMEM40A | 0.426 | 0.428 | 3.278 | 2.749 | 0.249 | 2.2E-16 |
| 3 | TMEM43 | 0.143 | 0.124 | 0.743 | 0.637 | 0.241 | 4.1E-05 |
| 3 | TMEM62 | 0.128 | 0.104 | 0.623 | 0.44 | 0.18 | 0.001 |
| 3 | TMEM87B | 0.11 | 0.091 | 0.418 | 0.444 | 0.223 | 0.00214 |
| 3 | TMPO | 0.294 | 0.249 | 1.482 | 1.436 | 0.14 | 8.7E-06 |
| 3 | TMSB4X | 0.988 | 0.938 | 31.903 | 27.724 | 0.203 | 7E-37 |
| 3 | TMX1 | 0.212 | 0.174 | 1.049 | 0.904 | 0.228 | 1.5E-05 |
| 3 | TOMM20 | 0.204 | 0.172 | 0.984 | 0.888 | 0.141 | 0.00041 |
| 3 | TOMM4 | 0.229 | 0.198 | 1.119 | 1.029 | 0.121 | 0.00088 |
| 3 | TOMM7 | 0.411 | 0.424 | 3.269 | 2.71 | 0.271 | 1.1E-14 |
| 3 | TOX4 | 0.266 | 0.224 | 1.306 | 1.206 | 0.114 | 4.5E-05 |
| 3 | TPRG1 | 0.111 | 0.016 | 0.469 | 0.084 | 2.744 | 2E-143 |
| 3 | TRAFD1 | 0.164 | 0.128 | 0.838 | 0.706 | 0.247 | 6.5E-07 |
| 3 | TRANK1 | 0.396 | 0.332 | 2.619 | 2.298 | 0.189 | 1.8E-08 |
| 3 | TRAPPC1 | 0.243 | 0.23 | 1.304 | 1.212 | 0.106 | 0.00999 |
| 3 | TRAPPC2 | 0.141 | 0.116 | 0.696 | 0.499 | 0.214 | 0.00065 |
| 3 | TRBC1 | 0.641 | 0.442 | 4.782 | 3.663 | 0.648 | 9.1E-89 |
| 3 | TRBC2 | 0.441 | 0.446 | 3.708 | 3.123 | 0.248 | 7.4E-14 |
| 3 | TRDC | 0.372 | 0.307 | 2.164 | 1.944 | 0.147 | 3E-08 |
| 3 | TRERF1 | 0.141 | 0.113 | 0.729 | 0.637 | 0.194 | 2.8E-07 |
| 3 | TRG-AS1 | 0.473 | 0.218 | 3.692 | 1.408 | 1.391 | 2E-176 |
| 3 | TRGV7 | 0.27 | 0.204 | 1.423 | 1.169 | 0.284 | 4.8E-12 |
| 3 | TRIM14 | 0.203 | 0.166 | 1.074 | 0.944 | 0.171 | 2.2E-05 |
| 3 | TRIM21 | 0.104 | 0.084 | 0.407 | 0.43 | 0.237 | 0.003 |
| 3 | TRIM22 | 0.462 | 0.448 | 4.631 | 4.411 | 0.32 | 8.1E-24 |
| 3 | TRIM23 | 0.101 | 0.082 | 0.446 | 0.417 | 0.128 | 0.00262 |
| 3 | TRIM24 | 0.164 | 0.124 | 0.849 | 0.674 | 0.329 | 4.4E-08 |
| 3 | TRIM38 | 0.271 | 0.222 | 1.431 | 1.291 | 0.148 | 1.8E-06 |
| 3 | TRIM4 | 0.236 | 0.199 | 1.231 | 1.097 | 0.164 | 4.1E-05 |
| 3 | TRIM4 | 0.122 | 0.09 | 0.447 | 0.441 | 0.303 | 1.6E-06 |
| 3 | TRIP11 | 0.196 | 0.167 | 0.999 | 0.93 | 0.103 | 0.00172 |
| 3 | TRMT112 | 0.262 | 0.228 | 1.323 | 1.209 | 0.129 | 0.00044 |
| 3 | TRPC4AP | 0.192 | 0.149 | 0.96 | 0.888 | 0.112 | 0.00012 |
| 3 | TSC22D1 | 0.132 | 0.112 | 0.64 | 0.494 | 0.109 | 0.00762 |
| 3 | TSEN44 | 0.17 | 0.144 | 0.816 | 0.744 | 0.13 | 0.001 |

| | | | | | | | |
|---|---|---|---|---|---|---|---|
| 3 | TSPAN2 | 0.134 | 0.09 | 0.636 | 0.447 | 0.409 | 4.2E-12 |
| 3 | TSR2 | 0.138 | 0.116 | 0.632 | 0.478 | 0.131 | 0.00331 |
| 3 | TTC1 | 0.227 | 0.19 | 1.106 | 0.998 | 0.147 | 4.7E-05 |
| 3 | TTC13 | 0.12 | 0.099 | 0.469 | 0.431 | 0.101 | 0.00144 |
| 3 | TTC17 | 0.413 | 0.343 | 2.404 | 2.314 | 0.114 | 1.8E-06 |
| 3 | TTC33 | 0.118 | 0.094 | 0.444 | 0.48 | 0.183 | 0.00064 |
| 3 | TTC39C | 0.4 | 0.31 | 2.41 | 2.018 | 0.246 | 8.2E-16 |
| 3 | TUT7 | 0.424 | 0.363 | 2.614 | 2.348 | 0.149 | 2.8E-07 |
| 3 | TXNDC9 | 0.128 | 0.104 | 0.612 | 0.417 | 0.243 | 0.0004 |
| 3 | TYMP | 0.121 | 0.098 | 0.603 | 0.418 | 0.219 | 0.00076 |
| 3 | TYROBP | 0.878 | 0.803 | 11.142 | 10.263 | 0.12 | 3.3E-08 |
| 3 | UBA42 | 0.797 | 0.707 | 7.684 | 6.923 | 0.141 | 2.1E-10 |
| 3 | UBE2A | 0.18 | 0.148 | 0.844 | 0.791 | 0.112 | 0.00497 |
| 3 | UBE2N | 0.17 | 0.147 | 0.848 | 0.744 | 0.17 | 0.00197 |
| 3 | UBE2W | 0.164 | 0.137 | 0.813 | 0.711 | 0.194 | 0.00034 |
| 3 | UBL3 | 0.267 | 0.186 | 1.423 | 1.033 | 0.462 | 3.5E-20 |
| 3 | UBR2 | 0.344 | 0.282 | 1.992 | 1.726 | 0.207 | 2.3E-11 |
| 3 | UBXN4 | 0.477 | 0.413 | 3.038 | 2.713 | 0.163 | 2.1E-07 |
| 3 | UBXN7 | 0.171 | 0.134 | 0.829 | 0.734 | 0.173 | 5E-06 |
| 3 | UCHL4 | 0.224 | 0.197 | 1.142 | 1.048 | 0.11 | 0.00409 |
| 3 | UFL1 | 0.209 | 0.179 | 1.033 | 0.944 | 0.114 | 0.0012 |
| 3 | UHMK1 | 0.347 | 0.294 | 1.848 | 1.707 | 0.123 | 4.5E-06 |
| 3 | UMAD1 | 0.217 | 0.181 | 1.143 | 1.037 | 0.141 | 4.8E-05 |
| 3 | UPF3A | 0.199 | 0.173 | 1.008 | 0.939 | 0.102 | 0.00429 |
| 3 | UPP1 | 0.288 | 0.22 | 1.44 | 1.244 | 0.317 | 9.4E-13 |
| 3 | UQCRB | 0.439 | 0.349 | 2.401 | 2.114 | 0.241 | 3E-14 |
| 3 | USP1 | 0.169 | 0.141 | 0.879 | 0.738 | 0.242 | 0.00024 |
| 3 | USP18 | 0.148 | 0.112 | 0.807 | 0.616 | 0.389 | 1.4E-07 |
| 3 | USP28 | 0.341 | 0.284 | 1.911 | 1.69 | 0.178 | 1.6E-07 |
| 3 | USP42 | 0.138 | 0.112 | 0.668 | 0.49 | 0.178 | 0.0004 |
| 3 | USP9Y | 0.183 | 0.138 | 0.963 | 0.788 | 0.289 | 6.2E-09 |
| 3 | UTY | 0.169 | 0.134 | 0.824 | 0.76 | 0.119 | 1.5E-05 |
| 3 | UXT | 0.232 | 0.204 | 1.148 | 1.049 | 0.129 | 0.00472 |
| 3 | VAMP2 | 0.446 | 0.486 | 3.491 | 3.326 | 0.111 | 6E-06 |
| 3 | VCPIP1 | 0.194 | 0.144 | 0.943 | 0.818 | 0.206 | 2.1E-06 |
| 3 | VEZF1 | 0.273 | 0.234 | 1.431 | 1.292 | 0.147 | 9E-05 |
| 3 | VIM | 0.427 | 0.376 | 4.4 | 3.233 | 0.477 | 8.6E-44 |
| 3 | VIM-AS1 | 0.111 | 0.092 | 0.421 | 0.466 | 0.162 | 0.00338 |
| 3 | VPS13A | 0.329 | 0.268 | 1.848 | 1.601 | 0.207 | 2.8E-09 |
| 3 | VPS28 | 0.311 | 0.274 | 1.649 | 1.412 | 0.124 | 0.00047 |
| 3 | VPS34 | 0.146 | 0.131 | 0.712 | 0.662 | 0.107 | 0.00184 |
| 3 | VPS41 | 0.164 | 0.138 | 0.806 | 0.734 | 0.134 | 0.00115 |
| 3 | VRK2 | 0.202 | 0.164 | 1.02 | 0.904 | 0.174 | 1.2E-05 |
| 3 | WAPL | 0.244 | 0.222 | 1.336 | 1.236 | 0.113 | 0.00091 |
| 3 | WASHC3 | 0.141 | 0.111 | 0.736 | 0.444 | 0.41 | 4.8E-09 |
| 3 | WDR44 | 0.12 | 0.094 | 0.474 | 0.493 | 0.22 | 0.0001 |
| 3 | WNK1 | 0.333 | 0.244 | 1.809 | 1.408 | 0.262 | 7.4E-14 |
| 3 | XAF1 | 0.448 | 0.444 | 4.076 | 4.178 | 0.281 | 1.1E-18 |
| 3 | XIAP | 0.234 | 0.208 | 1.248 | 1.144 | 0.113 | 0.00447 |
| 3 | XPC | 0.264 | 0.224 | 1.397 | 1.277 | 0.13 | 3.7E-05 |
| 3 | XPO4 | 0.183 | 0.141 | 0.88 | 0.762 | 0.209 | 4.2E-07 |
| 3 | XRN1 | 0.644 | 0.447 | 4.714 | 4.116 | 0.16 | 1.8E-13 |
| 3 | YIPF4 | 0.109 | 0.09 | 0.42 | 0.449 | 0.18 | 0.00319 |
| 3 | YPEL3 | 0.374 | 0.313 | 2.131 | 1.918 | 0.142 | 4.8E-08 |

| | | | | | | | |
|---|---|---|---|---|---|---|---|
| 3 | YTHDF2 | 0.178 | 0.149 | 0.847 | 0.767 | 0.142 | 0.00036 |
| 3 | YWHAH | 0.107 | 0.08 | 0.4 | 0.378 | 0.401 | 9.9E-06 |
| 3 | YWHAQ | 0.479 | 0.39 | 3.004 | 2.443 | 0.24 | 3.1E-14 |
| 3 | ZBP1 | 0.334 | 0.237 | 1.999 | 1.414 | 0.401 | 1.6E-22 |
| 3 | ZBTB2 | 0.124 | 0.104 | 0.602 | 0.461 | 0.1 | 0.00786 |
| 3 | ZBTB20 | 0.72 | 0.616 | 7.882 | 6.67 | 0.241 | 1.1E-20 |
| 3 | ZBTB38 | 0.417 | 0.234 | 2.497 | 1.343 | 0.884 | 4.4E-86 |
| 3 | ZBTB44 | 0.212 | 0.181 | 1.066 | 0.989 | 0.108 | 0.0004 |
| 3 | ZBTB7A | 0.224 | 0.171 | 1.149 | 0.934 | 0.31 | 3.9E-10 |
| 3 | ZC3H6 | 0.162 | 0.12 | 0.803 | 0.649 | 0.306 | 1.4E-08 |
| 3 | ZC3HAV1 | 0.491 | 0.424 | 3.29 | 3.02 | 0.123 | 1.2E-07 |
| 3 | ZCCHC8 | 0.112 | 0.088 | 0.499 | 0.434 | 0.199 | 0.00021 |
| 3 | ZEB2 | 0.838 | 0.734 | 9.878 | 8.418 | 0.214 | 4.4E-27 |
| 3 | ZFP90 | 0.118 | 0.099 | 0.442 | 0.402 | 0.136 | 0.00438 |
| 3 | ZFR | 0.369 | 0.319 | 2.07 | 1.914 | 0.112 | 1.8E-05 |
| 3 | ZFYVE28 | 0.211 | 0.149 | 1.109 | 0.841 | 0.381 | 2.3E-14 |
| 3 | ZKSCAN1 | 0.124 | 0.107 | 0.627 | 0.443 | 0.207 | 0.00641 |
| 3 | ZNF107 | 0.14 | 0.103 | 0.682 | 0.444 | 0.328 | 1.1E-07 |
| 3 | ZNF146 | 0.131 | 0.113 | 0.636 | 0.466 | 0.168 | 0.00868 |
| 3 | ZNF217 | 0.284 | 0.241 | 1.481 | 1.387 | 0.189 | 8.4E-06 |
| 3 | ZNF224 | 0.106 | 0.084 | 0.473 | 0.427 | 0.147 | 0.00066 |
| 3 | ZNF24 | 0.176 | 0.144 | 0.849 | 0.782 | 0.12 | 0.00822 |
| 3 | ZNF407 | 0.141 | 0.12 | 0.684 | 0.63 | 0.118 | 0.00441 |
| 3 | ZNF638 | 0.369 | 0.322 | 2.192 | 2 | 0.132 | 2.2E-05 |
| 3 | ZNF642 | 0.238 | 0.208 | 1.271 | 1.164 | 0.126 | 0.00144 |
| 3 | ZNF721 | 0.198 | 0.174 | 1.018 | 0.934 | 0.123 | 0.00715 |
| 3 | ZNF737 | 0.109 | 0.084 | 0.499 | 0.438 | 0.189 | 0.00025 |
| 3 | ZNF827 | 0.131 | 0.102 | 0.632 | 0.449 | 0.202 | 2.5E-05 |
| 3 | ZNF91 | 0.232 | 0.203 | 1.187 | 1.103 | 0.104 | 0.002 |
| 3 | ZNFX1 | 0.232 | 0.186 | 1.244 | 1.083 | 0.2 | 4.8E-07 |
| 3 | ZNRF2 | 0.189 | 0.144 | 0.943 | 0.862 | 0.131 | 6.2E-05 |
| 3 | ZZEF1 | 0.201 | 0.163 | 0.98 | 0.908 | 0.111 | 2.3E-05 |
| 4 | ABCE1 | 0.13 | 0.084 | 0.4 | 0.408 | 0.29 | 9.8E-05 |
| 4 | ABHD17B | 0.104 | 0.072 | 0.449 | 0.349 | 0.321 | 0.00289 |
| 4 | ABHD2 | 0.337 | 0.222 | 1.43 | 1.31 | 0.224 | 4.3E-09 |
| 4 | ABI2 | 0.117 | 0.073 | 0.441 | 0.373 | 0.242 | 4.9E-05 |
| 4 | ABLIM1 | 0.341 | 0.227 | 1.7 | 1.346 | 0.326 | 1.1E-10 |
| 4 | AC004448.2 | 0.111 | 0.046 | 0.424 | 0.223 | 0.927 | 2.8E-14 |
| 4 | AC004462.1 | 0.121 | 0.079 | 0.423 | 0.397 | 0.394 | 0.00014 |
| 4 | AC007384.1 | 0.179 | 0.11 | 0.797 | 0.481 | 0.446 | 1E-07 |
| 4 | AC008014.1 | 0.263 | 0.172 | 1.301 | 1.046 | 0.314 | 4.5E-08 |
| 4 | AC008104.3 | 0.129 | 0.061 | 0.478 | 0.31 | 0.624 | 4.8E-12 |
| 4 | AC018816.1 | 0.146 | 0.049 | 0.692 | 0.304 | 1.187 | 4.9E-20 |
| 4 | AC022074.1 | 0.241 | 0.167 | 1.003 | 0.88 | 0.189 | 1.5E-05 |
| 4 | AC024164.1 | 0.171 | 0.072 | 0.798 | 0.371 | 1.104 | 3.2E-21 |
| 4 | AC024171.2 | 0.127 | 0.044 | 0.483 | 0.266 | 0.849 | 4.6E-14 |
| 4 | AC044849.1 | 0.176 | 0.098 | 0.843 | 0.429 | 0.671 | 8.8E-11 |
| 4 | AC103491.3 | 0.137 | 0.071 | 1.601 | 0.704 | 1.184 | 4.2E-11 |
| 4 | AC118449.1 | 0.193 | 0.138 | 0.796 | 0.732 | 0.122 | 0.00065 |
| 4 | AC138123.1 | 0.131 | 0.063 | 0.437 | 0.301 | 0.834 | 4.2E-12 |
| 4 | AC239799.2 | 0.101 | 0.036 | 0.744 | 0.248 | 1.604 | 4.3E-18 |
| 4 | AC243472.2 | 0.224 | 0.124 | 1.966 | 1.111 | 0.823 | 3.1E-14 |
| 4 | AC244014.3 | 0.109 | 0.062 | 0.47 | 0.409 | 0.477 | 2.1E-06 |
| 4 | AC244402.1 | 0.226 | 0.148 | 0.933 | 0.816 | 0.194 | 7.2E-05 |

| | | | | | | | |
|---|---|---|---|---|---|---|---|
| 4 | ACAP1 | 0.279 | 0.14 | 1.247 | 0.794 | 0.661 | 1E-17 |
| 4 | ACBD6 | 0.246 | 0.174 | 1.048 | 0.963 | 0.122 | 6.3E-05 |
| 4 | ACTG1 | 0.741 | 0.614 | 7.232 | 4.931 | 0.286 | 7.9E-07 |
| 4 | ADAM28 | 0.13 | 0.091 | 0.469 | 0.42 | 0.13 | 0.00162 |
| 4 | ADGRE4 | 0.46 | 0.342 | 2.923 | 2.447 | 0.247 | 1.5E-06 |
| 4 | ADH4 | 0.243 | 0.171 | 1.014 | 0.872 | 0.217 | 2.4E-06 |
| 4 | ADK | 0.337 | 0.21 | 1.716 | 1.244 | 0.464 | 1.6E-12 |
| 4 | AF111167.1 | 0.141 | 0.094 | 0.842 | 0.462 | 0.499 | 9.5E-07 |
| 4 | AFF1 | 0.269 | 0.193 | 1.187 | 1.098 | 0.112 | 8.3E-05 |
| 4 | AFF3 | 0.281 | 0.011 | 1.444 | 0.064 | 4.476 | 0 |
| 4 | AFG3L2 | 0.213 | 0.124 | 0.87 | 0.627 | 0.472 | 1.7E-10 |
| 4 | AGK | 0.219 | 0.144 | 0.834 | 0.764 | 0.127 | 4.2E-06 |
| 4 | AGO2 | 0.206 | 0.133 | 0.841 | 0.731 | 0.22 | 1.4E-06 |
| 4 | AGPAT4 | 0.174 | 0.034 | 0.777 | 0.168 | 2.207 | 4.4E-76 |
| 4 | AHI1 | 0.479 | 0.163 | 4.42 | 0.984 | 2.168 | 1E-109 |
| 4 | AHR | 0.343 | 0.044 | 1.879 | 0.232 | 3.016 | 5E-242 |
| 4 | AHSA1 | 0.107 | 0.071 | 0.421 | 0.338 | 0.313 | 0.00072 |
| 4 | AIMP1 | 0.211 | 0.124 | 0.873 | 0.629 | 0.474 | 4.9E-10 |
| 4 | AK2 | 0.179 | 0.112 | 0.693 | 0.444 | 0.324 | 7.4E-07 |
| 4 | AKAP11 | 0.186 | 0.127 | 0.779 | 0.649 | 0.241 | 6.4E-05 |
| 4 | AKAP17A | 0.229 | 0.113 | 0.967 | 0.48 | 0.736 | 6.1E-18 |
| 4 | AKAP8 | 0.104 | 0.067 | 0.424 | 0.344 | 0.302 | 0.00026 |
| 4 | AKAP8L | 0.177 | 0.134 | 0.782 | 0.684 | 0.193 | 0.00721 |
| 4 | AKIRIN1 | 0.23 | 0.143 | 0.964 | 0.738 | 0.386 | 1.4E-08 |
| 4 | AKR1A1 | 0.13 | 0.069 | 0.446 | 0.327 | 0.449 | 8.7E-09 |
| 4 | AL049869.3 | 0.11 | 0.068 | 0.443 | 0.343 | 0.368 | 4.9E-05 |
| 4 | AL118416.1 | 0.119 | 0.084 | 0.443 | 0.444 | 0.289 | 0.00333 |
| 4 | AL136444.1 | 0.409 | 0.238 | 2.047 | 1.294 | 0.662 | 1.1E-20 |
| 4 | AL136446.1 | 0.284 | 0.02 | 2.74 | 0.123 | 4.478 | 0 |
| 4 | AL138963.4 | 0.337 | 0.244 | 1.473 | 1.418 | 0.14 | 1.6E-05 |
| 4 | AL390728.4 | 0.234 | 0.141 | 1.018 | 0.718 | 0.404 | 4.3E-10 |
| 4 | AL691403.1 | 0.167 | 0.093 | 0.996 | 0.444 | 0.844 | 1E-10 |
| 4 | ALG13 | 0.233 | 0.172 | 1.044 | 0.94 | 0.138 | 0.00048 |
| 4 | ALKBH7 | 0.139 | 0.087 | 0.447 | 0.421 | 0.404 | 8.3E-06 |
| 4 | AMBRA1 | 0.784 | 0.432 | 6.606 | 3.99 | 0.727 | 4.7E-43 |
| 4 | AMD1 | 0.201 | 0.142 | 0.929 | 0.842 | 0.124 | 0.00233 |
| 4 | ANAPC16 | 0.407 | 0.284 | 1.946 | 1.613 | 0.278 | 2.3E-08 |
| 4 | ANK3 | 0.14 | 0.039 | 0.866 | 0.224 | 1.948 | 1.7E-38 |
| 4 | ANKLE2 | 0.246 | 0.149 | 1.043 | 0.828 | 0.334 | 1.3E-07 |
| 4 | ANKRD13A | 0.137 | 0.097 | 0.414 | 0.473 | 0.123 | 0.00215 |
| 4 | ANP32A | 0.323 | 0.249 | 1.404 | 1.372 | 0.133 | 0.00027 |
| 4 | ANP32B | 0.433 | 0.299 | 2.174 | 1.737 | 0.324 | 1.2E-09 |
| 4 | ANTKMT | 0.126 | 0.066 | 0.444 | 0.316 | 0.423 | 6.6E-09 |
| 4 | ANTXR2 | 0.113 | 0.028 | 0.404 | 0.131 | 1.946 | 4E-36 |
| 4 | ANXA4 | 0.16 | 0.071 | 0.744 | 0.382 | 0.96 | 1.7E-17 |
| 4 | ANXA6 | 0.241 | 0.177 | 1.084 | 0.98 | 0.144 | 1.8E-05 |
| 4 | ANXA7 | 0.269 | 0.14 | 1.113 | 0.716 | 0.636 | 2.9E-18 |
| 4 | AP1S2 | 0.166 | 0.111 | 0.641 | 0.478 | 0.172 | 6.9E-05 |
| 4 | APEX1 | 0.166 | 0.086 | 0.614 | 0.423 | 0.441 | 7.5E-12 |
| 4 | API4 | 0.231 | 0.178 | 1.004 | 0.927 | 0.114 | 0.0032 |
| 4 | APIP | 0.101 | 0.064 | 0.376 | 0.314 | 0.262 | 0.00038 |
| 4 | APOO | 0.267 | 0.124 | 1.103 | 0.626 | 0.818 | 3.9E-24 |
| 4 | APRT | 0.343 | 0.217 | 1.492 | 1.136 | 0.487 | 9.9E-14 |
| 4 | ARCN1 | 0.237 | 0.173 | 0.976 | 0.894 | 0.124 | 0.00032 |

| | | | | | | | |
|---|---|---|---|---|---|---|---|
| 4 | AREG | 0.344 | 0.086 | 3.436 | 0.474 | 2.482 | 3E-128 |
| 4 | ARF4 | 0.33 | 0.22 | 1.724 | 1.284 | 0.426 | 1.5E-09 |
| 4 | ARF6 | 0.339 | 0.231 | 1.442 | 1.243 | 0.32 | 2.9E-08 |
| 4 | ARFIP1 | 0.191 | 0.124 | 0.746 | 0.641 | 0.216 | 4.4E-06 |
| 4 | ARHGAP12 | 0.209 | 0.114 | 0.878 | 0.617 | 0.409 | 4.3E-12 |
| 4 | ARHGAP14 | 0.781 | 0.646 | 7.348 | 6.096 | 0.271 | 7.5E-09 |
| 4 | ARHGAP4 | 0.2 | 0.132 | 0.946 | 0.73 | 0.374 | 3.1E-06 |
| 4 | ARHGDIA | 0.224 | 0.144 | 0.94 | 0.803 | 0.243 | 1.7E-05 |
| 4 | ARHGEF6 | 0.297 | 0.199 | 1.368 | 1.07 | 0.344 | 1.1E-07 |
| 4 | ARL1 | 0.101 | 0.064 | 0.386 | 0.321 | 0.267 | 0.00028 |
| 4 | ARL14EP | 0.137 | 0.064 | 0.479 | 0.324 | 0.462 | 2.5E-12 |
| 4 | ARL16 | 0.116 | 0.08 | 0.463 | 0.394 | 0.229 | 0.00175 |
| 4 | ARL2BP | 0.211 | 0.143 | 0.864 | 0.786 | 0.137 | 0.00049 |
| 4 | ARL4A | 0.124 | 0.042 | 0.463 | 0.267 | 1.076 | 6.6E-16 |
| 4 | ARMC9 | 0.133 | 0.06 | 0.437 | 0.284 | 0.913 | 7E-14 |
| 4 | ARRDC2 | 0.119 | 0.068 | 0.494 | 0.342 | 0.433 | 1E-06 |
| 4 | ASL | 0.106 | 0.049 | 0.407 | 0.301 | 0.432 | 8.1E-07 |
| 4 | ASTE1 | 0.149 | 0.091 | 0.622 | 0.462 | 0.43 | 1.7E-06 |
| 4 | ATF4 | 0.189 | 0.104 | 0.764 | 0.432 | 0.424 | 4.3E-11 |
| 4 | ATIC | 0.119 | 0.048 | 0.424 | 0.273 | 0.634 | 4E-10 |
| 4 | ATP4F1D | 0.447 | 0.341 | 2.177 | 1.974 | 0.142 | 1.7E-05 |
| 4 | ATP4F1E | 0.806 | 0.681 | 7.016 | 6.126 | 0.196 | 4.9E-07 |
| 4 | ATP4MC1 | 0.173 | 0.119 | 0.663 | 0.478 | 0.197 | 0.00018 |
| 4 | ATP4MC2 | 0.701 | 0.484 | 4.914 | 4.447 | 0.141 | 4.9E-05 |
| 4 | ATP4MG | 0.626 | 0.436 | 3.692 | 2.791 | 0.403 | 7.2E-16 |
| 4 | ATP4PB | 0.304 | 0.217 | 1.267 | 1.142 | 0.149 | 1.9E-05 |
| 4 | ATP4PD | 0.249 | 0.184 | 1.041 | 0.97 | 0.102 | 0.00065 |
| 4 | ATP4PF | 0.413 | 0.324 | 2.026 | 1.838 | 0.14 | 0.00035 |
| 4 | ATP4PO | 0.317 | 0.213 | 1.388 | 1.106 | 0.327 | 2.1E-08 |
| 4 | ATP6V0C | 0.47 | 0.264 | 2.326 | 1.473 | 0.66 | 1.5E-26 |
| 4 | ATP6V1G1 | 0.441 | 0.341 | 2.184 | 2.029 | 0.106 | 0.00029 |
| 4 | ATP8B4 | 0.396 | 0.11 | 2.226 | 0.648 | 1.78 | 2E-110 |
| 4 | ATXN7L3B | 0.424 | 0.318 | 2.131 | 1.903 | 0.163 | 2E-05 |
| 4 | B3GNT2 | 0.373 | 0.247 | 1.86 | 1.384 | 0.426 | 4.3E-11 |
| 4 | B4GALT4 | 0.144 | 0.094 | 0.602 | 0.499 | 0.271 | 1.4E-06 |
| 4 | BACH2 | 0.401 | 0.146 | 3.24 | 0.947 | 1.78 | 3E-127 |
| 4 | BAIAP2L1 | 0.123 | 0.041 | 0.444 | 0.244 | 0.894 | 2.7E-14 |
| 4 | BANF1 | 0.226 | 0.171 | 1.006 | 0.882 | 0.189 | 0.00137 |
| 4 | BAZ2B | 0.33 | 0.144 | 1.701 | 0.889 | 0.936 | 2.7E-32 |
| 4 | BBIP1 | 0.11 | 0.079 | 0.434 | 0.4 | 0.118 | 0.00672 |
| 4 | BCAP29 | 0.141 | 0.108 | 0.609 | 0.44 | 0.173 | 0.0014 |
| 4 | BCCIP | 0.166 | 0.116 | 0.664 | 0.474 | 0.208 | 0.0003 |
| 4 | BCL10 | 0.244 | 0.181 | 1.046 | 0.961 | 0.123 | 6E-05 |
| 4 | BCL2 | 0.194 | 0.093 | 1.086 | 0.434 | 1.024 | 3.5E-18 |
| 4 | BCL3 | 0.116 | 0.041 | 0.444 | 0.187 | 1.246 | 1.5E-20 |
| 4 | BCL6 | 0.183 | 0.108 | 0.761 | 0.476 | 0.402 | 8.4E-09 |
| 4 | BCL7B | 0.124 | 0.088 | 0.404 | 0.442 | 0.189 | 0.00309 |
| 4 | BCLAF1 | 0.499 | 0.433 | 3.666 | 2.984 | 0.297 | 2.6E-11 |
| 4 | BEX2 | 0.126 | 0.041 | 0.417 | 0.198 | 1.381 | 2.3E-24 |
| 4 | BEX4 | 0.144 | 0.073 | 0.443 | 0.341 | 0.644 | 2.5E-11 |
| 4 | BHLHE40 | 0.437 | 0.312 | 3.906 | 1.986 | 0.976 | 3.1E-37 |
| 4 | BHLHE40-AS1 | 0.119 | 0.084 | 0.431 | 0.442 | 0.264 | 0.00498 |
| 4 | BICRAL | 0.189 | 0.143 | 0.922 | 0.778 | 0.244 | 0.00249 |
| 4 | BID | 0.147 | 0.087 | 0.674 | 0.431 | 0.644 | 1.6E-09 |

| | | | | | | | |
|---|---|---|---|---|---|---|---|
| 4 | BIRC3 | 0.249 | 0.168 | 1.894 | 1.028 | 0.882 | 6.5E-09 |
| 4 | BMI1 | 0.18 | 0.131 | 0.728 | 0.677 | 0.104 | 0.00142 |
| 4 | BMS1 | 0.146 | 0.084 | 0.497 | 0.434 | 0.461 | 1.4E-07 |
| 4 | BORCS7 | 0.121 | 0.089 | 0.416 | 0.442 | 0.191 | 0.00609 |
| 4 | BRCC3 | 0.134 | 0.073 | 0.416 | 0.34 | 0.461 | 9.2E-09 |
| 4 | BRD2 | 0.336 | 0.234 | 1.489 | 1.324 | 0.169 | 3.4E-06 |
| 4 | BRIX1 | 0.144 | 0.089 | 0.437 | 0.443 | 0.277 | 6.7E-06 |
| 4 | BRMS1 | 0.104 | 0.068 | 0.398 | 0.333 | 0.246 | 0.00042 |
| 4 | BST2 | 0.416 | 0.428 | 3.631 | 3.272 | 0.14 | 0.0005 |
| 4 | BTF3 | 0.749 | 0.427 | 4.77 | 3.834 | 0.489 | 1.4E-27 |
| 4 | BTF3L4 | 0.133 | 0.074 | 0.423 | 0.361 | 0.434 | 7.3E-08 |
| 4 | BTG1 | 0.924 | 0.824 | 13.706 | 11.04 | 0.311 | 7.9E-14 |
| 4 | BTG2 | 0.833 | 0.694 | 12.94 | 9.241 | 0.484 | 7.2E-20 |
| 4 | BUD23 | 0.114 | 0.082 | 0.483 | 0.397 | 0.282 | 0.00431 |
| 4 | BZW1 | 0.721 | 0.61 | 4.466 | 4.044 | 0.113 | 0.00294 |
| 4 | BZW2 | 0.113 | 0.048 | 0.443 | 0.249 | 0.834 | 1.1E-13 |
| 4 | C11orf64 | 0.116 | 0.079 | 0.429 | 0.394 | 0.12 | 0.00121 |
| 4 | C12orf47 | 0.389 | 0.262 | 2.031 | 1.732 | 0.23 | 2.9E-09 |
| 4 | C14orf61 | 0.11 | 0.074 | 0.426 | 0.362 | 0.237 | 0.001 |
| 4 | C16orf44 | 0.131 | 0.049 | 0.447 | 0.292 | 0.906 | 3.1E-14 |
| 4 | C1orf112 | 0.103 | 0.06 | 0.379 | 0.299 | 0.342 | 1.1E-05 |
| 4 | C1orf162 | 0.294 | 0.144 | 1.348 | 0.888 | 0.613 | 1.7E-19 |
| 4 | C1QBP | 0.244 | 0.129 | 0.991 | 0.637 | 0.636 | 4.5E-16 |
| 4 | C9orf16 | 0.377 | 0.287 | 1.702 | 1.474 | 0.112 | 0.00024 |
| 4 | CA4B | 0.261 | 0.142 | 1.274 | 0.836 | 0.607 | 7.4E-14 |
| 4 | CALCOCO2 | 0.277 | 0.192 | 1.207 | 1.016 | 0.248 | 3.1E-06 |
| 4 | CALM2 | 0.481 | 0.364 | 2.649 | 2.187 | 0.282 | 3.3E-07 |
| 4 | CANX | 0.441 | 0.361 | 2.494 | 2.242 | 0.147 | 0.00034 |
| 4 | CAPG | 0.283 | 0.004 | 1.349 | 0.029 | 4.441 | 0 |
| 4 | CAPN12 | 0.14 | 0.034 | 0.481 | 0.172 | 1.742 | 4.4E-44 |
| 4 | CAPRIN2 | 0.11 | 0.071 | 0.431 | 0.348 | 0.266 | 0.00034 |
| 4 | CARD19 | 0.146 | 0.076 | 0.707 | 0.383 | 0.884 | 1.5E-13 |
| 4 | CARS | 0.124 | 0.077 | 0.481 | 0.388 | 0.309 | 2.4E-05 |
| 4 | CASD1 | 0.149 | 0.117 | 0.687 | 0.61 | 0.171 | 0.00344 |
| 4 | CASK | 0.326 | 0.09 | 1.64 | 0.491 | 1.741 | 9E-90 |
| 4 | CASP8AP2 | 0.18 | 0.123 | 0.8 | 0.646 | 0.308 | 4.5E-05 |
| 4 | CBLL1 | 0.224 | 0.147 | 0.972 | 0.781 | 0.316 | 8.3E-07 |
| 4 | CBWD2 | 0.111 | 0.074 | 0.432 | 0.378 | 0.192 | 0.00064 |
| 4 | CBX3 | 0.464 | 0.324 | 2.407 | 1.927 | 0.321 | 2.9E-10 |
| 4 | CBX4 | 0.197 | 0.139 | 0.827 | 0.739 | 0.161 | 0.00018 |
| 4 | CCDC173 | 0.121 | 0.062 | 0.464 | 0.339 | 0.734 | 1.6E-09 |
| 4 | CCDC200 | 0.243 | 0.104 | 1.847 | 0.883 | 1.073 | 1.4E-28 |
| 4 | CCDC40 | 0.129 | 0.064 | 0.446 | 0.332 | 0.718 | 2.3E-10 |
| 4 | CCDC49 | 0.181 | 0.126 | 0.747 | 0.663 | 0.192 | 0.00014 |
| 4 | CCDC84B | 0.306 | 0.226 | 1.3 | 1.204 | 0.111 | 0.0003 |
| 4 | CCND2 | 0.374 | 0.301 | 2.91 | 1.873 | 0.634 | 4.5E-05 |
| 4 | CCNG2 | 0.173 | 0.092 | 0.736 | 0.46 | 0.678 | 1.3E-11 |
| 4 | CCNI | 0.476 | 0.374 | 2.488 | 2.34 | 0.144 | 0.00068 |
| 4 | CCNL1 | 0.489 | 0.367 | 3.363 | 2.46 | 0.394 | 1.4E-09 |
| 4 | CCR7 | 0.136 | 0.012 | 0.482 | 0.062 | 3.224 | 2E-134 |
| 4 | CCS | 0.104 | 0.074 | 0.404 | 0.364 | 0.141 | 0.00884 |
| 4 | CCSER1 | 0.139 | 0.078 | 0.426 | 0.387 | 0.444 | 6.4E-08 |
| 4 | CCT2 | 0.336 | 0.214 | 1.478 | 1.144 | 0.346 | 2.1E-10 |
| 4 | CCT4 | 0.33 | 0.217 | 1.392 | 1.163 | 0.249 | 9.8E-09 |

| | | | | | | | |
|---|---|---|---|---|---|---|---|
| 4 | CCT4 | 0.219 | 0.138 | 0.93 | 0.681 | 0.448 | 4.7E-08 |
| 4 | CCT6A | 0.324 | 0.214 | 1.467 | 1.141 | 0.363 | 4.7E-09 |
| 4 | CCT7 | 0.2 | 0.139 | 0.844 | 0.691 | 0.307 | 2.2E-05 |
| 4 | CCT8 | 0.263 | 0.188 | 1.074 | 0.989 | 0.119 | 0.00012 |
| 4 | CD2 | 0.476 | 0.188 | 3.422 | 1.213 | 1.497 | 1.2E-77 |
| 4 | CD2AP | 0.344 | 0.174 | 1.682 | 0.974 | 0.786 | 3E-26 |
| 4 | CD300A | 0.264 | 0.184 | 1.248 | 1.019 | 0.292 | 3.7E-06 |
| 4 | CD37 | 0.409 | 0.394 | 2.927 | 2.464 | 0.248 | 2.7E-06 |
| 4 | CD3E | 0.333 | 0.176 | 2.06 | 1.32 | 0.642 | 2.8E-22 |
| 4 | CD42 | 0.649 | 0.327 | 4.34 | 2.984 | 0.842 | 6.1E-60 |
| 4 | CD44 | 0.823 | 0.431 | 9.314 | 3.262 | 1.414 | 2E-143 |
| 4 | CD44 | 0.374 | 0.181 | 2.098 | 1.084 | 0.941 | 1.2E-34 |
| 4 | CD63 | 0.447 | 0.284 | 2.416 | 1.646 | 0.444 | 4.4E-18 |
| 4 | CD69 | 0.671 | 0.424 | 8.448 | 6.726 | 0.347 | 2.6E-11 |
| 4 | CD7 | 0.647 | 0.494 | 4.028 | 3.766 | 0.417 | 3.8E-12 |
| 4 | CD74 | 0.801 | 0.704 | 9.344 | 8.301 | 0.171 | 9.5E-05 |
| 4 | CD82 | 0.181 | 0.093 | 0.891 | 0.421 | 0.773 | 8.8E-14 |
| 4 | CD83 | 0.231 | 0.121 | 1.499 | 0.714 | 1.07 | 9.2E-17 |
| 4 | CDC14A | 0.293 | 0.194 | 1.641 | 1.123 | 0.447 | 4.7E-09 |
| 4 | CDC42 | 0.874 | 0.741 | 9.638 | 8.296 | 0.216 | 7.6E-08 |
| 4 | CDK6 | 0.224 | 0.119 | 1.169 | 0.668 | 0.808 | 1.2E-14 |
| 4 | CDK9 | 0.101 | 0.062 | 0.421 | 0.308 | 0.44 | 6.7E-05 |
| 4 | CDKN2D | 0.296 | 0.196 | 1.333 | 1.094 | 0.283 | 8.3E-08 |
| 4 | CENPJ | 0.107 | 0.048 | 0.44 | 0.241 | 0.871 | 4.4E-12 |
| 4 | CEP83 | 0.136 | 0.044 | 0.614 | 0.279 | 1.14 | 2.9E-19 |
| 4 | CERS6 | 0.209 | 0.104 | 0.888 | 0.477 | 0.622 | 3E-16 |
| 4 | CFAP298 | 0.141 | 0.097 | 0.434 | 0.49 | 0.124 | 0.00078 |
| 4 | CFL2 | 0.121 | 0.061 | 0.462 | 0.317 | 0.444 | 7.7E-10 |
| 4 | CGGBP1 | 0.473 | 0.347 | 2.412 | 2.219 | 0.121 | 3.8E-05 |
| 4 | CGRRF1 | 0.113 | 0.07 | 0.462 | 0.341 | 0.439 | 4.1E-05 |
| 4 | CHCHD7 | 0.101 | 0.069 | 0.361 | 0.336 | 0.103 | 0.00262 |
| 4 | CHD2 | 0.624 | 0.499 | 4.268 | 3.742 | 0.19 | 4.7E-06 |
| 4 | CHD9 | 0.443 | 0.24 | 2.344 | 1.442 | 0.707 | 1.6E-24 |
| 4 | CHKA | 0.116 | 0.029 | 0.419 | 0.163 | 1.671 | 4.1E-36 |
| 4 | CHM | 0.123 | 0.082 | 0.447 | 0.41 | 0.146 | 0.0004 |
| 4 | CHMP2B | 0.146 | 0.103 | 0.49 | 0.417 | 0.189 | 7.2E-05 |
| 4 | CHMP3 | 0.13 | 0.094 | 0.474 | 0.473 | 0.282 | 0.00319 |
| 4 | CHMP4B | 0.144 | 0.108 | 0.667 | 0.434 | 0.319 | 0.0004 |
| 4 | CHRAC1 | 0.176 | 0.136 | 0.81 | 0.691 | 0.23 | 0.00647 |
| 4 | CHURC1 | 0.244 | 0.193 | 1.207 | 1.022 | 0.239 | 0.00067 |
| 4 | CISD2 | 0.197 | 0.134 | 0.811 | 0.704 | 0.202 | 4.5E-05 |
| 4 | CKLF | 0.407 | 0.309 | 3.72 | 1.861 | 0.999 | 2.8E-30 |
| 4 | CLCN3 | 0.181 | 0.126 | 0.764 | 0.641 | 0.233 | 0.00014 |
| 4 | CLDND1 | 0.441 | 0.274 | 2.893 | 1.803 | 0.682 | 4E-20 |
| 4 | CLIC1 | 0.744 | 0.614 | 4.907 | 4.028 | 0.232 | 4.3E-07 |
| 4 | CLNS1A | 0.249 | 0.148 | 1.023 | 0.81 | 0.336 | 1.7E-09 |
| 4 | CLPP | 0.123 | 0.066 | 0.433 | 0.319 | 0.443 | 4.7E-08 |
| 4 | CLSTN3 | 0.13 | 0.074 | 0.482 | 0.367 | 0.394 | 2E-07 |
| 4 | CMC1 | 0.417 | 0.278 | 2.466 | 1.704 | 0.49 | 1.9E-13 |
| 4 | CMC2 | 0.219 | 0.146 | 0.894 | 0.824 | 0.118 | 0.00017 |
| 4 | CMPK1 | 0.4 | 0.307 | 1.944 | 1.79 | 0.119 | 0.00043 |
| 4 | CMSS1 | 0.114 | 0.047 | 0.443 | 0.306 | 0.432 | 1.7E-09 |
| 4 | CMTM3 | 0.106 | 0.076 | 0.43 | 0.37 | 0.218 | 0.0085 |
| 4 | CMTM6 | 0.401 | 0.237 | 1.964 | 1.264 | 0.634 | 1.7E-19 |

| | | | | | | | |
|---|---|---|---|---|---|---|---|
| 4 | CNBP | 0.447 | 0.414 | 3.024 | 2.477 | 0.232 | 1.3E-06 |
| 4 | CNDP2 | 0.14 | 0.093 | 0.419 | 0.474 | 0.129 | 1E-05 |
| 4 | CNIH1 | 0.143 | 0.093 | 0.428 | 0.461 | 0.198 | 7.9E-05 |
| 4 | CNN2 | 0.276 | 0.114 | 1.244 | 0.602 | 1.049 | 2.7E-34 |
| 4 | CNOT9 | 0.141 | 0.114 | 0.642 | 0.489 | 0.124 | 0.00945 |
| 4 | CNR2 | 0.116 | 0.007 | 0.494 | 0.036 | 3.788 | 7E-146 |
| 4 | CNST | 0.269 | 0.188 | 1.204 | 1.014 | 0.246 | 4.8E-06 |
| 4 | COMMD10 | 0.249 | 0.17 | 1.089 | 0.98 | 0.143 | 4.5E-07 |
| 4 | COMMD2 | 0.161 | 0.114 | 0.642 | 0.469 | 0.196 | 0.00092 |
| 4 | COMMD3 | 0.114 | 0.079 | 0.462 | 0.383 | 0.272 | 0.00164 |
| 4 | COMMD6 | 0.697 | 0.414 | 4.891 | 3.679 | 0.411 | 1.7E-16 |
| 4 | COMMD7 | 0.241 | 0.178 | 1.061 | 0.919 | 0.208 | 3.2E-05 |
| 4 | COPB2 | 0.234 | 0.166 | 0.947 | 0.868 | 0.14 | 0.0001 |
| 4 | COPS2 | 0.223 | 0.149 | 0.971 | 0.838 | 0.213 | 0.00011 |
| 4 | COTL1 | 0.417 | 0.03 | 2.843 | 0.161 | 4.138 | 0 |
| 4 | COX17 | 0.276 | 0.219 | 1.314 | 1.166 | 0.173 | 0.00664 |
| 4 | COX20 | 0.207 | 0.133 | 0.914 | 0.69 | 0.408 | 3.5E-07 |
| 4 | COX4A | 0.366 | 0.24 | 1.484 | 1.329 | 0.243 | 4.9E-08 |
| 4 | COX4I1 | 0.726 | 0.638 | 4.788 | 4.291 | 0.13 | 0.00363 |
| 4 | COX7A2 | 0.403 | 0.379 | 2.447 | 2.231 | 0.191 | 4.1E-06 |
| 4 | COX7A2L | 0.239 | 0.162 | 0.94 | 0.814 | 0.221 | 6.8E-06 |
| 4 | COX7B | 0.334 | 0.224 | 1.406 | 1.144 | 0.284 | 3.1E-08 |
| 4 | COX7C | 0.709 | 0.444 | 4.239 | 4.109 | 0.341 | 2.1E-12 |
| 4 | CPB2-AS1 | 0.337 | 0.261 | 1.602 | 1.469 | 0.124 | 0.00082 |
| 4 | CPNE1 | 0.183 | 0.134 | 0.748 | 0.694 | 0.127 | 0.00304 |
| 4 | CRACR2A | 0.193 | 0.124 | 0.766 | 0.688 | 0.144 | 3.4E-06 |
| 4 | CRELD2 | 0.107 | 0.049 | 0.417 | 0.238 | 0.808 | 3.8E-11 |
| 4 | CRIP1 | 0.48 | 0.443 | 6.08 | 3.403 | 0.796 | 1.4E-14 |
| 4 | CROCC | 0.31 | 0.206 | 2.146 | 1.442 | 0.477 | 6.3E-10 |
| 4 | CROT | 0.123 | 0.079 | 0.432 | 0.393 | 0.437 | 7.4E-05 |
| 4 | CRTAM | 0.104 | 0.013 | 0.416 | 0.068 | 2.922 | 2.6E-73 |
| 4 | CRYBG1 | 0.149 | 0.064 | 0.644 | 0.343 | 0.89 | 3.6E-21 |
| 4 | CSDE1 | 0.497 | 0.38 | 2.648 | 2.378 | 0.144 | 1.1E-05 |
| 4 | CSGALNACT1 | 0.109 | 0.04 | 0.436 | 0.234 | 0.889 | 1.9E-17 |
| 4 | CSNK1G3 | 0.424 | 0.274 | 2.047 | 1.611 | 0.342 | 4.3E-12 |
| 4 | CSPP1 | 0.144 | 0.102 | 0.668 | 0.447 | 0.288 | 6.8E-05 |
| 4 | CSRNP1 | 0.189 | 0.082 | 0.917 | 0.428 | 1.1 | 3.4E-22 |
| 4 | CST3 | 0.104 | 0.028 | 0.476 | 0.137 | 1.794 | 4.4E-29 |
| 4 | CTNNB1 | 0.32 | 0.234 | 1.446 | 1.33 | 0.227 | 3.1E-05 |
| 4 | CTNNBIP1 | 0.106 | 0.044 | 0.409 | 0.27 | 0.6 | 4.2E-08 |
| 4 | CTSW | 0.884 | 0.706 | 14.489 | 9.102 | 0.671 | 7.5E-48 |
| 4 | CUTA | 0.343 | 0.224 | 1.69 | 1.227 | 0.462 | 7.7E-12 |
| 4 | CUX1 | 0.287 | 0.174 | 1.212 | 0.988 | 0.294 | 4.6E-11 |
| 4 | CWC22 | 0.167 | 0.122 | 0.704 | 0.634 | 0.141 | 0.00223 |
| 4 | CXCR3 | 0.239 | 0.1 | 1.202 | 0.464 | 1.09 | 1.9E-29 |
| 4 | CXXC4 | 0.249 | 0.174 | 1.094 | 0.944 | 0.213 | 2.6E-06 |
| 4 | CYP41A1 | 0.147 | 0.112 | 0.634 | 0.491 | 0.104 | 0.00141 |
| 4 | CYSLTR1 | 0.197 | 0.037 | 1.048 | 0.182 | 2.427 | 2E-91 |
| 4 | CYTH3 | 0.173 | 0.102 | 0.716 | 0.446 | 0.391 | 3.6E-08 |
| 4 | DAB1 | 0.147 | 0.108 | 0.647 | 0.448 | 0.237 | 0.0002 |
| 4 | DAD1 | 0.327 | 0.242 | 1.444 | 1.288 | 0.164 | 8.1E-05 |
| 4 | DAP | 0.104 | 0.073 | 0.394 | 0.363 | 0.121 | 0.00479 |
| 4 | DAP3 | 0.171 | 0.118 | 0.649 | 0.497 | 0.143 | 0.00024 |
| 4 | DARS | 0.244 | 0.124 | 1.039 | 0.634 | 0.71 | 1.1E-18 |

| | | | | | | | |
|---|---|---|---|---|---|---|---|
| 4 | DAZAP2 | 0.447 | 0.417 | 3.022 | 2.703 | 0.161 | 4.9E-06 |
| 4 | DBNL | 0.146 | 0.101 | 0.617 | 0.404 | 0.29 | 1.3E-05 |
| 4 | DCUN1D1 | 0.17 | 0.113 | 0.671 | 0.486 | 0.194 | 4.6E-05 |
| 4 | DDIT3 | 0.131 | 0.091 | 0.443 | 0.464 | 0.227 | 0.00089 |
| 4 | DDIT4 | 0.37 | 0.224 | 2.636 | 1.407 | 0.906 | 3.2E-17 |
| 4 | DDT | 0.234 | 0.141 | 0.961 | 0.764 | 0.329 | 1E-07 |
| 4 | DDX18 | 0.323 | 0.217 | 1.434 | 1.188 | 0.271 | 6E-08 |
| 4 | DDX19A | 0.124 | 0.073 | 0.463 | 0.363 | 0.341 | 2.3E-06 |
| 4 | DDX21 | 0.373 | 0.236 | 1.949 | 1.382 | 0.403 | 4.9E-13 |
| 4 | DDX3X | 0.617 | 0.47 | 4.148 | 3.344 | 0.31 | 3.3E-10 |
| 4 | DDX4 | 0.744 | 0.622 | 4.808 | 4.334 | 0.123 | 0.00024 |
| 4 | DDX40 | 0.234 | 0.163 | 0.971 | 0.864 | 0.166 | 2.7E-05 |
| 4 | DEGS1 | 0.21 | 0.1 | 0.906 | 0.411 | 0.827 | 8.7E-19 |
| 4 | DELE1 | 0.147 | 0.103 | 0.498 | 0.417 | 0.212 | 4.7E-05 |
| 4 | DENND1B | 0.347 | 0.248 | 1.723 | 1.468 | 0.231 | 6.4E-08 |
| 4 | DENND4A | 0.466 | 0.327 | 2.847 | 2.124 | 0.422 | 1.5E-11 |
| 4 | DESI2 | 0.166 | 0.122 | 0.684 | 0.612 | 0.16 | 0.00225 |
| 4 | DGKA | 0.11 | 0.08 | 0.434 | 0.399 | 0.122 | 0.00839 |
| 4 | DGKE | 0.104 | 0.069 | 0.421 | 0.346 | 0.28 | 0.00047 |
| 4 | DGUOK | 0.227 | 0.147 | 0.943 | 0.821 | 0.214 | 1.9E-05 |
| 4 | DHRSX | 0.149 | 0.103 | 0.621 | 0.444 | 0.188 | 0.00043 |
| 4 | DICER1 | 0.271 | 0.19 | 1.19 | 1.071 | 0.141 | 1.7E-05 |
| 4 | DIMT1 | 0.104 | 0.073 | 0.397 | 0.364 | 0.122 | 0.00398 |
| 4 | DIPK2A | 0.133 | 0.099 | 0.473 | 0.417 | 0.14 | 0.0089 |
| 4 | DLST | 0.137 | 0.1 | 0.448 | 0.41 | 0.131 | 0.00383 |
| 4 | DMAC1 | 0.12 | 0.048 | 0.416 | 0.284 | 0.848 | 7.8E-11 |
| 4 | DNAAF2 | 0.113 | 0.043 | 0.411 | 0.242 | 0.704 | 4.7E-11 |
| 4 | DNAJA1 | 0.494 | 0.402 | 4.099 | 2.743 | 0.474 | 1.1E-21 |
| 4 | DNAJC10 | 0.247 | 0.181 | 1.189 | 0.966 | 0.299 | 1E-05 |
| 4 | DNAJC19 | 0.184 | 0.133 | 0.737 | 0.684 | 0.104 | 0.00109 |
| 4 | DNAJC4 | 0.104 | 0.072 | 0.399 | 0.344 | 0.212 | 0.0025 |
| 4 | DNAJC9 | 0.133 | 0.092 | 0.469 | 0.449 | 0.309 | 0.00067 |
| 4 | DNMT3A | 0.116 | 0.047 | 0.47 | 0.233 | 1.011 | 8.1E-16 |
| 4 | DOCK10 | 0.669 | 0.444 | 4.834 | 4.363 | 0.148 | 1E-05 |
| 4 | DPH3 | 0.187 | 0.119 | 0.829 | 0.496 | 0.474 | 4.3E-07 |
| 4 | DPM3 | 0.183 | 0.111 | 0.702 | 0.447 | 0.349 | 1.1E-07 |
| 4 | DPYSL2 | 0.109 | 0.064 | 0.412 | 0.312 | 0.403 | 9.4E-06 |
| 4 | DRAM2 | 0.109 | 0.078 | 0.43 | 0.397 | 0.113 | 0.00879 |
| 4 | DSE | 0.119 | 0.064 | 0.474 | 0.324 | 0.441 | 3.8E-08 |
| 4 | DST | 0.127 | 0.017 | 0.644 | 0.091 | 2.843 | 2.2E-84 |
| 4 | DUSP1 | 0.427 | 0.279 | 4.039 | 2.468 | 0.643 | 2.8E-17 |
| 4 | DUSP2 | 0.61 | 0.374 | 9.323 | 3.363 | 1.471 | 9.3E-44 |
| 4 | DUT | 0.217 | 0.144 | 0.978 | 0.81 | 0.272 | 4.4E-05 |
| 4 | DYNLT3 | 0.204 | 0.129 | 0.903 | 0.643 | 0.468 | 9.3E-08 |
| 4 | DYRK2 | 0.119 | 0.074 | 0.426 | 0.373 | 0.191 | 9.1E-05 |
| 4 | DZIP3 | 0.12 | 0.041 | 0.464 | 0.209 | 1.149 | 3.8E-22 |
| 4 | E2F3 | 0.193 | 0.128 | 0.839 | 0.696 | 0.27 | 4.9E-06 |
| 4 | ECE1 | 0.124 | 0.079 | 0.424 | 0.421 | 0.317 | 4.4E-05 |
| 4 | ECHS1 | 0.144 | 0.084 | 0.443 | 0.404 | 0.442 | 2.1E-09 |
| 4 | ECPAS | 0.203 | 0.147 | 0.924 | 0.838 | 0.142 | 0.00494 |
| 4 | EDEM3 | 0.119 | 0.082 | 0.482 | 0.428 | 0.173 | 0.00199 |
| 4 | EDF1 | 0.401 | 0.307 | 1.942 | 1.732 | 0.173 | 0.0001 |
| 4 | EDRF1 | 0.176 | 0.134 | 0.747 | 0.691 | 0.112 | 0.0065 |
| 4 | EEA1 | 0.21 | 0.146 | 0.974 | 0.873 | 0.16 | 0.00088 |

| | | | | | | | |
|---|---|---|---|---|---|---|---|
| 4 | EEF1A1 | 1 | 0.948 | 47.34 | 26.188 | 0.844 | 4E-174 |
| 4 | EEF1B2 | 0.767 | 0.49 | 6.44 | 3.486 | 0.869 | 1.9E-46 |
| 4 | EEF1D | 0.76 | 0.604 | 4.948 | 4.836 | 0.301 | 9.3E-11 |
| 4 | EEF1G | 0.47 | 0.338 | 3.201 | 2.003 | 0.677 | 1.3E-31 |
| 4 | EEF2 | 0.381 | 0.247 | 1.827 | 1.44 | 0.344 | 7.5E-10 |
| 4 | EIF1 | 0.91 | 0.806 | 17.363 | 12.239 | 0.404 | 3E-20 |
| 4 | EIF1AX | 0.34 | 0.224 | 1.699 | 1.269 | 0.422 | 3.5E-11 |
| 4 | EIF2A | 0.241 | 0.146 | 1.061 | 0.761 | 0.479 | 4E-12 |
| 4 | EIF2D | 0.131 | 0.09 | 0.42 | 0.462 | 0.171 | 0.00078 |
| 4 | EIF2S1 | 0.163 | 0.119 | 0.642 | 0.494 | 0.133 | 0.00226 |
| 4 | EIF2S2 | 0.28 | 0.213 | 1.314 | 1.176 | 0.161 | 0.00064 |
| 4 | EIF2S3 | 0.241 | 0.166 | 0.944 | 0.884 | 0.11 | 2.5E-05 |
| 4 | EIF3D | 0.189 | 0.114 | 0.737 | 0.462 | 0.39 | 4.3E-08 |
| 4 | EIF3E | 0.284 | 0.168 | 1.168 | 0.881 | 0.406 | 3E-12 |
| 4 | EIF3F | 0.289 | 0.197 | 1.184 | 1.034 | 0.194 | 1.8E-06 |
| 4 | EIF3H | 0.449 | 0.324 | 2.464 | 1.894 | 0.38 | 4.2E-11 |
| 4 | EIF3I | 0.201 | 0.134 | 0.798 | 0.644 | 0.287 | 1.5E-05 |
| 4 | EIF3J | 0.32 | 0.217 | 1.428 | 1.204 | 0.246 | 1.4E-07 |
| 4 | EIF3K | 0.447 | 0.429 | 3.047 | 2.716 | 0.164 | 4.1E-06 |
| 4 | EIF4 | 0.444 | 0.291 | 2.332 | 1.719 | 0.44 | 1.5E-13 |
| 4 | EIF4A | 0.4 | 0.264 | 2.13 | 1.488 | 0.418 | 1.7E-12 |
| 4 | EIF4A1 | 0.326 | 0.214 | 1.492 | 1.163 | 0.36 | 3.9E-09 |
| 4 | EIF4A2 | 0.311 | 0.218 | 1.284 | 1.197 | 0.103 | 8.9E-06 |
| 4 | EIF4A3 | 0.147 | 0.112 | 0.64 | 0.49 | 0.116 | 0.00122 |
| 4 | EIF4B | 0.429 | 0.364 | 2.806 | 2.297 | 0.289 | 1.7E-11 |
| 4 | EIF4B | 0.289 | 0.199 | 1.263 | 1.124 | 0.167 | 2.1E-06 |
| 4 | EIF4E | 0.124 | 0.09 | 0.477 | 0.432 | 0.142 | 0.00462 |
| 4 | EIF4G1 | 0.214 | 0.144 | 0.898 | 0.747 | 0.266 | 4.8E-06 |
| 4 | EIF4G2 | 0.483 | 0.394 | 3.309 | 2.4 | 0.404 | 6E-14 |
| 4 | EIF6 | 0.177 | 0.133 | 0.719 | 0.64 | 0.147 | 0.00388 |
| 4 | ELAVL1 | 0.141 | 0.099 | 0.639 | 0.49 | 0.384 | 3.8E-05 |
| 4 | ELK3 | 0.231 | 0.16 | 1.009 | 0.884 | 0.188 | 2.8E-05 |
| 4 | ELL2 | 0.213 | 0.141 | 0.974 | 0.901 | 0.113 | 2.7E-06 |
| 4 | ELOA | 0.121 | 0.08 | 0.447 | 0.406 | 0.17 | 0.0003 |
| 4 | ELOVL4 | 0.249 | 0.147 | 1.174 | 0.849 | 0.442 | 6.3E-11 |
| 4 | ELOVL6 | 0.146 | 0.063 | 0.498 | 0.333 | 0.846 | 8.2E-17 |
| 4 | ELP1 | 0.13 | 0.091 | 0.44 | 0.463 | 0.24 | 0.00112 |
| 4 | EMC4 | 0.184 | 0.136 | 0.723 | 0.673 | 0.103 | 0.0019 |
| 4 | EML4 | 0.693 | 0.474 | 4.087 | 3.474 | 0.409 | 1.2E-23 |
| 4 | EMP3 | 0.699 | 0.487 | 4.044 | 3.69 | 0.444 | 1.5E-20 |
| 4 | ENO1 | 0.449 | 0.334 | 2.443 | 1.977 | 0.369 | 7.1E-09 |
| 4 | ENOPH1 | 0.131 | 0.091 | 0.482 | 0.44 | 0.1 | 0.00136 |
| 4 | EPHA4 | 0.224 | 0.104 | 1.04 | 0.462 | 0.887 | 4.4E-22 |
| 4 | EPM2AIP1 | 0.106 | 0.07 | 0.373 | 0.348 | 0.102 | 0.00085 |
| 4 | EPRS | 0.244 | 0.149 | 1.044 | 0.864 | 0.274 | 2E-07 |
| 4 | ERCC1 | 0.173 | 0.091 | 0.814 | 0.48 | 0.762 | 2.1E-12 |
| 4 | ERCC6 | 0.167 | 0.12 | 0.712 | 0.644 | 0.142 | 0.00108 |
| 4 | ERGIC1 | 0.137 | 0.064 | 0.479 | 0.341 | 0.72 | 7.2E-13 |
| 4 | ERGIC3 | 0.191 | 0.142 | 0.798 | 0.714 | 0.148 | 0.00207 |
| 4 | ERMP1 | 0.123 | 0.076 | 0.498 | 0.39 | 0.343 | 2E-05 |
| 4 | ERO1A | 0.12 | 0.073 | 0.464 | 0.367 | 0.342 | 1.6E-05 |
| 4 | ERO1B | 0.266 | 0.178 | 1.221 | 0.976 | 0.324 | 5E-07 |
| 4 | ESD | 0.163 | 0.064 | 0.666 | 0.311 | 1.097 | 4E-23 |
| 4 | ESF1 | 0.127 | 0.071 | 0.469 | 0.364 | 0.363 | 1.9E-07 |

| | | | | | | | |
|---|---|---|---|---|---|---|---|
| 4 | ESYT2 | 0.313 | 0.207 | 1.741 | 1.404 | 0.31 | 3.5E-09 |
| 4 | ETFA | 0.21 | 0.142 | 0.842 | 0.776 | 0.118 | 0.00087 |
| 4 | ETFB | 0.144 | 0.094 | 0.462 | 0.483 | 0.219 | 7.6E-05 |
| 4 | ETV3 | 0.219 | 0.142 | 0.994 | 0.736 | 0.434 | 1.7E-07 |
| 4 | ETV6 | 0.21 | 0.124 | 0.909 | 0.702 | 0.373 | 2.3E-09 |
| 4 | EXOC4 | 0.173 | 0.117 | 0.641 | 0.496 | 0.104 | 0.00011 |
| 4 | EXT1 | 0.129 | 0.068 | 0.498 | 0.396 | 0.33 | 7.8E-09 |
| 4 | EZH2 | 0.164 | 0.102 | 0.681 | 0.492 | 0.202 | 2.5E-06 |
| 4 | FAM102A | 0.123 | 0.067 | 0.498 | 0.341 | 0.447 | 4.6E-08 |
| 4 | FAM117B | 0.116 | 0.062 | 0.472 | 0.336 | 0.491 | 4.4E-08 |
| 4 | FAM177A1 | 0.273 | 0.194 | 1.282 | 1.044 | 0.296 | 4.1E-06 |
| 4 | FAM241A | 0.107 | 0.019 | 0.47 | 0.098 | 2.246 | 1.2E-43 |
| 4 | FAU | 0.987 | 0.927 | 24.13 | 19.131 | 0.393 | 4.3E-34 |
| 4 | FBL | 0.196 | 0.094 | 0.802 | 0.466 | 0.784 | 3.8E-17 |
| 4 | FCER1G | 0.741 | 0.418 | 6.384 | 4.989 | 0.346 | 4.2E-22 |
| 4 | FCF1 | 0.109 | 0.076 | 0.414 | 0.383 | 0.114 | 0.00294 |
| 4 | FCHSD2 | 0.3 | 0.198 | 1.438 | 1.148 | 0.312 | 1.5E-08 |
| 4 | FER | 0.189 | 0.118 | 0.781 | 0.648 | 0.249 | 6.5E-07 |
| 4 | FGFR1OP2 | 0.371 | 0.26 | 1.766 | 1.407 | 0.229 | 2E-07 |
| 4 | FKBP1A | 0.17 | 0.104 | 0.677 | 0.404 | 0.423 | 4.2E-07 |
| 4 | FKBP4 | 0.163 | 0.123 | 0.703 | 0.644 | 0.103 | 0.00485 |
| 4 | FLOT1 | 0.173 | 0.078 | 0.744 | 0.383 | 0.946 | 2.4E-18 |
| 4 | FOS | 0.449 | 0.429 | 47.431 | 14.494 | 1.608 | 1.4E-32 |
| 4 | FOSB | 0.123 | 0.046 | 0.616 | 0.247 | 1.262 | 6.2E-20 |
| 4 | FPGS | 0.117 | 0.061 | 0.424 | 0.308 | 0.464 | 1.9E-08 |
| 4 | FSD1L | 0.161 | 0.107 | 0.649 | 0.447 | 0.247 | 4.5E-05 |
| 4 | FTO | 0.179 | 0.123 | 0.742 | 0.671 | 0.164 | 0.00011 |
| 4 | FUT8 | 0.287 | 0.099 | 1.461 | 0.44 | 1.437 | 1.1E-43 |
| 4 | FXR1 | 0.297 | 0.228 | 1.348 | 1.248 | 0.11 | 0.0007 |
| 4 | FXYD2 | 0.117 | 0.002 | 0.617 | 0.01 | 4.988 | 4E-304 |
| 4 | FXYD4 | 0.841 | 0.411 | 10.226 | 3.819 | 1.421 | 8E-114 |
| 4 | FXYD7 | 0.146 | 0.011 | 0.876 | 0.062 | 3.816 | 4E-160 |
| 4 | FYTTD1 | 0.344 | 0.248 | 1.633 | 1.391 | 0.231 | 6.5E-06 |
| 4 | G3BP2 | 0.446 | 0.331 | 2.213 | 1.937 | 0.192 | 2.8E-06 |
| 4 | GAB1 | 0.309 | 0.026 | 1.408 | 0.141 | 3.414 | 0 |
| 4 | GADD44B | 0.487 | 0.443 | 6.336 | 3.81 | 0.734 | 4.1E-17 |
| 4 | GADD44GIP1 | 0.143 | 0.093 | 0.419 | 0.442 | 0.232 | 4E-05 |
| 4 | GALNT11 | 0.194 | 0.126 | 0.784 | 0.672 | 0.222 | 4.9E-06 |
| 4 | GAPDH | 0.837 | 0.703 | 9.149 | 7.344 | 0.317 | 1.4E-10 |
| 4 | GARS | 0.201 | 0.14 | 0.92 | 0.72 | 0.342 | 2.6E-05 |
| 4 | GART | 0.139 | 0.098 | 0.444 | 0.491 | 0.147 | 0.00182 |
| 4 | GAS4 | 0.373 | 0.141 | 1.789 | 0.724 | 1.306 | 1.4E-49 |
| 4 | GAS7 | 0.317 | 0.184 | 1.649 | 1.093 | 0.602 | 7.1E-16 |
| 4 | GATA3 | 0.436 | 0.191 | 3.004 | 1.087 | 1.467 | 1.4E-48 |
| 4 | GBE1 | 0.134 | 0.064 | 0.437 | 0.334 | 0.682 | 4.3E-12 |
| 4 | GDI2 | 0.401 | 0.3 | 1.844 | 1.674 | 0.139 | 4.7E-05 |
| 4 | GIMAP6 | 0.116 | 0.084 | 0.401 | 0.424 | 0.237 | 0.00424 |
| 4 | GINM1 | 0.173 | 0.117 | 0.694 | 0.604 | 0.199 | 9.1E-05 |
| 4 | GLRX3 | 0.141 | 0.086 | 0.446 | 0.444 | 0.324 | 2E-06 |
| 4 | GLUD1 | 0.264 | 0.182 | 1.1 | 0.973 | 0.178 | 6.4E-06 |
| 4 | GMDS | 0.233 | 0.146 | 1.038 | 0.891 | 0.22 | 2.4E-06 |
| 4 | GMFG | 0.63 | 0.411 | 3.906 | 3.626 | 0.108 | 0.00068 |
| 4 | GNAQ | 0.244 | 0.044 | 1.131 | 0.297 | 1.931 | 2E-99 |
| 4 | GNAS | 0.443 | 0.346 | 2.383 | 2.187 | 0.124 | 0.00017 |

| | | | | | | | |
|---|---|---|---|---|---|---|---|
| 4 | GNG10 | 0.111 | 0.069 | 0.4 | 0.332 | 0.264 | 7.7E-05 |
| 4 | GNL2 | 0.16 | 0.094 | 0.603 | 0.494 | 0.288 | 2.9E-07 |
| 4 | GNL3 | 0.167 | 0.069 | 0.627 | 0.348 | 0.849 | 6.4E-21 |
| 4 | GNLY | 0.926 | 0.949 | 79.844 | 40.244 | 0.668 | 1.8E-32 |
| 4 | GOLGA3 | 0.14 | 0.098 | 0.434 | 0.497 | 0.103 | 0.00098 |
| 4 | GOLGA8B | 0.316 | 0.18 | 1.349 | 0.993 | 0.443 | 2.9E-14 |
| 4 | GOLGB1 | 0.43 | 0.326 | 2.48 | 2.101 | 0.239 | 2.6E-06 |
| 4 | GOLIM4 | 0.231 | 0.083 | 1.016 | 0.446 | 1.189 | 4.6E-39 |
| 4 | GPAA1 | 0.106 | 0.076 | 0.438 | 0.364 | 0.264 | 0.00842 |
| 4 | GPR183 | 0.37 | 0.013 | 2.01 | 0.073 | 4.774 | 0 |
| 4 | GPR64 | 0.401 | 0.292 | 2.044 | 1.816 | 0.171 | 3.2E-06 |
| 4 | GPR68 | 0.116 | 0.048 | 0.496 | 0.244 | 1.014 | 8.4E-14 |
| 4 | GPRIN3 | 0.299 | 0.2 | 1.383 | 1.179 | 0.231 | 1.5E-07 |
| 4 | GPSM3 | 0.361 | 0.267 | 1.702 | 1.499 | 0.184 | 1.7E-05 |
| 4 | GPX4 | 0.289 | 0.204 | 1.288 | 1.092 | 0.238 | 8.2E-06 |
| 4 | GRAMD1B | 0.109 | 0.024 | 0.442 | 0.13 | 2.049 | 1E-37 |
| 4 | GRHPR | 0.111 | 0.07 | 0.414 | 0.331 | 0.327 | 8.3E-05 |
| 4 | GSN | 0.149 | 0.044 | 0.62 | 0.271 | 1.194 | 2E-28 |
| 4 | GSPT1 | 0.277 | 0.176 | 1.174 | 0.943 | 0.317 | 4.5E-09 |
| 4 | GSTO1 | 0.179 | 0.124 | 0.747 | 0.628 | 0.269 | 0.00011 |
| 4 | GTF2F2 | 0.144 | 0.096 | 0.606 | 0.496 | 0.288 | 3.6E-06 |
| 4 | GTF2H1 | 0.174 | 0.12 | 0.761 | 0.609 | 0.321 | 7.5E-05 |
| 4 | GTF2IRD2B | 0.134 | 0.073 | 0.494 | 0.361 | 0.444 | 2E-08 |
| 4 | GTF3C6 | 0.126 | 0.092 | 0.416 | 0.44 | 0.198 | 0.00795 |
| 4 | GTPBP10 | 0.149 | 0.106 | 0.496 | 0.441 | 0.141 | 0.00013 |
| 4 | GTPBP4 | 0.133 | 0.084 | 0.442 | 0.43 | 0.362 | 2.2E-05 |
| 4 | GTPBP8 | 0.111 | 0.064 | 0.444 | 0.326 | 0.444 | 4.7E-06 |
| 4 | GUCY1A1 | 0.116 | 0.016 | 0.477 | 0.079 | 2.491 | 3.4E-74 |
| 4 | GZMK | 0.487 | 0.029 | 4.703 | 0.201 | 4.44 | 0 |
| 4 | H2AFJ | 0.173 | 0.096 | 0.792 | 0.404 | 0.64 | 1.2E-10 |
| 4 | H2AFY | 0.221 | 0.121 | 0.914 | 0.609 | 0.486 | 1.8E-13 |
| 4 | H2AFZ | 0.276 | 0.204 | 1.23 | 1.089 | 0.176 | 0.00022 |
| 4 | H3F3B | 0.941 | 0.881 | 22.242 | 16.827 | 0.403 | 7.7E-16 |
| 4 | HADHA | 0.436 | 0.321 | 2.167 | 1.884 | 0.201 | 1.2E-06 |
| 4 | HADHB | 0.199 | 0.149 | 0.841 | 0.776 | 0.116 | 0.00248 |
| 4 | HAPLN3 | 0.121 | 0.014 | 0.434 | 0.076 | 2.412 | 8.1E-84 |
| 4 | HAX1 | 0.134 | 0.083 | 0.471 | 0.414 | 0.461 | 7.3E-06 |
| 4 | HBS1L | 0.246 | 0.176 | 1.11 | 0.938 | 0.243 | 3.8E-06 |
| 4 | HDGFL3 | 0.127 | 0.048 | 0.421 | 0.292 | 0.833 | 2.1E-13 |
| 4 | HEBP2 | 0.187 | 0.118 | 0.822 | 0.494 | 0.464 | 2.4E-07 |
| 4 | HECTD1 | 0.311 | 0.249 | 1.429 | 1.422 | 0.104 | 0.00346 |
| 4 | HELB | 0.224 | 0.126 | 0.914 | 0.669 | 0.442 | 6.8E-12 |
| 4 | HERPUD1 | 0.371 | 0.213 | 2.044 | 1.209 | 0.764 | 1E-20 |
| 4 | HIBCH | 0.164 | 0.116 | 0.706 | 0.622 | 0.182 | 0.00064 |
| 4 | HIF1A | 0.761 | 0.481 | 6.971 | 4.492 | 0.344 | 6.3E-17 |
| 4 | HIKESHI | 0.114 | 0.081 | 0.438 | 0.394 | 0.147 | 0.0038 |
| 4 | HINT1 | 0.701 | 0.443 | 4.896 | 2.937 | 0.737 | 9.4E-40 |
| 4 | HIP1 | 0.244 | 0.124 | 1.119 | 0.694 | 0.69 | 6.1E-21 |
| 4 | HIST1H4C | 0.731 | 0.603 | 6.248 | 4.442 | 0.202 | 2.1E-06 |
| 4 | HIVEP1 | 0.204 | 0.12 | 0.899 | 0.648 | 0.471 | 7E-10 |
| 4 | HIVEP2 | 0.269 | 0.187 | 1.176 | 1.089 | 0.111 | 1.4E-05 |
| 4 | HLA-DRB1 | 0.2 | 0.108 | 0.993 | 0.647 | 0.618 | 1E-12 |
| 4 | HLA-DRB4 | 0.101 | 0.063 | 0.499 | 0.343 | 0.497 | 8.2E-05 |
| 4 | HMG20A | 0.111 | 0.078 | 0.427 | 0.387 | 0.143 | 0.00298 |

| | | | | | | | |
|---|---|---|---|---|---|---|---|
| 4 | HMGA1 | 0.113 | 0.073 | 0.42 | 0.349 | 0.268 | 0.00019 |
| 4 | HMGCR | 0.133 | 0.073 | 0.442 | 0.346 | 0.606 | 1.9E-08 |
| 4 | HMGCS1 | 0.24 | 0.141 | 1.146 | 0.82 | 0.483 | 3.3E-09 |
| 4 | HMGN1 | 0.443 | 0.426 | 3.187 | 2.798 | 0.188 | 4.9E-06 |
| 4 | HNRNPA0 | 0.406 | 0.293 | 2.044 | 1.649 | 0.307 | 3.7E-07 |
| 4 | HNRNPAB | 0.34 | 0.249 | 1.601 | 1.466 | 0.127 | 0.0005 |
| 4 | HNRNPH2 | 0.207 | 0.143 | 0.883 | 0.706 | 0.324 | 1.4E-05 |
| 4 | HNRNPH3 | 0.227 | 0.16 | 0.932 | 0.847 | 0.138 | 7.7E-05 |
| 4 | HNRNPM | 0.467 | 0.433 | 3.294 | 2.89 | 0.189 | 2.7E-06 |
| 4 | HOOK2 | 0.19 | 0.137 | 0.879 | 0.78 | 0.173 | 0.00047 |
| 4 | HOPX | 0.614 | 0.443 | 4.414 | 3.484 | 0.333 | 4.7E-12 |
| 4 | HPRT1 | 0.129 | 0.074 | 0.487 | 0.346 | 0.441 | 3.5E-07 |
| 4 | HSP90AA1 | 0.634 | 0.418 | 4.274 | 3.909 | 0.129 | 9.9E-05 |
| 4 | HSP90AB1 | 0.843 | 0.467 | 11.348 | 4.144 | 1.139 | 7.1E-77 |
| 4 | HSP90B1 | 0.681 | 0.468 | 4.614 | 4.89 | 0.2 | 0.00013 |
| 4 | HSPA13 | 0.103 | 0.049 | 0.416 | 0.284 | 0.444 | 4.1E-06 |
| 4 | HSPA4 | 0.714 | 0.469 | 6.676 | 4.492 | 0.246 | 9.5E-09 |
| 4 | HSPA9 | 0.336 | 0.221 | 1.443 | 1.222 | 0.336 | 3.8E-09 |
| 4 | HSPB1 | 0.141 | 0.069 | 0.684 | 0.337 | 1.019 | 4E-16 |
| 4 | HSPD1 | 0.434 | 0.209 | 2.363 | 1.187 | 0.994 | 2.6E-41 |
| 4 | HSPE1 | 0.263 | 0.141 | 1.266 | 0.743 | 0.748 | 4.9E-18 |
| 4 | HSPH1 | 0.183 | 0.124 | 0.783 | 0.691 | 0.181 | 4.7E-05 |
| 4 | HTATSF1 | 0.249 | 0.167 | 1.077 | 0.884 | 0.284 | 1.4E-06 |
| 4 | HUWE1 | 0.343 | 0.238 | 1.623 | 1.342 | 0.264 | 1.2E-07 |
| 4 | HVCN1 | 0.23 | 0.029 | 1.11 | 0.144 | 2.94 | 8E-164 |
| 4 | IAH1 | 0.119 | 0.074 | 0.427 | 0.362 | 0.239 | 4.4E-05 |
| 4 | IARS | 0.146 | 0.088 | 0.646 | 0.46 | 0.413 | 8.1E-07 |
| 4 | ICAM3 | 0.21 | 0.13 | 0.882 | 0.643 | 0.433 | 2.2E-08 |
| 4 | ID2 | 0.449 | 0.306 | 2.794 | 2.042 | 0.446 | 1.8E-13 |
| 4 | IER2 | 0.764 | 0.647 | 14.944 | 10.223 | 0.641 | 2.5E-20 |
| 4 | IER3 | 0.143 | 0.044 | 0.821 | 0.242 | 1.762 | 1.2E-39 |
| 4 | IER4 | 0.106 | 0.044 | 0.444 | 0.28 | 0.671 | 3.3E-08 |
| 4 | IFITM2 | 0.941 | 0.774 | 16.64 | 10.102 | 0.721 | 1.6E-64 |
| 4 | IFITM3 | 0.923 | 0.747 | 21.171 | 12.173 | 0.798 | 4.4E-68 |
| 4 | IFNG-AS1 | 0.2 | 0.023 | 1.11 | 0.127 | 3.127 | 5E-144 |
| 4 | IFRD1 | 0.424 | 0.228 | 2.94 | 1.44 | 0.933 | 4.7E-32 |
| 4 | IFT47 | 0.106 | 0.044 | 0.391 | 0.269 | 0.438 | 7.1E-08 |
| 4 | IGFLR1 | 0.199 | 0.121 | 0.797 | 0.497 | 0.416 | 2.3E-08 |
| 4 | IL10RA | 0.436 | 0.449 | 3.388 | 3.043 | 0.14 | 0.00099 |
| 4 | IL12RB2 | 0.344 | 0.092 | 2.134 | 0.448 | 1.961 | 4E-110 |
| 4 | IL18R1 | 0.361 | 0.144 | 2.087 | 0.804 | 1.374 | 1.9E-44 |
| 4 | IL2RA | 0.12 | 0.008 | 0.834 | 0.042 | 4.312 | 2E-144 |
| 4 | IL4R | 0.21 | 0.046 | 0.941 | 0.29 | 1.698 | 1E-49 |
| 4 | IL7R | 0.631 | 0.041 | 7.737 | 0.343 | 4.444 | 0 |
| 4 | ILF2 | 0.244 | 0.18 | 1.087 | 0.917 | 0.246 | 1.4E-05 |
| 4 | ILF3 | 0.343 | 0.242 | 1.739 | 1.443 | 0.249 | 1.6E-06 |
| 4 | ILF3-DT | 0.166 | 0.11 | 0.664 | 0.461 | 0.244 | 4.1E-05 |
| 4 | IMMP2L | 0.399 | 0.163 | 2.223 | 0.996 | 1.149 | 8E-46 |
| 4 | INPP4B | 0.301 | 0.011 | 1.988 | 0.063 | 4.971 | 0 |
| 4 | INPP4F | 0.106 | 0.044 | 0.399 | 0.273 | 0.44 | 4.4E-08 |
| 4 | INTS8 | 0.14 | 0.094 | 0.619 | 0.493 | 0.33 | 6.7E-06 |
| 4 | IPMK | 0.117 | 0.072 | 0.429 | 0.362 | 0.244 | 3.4E-05 |
| 4 | IPO4 | 0.116 | 0.069 | 0.447 | 0.341 | 0.381 | 6.3E-06 |
| 4 | IRAK3 | 0.107 | 0.006 | 0.477 | 0.032 | 3.899 | 2E-148 |

| | | | | | | | |
|---|---|---|---|---|---|---|---|
| 4 | IRF2BP2 | 0.149 | 0.113 | 0.683 | 0.492 | 0.207 | 0.00481 |
| 4 | ITGA1 | 0.131 | 0.092 | 0.63 | 0.42 | 0.279 | 0.00121 |
| 4 | ITGA4 | 0.643 | 0.41 | 4.606 | 3.937 | 0.227 | 6.9E-08 |
| 4 | ITGAM | 0.117 | 0.074 | 0.476 | 0.404 | 0.236 | 0.00013 |
| 4 | ITGAX | 0.247 | 0.096 | 1.42 | 0.447 | 1.341 | 1E-41 |
| 4 | ITGB1 | 0.727 | 0.448 | 4.993 | 4.467 | 0.424 | 1.1E-14 |
| 4 | ITM2A | 0.243 | 0.112 | 1.04 | 0.484 | 0.847 | 2.7E-23 |
| 4 | ITM2C | 0.249 | 0.073 | 1.294 | 0.42 | 1.623 | 4E-61 |
| 4 | ITPRID2 | 0.106 | 0.074 | 0.441 | 0.382 | 0.238 | 0.00343 |
| 4 | IVD | 0.134 | 0.093 | 0.411 | 0.47 | 0.121 | 0.00108 |
| 4 | JADE1 | 0.133 | 0.086 | 0.416 | 0.447 | 0.209 | 7.2E-05 |
| 4 | JUN | 0.449 | 0.498 | 11.223 | 9.076 | 0.306 | 0.00018 |
| 4 | JUNB | 0.469 | 0.397 | 6.197 | 3.902 | 0.667 | 4E-19 |
| 4 | JUND | 0.243 | 0.144 | 1.329 | 0.917 | 0.434 | 4.4E-11 |
| 4 | KARS | 0.147 | 0.104 | 0.606 | 0.433 | 0.184 | 0.00014 |
| 4 | KCNQ4 | 0.176 | 0.064 | 0.89 | 0.419 | 1.086 | 1.4E-28 |
| 4 | KDM4B | 0.299 | 0.183 | 1.376 | 1.026 | 0.424 | 1.2E-11 |
| 4 | KDM6A | 0.447 | 0.324 | 2.404 | 2.124 | 0.179 | 2.6E-08 |
| 4 | KDM6B | 0.106 | 0.074 | 0.421 | 0.387 | 0.12 | 0.00461 |
| 4 | KDSR | 0.11 | 0.029 | 0.473 | 0.146 | 1.604 | 8.4E-32 |
| 4 | KHDRBS1 | 0.466 | 0.329 | 2.308 | 1.942 | 0.249 | 9.6E-09 |
| 4 | KIAA0319L | 0.186 | 0.103 | 0.827 | 0.441 | 0.613 | 3.7E-11 |
| 4 | KIR2DL4 | 0.177 | 0.036 | 0.924 | 0.188 | 2.297 | 1.6E-74 |
| 4 | KIT | 0.12 | 0.004 | 0.608 | 0.023 | 4.701 | 9E-220 |
| 4 | KLF10 | 0.306 | 0.161 | 1.891 | 0.911 | 1.044 | 2E-23 |
| 4 | KLF2 | 0.184 | 0.134 | 1.012 | 0.812 | 0.317 | 0.00035 |
| 4 | KLF6 | 0.743 | 0.611 | 9.68 | 7.242 | 0.419 | 1.9E-11 |
| 4 | KLF7 | 0.113 | 0.068 | 0.448 | 0.344 | 0.336 | 1.6E-05 |
| 4 | KLHL28 | 0.199 | 0.142 | 0.831 | 0.747 | 0.144 | 0.00034 |
| 4 | KLHL4 | 0.177 | 0.103 | 0.862 | 0.436 | 0.686 | 1.3E-09 |
| 4 | KLHL6 | 0.141 | 0.096 | 0.498 | 0.411 | 0.227 | 1.4E-05 |
| 4 | KLRC1 | 0.441 | 0.244 | 4.463 | 2.126 | 1.07 | 4.8E-64 |
| 4 | KLRC2 | 0.631 | 0.449 | 6.704 | 4.31 | 0.337 | 1.4E-13 |
| 4 | KLRC3 | 0.614 | 0.462 | 6.113 | 4.686 | 0.383 | 3.3E-12 |
| 4 | KLRK1 | 0.747 | 0.64 | 6.792 | 4.969 | 0.186 | 4.5E-05 |
| 4 | KRCC1 | 0.191 | 0.139 | 0.829 | 0.724 | 0.194 | 0.00085 |
| 4 | KRI1 | 0.106 | 0.067 | 0.394 | 0.332 | 0.248 | 0.00024 |
| 4 | KRR1 | 0.229 | 0.144 | 0.92 | 0.802 | 0.198 | 9.9E-06 |
| 4 | KRT10 | 0.33 | 0.243 | 1.498 | 1.364 | 0.134 | 0.00047 |
| 4 | LACTB | 0.134 | 0.068 | 0.431 | 0.347 | 0.614 | 1.9E-10 |
| 4 | LAGE3 | 0.119 | 0.076 | 0.436 | 0.361 | 0.27 | 0.00012 |
| 4 | LAMTOR1 | 0.263 | 0.193 | 1.132 | 1 | 0.178 | 0.00016 |
| 4 | LAMTOR3 | 0.111 | 0.077 | 0.439 | 0.373 | 0.234 | 0.00239 |
| 4 | LAMTOR4 | 0.203 | 0.134 | 0.806 | 0.664 | 0.28 | 4.1E-06 |
| 4 | LARP1 | 0.12 | 0.067 | 0.472 | 0.348 | 0.442 | 2.9E-07 |
| 4 | LARP1B | 0.103 | 0.046 | 0.372 | 0.238 | 0.643 | 2.8E-11 |
| 4 | LARS | 0.204 | 0.131 | 0.842 | 0.691 | 0.302 | 7.8E-07 |
| 4 | LATS2 | 0.229 | 0.169 | 1.083 | 0.929 | 0.222 | 0.00023 |
| 4 | LDHB | 0.419 | 0.211 | 2.113 | 1.134 | 0.897 | 2.1E-34 |
| 4 | LDLRAD4 | 0.237 | 0.147 | 1.074 | 0.924 | 0.216 | 1.2E-06 |
| 4 | LEF1 | 0.109 | 0.014 | 0.44 | 0.073 | 2.63 | 3.2E-77 |
| 4 | LENG8 | 0.119 | 0.082 | 0.488 | 0.424 | 0.201 | 0.00145 |
| 4 | LEPROTL1 | 0.439 | 0.293 | 2.239 | 1.696 | 0.4 | 9.6E-13 |
| 4 | LETMD1 | 0.103 | 0.073 | 0.424 | 0.344 | 0.249 | 0.00434 |

| | | | | | | | |
|---|---|---|---|---|---|---|---|
| 4 | LIN44 | 0.146 | 0.106 | 0.477 | 0.434 | 0.107 | 0.00324 |
| 4 | LINC00910 | 0.227 | 0.144 | 1.371 | 1.048 | 0.387 | 1.5E-06 |
| 4 | LINC00996 | 0.363 | 0.129 | 3.489 | 0.976 | 1.878 | 1.4E-73 |
| 4 | LINC01138 | 0.486 | 0.469 | 3.817 | 3.404 | 0.164 | 4.9E-05 |
| 4 | LINC01473 | 0.137 | 0.08 | 0.608 | 0.464 | 0.387 | 3E-07 |
| 4 | LINC01478 | 0.183 | 0.133 | 0.764 | 0.714 | 0.1 | 0.00107 |
| 4 | LINC01684 | 0.11 | 0.049 | 0.404 | 0.298 | 0.443 | 1.2E-07 |
| 4 | LINC01871 | 0.146 | 0.096 | 0.747 | 0.423 | 0.434 | 8.4E-07 |
| 4 | LINC02246 | 0.363 | 0.273 | 1.796 | 1.612 | 0.146 | 8.8E-05 |
| 4 | LMNA | 0.133 | 0.024 | 0.704 | 0.124 | 2.408 | 4E-63 |
| 4 | LPXN | 0.3 | 0.199 | 1.343 | 1.072 | 0.337 | 4.4E-08 |
| 4 | LRCH1 | 0.197 | 0.136 | 0.847 | 0.722 | 0.231 | 3.7E-05 |
| 4 | LRIG2 | 0.111 | 0.074 | 0.479 | 0.364 | 0.391 | 0.00072 |
| 4 | LRPPRC | 0.249 | 0.167 | 1.042 | 0.886 | 0.247 | 1.2E-06 |
| 4 | LRRC8D | 0.101 | 0.048 | 0.338 | 0.293 | 0.206 | 9.1E-06 |
| 4 | LRRFIP1 | 0.643 | 0.493 | 4.431 | 3.646 | 0.314 | 1.3E-09 |
| 4 | LSM2 | 0.187 | 0.132 | 0.773 | 0.643 | 0.264 | 0.00016 |
| 4 | LSM4 | 0.177 | 0.092 | 0.746 | 0.444 | 0.718 | 7.8E-13 |
| 4 | LSM4 | 0.173 | 0.107 | 0.619 | 0.424 | 0.241 | 1.3E-06 |
| 4 | LSM6 | 0.14 | 0.101 | 0.439 | 0.49 | 0.136 | 0.00294 |
| 4 | LSM7 | 0.28 | 0.197 | 1.14 | 1.013 | 0.183 | 1.8E-05 |
| 4 | LTA | 0.107 | 0.017 | 0.611 | 0.083 | 2.874 | 7.5E-60 |
| 4 | LTA4H | 0.23 | 0.103 | 1.141 | 0.418 | 1.141 | 1.1E-24 |
| 4 | LTB | 0.293 | 0.024 | 2.273 | 0.141 | 4.014 | 1E-304 |
| 4 | LTV1 | 0.107 | 0.076 | 0.433 | 0.389 | 0.143 | 0.00444 |
| 4 | LY74 | 0.243 | 0.2 | 1.203 | 1.116 | 0.109 | 0.00419 |
| 4 | LYPLA1 | 0.193 | 0.13 | 0.743 | 0.662 | 0.184 | 2.7E-05 |
| 4 | LYRM7 | 0.104 | 0.061 | 0.426 | 0.303 | 0.493 | 8.1E-06 |
| 4 | MADD | 0.121 | 0.086 | 0.414 | 0.464 | 0.144 | 0.00228 |
| 4 | MAFF | 0.27 | 0.101 | 1.406 | 0.462 | 1.324 | 1.6E-43 |
| 4 | MAGED1 | 0.147 | 0.092 | 0.634 | 0.49 | 0.374 | 1.4E-07 |
| 4 | MAGED2 | 0.107 | 0.074 | 0.404 | 0.347 | 0.179 | 0.00269 |
| 4 | MAGOHB | 0.14 | 0.064 | 0.622 | 0.327 | 0.929 | 6.9E-14 |
| 4 | MAGT1 | 0.231 | 0.146 | 1.018 | 0.827 | 0.3 | 2.5E-06 |
| 4 | MALT1 | 0.214 | 0.134 | 0.897 | 0.723 | 0.312 | 8.5E-08 |
| 4 | MAML2 | 0.446 | 0.207 | 2.668 | 1.323 | 1.012 | 3.1E-44 |
| 4 | MAML3 | 0.271 | 0.03 | 1.388 | 0.178 | 2.964 | 6E-218 |
| 4 | MAP3K1 | 0.17 | 0.094 | 0.7 | 0.401 | 0.482 | 1.1E-09 |
| 4 | MAP3K8 | 0.671 | 0.42 | 7.306 | 3.434 | 1.048 | 6E-40 |
| 4 | MAPK1IP1L | 0.296 | 0.214 | 1.347 | 1.242 | 0.127 | 6.7E-05 |
| 4 | MAPK6 | 0.141 | 0.096 | 0.601 | 0.49 | 0.293 | 8.4E-06 |
| 4 | MARK2 | 0.14 | 0.097 | 0.616 | 0.492 | 0.326 | 1.9E-05 |
| 4 | MATK | 0.263 | 0.181 | 1.187 | 1.003 | 0.243 | 2.7E-06 |
| 4 | MBD1 | 0.109 | 0.071 | 0.414 | 0.346 | 0.219 | 0.00049 |
| 4 | MBD2 | 0.111 | 0.073 | 0.442 | 0.372 | 0.248 | 0.00039 |
| 4 | MCCC2 | 0.113 | 0.074 | 0.396 | 0.37 | 0.1 | 0.00044 |
| 4 | MCFD2 | 0.13 | 0.092 | 0.411 | 0.46 | 0.142 | 0.00218 |
| 4 | MCRIP1 | 0.124 | 0.087 | 0.444 | 0.42 | 0.117 | 0.00208 |
| 4 | MCUB | 0.383 | 0.199 | 2.033 | 1.14 | 0.834 | 4.5E-29 |
| 4 | MDFIC | 0.309 | 0.117 | 1.734 | 0.631 | 1.449 | 3.7E-40 |
| 4 | MDH1 | 0.223 | 0.166 | 0.93 | 0.844 | 0.121 | 0.00071 |
| 4 | MDN1 | 0.146 | 0.084 | 0.481 | 0.43 | 0.434 | 9.2E-08 |
| 4 | MECOM | 0.134 | 0.081 | 0.447 | 0.386 | 0.43 | 1.6E-06 |
| 4 | MEF2A | 0.419 | 0.208 | 2.148 | 1.183 | 0.867 | 2.5E-34 |

| | | | | | | | |
|---|---|---|---|---|---|---|---|
| 4 | METAP2 | 0.231 | 0.141 | 0.963 | 0.817 | 0.238 | 4.7E-07 |
| 4 | METTL16 | 0.147 | 0.114 | 0.64 | 0.604 | 0.104 | 0.00212 |
| 4 | METTL26 | 0.171 | 0.104 | 0.699 | 0.424 | 0.416 | 2.8E-07 |
| 4 | MGAT4 | 0.21 | 0.101 | 0.982 | 0.464 | 0.798 | 2.2E-18 |
| 4 | MGRN1 | 0.107 | 0.064 | 0.399 | 0.332 | 0.264 | 2.4E-05 |
| 4 | MIB1 | 0.147 | 0.104 | 0.64 | 0.444 | 0.231 | 4.7E-05 |
| 4 | MIF | 0.636 | 0.417 | 4.044 | 2.647 | 0.614 | 3.8E-27 |
| 4 | MIR181A1HG | 0.213 | 0.102 | 1.077 | 0.468 | 0.924 | 1.9E-19 |
| 4 | MIR222HG | 0.107 | 0.036 | 0.403 | 0.216 | 1.218 | 7.5E-21 |
| 4 | MIR924HG | 0.363 | 0.213 | 1.83 | 1.14 | 0.67 | 1.8E-17 |
| 4 | MIS18BP1 | 0.283 | 0.174 | 1.24 | 0.949 | 0.382 | 1.5E-10 |
| 4 | MLH1 | 0.13 | 0.081 | 0.413 | 0.414 | 0.311 | 2.7E-05 |
| 4 | MMP24-AS1 | 0.21 | 0.06 | 0.876 | 0.309 | 1.404 | 9.4E-43 |
| 4 | MOB1B | 0.131 | 0.094 | 0.434 | 0.476 | 0.167 | 0.00369 |
| 4 | MOB3A | 0.14 | 0.091 | 0.629 | 0.484 | 0.374 | 7.6E-07 |
| 4 | MOB4 | 0.311 | 0.212 | 1.268 | 1.13 | 0.167 | 8.4E-07 |
| 4 | MORF4L2 | 0.286 | 0.167 | 1.337 | 0.921 | 0.437 | 7.7E-14 |
| 4 | MPC2 | 0.191 | 0.124 | 0.83 | 0.622 | 0.414 | 3.4E-06 |
| 4 | MPG | 0.161 | 0.083 | 0.698 | 0.404 | 0.789 | 3.5E-12 |
| 4 | MPHOSPH10 | 0.13 | 0.092 | 0.601 | 0.46 | 0.387 | 0.0012 |
| 4 | MPZL3 | 0.189 | 0.038 | 0.886 | 0.194 | 2.184 | 2.4E-78 |
| 4 | MRPL14 | 0.127 | 0.064 | 0.44 | 0.294 | 0.616 | 3.4E-10 |
| 4 | MRPL18 | 0.106 | 0.061 | 0.389 | 0.293 | 0.406 | 6.8E-06 |
| 4 | MRPL28 | 0.107 | 0.074 | 0.433 | 0.34 | 0.308 | 0.00172 |
| 4 | MRPL3 | 0.164 | 0.09 | 0.602 | 0.442 | 0.446 | 6.6E-10 |
| 4 | MRPL33 | 0.171 | 0.119 | 0.724 | 0.609 | 0.24 | 0.00015 |
| 4 | MRPL41 | 0.14 | 0.107 | 0.604 | 0.416 | 0.227 | 0.00112 |
| 4 | MRPL42 | 0.19 | 0.121 | 0.72 | 0.487 | 0.293 | 2.2E-06 |
| 4 | MRPL42 | 0.144 | 0.092 | 0.444 | 0.444 | 0.292 | 4.9E-07 |
| 4 | MRPL44 | 0.144 | 0.083 | 0.474 | 0.393 | 0.449 | 8.3E-10 |
| 4 | MRPL47 | 0.231 | 0.166 | 0.948 | 0.844 | 0.182 | 0.00015 |
| 4 | MRPS12 | 0.116 | 0.071 | 0.383 | 0.333 | 0.203 | 5E-05 |
| 4 | MRPS18C | 0.183 | 0.134 | 0.768 | 0.684 | 0.164 | 0.00142 |
| 4 | MRPS21 | 0.289 | 0.216 | 1.284 | 1.144 | 0.166 | 0.00026 |
| 4 | MRPS24 | 0.109 | 0.078 | 0.414 | 0.366 | 0.179 | 0.00697 |
| 4 | MRPS27 | 0.126 | 0.092 | 0.487 | 0.444 | 0.101 | 0.00661 |
| 4 | MRPS33 | 0.124 | 0.061 | 0.48 | 0.299 | 0.684 | 1.1E-10 |
| 4 | MRPS34 | 0.144 | 0.108 | 0.617 | 0.436 | 0.203 | 0.00045 |
| 4 | MRPS6 | 0.384 | 0.284 | 2.114 | 1.714 | 0.303 | 1E-06 |
| 4 | MSMO1 | 0.137 | 0.094 | 0.44 | 0.471 | 0.224 | 0.0004 |
| 4 | MTAP | 0.134 | 0.069 | 0.443 | 0.369 | 0.447 | 3.4E-10 |
| 4 | MTCH2 | 0.136 | 0.1 | 0.448 | 0.408 | 0.111 | 0.0071 |
| 4 | MTHFD2 | 0.146 | 0.096 | 0.748 | 0.487 | 0.638 | 1.1E-06 |
| 4 | MTHFS | 0.146 | 0.066 | 0.488 | 0.32 | 0.878 | 3E-14 |
| 4 | MTM1 | 0.146 | 0.033 | 0.614 | 0.17 | 1.847 | 1.5E-42 |
| 4 | MTMR2 | 0.167 | 0.109 | 0.67 | 0.466 | 0.244 | 1.9E-05 |
| 4 | MT-ND1 | 0.683 | 0.447 | 4.806 | 4.312 | 0.147 | 6.8E-07 |
| 4 | MT-ND2 | 0.917 | 0.814 | 13.128 | 10.474 | 0.312 | 8.9E-18 |
| 4 | MTRF1L | 0.137 | 0.097 | 0.431 | 0.486 | 0.126 | 0.00193 |
| 4 | MXD1 | 0.163 | 0.108 | 0.674 | 0.444 | 0.281 | 3.5E-05 |
| 4 | MYADM | 0.247 | 0.093 | 1.669 | 0.427 | 1.663 | 7.9E-46 |
| 4 | MYD88 | 0.106 | 0.064 | 0.382 | 0.312 | 0.291 | 8E-05 |
| 4 | MZT2B | 0.196 | 0.143 | 0.784 | 0.71 | 0.146 | 0.0011 |
| 4 | N4BP2 | 0.171 | 0.114 | 0.718 | 0.609 | 0.239 | 4.2E-05 |

| | | | | | | | |
|---|---|---|---|---|---|---|---|
| 4 | NAA16 | 0.19 | 0.122 | 0.742 | 0.643 | 0.227 | 2.4E-06 |
| 4 | NAA20 | 0.143 | 0.092 | 0.432 | 0.442 | 0.267 | 2.7E-05 |
| 4 | NAA34 | 0.199 | 0.134 | 0.804 | 0.712 | 0.176 | 1.5E-05 |
| 4 | NACA | 0.903 | 0.744 | 10.364 | 7.714 | 0.426 | 9.8E-24 |
| 4 | NAMPT | 0.22 | 0.119 | 1.028 | 0.644 | 0.674 | 4.9E-14 |
| 4 | NAP1L1 | 0.443 | 0.344 | 3.299 | 2.11 | 0.644 | 6.7E-26 |
| 4 | NARF | 0.276 | 0.174 | 1.179 | 0.972 | 0.279 | 4.3E-09 |
| 4 | NBDY | 0.243 | 0.109 | 1.219 | 0.442 | 1.144 | 6.2E-30 |
| 4 | NBEAL1 | 0.361 | 0.247 | 1.764 | 1.467 | 0.266 | 4.7E-07 |
| 4 | NBPF10 | 0.109 | 0.076 | 0.433 | 0.386 | 0.167 | 0.00291 |
| 4 | NBPF14 | 0.18 | 0.131 | 0.742 | 0.684 | 0.137 | 0.00114 |
| 4 | NCALD | 0.496 | 0.369 | 2.998 | 2.644 | 0.181 | 1.1E-07 |
| 4 | NCAM1 | 0.474 | 0.233 | 3.249 | 1.493 | 1.122 | 4.8E-48 |
| 4 | NCL | 0.807 | 0.707 | 7.44 | 6.779 | 0.143 | 0.00042 |
| 4 | NCOA4 | 0.247 | 0.193 | 1.101 | 1.014 | 0.118 | 0.00128 |
| 4 | NDUFA10 | 0.203 | 0.139 | 0.844 | 0.711 | 0.246 | 3.2E-05 |
| 4 | NDUFA4 | 0.201 | 0.144 | 0.79 | 0.73 | 0.114 | 0.00041 |
| 4 | NDUFAB1 | 0.207 | 0.144 | 0.834 | 0.766 | 0.124 | 0.00232 |
| 4 | NDUFAF3 | 0.207 | 0.142 | 0.808 | 0.713 | 0.18 | 4.1E-05 |
| 4 | NDUFAF8 | 0.149 | 0.088 | 0.444 | 0.44 | 0.307 | 6.7E-07 |
| 4 | NDUFB3 | 0.14 | 0.089 | 0.44 | 0.414 | 0.382 | 1.9E-05 |
| 4 | NDUFB9 | 0.314 | 0.224 | 1.332 | 1.224 | 0.122 | 4.3E-05 |
| 4 | NDUFC2 | 0.361 | 0.281 | 1.708 | 1.421 | 0.167 | 0.00046 |
| 4 | NDUFS7 | 0.139 | 0.094 | 0.49 | 0.442 | 0.118 | 0.00046 |
| 4 | NDUFS8 | 0.213 | 0.136 | 0.837 | 0.662 | 0.34 | 3.4E-07 |
| 4 | NDUFV3 | 0.149 | 0.107 | 0.482 | 0.432 | 0.131 | 0.00014 |
| 4 | NECAP2 | 0.136 | 0.097 | 0.444 | 0.479 | 0.183 | 0.00209 |
| 4 | NELFCD | 0.116 | 0.08 | 0.442 | 0.386 | 0.194 | 0.00178 |
| 4 | NEPRO | 0.13 | 0.09 | 0.431 | 0.443 | 0.23 | 0.0008 |
| 4 | NFE2L2 | 0.389 | 0.273 | 2.004 | 1.643 | 0.287 | 7.3E-09 |
| 4 | NFKB1 | 0.489 | 0.394 | 4.84 | 3.066 | 0.649 | 1.3E-23 |
| 4 | NFKB2 | 0.111 | 0.063 | 0.462 | 0.312 | 0.466 | 6.1E-07 |
| 4 | NFKBIA | 0.803 | 0.724 | 31.382 | 12.31 | 1.34 | 2.3E-38 |
| 4 | NFKBIZ | 0.37 | 0.223 | 2.634 | 1.603 | 0.716 | 1.1E-19 |
| 4 | NHP2 | 0.173 | 0.084 | 0.631 | 0.402 | 0.641 | 1.2E-14 |
| 4 | NIBAN1 | 0.341 | 0.221 | 1.809 | 1.279 | 0.401 | 1.9E-13 |
| 4 | NIFK | 0.149 | 0.083 | 0.477 | 0.408 | 0.4 | 1E-08 |
| 4 | NINJ1 | 0.141 | 0.077 | 0.613 | 0.384 | 0.673 | 3.4E-09 |
| 4 | NIPSNAP2 | 0.173 | 0.113 | 0.648 | 0.481 | 0.18 | 2E-05 |
| 4 | NIT2 | 0.101 | 0.047 | 0.342 | 0.284 | 0.309 | 4.2E-06 |
| 4 | NMD3 | 0.114 | 0.046 | 0.461 | 0.283 | 0.704 | 6.5E-10 |
| 4 | NME2 | 0.423 | 0.21 | 3.14 | 1.163 | 1.432 | 1.9E-84 |
| 4 | NME3 | 0.16 | 0.119 | 0.672 | 0.482 | 0.207 | 0.00262 |
| 4 | NOA1 | 0.109 | 0.064 | 0.428 | 0.312 | 0.449 | 1.6E-05 |
| 4 | NOCT | 0.117 | 0.074 | 0.494 | 0.388 | 0.347 | 6.1E-05 |
| 4 | NOL10 | 0.109 | 0.077 | 0.43 | 0.393 | 0.132 | 0.00489 |
| 4 | NOL11 | 0.121 | 0.07 | 0.446 | 0.349 | 0.387 | 1.4E-06 |
| 4 | NOL7 | 0.241 | 0.168 | 0.969 | 0.887 | 0.127 | 4.7E-05 |
| 4 | NOLC1 | 0.204 | 0.126 | 0.891 | 0.632 | 0.494 | 2.4E-08 |
| 4 | NOP10 | 0.317 | 0.233 | 1.476 | 1.244 | 0.236 | 3.7E-05 |
| 4 | NOP14 | 0.161 | 0.099 | 0.637 | 0.404 | 0.334 | 7.4E-07 |
| 4 | NOP43 | 0.321 | 0.184 | 1.464 | 0.98 | 0.674 | 3.8E-17 |
| 4 | NOP46 | 0.129 | 0.077 | 0.481 | 0.387 | 0.314 | 3.8E-06 |
| 4 | NORAD | 0.446 | 0.392 | 3.14 | 2.467 | 0.342 | 1.6E-11 |

| | | | | | | | |
|---|---|---|---|---|---|---|---|
| 4 | NOSIP | 0.176 | 0.101 | 0.734 | 0.491 | 0.481 | 2.6E-09 |
| 4 | NOTCH2 | 0.214 | 0.146 | 0.924 | 0.837 | 0.144 | 0.00046 |
| 4 | NPM1 | 0.69 | 0.469 | 4.064 | 3.282 | 0.626 | 6.1E-29 |
| 4 | NR4A1 | 0.13 | 0.043 | 0.844 | 0.239 | 1.822 | 1.5E-26 |
| 4 | NR4A2 | 0.249 | 0.18 | 1.429 | 1.174 | 0.38 | 9.4E-06 |
| 4 | NRIP1 | 0.294 | 0.188 | 1.418 | 1.063 | 0.414 | 4.3E-10 |
| 4 | NSA2 | 0.363 | 0.266 | 1.746 | 1.414 | 0.214 | 9E-06 |
| 4 | NSD3 | 0.417 | 0.322 | 2.11 | 1.941 | 0.121 | 0.00037 |
| 4 | NSMCE1 | 0.201 | 0.073 | 0.919 | 0.348 | 1.401 | 9.9E-34 |
| 4 | NSUN2 | 0.187 | 0.127 | 0.733 | 0.674 | 0.121 | 6.8E-05 |
| 4 | NSUN6 | 0.107 | 0.068 | 0.463 | 0.346 | 0.418 | 0.00013 |
| 4 | NT4C | 0.187 | 0.131 | 0.718 | 0.641 | 0.14 | 0.00021 |
| 4 | NT4DC1 | 0.209 | 0.114 | 0.919 | 0.602 | 0.609 | 1.1E-12 |
| 4 | NUCB2 | 0.303 | 0.187 | 1.443 | 1.044 | 0.473 | 6.3E-13 |
| 4 | NUDCD2 | 0.119 | 0.074 | 0.488 | 0.361 | 0.433 | 3.4E-05 |
| 4 | NUDT4 | 0.143 | 0.102 | 0.478 | 0.416 | 0.163 | 0.00014 |
| 4 | NUP160 | 0.117 | 0.08 | 0.446 | 0.414 | 0.103 | 0.00169 |
| 4 | NUP48 | 0.201 | 0.14 | 0.82 | 0.744 | 0.138 | 9.5E-05 |
| 4 | NUS1 | 0.134 | 0.087 | 0.441 | 0.446 | 0.279 | 4.4E-05 |
| 4 | NUTF2 | 0.14 | 0.082 | 0.489 | 0.388 | 0.604 | 2.3E-07 |
| 4 | NXT1 | 0.21 | 0.12 | 0.943 | 0.632 | 0.493 | 2.7E-11 |
| 4 | OCIAD1 | 0.443 | 0.332 | 2.211 | 1.968 | 0.168 | 2.3E-05 |
| 4 | OCIAD2 | 0.293 | 0.183 | 1.299 | 0.974 | 0.414 | 6.7E-11 |
| 4 | OGDH | 0.141 | 0.081 | 0.449 | 0.409 | 0.426 | 1.2E-07 |
| 4 | OGFRL1 | 0.137 | 0.062 | 0.416 | 0.303 | 0.768 | 2.1E-14 |
| 4 | OIP4-AS1 | 0.239 | 0.167 | 0.99 | 0.874 | 0.178 | 4.8E-05 |
| 4 | OOEP | 0.441 | 0.309 | 3.034 | 1.811 | 0.744 | 1.5E-31 |
| 4 | ORAI2 | 0.167 | 0.063 | 0.724 | 0.319 | 1.184 | 3.4E-26 |
| 4 | OST4 | 0.494 | 0.366 | 2.464 | 2.149 | 0.248 | 4.8E-07 |
| 4 | OSTC | 0.26 | 0.161 | 1.043 | 0.818 | 0.364 | 9.9E-10 |
| 4 | OTUB1 | 0.187 | 0.136 | 0.714 | 0.667 | 0.1 | 0.00124 |
| 4 | OTULINL | 0.246 | 0.172 | 1.264 | 0.984 | 0.36 | 3E-05 |
| 4 | OXNAD1 | 0.147 | 0.1 | 0.63 | 0.427 | 0.246 | 0.00001 |
| 4 | OXR1 | 0.213 | 0.124 | 0.876 | 0.664 | 0.398 | 8.8E-10 |
| 4 | PA2G4 | 0.449 | 0.334 | 2.304 | 1.948 | 0.234 | 4.4E-06 |
| 4 | PABPC1 | 0.871 | 0.681 | 11.29 | 6.444 | 0.784 | 2.7E-48 |
| 4 | PABPC4 | 0.127 | 0.082 | 0.434 | 0.416 | 0.349 | 7.3E-05 |
| 4 | PACSIN2 | 0.194 | 0.126 | 0.841 | 0.644 | 0.361 | 1.3E-06 |
| 4 | PATJ | 0.183 | 0.046 | 0.882 | 0.234 | 1.91 | 1.3E-46 |
| 4 | PBXIP1 | 0.186 | 0.117 | 0.901 | 0.618 | 0.443 | 2.2E-07 |
| 4 | PCIF1 | 0.14 | 0.104 | 0.644 | 0.436 | 0.286 | 0.00041 |
| 4 | PCMT1 | 0.203 | 0.147 | 0.876 | 0.743 | 0.237 | 0.00023 |
| 4 | PCMTD1 | 0.413 | 0.301 | 2.248 | 1.799 | 0.321 | 3.1E-07 |
| 4 | PCNX2 | 0.197 | 0.098 | 0.793 | 0.424 | 0.494 | 1.8E-14 |
| 4 | PDCD4 | 0.609 | 0.482 | 4.182 | 3.607 | 0.213 | 8E-07 |
| 4 | PDCD4 | 0.19 | 0.129 | 0.76 | 0.644 | 0.216 | 4.9E-05 |
| 4 | PDCD6IP | 0.377 | 0.289 | 1.833 | 1.684 | 0.121 | 0.00027 |
| 4 | PDE4B | 0.229 | 0.047 | 1.162 | 0.243 | 2.202 | 6.4E-93 |
| 4 | PDE4DIP | 0.19 | 0.092 | 0.761 | 0.468 | 0.702 | 4.5E-16 |
| 4 | PDE7B | 0.229 | 0.043 | 2.346 | 0.236 | 3.312 | 4E-109 |
| 4 | PDE8A | 0.19 | 0.112 | 0.882 | 0.616 | 0.416 | 3.1E-09 |
| 4 | PEBP1 | 0.227 | 0.147 | 0.914 | 0.73 | 0.324 | 2.4E-07 |
| 4 | PECAM1 | 0.107 | 0.069 | 0.466 | 0.373 | 0.322 | 0.00021 |
| 4 | PER1 | 0.12 | 0.074 | 0.4 | 0.413 | 0.277 | 3.1E-05 |

| | | | | | | | |
|---|---|---|---|---|---|---|---|
| 4 | PEX2 | 0.171 | 0.119 | 0.671 | 0.496 | 0.17 | 0.00028 |
| 4 | PFDN2 | 0.241 | 0.147 | 1.01 | 0.806 | 0.326 | 2.6E-07 |
| 4 | PFDN4 | 0.67 | 0.491 | 4.477 | 3.378 | 0.406 | 2.4E-14 |
| 4 | PFDN4 | 0.13 | 0.081 | 0.496 | 0.391 | 0.344 | 1.6E-05 |
| 4 | PGK1 | 0.433 | 0.29 | 2.14 | 1.673 | 0.363 | 8.9E-12 |
| 4 | PGLS | 0.23 | 0.146 | 1.002 | 0.739 | 0.44 | 1.4E-08 |
| 4 | PGM2 | 0.109 | 0.044 | 0.474 | 0.249 | 0.87 | 1.4E-09 |
| 4 | PGRMC2 | 0.121 | 0.088 | 0.413 | 0.448 | 0.196 | 0.00421 |
| 4 | PHACTR2 | 0.281 | 0.184 | 1.346 | 1.097 | 0.304 | 1E-08 |
| 4 | PHB | 0.103 | 0.047 | 0.347 | 0.263 | 0.402 | 1.9E-06 |
| 4 | PHB2 | 0.181 | 0.117 | 0.671 | 0.473 | 0.228 | 4.4E-06 |
| 4 | PHF4A | 0.143 | 0.093 | 0.486 | 0.463 | 0.34 | 1E-06 |
| 4 | PHTF1 | 0.143 | 0.107 | 0.492 | 0.443 | 0.123 | 0.0079 |
| 4 | PIK3CA | 0.266 | 0.191 | 1.168 | 1.07 | 0.126 | 0.00011 |
| 4 | PIKFYVE | 0.181 | 0.136 | 0.782 | 0.712 | 0.134 | 0.00283 |
| 4 | PIM3 | 0.161 | 0.04 | 0.747 | 0.24 | 1.636 | 1.4E-34 |
| 4 | PKM | 0.201 | 0.147 | 0.894 | 0.78 | 0.199 | 0.00044 |
| 4 | PLA2G6 | 0.12 | 0.031 | 0.444 | 0.146 | 1.413 | 6.9E-34 |
| 4 | PLCL2 | 0.434 | 0.298 | 2.287 | 1.841 | 0.304 | 2.1E-10 |
| 4 | PLEKHA3 | 0.126 | 0.088 | 0.468 | 0.431 | 0.118 | 0.00222 |
| 4 | PLEKHA4 | 0.107 | 0.044 | 0.418 | 0.304 | 0.444 | 3.4E-08 |
| 4 | PLEKHA7 | 0.214 | 0.094 | 0.842 | 0.448 | 0.878 | 7.1E-23 |
| 4 | PLEKHF2 | 0.209 | 0.138 | 0.87 | 0.704 | 0.306 | 2.6E-06 |
| 4 | PLEKHO1 | 0.136 | 0.091 | 0.491 | 0.484 | 0.29 | 0.00028 |
| 4 | PLK2 | 0.134 | 0.037 | 0.749 | 0.216 | 1.813 | 2.3E-36 |
| 4 | PLP2 | 0.306 | 0.102 | 1.616 | 0.441 | 1.48 | 2.1E-62 |
| 4 | PLRG1 | 0.147 | 0.107 | 0.61 | 0.434 | 0.191 | 0.00016 |
| 4 | PLSCR1 | 0.48 | 0.364 | 4.392 | 4.077 | 0.107 | 1.1E-05 |
| 4 | PLXDC2 | 0.124 | 0.047 | 0.703 | 0.273 | 1.368 | 4.9E-20 |
| 4 | PLXNC1 | 0.106 | 0.037 | 0.396 | 0.189 | 1.064 | 3.6E-19 |
| 4 | PMAIP1 | 0.22 | 0.161 | 1.374 | 1.04 | 0.387 | 0.00015 |
| 4 | PNKD | 0.174 | 0.129 | 0.732 | 0.641 | 0.17 | 0.00204 |
| 4 | PNPLA2 | 0.147 | 0.106 | 0.604 | 0.437 | 0.169 | 0.00014 |
| 4 | PNRC1 | 0.876 | 0.823 | 12.733 | 11.198 | 0.184 | 0.00018 |
| 4 | POC1B | 0.176 | 0.12 | 0.712 | 0.632 | 0.172 | 0.00012 |
| 4 | POLG2 | 0.113 | 0.064 | 0.421 | 0.32 | 0.394 | 1.2E-06 |
| 4 | POLK | 0.141 | 0.104 | 0.632 | 0.444 | 0.214 | 0.00029 |
| 4 | POLR2E | 0.19 | 0.126 | 0.736 | 0.618 | 0.243 | 9.3E-06 |
| 4 | POLR2K | 0.22 | 0.131 | 1.037 | 0.682 | 0.604 | 1.9E-10 |
| 4 | POLR2L | 0.41 | 0.333 | 2.691 | 1.923 | 0.484 | 1.6E-17 |
| 4 | POLR3D | 0.107 | 0.064 | 0.401 | 0.316 | 0.341 | 1.9E-05 |
| 4 | PPA1 | 0.101 | 0.042 | 0.394 | 0.199 | 0.988 | 1.5E-13 |
| 4 | PPCS | 0.236 | 0.183 | 1.093 | 0.977 | 0.162 | 0.00228 |
| 4 | PPIA | 0.716 | 0.479 | 4.441 | 4.647 | 0.243 | 1.7E-09 |
| 4 | PPIB | 0.467 | 0.447 | 3.339 | 3.061 | 0.126 | 8.7E-05 |
| 4 | PPIG | 0.313 | 0.214 | 1.431 | 1.203 | 0.24 | 2.1E-07 |
| 4 | PPIL4 | 0.22 | 0.144 | 1 | 0.821 | 0.284 | 3.2E-05 |
| 4 | PPIP4K2 | 0.29 | 0.211 | 1.283 | 1.179 | 0.121 | 9.8E-05 |
| 4 | PPP1CB | 0.349 | 0.23 | 1.814 | 1.291 | 0.491 | 8E-11 |
| 4 | PPP1R10 | 0.407 | 0.28 | 2.204 | 1.76 | 0.324 | 2.2E-09 |
| 4 | PPP1R14A | 0.401 | 0.293 | 6.247 | 2.804 | 1.144 | 1.4E-38 |
| 4 | PPP1R14B | 0.219 | 0.128 | 0.983 | 0.687 | 0.416 | 4.4E-11 |
| 4 | PPP1R14B | 0.121 | 0.043 | 0.634 | 0.316 | 1.008 | 3.8E-14 |
| 4 | PPP1R2 | 0.43 | 0.322 | 2.431 | 1.941 | 0.324 | 1.5E-06 |

| | | | | | | | |
|---|---|---|---|---|---|---|---|
| 4 | PPP1R9A | 0.209 | 0.013 | 1.042 | 0.073 | 3.847 | 7E-270 |
| 4 | PPP2CA | 0.417 | 0.319 | 2.083 | 1.843 | 0.169 | 7.4E-05 |
| 4 | PPP3CA | 0.481 | 0.436 | 3.694 | 3.069 | 0.268 | 6.9E-09 |
| 4 | PPT1 | 0.11 | 0.064 | 0.433 | 0.31 | 0.483 | 7.5E-06 |
| 4 | PRELID1 | 0.287 | 0.209 | 1.224 | 1.108 | 0.144 | 8E-05 |
| 4 | PRELID3B | 0.12 | 0.071 | 0.464 | 0.346 | 0.423 | 4.8E-06 |
| 4 | PRKRA | 0.129 | 0.073 | 0.429 | 0.367 | 0.424 | 1.8E-07 |
| 4 | PRMT1 | 0.183 | 0.109 | 0.744 | 0.431 | 0.487 | 1.4E-08 |
| 4 | PRNP | 0.134 | 0.099 | 0.494 | 0.413 | 0.214 | 0.00475 |
| 4 | PRPF38A | 0.149 | 0.096 | 0.46 | 0.476 | 0.234 | 3.1E-05 |
| 4 | PRPF6 | 0.309 | 0.184 | 1.397 | 0.993 | 0.493 | 3.4E-13 |
| 4 | PRPS1 | 0.12 | 0.067 | 0.464 | 0.329 | 0.497 | 4.4E-07 |
| 4 | PRXL2C | 0.126 | 0.089 | 0.404 | 0.441 | 0.164 | 0.00241 |
| 4 | PSD4 | 0.101 | 0.069 | 0.387 | 0.342 | 0.134 | 0.00189 |
| 4 | PSIP1 | 0.374 | 0.277 | 1.78 | 1.497 | 0.147 | 1.3E-05 |
| 4 | PSMA1 | 0.306 | 0.224 | 1.402 | 1.267 | 0.146 | 6.6E-05 |
| 4 | PSMA3 | 0.206 | 0.144 | 0.901 | 0.81 | 0.143 | 0.0017 |
| 4 | PSMB1 | 0.499 | 0.363 | 2.441 | 2.136 | 0.241 | 9.4E-08 |
| 4 | PSMB4 | 0.176 | 0.123 | 0.731 | 0.491 | 0.306 | 0.0002 |
| 4 | PSMD11 | 0.123 | 0.084 | 0.478 | 0.422 | 0.181 | 0.00122 |
| 4 | PSMD13 | 0.16 | 0.106 | 0.486 | 0.42 | 0.171 | 7.5E-05 |
| 4 | PSMD14 | 0.164 | 0.114 | 0.644 | 0.486 | 0.147 | 0.00035 |
| 4 | PSMD6 | 0.22 | 0.166 | 0.937 | 0.863 | 0.118 | 0.00215 |
| 4 | PSTPIP1 | 0.106 | 0.061 | 0.407 | 0.3 | 0.44 | 4.3E-06 |
| 4 | PTCH2 | 0.174 | 0.079 | 1.412 | 0.499 | 1.337 | 3.2E-19 |
| 4 | PTGDR | 0.249 | 0.172 | 1.188 | 0.948 | 0.327 | 3.4E-06 |
| 4 | PTGES3 | 0.443 | 0.348 | 2.344 | 2.042 | 0.199 | 2.4E-05 |
| 4 | PTK2 | 0.121 | 0.022 | 0.424 | 0.12 | 2.127 | 2.4E-46 |
| 4 | PTMA | 0.986 | 0.942 | 27.476 | 24.264 | 0.126 | 0.00044 |
| 4 | PUF60 | 0.149 | 0.094 | 0.414 | 0.443 | 0.183 | 1.6E-05 |
| 4 | PUM3 | 0.194 | 0.114 | 0.766 | 0.612 | 0.323 | 1E-08 |
| 4 | PWWP3A | 0.169 | 0.104 | 0.668 | 0.432 | 0.329 | 8.2E-07 |
| 4 | PYCR2 | 0.114 | 0.072 | 0.441 | 0.347 | 0.346 | 7.9E-05 |
| 4 | QARS | 0.121 | 0.077 | 0.444 | 0.374 | 0.249 | 6.6E-05 |
| 4 | R3HDM4 | 0.121 | 0.067 | 0.476 | 0.32 | 0.474 | 1.1E-07 |
| 4 | RAB11A | 0.33 | 0.216 | 1.464 | 1.204 | 0.281 | 4.8E-09 |
| 4 | RAB11B | 0.139 | 0.099 | 0.443 | 0.492 | 0.143 | 0.00234 |
| 4 | RAB21 | 0.241 | 0.17 | 1.016 | 0.904 | 0.169 | 2.9E-05 |
| 4 | RAB37 | 0.139 | 0.087 | 0.486 | 0.446 | 0.394 | 1.1E-05 |
| 4 | RAB3GAP1 | 0.244 | 0.116 | 1.142 | 0.641 | 0.834 | 6.9E-24 |
| 4 | RAB4A | 0.261 | 0.189 | 1.101 | 1.024 | 0.102 | 0.00012 |
| 4 | RACK1 | 0.823 | 0.617 | 7.713 | 4.068 | 0.606 | 4.4E-36 |
| 4 | RAD23A | 0.249 | 0.164 | 1.088 | 0.849 | 0.342 | 2E-07 |
| 4 | RALA | 0.301 | 0.213 | 1.442 | 1.3 | 0.14 | 3.3E-06 |
| 4 | RALB | 0.163 | 0.122 | 0.718 | 0.632 | 0.184 | 0.00446 |
| 4 | RALGPS2 | 0.12 | 0.072 | 0.407 | 0.377 | 0.426 | 6.2E-06 |
| 4 | RANBP1 | 0.139 | 0.088 | 0.469 | 0.424 | 0.426 | 1.5E-05 |
| 4 | RANBP2 | 0.413 | 0.291 | 2.114 | 1.763 | 0.262 | 1.2E-08 |
| 4 | RANBP6 | 0.103 | 0.07 | 0.381 | 0.344 | 0.148 | 0.0018 |
| 4 | RAP2C-AS1 | 0.101 | 0.066 | 0.387 | 0.334 | 0.214 | 0.00061 |
| 4 | RAPGEF2 | 0.17 | 0.121 | 0.747 | 0.69 | 0.134 | 0.00048 |
| 4 | RAPGEF6 | 0.33 | 0.227 | 1.476 | 1.31 | 0.172 | 7.8E-07 |
| 4 | RARA | 0.189 | 0.074 | 0.827 | 0.374 | 1.143 | 3.5E-27 |
| 4 | RASGEF1B | 0.277 | 0.181 | 1.312 | 1.03 | 0.349 | 6.1E-09 |

| | | | | | | | |
|---|---|---|---|---|---|---|---|
| 4 | RB1 | 0.349 | 0.26 | 1.649 | 1.443 | 0.104 | 9.3E-05 |
| 4 | RBFOX2 | 0.277 | 0.14 | 1.18 | 0.772 | 0.612 | 4.4E-17 |
| 4 | RBIS | 0.279 | 0.211 | 1.282 | 1.133 | 0.178 | 0.00042 |
| 4 | RBM19 | 0.116 | 0.079 | 0.479 | 0.4 | 0.26 | 0.00107 |
| 4 | RBM3 | 0.306 | 0.194 | 1.37 | 1.063 | 0.366 | 3.3E-10 |
| 4 | RBM8A | 0.419 | 0.418 | 2.974 | 2.682 | 0.14 | 0.00012 |
| 4 | RBMS1 | 0.34 | 0.242 | 1.761 | 1.402 | 0.329 | 1.7E-08 |
| 4 | RBMX | 0.244 | 0.184 | 1.144 | 0.992 | 0.207 | 0.00014 |
| 4 | RBPJ | 0.299 | 0.231 | 1.481 | 1.397 | 0.179 | 0.00048 |
| 4 | RCAN3 | 0.18 | 0.016 | 0.87 | 0.077 | 3.499 | 1E-178 |
| 4 | RCBTB2 | 0.176 | 0.094 | 0.843 | 0.494 | 0.767 | 6.2E-12 |
| 4 | RCSD1 | 0.416 | 0.281 | 2.199 | 1.646 | 0.418 | 6.3E-11 |
| 4 | RDX | 0.116 | 0.074 | 0.474 | 0.384 | 0.301 | 0.00019 |
| 4 | REEP3 | 0.116 | 0.066 | 0.496 | 0.32 | 0.896 | 4.1E-07 |
| 4 | REL | 0.46 | 0.441 | 3.478 | 3.08 | 0.216 | 4.3E-06 |
| 4 | RELL1 | 0.246 | 0.088 | 1.042 | 0.462 | 1.187 | 1.5E-41 |
| 4 | REV3L | 0.234 | 0.164 | 1.023 | 0.917 | 0.147 | 3E-05 |
| 4 | RGCC | 0.341 | 0.223 | 2.48 | 1.469 | 0.812 | 3.2E-12 |
| 4 | RGS10 | 0.143 | 0.083 | 0.643 | 0.434 | 0.486 | 9.5E-10 |
| 4 | RGS19 | 0.11 | 0.08 | 0.431 | 0.394 | 0.124 | 0.009 |
| 4 | RHBDD1 | 0.103 | 0.072 | 0.408 | 0.374 | 0.124 | 0.00378 |
| 4 | RHOB | 0.109 | 0.047 | 0.427 | 0.246 | 1.094 | 4.3E-13 |
| 4 | RHOH | 0.341 | 0.227 | 1.944 | 1.338 | 0.44 | 1.4E-11 |
| 4 | RIN3 | 0.387 | 0.228 | 1.944 | 1.39 | 0.483 | 2.8E-18 |
| 4 | RIOK3 | 0.401 | 0.318 | 2.089 | 1.912 | 0.127 | 0.00073 |
| 4 | RIPOR2 | 0.681 | 0.44 | 4.242 | 4.221 | 0.314 | 6.9E-10 |
| 4 | RLIM | 0.129 | 0.094 | 0.419 | 0.48 | 0.113 | 0.00628 |
| 4 | RNASEH1 | 0.121 | 0.07 | 0.476 | 0.347 | 0.414 | 1.1E-06 |
| 4 | RNASET2 | 0.176 | 0.11 | 0.767 | 0.463 | 0.447 | 3.7E-07 |
| 4 | RNF130 | 0.261 | 0.102 | 1.21 | 0.46 | 1.111 | 2E-37 |
| 4 | RNF144A | 0.131 | 0.041 | 0.438 | 0.269 | 1.002 | 2.6E-19 |
| 4 | RNF24 | 0.144 | 0.072 | 0.606 | 0.373 | 0.702 | 2.2E-14 |
| 4 | RNH1 | 0.149 | 0.097 | 0.608 | 0.484 | 0.331 | 4.2E-05 |
| 4 | RNPS1 | 0.413 | 0.288 | 1.924 | 1.629 | 0.239 | 3.5E-08 |
| 4 | ROMO1 | 0.31 | 0.224 | 1.471 | 1.178 | 0.32 | 4.3E-06 |
| 4 | RPIA | 0.117 | 0.084 | 0.461 | 0.427 | 0.109 | 0.00676 |
| 4 | RPL10 | 1 | 0.969 | 64.249 | 40.244 | 0.697 | 3E-138 |
| 4 | RPL10A | 0.827 | 0.432 | 8.349 | 4.171 | 1.003 | 8.4E-83 |
| 4 | RPL11 | 0.997 | 0.936 | 34.411 | 21.131 | 0.708 | 4E-114 |
| 4 | RPL12 | 0.986 | 0.901 | 24.484 | 16.707 | 0.614 | 1.2E-72 |
| 4 | RPL13 | 0.941 | 0.781 | 16.244 | 9.333 | 0.8 | 4.2E-86 |
| 4 | RPL13A | 0.921 | 0.76 | 14.177 | 8.394 | 0.746 | 1.1E-66 |
| 4 | RPL14 | 0.943 | 0.849 | 17.88 | 12.666 | 0.497 | 4.5E-44 |
| 4 | RPL14 | 0.824 | 0.662 | 8.194 | 6.101 | 0.424 | 8.2E-21 |
| 4 | RPL17 | 0.847 | 0.67 | 8.969 | 6.163 | 0.441 | 4.4E-33 |
| 4 | RPL18 | 0.993 | 0.907 | 28.736 | 17.314 | 0.731 | 2E-109 |
| 4 | RPL18A | 0.911 | 0.779 | 12.701 | 9.087 | 0.483 | 3.2E-29 |
| 4 | RPL19 | 0.987 | 0.913 | 27.106 | 17.794 | 0.607 | 1.2E-79 |
| 4 | RPL21 | 0.899 | 0.746 | 10.887 | 8.247 | 0.399 | 1.1E-22 |
| 4 | RPL22 | 0.934 | 0.748 | 13.399 | 8.216 | 0.706 | 2E-47 |
| 4 | RPL22L1 | 0.244 | 0.122 | 1.192 | 0.612 | 0.963 | 1.2E-23 |
| 4 | RPL23 | 0.623 | 0.432 | 3.833 | 2.817 | 0.444 | 4.7E-18 |
| 4 | RPL23A | 0.777 | 0.491 | 6.649 | 4.822 | 0.464 | 3.9E-22 |
| 4 | RPL24 | 0.843 | 0.692 | 9.702 | 6.663 | 0.442 | 2.5E-31 |

| | | | | | | | |
|---|---|---|---|---|---|---|---|
| 4 | RPL26 | 0.876 | 0.694 | 9.846 | 6.744 | 0.444 | 1.4E-36 |
| 4 | RPL27 | 0.66 | 0.472 | 4.412 | 3.221 | 0.486 | 3E-20 |
| 4 | RPL27A | 0.493 | 0.413 | 3.73 | 2.682 | 0.476 | 1.4E-16 |
| 4 | RPL28 | 0.994 | 0.921 | 32.063 | 19.418 | 0.724 | 8E-114 |
| 4 | RPL29 | 0.92 | 0.78 | 14.184 | 9.132 | 0.634 | 8.9E-44 |
| 4 | RPL3 | 0.74 | 0.434 | 6.341 | 4.034 | 0.642 | 4.2E-31 |
| 4 | RPL30 | 0.991 | 0.934 | 30.499 | 21.8 | 0.484 | 1.4E-46 |
| 4 | RPL31 | 0.837 | 0.642 | 8.227 | 4.714 | 0.426 | 1.6E-29 |
| 4 | RPL32 | 0.971 | 0.88 | 24.334 | 14.276 | 0.769 | 7E-106 |
| 4 | RPL34 | 0.99 | 0.904 | 28.011 | 17.444 | 0.674 | 8.5E-90 |
| 4 | RPL34 | 0.86 | 0.67 | 9.709 | 6.144 | 0.648 | 4.6E-41 |
| 4 | RPL34A | 0.973 | 0.868 | 20.962 | 13.648 | 0.618 | 8.8E-68 |
| 4 | RPL36 | 0.887 | 0.706 | 10.462 | 6.789 | 0.624 | 1.3E-44 |
| 4 | RPL36A | 0.844 | 0.631 | 9.483 | 4.498 | 0.802 | 6.1E-60 |
| 4 | RPL37 | 0.963 | 0.862 | 19.67 | 13.414 | 0.442 | 9.9E-42 |
| 4 | RPL37A | 0.847 | 0.686 | 8.743 | 6.409 | 0.427 | 2.4E-21 |
| 4 | RPL38 | 0.84 | 0.644 | 8.496 | 4.947 | 0.431 | 2.1E-30 |
| 4 | RPL39 | 0.979 | 0.826 | 17.942 | 11.326 | 0.664 | 3.4E-68 |
| 4 | RPL4 | 0.96 | 0.833 | 17.6 | 11.894 | 0.464 | 3.9E-44 |
| 4 | RPL4 | 0.47 | 0.347 | 3.341 | 2.112 | 0.666 | 1.5E-29 |
| 4 | RPL41 | 0.993 | 0.943 | 43.624 | 28.44 | 0.612 | 1E-101 |
| 4 | RPL6 | 0.947 | 0.86 | 17.867 | 12.997 | 0.449 | 1.1E-32 |
| 4 | RPL7 | 0.764 | 0.49 | 6.66 | 4.648 | 0.416 | 4E-22 |
| 4 | RPL7A | 0.986 | 0.903 | 23.433 | 16.234 | 0.436 | 2.1E-62 |
| 4 | RPL7L1 | 0.2 | 0.126 | 0.782 | 0.642 | 0.286 | 4.1E-07 |
| 4 | RPL8 | 0.981 | 0.826 | 21.131 | 10.601 | 0.994 | 4E-137 |
| 4 | RPL9 | 0.803 | 0.442 | 7.049 | 4.217 | 0.741 | 4.2E-41 |
| 4 | RPLP0 | 0.937 | 0.76 | 14.381 | 8.47 | 0.844 | 8.3E-86 |
| 4 | RPLP1 | 0.999 | 0.949 | 49.449 | 36.267 | 0.713 | 8.6E-96 |
| 4 | RPLP2 | 0.999 | 0.946 | 38.914 | 24.814 | 0.649 | 9E-114 |
| 4 | RPP21 | 0.101 | 0.073 | 0.408 | 0.347 | 0.19 | 0.00848 |
| 4 | RPS11 | 0.719 | 0.48 | 4.304 | 4.48 | 0.244 | 3.4E-08 |
| 4 | RPS12 | 0.996 | 0.943 | 42.967 | 24.986 | 0.782 | 7E-148 |
| 4 | RPS13 | 0.993 | 0.917 | 30.094 | 18.296 | 0.718 | 2E-103 |
| 4 | RPS14 | 0.977 | 0.891 | 24.744 | 16.037 | 0.626 | 2.1E-78 |
| 4 | RPS14 | 0.911 | 0.763 | 11.264 | 8.419 | 0.42 | 1.1E-26 |
| 4 | RPS14A | 0.973 | 0.873 | 23.428 | 14.428 | 0.699 | 1.8E-84 |
| 4 | RPS16 | 0.79 | 0.498 | 6.964 | 4.878 | 0.414 | 1E-27 |
| 4 | RPS17 | 0.416 | 0.361 | 2.882 | 2.16 | 0.416 | 7.5E-12 |
| 4 | RPS18 | 0.873 | 0.661 | 10.788 | 6.068 | 0.83 | 4E-67 |
| 4 | RPS19 | 0.94 | 0.786 | 14.724 | 9.947 | 0.466 | 9.7E-48 |
| 4 | RPS2 | 0.783 | 0.413 | 7.293 | 3.888 | 0.908 | 7.7E-61 |
| 4 | RPS20 | 0.443 | 0.337 | 2.442 | 1.986 | 0.298 | 3.9E-07 |
| 4 | RPS21 | 0.947 | 0.824 | 18.339 | 11.37 | 0.69 | 1.2E-73 |
| 4 | RPS23 | 0.961 | 0.813 | 18.968 | 10.441 | 0.848 | 9E-103 |
| 4 | RPS24 | 0.996 | 0.946 | 40.981 | 27.842 | 0.873 | 5E-184 |
| 4 | RPS24 | 0.767 | 0.429 | 6.164 | 3.924 | 0.642 | 9.6E-36 |
| 4 | RPS27 | 0.984 | 0.906 | 30.044 | 19.98 | 0.489 | 2.5E-69 |
| 4 | RPS27A | 1 | 0.949 | 43.747 | 29.489 | 0.464 | 4.5E-96 |
| 4 | RPS28 | 0.984 | 0.893 | 26.994 | 16.316 | 0.726 | 2.3E-93 |
| 4 | RPS29 | 0.926 | 0.774 | 14.346 | 9.736 | 0.449 | 2.1E-42 |
| 4 | RPS3 | 0.98 | 0.917 | 26.341 | 18.749 | 0.49 | 8.8E-44 |
| 4 | RPS3A | 0.977 | 0.844 | 19.974 | 12.212 | 0.71 | 3.6E-77 |
| 4 | RPS4 | 0.936 | 0.723 | 13.874 | 7.342 | 0.918 | 2E-104 |

| | | | | | | | |
|---|---|---|---|---|---|---|---|
| 4 | RPS4X | 0.939 | 0.793 | 14.768 | 9.421 | 0.728 | 1.8E-69 |
| 4 | RPS6 | 0.929 | 0.772 | 14.008 | 9.141 | 0.714 | 7.1E-67 |
| 4 | RPS6KA2 | 0.113 | 0.02 | 0.442 | 0.101 | 2.164 | 2.4E-46 |
| 4 | RPS6KA3 | 0.386 | 0.294 | 2.074 | 1.788 | 0.214 | 6.6E-05 |
| 4 | RPS6KA4 | 0.293 | 0.173 | 1.276 | 0.986 | 0.371 | 1.1E-12 |
| 4 | RPS7 | 0.986 | 0.898 | 24.204 | 16.493 | 0.612 | 1.3E-64 |
| 4 | RPS8 | 0.993 | 0.944 | 47.44 | 27.192 | 0.803 | 2E-133 |
| 4 | RPS9 | 0.479 | 0.372 | 3.623 | 2.321 | 0.642 | 1.3E-24 |
| 4 | RPSA | 0.84 | 0.484 | 9.248 | 4.79 | 0.949 | 8.5E-74 |
| 4 | RRBP1 | 0.337 | 0.207 | 1.988 | 1.178 | 0.744 | 6.6E-15 |
| 4 | RRM2B | 0.117 | 0.069 | 0.47 | 0.346 | 0.444 | 3.7E-06 |
| 4 | RSL1D1 | 0.311 | 0.163 | 1.484 | 0.842 | 0.802 | 1.1E-22 |
| 4 | RSL24D1 | 0.289 | 0.216 | 1.347 | 1.176 | 0.196 | 0.00014 |
| 4 | RSRC2 | 0.339 | 0.247 | 1.667 | 1.468 | 0.183 | 0.00012 |
| 4 | RTCB | 0.104 | 0.074 | 0.418 | 0.344 | 0.236 | 0.00469 |
| 4 | RTF2 | 0.203 | 0.137 | 0.841 | 0.704 | 0.272 | 1.9E-05 |
| 4 | RTRAF | 0.241 | 0.168 | 0.977 | 0.842 | 0.198 | 2.7E-05 |
| 4 | RUFY3 | 0.143 | 0.073 | 0.496 | 0.374 | 0.673 | 9.7E-11 |
| 4 | RUNX2 | 0.216 | 0.03 | 0.924 | 0.144 | 2.483 | 4E-138 |
| 4 | RUNX3 | 0.404 | 0.293 | 2.942 | 1.862 | 0.66 | 3.1E-27 |
| 4 | RWDD1 | 0.344 | 0.244 | 1.623 | 1.411 | 0.202 | 4.2E-06 |
| 4 | S100A10 | 0.679 | 0.461 | 4.434 | 3.273 | 0.731 | 8.9E-31 |
| 4 | S100A11 | 0.413 | 0.276 | 3.067 | 1.644 | 0.891 | 8.6E-17 |
| 4 | S1PR1 | 0.146 | 0.107 | 0.639 | 0.449 | 0.193 | 0.00021 |
| 4 | SACM1L | 0.334 | 0.242 | 1.471 | 1.374 | 0.194 | 4.4E-06 |
| 4 | SAE1 | 0.109 | 0.071 | 0.418 | 0.364 | 0.194 | 0.00041 |
| 4 | SAR1B | 0.167 | 0.111 | 0.71 | 0.44 | 0.37 | 2.2E-05 |
| 4 | SATB1-AS1 | 0.221 | 0.138 | 1.167 | 0.797 | 0.449 | 2.9E-09 |
| 4 | SBDS | 0.323 | 0.243 | 1.768 | 1.412 | 0.324 | 1.7E-05 |
| 4 | SCAMP2 | 0.143 | 0.111 | 0.634 | 0.444 | 0.224 | 0.00212 |
| 4 | SCFD2 | 0.141 | 0.084 | 0.489 | 0.484 | 0.28 | 2.6E-08 |
| 4 | SCMH1 | 0.101 | 0.042 | 0.424 | 0.229 | 0.891 | 4.5E-13 |
| 4 | SCOC | 0.111 | 0.079 | 0.416 | 0.383 | 0.117 | 0.00416 |
| 4 | SCRN1 | 0.116 | 0.061 | 0.471 | 0.31 | 0.604 | 2.2E-08 |
| 4 | SCYL3 | 0.104 | 0.044 | 0.398 | 0.266 | 0.483 | 1.2E-07 |
| 4 | SDCCAG8 | 0.146 | 0.084 | 0.674 | 0.432 | 0.644 | 2.4E-10 |
| 4 | SDHB | 0.211 | 0.144 | 0.824 | 0.741 | 0.143 | 3E-05 |
| 4 | SDHD | 0.27 | 0.183 | 1.103 | 0.934 | 0.24 | 1.8E-06 |
| 4 | SEC11A | 0.391 | 0.241 | 2.104 | 1.322 | 0.671 | 9.3E-18 |
| 4 | SEC14L1 | 0.321 | 0.197 | 1.444 | 1.044 | 0.442 | 1.2E-13 |
| 4 | SEC61B | 0.467 | 0.376 | 2.708 | 2.271 | 0.244 | 0.00014 |
| 4 | SEC61G | 0.367 | 0.293 | 1.944 | 1.608 | 0.274 | 0.00045 |
| 4 | SELENOH | 0.379 | 0.234 | 1.769 | 1.239 | 0.413 | 2.5E-14 |
| 4 | SELENOS | 0.289 | 0.17 | 1.417 | 0.878 | 0.69 | 3.4E-14 |
| 4 | SELL | 0.741 | 0.114 | 8.646 | 0.778 | 3.476 | 0 |
| 4 | SEPTIN1 | 0.287 | 0.204 | 1.26 | 1.11 | 0.182 | 4E-05 |
| 4 | SEPTIN11 | 0.289 | 0.123 | 1.29 | 0.638 | 1.014 | 1E-34 |
| 4 | SEPTIN6 | 0.447 | 0.394 | 3.324 | 2.472 | 0.37 | 6.8E-12 |
| 4 | SERBP1 | 0.473 | 0.436 | 3.414 | 2.933 | 0.219 | 7.2E-08 |
| 4 | SERP1 | 0.466 | 0.328 | 2.427 | 1.902 | 0.341 | 2.6E-10 |
| 4 | SERPINB9 | 0.18 | 0.114 | 0.762 | 0.499 | 0.347 | 1.2E-06 |
| 4 | SERTAD2 | 0.166 | 0.098 | 0.646 | 0.411 | 0.339 | 7.4E-08 |
| 4 | SESN1 | 0.293 | 0.203 | 1.349 | 1.14 | 0.231 | 1.5E-06 |
| 4 | SET | 0.701 | 0.6 | 4.263 | 4.807 | 0.131 | 0.00044 |

| | | | | | | | |
|---|---|---|---|---|---|---|---|
| 4 | SETDB1 | 0.12 | 0.076 | 0.469 | 0.376 | 0.317 | 6.4E-05 |
| 4 | SF1 | 0.479 | 0.428 | 3.497 | 2.794 | 0.323 | 2.9E-09 |
| 4 | SF3A2 | 0.149 | 0.091 | 0.472 | 0.442 | 0.371 | 1.8E-06 |
| 4 | SF3B4 | 0.24 | 0.172 | 0.972 | 0.867 | 0.164 | 0.00017 |
| 4 | SF3B4 | 0.209 | 0.141 | 0.764 | 0.701 | 0.126 | 2.7E-05 |
| 4 | SFPQ | 0.431 | 0.288 | 2.07 | 1.696 | 0.288 | 8.6E-11 |
| 4 | SHLD2 | 0.173 | 0.112 | 0.704 | 0.612 | 0.202 | 8.2E-06 |
| 4 | SIK3 | 0.441 | 0.341 | 3.001 | 2.314 | 0.374 | 1.3E-07 |
| 4 | SINHCAF | 0.227 | 0.104 | 0.932 | 0.443 | 0.778 | 1.5E-21 |
| 4 | SIPA1L1 | 0.43 | 0.284 | 2.284 | 1.84 | 0.304 | 2E-11 |
| 4 | SIVA1 | 0.149 | 0.104 | 0.607 | 0.414 | 0.238 | 3.7E-05 |
| 4 | SKAP2 | 0.311 | 0.221 | 1.416 | 1.244 | 0.174 | 7.3E-06 |
| 4 | SLC16A3 | 0.127 | 0.067 | 0.489 | 0.334 | 0.44 | 4.2E-09 |
| 4 | SLC24A24 | 0.12 | 0.082 | 0.434 | 0.418 | 0.344 | 0.00081 |
| 4 | SLC24A3 | 0.381 | 0.272 | 1.737 | 1.424 | 0.188 | 1.3E-06 |
| 4 | SLC24A36 | 0.294 | 0.188 | 1.392 | 1.02 | 0.449 | 2.8E-10 |
| 4 | SLC24A39 | 0.126 | 0.048 | 0.474 | 0.273 | 0.794 | 1.3E-12 |
| 4 | SLC24A4 | 0.39 | 0.28 | 2.014 | 1.622 | 0.313 | 9.4E-08 |
| 4 | SLC24A6 | 0.449 | 0.388 | 3.306 | 2.448 | 0.428 | 4E-13 |
| 4 | SLC30A9 | 0.18 | 0.13 | 0.747 | 0.679 | 0.138 | 0.00112 |
| 4 | SLC34D1 | 0.17 | 0.106 | 0.764 | 0.47 | 0.423 | 4.4E-07 |
| 4 | SLC34F1 | 0.233 | 0.123 | 0.981 | 0.611 | 0.684 | 7.7E-16 |
| 4 | SLC34F4 | 0.133 | 0.098 | 0.494 | 0.411 | 0.219 | 0.00423 |
| 4 | SLC38A1 | 0.683 | 0.439 | 4.67 | 4.279 | 0.126 | 2.2E-06 |
| 4 | SLC38A2 | 0.469 | 0.343 | 2.637 | 2.273 | 0.214 | 7.1E-06 |
| 4 | SLC39A10 | 0.307 | 0.161 | 1.476 | 0.878 | 0.843 | 1.8E-22 |
| 4 | SLC39A8 | 0.123 | 0.06 | 0.481 | 0.303 | 0.667 | 9.6E-11 |
| 4 | SLC40A1 | 0.127 | 0.092 | 0.434 | 0.446 | 0.249 | 0.0038 |
| 4 | SLC44A1 | 0.133 | 0.039 | 0.463 | 0.196 | 1.422 | 4E-32 |
| 4 | SLC7A6 | 0.13 | 0.041 | 0.421 | 0.203 | 1.36 | 2E-27 |
| 4 | SLFN13 | 0.26 | 0.197 | 1.197 | 1.11 | 0.109 | 0.00144 |
| 4 | SMAD3 | 0.146 | 0.1 | 0.494 | 0.448 | 0.118 | 0.0006 |
| 4 | SMAP2 | 0.481 | 0.284 | 2.867 | 1.682 | 0.77 | 2.2E-27 |
| 4 | SMARCA4 | 0.119 | 0.083 | 0.492 | 0.424 | 0.214 | 0.00247 |
| 4 | SMARCC1 | 0.339 | 0.249 | 1.649 | 1.416 | 0.121 | 0.00025 |
| 4 | SMC4 | 0.174 | 0.131 | 0.776 | 0.714 | 0.12 | 0.00467 |
| 4 | SMDT1 | 0.374 | 0.208 | 1.746 | 1.119 | 0.64 | 3.8E-21 |
| 4 | SMIM14 | 0.119 | 0.084 | 0.467 | 0.413 | 0.174 | 0.00317 |
| 4 | SMS | 0.176 | 0.104 | 0.734 | 0.43 | 0.473 | 1.7E-08 |
| 4 | SNAP29 | 0.276 | 0.2 | 1.244 | 1.048 | 0.247 | 4.3E-05 |
| 4 | SNF8 | 0.12 | 0.083 | 0.442 | 0.403 | 0.164 | 0.00178 |
| 4 | SNHG14 | 0.144 | 0.104 | 0.633 | 0.44 | 0.202 | 0.00018 |
| 4 | SNHG14 | 0.143 | 0.08 | 0.63 | 0.418 | 0.493 | 1.1E-08 |
| 4 | SNHG16 | 0.13 | 0.046 | 0.41 | 0.218 | 1.224 | 1.3E-22 |
| 4 | SNHG24 | 0.16 | 0.088 | 0.668 | 0.447 | 0.448 | 6.2E-10 |
| 4 | SNHG29 | 0.447 | 0.293 | 3.123 | 1.714 | 0.866 | 2.8E-42 |
| 4 | SNHG32 | 0.111 | 0.068 | 0.441 | 0.324 | 0.444 | 2.4E-05 |
| 4 | SNHG4 | 0.214 | 0.124 | 0.847 | 0.664 | 0.342 | 2.1E-10 |
| 4 | SNHG8 | 0.306 | 0.116 | 1.432 | 0.496 | 1.266 | 1.3E-47 |
| 4 | SNHG9 | 0.183 | 0.131 | 0.804 | 0.669 | 0.268 | 0.00034 |
| 4 | SNRPA | 0.13 | 0.09 | 0.493 | 0.442 | 0.147 | 0.00125 |
| 4 | SNRPC | 0.203 | 0.141 | 0.862 | 0.769 | 0.164 | 0.00145 |
| 4 | SNRPD2 | 0.613 | 0.464 | 3.664 | 3.074 | 0.243 | 9.1E-08 |
| 4 | SNRPF | 0.183 | 0.097 | 0.729 | 0.448 | 0.669 | 2.7E-12 |

| | | | | | | | |
|---|---|---|---|---|---|---|---|
| 4 | SNU13 | 0.24 | 0.174 | 1.036 | 0.919 | 0.173 | 1.9E-05 |
| 4 | SNX10 | 0.187 | 0.094 | 0.739 | 0.482 | 0.617 | 9.8E-14 |
| 4 | SOCS1 | 0.209 | 0.084 | 1.067 | 0.434 | 1.298 | 1.3E-28 |
| 4 | SOD2 | 0.204 | 0.141 | 0.848 | 0.792 | 0.1 | 0.00146 |
| 4 | SOX4 | 0.231 | 0.044 | 1.739 | 0.244 | 2.776 | 8E-106 |
| 4 | SPAG1 | 0.147 | 0.076 | 0.619 | 0.396 | 0.646 | 1.4E-13 |
| 4 | SPATS2L | 0.126 | 0.08 | 0.444 | 0.442 | 0.331 | 4.4E-05 |
| 4 | SPCS1 | 0.44 | 0.347 | 2.229 | 1.997 | 0.149 | 0.00113 |
| 4 | SPECC1 | 0.141 | 0.037 | 0.668 | 0.187 | 1.838 | 1.5E-48 |
| 4 | SPG21 | 0.124 | 0.089 | 0.474 | 0.439 | 0.112 | 0.00478 |
| 4 | SPINK2 | 0.2 | 0.001 | 1.006 | 0.008 | 7.043 | 0 |
| 4 | SPRY1 | 0.107 | 0.01 | 0.464 | 0.044 | 3.379 | 9E-103 |
| 4 | SPTBN1 | 0.4 | 0.181 | 2.074 | 1.029 | 1.012 | 4E-43 |
| 4 | SPTLC2 | 0.349 | 0.229 | 1.649 | 1.278 | 0.376 | 9.4E-11 |
| 4 | SPTSSB | 0.181 | 0.002 | 0.864 | 0.012 | 6.144 | 0 |
| 4 | SRFBP1 | 0.114 | 0.064 | 0.443 | 0.329 | 0.449 | 7.8E-07 |
| 4 | SRGAP3 | 0.17 | 0.016 | 0.87 | 0.08 | 3.434 | 2E-148 |
| 4 | SRP44 | 0.24 | 0.174 | 1.01 | 0.942 | 0.101 | 0.00035 |
| 4 | SRP72 | 0.34 | 0.237 | 1.449 | 1.307 | 0.244 | 6.8E-08 |
| 4 | SRRM1 | 0.49 | 0.44 | 3.38 | 3.038 | 0.144 | 7.9E-06 |
| 4 | SRSF10 | 0.47 | 0.447 | 3.326 | 2.934 | 0.181 | 2.8E-05 |
| 4 | SRSF3 | 0.441 | 0.319 | 2.479 | 1.987 | 0.319 | 1.4E-09 |
| 4 | SRSF4 | 0.647 | 0.42 | 4.373 | 3.897 | 0.166 | 7.5E-06 |
| 4 | SRSF6 | 0.12 | 0.069 | 0.478 | 0.343 | 0.478 | 9.4E-07 |
| 4 | SSBP2 | 0.471 | 0.096 | 3.082 | 0.43 | 2.441 | 3E-214 |
| 4 | SSR2 | 0.393 | 0.293 | 1.804 | 1.648 | 0.121 | 6.1E-05 |
| 4 | SSR3 | 0.28 | 0.194 | 1.32 | 1.029 | 0.36 | 7.3E-07 |
| 4 | SSR4 | 0.42 | 0.342 | 2.909 | 2.098 | 0.472 | 6E-14 |
| 4 | ST13 | 0.461 | 0.303 | 2.43 | 1.748 | 0.467 | 8.3E-14 |
| 4 | ST3GAL1 | 0.207 | 0.09 | 0.934 | 0.476 | 0.973 | 2.4E-23 |
| 4 | STAM | 0.299 | 0.134 | 1.379 | 0.718 | 0.941 | 4.1E-31 |
| 4 | STAMBPL1 | 0.144 | 0.074 | 0.624 | 0.384 | 0.704 | 1.4E-13 |
| 4 | STAP1 | 0.119 | 0.03 | 0.41 | 0.144 | 1.411 | 1.8E-34 |
| 4 | STARD4 | 0.126 | 0.091 | 0.499 | 0.441 | 0.148 | 0.00462 |
| 4 | STEAP1B | 0.161 | 0.078 | 0.641 | 0.383 | 0.764 | 4.2E-14 |
| 4 | STK17A | 0.817 | 0.712 | 9.044 | 7.432 | 0.284 | 4.4E-11 |
| 4 | STK17B | 0.88 | 0.726 | 10.327 | 7.384 | 0.484 | 3.8E-26 |
| 4 | STN1 | 0.139 | 0.084 | 0.484 | 0.412 | 0.403 | 2.9E-06 |
| 4 | STRBP | 0.144 | 0.062 | 0.624 | 0.33 | 0.921 | 1.4E-20 |
| 4 | STX17 | 0.219 | 0.139 | 0.886 | 0.741 | 0.248 | 2.1E-07 |
| 4 | STX4 | 0.137 | 0.091 | 0.474 | 0.473 | 0.284 | 0.00012 |
| 4 | SUCLG2 | 0.331 | 0.214 | 1.391 | 1.242 | 0.164 | 8.3E-09 |
| 4 | SUN1 | 0.127 | 0.08 | 0.473 | 0.396 | 0.244 | 3.3E-05 |
| 4 | SUPT3H | 0.313 | 0.166 | 1.483 | 0.947 | 0.648 | 1.2E-20 |
| 4 | SVIL | 0.176 | 0.042 | 0.919 | 0.229 | 2.004 | 2.2E-48 |
| 4 | SVIP | 0.141 | 0.094 | 0.66 | 0.478 | 0.467 | 1.7E-06 |
| 4 | SWAP70 | 0.12 | 0.047 | 0.422 | 0.289 | 0.844 | 9.7E-12 |
| 4 | SYNCRIP | 0.367 | 0.244 | 1.678 | 1.371 | 0.292 | 2.3E-09 |
| 4 | SYNRG | 0.447 | 0.327 | 2.61 | 1.947 | 0.414 | 2.4E-09 |
| 4 | SYPL1 | 0.243 | 0.04 | 1.034 | 0.2 | 2.373 | 1E-128 |
| 4 | TAB2 | 0.243 | 0.144 | 1.139 | 0.844 | 0.431 | 4.3E-10 |
| 4 | TAF13 | 0.141 | 0.093 | 0.648 | 0.466 | 0.474 | 1.1E-06 |
| 4 | TAF9B | 0.107 | 0.026 | 0.384 | 0.121 | 1.676 | 1.3E-34 |
| 4 | TAGLN2 | 0.674 | 0.386 | 4.228 | 2.421 | 1.042 | 4.9E-49 |

| | | | | | | | |
|---|---|---|---|---|---|---|---|
| 4 | TALDO1 | 0.209 | 0.146 | 0.829 | 0.73 | 0.182 | 0.0001 |
| 4 | TARS | 0.197 | 0.138 | 0.934 | 0.73 | 0.347 | 6.9E-05 |
| 4 | TARSL2 | 0.223 | 0.13 | 0.927 | 0.687 | 0.432 | 1.8E-10 |
| 4 | TASP1 | 0.139 | 0.1 | 0.484 | 0.424 | 0.148 | 0.00273 |
| 4 | TATDN1 | 0.146 | 0.082 | 0.667 | 0.407 | 0.711 | 4.2E-11 |
| 4 | TC2N | 0.346 | 0.14 | 1.987 | 0.822 | 1.273 | 1.5E-49 |
| 4 | TCERG1 | 0.289 | 0.2 | 1.313 | 1.102 | 0.243 | 1.4E-06 |
| 4 | TCF4 | 0.127 | 0.034 | 0.444 | 0.174 | 1.64 | 7.4E-36 |
| 4 | TCF7 | 0.337 | 0.049 | 1.661 | 0.267 | 2.639 | 5E-208 |
| 4 | TCIRG1 | 0.141 | 0.066 | 0.439 | 0.331 | 0.704 | 1.5E-13 |
| 4 | TCP1 | 0.243 | 0.16 | 1.046 | 0.827 | 0.342 | 1.3E-08 |
| 4 | TCP11L1 | 0.111 | 0.074 | 0.43 | 0.374 | 0.203 | 0.0008 |
| 4 | TEC | 0.236 | 0.024 | 1.111 | 0.131 | 3.082 | 1E-193 |
| 4 | TECR | 0.171 | 0.129 | 0.734 | 0.648 | 0.182 | 0.00462 |
| 4 | TENT4C | 0.209 | 0.143 | 0.943 | 0.841 | 0.163 | 0.0004 |
| 4 | TESPA1 | 0.117 | 0.076 | 0.488 | 0.38 | 0.361 | 0.00014 |
| 4 | TEX10 | 0.109 | 0.068 | 0.419 | 0.342 | 0.241 | 9.1E-05 |
| 4 | TEX14 | 0.204 | 0.12 | 1.403 | 0.946 | 0.668 | 1.7E-10 |
| 4 | TFAM | 0.179 | 0.109 | 0.7 | 0.441 | 0.344 | 1.8E-07 |
| 4 | TFG | 0.199 | 0.134 | 0.806 | 0.674 | 0.246 | 1.6E-05 |
| 4 | THEM4 | 0.181 | 0.061 | 0.792 | 0.299 | 1.404 | 7.1E-34 |
| 4 | THOC7 | 0.249 | 0.161 | 1.049 | 0.837 | 0.327 | 1.2E-07 |
| 4 | TIAM1 | 0.244 | 0.023 | 1.242 | 0.124 | 3.326 | 1E-248 |
| 4 | TIMM13 | 0.104 | 0.044 | 0.361 | 0.244 | 0.41 | 8.8E-08 |
| 4 | TIMP1 | 0.247 | 0.139 | 1.436 | 0.812 | 0.823 | 3E-17 |
| 4 | TKT | 0.123 | 0.081 | 0.464 | 0.397 | 0.227 | 0.00024 |
| 4 | TLK1 | 0.349 | 0.271 | 1.676 | 1.449 | 0.104 | 0.00023 |
| 4 | TMA16 | 0.101 | 0.041 | 0.362 | 0.202 | 0.843 | 7.4E-14 |
| 4 | TMC6 | 0.123 | 0.083 | 0.486 | 0.411 | 0.244 | 0.00067 |
| 4 | TMED10 | 0.417 | 0.387 | 2.864 | 2.422 | 0.242 | 5E-07 |
| 4 | TMEM106B | 0.261 | 0.17 | 1.114 | 0.894 | 0.317 | 4.2E-08 |
| 4 | TMEM123 | 0.77 | 0.37 | 7.442 | 2.743 | 1.434 | 3E-120 |
| 4 | TMEM14B | 0.24 | 0.167 | 1.027 | 0.908 | 0.177 | 2.3E-06 |
| 4 | TMEM14C | 0.169 | 0.033 | 0.744 | 0.167 | 2.142 | 1.1E-73 |
| 4 | TMEM167A | 0.17 | 0.117 | 0.677 | 0.6 | 0.173 | 0.00017 |
| 4 | TMEM230 | 0.236 | 0.179 | 1.023 | 0.934 | 0.129 | 0.00174 |
| 4 | TMEM243 | 0.146 | 0.091 | 0.617 | 0.442 | 0.448 | 2.9E-06 |
| 4 | TMEM248 | 0.313 | 0.236 | 1.392 | 1.228 | 0.181 | 0.00025 |
| 4 | TMEM34B | 0.146 | 0.108 | 0.492 | 0.438 | 0.138 | 0.00465 |
| 4 | TMEM87B | 0.139 | 0.091 | 0.439 | 0.442 | 0.244 | 0.00012 |
| 4 | TMX2 | 0.103 | 0.063 | 0.379 | 0.31 | 0.29 | 6.4E-05 |
| 4 | TNFAIP3 | 0.63 | 0.472 | 6.669 | 4.003 | 0.414 | 0.00064 |
| 4 | TNFAIP8 | 0.364 | 0.234 | 1.946 | 1.371 | 0.412 | 1.7E-12 |
| 4 | TNFRSF11A | 0.12 | 0.003 | 0.434 | 0.017 | 4.672 | 3E-241 |
| 4 | TNFRSF14-AS1 | 0.101 | 0.048 | 0.471 | 0.239 | 0.981 | 4.5E-10 |
| 4 | TNFRSF18 | 0.244 | 0.049 | 1.931 | 0.334 | 2.43 | 1.9E-92 |
| 4 | TNFRSF4 | 0.179 | 0.032 | 1.263 | 0.164 | 2.944 | 1.2E-87 |
| 4 | TNFSF10 | 0.31 | 0.193 | 4.113 | 1.64 | 1.318 | 3.2E-14 |
| 4 | TOB1 | 0.133 | 0.073 | 0.664 | 0.366 | 0.849 | 6.5E-09 |
| 4 | TOMM20 | 0.221 | 0.174 | 1.044 | 0.896 | 0.234 | 0.00739 |
| 4 | TOMM22 | 0.177 | 0.113 | 0.704 | 0.443 | 0.341 | 2.3E-06 |
| 4 | TOMM7 | 0.473 | 0.432 | 3.267 | 2.773 | 0.237 | 1.4E-07 |
| 4 | TOR1AIP2 | 0.166 | 0.12 | 0.7 | 0.616 | 0.186 | 0.0014 |
| 4 | TP43BP1 | 0.173 | 0.113 | 0.716 | 0.494 | 0.268 | 9.3E-06 |

| | | | | | | | |
|---|---|---|---|---|---|---|---|
| 4 | TPI1 | 0.613 | 0.473 | 3.708 | 3.178 | 0.223 | 4.2E-07 |
| 4 | TPK1 | 0.183 | 0.097 | 0.828 | 0.434 | 0.629 | 1.9E-12 |
| 4 | TPM4 | 0.373 | 0.191 | 2.148 | 1.202 | 0.844 | 2.3E-28 |
| 4 | TPST2 | 0.3 | 0.231 | 1.442 | 1.306 | 0.239 | 0.00018 |
| 4 | TPT1 | 1 | 0.948 | 40.402 | 23.838 | 1.08 | 1E-232 |
| 4 | TRA2B | 0.471 | 0.387 | 3.381 | 2.406 | 0.432 | 6.5E-17 |
| 4 | TRABD | 0.161 | 0.104 | 0.631 | 0.44 | 0.226 | 2.4E-05 |
| 4 | TRABD2A | 0.186 | 0.118 | 0.808 | 0.611 | 0.403 | 4.2E-07 |
| 4 | TRAF1 | 0.137 | 0.074 | 0.603 | 0.393 | 0.619 | 4.4E-09 |
| 4 | TRAF4 | 0.376 | 0.136 | 1.777 | 0.728 | 1.287 | 4.2E-64 |
| 4 | TRAPPC8 | 0.241 | 0.177 | 1.039 | 0.963 | 0.109 | 0.00042 |
| 4 | TRDC | 0.417 | 0.308 | 3.442 | 1.921 | 0.841 | 2.4E-29 |
| 4 | TRGC1 | 0.28 | 0.192 | 1.291 | 1.041 | 0.297 | 8.8E-07 |
| 4 | TRGV9 | 0.14 | 0.09 | 0.477 | 0.449 | 0.361 | 3.9E-05 |
| 4 | TRIAP1 | 0.101 | 0.04 | 0.341 | 0.236 | 0.434 | 6.2E-09 |
| 4 | TRIM24 | 0.134 | 0.094 | 0.461 | 0.479 | 0.228 | 0.00142 |
| 4 | TRIM73 | 0.107 | 0.067 | 0.377 | 0.326 | 0.209 | 0.00016 |
| 4 | TRIM8 | 0.116 | 0.082 | 0.414 | 0.408 | 0.337 | 0.0029 |
| 4 | TRIO | 0.139 | 0.064 | 0.613 | 0.348 | 0.774 | 1.9E-13 |
| 4 | TRMO | 0.129 | 0.08 | 0.419 | 0.407 | 0.349 | 2.3E-05 |
| 4 | TRMT10C | 0.104 | 0.072 | 0.411 | 0.34 | 0.274 | 0.00212 |
| 4 | TRMT11 | 0.103 | 0.068 | 0.381 | 0.347 | 0.132 | 0.00133 |
| 4 | TRPS1 | 0.131 | 0.089 | 0.473 | 0.499 | 0.199 | 0.00038 |
| 4 | TSC22D3 | 0.379 | 0.243 | 3.206 | 1.949 | 0.718 | 4.8E-14 |
| 4 | TSPAN2 | 0.146 | 0.094 | 0.64 | 0.468 | 0.442 | 1.4E-05 |
| 4 | TSPO | 0.28 | 0.161 | 1.227 | 0.826 | 0.472 | 4.5E-14 |
| 4 | TSSC4 | 0.103 | 0.074 | 0.389 | 0.344 | 0.134 | 0.00873 |
| 4 | TSTD1 | 0.291 | 0.187 | 1.347 | 0.994 | 0.439 | 8E-10 |
| 4 | TSTD2 | 0.124 | 0.077 | 0.421 | 0.383 | 0.444 | 1.2E-05 |
| 4 | TTC19 | 0.146 | 0.108 | 0.614 | 0.444 | 0.149 | 0.00061 |
| 4 | TTC39C | 0.426 | 0.319 | 2.44 | 2.046 | 0.304 | 8.9E-07 |
| 4 | TTC7A | 0.149 | 0.064 | 0.604 | 0.343 | 0.819 | 3.7E-17 |
| 4 | TTN | 0.191 | 0.023 | 1.127 | 0.124 | 3.167 | 8E-146 |
| 4 | TUBA1A | 0.389 | 0.26 | 2.176 | 1.664 | 0.387 | 2.1E-11 |
| 4 | TUBA1B | 0.293 | 0.184 | 1.482 | 1.003 | 0.463 | 1.3E-11 |
| 4 | TUBA1C | 0.14 | 0.073 | 0.472 | 0.361 | 0.662 | 2.7E-10 |
| 4 | TUBA4A | 0.147 | 0.079 | 0.676 | 0.402 | 0.749 | 1.4E-12 |
| 4 | TUBB | 0.236 | 0.146 | 1.069 | 0.76 | 0.493 | 2.2E-09 |
| 4 | TUBB4B | 0.207 | 0.109 | 0.982 | 0.473 | 0.776 | 1.2E-14 |
| 4 | TUFM | 0.207 | 0.141 | 0.809 | 0.714 | 0.18 | 2.6E-05 |
| 4 | TWISTNB | 0.121 | 0.062 | 0.492 | 0.314 | 0.643 | 1.3E-09 |
| 4 | TXN | 0.303 | 0.168 | 1.429 | 0.846 | 0.837 | 3.1E-18 |
| 4 | TXNDC11 | 0.139 | 0.083 | 0.442 | 0.448 | 0.244 | 1.4E-06 |
| 4 | TXNRD1 | 0.284 | 0.186 | 1.344 | 1.004 | 0.43 | 8.2E-09 |
| 4 | TYMP | 0.149 | 0.099 | 0.604 | 0.427 | 0.198 | 0.00014 |
| 4 | TYW1 | 0.196 | 0.138 | 0.866 | 0.739 | 0.23 | 0.00012 |
| 4 | UBA42 | 0.846 | 0.714 | 8.998 | 6.941 | 0.372 | 1.5E-18 |
| 4 | UBAC2 | 0.334 | 0.206 | 1.626 | 1.163 | 0.484 | 1.1E-13 |
| 4 | UBAP2 | 0.121 | 0.079 | 0.444 | 0.414 | 0.13 | 0.00022 |
| 4 | UBE2D1 | 0.134 | 0.086 | 0.428 | 0.424 | 0.314 | 3.8E-05 |
| 4 | UBE2D3 | 0.461 | 0.47 | 3.412 | 3.198 | 0.134 | 0.00198 |
| 4 | UBE2G2 | 0.244 | 0.181 | 1.103 | 0.944 | 0.208 | 3.8E-05 |
| 4 | UBE2I | 0.303 | 0.217 | 1.284 | 1.144 | 0.143 | 2.8E-05 |
| 4 | UBE2J1 | 0.266 | 0.146 | 1.208 | 0.813 | 0.471 | 4E-13 |

| | | | | | | | |
|---|---|---|---|---|---|---|---|
| 4 | UBE2Q2 | 0.269 | 0.192 | 1.164 | 1.043 | 0.144 | 3.7E-05 |
| 4 | UBE2S | 0.109 | 0.076 | 0.447 | 0.349 | 0.319 | 0.00247 |
| 4 | UBQLN2 | 0.13 | 0.043 | 0.624 | 0.24 | 1.319 | 8E-18 |
| 4 | UBTF | 0.297 | 0.191 | 1.277 | 1.039 | 0.297 | 8.4E-09 |
| 4 | UCHL3 | 0.123 | 0.083 | 0.461 | 0.424 | 0.121 | 0.00068 |
| 4 | UCP2 | 0.344 | 0.269 | 1.846 | 1.621 | 0.188 | 0.00093 |
| 4 | UFM1 | 0.343 | 0.227 | 1.604 | 1.228 | 0.386 | 1.5E-09 |
| 4 | UGCG | 0.107 | 0.066 | 0.476 | 0.331 | 0.424 | 4.4E-05 |
| 4 | UQCRH | 0.397 | 0.269 | 1.824 | 1.494 | 0.288 | 2.8E-09 |
| 4 | UQCRQ | 0.346 | 0.247 | 1.41 | 1.38 | 0.13 | 0.00018 |
| 4 | UROS | 0.121 | 0.066 | 0.449 | 0.314 | 0.416 | 3.9E-08 |
| 4 | USP1 | 0.196 | 0.143 | 0.814 | 0.747 | 0.104 | 0.00093 |
| 4 | USP10 | 0.149 | 0.092 | 0.474 | 0.473 | 0.282 | 8.6E-08 |
| 4 | USP12 | 0.177 | 0.127 | 0.706 | 0.647 | 0.104 | 0.00114 |
| 4 | USP38 | 0.149 | 0.094 | 0.624 | 0.483 | 0.37 | 2.3E-07 |
| 4 | USP43 | 0.144 | 0.086 | 0.64 | 0.444 | 0.44 | 2.6E-09 |
| 4 | USP46 | 0.111 | 0.06 | 0.397 | 0.292 | 0.444 | 2.3E-07 |
| 4 | UST | 0.146 | 0.049 | 0.688 | 0.347 | 0.987 | 1.1E-19 |
| 4 | UTP23 | 0.161 | 0.107 | 0.604 | 0.441 | 0.161 | 6.7E-05 |
| 4 | UXS1 | 0.136 | 0.081 | 0.43 | 0.414 | 0.344 | 1.7E-06 |
| 4 | UXT | 0.297 | 0.204 | 1.32 | 1.063 | 0.313 | 4.2E-07 |
| 4 | VAPA | 0.374 | 0.261 | 1.794 | 1.484 | 0.273 | 3.1E-08 |
| 4 | VAV3 | 0.411 | 0.392 | 3.031 | 2.826 | 0.101 | 3.6E-05 |
| 4 | VDAC1 | 0.293 | 0.206 | 1.284 | 1.07 | 0.263 | 2.4E-06 |
| 4 | VEZF1 | 0.313 | 0.237 | 1.497 | 1.304 | 0.199 | 0.00014 |
| 4 | VEZT | 0.147 | 0.084 | 0.614 | 0.437 | 0.491 | 3.4E-10 |
| 4 | VIM | 0.611 | 0.39 | 6.94 | 3.269 | 1.086 | 7E-39 |
| 4 | VIM-AS1 | 0.143 | 0.093 | 0.494 | 0.469 | 0.339 | 4.5E-05 |
| 4 | VIRMA | 0.273 | 0.194 | 1.118 | 1.036 | 0.11 | 7.1E-05 |
| 4 | VPS29 | 0.247 | 0.179 | 1.126 | 0.942 | 0.247 | 9.9E-06 |
| 4 | VSIR | 0.123 | 0.066 | 0.447 | 0.333 | 0.716 | 1.4E-08 |
| 4 | VTA1 | 0.144 | 0.11 | 0.617 | 0.461 | 0.134 | 0.00119 |
| 4 | VWA8 | 0.181 | 0.124 | 0.823 | 0.701 | 0.232 | 7.6E-05 |
| 4 | WARS | 0.13 | 0.06 | 1.019 | 0.313 | 1.704 | 9.5E-14 |
| 4 | WASHC2A | 0.16 | 0.114 | 0.642 | 0.486 | 0.131 | 0.00112 |
| 4 | WASHC3 | 0.18 | 0.114 | 0.686 | 0.476 | 0.241 | 2.2E-06 |
| 4 | WBP11 | 0.327 | 0.238 | 1.473 | 1.34 | 0.137 | 3.4E-05 |
| 4 | WBP2 | 0.107 | 0.064 | 0.393 | 0.308 | 0.34 | 3.7E-05 |
| 4 | WDFY2 | 0.183 | 0.109 | 0.826 | 0.6 | 0.461 | 1.1E-08 |
| 4 | WDR41 | 0.123 | 0.067 | 0.42 | 0.338 | 0.619 | 4.8E-08 |
| 4 | WDR43 | 0.249 | 0.14 | 1.134 | 0.807 | 0.492 | 1E-12 |
| 4 | WDR74 | 0.233 | 0.141 | 1.237 | 0.92 | 0.427 | 1.1E-09 |
| 4 | WDR83OS | 0.34 | 0.244 | 1.447 | 1.308 | 0.243 | 4.7E-07 |
| 4 | WHAMM | 0.301 | 0.187 | 1.463 | 1.038 | 0.494 | 4.5E-12 |
| 4 | WRNIP1 | 0.101 | 0.071 | 0.372 | 0.347 | 0.102 | 0.00493 |
| 4 | WSB1 | 0.287 | 0.222 | 1.393 | 1.243 | 0.143 | 0.00108 |
| 4 | WTAP | 0.366 | 0.262 | 1.696 | 1.4 | 0.177 | 2.5E-06 |
| 4 | XBP1 | 0.34 | 0.189 | 1.722 | 1.042 | 0.712 | 8.8E-20 |
| 4 | XCL1 | 0.8 | 0.164 | 14.003 | 1.133 | 3.627 | 0 |
| 4 | XCL2 | 0.647 | 0.183 | 7.683 | 1.22 | 2.644 | 6E-233 |
| 4 | XYLT1 | 0.129 | 0.078 | 0.487 | 0.442 | 0.138 | 6.8E-06 |
| 4 | YBX1 | 0.727 | 0.474 | 4.447 | 4.449 | 0.292 | 6.8E-10 |
| 4 | YME1L1 | 0.41 | 0.304 | 2.127 | 1.861 | 0.193 | 4.5E-06 |
| 4 | YPEL4 | 0.443 | 0.368 | 2.644 | 2.466 | 0.101 | 0.00089 |

| | | | | | | | |
|---|---|---|---|---|---|---|---|
| 4 | YWHAE | 0.324 | 0.208 | 1.469 | 1.134 | 0.374 | 1.5E-10 |
| 4 | YWHAG | 0.124 | 0.074 | 0.492 | 0.38 | 0.371 | 3.4E-06 |
| 4 | YWHAH | 0.143 | 0.082 | 0.404 | 0.392 | 0.367 | 1.5E-07 |
| 4 | YWHAQ | 0.41 | 0.399 | 2.938 | 2.499 | 0.177 | 1.8E-05 |
| 4 | Z93241.1 | 0.141 | 0.072 | 0.802 | 0.461 | 0.798 | 2.1E-11 |
| 4 | ZBED4 | 0.139 | 0.068 | 0.447 | 0.348 | 0.642 | 9.4E-12 |
| 4 | ZBTB21 | 0.114 | 0.077 | 0.404 | 0.394 | 0.341 | 0.00061 |
| 4 | ZC3H12A | 0.149 | 0.097 | 0.724 | 0.407 | 0.416 | 4.4E-07 |
| 4 | ZC3H7A | 0.364 | 0.229 | 1.646 | 1.307 | 0.342 | 9.8E-12 |
| 4 | ZCCHC8 | 0.134 | 0.089 | 0.409 | 0.442 | 0.202 | 0.0002 |
| 4 | ZDHHC21 | 0.183 | 0.118 | 0.724 | 0.627 | 0.207 | 4.8E-06 |
| 4 | ZEB1 | 0.447 | 0.102 | 2.48 | 0.486 | 2.082 | 9E-168 |
| 4 | ZFAND1 | 0.13 | 0.064 | 0.474 | 0.327 | 0.436 | 2.5E-10 |
| 4 | ZFAND4 | 0.277 | 0.197 | 1.296 | 1.088 | 0.242 | 4.4E-06 |
| 4 | ZFAS1 | 0.137 | 0.096 | 0.462 | 0.49 | 0.198 | 0.00108 |
| 4 | ZFP36 | 0.464 | 0.44 | 6.638 | 4.748 | 0.48 | 3E-10 |
| 4 | ZFP36L1 | 0.417 | 0.388 | 3.402 | 2.601 | 0.387 | 1.4E-08 |
| 4 | ZFP36L2 | 0.794 | 0.689 | 10.946 | 7.13 | 0.619 | 4.8E-17 |
| 4 | ZKSCAN1 | 0.144 | 0.107 | 0.614 | 0.443 | 0.141 | 0.00044 |
| 4 | ZMAT4 | 0.174 | 0.048 | 0.848 | 0.34 | 1.274 | 2.8E-34 |
| 4 | ZMIZ1 | 0.107 | 0.034 | 0.443 | 0.179 | 1.31 | 1.2E-21 |
| 4 | ZMPSTE24 | 0.13 | 0.087 | 0.418 | 0.43 | 0.27 | 0.00033 |
| 4 | ZMYM2 | 0.326 | 0.234 | 1.48 | 1.342 | 0.131 | 3.7E-05 |
| 4 | ZMYM4 | 0.246 | 0.169 | 1.122 | 0.908 | 0.304 | 4.2E-07 |
| 4 | ZNF217 | 0.389 | 0.241 | 1.864 | 1.396 | 0.417 | 3E-14 |
| 4 | ZNF280D | 0.177 | 0.128 | 0.741 | 0.668 | 0.168 | 0.00091 |
| 4 | ZNF281 | 0.12 | 0.083 | 0.493 | 0.413 | 0.246 | 0.00118 |
| 4 | ZNF32 | 0.101 | 0.072 | 0.384 | 0.342 | 0.129 | 0.00864 |
| 4 | ZNF322 | 0.104 | 0.064 | 0.43 | 0.314 | 0.44 | 0.0001 |
| 4 | ZNF639 | 0.113 | 0.074 | 0.448 | 0.363 | 0.303 | 0.00047 |
| 4 | ZNF74D | 0.101 | 0.07 | 0.473 | 0.36 | 0.392 | 0.00237 |
| 4 | ZNF770 | 0.209 | 0.131 | 0.871 | 0.644 | 0.414 | 6.3E-08 |
| 4 | ZNF791 | 0.133 | 0.091 | 0.403 | 0.469 | 0.101 | 0.00071 |
| 4 | ZNF808 | 0.179 | 0.122 | 0.711 | 0.622 | 0.193 | 0.00012 |
| 4 | ZNF92 | 0.143 | 0.1 | 0.662 | 0.412 | 0.372 | 3.3E-05 |
| 4 | ZNRD2 | 0.161 | 0.098 | 0.621 | 0.486 | 0.343 | 4.7E-07 |

| Table S4 \| Signature genes for gene set analysis | | |
|---|---|---|
| ISG signatures | NK cell activation signatures | Exhaustion signatures |
| IFIT1 | CCL3 | ADAM19 |
| IFIT3 | CCL4 | ADGRG1 |
| IFIT5 | CCL5 | BAIAP3 |
| IFIT2 | CCL28 | CD244 |
| IFI6 | CCL3L1 | CD38 |
| IFI44L | CCL4L2 | CD7 |
| IFI35 | GNLY | CELA1 |
| IFIH1 | GZMA | CXCL10 |
| IFI44 | GZMB | DAPK2 |
| IFI27 | GZMH | EOMES |
| IFI30 | GZMK | FGL2 |
| IFI16 | GZMM | GZMA |
| IFI27L1 | PRF1 | GZMK |
| IFI27L2 | LTB | HIP1 |
| IFITM2 | NKG7 | IFIT3 |
| IFITM3 | CST7 | IKZF3 |
| IFITM10 | ZAP70 | IRF7 |
| MX2 | TYROBP | ISG15 |
| MX1 | XCL1 | ISG20 |
| TBK1 | XCL2 | LAG3 |
| ISG15 | CSF1 | LRRK1 |
| ISG20 | EOMES | MAF |
| ISG20L2 | IL32 | NR4A2 |
| IRF1 | FCGR3A | OAS3 |
| IRF2 | FCER1G | PTGER4 |
| IRF2BP1 | IFNG | SLC27A4 |
| IRF2BP2 | KLRB1 | TOX |
| IRF2BPL | KLRC2 | TOX2 |
| IRF3 | KLRC3 | VMP1 |
| IRF4 | KLRC4 | ZBTB4 |
| IRF5 | KLRD1 | |
| IRF7 | KLRG1 | |
| IRF8 | KLRK1 | |
| IRF9 | TBX21 | |
| OAS1 | CX3CR1 | |
| OAS2 | CXCR2 | |
| OAS3 | CXCR3 | |
| OASL | CXCR4 | |
| STAT1 | TNF | |
| STAT2 | | |
| APOBEC3C | | |
| APOBEC3D | | |
| APOBEC3F | | |
| APOBEC3G | | |
| APOBEC3H | | |
| GBP1 | | |
| GBP2 | | |
| GBP3 | | |
| GBP4 | | |
| GBP5 | | |

| Table S5 \| DEgenes of IFN-β treated vs untreated NK cells | | |
|---|---|---|
| gene name | log2FC | p value |
| IFIT1 | 7.966835142 | 9.26E-14 |
| IFI27 | 7.903002407 | 4.12E-29 |
| IFI44L | 7.137006383 | 1.59E-26 |
| IFIT3 | 6.539272441 | 1.56E-11 |
| LINC00487 | 5.609296189 | 1.09E-09 |
| SPECC1L-ADORA2A | 5.477527367 | 0.002036822 |
| RPL23P3 | 5.444833716 | 5.93E-05 |
| LAMP3 | 5.313360555 | 1.32E-09 |
| MX1 | 5.299325962 | 5.02E-10 |
| IL6 | 5.049829588 | 0.03895924 |
| MX2 | 4.935051468 | 8.30E-11 |
| ERICH3 | 4.871257878 | 3.01E-06 |
| IFI44 | 4.863029569 | 3.74E-18 |
| SPAG6 | 4.861210784 | 0.000480732 |
| OTOF | 4.609779361 | 5.88E-16 |
| IFI6 | 4.566740201 | 1.40E-10 |
| USP41 | 4.486484062 | 0.00575979 |
| HES4 | 4.473284634 | 0.022186644 |
| CYP2J2 | 4.454723807 | 1.10E-11 |
| BCL2L14 | 4.438421224 | 0.013738 |
| RUFY4 | 4.302057439 | 0.003486276 |
| ARHGAP23 | 4.112444625 | 0.005176507 |
| OAS1 | 3.995902359 | 2.16E-14 |
| PLAAT2 | 3.954215203 | 0.000744661 |
| ISG15 | 3.947667447 | 6.52E-07 |
| SSTR3 | 3.89666879 | 1.40E-05 |
| TLR7 | 3.884682764 | 0.00493638 |
| NFE2 | 3.877709051 | 0.000803036 |
| GTF2IRD1 | 3.772562606 | 0.03576283 |
| RSAD2 | 3.771640214 | 7.49E-11 |
| C2orf66 | 3.713444768 | 0.008948349 |
| AXL | 3.621439999 | 0.03799672 |
| CD68 | 3.586365509 | 0.0006557 |
| GMPR | 3.581847857 | 1.00E-13 |
| CTSL | 3.515181909 | 0.008690507 |
| IFIT2 | 3.504824355 | 1.67E-05 |
| LINC02574 | 3.426151429 | 0.001503412 |
| POU5F1B | 3.394266429 | 0.003342819 |
| USP18 | 3.382459754 | 1.15E-09 |
| LGALS9 | 3.334496073 | 2.57E-10 |
| FBXO39 | 3.324008507 | 0.006574667 |
| RASGRP3 | 3.299354295 | 1.41E-09 |
| CMPK2 | 3.28696138 | 7.03E-11 |
| KLHDC7B | 3.279942532 | 1.92E-16 |
| PDCD1LG2 | 3.264730346 | 0.032659825 |
| GRM5-AS1 | 3.234247574 | 0.032304832 |
| HNRNPA3P5 | 3.210732877 | 0.041175498 |
| KCTD14 | 3.126020913 | 3.89E-05 |
| OAS3 | 3.09288409 | 3.39E-09 |

| | | |
|---|---|---|
| ODF3B | 3.027547241 | 1.36E-13 |
| SHISAL1 | 3.02091069 | 0.045211391 |
| HERC5 | 2.996041221 | 2.38E-16 |
| BATF2 | 2.982490079 | 6.27E-05 |
| HERC6 | 2.937794146 | 8.30E-07 |
| IRF7 | 2.936931485 | 2.10E-14 |
| ISG20 | 2.915098395 | 4.90E-06 |
| ETV7 | 2.914402601 | 0.000117362 |
| KLHDC7B-DT | 2.878240465 | 5.00E-06 |
| LGALS3BP | 2.821654434 | 1.05E-10 |
| CPLX3 | 2.764706894 | 0.034370788 |
| TTC21A | 2.725606889 | 0.044634421 |
| OR52N4 | 2.698063727 | 0.002355178 |
| TYMP | 2.696417359 | 9.01E-08 |
| PLSCR1 | 2.686085628 | 1.63E-18 |
| HOXB9 | 2.656634322 | 0.010308139 |
| CLEC7A | 2.645387728 | 0.000591672 |
| BTBD11 | 2.616118983 | 0.000240401 |
| DHRS9 | 2.604107359 | 0.030459684 |
| RASD1 | 2.594953101 | 0.015709361 |
| XAF1 | 2.568914001 | 6.99E-15 |
| CUEDC1 | 2.567168004 | 0.03371358 |
| TAF7L | 2.513236682 | 0.028393274 |
| UBQLNL | 2.489744298 | 0.001646441 |
| PDCD1 | 2.484705083 | 0.001120301 |
| GPR84 | 2.465089142 | 0.007501474 |
| SLC15A3 | 2.458354342 | 0.033971051 |
| EPSTI1 | 2.417850706 | 2.28E-10 |
| IL10 | 2.408307563 | 0.00392163 |
| TMEM119 | 2.40450446 | 0.043150174 |
| IL3RA | 2.375150463 | 0.007690761 |
| AK2P1 | 2.369564655 | 0.043734642 |
| ECE1 | 2.343077417 | 4.25E-06 |
| OAS2 | 2.338929541 | 9.41E-12 |
| HELZ2 | 2.28908088 | 1.05E-05 |
| LY6E | 2.276747286 | 5.68E-16 |
| CCL4L1 | 2.257773628 | 0.028107726 |
| CCL4L2 | 2.257773628 | 0.028107726 |
| NCF1C | 2.253163512 | 7.61E-07 |
| IFITM3 | 2.224187758 | 5.52E-06 |
| GDF15 | 2.206814565 | 0.015754831 |
| IRS1 | 2.136095716 | 0.012264771 |
| IL1RN | 2.12763409 | 0.004454393 |
| NCF1 | 2.117647356 | 2.97E-06 |
| H1-0 | 2.100465826 | 2.27E-07 |
| SH2B2 | 2.072810839 | 0.025068103 |
| C15orf48 | 2.030556483 | 4.83E-07 |
| FAM171A1 | 2.02779464 | 0.003563849 |
| MILR1 | 2.000185187 | 0.001996669 |
| OASL | 1.978224374 | 2.36E-10 |
| IFI35 | 1.950581195 | 6.60E-07 |

| | | |
|---|---|---|
| DHX58 | 1.934529204 | 8.40E-10 |
| CHN1 | 1.931890685 | 0.026523606 |
| RPL32P27 | 1.881349571 | 0.025203079 |
| SAMD9L | 1.87023555 | 1.16E-06 |
| PARP9 | 1.858177906 | 3.47E-09 |
| DDX60 | 1.843089598 | 1.17E-07 |
| IFIH1 | 1.836366345 | 5.66E-08 |
| RPS2P36 | 1.835928179 | 0.015919326 |
| AGRN | 1.827976772 | 6.02E-05 |
| IFIT5 | 1.814304983 | 9.33E-07 |
| NCF2 | 1.813370324 | 0.004456176 |
| TNFSF13B | 1.808307531 | 0.000102708 |
| DDX58 | 1.79760812 | 0.001280769 |
| NT5C3A | 1.779876159 | 8.54E-06 |
| STAT1 | 1.778457036 | 1.41E-07 |
| TRIM6 | 1.740104617 | 0.008125506 |
| EIF2AK2 | 1.729410084 | 1.37E-11 |
| NPDC1 | 1.725190075 | 0.026799359 |
| KIAA1217 | 1.719591317 | 0.022539263 |
| KIAA1614-AS1 | 1.717462134 | 0.045376023 |
| PLEKHD1 | 1.714165416 | 0.012214376 |
| LOC101927741 | 1.663400784 | 4.07E-06 |
| NCF1B | 1.661770424 | 0.000647407 |
| TTYH2 | 1.646893733 | 0.007164719 |
| SYTL5 | 1.628373355 | 0.010807598 |
| LAP3 | 1.619717095 | 0.002128765 |
| ASCL2 | 1.605151582 | 0.012599262 |
| LUCAT1 | 1.597597458 | 0.001105395 |
| HMOX1 | 1.593711025 | 0.000197834 |
| AK8 | 1.590996058 | 0.009338999 |
| SCO2 | 1.58215074 | 0.004754891 |
| SPATS2L | 1.56646082 | 2.56E-08 |
| PARP12 | 1.52875259 | 2.76E-07 |
| LRRC37A17P | 1.522501963 | 0.041985139 |
| AIM2 | 1.491822426 | 0.000180421 |
| STAT2 | 1.482021516 | 1.89E-05 |
| CD38 | 1.479876043 | 5.18E-05 |
| SHB | 1.471754028 | 0.000589289 |
| RAB20 | 1.461781209 | 0.024631022 |
| BEST1 | 1.461736645 | 0.016386559 |
| TMEM238 | 1.392704871 | 0.044220772 |
| TREX1 | 1.384358966 | 1.83E-05 |
| ATP6V0A1 | 1.382352334 | 0.017669279 |
| GALM | 1.380357044 | 2.74E-05 |
| TRIM25 | 1.375386482 | 2.60E-07 |
| UCHL1 | 1.365940738 | 0.047038592 |
| DDX60L | 1.365516143 | 7.35E-05 |
| PGAP1 | 1.362885565 | 8.94E-06 |
| ZCCHC2 | 1.356625299 | 2.20E-05 |
| CTTN | 1.356298584 | 0.045346162 |
| SBK1 | 1.354677877 | 1.75E-07 |

| | | |
|---|---|---|
| TLR3 | 1.352345041 | 0.006005848 |
| RRAD | 1.343287619 | 0.045955764 |
| RAB44 | 1.342396006 | 3.30E-11 |
| HDX | 1.334961218 | 0.002820112 |
| TASL | 1.313728361 | 0.004033454 |
| WARS1 | 1.296419371 | 0.00816166 |
| MTMR11 | 1.293819105 | 0.0039106 |
| SELL | 1.280660448 | 5.42E-05 |
| NKX3-1 | 1.275915234 | 0.029405415 |
| SERPINE1 | 1.274319166 | 0.008133737 |
| UBE2L6 | 1.27407363 | 1.15E-07 |
| CSF1 | 1.26694552 | 0.004523487 |
| LINC01260 | 1.25231298 | 0.004636615 |
| C5AR2 | 1.251846588 | 0.044376462 |
| SP110 | 1.244347845 | 2.06E-06 |
| IRF9 | 1.22899212 | 1.15E-13 |
| GCNT4 | 1.226176568 | 0.010080186 |
| CCDC194 | 1.225618398 | 0.001717635 |
| BST2 | 1.224977453 | 1.32E-06 |
| ZNFX1 | 1.213199507 | 0.000815496 |
| OSGIN1 | 1.203613509 | 0.042096065 |
| HFE | 1.199643921 | 1.18E-05 |
| SAMD9 | 1.182954522 | 0.000140213 |
| SOBP | 1.182061796 | 0.00068721 |
| CHMP5 | 1.178623146 | 5.55E-09 |
| TRIM22 | 1.174467109 | 3.85E-05 |
| GBP1 | 1.171506364 | 0.000659477 |
| ENDOD1 | 1.167718534 | 0.002603878 |
| FMNL2 | 1.166266874 | 0.017267023 |
| PPM1K | 1.157374894 | 9.34E-05 |
| CARMIL3 | 1.149934914 | 0.000632555 |
| GLP1R | 1.146514326 | 1.49E-07 |
| ZBP1 | 1.146087407 | 0.00028476 |
| KLF2 | 1.126816905 | 0.000248387 |
| DGAT2 | 1.126406882 | 0.000168851 |
| CYB561 | 1.122155889 | 0.002505298 |
| SERPINB9 | 1.121295275 | 0.001010288 |
| IL15RA | 1.118556588 | 0.000441652 |
| SMAGP | 1.108221764 | 3.71E-10 |
| C3 | 1.099738296 | 0.000120955 |
| SPATA1 | 1.097331059 | 0.023491704 |
| NINJ1 | 1.095748971 | 0.00201415 |
| LINC00504 | 1.08687806 | 0.000427514 |
| RASGEF1B | 1.082808667 | 0.025686567 |
| SP100 | 1.080098814 | 4.37E-07 |
| CDK18 | 1.075081655 | 0.014737498 |
| IFI30 | 1.068908062 | 8.50E-05 |
| DTX3L | 1.054990905 | 0.000343957 |
| TRIM69 | 1.050114056 | 8.33E-06 |
| ALPK1 | 1.049163831 | 7.64E-06 |
| TSPAN18 | 1.039740073 | 0.000972125 |

| | | |
|---|---|---|
| SAT1 | 1.036563226 | 2.30E-05 |
| PML | 1.035566145 | 3.00E-06 |
| APBB2 | 1.027304399 | 0.001623222 |
| PALM2AKAP2 | 1.021318094 | 9.57E-06 |
| BACE2 | 1.018749015 | 0.029738045 |
| METRNL | 1.016655715 | 3.64E-06 |
| SYP | 1.01267303 | 0.017654367 |
| TREML2 | 1.007619161 | 2.20E-07 |
| PHYHD1 | 1.006092991 | 0.041371581 |
| CCDC96 | 1.005112843 | 0.045353546 |
| ALOX5AP | 1.002010201 | 2.03E-05 |
| SPSB1 | 0.998036433 | 0.048075117 |
| RNASE4 | 0.992636028 | 0.041857725 |
| FOSL1 | 0.991860426 | 0.004384173 |
| PARP10 | 0.985047461 | 5.46E-05 |
| GDPD5 | 0.984506621 | 0.024124748 |
| RTP4 | 0.980413488 | 0.000550674 |
| TENT5A | 0.977529661 | 5.50E-05 |
| PRRG4 | 0.972184035 | 0.00182715 |
| CXCR1 | 0.969897877 | 0.006206486 |
| GRN | 0.966230832 | 0.000209787 |
| HM13 | 0.96335397 | 0.002112557 |
| PARP14 | 0.957810021 | 0.00080179 |
| HNRNPLL | 0.955948438 | 4.33E-10 |
| PATL2 | 0.955038735 | 6.65E-05 |
| CAPN3 | 0.95087512 | 0.006997455 |
| NPTX1 | 0.947823811 | 0.016382516 |
| FRMD4B | 0.945379632 | 0.000640257 |
| SERPINB1 | 0.94464928 | 1.68E-05 |
| TTC22 | 0.94276729 | 1.26E-05 |
| ARPIN-AP3S2 | 0.942137774 | 0.000416839 |
| TAP1 | 0.930688273 | 1.06E-05 |
| ADAR | 0.927994311 | 0.00016161 |
| CAPN2 | 0.927815393 | 4.84E-09 |
| LEXM | 0.927503531 | 0.001431819 |
| ZC3H12A | 0.926120426 | 0.005474885 |
| IFITM1 | 0.925207402 | 0.00018704 |
| ARL11 | 0.920102975 | 0.005749787 |
| MOV10 | 0.916453833 | 0.000265026 |
| RAP1GAP2 | 0.914365144 | 0.000178505 |
| IRAK2 | 0.908241879 | 0.002205889 |
| ABCB6 | 0.905216535 | 0.013695242 |
| CCL3 | 0.904284103 | 0.003166857 |
| CD274 | 0.900044883 | 0.002952946 |
| CHRNA10 | 0.898726947 | 0.025572519 |
| CD9 | 0.893436634 | 0.000389926 |
| SLC38A5 | 0.890142665 | 0.00064367 |
| HVCN1 | 0.88231764 | 0.000161505 |
| EPOP | 0.88081036 | 0.000101471 |
| CCR1 | 0.875939529 | 0.000125654 |
| ADAP1 | 0.869243089 | 0.034307828 |

| | | |
|---|---|---|
| SCARB2 | 0.867468801 | 6.76E-06 |
| ELF3 | 0.865628684 | 0.035400629 |
| PLPPR2 | 0.862788195 | 0.00812861 |
| PGD | 0.856833675 | 0.001397993 |
| ARRDC3 | 0.848066395 | 0.001309539 |
| PIK3AP1 | 0.845962964 | 0.001530641 |
| CLIC4 | 0.843169809 | 0.022074741 |
| CEBPB | 0.842392295 | 0.000802336 |
| VAMP5 | 0.841511769 | 0.001102115 |
| GTPBP1 | 0.8413457 | 0.000174191 |
| CCL4 | 0.839566698 | 0.010332153 |
| CPT1B | 0.839538839 | 7.17E-05 |
| CLDN12 | 0.838473955 | 0.028419676 |
| TGFBR3 | 0.837544144 | 0.005248046 |
| FHDC1 | 0.835504108 | 0.035041246 |
| PSMB9 | 0.833262917 | 1.12E-06 |
| GPR153 | 0.823331845 | 0.003429125 |
| ZNF273 | 0.821487956 | 7.19E-05 |
| PHF11 | 0.814644055 | 1.58E-05 |
| RBM47 | 0.813582151 | 0.0331437 |
| GPAT3 | 0.813053571 | 0.021175392 |
| GCH1 | 0.81183271 | 0.000493565 |
| ISM1 | 0.806879997 | 0.005548021 |
| NAPA | 0.806376345 | 0.000717732 |
| AKR1C7P | 0.806054171 | 0.03003848 |
| SP140 | 0.80428399 | 0.000211301 |
| CREG1 | 0.80368637 | 0.000713014 |
| BFSP1 | 0.80074191 | 0.001982813 |
| TRANK1 | 0.799423054 | 0.004257812 |
| NMI | 0.798015403 | 0.000763833 |
| STAC3 | 0.795836048 | 0.002579405 |
| SEMA4A | 0.795578186 | 0.001298391 |
| CNP | 0.79507438 | 0.00021523 |
| IL17RA | 0.793970563 | 0.000661491 |
| ZNF20 | 0.790755257 | 0.010040053 |
| NAT14 | 0.784872403 | 0.014917803 |
| ZNF334 | 0.777630114 | 0.026099753 |
| LONRF3 | 0.777226132 | 0.007105223 |
| RNF19B | 0.776249924 | 0.006856087 |
| LRRC3 | 0.769240425 | 0.030938228 |
| IRF4 | 0.766523307 | 0.000462958 |
| ITGA5 | 0.766252123 | 1.33E-05 |
| DRAP1 | 0.757353526 | 6.28E-05 |
| SHFL | 0.756394977 | 0.000821035 |
| RELL2 | 0.754704338 | 0.002133982 |
| PHACTR1 | 0.752186262 | 0.001337351 |
| N4BP1 | 0.751923077 | 0.000343322 |
| ZNF311 | 0.751491601 | 0.024112371 |
| GNG5 | 0.75002914 | 2.20E-05 |
| BLZF1 | 0.748082947 | 0.006347859 |
| MYD88 | 0.747466008 | 0.000349329 |

| | | |
|---|---|---|
| IFI16 | 0.742214238 | 2.71E-05 |
| PLAAT4 | 0.738942414 | 0.007485125 |
| RHEBL1 | 0.735772002 | 0.000142578 |
| GZMB | 0.733026797 | 0.002750365 |
| CMTR1 | 0.731827544 | 3.12E-05 |
| H1-10 | 0.731087391 | 0.003907928 |
| IFITM2 | 0.72286682 | 0.000207327 |
| DAPP1 | 0.722153143 | 0.004536775 |
| PNPT1 | 0.721832421 | 0.005778675 |
| SIGLEC6 | 0.72072638 | 0.028103391 |
| HLA-C | 0.717176074 | 0.004014782 |
| ACP2 | 0.716398684 | 4.98E-08 |
| IER3 | 0.716334814 | 0.004839791 |
| MYADM | 0.713361564 | 0.030331616 |
| PSME2 | 0.70770043 | 8.30E-08 |
| TAP2 | 0.705340513 | 0.007929349 |
| SNAI3 | 0.703873815 | 0.014973158 |
| L3HYPDH | 0.702598393 | 0.003577517 |
| S100A11 | 0.700456222 | 1.81E-05 |
| H2BC5 | 0.700038048 | 0.016185449 |
| OSM | 0.693015716 | 0.009389173 |
| TFEB | 0.690367324 | 0.00185852 |
| IRF5 | 0.689513812 | 0.000759238 |
| HIVEP3 | 0.688491791 | 0.024172645 |
| TMEM62 | 0.687244466 | 0.000352702 |
| KCTD12 | 0.685790506 | 0.01075329 |
| ACTN1 | 0.684857683 | 0.03192464 |
| EHD4 | 0.680323948 | 0.0242303 |
| PDLIM7 | 0.679077131 | 0.000533555 |
| FLVCR2 | 0.677818766 | 0.033430085 |
| REC8 | 0.676814445 | 0.002480472 |
| RAB3D | 0.675919089 | 0.001870403 |
| ACSL1 | 0.671473894 | 0.019444784 |
| BCL3 | 0.669729063 | 0.004236821 |
| NFKBIA | 0.669211928 | 0.012022511 |
| TRIM5 | 0.665811118 | 0.006077489 |
| TDRD7 | 0.663833141 | 0.000353533 |
| GBP2 | 0.659804851 | 0.008912373 |
| TNFSF10 | 0.657535347 | 0.000608706 |
| FAM89A | 0.656760387 | 0.041147058 |
| HLA-A | 0.656503043 | 0.000499588 |
| CEBPD | 0.656310792 | 0.024632067 |
| SQSTM1 | 0.650131546 | 0.008983908 |
| IKBKE | 0.649275273 | 0.010804938 |
| CX3CR1 | 0.644319125 | 0.023780161 |
| DUSP5 | 0.644150346 | 0.018095961 |
| PDIA5 | 0.644138948 | 4.02E-05 |
| H2BC21 | 0.642291425 | 0.026920841 |
| FTH1 | 0.639849685 | 0.000697108 |
| UBA7 | 0.639743308 | 0.002391802 |
| TSPAN2 | 0.636030046 | 5.03E-07 |

| | | |
|---|---|---|
| HOXA1 | 0.631040713 | 0.025677112 |
| HSPA1B | 0.628613153 | 0.002844626 |
| CHROMR | 0.628494277 | 0.007234657 |
| MICB | 0.62688065 | 2.35E-05 |
| PLAC8 | 0.625491097 | 0.025282974 |
| MYDGF | 0.624869626 | 0.004430714 |
| PPIF | 0.624187607 | 0.003603178 |
| CD164 | 0.621706313 | 0.000265663 |
| TMEM123 | 0.618119182 | 0.006001973 |
| NAPSA | 0.615975925 | 0.025776282 |
| FGFRL1 | 0.615396329 | 0.023648602 |
| E2F2 | 0.612604258 | 1.02E-10 |
| B2M | 0.607914829 | 1.12E-05 |
| SESN2 | 0.603859404 | 0.02399576 |
| TLDC2 | 0.596460387 | 0.007006835 |
| ITGA3 | 0.596068783 | 0.005455156 |
| JUNB | 0.595910593 | 9.36E-05 |
| ATP6V0C | 0.59483673 | 0.003157116 |
| HLA-B | 0.594190925 | 0.000558769 |
| TRIM34 | 0.592741884 | 6.03E-05 |
| GALNT3 | 0.592708249 | 0.002009911 |
| TKTL1 | 0.591568588 | 0.006659089 |
| NFE2L2 | 0.591026967 | 0.000634431 |
| MXD1 | 0.588690178 | 0.003999537 |
| FTL | 0.587766849 | 0.016535352 |
| C1R | 0.582150932 | 0.041141198 |
| CHCHD10 | 0.579833734 | 0.014541712 |
| DYNLT1 | 0.579390521 | 0.000651075 |
| GRK5 | 0.578381164 | 0.011480636 |
| HLA-DPA1 | 0.574232515 | 0.018443602 |
| JUN | 0.573932708 | 0.000189116 |
| SHISA5 | 0.572975089 | 0.000709945 |
| QSER1 | 0.572970414 | 0.000277421 |
| TNFSF13 | 0.57175945 | 0.018382109 |
| MLKL | 0.570874708 | 0.004952052 |
| TRIM38 | 0.568879469 | 0.00618712 |
| MASTL | 0.567681641 | 0.001917858 |
| ANXA1 | 0.566746058 | 0.001465886 |
| MARCHF3 | 0.566116353 | 0.008366218 |
| TRADD | 0.566020989 | 0.02228176 |
| JAK2 | 0.565050234 | 0.038312931 |
| FBXO6 | 0.563664621 | 0.005341311 |
| NFKBIE | 0.561385814 | 0.017110541 |
| YEATS2 | 0.554528528 | 0.003808661 |
| HLA-F | 0.554471605 | 0.012836317 |
| VWA7 | 0.553567265 | 0.016201285 |
| DDIT3 | 0.551653018 | 0.009788852 |
| C16orf54 | 0.550842061 | 0.008269985 |
| GAREM2 | 0.547662057 | 0.033991661 |
| SH3BP5 | 0.547255358 | 0.006990349 |
| SQOR | 0.546820802 | 0.00440444 |

| | | |
|---|---|---|
| TRIM14 | 0.545194577 | 0.001558228 |
| FBXO8 | 0.544842835 | 0.00079366 |
| GNA15 | 0.543106678 | 0.002929216 |
| APOL3 | 0.541595198 | 0.01595044 |
| MUC1 | 0.539957508 | 0.005438645 |
| CTSC | 0.539495858 | 0.000265337 |
| PMAIP1 | 0.536345741 | 0.036473828 |
| MZB1 | 0.534671806 | 0.033465691 |
| RBCK1 | 0.534621676 | 0.000249309 |
| CTSS | 0.533141083 | 0.024026776 |
| FGD2 | 0.532244593 | 0.007218519 |
| CASP7 | 0.530422789 | 0.018889372 |
| SLC16A3 | 0.528035723 | 0.013762358 |
| PSMB8 | 0.527466337 | 1.01E-07 |
| VPS18 | 0.527269211 | 0.004210607 |
| PARS2 | 0.527162308 | 0.015000058 |
| FAM241B | 0.526774182 | 0.04838318 |
| ZNF267 | 0.522645446 | 0.026616742 |
| ADAMTS14 | 0.520856388 | 0.002787234 |
| TRIM21 | 0.519292507 | 0.015174964 |
| RARG | 0.518439473 | 0.002710575 |
| ULBP2 | 0.51817495 | 0.004309575 |
| LIMK2 | 0.518069103 | 0.026747831 |
| PIM3 | 0.516791117 | 0.025963534 |
| CCR5 | 0.51518231 | 0.049106703 |
| RNF31 | 0.514867262 | 0.000592743 |
| APOL1 | 0.512014304 | 0.011539696 |
| SRGAP2C | 0.511782218 | 0.022563892 |
| NECTIN3 | 0.510343417 | 0.02824814 |
| ANXA4 | 0.508487237 | 0.009904301 |
| SRGAP2 | 0.508099389 | 0.012161154 |
| PTPRN2 | 0.507730798 | 0.01021768 |
| PSMB10 | 0.503662055 | 0.0126719 |
| RIPOR2 | 0.503178703 | 0.000254153 |
| PDIA4 | 0.501573116 | 0.027486232 |
| SOX13 | 0.500157059 | 0.011061445 |
| MBOAT2 | -0.500623006 | 0.02159505 |
| GPATCH11 | -0.501292101 | 0.00091396 |
| MTCO1P12 | -0.501388786 | 0.02278951 |
| ATP8A1 | -0.502748187 | 0.013778495 |
| GAN | -0.503264315 | 0.000128525 |
| DTWD2 | -0.503385247 | 0.017496668 |
| GADD45A | -0.50469837 | 0.016054601 |
| ZNF577 | -0.504709769 | 0.013772272 |
| MIAT | -0.505318957 | 0.014077563 |
| ATP8 | -0.506813538 | 0.001635195 |
| LINC00205 | -0.506841094 | 0.010609582 |
| USP53 | -0.507212427 | 0.002858502 |
| MSI2 | -0.510259667 | 0.003416474 |
| ZNF660 | -0.510270531 | 0.036903689 |
| MORC4 | -0.511223702 | 0.040318481 |

| | | |
|---|---|---|
| ACACA | -0.512361859 | 0.031091435 |
| DRAXIN | -0.517436296 | 0.035813366 |
| TRIM74 | -0.517620864 | 0.047562182 |
| L3MBTL1 | -0.518131541 | 0.027520112 |
| AKT3 | -0.518901852 | 0.002408708 |
| PDE4B | -0.519174 | 0.027865442 |
| NCR2 | -0.520677009 | 0.015216771 |
| SLC46A1 | -0.522766832 | 0.001131118 |
| BCAS3 | -0.525650915 | 0.038110672 |
| GOLGA6L4 | -0.529411342 | 0.047987794 |
| NUDT3 | -0.531243937 | 4.42E-06 |
| LINC00954 | -0.531961425 | 0.044807567 |
| SYNE2 | -0.532435119 | 0.001121775 |
| ALDOC | -0.532810969 | 0.0493855 |
| CBX5 | -0.532925261 | 0.000929466 |
| BTBD3 | -0.533582734 | 0.001457074 |
| FAM168B | -0.534498091 | 0.000192151 |
| PAIP2B | -0.536610111 | 0.035313685 |
| SMIM11 | -0.537144545 | 0.008499051 |
| GOLGA8O | -0.537426921 | 0.036479838 |
| KLRF1 | -0.537534619 | 0.007564857 |
| OPN3 | -0.541758898 | 0.010801728 |
| ND4L | -0.543754869 | 4.86E-05 |
| STAU2 | -0.545783705 | 0.011343231 |
| LENG8-AS1 | -0.546879002 | 0.046692569 |
| SORL1 | -0.548662112 | 0.005443768 |
| APBA2 | -0.550371584 | 0.000928667 |
| CMTM4 | -0.553251597 | 0.04100479 |
| RCAN3 | -0.556385127 | 0.046735865 |
| DANCR | -0.556948765 | 0.001316091 |
| LINC00892 | -0.557400923 | 0.032180455 |
| PLCE1 | -0.558455974 | 0.016175486 |
| RBL2 | -0.568153801 | 0.000211064 |
| UBXN7 | -0.56884016 | 4.89E-05 |
| JMJD7-PLA2G4B | -0.570098269 | 0.008551288 |
| LOC103344931 | -0.570897532 | 0.013942604 |
| SLC2A13 | -0.57101297 | 0.046801644 |
| LYRM7 | -0.573165905 | 5.61E-05 |
| SLC25A15 | -0.573381729 | 0.025611735 |
| RPS3A | -0.575429142 | 0.001051166 |
| GOLGA8B | -0.575688179 | 1.50E-06 |
| EIF3L | -0.577579509 | 0.000750972 |
| PIAS2 | -0.58013827 | 0.01722617 |
| SOCS2-AS1 | -0.581127403 | 0.038691588 |
| HILPDA | -0.582876513 | 0.049637273 |
| LOC101927060 | -0.583082314 | 0.004511507 |
| NBPF15 | -0.586235367 | 0.000156811 |
| CHML | -0.589169495 | 0.021955991 |
| KIAA0825 | -0.589938033 | 0.01337279 |
| PDCD4 | -0.590344643 | 0.001161216 |
| MACF1 | -0.592373375 | 0.001525738 |

| | | |
|---|---|---|
| ND5 | -0.592997114 | 2.07E-16 |
| MASP2 | -0.595675631 | 0.014772768 |
| CD93 | -0.59879798 | 0.018733509 |
| PC | -0.600317451 | 0.02449108 |
| ZNF232-AS1 | -0.601213182 | 0.030810976 |
| GPR157 | -0.602629469 | 0.019408944 |
| LDLRAD4 | -0.603711193 | 0.008576768 |
| CBX7 | -0.603982013 | 0.045716462 |
| LOC729296 | -0.604381859 | 0.049708719 |
| MIR646HG | -0.604381859 | 0.049708719 |
| CEP170 | -0.605681895 | 0.013385025 |
| CEP170P1 | -0.605681895 | 0.013385025 |
| GOLGA8N | -0.611915049 | 0.009679006 |
| CCDC74A | -0.615466958 | 0.032629333 |
| ACVR2A | -0.618024206 | 0.013717799 |
| H4C8 | -0.619082063 | 0.02804518 |
| ADAMTS17 | -0.619375083 | 0.004086721 |
| IGF1R | -0.622888684 | 0.020131282 |
| SCAI | -0.626844794 | 0.013430549 |
| ATP13A3 | -0.627914193 | 0.000803221 |
| RNF144A | -0.628273347 | 0.034256949 |
| PCYOX1 | -0.628403507 | 0.000133177 |
| SH3YL1 | -0.630000281 | 0.000338323 |
| GOLGA8A | -0.630511483 | 1.44E-05 |
| DLG5 | -0.630731595 | 1.50E-05 |
| GEM | -0.631556857 | 6.69E-05 |
| TSPAN3 | -0.631646816 | 0.002762917 |
| LDB2 | -0.632784389 | 0.033176299 |
| FAM102A | -0.634586572 | 0.027902213 |
| FITM2 | -0.635882413 | 0.000188632 |
| MYO9A | -0.643688577 | 0.001156776 |
| LINC01963 | -0.6438912 | 5.25E-06 |
| SPRY2 | -0.649116634 | 0.031954549 |
| CPQ | -0.649924887 | 0.048898409 |
| MAP3K9 | -0.650231702 | 0.001003592 |
| NCR1 | -0.652010848 | 0.002845233 |
| CDCA7L | -0.654235395 | 0.013797473 |
| RASSF4 | -0.655491587 | 0.001760693 |
| CYTB | -0.663083678 | 0.010817239 |
| ZSWIM4 | -0.664982783 | 0.022044948 |
| SOCS7 | -0.665018462 | 2.05E-05 |
| F2R | -0.665316919 | 0.008519902 |
| SERP2 | -0.667914857 | 0.002707797 |
| C1orf198 | -0.66929375 | 0.008128511 |
| CCR2 | -0.67637543 | 0.027996016 |
| SPEF2 | -0.676805712 | 0.000777005 |
| IKBKGP1 | -0.677592671 | 0.016943636 |
| ZDHHC20P1 | -0.681824071 | 0.00464783 |
| TMEM200A | -0.685034452 | 0.030565502 |
| LINC01814 | -0.686705928 | 0.009928435 |
| ZNF154 | -0.687483215 | 0.000802854 |

| | | |
|---|---|---|
| MIB1 | -0.688759172 | 5.45E-05 |
| BRI3BP | -0.69014389 | 0.001680873 |
| GOLGA8Q | -0.690993838 | 0.013476076 |
| ST8SIA6 | -0.691991861 | 0.002954748 |
| ABLIM1 | -0.693673388 | 0.013306797 |
| CELSR2 | -0.69535075 | 0.018740946 |
| ZNF567-DT | -0.698234795 | 0.011183559 |
| PRKCA | -0.700986566 | 3.68E-05 |
| CAMK1 | -0.70154426 | 7.51E-05 |
| SACS | -0.70254364 | 0.002905691 |
| LRRC70 | -0.706899346 | 0.013816579 |
| RIMKLB | -0.708939143 | 0.036173319 |
| USP13 | -0.709364813 | 0.003793315 |
| AFDN | -0.710720213 | 1.11E-06 |
| ARMCX2 | -0.711739088 | 0.033074101 |
| MSANTD3-TMEFF1 | -0.71180437 | 0.024283725 |
| LIPG | -0.714114386 | 0.048184089 |
| SUCLG2-DT | -0.714383291 | 0.00311461 |
| EIF4B | -0.716490104 | 6.54E-10 |
| KLRC4 | -0.718594206 | 0.038932459 |
| FAM210B | -0.719181017 | 0.002368251 |
| CACNA1C-AS1 | -0.720308503 | 0.039792191 |
| KLRK1-AS1 | -0.721558059 | 0.000509605 |
| RN7SL1 | -0.721560018 | 0.036129884 |
| ABCC4 | -0.72250851 | 0.003653036 |
| CXCR6 | -0.725017162 | 0.044053106 |
| FGD4 | -0.730005622 | 0.015222874 |
| SLC14A1 | -0.730601902 | 0.00911693 |
| DAAM1 | -0.732305512 | 0.003562665 |
| ZNF827 | -0.734873653 | 6.64E-05 |
| RANBP17 | -0.735720348 | 0.04913982 |
| AFAP1L2 | -0.739898513 | 0.034073733 |
| BACE1 | -0.743901737 | 0.002653867 |
| ZNF512B | -0.745560419 | 0.004963006 |
| ZHX3 | -0.747175167 | 0.000718744 |
| LINC02458 | -0.748099816 | 0.034589985 |
| LINC00861 | -0.751069501 | 0.003727105 |
| PLXNA1 | -0.752050156 | 0.002816648 |
| GP5 | -0.752790278 | 0.049584635 |
| PFKFB2 | -0.753400199 | 5.70E-06 |
| SLC12A6 | -0.754683372 | 2.53E-06 |
| SEMA3A | -0.75469516 | 0.025868753 |
| MAL | -0.761510756 | 0.018084013 |
| TTN | -0.768468052 | 0.029682293 |
| ASTN2 | -0.770558739 | 0.045586592 |
| AHRR | -0.772146623 | 0.038927174 |
| VASH1 | -0.784810496 | 0.002046502 |
| FAM135A | -0.784842149 | 0.027511682 |
| EPAS1 | -0.78585905 | 0.01134597 |
| ZNF337-AS1 | -0.789244949 | 0.007246044 |
| MAP1LC3A | -0.790081648 | 0.040066526 |

| | | |
|---|---|---|
| SIX4 | -0.790649494 | 0.024726646 |
| BOLA3-DT | -0.791032953 | 0.006063618 |
| ABCB1 | -0.792512277 | 1.62E-05 |
| BCL6 | -0.793128569 | 0.013497143 |
| ARHGAP42 | -0.794946881 | 0.000740769 |
| DOCK7 | -0.799295746 | 0.004687649 |
| KLF7 | -0.805133756 | 0.000105724 |
| ACO1 | -0.806197556 | 0.010867287 |
| GLI1 | -0.807245069 | 0.030226232 |
| PNPLA7 | -0.810201678 | 0.032089043 |
| CAMK4 | -0.811556725 | 0.034068962 |
| GNAQ | -0.817837207 | 0.000327677 |
| TNFSF4 | -0.819810518 | 0.001013044 |
| SPIRE1 | -0.821090478 | 0.000370989 |
| HSD17B8 | -0.823549036 | 0.019490012 |
| ZFHX3 | -0.825152985 | 0.017395824 |
| LINC01132 | -0.827010489 | 0.015106014 |
| IMMP2L | -0.834191301 | 0.036667309 |
| ADAM22 | -0.841159117 | 0.015156668 |
| SOGA1 | -0.843240806 | 0.002119754 |
| FAM124B | -0.843808843 | 0.006498562 |
| GUCY1B1 | -0.847947674 | 0.004369764 |
| EML6 | -0.848647546 | 0.000149679 |
| CCDC144B | -0.856114196 | 0.027440011 |
| TNIP3 | -0.86615318 | 0.031691006 |
| TANC2 | -0.866284649 | 0.000156563 |
| DST | -0.867236377 | 0.008703462 |
| SH3D21 | -0.872709222 | 0.000158143 |
| SLC4A4 | -0.890375878 | 0.000348628 |
| RTKN2 | -0.894584955 | 0.003336809 |
| VEGFA | -0.896616736 | 0.001139286 |
| YPEL2 | -0.898555628 | 8.38E-05 |
| C20orf204 | -0.905440648 | 0.039349723 |
| CES4A | -0.907609429 | 7.26E-05 |
| SMPD3 | -0.913789502 | 0.000148003 |
| MTURN | -0.92163077 | 0.01666571 |
| PDGFD | -0.921705707 | 1.19E-06 |
| BCAS4 | -0.922147501 | 0.000746008 |
| CASS4 | -0.927205953 | 0.000129001 |
| SOX4 | -0.928072629 | 0.010784654 |
| CEP126 | -0.93285288 | 8.38E-06 |
| AMOTL1 | -0.933131237 | 1.70E-06 |
| JMY | -0.954095732 | 0.008067056 |
| USP43 | -0.954216006 | 0.001793982 |
| ARRB1 | -0.955864217 | 1.07E-09 |
| CYP1A1 | -0.95682636 | 0.004355131 |
| LOC651337 | -0.971748967 | 0.014257253 |
| CRTAM | -0.976166548 | 0.000671267 |
| SESN3 | -0.979444683 | 0.014999785 |
| ARL10 | -0.986492055 | 2.70E-09 |
| GREM2 | -0.987591771 | 0.04515075 |

| | | |
|---|---|---|
| PKIA | -0.991060623 | 0.029871132 |
| ITM2A | -0.995022595 | 1.48E-06 |
| CDC42BPA | -1.005409742 | 0.021126712 |
| CREB3L3 | -1.008286718 | 0.03822445 |
| NETO2 | -1.011209131 | 0.008102614 |
| GALNT12 | -1.013763703 | 0.012791699 |
| CRYBG3 | -1.013941717 | 0.007340592 |
| GRK3 | -1.016664046 | 5.33E-06 |
| CCDC88A | -1.024703305 | 0.000598393 |
| MTSS2 | -1.030745586 | 0.015690762 |
| FBN1 | -1.03230366 | 0.002682491 |
| LINC02175 | -1.035108161 | 0.005606077 |
| BCL2L2-PABPN1 | -1.041457548 | 0.004004308 |
| PLAU | -1.04454467 | 0.043301324 |
| LEF1 | -1.045313346 | 0.000140571 |
| PLAG1 | -1.046758719 | 0.01744656 |
| ALDH5A1 | -1.051202698 | 0.012611017 |
| KLRB1 | -1.051778017 | 0.000385129 |
| MYO18B | -1.053553831 | 0.043064023 |
| STARD4-AS1 | -1.05754414 | 0.009844776 |
| SMIM10L2A | -1.058895423 | 0.01502296 |
| PITPNA-AS1 | -1.064318427 | 0.009550106 |
| FAM117B | -1.072612834 | 0.010487669 |
| MYO6 | -1.077872087 | 0.027180169 |
| SALL2 | -1.079577867 | 0.036279695 |
| RAB6B | -1.08339826 | 0.006145047 |
| CLIC5 | -1.088260629 | 0.030641225 |
| PDK4 | -1.099917574 | 0.004935849 |
| CYTH3 | -1.113511251 | 1.23E-06 |
| UBASH3A | -1.123977357 | 5.17E-05 |
| KANK2 | -1.125442082 | 0.003781669 |
| ADGRA3 | -1.128359436 | 0.000113299 |
| TLR4 | -1.133377019 | 0.00347192 |
| INPP4B | -1.141535138 | 0.002712017 |
| TRAV22 | -1.146351754 | 0.039786943 |
| BEX2 | -1.148069689 | 0.008044536 |
| PPP1R9A | -1.149653167 | 0.033154803 |
| ANKRD6 | -1.153863558 | 0.004642188 |
| IL7R | -1.157103355 | 0.011564652 |
| NEO1 | -1.158772121 | 0.007901703 |
| SMIM24 | -1.165909648 | 0.01357956 |
| MIR600HG | -1.171402513 | 0.00010613 |
| EGR3 | -1.180947038 | 0.00830548 |
| MCC | -1.197759957 | 0.003310167 |
| GLI2 | -1.200644898 | 0.000666123 |
| THEMIS | -1.202760893 | 3.33E-05 |
| ADAMTS10 | -1.207362578 | 0.016812866 |
| TAPT1-AS1 | -1.215849392 | 0.018450106 |
| TIE1 | -1.21953874 | 0.015610194 |
| KIF5A | -1.230039666 | 0.011306817 |
| JAML | -1.231581282 | 0.030205401 |

| | | |
|---|---|---|
| OBSCN | -1.23857073 | 8.74E-05 |
| APBA1 | -1.25360544 | 4.91E-10 |
| SNHG9 | -1.259261832 | 0.005538735 |
| IL9R | -1.26136019 | 0.008366843 |
| CDH17 | -1.271388706 | 0.015372186 |
| RUNDC3B | -1.273431054 | 0.039463101 |
| WASL-DT | -1.279796157 | 0.007331461 |
| PCSK5 | -1.281114666 | 0.018838256 |
| ZNF540 | -1.282678087 | 0.000112254 |
| ZNF883 | -1.298368799 | 0.01247162 |
| TXNRD3 | -1.304047768 | 0.005294534 |
| NHS | -1.308747708 | 0.029069783 |
| RPS6KA2 | -1.323613678 | 0.009464129 |
| MFSD6L | -1.324173973 | 0.048735688 |
| TMEM170B | -1.361002431 | 8.49E-07 |
| CADM1 | -1.363246701 | 0.020776881 |
| KLHL33 | -1.364733375 | 0.01641242 |
| NELL2 | -1.381459293 | 0.015445722 |
| FCRL6 | -1.38999332 | 0.048600464 |
| FCRL3 | -1.391066292 | 0.002563707 |
| AXIN2 | -1.408177966 | 0.042033825 |
| SPTLC3 | -1.413889743 | 0.001286594 |
| ACBD7 | -1.415264246 | 0.000835678 |
| EEF1A1P14 | -1.420142533 | 0.001356405 |
| CD27 | -1.428498561 | 0.012840493 |
| SRGAP3 | -1.451862584 | 0.015810245 |
| AEBP1 | -1.46216121 | 0.030277285 |
| ENPP1 | -1.481013505 | 0.009050035 |
| P2RY14 | -1.491832888 | 3.59E-06 |
| MYO5B | -1.49199338 | 0.000966076 |
| KCNC1 | -1.49241043 | 0.040827309 |
| PLXDC2 | -1.493036973 | 0.000444379 |
| KLRF2 | -1.505618784 | 0.00038635 |
| COL24A1 | -1.506042926 | 0.048862093 |
| CDH2 | -1.516960418 | 0.001935356 |
| NTRK2 | -1.52435815 | 0.013918685 |
| ANKRD50 | -1.541704477 | 0.017776448 |
| ZNF365 | -1.556688111 | 0.010563386 |
| PTPRG | -1.556717342 | 0.004549544 |
| ZNF497 | -1.564768167 | 0.011366636 |
| FHAD1 | -1.574393464 | 0.046570689 |
| COL4A4 | -1.576180116 | 0.005029597 |
| KIT | -1.614793721 | 1.56E-05 |
| LOC105375743 | -1.620404156 | 0.023236954 |
| ZDHHC23 | -1.626385068 | 8.52E-11 |
| SH2D4A | -1.644916816 | 0.007119811 |
| AFAP1L1 | -1.684359426 | 0.004572281 |
| ABLIM2 | -1.684461751 | 0.015437468 |
| MGLL | -1.717280932 | 0.000959156 |
| CAPN5 | -1.768885292 | 0.03747957 |
| NRXN3 | -1.779318289 | 0.007622867 |

| | | |
|---|---|---|
| RNF217 | -1.862065539 | 0.006760918 |
| PDE7B | -1.872951619 | 0.008351786 |
| TRMT9B | -1.918978791 | 0.033244696 |
| BMP2 | -1.941102451 | 0.016969065 |
| VCAN | -1.972423013 | 0.000601557 |
| CDHR1 | -1.975631936 | 0.015782929 |
| KRT8P46 | -2.000126916 | 0.04977048 |
| KLHL13 | -2.00589355 | 0.030369619 |
| SERPIND1 | -2.036094599 | 0.008979281 |
| MTCL1 | -2.069304495 | 0.005842123 |
| SLC16A10 | -2.124216301 | 0.008273981 |
| TNR | -2.124838755 | 0.000150224 |
| F2RL1 | -2.132700034 | 0.008914587 |
| TP53TG5 | -2.143632504 | 0.018811345 |
| TMEM139 | -2.157794081 | 0.045287461 |
| ADGRL2 | -2.197847388 | 0.000116802 |
| SNPH | -2.281889089 | 0.000510394 |
| SLC2A10 | -2.296809786 | 0.015303888 |
| TNFRSF8 | -2.341529357 | 0.002673436 |
| FAM110C | -2.353823706 | 1.32E-09 |
| RPL7P57 | -2.419207455 | 0.021150909 |
| MID2 | -2.508364301 | 0.005244131 |
| ACBD3-AS1 | -2.55453664 | 0.025409746 |
| AFF3 | -2.607557111 | 0.003931195 |
| ZSWIM5 | -2.61608675 | 0.003579428 |
| SCN2A | -2.650468574 | 9.04E-12 |
| TAB2-AS1 | -2.776855964 | 0.042096884 |
| RHPN2 | -3.004986992 | 0.031403848 |
| LAMC1 | -3.222902481 | 0.033920041 |
| LOC102724474 | -3.306472489 | 0.032211898 |
| LRIT2 | -3.326629242 | 0.03356152 |
| PRSS33 | -3.338774538 | 0.047265762 |
